# Laboratory evolution of *Escherichia coli* enables life based on fluorinated amino acids

**DOI:** 10.1101/665950

**Authors:** Federica Agostini, Ludwig Sinn, Daniel Petras, Christian J. Schipp, Vladimir Kubyshkin, Allison Ann Berger, Pieter C. Dorrestein, Juri Rappsilber, Nediljko Budisa, Beate Koksch

**Affiliations:** Institute of Chemistry and Biochemistry – Organic Chemistry, Freie Universität Berlin, Berlin, Germany; Institute of Chemistry – Biocatalysis, Technische Universität Berlin, Berlin, Germany; Institute of Biotechnology – Bioanalytics, Technische Universität Berlin, Berlin, Germany; Wellcome Centre for Cell Biology, University of Edinburgh, Edinburgh EH9 3BF, UK; Skaggs School of Pharmacy and Pharmaceutical Sciences, University of California, San Diego, USA

## Abstract

Organofluorine compounds are toxic to various living beings in different habitats. On the other hand, fluorine incorporation into single proteins via related amino acid analogues has become common practice in protein engineering. Thus, an essential question remains: can fluorinated amino acids generally be used as xeno-nutrients to build up biomass, or do large amounts of fluorine in the cells render them nonviable? To gain information about the effect of long-term exposure of a cellular proteome to fluorinated organic compounds, we constructed an experiment based on bacterial adaptation in artificial fluorinated habitats. We propagated *Escherichia coli* (*E. coli*) in the presence of either 4- or 5-fluoroindole as essential precursors for the in situ synthesis of tryptophan (Trp) analogues. We found that full adaptation requires astonishingly few genetic mutations but is accompanied by large rearrangements in regulatory networks, membrane integrity and quality control of protein folding. These findings highlight the cellular mechanisms of the evolutionary adaption process to unnatural amino acids and provide the molecular foundation for novel and innovative bioengineering of microbial strains with potential for biotechnological applications.

**One Sentence Summary:** Laboratory evolution enabled for the first time *Escherichia coli* to use fluorinated indoles as essential precursors for protein synthesis by introducing few genetic mutations but large rearrangements in regulatory networks, membrane integrity and quality control of protein folding.

## Main Text

For billions of years, living organisms have used mainly six chemical elements (carbon, hydrogen, nitrogen, oxygen, phosphorus and sulfur) out of the 118 available on Earth for the synthesis of the core macromolecules of life (DNA, RNA, proteins, lipids and carbohydrates). Fluorine is used almost exclusively by humans for various innovative syntheses while evolution has produced only a few chemical molecules and reactions with fluorine as a building block.(*1*) The high strength and polarization of the C-F bond alter the geometry, conformation and interactions of molecules, hence its incorporation has proven to be an extremely powerful method to modulate stability and/or activity of a vast variety of target materials, fine chemicals, drugs and pesticides.(*2–4*) Similarly, complex biological macromolecules such as peptides and proteins can be artificially labelled with fluorine by incorporation of fluorinated amino acids.(*5*). Highly fluorinated domains can drive the processes of protein-protein interaction and folding (*6*) or tune resistance against proteolytic degradation.(*7*) Thus, fluorinated substances have become indispensable in our daily lives. However, the properties that make fluorine so attractive for chemical synthesis often represent a threat in the cellular environment, where fluorine-containing organic compounds behave as environmental stressors.(*8*)

In this study, we investigated which biochemical factors limit the tolerance of natural systems towards fluorinated compounds. In particular, we tested whether the promiscuity of enzymatic reactions towards unnatural substrates (*9, 10*) can be exploited in order to convert toxic organofluorine compounds into essential metabolic precursors supporting the creation of cellular biomass. Inspired by reports from the 1960s, in which scientists had attempted to introduce fluorinated amino acids into proteins *in vivo* (*11*) but had observed growth inhibition, we opted to exploit evolutionary mechanisms to accomplish a proteome-wide incorporation of fluorinated amino acids. Specifically, we carried out two parallel adaptive laboratory evolution (ALE) experiments that coupled for the first time reprogrammed protein synthesis in *Escherichia coli* (*E. coli*) with *in situ* synthesis of fluorinated amino acid analogues via enzymatic conversion of fluorinated precursors. In particular, we created two metabolic prototypes with stable genetic architectures that were able to convert 4- and 5-fluoroindole into 4- and 5-fluorotryptophan, respectively, in a single-step reaction catalyzed by the endogenous enzyme Trp synthase (TrpS). Trp is an attractive target because it is encoded by a single codon (UGG) in the genetic code and there are no salvage pathways for its biosynthesis in *E. coli*. The replacement of Trp by 4-fluorotryptohan into the proteome of *E. coli* via long-term cultivation had been attempted before by Bacher and Ellington.(*12*) However, the authors reported that the fluorinated amino acid acted as toxic metabolite for the cells and they eventually achieved only partial incorporation, while the viability of their strain decreased over time.

In contrast, we report here the selection of clones that gained the ability to live under progressively more stringent conditions, characterized by the absence of Trp. We essentially adapted a procedure (*13*) that had previously enabled the trophic replacement of Trp by a sulfurated analogue by feeding the corresponding indole precursor to an *E. coli* strain lacking the *trp* operon. In our ALE experiments reported here, we generated a new Trp auxotrophic strain and then exposed it to 4- and 5-fluoroindole. We applied increasing selection pressure by decreasing the availability of Trp precursor (indole) until complete depletion. This method enabled the natural adaptation of *E. coli* to two different fluorinated Trp analogues and thus proved efficient for the generation of new synthetic life forms. It must be noted, that indole and its fluorinated analogues are not less toxic than the corresponding amino acids derivatives. In fact, they exert a concentration-dependent inhibition of cell division, by altering the electrostatic potential of cellular membranes.(*14*) Nevertheless, our long-term adaptation setup enabled cells to repurpose these toxic metabolites into substrates for the synthesis of an essential amino acid and subsequent proteome-wide incorporation. At the end of the ALE, the adapted strains had become facultative fluorotryptophan/Trp users, as they retained the genes encoding TrpRS (tryptophanyl-tRNA synthetase, *trpS*) able to recognize Trp and its analogues with similar efficiency.

In order to monitor whether the adaptation process had involved the simultaneous rearrangement of more than one class of biomolecules, isolates from different time points of the ALE experiment were studied by means of genomic, proteomic and metabolomic analyses which allowed us to integrate different layers of biochemical information. An adaptation model is proposed based on these data, highlighting that stress response mechanisms, quality of protein folding and membrane integrity were the main mechanisms where adaptation was most required in order to adjust the cellular metabolism to the fluorinated proteome and, thus, to create artificial biodiversity involving fluorinated substrates.

## Results

Based on previous literature reports and our own determination of the catalytic parameters of TrpRS, we knew that 4- and 5-fluorotryptophan would be suitable substrates for proteome synthesis (see Table S1). In preliminary experiments, we had observed that 4- and 5- fluoroindole were almost isosteric to indole but exhibited higher lipophilicity and polarity (Fig. S1A-C). In order to force *E. coli* to use the fluoroindoles as essential nutrients for growth, stable Trp-auxotrophy was achieved by deletion of genes from both biosynthetic (*trpLEDC; trp* operon) and degradation (*tnaA*; encoding the enzyme tryptophanase) pathways. The new strain was named TUB00 (*ΔtrpLEDC, ΔtnaA*, Fig. 1 and S2).

**Fig. 1.**
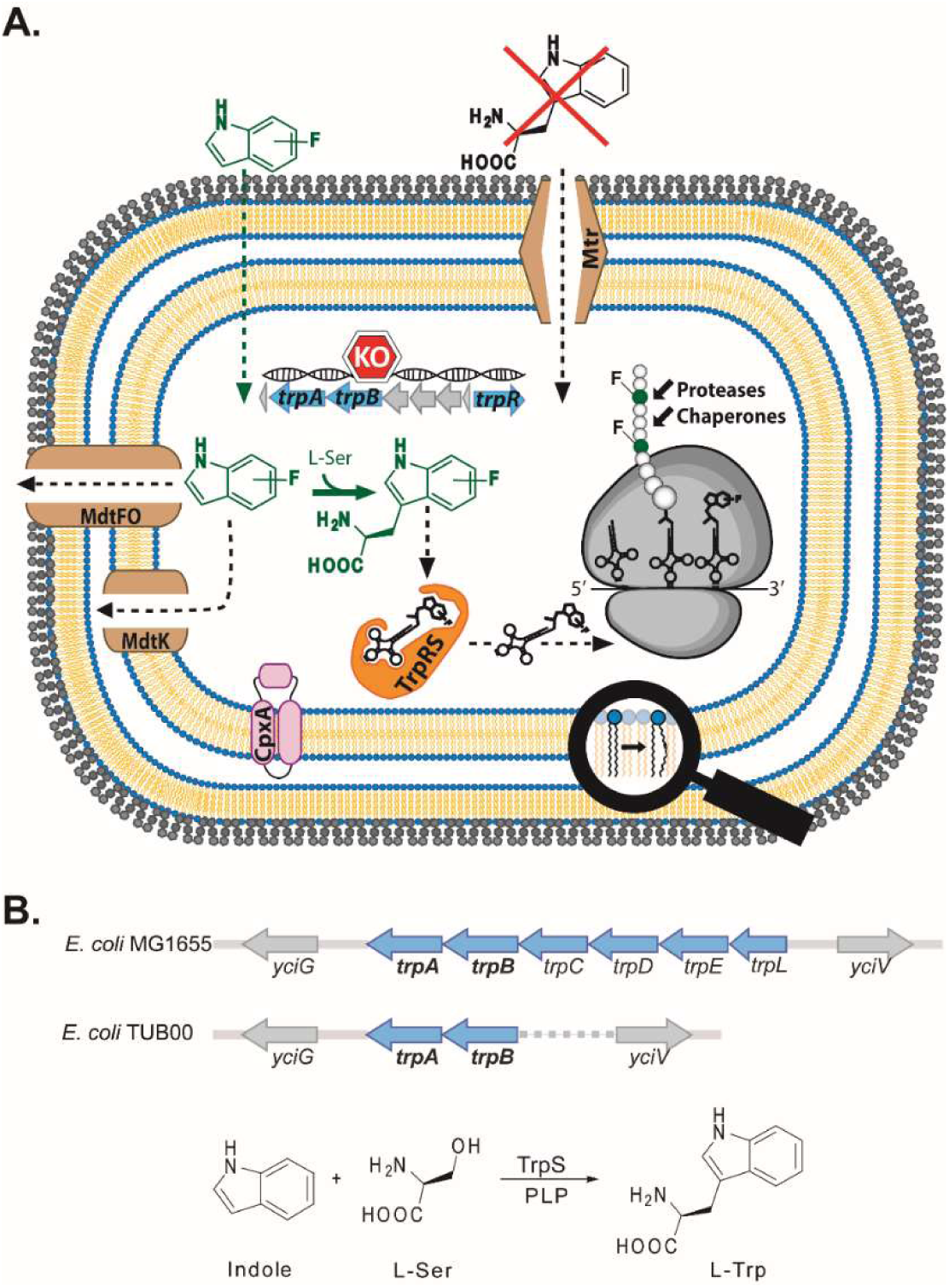
**(A)** ALE experimental setup and overview of the changes observed in the *E. coli* strains adapted to fluoroindoles that marked a divergence from TUB00. Abbreviations include: *trpA* (Trp synthase α subunit); *trpB* (Trp synthase β subunit); *trpR* (*trp* operon transcriptional repressor); TrpRS (tryptophanyl-tRNA synthetase); Mtr (Trp permease); MdtF, MdtO, MdtK (multi-drug efflux pumps); CpxA (sensor histidine kinase). Under natural conditions, cells can uptake extracellular Trp. However, during the ALE cultivation Trp was not supplied, hence its structure is crossed. (**B**) The *trp* operon of *E. coli* in the parent strain MG1655 and in the Trp-auxotrophic derivative TUB00 (used for ALE cultivation) after deletion of the genes *trpLEDC*. The PLP-dependent reaction catalyzed by TrpS is showed below.

The ALE experiment included continuous cultivation in shake flasks and serial re-inoculation of parallel cultures (see Method 5 in SI). Thereby, the concentration of the major growth nutrient indole was gradually decreased from 0.5 – 1 μM to zero, while keeping the levels of fluoroindoles and other nutrients (in particular glucose, nitrogen and phosphate) constant (Tables S3 and S4). The cells remained viable for over one year of cultivation and eventually acquired the ability of growing in NMM0 containing no amino acids and supplied exclusively with 4- or 5-fluoroindole. This demonstrated that *E. coli* had developed the ability to use these fluorinated substrates as metabolic intermediates, i.e. “xeno-nutrients”. Overall, we propagated parallel serial cultures for 825 generations and 93 serial re-inoculation steps in the case of 4-fluoroindole ALE (Fig. 2A) and 678 generations and 83 serial re-inoculation steps in the case of 5-fluoroindole ALE (Fig. 2B). The strains adapted to 4-fluoroindole are referred to below as “4TUBX” and those adapted to 5-fluoroindole as “5TUBX”, where X corresponds to the respective serial re-inoculation step number (“passage”).

**Fig. 2.**
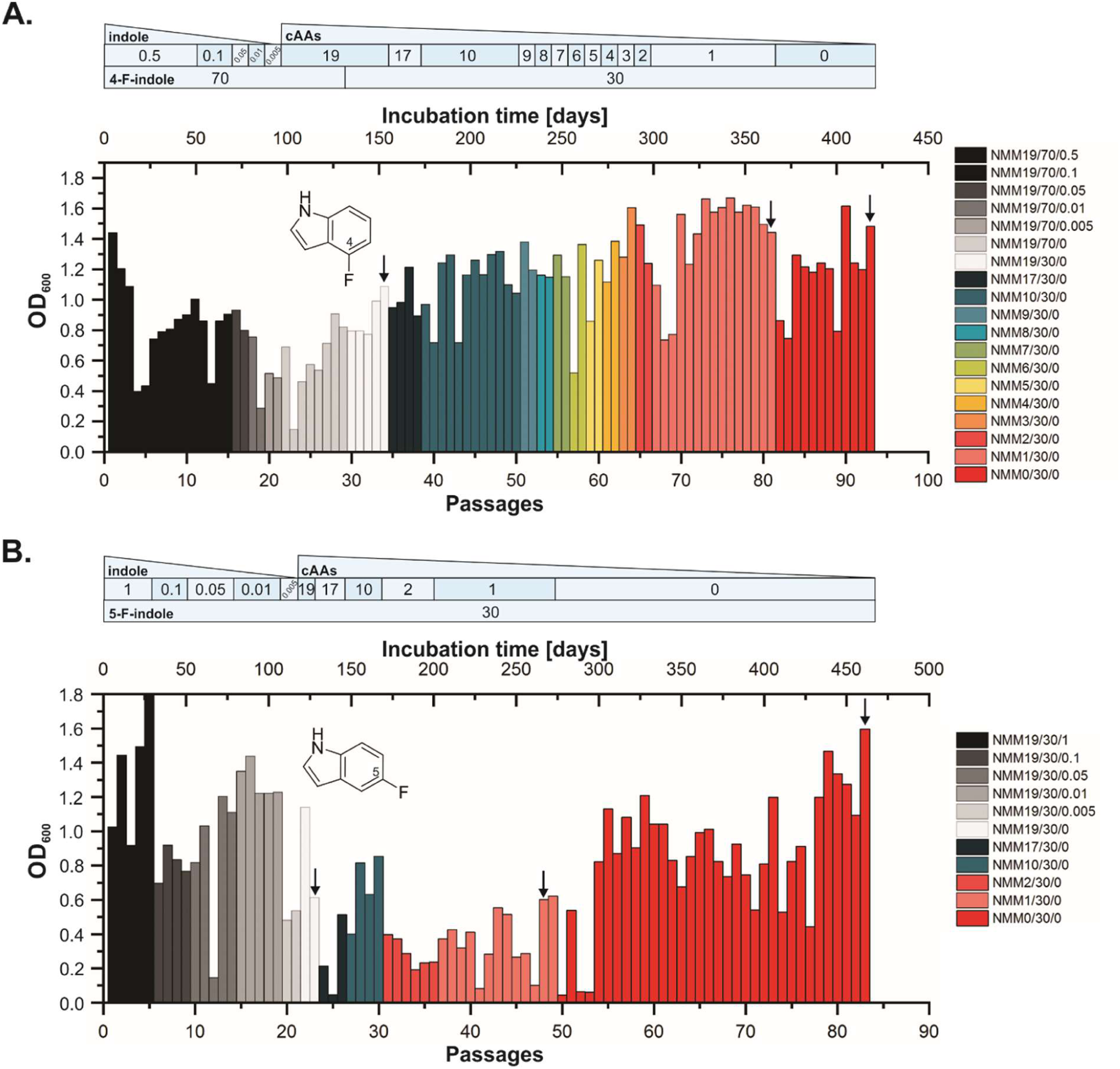
Fermentation schemes of *E. coli* ALE experiments towards usage of (**A**) 4-fluoroindole; (**B**) 5-fluoroindole as precursors for the synthesis of Trp analogues. Optical density (OD_600_) at the re-inoculation step (“passage”) is plotted against days of incubation and number of passages. The color code refers to the composition of New Minimal Medium (NMM, Tables S3 and S4), NMM*a/b/c* where “*a*” is the number of amino acids supplied, *“b”* is the concentration of fluoroindole and “*c*” is the concentration of indole, both µM. Black arrows indicate the isolates used for strain characterization, referred to as: early (4TUB34 and 5TUB23), intermediate (4TUB81 and 5TUB48) and final time points (4TUB93 and 5TUB83).

To elucidate the molecular mechanisms that had allowed the adaptation of 4TUB and 5TUB towards fluoroindoles as xeno-nutrients, we characterized isolates from the ALE experiments by means of whole genome sequencing, quantitative proteomic and non-targeted metabolomic analyses. Firstly, genomic mutations accumulated during the two parallel ALE experiments were identified by whole-genome sequencing of isolates of 4TUB and 5TUB at early, intermediate and final time points by means of an Illumina HiSeq 4000 sequencing platform (Table S5 and supplementary data). The impact of mutations on structure and function of the corresponding protein product was predicted by PROVEAN v1.1 (Protein Variation Effect Analyzer, Table S6).(*15*) Notably, *E. coli* adapted to whole-proteome fluorination by a relatively low number of mutations. Of more than 20,000 TGG (Trp) codons present in the genome of *E. coli*, only one was mutated during 5-fluoroindole ALE (in the gene *mrr*, encoding methylated adenine and cytosine restriction protein, Trp105Stop). This suggests a remarkable tolerance of the bacterial proteome towards global incorporation of fluorotryptophans. No major changes of ribosomal proteins were required to catalyze protein synthesis with 4- and 5-fluorotryptophan, which is in agreement with the well-known tolerance of the ribosome towards different amino acid analogues.(*16*) Only during 4-fluoroindole ALE single point mutations occurred at the level of the 30S ribosomal proteins S1 (*rpsA*, Table 1) and S10 (*rpsJ*, Table 1). Mutations of other genes involved in protein biosynthesis included isoleucyl-tRNA synthetase (ileS, Table 1) in 5-fluoroindole ALE and glycyl-tRNA synthetase (*glyQ*) in 4-fluoroindole ALE (this latter categorized by PROVEAN as neutral, Table S6). Two genes encoding proteases (*ptrA* and *ftsH*, Tables 1 and S6) were mutated over the course of 5-fluoroindole ALE, which might increase the cellular tolerance towards fluorinated proteomes.

**Table 1.**
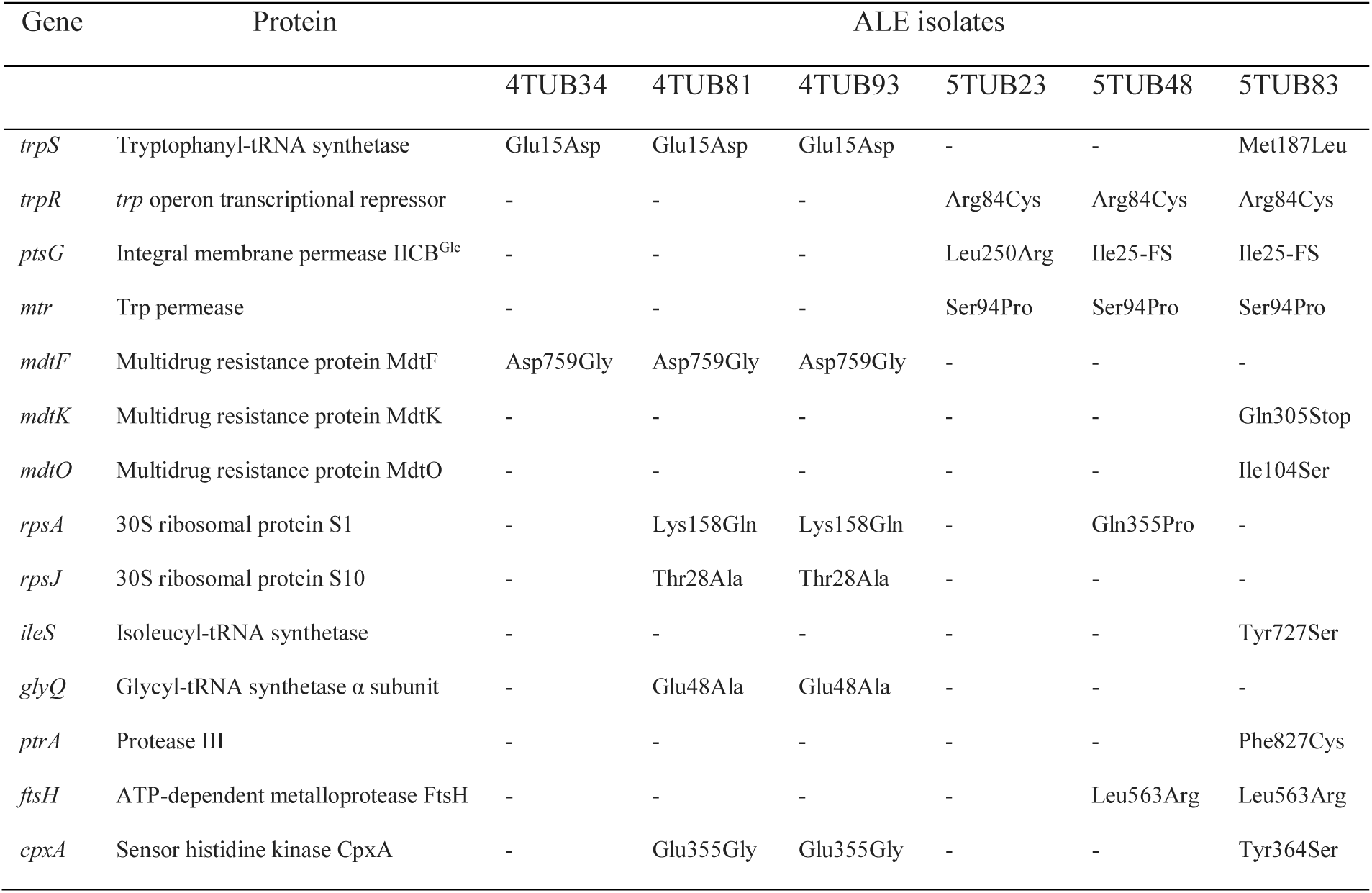
Genomic mutations in genes involved in indole, Trp and protein metabolism detected at early, intermediate and final time points of 4-fluoroindole ALE (4TUB34, 4TUB81 and 4TUB93) and of 5-fluoroindole ALE (5TUB23, 5TUB48 and 5TUB83). Deletion of a nucleobase in *ptsG* induces a translational frameshift (FS).

The gene *trpS*, encoding TrpRS, was mutated in both ALE experiments. We reasoned that the mutant enzymes might improve the usage of fluorinated Trp analogues for protein synthesis, especially in the case of 5-fluorotryptophan, where mutation of *trpS* correlates with the emergence of fully adapted phenotypes. The mutation of *trpS* in 4-fluoroindole ALE was categorized as neutral (Table S6), which is consistent with our preliminary observation that 4-fluorotryptophan is relatively efficiently activated by TrpRS (Table S1), hence adaptation did not require substantial modification of the enzyme.

Notably, during 5-fluoroindole ALE, a mutation in *trpR*, encoding the transcriptional repressor TrpR of the *trp* operon, altered the conserved residue Arg84, which is important for binding to DNA via electrostatic interactions (Table 1, Fig. S6).(*17*) TrpR regulates the catabolite-repression of Trp biosynthesis and downregulates the expression of *trpA* and *trpB*, encoding the Trp synthase. Replacement of the positively charged guanidinium group of Arg84 by the uncharged side-chain of Cys is expected to decrease the efficiency of TrpR as transcriptional repressor and to upregulate Trp biosynthesis. During the adaptation towards 4-fluoroindole, wild type TrpR is conserved and TrpS is downregulated (see supplementary data). Another mutation possibly involved in Trp biosynthesis is a frameshift truncation of the membrane glucose transporter PtsG/IICB^Glc^ (ptsG, Table 1) that appeared at the intermediate time point of 5-fluoroindole ALE. PtsG/IICB^Glc^is downregulated in all ALE isolates (see supplementary data) and the inactivation of this transporter in E. coli is associated with higher metabolic flux in the biosynthetic pathway of Ser,(*18*) that is required for the synthesis of (fluoro)tryptophan. Although the precise mechanism of indole uptake in the cells is still a matter of controversy, indole and its analogues are known to passively diffuse through cellular membranes.(*19*) Nonetheless, we found a mutation in mtr, encoding the high-affinity transporter Trp permease in all isolates from the 5-fluoroindole ALE experiment (Table 1). Since Trp is absent from the ALE medium, this mutation could inactivate an unused transporter (and thus be neutral) or facilitate 5-fluoroindole uptake into the cytoplasm and thus be beneficial for enlarging the pool of substrate for 5-fluorotryptophan synthesis. Proteomics data suggest that Mtr played a functional role in the adaptation towards fluoroindoles as xeno-nutrients, as it is upregulated throughout the whole 5-fluoroindole ALE as well as at the final time point of 4-fluoroindole ALE (4TUB93). Mutation of genes encoding multi-drug efflux pumps (*mdtK, mdtO*, Table 1) was observed in the final isolate of 5-fluoroindole ALE (5TUB83) and at early stages of 4-fluoroindole ALE (4TUB34, *mdtF*, Table 1). Multi-drug efflux pumps carry out detoxification in presence of xeno-compounds, such as the fluoroindoles.(*20, 21*) However, in our ALE setup, precisely these compounds must be accumulated, as they are essential precursors for Trp and protein synthesis. For this reason, we believe that mutation of this class of transporters was crucial for the adaptation to fluoroindoles.

After genomics, we investigated what changes had occurred in the proteome of *E. coli* upon global incorporation of fluorotryptophans. The fluorinated proteomes of the isolates from early and final time points of 4TUB and 5TUB were compared to the standard proteome of TUB00. Specifically, we quantified changes in abundance of single proteins by means of Stable Isotope Labelling by Amino acids in Cell culture analysis (SILAC, Fig. S3 and supplementary data).(*22*) Contrary to our expectation, relatively few proteins showed abundance change (see Fig. S4). However, functional proteins and enzymes directly involved in the quality control of protein folding were strongly affected. In particular, protein chaperones and proteases that assist folding and degradation of misfolded proteins were upregulated at the early time points of both ALEs (4TUB34 and 5TUB23, Table 2), thus suggesting that a stress response associated with the presence of a large number of misfolded proteins was underway at the beginning of the adaptation process. This might be a specific effect of the incorporation of the fluorotryptophans into the proteome. Remarkably, at the end of the ALE experiments (4TUB93 and 5TUB83; Table 2), the same proteins are downregulated, meaning that the situation of stress was resolved.

**Table 2.**
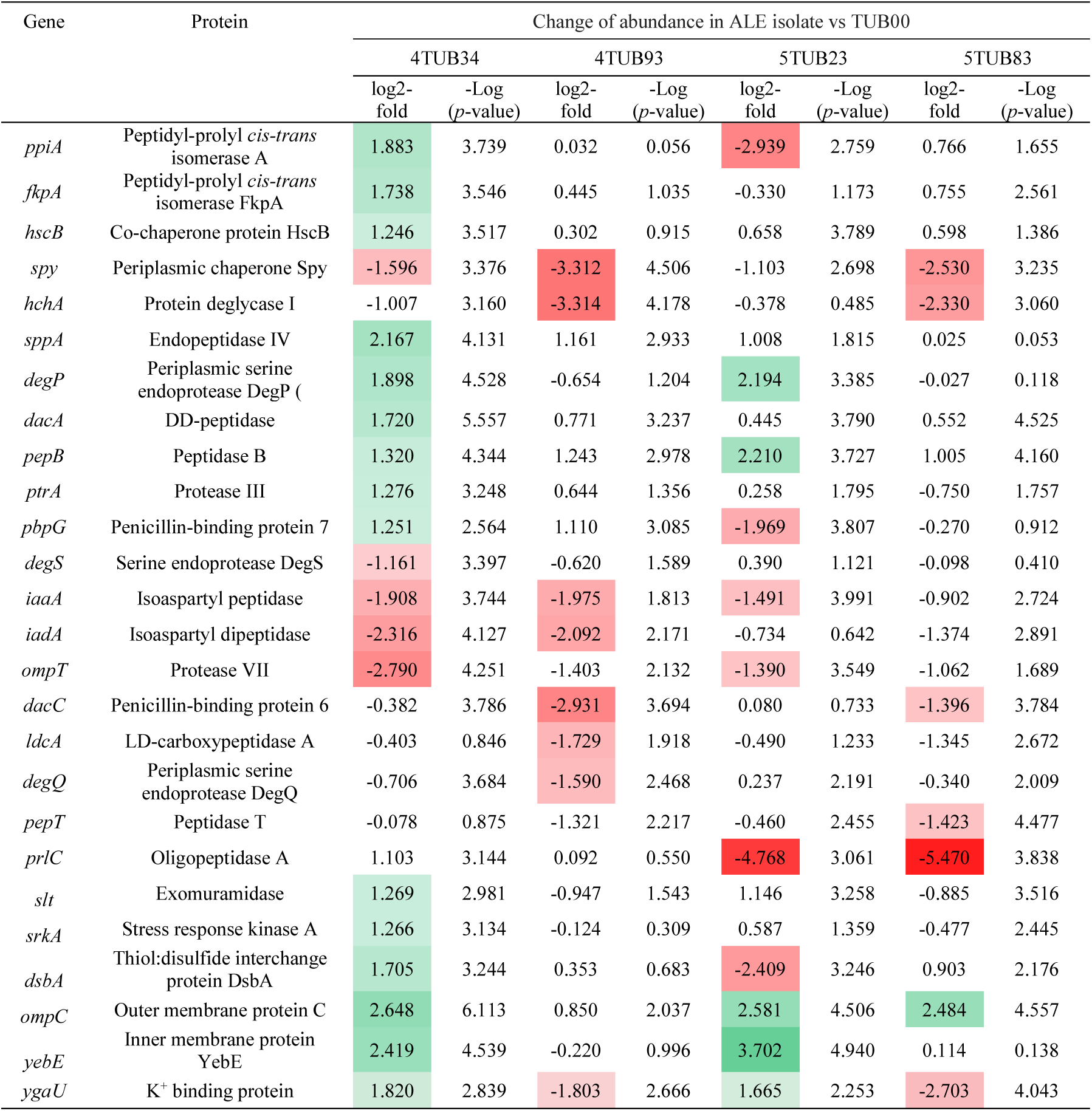
Differential abundance of protein chaperones and proteases at early and final time-points of 4-fluoroindole ALE (4TUB34 and 4TUB93, respectively) and 5-fluoroindole ALE (5TUB23 and 5TUB83, respectively), compared to TUB00. The change in abundance is reported as log2-fold and is visualized in the form of heat map, where green indicates upregulation (protein more abundant in the ALE isolates) and red indicates downregulation (protein more abundant in the TUB00 ancestor strain); white indicates a not significant abundance change.

| Gene | Protein | Change of abundance in ALE isolate vs TUB00 |  |  |  |  |  |  |  |
| --- | --- | --- | --- | --- | --- | --- | --- | --- | --- |
|  |  | 4TUB34 |  | 4TUB93 |  | 5TUB23 |  | 5TUB83 |  |
|  |  | log2-fold | -Log (p-value) | log2-fold | -Log (p-value) | log2-fold | -Log (p-value) | log2-fold | -Log (p-value) |
| *ppiA* | Peptidyl-prolyl *cis-trans* isomerase A | 1.883 | 3.739 | 0.032 | 0.056 | -2.939 | 2.759 | 0.766 | 1.655 |
| *fkpA* | Peptidyl-prolyl *cis-trans* isomerase FkpA | 1.738 | 3.546 | 0.445 | 1.035 | -0.330 | 1.173 | 0.755 | 2.561 |
| *hscB* | Co-chaperone protein HscB | 1.246 | 3.517 | 0.302 | 0.915 | 0.658 | 3.789 | 0.598 | 1.386 |
| *spy* | Periplasmic chaperone Spy | -1.596 | 3.376 | -3.312 | 4.506 | -1.103 | 2.698 | -2.530 | 3.235 |
| *hchA* | Protein deglycase I | -1.007 | 3.160 | -3.314 | 4.178 | -0.378 | 0.485 | -2.330 | 3.060 |
| *sppA* | Endopeptidase IV | 2.167 | 4.131 | 1.161 | 2.933 | 1.008 | 1.815 | 0.025 | 0.053 |
| *degP* | Periplasmic serine endoprotease DegP ( | 1.898 | 4.528 | -0.654 | 1.204 | 2.194 | 3.385 | -0.027 | 0.118 |
| *dacA* | DD-peptidase | 1.720 | 5.557 | 0.771 | 3.237 | 0.445 | 3.790 | 0.552 | 4.525 |
| *pepB* | Peptidase B | 1.320 | 4.344 | 1.243 | 2.978 | 2.210 | 3.727 | 1.005 | 4.160 |
| *ptrA* | Protease III | 1.276 | 3.248 | 0.644 | 1.356 | 0.258 | 1.795 | -0.750 | 1.757 |
| *pbpG* | Penicillin-binding protein 7 | 1.251 | 2.564 | 1.110 | 3.085 | -1.969 | 3.807 | -0.270 | 0.912 |
| *degS* | Serine endoprotease DegS | -1.161 | 3.397 | -0.620 | 1.589 | 0.390 | 1.121 | -0.098 | 0.410 |
| *iaaA* | Isoaspartyl peptidase | -1.908 | 3.744 | -1.975 | 1.813 | -1.491 | 3.991 | -0.902 | 2.724 |
| *iadA* | Isoaspartyl dipeptidase | -2.316 | 4.127 | -2.092 | 2.171 | -0.734 | 0.642 | -1.374 | 2.891 |
| *ompT* | Protease VII | -2.790 | 4.251 | -1.403 | 2.132 | -1.390 | 3.549 | -1.062 | 1.689 |
| *dacC* | Penicillin-binding protein 6 | -0.382 | 3.786 | -2.931 | 3.694 | 0.080 | 0.733 | -1.396 | 3.784 |
| *ldcA* | LD-carboxypeptidase A | -0.403 | 0.846 | -1.729 | 1.918 | -0.490 | 1.233 | -1.345 | 2.672 |
| *degQ* | Periplasmic serine endoprotease DegQ | -0.706 | 3.684 | -1.590 | 2.468 | 0.237 | 2.191 | -0.340 | 2.009 |
| *pepT* | Peptidase T | -0.078 | 0.875 | -1.321 | 2.217 | -0.460 | 2.455 | -1.423 | 4.477 |
| *prlC* | Oligopeptidase A | 1.103 | 3.144 | 0.092 | 0.550 | -4.768 | 3.061 | -5.470 | 3.838 |
| *slt* | Exomuramidase | 1.269 | 2.981 | -0.947 | 1.543 | 1.146 | 3.258 | -0.885 | 3.516 |
| *srkA* | Stress response kinase A | 1.266 | 3.134 | -0.124 | 0.309 | 0.587 | 1.359 | -0.477 | 2.445 |
| *dsbA* | Thiol:disulfide interchange protein DsbA | 1.705 | 3.244 | 0.353 | 0.683 | -2.409 | 3.246 | 0.903 | 2.176 |
| *ompC* | Outer membrane protein C | 2.648 | 6.113 | 0.850 | 2.037 | 2.581 | 4.506 | 2.484 | 4.557 |
| *yebE* | Inner membrane protein YebE | 2.419 | 4.539 | -0.220 | 0.996 | 3.702 | 4.940 | 0.114 | 0.138 |
| *ygaU* | K<sup>+</sup> binding protein | 1.820 | 2.839 | -1.803 | 2.666 | 1.665 | 2.253 | -2.703 | 4.043 |

Moreover, a subset of proteases and chaperones including DegP, PpiA, DsbA and Spy (*23*) was found to be under the common regulation of the two-component signal transduction system CpxAR, that controls the stress response against misfolded proteins in the periplasm of *E. coli*. This system is composed of the sensor histidine kinase *CpxA* and the transcriptional regulator CpxR, that activates the expression of genes encoding proteases and chaperones. Notably, at the end of both 4- and 5-fluoroindole ALE, DegP, PPiA, DsbA, Spy and other CpxAR-regulated proteins such as Slt, YgaU,(*24*) SrkA,(*25*) OmpC (*26*) and YebE (*27*) are not upregulated, concomitant with a mutation of cpxA in the sequence encoding the histidine kinase domain required for cross-talking with CpxR (Table 1). We conclude that, although Trp is the least-abundant amino acid in the proteome of *E. coli* (∼ 1%), its global fluorination induces a stress response, most likely associated with the partial misfolding of a large number of proteins. Cells coped with this condition by loosening the protein quality check mechanisms that normally ensure correct folding, such as protein chaperones and proteases.

Finally, we investigated whether the adaptation to fluoroindoles had altered the chemical composition of TUB00. The metabolomes of all relevant isolates were extracted and analyzed by non-targeted tandem-mass spectrometry and molecular networking with Global Natural Products Social Molecular Networking (GNPS, supplementary data).(*28*) Multivariate statistics of Principal Coordinate Analysis (PCoA) indicated that the metabolomes of TUB00 and the ALE isolates had increasingly diversified (Fig. 3A, B). Moreover, investigation of GNPS molecular networks revealed that the most significant changes had occurred at the level of Trp, biotin and lipid metabolites (Fig. 3C, D). In particular, Trp was present exclusively in TUB00, while fluorotryptophan was detected in the ALE isolates (Fig. 3C, E; Table S10).

**Fig. 3.**
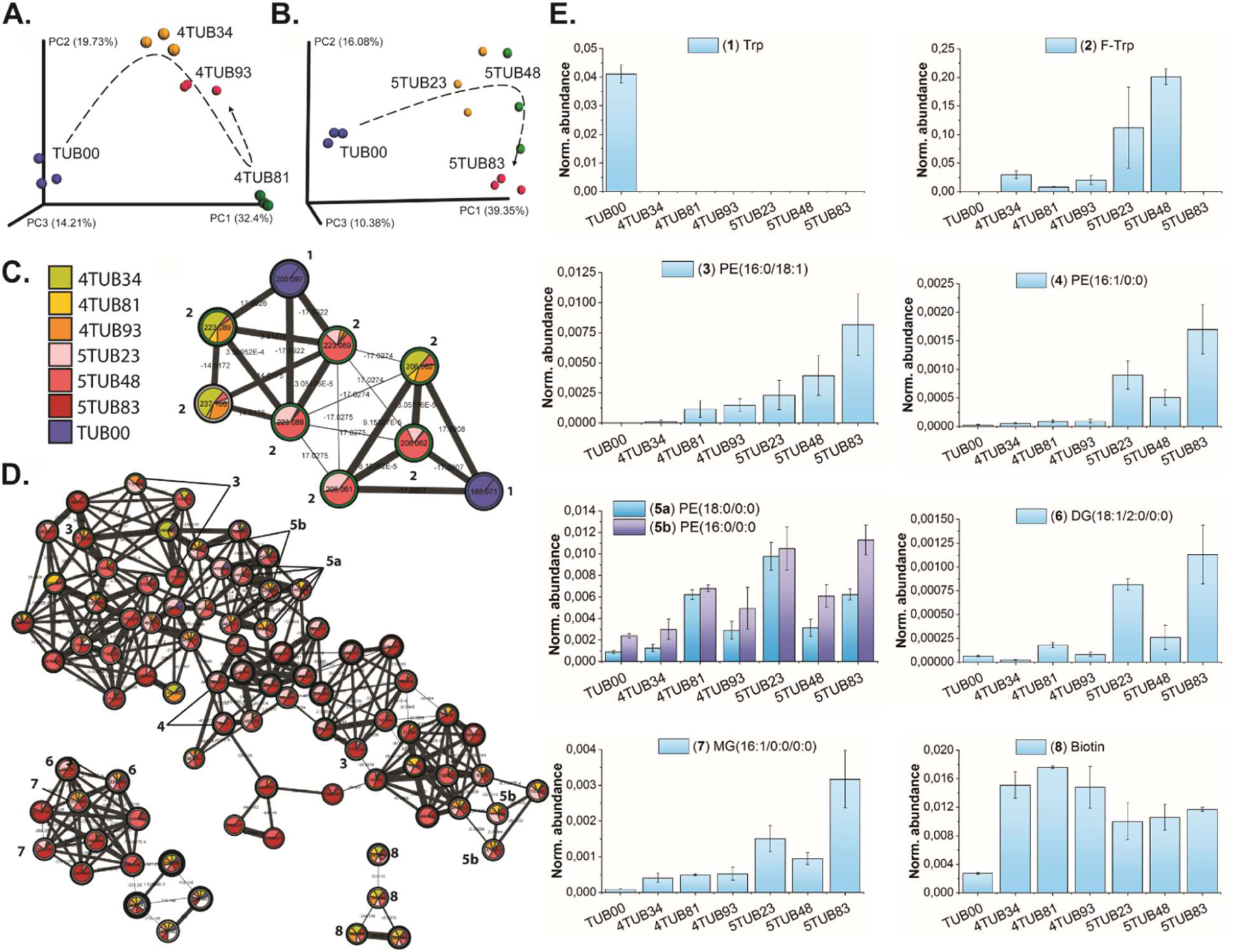
Analysis of metabolites produced during 4- and 5-fluoroindole ALE. (**A**) Principal Coordinate Analysis (PCoA) plots with Canberra distance metric of the metabolomes from 4-fluoroindole ALE and (**B**) 5-fluoroindole ALE. Each point represents the metabolome extracted from three independent cultures at early (yellow), intermediate (green) and final (red) time points of ALE and TUB00 (blue). The spatial distance in the plot is proportional to the chemical diversity between the samples and evolutionary trajectories are shown (dashed arrows). (**C**) Molecular subnetworks of Trp and fluorotryptophan and of (**D**) lipids and biotin. The nodes represent metabolites with unique retention time and m/z identifiers: (**1**) Trp; (**2**) fluorotryptophan; (**3**) 1-palmitoyl-2-oleyl-sn-glycero-3-phosphoethanolamine (PE(16:0/18:1)); (4) 1-palmitoleoyl-sn-glycero-3-phosphoethanolamine (PE(16:1/0:0); **5a**) 1-oleyl-2-hydroxy-sn-glycero-3-phosphoethanolamine (PE(18:0/0:0)); **5b**) 1-palmitoyl-2-hydroxy-sn-glycero-3-phosphoethanolamine (PE(16:0/0:0)); (**6**) 1-oleoyl-2-acetyl-sn-glycerol (DG(18:1/2:0/0:0)); (**7**) monopalmitolein (MG(16:0/0:0/0:0)); (**8**) biotin. The pie chart representation illustrates the relative abundance of each feature across the samples. (**E**) Normalized abundance of the annotated metabolites.

Remarkably, during the adaptation to 5-fluoroindole, (**2**) 5-fluorotryptophan^1^became up to 5-times more abundant than Trp in TUB00 (Fig. 3E, Table S10), which correlates with our hypothesis that mutation of TrpR upregulates 5-fluorotryptophan biosynthesis. However, it might also indicate that this analogue accumulates in the cytoplasm due to low kinetics of TrpRS. The abundance of 5-fluorotryptophan abruptly dropped by 10^3^-fold at the end of ALE (5TUB83) concomitant with a mutation in *trpS*. The mutant TrpRS (Met187Leu, Table 1) might improve kinetics and increase the usage of 5-fluorotryptophan for protein biosynthesis.

Besides the presence of fluorotryptophan, another distinctive trait of the fluoroindole-adapted strains was the presence of a large excess of biotin, glycerol- and phospholipids in comparison to the ancestor strain TUB00 (Fig. 3D, E). At the final time point of 5-fluoroindole ALE (5TUB83), a high abundance of phosphatidylethanolamines (the main components of *E. coli* membranes) carrying unsaturated palmitic (C16) and oleic (C18) fatty acid chains was observed. It is known that indole naturally acts as a stressor for cells by acting as a membrane ionophore (*14*) and this effect was expected to be stronger in the case of fluoroindoles, and especially 5-fluoroindole, being more lipophilic (thus having higher affinity for cell membranes) as well as more polar (thus increasing the flux of electrolytes across the membrane) than indole itself.

For these reasons, we hypothesized that the cell membrane rearrangement is required for the cells in the process of their specialization to fluoroindoles. Enrichment of membranes with lipids carrying unsaturated fatty acids generally increases the fluidity of the lipid bilayer as it perturbs the stacking of adjacent chains and introduces disorder in the overall structure.(*29*) We investigated the properties of the cell membranes in 4TUB93, 5TUB83 and TUB00 by fluorescence microscopy. Cells were treated with Nile Red, a stain and hydrophobic probe that intercalates between membrane lipids and whose fluorescence is quenched when it is exposed to polar solvents such as water (Fig. 4A, B). Both adapted strains were significantly less fluorescent than the ancestor strain (Fig. 4C, D) and particularly in the case of 5TUB83, the vast majority of cells were not fluorescent at all (Fig. 4C, D), thus suggesting that the membrane retention of Nile Red had decreased.

**Fig. 4.**
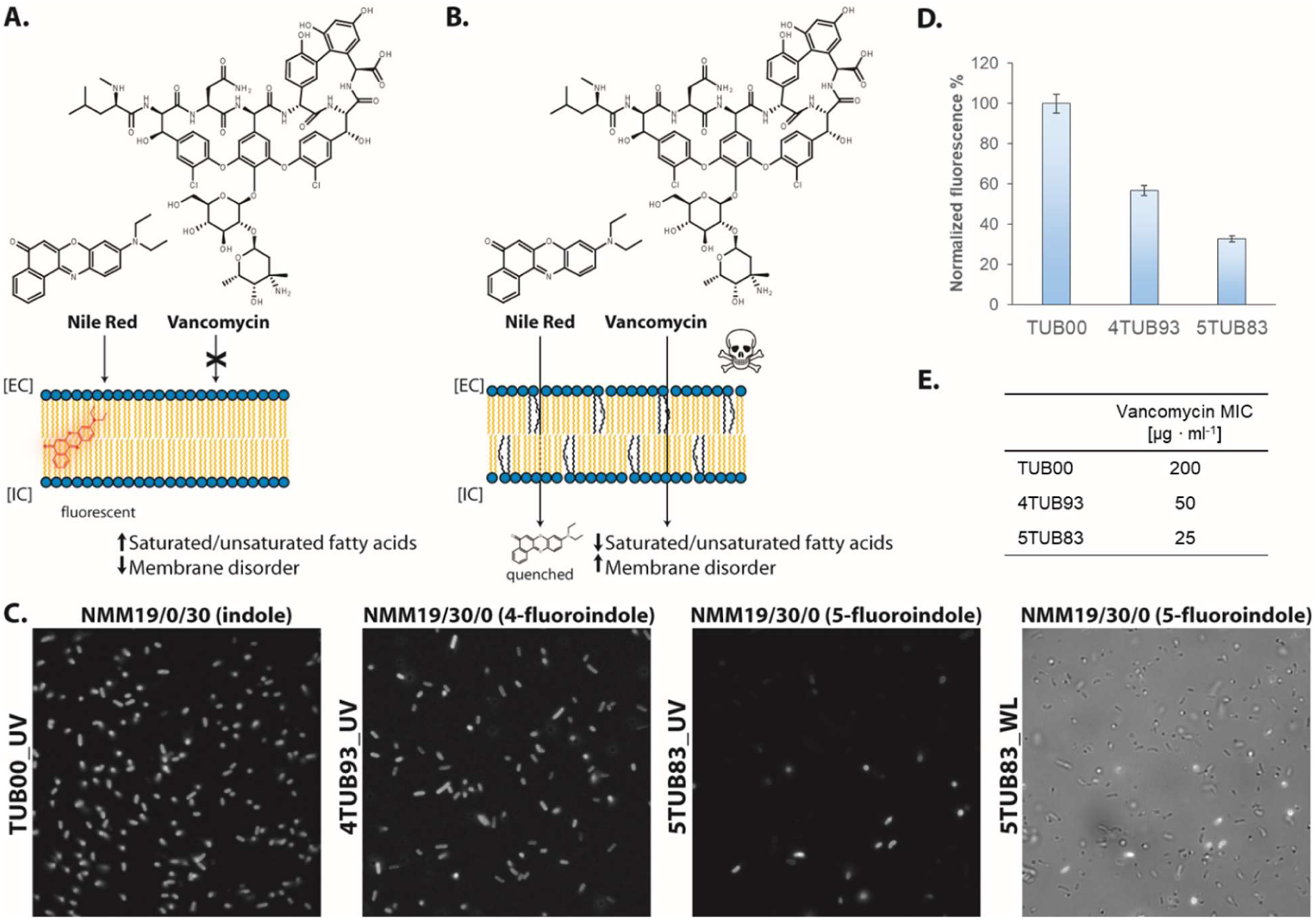
Cell membrane rearrangement during 4- and 5-fluoroindole ALE. (**A, B**) Role of lipid composition of *E. coli* membrane in retaining the hydrophobic stain Nile Red and preventing the penetration of the antibiotic vancomycin were investigated (**C**) Fluorescent micrographs of the ancestor strain TUB00 and of the final time-points of 4- and 5-fluoroindole ALE (4TUB93 and 5TUB83, respectively) stained with Nile red. The last panel reports 5TUB83 irradiated with white light (WL) for cell count comparison. (**D**) Total fluorescence normalized to TUB00. (**E**) Minimal inhibitory concentration (MIC) of vancomycin.

The rearrangement of the cell membrane might favor the uptake of fluoroindoles or reduce the toxic effect when these analogues accumulate in the lipid bilayer. We further tested the permeability of the cell membrane by measuring the susceptibility of TUB00, 4TUB93 and 5TUB83 to vancomycin, a high molecular weight antibiotic that is usually ineffective against *E. coli* as its passage through cell membranes is negligible (Fig. 4A).(*30*) We found that both adapted strains are less tolerant towards vancomycin than TUB00 (Fig. 4E, Table S11). This confirmed that adaptation towards fluoroindoles affected the organization of the cell membrane and increased its permeability to extracellular solutes.

## Discussion

We performed two parallel adaptive laboratory experiments (ALE) that enabled *E. coli* to use two fluorinated indole analogues as xeno-nutrients, i.e. essential building blocks for the synthesis of all endogenous proteins. In particular, we established the *in vivo* synthesis of 4- and 5-fluorotryptophan from 4- and 5-fluoroindole via intracellular metabolic conversion and subsequent proteome-wide translation in response to more than 20,000 UGG codons in *E. coli*. The 4- and 5-fluoroindole-adapted strains exhibit modified vital processes such as protein folding, membrane dynamics, stress and nutrient uptake. Genomic, proteomic and metabolomic analyses revealed that evolutionary mechanisms originated from a small number of mutations and were accompanied by only minor changes in the protein synthesis machinery. At the initial adaptation stages, the key features of adaptation included upregulation of proteases and protein chaperones as well as activation of the CpxAR-mediated stress response. As cells became proficient in utilizing fluoroindoles, these processes were attenuated. *E. coli* exhibited a remarkable ability to balance quality control of protein folding with enforced nutritional requirements by accepting fluorine-containing substrates. At the same time, this is strong evidence that the interplay between protein mutational robustness, protein folding and environmental stress is a key factor that determines the evolution of new traits in habitats containing fluorinated compounds. The challenge of continuous exposure to fluorinated stressors was responded to primarily at the level of the cell membrane, as demonstrated by evidences of membrane rearrangement and diversification of the lipid production, especially during the adaptation to 5-fluoroindole. The change in lipid composition did not compromise the viability of the strains and reflected the difference in lipophilicity and polarity between the two fluorinated xeno-nutrients. Upregulated production of unsaturated phosphatidylethanolamines changed the membrane properties and rendered the adapted strains more susceptible towards the antibiotic vancomycin. Although the 4TUB and 5TUB strains developed the ability to grow in a fluorinated habitat, under conditions that are not permissive to the parent strain TUB00 (Fig. S7) their growth behavior suffered by significant elongation of the generation time (Table S12). This hints that the strain robustness did not increase at the same pace of the adaptation to xeno-nutrients and that limitations in key physiological mechanisms might still exist. It is reasonable that adaptation was facilitated by mutations in transporter proteins such as Mtr and multi-drug efflux pumps (MdtF, MdtO, MdtK) that favored intracellular accumulation of the fluorinated xenobiotic compounds.

Here we report the first laboratory adaptation of *E. coli* to two different fluorinated indole analogues as precursors for endogenous protein synthesis that eventually led to the complete inclusion of 4- and 5-fluorotryptophan into the proteome of actively proliferating cells. An integrative multi-omics analysis of the adaptation process allowed us to prove that our method enabled the selection of clones with reconfigured regulatory networks, albeit carrying surprisingly few genetic mutations. With the help of these two new bacterial strains we are ready to usher in a new era that will focus on the uptake, toxicity, and metabolism of fluorine-carrying building blocks as, for the first time, it becomes possible to study the way in which whole living organisms accommodate fluorine.

The established fluoroindole-adapted strains provide us with a solid basis for 1) further investigations that are needed to identify the core biological barriers that control microbial adaptation to unnatural chemistries, e.g. antibiotics and pollutants; 2) developing biocontained strains, whose survival and proliferation is strictly dependent on the presence of the xeno-nutrients and will ensure environmental biosafety, and 3) engineering microbial strains with improved growth behavior for biotechnological applications, e.g. facile fluorine-labelling of bioactive peptides. Following these new avenues will lead us to what we see as next main breakthrough i.e. to explore alternative life forms based on xeno-elements.

## Supporting information

Auxiliary Table 1_Genomics

Auxiliary Table 2_Proteomics_4TUB34

Auxiliary Table 2_Proteomics_4TUB93

Auxiliary Table 2_Proteomics_5TUB23

Auxiliary Table 2_Proteomics_5TUB83

Auxiliary Table 3_Metabolomics

## Acknowledgments

The authors thank Dr. Franz-Josef Schmitt (TU Berlin) for cell imaging by fluorescence microscopy and Dr. Torsten Semmler (Robert Koch Institut, Berlin) for revision of sequencing data.

## Author contributions

F. A. designed the experimental setup, performed strain cultivation and all other experiments and wrote the manuscript. D. P. performed metabolomics MS data acquisition, supervised molecular networking data analysis and assisted in writing the manuscript. L. S. performed shotgun proteomics data acquisition and SILAC raw data analysis. C. S. performed the measurements of ATP/[^32^P]pyrophosphate exchange assay. V. K. performed the measurements of fluoroindoles dipole moment and lipophilicity. A. B. assisted in designing experiments and in writing the manuscript. P. D. directed metabolomics experiments. J. R. directed proteomics experiments and assisted in writing the manuscript. N. B. conceived project, directed experimental work and wrote the manuscript. B. K. designed experiments and wrote the manuscript.

## Data and materials availability

The complete genomic, proteomic and metabolomic datasets are available as auxiliary supplementary tables.

All metabolomics MS/MS data can be found on the Mass spectrometry Interactive Virtual Environment (MassIVE) at https://massive.ucsd.edu/ with the accession number MSV000083134. Molecular Networking and Spectrum Library Matching results can be found online at GNPS under the following links:

https://gnps.ucsd.edu/ProteoSAFe/status.jsp?task=628bd0896fad4eae91f6be5f12373d25 (analog search on); https://gnps.ucsd.edu/ProteoSAFe/status.jsp?task=7b23638cf23a4cb5ab71abbbd6a01248 (analog search off).

## Footnotes

1 The isomers 4- and 5-fluorotryptophan cannot be distinguished by LC-MS/MS and GNPS. It was assumed that 4-fluorotryptophan was present in 4TUB and 5-fluorotryptophan in 5TUB, corresponding to the precursors supplied in the cultivation medium, 4- and 5-fluoroindole, respectively.

