## Supplementary Material for "Laboratory evolution of *Escherichia coli* enables life based on fluorinated amino acids"

**Materials and Methods:**

**External databases:**

Auxiliary table 1. Genomics

Auxiliary table 2. Proteomics

Auxiliary table 3. Metabolomics

**Materials and Methods:**

**1. Measurement of dipole moment and lipophilicity of indole and fluoroindoles**

It had been previously reported in literature that *E.coli* can metabolically convert indole and several derivatives into Trp analogues by using genomic levels of tryptophan synthase (TrpS). To investigate the physicochemical properties of indoles carrying alternative fluorine substitutions, we determined polarity and lipophilicity of indole (log*P*) and 4-, 5-, 6-, 7-fluoroindole by measuring dipole moment (D) and octanol/water partition. All fluoroindoles are significantly more lipophilic than indole and 4-, 5- and 6-fluoroindole exhibit larger dipole moments.

Molecular modelling was performed in Scigress Modelling 3.2 (Fujitsu, Poland). COSMO volume was calculated using PM6 algorithm from the MOPAC package assuming aqueous medium. Dipole moments and electrostatic potential map were calculated using DFT B88-PW91 functional with the DZVP basis set. Lipophilicity was accessed by partitioning the substances between octan-1-ol and deionized water. 4-7 mg of a compound was shaken with the solvents, 1.00 ml each phase, for ≥ 18 hours at 298±2 K. Samples, 0.30 ml each were taken, and these were mixed with 0.30 ml of acetonitrile-d3 in the identical type NMR tubes. 1H NMR spectra were measured in 30-degree pulse multiscan experiments with 3.3+3 s acquisition+recycling time at 700 MHz frequency and 298 K. 19F{1H} NMR spectra were measured in 30-degree pulse multiscan experiments with 0.5+3.5 s acquisition+recycling time at 471 MHz frequency and 298 K. The spectra of the phases were overlaid for intensity comparison, and intensity ratios were taken as partitioning constants. Repetition of the partitioning was the main source of the errors, the measurements/sample tuning error is also included in the final values.

**2. ATP/[$^{32}$P]pyrophosphate exchange assay and tryptophanyl-tRNA synthetase kinetics determination**

The AMP-adenylation reaction (amino acid activation) of Trp, 4- and 5-fluorotryptophan by *E. coli* tryptophanyl-tRNA synthetase (TrpRS) was investigated by ATP/[$^{32}$P]pyrophosphate ($PP_i$) exchange assay. The reaction was performed at 37 °C in 100 mM Tris/HCl (pH 8.0), 80 mM $MgCl_2$, 5 mM KF, 2.2 mM $NaP_2O_7$, 1.25 mM β-mercaptoethanol, 5.5 mM ATP and 0.1 mg/mL BSA. $^{32}$P-labeled $PP_i$ was added to 10 μCi in each reaction. The concentration of TrpRS varied from 0.5 to 2 μM and the concentration of tryptophan, 4- and 5-fluorotryptophan ranged from 0.025 mM to 2 mM. The total volume of the reaction was 600 μL. Michaelis-Menten kinetics were obtained by quenching the reaction in 600 μL of 240 mM sodium pyrophosphate containing 70% (v/v) perchloric acid and 7.5% (w/v) activated charcoal at 30 s, 1 min, 2 min, 5 min and 10 min time-points. The suspension was vortexed and filtered through a filter paper. The filter was then washed twice with 10 mL of distilled water and soaked into 4 mL of scintillation solution, then the adsorbed [$^{32}$P]-labeled ATP was analyzed by scintillation counting.

### 3. Recombinant expression of tryptophanyl-tRNA synthetase

The tryptophanyl-tRNA synthetase from *E. coli* (TrpRS) was expressed in *E. coli* BL21(DE3) (New England Biolabs, Germany). The strain was rendered chemically competent and transformed with the expression plasmid pQE80L-6H-ecTrpRS (ori ColEI, AmpR) carrying the sequence of TrpRS with an additional N-terminal hexahistidine purification tag. An overnight culture (LB, 1 mM Amp) was used to inoculate (1:1,000) 1L of LB, 1 mM Amp. The culture was incubated at 37 °C, 250 rpm shaking until it reached $OD_{600}$ 0.6 - 0 .7, then heterologous gene expression was induced by adding isopropil-β-D-1-tiogalattopiranoside (IPTG, 1 mM) and incubating for 4 additional hours at 30 °C, 250 rpm shaking. Cells were harvested, resuspended in 50 mM TRIS/HCl, 100 mM NaCl, 20 mM imidazole (pH 8 at 4 °C), then lysozyme, DNAse and RNAse were added. Cells were lysed by sonication and cell debris was pelleted by centrifugation (40 min, 16,000 g, 4 °C). The soluble phase was filtered (Millex GV Filter Millipore, 0.45 µm pore diameter) and TrpRS purification was achieved via affinity chromatography with a Protino Ni-NTA column, 5ml Fast Flow (Macherey-Nagel). A linear gradient of mobile phase A (50 mM TRIS/HCl, 100 mM NaCl, 20 mM imidazole, pH 8 at 4 °C) and phase B (50 mM TRIS/HCl, 100 mM NaCl, 250 mM imidazole, pH 8 at 4 °C) with 0-100% B over 5 column volumes was applied. Protein elution was followed by UV absorbance at 280 nm, then the eluted fractions were collected and imidazole was removed by dialysis (Sigma-Aldrich dialysis sack, cutoff 3 kDa) against 50 mM TRIS/HCl, 100 mM NaCl (pH 8 at 4 °C). TrpRS identity was confirmed by LC-ESI-Q-TOF mass spectrometry and sodium dodecylsulphate polyacrylamide gel elecrophoresis (SDS-PAGE). The purity of the final product was > 95%.

### 4. Minimal Inhibitory Concentration measurement

Triplicate cultures of 4TUB93, 5TUB83 and TUB00 in LB medium (1 ml) were inoculated from cryostocks and incubated at 30 °C, 180 rpm until the optical density corresponding to 0.5 McFarland standard was reached. Serial dilutions of LB medium supplemented with indole, 4- and 5-fluoroindole (1 - 10 mM) were aliquoted onto 96-wells plates, inoculated 1:100 to a final volume 200 µl and incubated 16 hours at 30 °C, 180 rpm shaking. $OD_{600}$ was measured on a Tecan Infinite Microplate Reader (Männedorf, Switzerland).

### 5. Adaptive Laboratory Evolution experiments (ALE)

The adaptation of *E. coli* towards 4- and 5-fluoroindole was achieved by continuous aerobic cultivation in shake flasks where microbial cells were propagated in parallel serial cultures. The synthetic medium used for ALE was New Minimal Medium (NMM) containing: 22.5 mM $KH_2PO_4$, 50 mM $K_2HPO_4$, 7.5 mM $(NH_4)_2SO_4$, 8.5 mM NaCl, 1 mM $MgSO_4$, 1 mg/L $Ca^{2+}$ (as $CaCl_2$), 1 mg/L $Fe^{2+}$ (as $FeCl_2$), 10 ng/L trace elements ($Cu^{2+}$, $Zn^{2+}$, Mn2+, and $MoO_4^{2-}$), 20 mM glucose, 10 mg/L biotin and thiamine. 50 mg/L of canonical amino acids, indole, 4- and 5-fluoroindole were additionally added as described in Table S3, S4. The strain propagation was performed by cultivating biological triplicates (three independent cultures) for each ALE experiment (supplemented with either 4- or 5-fluoroindole) in a volume of 10 ml. At the beginning of ALE, TUB00 was cultivated in New Minimal Medium (NMM) supplied with all amino acids (except Trp). The strain was provided with the highest concentration of 4- or 5-fluoroindole (70 and 30 μM, respectively) and the lowest concentration of indole required to reach $OD_{600}$ of 1 within three days. Cultures were incubated at 30 °C and 180 rpm shaking until the early stationary phase ($OD_{600}$ 0.8 - 1.0) was reached. The volume corresponding to $OD_{600}$ 0.02 was then transferred to a new batch of fresh medium, this serial dilution being indicated as "passage". Cryostocks in 7% DMSO were prepared at each passage and stored at – 80 °C for further analyses. During ALE, the concentration of 4-fluoroindole was decreased from 70 to 30 μM because it was observed that this led cells to grow to higher density. After the cells became independent from indole for growth, all amino acids were gradually removed from the cultivation medium. This step was necessary in order to rule out possible contamination by Trp coming from commercially available preparations.

### 6. Genome sequencing

Isolates from early (4TUB34, 5TUB23), intermediate (4TUB81, 5TUB48) and final time-points of ALE experiments (4TUB93, 5TUB83) were selected for whole genome sequencing. Cells were revitalize from cryostocks and streaked twice on LB, 1.5% agar plates, then a single CFU was used to inoculate 5 ml of LB medium, subsequently incubated overnight at 30 °C, 180 rpm shaking. Chromosomal DNA was extracted with Genome Wizard Kit (Promega; Madison, USA), precipitated with ethanol absolute and dried at room temperature for 15 minutes. Chromosomal DNA was sequenced by Illumina HiSeq 4000 sequencing platform (BGI, Hong Kong) using the genome of *E. coli* K-12 MG1655 (NC_000913.3, as downloaded from https://www.ncbi.nlm.nih.gov) as reference sequence.

### 7. Prediction of mutations impact on proteins biological function by PROVEAN v1.1

The biological relevance of genomic mutations was estimated by predicting the impact of amino acid substitutions or insertions/deletions (InDel) on the biological function of the targeted proteins by the tool PROVEAN Protein (Protein Variation Effect Analyzer) v1.1. The default PROVEAN score threshold -2.5 for binary classification (i.e. deleterious vs neutral) was used.

### 8. Sample preparation for SILAC proteomic analysis

Proteomics sample preparation was based on a modified protocol of the Filter-aided Sample Preparation (FASP). TUB00 and isolates from early (4TUB34, 5TUB23) and final time-points of ALE (4TUB93, 5TUB83) were cultivated at 30 °C, 180 rpm in light SILAC media (TUB00 in NMM(19/0/30); 4TUB34 and 4TUB93 in NMM(19/30/1) 4-fluoroindole; 5TUB23 and 5TUB83 in NMM(19/30/1) 5-fluoroindole). The reference strain TUB00-dKO (TUB00*ΔargA::FRT, ΔlysA::FRT*, Budisa Lab, TU Berlin) was cultivated in heavy SILAC medium NMM(17/30/1) containing all amino acids except Trp, Arg or Lys and supplemented with 0.5 g/L $^{13}C_6{}^{15}N_4$-Arg, $^{13}C_6{}^{15}N_2$-Lys (Silantes, Munich, Germany). Arg and Lys auxotrophy ensured the complete isotope labelling of the proteome. Two parallel cultivations of TUB00-dKO were grown in presence of 1 μM indole and 30 μM of either 4- or 5-fluoroindole. All cultures were cultivated in the corresponding SILAC medium until they reached the stationary phase of growth and then reinoculated 1:100 in fresh medium. This procedure was repeated two times in order to ensure complete labeling of the proteome. After the third cultivation, cells were harvested by centrifugation (3,000 g, 5 minutes) during the early stationary phase ($OD_{600}$ 0.8 - 1). Wet cell mass was weighted and adjusted to 20 mg/ml with lysis buffer (0.5 % SDS, 0.1 M Tris pH 8.5 at 25 °C, 0.1 M DTT). The resuspended samples were heated at 95 °C for 5 minutes and cleared from insoluble material by spinning at 20,000 g for 15 minutes. 50 µl of cleared lysates from the adapted strains and TUB00 were mixed in equal ratio with the heavily-labeled reference TUB00-dKO. The samples were mixed and diluted with Urea buffer (8 M in 100 mM TRIS pH 8.5 at 25 °C), applied on the filter and centrifuged at 13,000 g for 20 minutes, 25°C. Following this step, 50 mM iodoacetamide in 8 M urea was added, mixed shortly and incubated for 20 minutes in the dark. After that, the iodoacetamide was depleted by spinning again and three wash steps with Urea buffer were applied. The buffer was exchanged to 1 M Urea in 100 mM Tris, pH 8.0 at 25 °C by three iterations of spinning and reapplication of this buffer. Tryptic digestion of the samples was achieved by adding 2.5 µg of trypsin per sample (estimated ratio 1:80 (m/m)) and incubation for 16 hours at 37 °C. The tryptic digestion was stopped by adding 10 % trifluoroacetic acid (TFA) ad 0.5 % and centrifuged at 13,000 g for 15 minutes, 25°C. The eluate was collected and pooled with two further elution steps using the digestion buffer with 0.5 % TFA. The samples were finally cleaned by following the protocol of Stage-tipping by Rappsilber et. al. The C18 solid phase was activated with 20 µl MeOH and equilibrated with 2x20 µl 0.1% TFA, prior loading 150 µl sample. The sample was washed twice with 20 µl 0.1% TFA and stored at -20 °C until further use.

**9. MS data acquisition and analysis for SILAC**

Two µg of peptides dissolved in 3.2 % acetonitrile with 0.1% formic acid were loaded directly on 50 cm EASY-Spray column (Thermo Fisher Scientific) packed with C18-stationary phase and equilibrated to 3 % B at a flow of 0.3 ml/min. Mobile phase A consisted of water with 0.1 % formic acid, while mobile phase B contained 80 % (v/v) acetonitrile with 0.1% formic acid. Peptide elution was achieved through gradient elution at 0.25 ml/min flow ramping up linearly 4 - 35 % B over 122 minutes, increasing to 45 % in 10 minutes and to 95 % in 8 minutes. Eluting peptides were sprayed into a hybrid Orbitrap-linear ion trap mass spectrometer (Fusion Lumos Tribrid, Thermo Fisher Scientific), quadrupole-isolated from 375 - 1500 m/z and recorded in a MS1 scan using the Orbitrap detector (R = 120,000). Peptides fulfilling isotopic compositional constraints, an intensity threshold, possessing a charge state of 2 - 6 and a recognized mass difference of the corresponding SILAC-pair (Lys8: 8.0142 Da; Arg10: 9.9874 Da) were then isolated using a window of 1.6 Da. Fragmentation was accomplished using CID with stepped 35 % normalized collision energy +/- 1 % and resulting fragments were recorded in the linear ion trap set to scanning rate "rapid". Dynamic exclusion was enabled for a duration of 60 seconds after single observation within boundaries of 5 ppm each. Processing of raw-files was performed using MaxQuant 1.5.7.4 with reviewed protein entries downloaded from SwissProt database for *E. coli* K-12 MG1655 and F-plasmid, as well as modified FASTAs containing the mutations found in genomics experiments (in total 4413 entries) plus frequently detected contaminants. For the main search, Lys8 and Arg10 were set as heavy labels of which a maximum of 3 per peptide were allowed; missed cleavages were set to 2 using trypsin as protease with restraint on N-terminally proline-containing peptides. Allowed variable modification included N-terminal acetylation, methionine-oxidation and replacement of tryptophan by fluorotryptophan (mass shift of +17.9905781679 Da). The re-quantify option for quantitation of missing SILAC-pairs was enabled. Protein FDR for identification was set to 0.01. Matching between runs was enabled with standard settings. Minimum of labeled SILAC-pairs for quantitation of protein ratio was set to 1 using unique and razor peptides employing the advanced ratio estimation option of MaxQuant. All other settings were kept default. Further analysis was conducted using Perseus 1.5.7.0. The data was cleared from decoy hits, proteins identified by site only and contaminants first. Then, the data was transformed to log2 ratios of light to heavy isotopic species (L/H) and the technical replicates of SILAC experiment TUB00 vs. TUB00-dKO were averaged using the median. Moreover, the data was filtered to contain a minimum of three valid values for each protein within an individual SILAC experiment. A two-sided two-sample t-test was performed comparing SILAC experiment TUB00 vs. TUB00-dKO with ALE isolates TUBn vs. TUB00-dKO to check for changes in the evolved strains. Artificial variance was set to a value of 2.5 and permutated FDR to 0.01. Only proteins found to exceed a combination of distinct minimum value of log2-fold change and *p*-value from t-test were assigned as significant (as visualized in volcano plots, see Fig.S4).

**10. Sample preparation for non-targeted metabolomics and MS data acquisition**

TUB00 and the isolates from early, intermediate and final time points of 4-fluoroindole (4TUB34, 4TUB81, 4TUB93) and 5-fluoroindole (5TUB23, 5TUB48, 5TUB83) ALE experiments were revitalized from cryostocks and used to inoculate 10 ml of the corresponding ALE medium (see Tables S3-S4). Biological triplicates of each isolate and TUB00 were prepared and incubated at 30 °C, 180 rpm, until they reached the exponential phase of growth ($OD_{600}$ 0.6 – 0.8), then an aliquot was used to inoculate (1:100) 200 ml of fresh medium. The cultures were incubated at 30 °C, 180 rpm and subsequently harvested at the early stationary phase ($OD_{600}$ 0.8 - 1) by 5 min centrifugation at 3,000 g, 4 °C. The metabolism was quenched by resuspending wet cell mass (wcm) in -80 °C cold methanol ad 250 mg/ml. Subsequently, 50 µl (corresponding to 15 mg wcm) of each sample were resuspended in 450 µl -80 °C methanol in a 96 Deepwell plate, sonicated 10 minutes and mixed by shaking on a tilting plate for 20 minutes at room temperature. Insoluble cell debris was pelleted by centrifugation, 10 min at 13,000 g. 450 µl of supernatant containing cytosolic metabolites was transferred to a new Deepwell plate and dried in a CentriVap Benchtop Vacuum Concentrator (LabConco, USA). The metabolites were then solved in 100 µl 50% methanol, 1% formic acid and submitted for MS measurement. Non-targeted LC-MS/MS analysis of the extracted metabolites was performed on a Q-Exactive Orbital IonTrap coupled to UHPLC System in Data Dependent Acquisition mode. The samples were first subjected chromatographic separation (UHPLC), during which the metabolites were eluted from a C18 core-shell column (Kinetex, 50 x 2.1 mm, 1.8 µm particle size, 100 A pore size, Phenomenex, Torrance, USA) with a linear gradient of water (A) and acetonitrile (B), both with 0.1% formic acid. The gradient started at 3% B for 1 min and then increased to 5% B over 1 min. From 2 to 8 min the mobile phase composition was increased to 60% B and then ultimately to 99% B at 10 min. After a 2 min washout phase at 99% B the column was re-equilibrated at 3 % B. Subsequently, the eluting metabolites were analyzed in positive electrospray MS/MS data depended acquisition (DDA) mode. ESI parameters were set to 35 L/min sheath gas flow, 10 L/min auxiliary gas flow, 2 L/min sweep gas flow and 400 °C auxiliary gas temperature. The spray voltage was set to 3.5 kV and the inlet capillary was set to 250 °C. S-lens voltage of 50 V was applied.

The acquisition mass range was set to 100 - 1500 m/z with a resolution of R=17,000 at 200 m/z with 1 micro scan and automatic gain control (AGC) of 5e5 and maximum C-trap fill time was set to 100 ms. Top5 of the most abundant peaks per precursor scan were submitted for MS/MS. Therefore, the quadrupole precursor selection windows were set to 1 m/z. Normalized collision energy was stepwise increased from 20 to 30 to 40 % with z = 1 as default charge state. MS/MS scans were automatically triggered at the apex of chromatographic peaks within 2 to 15 s from their first occurrence. Dynamic exclusion was set to 5s. Ion species with unassigned charge states and isotope peaks were excluded from MS/MS acquisition.

### 11. MS data analysis with Global Natural Products Social Molecular Networking (GNPS)

Raw spectra were converted to .mzXML GNPS-compatible format (as on documentation at https://gnps.ucsd.edu) using the software MSconvert (ProteoWizard). MS1 and MS2 feature detection was performed with the software Mzmine2.28. Metabolites were separated by means of their characteristic m/z ratio and retention time. Extracted Ion Chromatograms (XICs) of MS1 data were built and chromatographic peaks were assigned to all detected ions. The area of each ion peak in the different samples (i.e. metabolite abundance) was compared and aligned into a matrix. Subsequently, MS2 features were detected and correlated to the aligned MS1 features. MS1 features without MS2 features assigned were filtered out and the resulting matrix was exported as .csv file. Features that were less than 2-times more abundant in the samples than in the blank were manually removed and not considered for further analysis. A signal cut-off was set to $1 \cdot 10^5$ to eliminate noise signals generated during gap-filling, given that the Total Ion Current (TIC) of the samples was in the range of $10^{10}$. The matrix was exported as .mgf file and MS2 features were submitted for network analysis and database dereplication to GNPS. The Cosine Score Threshold (minimum value to define a correlation between features based on spectral similarity) was set to 0.7, the Precursor Ion Mass Tolerance to 0.01 Da, Fragment Ion Mass Tolerance to 0.01 Da, Minimum Matched Fragment Ions to 4, Minimum Cluster Size to 1 (MS Cluster off) and Library Search Minimum Matched Peaks to 4. When Analog Search was performed, the Cosine Score Threshold was se to 0.7 and the Maximum Analog Search Mass Difference was 100. Correlated features were clustered into “molecular subnetworks”, which were visualized with Cytoscape. Each MS2 feature was displayed as a node and nodes with spectral similarity were connected by edges, whose thickness is proportional to the cosine score. Multivariate statistical analysis was performed in order to determine relationships between the analyzed metabolomes. In particular, principal coordinates analysis (PCoA) was performed on the metabolome abundance matrix with the UCSD in-house tool ClusterApp using the Canberra distance metric and the grouping method HCA. The results were visualized in EMPeror.

After annotation, the abundances of redundant molecular features (e.g. different ion adducts and in source fragments annotated by GNPS as the same metabolite) was determined by summing the individual XIC peak areas and normalizing to the total abundance of features detected in each metabolome sample (TIC). For the tryptophan and lipids-related subnetworks (Fig. 3, main text), the average across the triplicate metabolomes was calculated. The abundances cutoff was set at 0.00001 for features to be considered significant. Overall, 866 "features" (i.e., molecules with unique m/z and retention time identifiers) were detected in TUB00, 4TUB and 5TUB. Based on the similarity of MS2 spectra, 636 features were clustered by GNPS into 91 subnetworks of chemically related molecules, while 230 molecules with unique spectral signatures were classified as "self-loops".

#### 12. Cell imaging by fluorescence microscopy

TUB00 and the final time points of the ALE experiments (4TUB93 and 5TUB83) were cultivated in the corresponding ALE medium (TUB00, NMM(19/0/30); ALE isolates, NMM(0/30/0) supplemented with either 4- or 5-fluoroindole), at 30 °C, 180 rpm. Cells in the exponential phase of growth ($OD_{600}$ 0.6 - 0.8) were used to inoculate (1:100) 5 ml cultures, which were then incubated overnight at 30 °C and 180 rpm shaking. Subsequently, cells were harvested by centrifugation (5,000 g, 1 min, 4 °C) and resuspended in 0.1 M $Na_2HPO_4$ PBS buffer pH 7.4. This step was repeated twice in order to remove the cultivation medium, then $OD_{600}$ was adjusted to 1.

Nile Red was solved in DMSO ad 10 mM, added 1:1,000 to the cell suspensions (final concentration 10 µM) and samples were incubated 5 min at room temperature, in the dark. A 30 µl aliquot was spotted onto a glass slide and incubated further 5 minutes in the dark, in order to let the cells deposit onto the visible surface. Fluorescence was excited at 530 nm and a 560 nm detection filter was applied. Images were recorded on a Nikon Eclipse Ti microscope (Nikon, Japan). Image processing was performed with ImageJ (NIH, USA) software.

#### 13. Cell membrane permeability assay with vancomycin

The same procedure as for MIC measurement (Method 4) was followed. As solely difference, cultures and vancomycin dilutions (25-400 ng/ml) were not prepared in rich LB medium but in ALE media (TUB00, NMM(19/0/30); 4TUB93 and 5TUB83, NMM(0/30/0) supplemented with either 4- or 5-fluoroindole) which reproduced the conditions of the evolution experiment. Triplicate cultures were incubated at 30 °C, 180 rpm until the optical density corresponding to 0.5 McFarland standard was reached. Dilutions of vancomycin (25 - 400 ng/ml) were distributed in a 96-wells plate, inoculated (1:100) with cell cultures to the final volume of 200 µl and incubated for 16 hours at 30 °C, 180 rpm shaking. $OD_{600}$ was measured with a Tecan Infinite Microplate Reader (Männedorf, Switzerland).

### Supplementary Figures

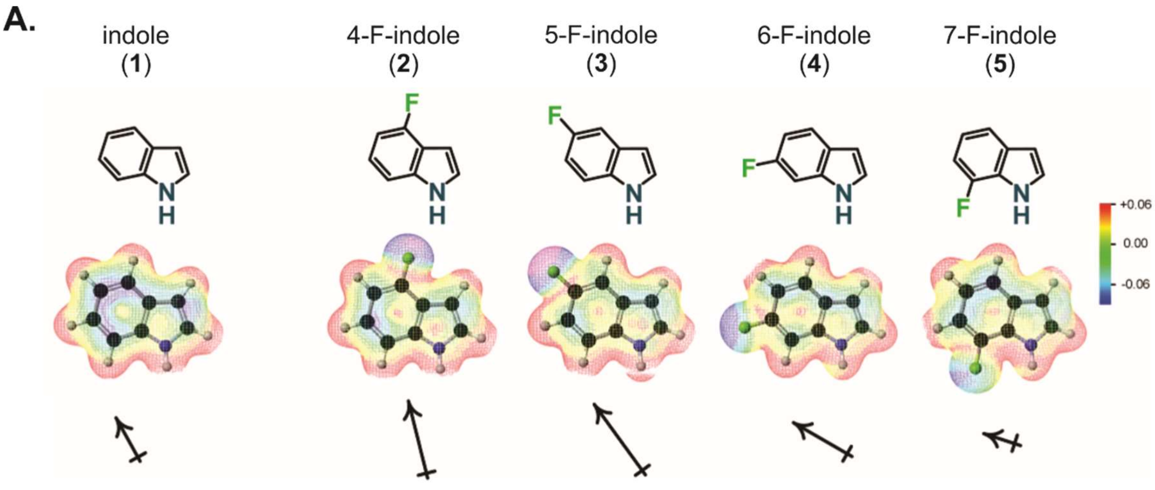

**B.**

| Substitution | Size [Å] | Lipophilicity [Log*P*] | Dipole moment [D] |
|---|---|---|---|
| (none) | 147 | 2.20 ± 0.04 | 2.24 |
| 4-F | 156 | 2.58 ± 0.13 | 3.59 |
| 5-F | 156 | 2.56 ± 0.07 | 3.93 |
| 6-F | 156 | 2.61 ± 0.06 | 3.05 |
| 7-F | 156 | 2.66 ± 0.05 | 1.74 |

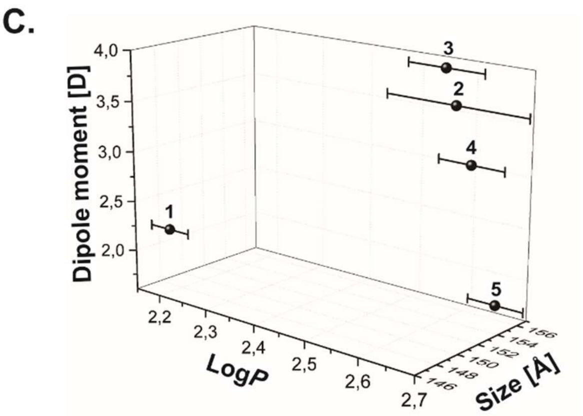

**Fig. S1.** Structure, size, lipophilicity (log*P*, octanol/water partition) and dipole moment (D) of 4-, 5-, 6-, 7-fluoroindole in comparison to indole. (**A**) The heat-map represents distribution of electron density across each molecule in a scale from blue (high electron density) to red (low electron density). The arrows below indicate the direction of the dipole moment and the size is proportional to the dipole magnitude (D). Table (**B**) and scatter plot (**C**) summarize the correlation between size, lipophilicity and dipole moment of indole and fluoroindoles.

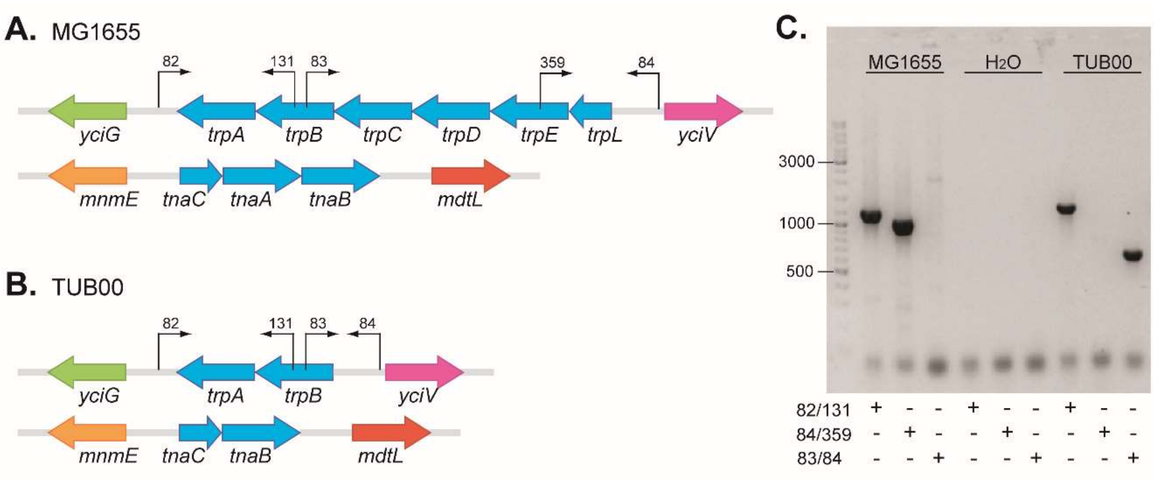

**Fig. S2.** Overview of *E. coli* genotype with regards of *trp* operon (*trpLEDCBA*) and tryptophanase locus (*tnaBAC*) (**A**) in the parent strain MG1655 and (**B**) in the derivative TUB00 after gene knock-out (*ΔtrpLEDC*, *ΔtnaA*); (**C**) *trp* operon amplification by colony PCR. The DNA ladder indicates the size of amplicons in kDa and the primer mixes reported below refer to the annealing regions reported in panels A and B.

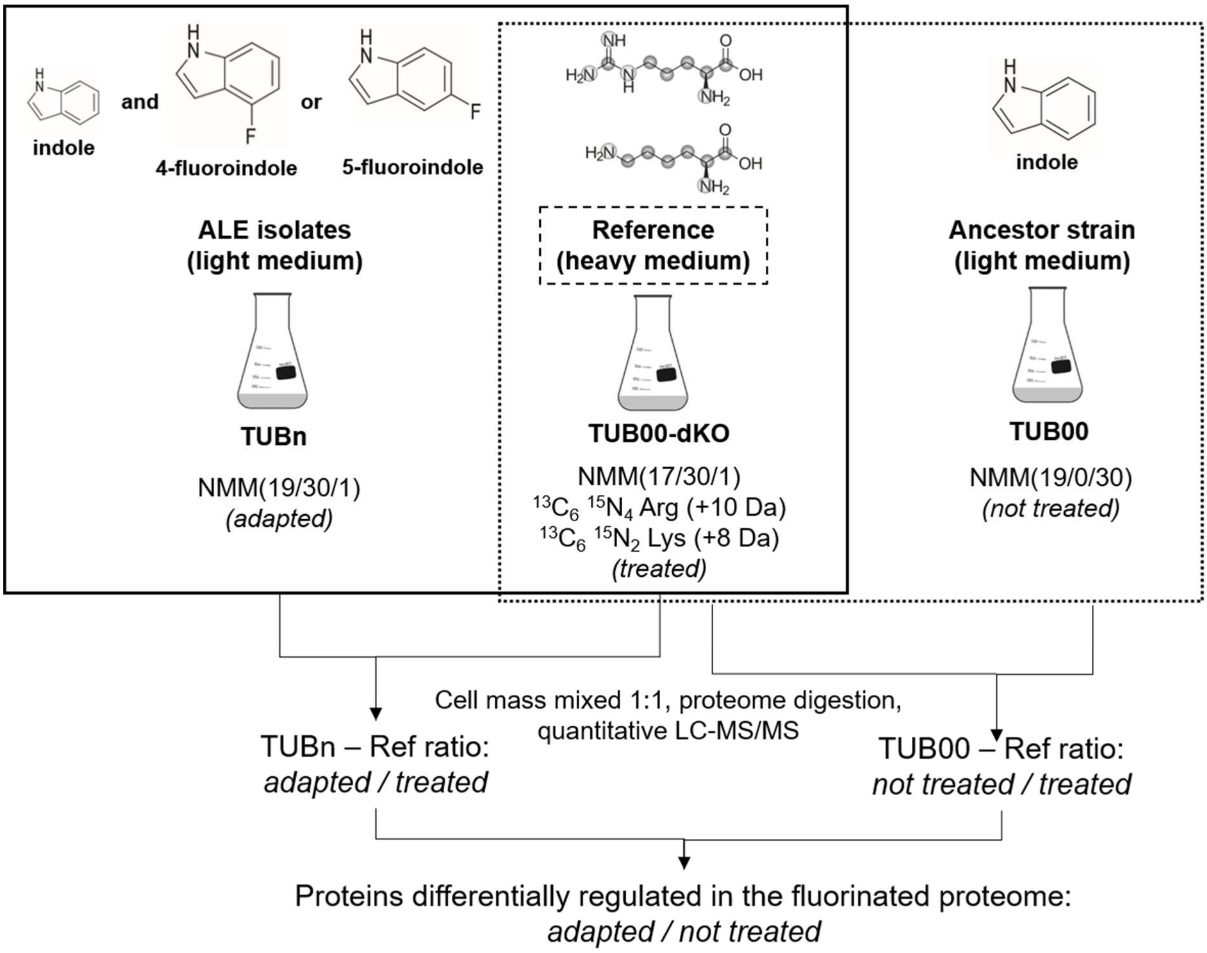

**Fig. S3.** SILAC experimental setup. Isolates from early- and final time-points of 4TUB (4TUB34 and 4TUB93, respectively) and 5TUB (4TUB23 and 5TUB83, respectively) were cultivated in “light” (L, not isotope-labeled) minimal medium NMM19/30/1 containing all amino acids except tryptophan, 30 μM of either 4- or 5-fluoroindole and 1 μM indole. The reference strain for quantification TUB00-dKO (i.e. TUB00 rendered genetically auxotrophic for biosynthesis of Lys and Arg) was cultivated in “heavy” (H, isotope-labelled) minimal medium NMM(17/30/1) supplemented with $^{13}C_6{}^{15}N_4$ Arg, $^{13}C_6{}^{15}N_2$ Lys, 30 μM of either 4- or 5-fluoroindole and 1 μM indole.

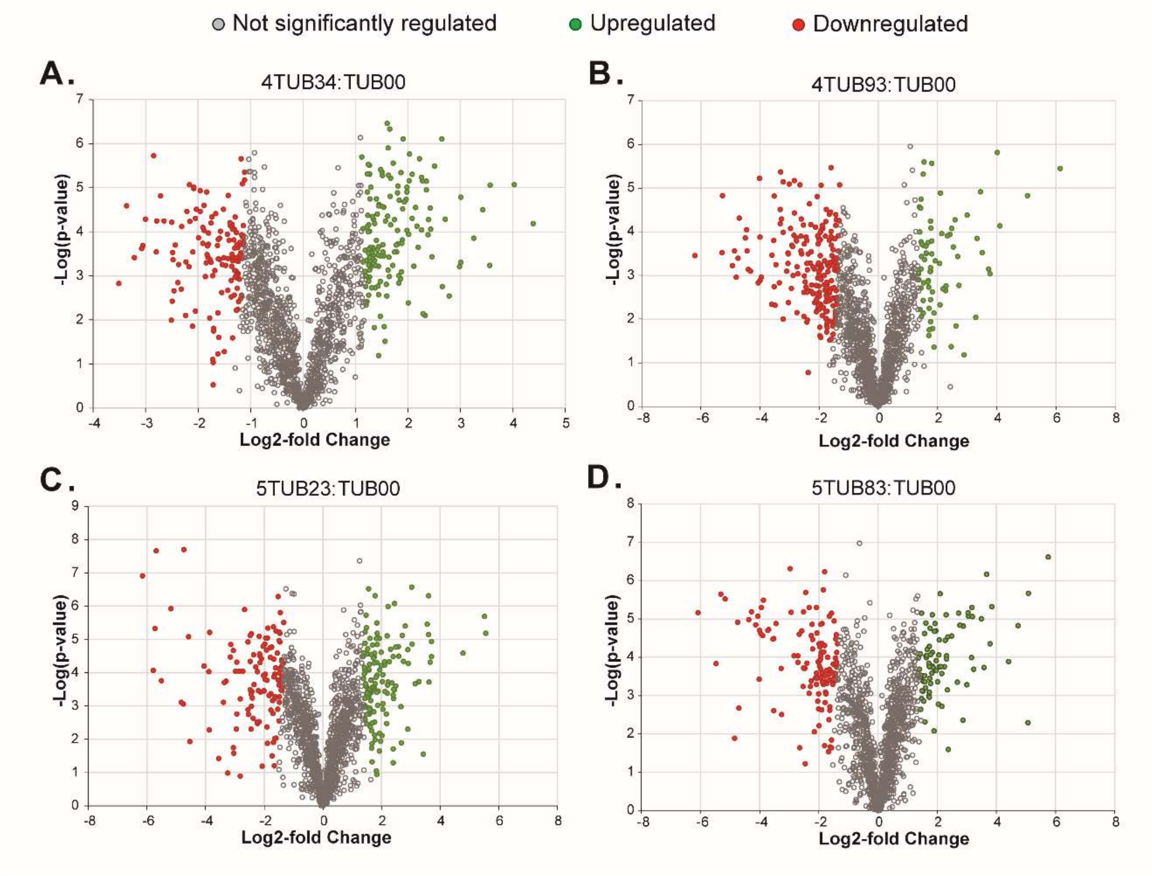

**Fig. S4.** Volcano plots of SILAC experiments showing changes in protein abundance between the ALE isolates and TUB00. The *adapted/treated* SILAC ratios of isolates (NMM(19/30/1)) from early- and final time-points of the adaptation towards (**A**, **B**) 4-fluoroindole (4TUB34 and 4TUB93, respectively) and (**C**, **D**) 5-fluoroindole (5TUB23 and 5TUB83, respectively) were compared to the H *not treated/treated* L ratios of TUB00 (NMM(19/0/30)). Red: downregulated proteins (less abundant in ALE isolates than in TUB00); green: upregulated proteins (more abundant in ALE isolates than in TUB00); grey: protein with same abundance in the ALE isolates and in TUB00.

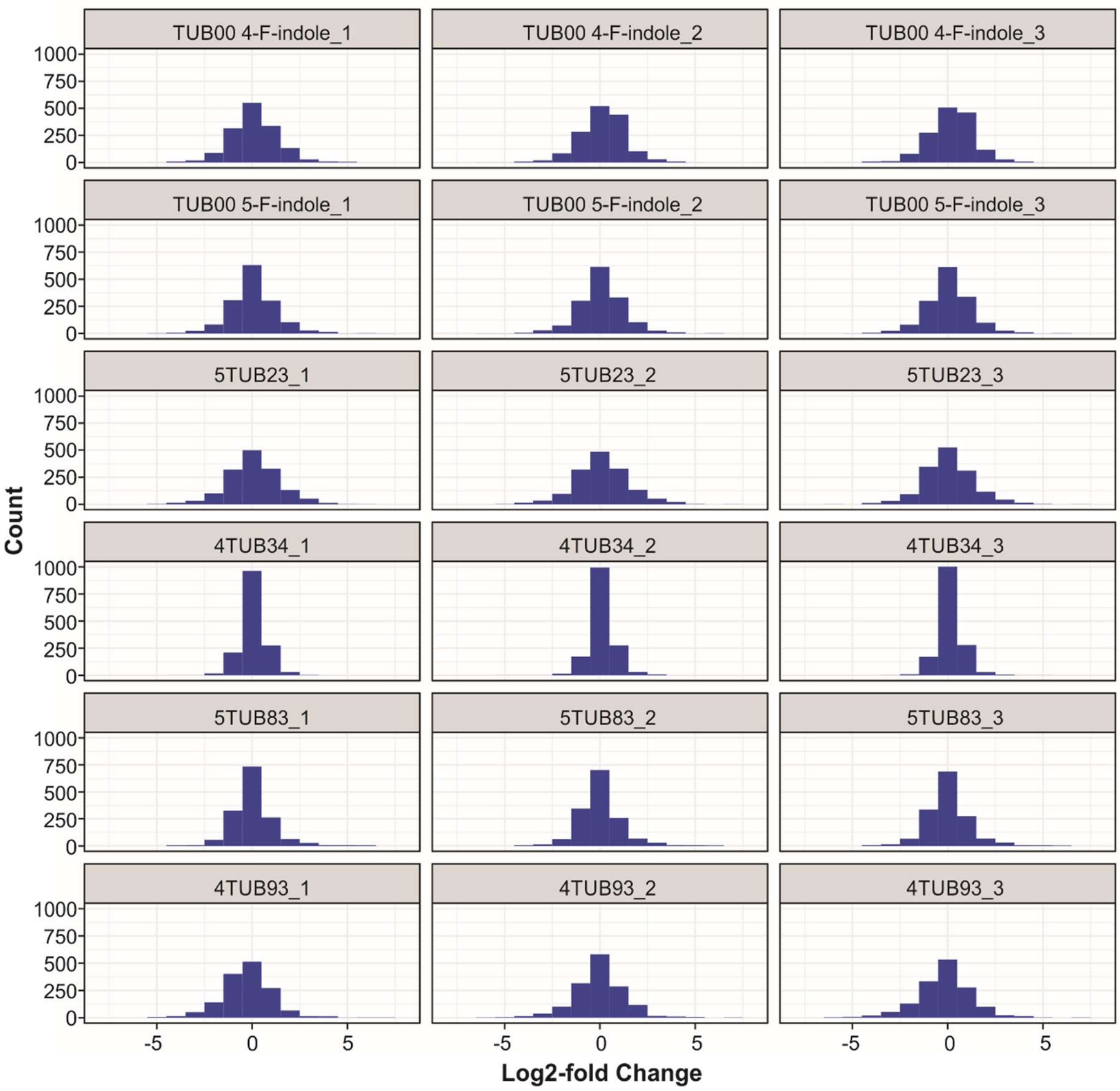

**Fig. S5.** Histograms of the log2-fold Change ratios of all proteins detected during the SILAC experiment. Results are shown for TUB00 (NMM(19/0/30)) in comparison to TUB00-dKO grown in NMM(17/30/1) supplemented with $^{13}C_6{}^{15}N_4$ Arg and , $^{13}C_6{}^{15}N_2$ Lys and either 4- or 5-fluoroindole; 4TUB34 and 4TUB93 (NMM(19/30/1) 4-fluoroindole) and 5TUB23 and 5TUB83 (NMM(19/30/1) 5-fluoroindole) in comparison to TUB00-dKO (NMM(17/30/1) supplemented with $^{13}C_6{}^{15}N_4$ Arg and $^{13}C_6{}^{15}N_2$ Lys and the corresponding fluoroindole.

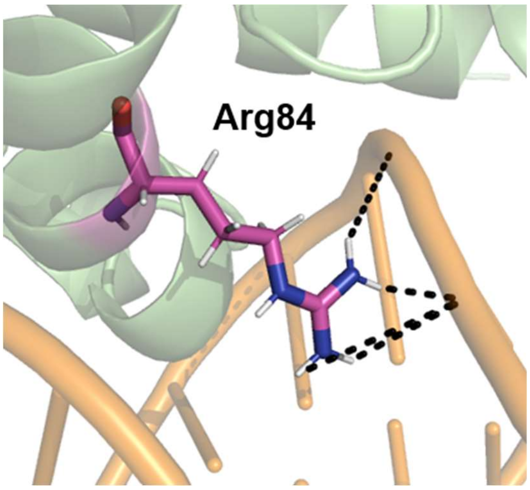

**Fig. S6.** Interaction of TrpR Arg84 with DNA. C atoms of Arg84 are represented in pink, O in red, N in blue and H in white color; double stranded DNA is showed orange. Arg positively charged guanidinium group interacts with the negatively charged sugar-phosphate backbone of DNA. The figure was generated with Pymol (DeLano Scientific LLC, Schrödinger LLC) from the structure 1WRT deposited in the RCSB Protein Data Bank (RCSB PDB, www.rcsb.org)

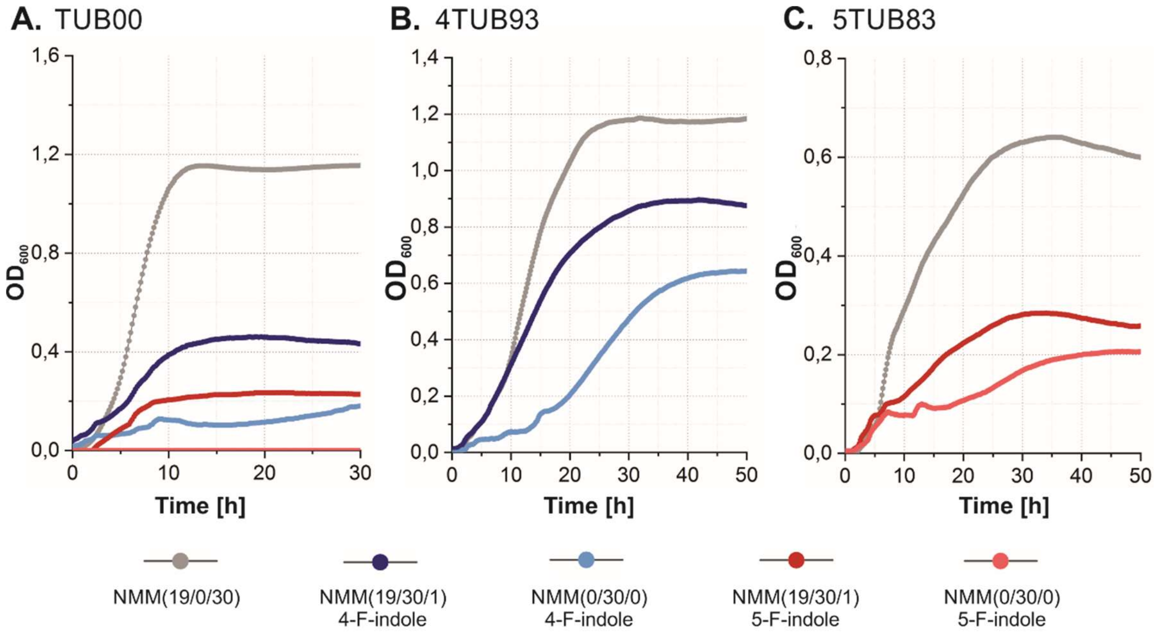

**Fig. S7.** Growth curves of (**A**) TUB00 and isolates from final time points of (**B**) 4-fluoroindole (4TUB93) and (**C**) 5-fluoroindole (5TUB83) ALE measured at 30 °C in different media, including NMM(19/0/30) supplemented with indole; NMM(19/30/1) supplemented with indole and either 4- or 5-fluoroindole; NMM(0/30/0) supplemented either 4- or 5-fluoroindole. Biological triplicates (of 4TUB93 and 5TUB83) and duplicates (of TUB00) were cultivated in a 96-wells plate and cell density was continuously measured with a microplate reader. For generation times, see Table S12. The optical density ($OD_{600}$) values are not comparable to those measured during ALE experiment (Fig. 2 in main text) due to different cultivation setups (shake flasks and microplates).

### Supplementary tables

**Table S1.** Kinetic parameters relative to AMP adenylation (amino acid activation) of tryptophan, 4- and 5-fluorotryptophan (Trp, 4-F-Trp and 5-F-Trp, respectively) by *E. coli* tryptophanyl-tRNA synthetase (TrpRS) as measured by ATP/[$^{32}$P]pyrophosphate exchange assay. Calculated Michaelis-Menten constant ($K_M$), turnover number ($k_{cat}$) kinetic efficiency ($K_M/k_{cat}$) and kinetic efficiency relative to the reaction with the natural substrate Trp ($K_M/k_{cat}$) are shown.

| Substrate | $K_M$ [μM] | $k_{cat}$ [$s^{-1}$] | $k_{cat}/ K_M$ [L·(mmol·s)$^{-1}$] | Rel $k_{cat}/ K_M$ |
|---|---|---|---|---|
| Trp | 3 ± 1 | 1.46 ± 0.10 | 466.91 ± 0.12 | 1 |
| 4-F-Trp | 7 ± 2 | 0.68 ± 0.04 | 106.56 ± 0.05 | 0.23 |
| 5-F-Trp | 44 ± 16 | 0.40 ± 0.03 | 9.03 ± 0.03 | 0.02 |

At concentrations 3-5 mM, indole is known to inhibit growth of *E. coli* and this effect was expected to be stronger in the case of the fluoroindoles. The tolerance of TUB00 towards indole, 4- and 5-fluoroindole was assessed by measuring the minimal inhibitory concentration (MIC).

**Table S2.** Minimal Inhibitory Concentration (MIC) measurement of indole, 4- and 5-fluoroindole for TUB00 and the final time points of 4- and 5-fluoroindole ALE (4TUB93 and 5TUB83, respectively). MICs were measured in lysogeny broth (LB) to estimate the growth-inhibiting effects of the three compounds prior and after the ALE experiment.

| | TUB00 | 4TUB93 | 5TUB83 |
|---|---|---|---|
| Indole | 10 mM | 10 mM | 7.5 mM |
| 4-fluoroindole | 2.5 mM | 2.5 mM | 1.25 mM |
| 5-fluoroindole | 1.88 mM | 3.75 mM | 3.75 mM |

The TUB00 resulted very tolerant towards indole (MIC 10 mM), while 4-times less tolerant towards 4-fluoroindole (MIC 2.5 mM) and about 5-times less towards 5-fluoroindole (MIC 1.88 mM). The 4-fluoroindole-adapted strain showed improved tolerance for 5-fluoroindole in comparison to TUB00, while the 5-fluoroindole-adapted strain is less tolerant against indole.

**Table S3.** New minimal medium (NMM) composition during 4-fluoroindole ALE. NMM(*a*/*b*/*c*) where *a* is the number of amino acids supplied, *b* is μM concentration of 4-fluoroindole and *c* is μM concentration of indole.

| Medium | Amino acids supplied | Indole [μM] | 4-fluoroindole [μM] |
|---|---|---|---|
| NMM19a | Tyr, Phe, Cys, Gly, Ser, Leu, Val, Ile, Ala, Asp, Glu, His, Met, Asn, Pro, Gln, Arg, Thr, Lys | 0.5 | 70 |
| NMM19b | *v.s.* | 0.1 | 70 |
| NMM19c | *v.s.* | 0.05 | 70 |
| NMM19d | *v.s.* | 0.01 | 70 |
| NMM19e | *v.s.* | 0.005 | 70 |
| NMM19f | *v.s.* | 0 | 70 |
| NMM19g | *v.s.* | 0 | 30 |
| NMM17 | Cys, Gly, Ser, Leu, Val, Ile, Ala, Asp, Glu, His, Met, Asn, Pro, Gln, Arg, Thr, Lys | 0 | 30 |
| NMM10 | Asp, Glu, His, Met, Asn, Pro, Gln, Arg, Thr, Lys | 0 | 30 |
| NMM9 | Glu, His, Met, Asn, Pro, Gln, Arg, Thr, Lys | 0 | 30 |
| NMM8 | Glu, His, Met, Pro, Gln, Arg, Thr, Lys | 0 | 30 |
| NMM7 | Glu, His, Met, Pro, Gln, Arg, Thr | 0 | 30 |
| NMM6 | Glu, His, Met, Gln, Arg, Thr | 0 | 30 |
| NMM5 | Glu, His, Met, Arg, Thr | 0 | 30 |
| NMM4 | His, Met, Arg, Thr | 0 | 30 |
| NMM3 | His, Met, Arg | 0 | 30 |
| NMM2 | His, Met | 0 | 30 |
| NMM1 | Met | 0 | 30 |
| NMM0 | - | 0 | 30 |

**Table S4.** New minimal medium (NMM) composition during 5-fluoroindole ALE. NMM(*a*/*b*/c*)* where *a* is the number of amino acids supplied, *b* is µM concentration of 5-fluoroindole and *c* is µM concentration of indole.

| Medium | Amino acids supplied | Indole [µM] | 5-fluoroindole [µM] |
|---|---|---|---|
| NMM19a | Tyr, Phe, Cys, Gly, Ser, Leu, Val, Ile, Ala, Asp, Glu, His, Met, Asn, Pro, Gln, Arg, Thr, Lys | 1 | 30 |
| NMM19b | *v.s.* | 0.1 | 30 |
| NMM19c | *v.s.* | 0.05 | 30 |
| NMM19d | *v.s.* | 0.01 | 30 |
| NMM19e | *v.s.* | 0.005 | 30 |
| NMM19f | *v.s.* | 0 | 30 |
| NMM17 | Cys, Gly, Ser, Leu, Val, Ile, Ala, Asp, Glu, His, Met, Asn, Pro, Gln, Arg, Thr, Lys | 0 | 30 |
| NMM13 | Cys, Gly, Ser, Asp, Glu, His, Met, Asn, Pro, Gln, Arg, Thr, Lys | 0 | 30 |
| NMM10 | Asp, Glu, His, Met, Asn, Pro, Gln, Arg, Thr, Lys | 0 | 30 |
| NMM2 | Asp, Glu | 0 | 30 |
| NMM1 | Glu | 0 | 30 |
| NMM0 | - | 0 | 30 |

**Table S5.** Number of genomic mutations detected at early-, intermediate- and final time-points of 4-fluoroindole ALE (4TUB34, 4TUB81 and 4TUB93) and of 5-fluoroindole ALE (5TUB23, 5TUB48 and 5TUB83). Abbreviations: Single nucleotide polymorphisms (SNPs), coding DNA sequence (CDS), insertions and deletions (InDel).

| | 4TUB34 | 4TUB81 | 4TUB93 | 5TUB23 | 5TUB48 | 5TUB83 |
|---|---|---|---|---|---|---|
| Total SNPs | 12 | 38 | 32 | 8 | 16 | 36 |
| SNPs in CDS | 11 | 31 | 31 | 7 | 11 | 31 |
| Synonymous SNPs | 0 | 3 | 3 | 1 | 0 | 1 |
| Nonsynonymous SNPs | 10 | 27 | 28 | 6 | 10 | 28 |
| SNPs in RNA | 0 | 1 | 0 | 0 | 0 | 0 |
| Intergenic SNPs | 1 | 7 | 1 | 1 | 5 | 5 |
| InDel in CDS | 0 | 0 | 0 | 2 | 3 | 3 |
| Insertions | 2 | 2 | 2 | 3 | 2 | 2 |
| Deletions | 0 | 0 | 0 | 1 | 2 | 2 |
| Premature stop | 1 | 1 | 1 | 2 | 2 | 2 |
| Frameshift | 0 | 0 | 0 | 2 | 3 | 3 |

**Table S6.** Effect of genomic mutations on protein functions as predicted by PROVEAN (Protein Variation Effect Analyzer) v1.1, an alignment-based software that assigns scores to genetic variations based on the assumption that alterations of evolutionary conserved regions are deleterious. A score threshold of < -2.5 was used.

| Gene | 4TUB | PROVEAN prediction | 5TUB | PROVEAN prediction |
|---|---|---|---|---|
| *trpS* | Glu15Asp | -1,361: neutral | Met187Leu | -2.996: deleterious |
| *trpR* | - | - | Arg84Cys | -7.994: deleterious |
| *ptsG* | - | - | Ile25-FS | -11.092: deleterious |
| *mtr* | - | - | Ser94Pro | -4.077: deleterious |
| *mdtF* | Asp759Gly | -6,412: deleterious | - | - |
| *mdtK* | - | - | Gln305Stop | -9.306: deleterious |
| *mdtO* | - | - | Ile104Ser | -3.125: deleterious |
| *rpsA* | Lys158Gln | -3,978: deleterious | Gln355Pro | -2.560: deleterious |
| *rpsJ* | Thr28Ala | -4,220: deleterious | - | - |
| *ileS* | - | - | Tyr727Ser | -8.569: deleterious |
| *glyQ* | Glu48Ala | 4,163: neutral | - | - |
| *ptrA* | - | - | Phe827Cys | -3.373: deleterious |
| *ftsH* | - | - | Leu563Arg | -4.556: deleterious |
| *cpxA* | Glu355Gly | -6,558: deleterious | Tyr364Ser | -8.950: deleterious |

**Table S7.** Overview of changes in protein abundance found in the comparison of adapted/not treated SILAC ratios. The isolates from the early (4TUB34, 5TUB23) and final (4TUB93, 5TUB83) time points of ALE (NMM(19/30/1) supplemented with either 4- or 5-fluoroindole) were compared to TUB00 (NMM(19/0/30)). Proteins significantly more abundant in the adapted strains than in TUB00 are classified as upregulated; proteins less abundant in the adapted strains than in TUB00 are classified as downregulated. 2439 proteins were identified and 1501 SILAC ratios were quantified. The complete list is provided in supplementary tables.

| | Significant | Upregulated | Downregulated | Not significant |
|---|---|---|---|---|
| 4TUB34 | 289 | 151 | 138 | 1212 |
| 4TUB93 | 287 | 75 | 212 | 1214 |
| 5TUB23 | 264 | 141 | 123 | 1237 |
| 5TUB83 | 209 | 83 | 126 | 1292 |

**Table S8.** "Library hit" metabolites annotated by Global Natural Products Social Molecular Networking (GNPS). Abbreviations include: (**1**) Trp, tryptophan; (**2**) F-Trp, fluorotryptophan; (**3**) PE(16:0/18:1), 1-palmitoyl-2-oleyl-sn-glycero-3-phosphoethanolamine; (**4**) PE(16:1/0:0), 1-palmitoleoyl-sn-glycero-3-phosphoethanolamine; (**5a**) PE(18:0/0:0), 1-oleyl-2-hydroxy-sn-glycero-3-phosphoethanolamine; (**5b**) PE(16:0/0:0), 1-palmitoyl-2-hydroxy-sn-glycero-3-phosphoethanolamine; (**6**) DG(18:1/2:0/0:0), 1-oleoyl-2-acetyl-sn-glycerol; (**7**) MG(16:0/0:0/0:0), monopalmitolein; (**8**) biotin.

| Annotation | m/z | RT [min] | Adduct | Mass Diff [m/z] | Cosine score |
|---|---|---|---|---|---|
| (**1**) Trp | 188.0708 | 1.31 | $[M+H-NH_3]^+$ | 0.0008 | 0.9879 |
| (**1**) Trp | 205.0967 | 1.31 | $[M+H]^+$ | 0.0003 | 0.9971 |
| (**2**) F-Trp | 223.0889 | 2.49 | $[M+H]^+$ | 0.0009 | 0.9984 |
| (**2**) F-Trp | 223.0890 | 2.40 | $[M+H]^+$ | 0.0010 | 0.9987 |
| (**2**) F-Trp | 223.0893 | 2.61 | $[M+H]^+$ | 0.0013 | 0.9718 |
| (**3**) PE(16:0/18:1) | 718.54 | 10.30 | $[M+H]^+$ | 0.0000 | 0.9000 |
| (**3**) PE(16:0/18:1) | 718.54 | 12.70 | $[M+H]^+$ | 0.0000 | 0.8900 |
| (**3**) PE(16:0/18:1) | 718.54 | 10.14 | $[M+H]^+$ | 0.0000 | 0.9000 |
| (**5a**) PE(18:0/0:0) | 482.32 | 9.06 | $[M+H]^+$ | 0.0000 | 0.8600 |
| (**5a**) PE(18:0/0:0) | 480.31 | 8.51 | $[M+H]^+$ | 0.0000 | 0.9500 |
| (**5b**) PE(16:0/0:0) | 454.29 | 8.19 | $[M+H]^+$ | 0.0000 | 0.8900 |
| (**5b**) PE(16:0/0:0) | 476.27 | 8.33 | $[M+Na]^+$ | 0.0100 | 0.9700 |
| (**5b**) PE(16:0/0:0) | 454.29 | 8.33 | $[M+H]^+$ | 0.0000 | 0.8800 |
| (**6**) DG(18:1/2:0/0:0) | 339.29 | 8.38 | $[M+H-H_2O]^+$ | 0.0000 | 0.9800 |
| (**6**) DG(18:1/2:0/0:0) | 339.29 | 8.56 | $[M+H-H_2O]^+$ | 0.0000 | 0.9800 |
| (**7**) MG(16:1/0:0/0:0) | 311.26 | 7.80 | $[M+H-H_2O]^+$ | 0.0000 | 0.9500 |
| (**7**) MG(16:1/0:0/0:0) | 311.26 | 7.68 | $[M+H-H_2O]^+$ | 0.0000 | 0.9500 |
| (**8**) Biotin | 245.10 | 3.60 | $[M+H]^+$ | 0.0000 | 1.0000 |
| (**8**) Biotin | 489.18 | 3.61 | $[2M+H]^+$ | 0.0000 | 0.9800 |

**Table S9.** “Analog hit” metabolites annotated by GNPS. For abbreviations, see Table S8. The features with m/z 206.06 were originally annotated by GNPS as 5-hydroxy-fluoro-tryptophan. However, these were later recognized as degradation products of fluorotrytophan, resulting from in-source deamination, in analogy to Trp (see Table S8).

| Annotation | m/z | RT [min] | Adduct | Mass Diff [m/z] | Cosine score |
|---|---|---|---|---|---|
| (**2**) F-Trp | 206.0614 | 2.39 | $[M+H-NH_3]^+$ | 17.0270 | 0.9779 |
| (**2**) F-Trp | 206.0615 | 2.46 | $[M+H-NH_3]^+$ | 17.0260 | 0.9777 |
| (**2**) F-Trp | 206.0615 | 2.61 | $[M+H-NH_3]^+$ | 17.0260 | 0.9804 |
| (**2**) F-Trp | 237.1065 | 4.09 | $[M+H+CH_3]^+$ | 14.0185 | 0.9178 |
| (**4**) PE(16:1/0:0) | 483.27 | 7.80 | - | 32.0000 | 0.7900 |
| (**4**) PE(16:1/0:0) | 500.30 | 7.79 | - | 49.0300 | 0.8000 |
| (**5a**) PE(18:0/0:0) | 440.28 | 7.96 | - | 40.0300 | 0.9500 |
| (**5a**) PE(18:0/0:0) | 466.29 | 8.21 | - | 14.0100 | 0.9700 |
| (**5a**) PE(18:0/0:0) | 452.28 | 7.81 | - | 28.0300 | 0.9700 |
| (**5b**) PE(16:0/0:0) | 474.26 | 7.81 | - | 2.0200 | 0.9700 |
| (**8**) Biotin | 227.08 | 3.61 | - | 18.0100 | 0.8900 |

**Table S10.** Normalized abundance of selected metabolites annotated by GNPS. All experiments were performed in biological triplicates corresponding to cell isolates before ALE (TUB00) and at early (4TUB34, 5TUB23), intermediate (4TUB81, 5TUB48) and final time points of ALE (4TUB93, 5TUB83). For abbreviations, see Table S8.

| Annotation | TUB00 | 4TUB34 | 4TUB81 | 4TUB93 | 5TUB23 | 5TUB48 | 5TUB83 |
|---|---|---|---|---|---|---|---|
| (**1**) Trp | 0.04110 ± 0.00310 | 0 | 0 | 0 | 0 | 0 | 0 |
| (**2**) F-Trp | 0 | 0.02988 ± 0.00685 | 0.00818 ± 0.00084 | 0.02042 ± 0.00784 | 0.11166 ± 0.07073 | 0.20081 ± 0.01400 | 0.00067 ± 0.00005 |
| (**3**) PE(16:0/18:1) | 0 | 0 | 0.00115 ± 0.00071 | 0.00146 ± 0.00053 | 0.00231 ± 0.00122 | 0.00393 ± 0.00164 | 0.00816 ± 0.00257 |
| (**4**) PE(16:1/0:0) | 0.00002 ± 0.00001 | 0.00005 ± 0.00001 | 0.00009 ± 0.00003 | 0.00009 ± 0.00003 | 0.00090 ± 0.00025 | 0.00051 ± 0.00014 | 0.00170 ± 0.00043 |
| (**5a**) PE(18:0/0:0) | 0.00088 ± 0.00012 | 0.00125 ± 0.00033 | 0.00622 ± 0.00044 | 0.00291 ± 0.00082 | 0.00978 ± 0.00131 | 0.00313 ± 0.00079 | 0.00623 ± 0.00051 |
| (**5b**) PE(16:0/0:0) | 0.00239 ± 0.00019 | 0.00299 ± 0.00095 | 0.00680 ± 0.00032 | 0.00495 ± 0.00194 | 0.01045 ± 0.00200 | 0.00609 ± 0.00105 | 0.01133 ± 0.00138 |
| (**6**) DG(18:1/2:0/0:0) | 0.00006 ± 0.00001 | 0.00002 ± 0.00001 | 0.00018 ± 0.00003 | 0.00008 ± 0.00002 | 0.00082 ± 0.00006 | 0.00026 ± 0.00013 | 0.00113 ± 0.00031 |
| (**7**) MG(16:1/0:0/0:0) | 0.00009 ± 0.00001 | 0.00041 ± 0.00013 | 0.00049 ± 0.00003 | 0.00053 ± 0.00018 | 0.00151 ± 0.00036 | 0.00095 ± 0.00016 | 0.00317 ± 0.00080 |
| (**8**) Biotin | 0.00273 ± 0.00013 | 0.01508 ± 0.00187 | 0.01762 ± 0.00018 | 0.01478 ± 0.00296 | 0.01004 ± 0.00258 | 0.01061 ± 0.00181 | 0.01175 ± 0.00028 |

**Table S11.** Minimal Inhibitory Concentration (MIC) of vancomycin for TUB00 and the final time points 4- and 5-fluoroindole ALE (4TUB93 and 5TUB83, respectively). MIC was measured in ALE medium (4TUB93, NMM(0/30/0) supplemented with 4-fluoroindole; 5TUB83, NMM(0/30/0) supplemented with 5-fluoroindole; TUB00, NMM(19/0/30).

| | TUB00 | 4TUB93 | 5TUB83 |
|---|---|---|---|
| Vancomycin | 200 $\mu g \cdot ml^{-1}$ | 50 $\mu g \cdot ml^{-1}$ | 25 $\mu g \cdot ml^{-1}$ |

**Table S12.** Generation times of TUB00 and the fluoroindole-adapted strains at the final time point of the ALE experiments (4TUB93 and 5TUB83) as calculated from the growth curves measured in 96-well plates at 30 °C in LB and NMM in different supplementation conditions.

| Cultivation medium | TUB00 [min] | 4TUB93 [min] | 5TUB83 [min] |
|---|---|---|---|
| LB | 31 ± 2 | 65 ± 4 | 61 ± 2 |
| NMM(19/0/30) | 126 ± 5 | 189 ± 15 | 171 ± 5 |
| NMM(19/30/1) 4-fluoroindole | 163 ± 6 | 158 ± 15 | NA |
| NMM(19/30/1) 5-fluoroindole | 145 ± 7 | NA | 115 ± 25 |
| NMM(0/30/0) 4-fluoroindole | NA | 278 ± 18 | NA |
| NMM(0/30/0) 5-fluoroindole | NA | NA | 1080 ± 26 |
