## Supplementary material for "Laboratory evolution of *Escherichia coli* enables life based on fluorinated amino acids": Auxiliary Table 1_Genomics

| Genomic position (bp) | Gene | 4TUB34 | 4TUB49 | 4TUB81 | 4TUB93 | 5TUB23 | 5TUB48 | 5TUB83 | Protein name |
|---|---|---|---|---|---|---|---|---|---|
| 209380 | *accA* | A → C (Ile254Leu) | A → C (Ile254Leu) | A → C (Ile254Leu) | A → C (Ile254Leu) | / | / | / | Acetyl-CoA carboxylase carboxytransferase α subunit |
| 3613075 | *acpT* | / | C → G (Ala36Gly) | C → G (Ala36Gly) | C → G (Ala36Gly) | / | / | / | 4'-phosphopantetheinyl transferase AcpT |
| 486024 | *acrR* | / | / | / | / | A → C (Syn) | / | / | HTH-type transcriptional regulator AcrR |
| 3352358 | *arcB* | / | / | / | / | / | T → G (Asn223Thr) | / | Aerobic respiration control sensor protein ArcB |
| 786267 | *aroG* | / | T → C (Val212Ala) | / | / | / | / | / | Phospho-2-dehydro-3-deoxyheptonate aldolase, Phe-sensitive |
| 1788874 | *aroH* | / | / | T → C (Val147Ala) | T → C (Val147Ala) | / | / | / | Phospho-2-dehydro-3-deoxyheptonate aldolase, Trp-sensitive |
| 3518868 | *aroK* | / | / | T → C (Val66Ala) | A → G (Val66Ala) | / | / | / | Shikimate kinase 1 |
| 4103912 | *cpxA* | / | T → C (Glu355Gly) | A → G (Glu355Gly) | T → C (Glu355Gly) | / | / | T → G (Tyr364Ser) | Sensor histidine kinase CpxA |
| 88479 | *cra* | / | / | / | / | / | / | T → G (Leu151Arg) | Catabolite repressor/activator |
| 2141759 | *dcd* | / | / | A → G (Met153Val) | T → C (Met153Val) | / | / | / | dCTP deaminase |
| 1252970 | *dhaR* | / | / | / | / | / | / | T → G (Phe635Leu) | PTS-dependent dihydroxyacetone kinase operon regulatory protein |
| 2702842 | *era* | / | / | / | / | / | / | T → G (Glu182Ala) | GTPase Era |
| 4629304 | *ettA* | / | / | G → T (Glu407Stop) | / | / | / | / | Energy-dependent translational throttle protein EttA |
| 2485362 | *evgS* | A → G (Asp330Gly) | A → G (Asp330Gly) | A → G (Asp330Gly) | A → G (Asp330Gly) | / | / | / | Hybrid sensory histidine kinase in two-component regulatory system with EvgA |
| 4026634 | *fre* | / | / | / | / | Del of T (Leu37 FS) | Del of T (Leu37 FS) | Del of T (Leu37 FS) | NAD(P)H-flavin reductase |
| 4092387 | *frvA* | / | / | T → C (Leu146Pro) | A → G (Leu146Pro) | / | / | / | PTS system fructose-like EIIA component |
| 103998 | *ftsA* | / | / | A → G (Asp6Gly) | A →G (Asp6Gly) | / | / | / | Cell division protein FtsA |
| 3325248 | *ftsH* | / | / | / | / | / | A → C (Leu563Arg) | A → C (Leu563Arg) | ATP-dependent zinc metalloprotease FtsH |
| 3125145 | *glcF* | / | / | A → C (Lys105Asn) | / | / | / | / | Glycolate oxidase iron-sulfur subunit |
| 3725176 | *glyQ* | / | / | A → C (Glu48Ala) | T → G (Glu48Ala) | / | / | / | Gly--tRNA ligase α subunit |
| 24570 | *ileS* | / | / | / | / | / | / | A → C (Tyr727Ser) | Ile--tRNA ligase |
| 4507362 | *insN-2* | / | / | / | / | / | / | A → C (Lys56Gln) | KpLE2 phage-like element |
| 3334127 | *ispB* | A → C (Asn140His) | A → C (Asn140His) | A → C (Asn140His) | A → C (Asn140His) | / | / | / | Octaprenyl diphosphate synthase |
| 1262762 | *ispE* | T → C (Asp39Gly) | / | / | / | / | / | / | 4-diphosphocytidyl-2-C-methylerythritol kinase |
| 3662689 | *mdtF* | / | A → G (Asp759Gly) | A → G (Asp759Gly) | A → G (Asp759Gly) | / | / | / | Multidrug resistance protein MdtF |
| 1744369 | *mdtK* | / | / | / | / | / | / | C → T (Gln305Stop) | Multidrug resistance protein MdtK |
| 4302768 | *mdtO* | / | / | / | / | / | / | A → C (Ile104Ser) | Multidrug resistance protein MdtO |
| 4128311 | *metJ* | / | / | / | T → G (Ile29Leu) | / | / | T → G (Glu56Ala) | Met repressor |
| 668036 | *mrdA* | A → C (Leu61Arg) | A → C (Leu61Arg) | T → G (Leu61Arg) | A → C (Leu61Arg) | A → C (Ile59Ser) | A → C (Ile59Ser) | A → C (Ile59Ser) | Penicillin-binding protein 2 transpeptidase |
| 4587617 | *mrr* | / | / | / | / | G → A (Trp223Stop) | G → A (Trp223Stop) | G → A (Trp223Stop) | Mrr restriction system protein |
| 3305538 | *mtr* | / | / | / | / | A → G (Ser94Pro) | A → G (Ser94Pro) | A → G (Ser94Pro) | Trp-specific transport protein |
| 2710548 | *nadB* | / | / | A → C (Glu43Asp) | / | / | / | / | L-Asp oxidase |
| 1164542 | *nagZ* | / | / | / | A → T (Ile150Phe) | / | / | / | β-hexosaminidase |
| 3370123 | *nanK* | / | / | T → C (Syn) | A → G (Syn) | / | / | / | N-acetylmannosamine kinase |
| 3615316 | *nikB* | / | A → C (Met26Leu) | / | / | / | / | / | Nickel transport system permease protein NikB |
| 2347844 | *nrdB* | T → C (Ile154Thr) | A → G (Ile61Val) | T → C (Ile154Thr) | T → C (Ile154Thr) | / | / | / | Ribonucleoside-diphosphate reductase 1 β subunit |
| 1677301 | *pntA* | / | / | / | / | / | / | T → G (Lys201Asn) | NAD(P) transhydrogenase α subunit |
| 2956405 | *ptrA* | / | / | / | / | / | / | A → C (Phe827Cys) | Protease 3 |
| 1158617 | *ptsG* | / | / | / | / | T → G (Leu250Arg) | Del of T (Ile25 FS) | Del of T (Ile25 FS) | PTS system glucose-specific EIICB component |
| 1756362 | *pykF* | / | / | / | A → C (Glu222Ala) | / | / | / | Pyruvate kinase I |
| 192080 | *pyrH* | G → A (Gly76Ser) | G → A (Gly76Ser) | G → A (Gly76Ser) | G → A (Gly76Ser) | / | / | / | Uridylate kinase |
| 3440239 | *rpoA* | / | A → G (Val264Ala) | T → C (Val264Ala) | A → G (Val264Ala) | / | / | / | DNA-directed RNA polymerase α subunit |
| 4188972 | *rpoC* | / | / | / | / | / | / | A → C (Asp1208Ala) | DNA-directed RNA polymerase β' subunit |
| 962466 | *rpsA* | / | / | A → C (Lys158Gln) | A → C (Lys158Gln) | / | A → C (Gln355Pro) | / | 30S ribosomal protein S1 |
| 3453189 | *rpsJ* | / | / | A → G (Thr28Ala) | T → C (Thr28Ala) | / | / | / | 30S ribosomal protein S10 |
| 4171475 | *rrlB* | / | A → G | / | / | / | / | / | 23S ribosomal RNA |
| 3942033 | *rrsC* | / | / | A → G | / | / | / | / | rrsC 16S ribosomal RNA |
| 3085637 | *speA* | / | / | / | / | / | / | A → C (Phe92Cys) | Biosynthetic Arg decarboxylase |
| 3082986 | *speB* | / | / | / | T → C (Asp271Gly) | / | / | / | Agmatinase |
| 3108699 | *speC* | / | / | / | T → C (Asn153Asp) | / | / | / | Constitutive ornithine decarboxylase |
| 3823565 | *spoT* | / | / | A → C (Tyr389Ser) | A → C (Tyr389Ser) | / | / | / | Bifunctional (p)ppGpp synthase/hydrolase SpoT |
| 2251 | *thrA* | / | / | / | / | / | T → G (Ser639Ala) | / | Bifunctional aspartokinase/homoserine dehydrogenase 1 |
| 1332759 | *topA* | / | / | / | / | / | / | T → G (Phe571Cys) | DNA topoisomerase 1 |
| 4111195 | *tpiA* | / | / | / | / | / | / | A → C (Ser105Ala) | Triosephosphate isomerase |
| 4633009 | *trpR* | / | / | / | / | C → T (Arg84Cys) | C → T (Arg84Cys) | C → T (Arg84Cys) | Trp operon repressor |
| 3513594 | *trpS* | T → G (Glu15Asp) | T → G (Glu15Asp) | A → C (Glu15Asp) | T → G (Glu15Asp) | / | / | T → G (Met187Leu) | Trp--tRNA ligase |
| 4253359 | *ubiA* | / | / | / | / | / | / | A → C (Asn115Thr) | 4-hydroxybenzoate octaprenyltransferase |
| 4418703 | *ulaG* | / | / | A → C (Tyr308Ser) | / | / | / | / | Probable L-ascorbate-6-phosphate lactonase UlaG |
| 1995086 | *uvrY* | / | T → C (Lys92Glu) | A → G (Lys92Glu) | T → C (Lys92Glu) | / | / | / | Response regulator UvrY |
| 3929293 | *viaA* | / | / | / | / | / | / | G → T (Ser104Tyr) | Protein ViaA |
| 1220485^1220486 | *ycgH_1* | / | / | / | / | In of T (Ser298 FS) | In of T (Ser298 FS) | In of T (Ser298 FS) | Putative transporter component, N-terminal fragment |
| 1812754 | *ydjO* | / | / | A → G (Ile127Val) | / | / | / | / | Uncharacterized protein YdjO |
| 1868406 | *yeaG* | / | A → G (Gln500Arg) | / | / | / | / | / | Uncharacterized protein YeaG |
| 2209715 | *yehQ* | / | / | / | / | / | G → T (Val206Phe) | G → T (Val206Phe) | Protein YehQ |
| 2412747 | *yfbV* | / | / | / | / | / | / | T → G (Tyr129Ser) | UPF0208 membrane protein YfbV |
| 2526188 | *yfeR* | A → C (Ile223Met) | A → C (Ile223Met) | T → G (Ile223Met) | A → C (Ile223Met) | / | / | / | Transcriptional regulator of yefH |
| 2762367 | *yfjK* | A → G (Ser392Pro) | / | / | / | / | / | / | Radiation resistance protein DEAD/H helicase-like protein CP4-57 |
| 2861088 | *ygbI* | / | / | / | / | / | / | A → C (Syn) | putative DeoR-type DNA-binding transcriptional regulator YgbI |
| 3096108 | *yggT* | / | / | / | / | / | / | A → C (Ile97Leu) | Uncharacterized protein YggT |
| 3385746 | *yhcN* | / | / | / | / | / | / | T → G (Ile70Ser) | Uncharacterized protein YhcN |

| Genomic position (bp) | Gene | 4TUB34 | 4TUB49 | 4TUB81 | 4TUB93 | 5TUB23 | 5TUB48 | 5TUB83 | Protein name |
|---|---|---|---|---|---|---|---|---|---|
| 3711747 | *yhjX* | / | / | A → G (Syn) | T → C (Syn) | / | / | / | Uncharacterized MFS-type transporter YhjX |
| 3771196 | *yibH* | / | / | / | / | / | / | T → G (Ile62Leu) | Inner membrane protein YibH |
| 4506186 | *yjhC* | / | A → G (Syn) | A → G (Syn) | A → G (Syn) | / | / | / | KpLE2 phage-like element; putative oxidoreductase YjhC |
| 1755461 | *IGR* | / | / | T → G (IGR) | / | / | A → C | A → C | Intergenic sequence 6 bp upstream of pykF |
| 1527823 | *IGR* | / | / | T → G (IGR) | / | / | A → C | A → C | Intergenic sequence, 67 bp upstream of rhsE |
| 3258210 | *IGR* | / | / | T → G (IGR) | / | / | A → C | T → G | Intergenic sequence, 75 bp upstream of tdcG |
| 3473535 | *IGR* | / | / | T → G (IGR) | / | / | A → C | T → G | Intergenic sequence, 21 bp upstream of fusA |
| 257908 | *IS1 in crl_2* | G → A (IGR) | G → A (IGR) | G → A (IGR) | G → A (IGR) | G → A (IGR) | G → A (IGR) | G → A (IGR) | Insertion element |
| 258675^258676 | *IS1 in crl_2* | In of G (IGR) | G → A (IGR) | G → A (IGR) | G → A (IGR) | G → A (IGR) | G → A (IGR) | G → A (IGR) | Insertion element |
| 4296380^4296381 | *REP321j* | In of CG | In of CG | In of CG | In of CG | In of CG | In of CG | In of CG | In REP321j 56 bp downstream of yjcO |
| 3883886 | *σ70 promoter of dnaA* | / | / | A → G | / | / | / | / | σ70 promoter, 157 bp upstream of dnaA |
| 3073817 | *σ70 promoter of epd* | / | / | / | / | / | A → C | / | σ70 promoter, Cra bindind site 121 bp downstream of epd |
| 2071319 | *σ70 promoter of isrC* | / | T → C | / | / | / | / | / | OxyR binding site in σ70 promoter of small RNA isrC |
| 1157621 | *σ70 promoter of ycfH* | / | / | / | / | / | / | T → G | σ70 promoter, bindind site 47 bp downstream of ycfH |
| Syn = synonymous | | | | | | | | | |
| In = insertion | | | | | | | | | |
| Del = deletion | | | | | | | | | |
| FS = frameshift | | | | | | | | | |
| IGR = intergenic region | | | | | | | | | |
