## Supplementary material for "Laboratory evolution of *Escherichia coli* enables life based on fluorinated amino acids": Auxiliary Table 2_Proteomics_5TUB83

| Gene names | og2(fold-Change | -Log(p-value) | Significant | Protein IDs | Majority protein IDs | Protein names |
|---|---|---|---|---|---|---|
| *dps* | -6.084 | 5.165 | + | P0ABT2 | P0ABT2 | DNA protection during starvation protein |
| *prlC* | -5.470 | 3.838 | + | P27298 | P27298 | Oligopeptidase A |
| *ptsG* | -5.318 | 5.648 | + | P69786 | P69786 | PTS system glucose-specific EIICB component |
| *osmY* | -5.163 | 5.519 | + | P0AFH8 | P0AFH8 | Osmotically-inducible protein Y |
| *yciG* | -4.854 | 1.892 | + | P21361 | P21361 | Uncharacterized protein YciG |
| *ybaY* | -4.752 | 4.921 | + | P77717 | P77717 | Uncharacterized lipoprotein YbaY |
| *yghA* | -4.703 | 2.674 | + | P0AG84 | P0AG84 | Uncharacterized oxidoreductase YghA |
| *psiF* | -4.366 | 4.986 | + | P0AFM4 | P0AFM4 | Phosphate starvation-inducible protein PsiF |
| *ynhG* | -4.282 | 5.187 | + | P76193 | P76193 | Probable L,D-transpeptidase YnhG |
| *yeaG* | -4.129 | 4.847 | + | P0ACY3 | P0ACY3 | Uncharacterized protein YeaG |
| *ygdI* | -4.066 | 5.078 | + | P65292 | P65292 | Uncharacterized lipoprotein YgdI |
| *yhbO* | -4.012 | 3.433 | + | P45470 | P45470 | Protein YhbO |
| *ecnB* | -4.011 | 4.699 | + | P0ADB7 | P0ADB7 | Entericidin B |
| *ahr* | -3.964 | 4.607 | + | P27250 | P27250 | Aldehyde reductase Ahr |
| *osmE* | -3.951 | 5.297 | + | P0ADB1 | P0ADB1 | Osmotically-inducible lipoprotein E |
| *wrbA* | -3.908 | 4.569 | + | P0A8G6 | P0A8G6 | NAD(P)H dehydrogenase (quinone) |
| *ilvC* | -3.870 | 5.488 | + | P05793 | P05793 | Ketol-acid reductoisomerase |
| *elaB* | -3.730 | 4.702 | + | P0AEH5 | P0AEH5 | Protein ElaB |
| *fbaB* | -3.694 | 4.730 | + | P0A991 | P0A991 | Fructose-bisphosphate aldolase class 1 |
| *ygaM* | -3.558 | 4.473 | + | P0ADQ7 | P0ADQ7 | Uncharacterized protein YgaM |
| *poxB* | -3.524 | 4.488 | + | P07003 | P07003 | Pyruvate dehydrogenase [ubiquinone] |
| *yncE* | -3.523 | 2.602 | + | P76116 | P76116 | Uncharacterized protein YncE |
| *stpA* | -3.452 | 4.881 | + | P0ACG1 | P0ACG1 | DNA-binding protein StpA |
| *talA* | -3.269 | 3.709 | + | P0A867 | P0A867 | Transaldolase A |
| *katE* | -3.265 | 2.507 | + | P21179 | P21179 | Catalase HPII |
| *ysgA* | -2.974 | 6.317 | + | P56262 | P56262 | Putative carboxymethylenebutenolidase |
| *yqjD* | -2.942 | 5.174 | + | P64581 | P64581 | Uncharacterized protein YqjD |
| *amyA* | -2.846 | 4.042 | + | P26612 | P26612 | Cytoplasmic alpha-amylase |
| *glnH* | -2.706 | 3.873 | + | P0AEQ3 | P0AEQ3 | Glutamine-binding periplasmic protein |
| *ygaU* | -2.703 | 4.043 | + | P0ADE6 | P0ADE6 | Uncharacterized protein YgaU |
| *osmC* | -2.667 | 4.598 | + | P0C0L2 | P0C0L2 | Peroxiredoxin OsmC |
| *gor* | -2.652 | 1.636 | + | P06715 | P06715 | Glutathione reductase |
| *dkgA* | -2.578 | 4.692 | + | Q46857 | Q46857 | 2,5-diketo-D-gluconic acid reductase A |
| *spy* | -2.530 | 3.235 | + | P77754 | P77754 | Spheroplast protein Y |
| *msyB* | -2.524 | 3.717 | + | P25738 | P25738 | Acidic protein MsyB |
| *glnQ* | -2.513 | 5.187 | + | P10346 | P10346 | Glutamine transport ATP-binding protein GlnQ |
| *ycaC* | -2.511 | 3.707 | + | P21367 | P21367 | Uncharacterized protein YcaC |
| *bfr* | -2.481 | 3.713 | + | P0ABD3 | P0ABD3 | Bacterioferritin |
| *aceA* | -2.445 | 5.690 | + | P0A9G6 | P0A9G6 | Isocitrate lyase |
| *tktB* | -2.409 | 3.841 | + | P33570 | P33570 | Transketolase 2 |
| *hchA* | -2.330 | 3.060 | + | P31658 | P31658 | Molecular chaperone Hsp31 and glyoxalase 3 |
| *ldhA* | -2.327 | 5.296 | + | P52643 | P52643 | D-lactate dehydrogenase |
| *erfK* | -2.290 | 3.233 | + | P39176 | P39176 | Probable L,D-transpeptidase ErfK/SrfK |
| *uspD* | -2.265 | 4.568 | + | P0AAB8 | P0AAB8 | Universal stress protein D |
| *acnA* | -2.259 | 3.386 | + | P25516 | P25516 | Aconitate hydratase A |
| *yegS* | -2.233 | 4.857 | + | P76407 | P76407 | Lipid kinase YegS |
| *bolA* | -2.215 | 4.222 | + | P0ABE2 | P0ABE2 | Protein BolA |
| *deoA* | -2.212 | 4.461 | + | P07650 | P07650 | Thymidine phosphorylase |
| *sra* | -2.155 | 2.059 | + | P68191 | P68191 | Stationary-phase-induced ribosome-associated protein |
| *ybeL* | -2.115 | 3.466 | + | P0AAT9 | P0AAT9 | Uncharacterized protein YbeL |
| *grxB* | -2.090 | 5.304 | + | P0AC59 | P0AC59 | Glutaredoxin-2 |
| *proP* | -2.087 | 3.296 | + | P0C0L7 | P0C0L7 | Proline/betaine transporter |
| *luxS* | -2.078 | 3.583 | + | P45578 | P45578 | S-ribosylhomocysteine lyase |

| Gene names | og2(fold-Change | -Log(p-value) | Significant | Protein IDs | Majority protein IDs | Protein names |
|---|---|---|---|---|---|---|
| *rho* | -2.070 | 4.638 | + | P0AG30 | P0AG30 | Transcription termination factor Rho |
| *curA* | -2.062 | 3.457 | + | P76113 | P76113 | NADPH-dependent curcumin reductase |
| *sstT* | -2.043 | 2.857 | + | P0AGE4 | P0AGE4 | Serine/threonine transporter SstT |
| *yciN* | -2.041 | 3.474 | + | P0AB61 | P0AB61 | Protein YciN |
| *ndh* | -2.018 | 3.885 | + | P00393 | P00393 | NADH dehydrogenase |
| *ybaL* | -2.015 | 3.044 | + | P39830 | P39830 | Inner membrane protein YbaL |
| *cdd* | -2.006 | 2.655 | + | P0ABF6 | P0ABF6 | Cytidine deaminase |
| *mscS* | -2.001 | 4.356 | + | P0C0S1 | P0C0S1 | Small-conductance mechanosensitive channel |
| *slp* | -1.996 | 2.209 | + | P37194 | P37194 | Outer membrane protein slp |
| *yecC* | -1.981 | 3.763 | + | P37774 | P37774 | Uncharacterized amino-acid ABC transporter ATP-binding protein YecC |
| *modF* | -1.970 | 4.270 | + | P31060 | P31060 | Putative molybdenum transport ATP-binding protein ModF |
| *mscL* | -1.964 | 3.919 | + | P0A742 | P0A742 | Large-conductance mechanosensitive channel |
| *ybjP* | -1.946 | 3.951 | + | P75818 | P75818 | Uncharacterized lipoprotein YbjP |
| *deoC* | -1.942 | 4.860 | + | P0A6L0 | P0A6L0 | Deoxyribose-phosphate aldolase |
| *dcyD* | -1.933 | 4.040 | + | P76316 | P76316 | D-cysteine desulfhydrase |
| *yjiM* | -1.931 | 3.684 | + | P39384 | P39384 | Uncharacterized protein YjiM |
| *ygdR* | -1.930 | 3.462 | + | P65294 | P65294 | Uncharacterized lipoprotein YgdR |
| *btuE* | -1.929 | 4.125 | + | P06610 | P06610 | Vitamin B12 transport periplasmic protein BtuE |
| *yibF* | -1.913 | 4.366 | + | P0ACA1 | P0ACA1 | Uncharacterized GST-like protein YibF |
| *ydjN* | -1.908 | 2.629 | + | P77529 | P77529 | Uncharacterized symporter YdjN |
| *uvrA* | -1.900 | 4.883 | + | P0A698 | P0A698 | UvrABC system protein A |
| *yaeP* | -1.892 | 3.571 | + | P0A8K5 | P0A8K5 | UPF0253 protein YaeP |
| *yibT* | -1.883 | 3.355 | + | Q2M7R5 | Q2M7R5 | Uncharacterized protein YibT |
| *fumC* | -1.855 | 3.300 | + | P05042 | P05042 | Fumarate hydratase class II |
| *dhaM* | -1.848 | 5.763 | + | P37349 | P37349 | PTS-dependent dihydroxyacetone kinase, phosphotransferase subunit DhaM |
| *ppsA* | -1.838 | 4.143 | + | P23538 | P23538 | Phosphoenolpyruvate synthase |
| *gabD* | -1.828 | 3.361 | + | P25526 | P25526 | Succinate-semialdehyde dehydrogenase [NADP(+)] GabD |
| *arcA* | -1.813 | 6.233 | + | P0A9Q1 | P0A9Q1 | Aerobic respiration control protein ArcA |
| *yahK* | -1.800 | 4.306 | + | P75691 | P75691 | Aldehyde reductase YahK |
| *ampD* | -1.799 | 1.689 | + | P13016 | P13016 | 1,6-anhydro-N-acetylmuramyl-L-alanine amidase AmpD |
| *ydbK* | -1.787 | 2.823 | + | P52647 | P52647 | Probable pyruvate-flavodoxin oxidoreductase |
| *livJ* | -1.785 | 3.503 | + | P0AD96 | P0AD96 | Leu/Ile/Val-binding protein |
| *gss* | -1.773 | 3.088 | + | P0AES0 | P0AES0 | Bifunctional glutathionylspermidine synthetase/amidase |
| *kgtP* | -1.769 | 3.050 | + | P0AEX3 | P0AEX3 | Alpha-ketoglutarate permease |
| *uspG* | -1.745 | 3.658 | + | P39177 | P39177 | Universal stress protein G |
| *gabT* | -1.728 | 3.313 | + | P22256 | P22256 | 4-aminobutyrate aminotransferase GabT |
| *gstB* | -1.706 | 4.007 | + | P0ACA7 | P0ACA7 | Glutathione S-transferase GstB |
| *nagA* | -1.701 | 3.651 | + | P0AF18 | P0AF18 | N-acetylglucosamine-6-phosphate deacetylase |
| *sufA* | -1.682 | 1.537 | + | P77667 | P77667 | Protein SufA |
| *yccX* | -1.667 | 3.553 | + | P0AB65 | P0AB65 | Acylphosphatase |
| *crl* | -1.664 | 3.398 | + | P24251 | P24251 | Sigma factor-binding protein Crl |
| *efeO* | -1.657 | 2.371 | + | P0AB24 | P0AB24 | Iron uptake system component EfeO |
| *icd* | -1.647 | 5.075 | + | P08200 | P08200 | Isocitrate dehydrogenase [NADP] |
| *yiiS* | -1.644 | 3.685 | + | P32162 | P32162 | UPF0381 protein YiiS |
| *hinT* | -1.634 | 2.623 | + | P0ACE7 | P0ACE7 | HIT-like protein HinT |
| *ydcY* | -1.626 | 2.710 | + | P64455 | P64455 | Uncharacterized protein YdcY |
| *fhuA* | -1.621 | 1.665 | + | P06971 | P06971 | Ferrichrome-iron receptor |
| *ppsR* | -1.620 | 3.404 | + | P0A8A4 | P0A8A4 | Phosphoenolpyruvate synthase regulatory protein |
| *lpoB* | -1.590 | 3.616 | + | P0AB38 | P0AB38 | Penicillin-binding protein activator LpoB |
| *osmF* | -1.574 | 1.637 | + | P33362 | P33362 | Putative osmoprotectant uptake system substrate-binding protein OsmF |
| *can* | -1.572 | 1.837 | + | P61517 | P61517 | Carbonic anhydrase 2 |
| *aroD* | -1.567 | 5.153 | + | P05194 | P05194 | 3-dehydroquinate dehydratase |
| *yccU* | -1.538 | 3.598 | + | P75874 | P75874 | Uncharacterized protein YccU |

| Gene names | og2(fold-Change | -Log(p-value) | Significant | Protein IDs | Majority protein IDs | Protein names |
|---|---|---|---|---|---|---|
| *ihfA* | -1.523 | 3.299 | + | P0A6X7 | P0A6X7 | Integration host factor subunit alpha |
| *yaeH* | -1.522 | 3.824 | + | P62768 | P62768 | UPF0325 protein YaeH |
| *yfaY* | -1.503 | 3.429 | + | P77808 | P77808 | NMN amidohydrolase-like protein YfaY |
| *glf* | -1.486 | 3.465 | + | P37747 | P37747 | UDP-galactopyranose mutase |
| *chbB* | -1.480 | 4.954 | + | P69795 | P69795 | N,N-diacetylchitobiose-specific phosphotransferase enzyme IIB component |
| *yciO* | -1.462 | 3.825 | + | P0AFR4 | P0AFR4 | Uncharacterized protein YciO |
| *yafJ* | -1.443 | 3.579 | + | Q47147 | Q47147 | Putative glutamine amidotransferase YafJ |
| *yceB* | -1.437 | 3.976 | + | P0AB26 | P0AB26 | Uncharacterized lipoprotein YceB |
| *ybhG* | -1.430 | 3.847 | + | P75777 | P75777 | UPF0194 membrane protein YbhG |
| *ihfB* | -1.425 | 3.398 | + | P0A6Y1 | P0A6Y1 | Integration host factor subunit beta |
| *pepT* | -1.423 | 4.477 | + | P29745 | P29745 | Peptidase T |
| *bioA* | -1.416 | 3.754 | + | P12995 | P12995 | Adenosylmethionine-8-amino-7-oxononanoate aminotransferase |
| *tesB* | -1.415 | 4.576 | + | P0AGG2 | P0AGG2 | Acyl-CoA thioesterase 2 |
| *yajO* | -1.413 | 4.584 | + | P77735 | P77735 | Uncharacterized oxidoreductase YajO |
| *yjfI* | -1.397 | 4.132 | + | P0AF76 | P0AF76 | Uncharacterized protein YjfI |
| *dacC* | -1.396 | 3.784 | + | P08506 | P08506 | D-alanyl-D-alanine carboxypeptidase DacC |
| *nadE* | -1.395 | 3.814 | + | P18843 | P18843 | NH(3)-dependent NAD(+) synthetase |
| *gstA* | -1.392 | 3.867 | + | P0A9D2 | P0A9D2 | Glutathione S-transferase GstA |
| *ycfP* | -1.372 | 4.358 | + | P0A8E1 | P0A8E1 | UPF0227 protein YcfP |
| *gcd* | -2.462 | 1.226 | + | P15877 | P15877 | Quinoprotein glucose dehydrogenase |
| *fklB* | 1.372 | 3.998 | + | P0A9L3 | P0A9L3 | FKBP-type 22 kDa peptidyl-prolyl cis-trans isomerase |
| *amiA* | 1.394 | 3.684 | + | P36548 | P36548 | N-acetylmuramoyl-L-alanine amidase AmiA |
| *glk* | 1.417 | 3.470 | + | P0A6V8 | P0A6V8 | Glucokinase |
| *miaB* | 1.424 | 4.098 | + | P0AEI1 | P0AEI1 | tRNA-2-methylthio-N(6)-dimethylallyladenosine synthase |
| *yggT* | 1.432 | 2.640 | + | P64564 | P64564 | Uncharacterized protein YggT |
| *gpt* | 1.439 | 3.528 | + | P0A9M5 | P0A9M5 | Xanthine phosphoribosyltransferase |
| *coaA* | 1.470 | 5.155 | + | P0A6I3 | P0A6I3 | Pantothenate kinase |
| *uvrD* | 1.472 | 2.917 | + | P03018 | P03018 | DNA helicase II |
| *eco* | 1.484 | 3.124 | + | P23827 | P23827 | Ecotin |
| *cspE* | 1.508 | 3.119 | + | P0A972 | P0A972 | Cold shock-like protein CspE |
| *pstS* | 1.533 | 3.368 | + | P0AG82 | P0AG82 | Phosphate-binding protein PstS |
| *leuD* | 1.538 | 2.856 | + | P30126 | P30126 | 3-isopropylmalate dehydratase small subunit |
| *ygiC* | 1.563 | 4.618 | + | P0ADT5 | P0ADT5 | Putative acid--amine ligase YgiC |
| *yhgF* | 1.569 | 3.518 | + | P46837 | P46837 | Protein YhgF |
| *ftnA* | 1.581 | 2.314 | + | P0A998 | P0A998 | Bacterial non-heme ferritin |
| *yjgM* | 1.591 | 3.360 | + | P39337 | P39337 | Uncharacterized N-acetyltransferase YjgM |
| *nuoE* | 1.597 | 3.199 | + | P0AFD1 | P0AFD1 | NADH-quinone oxidoreductase subunit E |
| *metC* | 1.609 | 4.034 | + | P06721 | P06721 | Cystathionine beta-lyase MetC |
| *rsuA* | 1.613 | 4.894 | + | P0AA43 | P0AA43 | Ribosomal small subunit pseudouridine synthase A |
| *yeiP* | 1.624 | 3.987 | + | P0A6N8 | P0A6N8 | Elongation factor P-like protein |
| *ampH* | 1.644 | 4.054 | + | P0AD70 | P0AD70 | D-alanyl-D-alanine-carboxypeptidase/endopeptidase AmpH |
| *upp* | 1.656 | 4.209 | + | P0A8F0 | P0A8F0 | Uracil phosphoribosyltransferase |
| *mdaB* | 1.714 | 3.843 | + | P0AEY5 | P0AEY5 | Modulator of drug activity B |
| *sseB* | 1.718 | 3.006 | + | P0AFZ1 | P0AFZ1 | Protein SseB |
| *rcnB* | 1.724 | 2.953 | + | P64534 | P64534 | Nickel/cobalt homeostasis protein RcnB |
| *sodB* | 1.725 | 3.086 | + | P0AGD3 | P0AGD3 | Superoxide dismutase [Fe] |
| *nrdB* | 1.728 | 3.759 | + | P69924 | P69924 | Ribonucleoside-diphosphate reductase 1 subunit beta |
| *metQ* | 1.728 | 4.284 | + | P28635 | P28635 | D-methionine-binding lipoprotein MetQ |
| *nrdA* | 1.740 | 3.447 | + | P00452 | P00452 | Ribonucleoside-diphosphate reductase 1 subunit alpha |
| *ybhA* | 1.755 | 4.756 | + | P21829 | P21829 | Pyridoxal phosphate phosphatase YbhA |
| *ybhC* | 1.770 | 3.320 | + | P46130 | P46130 | Putative acyl-CoA thioester hydrolase YbhC |
| *grxA* | 1.780 | 4.072 | + | P68688 | P68688 | Glutaredoxin-1 |
| *yhiI* | 1.819 | 3.587 | + | P37626 | P37626 | Uncharacterized protein YhiI |

| Gene names | og2(fold-Change | -Log(p-value) | Significant | Protein IDs | Majority protein IDs | Protein names |
|---|---|---|---|---|---|---|
| *thiG* | 1.827 | 4.123 | + | P30139 | P30139 | Thiazole synthase |
| *rsmD* | 1.834 | 5.102 | + | P0ADX9 | P0ADX9 | Ribosomal RNA small subunit methyltransferase D |
| *metL* | 1.835 | 3.772 | + | P00562 | P00562 | Bifunctional aspartokinase/homoserine dehydrogenase 2 |
| *recA* | 1.864 | 3.735 | + | P0A7G6 | P0A7G6 | Protein RecA |
| *tdcF* | 1.897 | 2.080 | + | P0AGL2 | P0AGL2 | Putative reactive intermediate deaminase TdcF |
| *nfsB* | 1.899 | 4.676 | + | P38489 | P38489 | Oxygen-insensitive NAD(P)H nitroreductase |
| *metN* | 1.925 | 4.886 | + | P30750 | P30750 | Methionine import ATP-binding protein MetN |
| *rsmG* | 1.950 | 4.694 | + | P0A6U5 | P0A6U5 | Ribosomal RNA small subunit methyltransferase G |
| *dcuA* | 1.961 | 3.919 | + | P0ABN5 | P0ABN5 | Anaerobic C4-dicarboxylate transporter DcuA |
| *ridA* | 1.969 | 3.651 | + | P0AF93 | P0AF93 | 2-iminobutanoate/2-iminopropanoate deaminase |
| *qseB* | 2.004 | 3.362 | + | P52076 | P52076 | Transcriptional regulatory protein QseB |
| *rihA* | 2.032 | 2.771 | + | P41409 | P41409 | Pyrimidine-specific ribonucleoside hydrolase RihA |
| *metJ* | 2.049 | 4.059 | + | P0A8U6 | P0A8U6 | Met repressor |
| *glpX* | 2.069 | 4.453 | + | P0A9C9 | P0A9C9 | Fructose-1,6-bisphosphatase 1 class 2 |
| *ribB* | 2.071 | 3.753 | + | P0A7J0 | P0A7J0 | 3,4-dihydroxy-2-butanone 4-phosphate synthase |
| *pckA* | 2.089 | 5.662 | + | P22259 | P22259 | Phosphoenolpyruvate carboxykinase [ATP] |
| *yigA* | 2.121 | 3.752 | + | P23305 | P23305 | Uncharacterized protein YigA |
| *deaD* | 2.148 | 3.571 | + | P0A9P6 | P0A9P6 | ATP-dependent RNA helicase DeaD |
| *yajD* | 2.153 | 2.714 | + | P0AAQ2 | P0AAQ2 | Uncharacterized protein YajD |
| *cydA* | 2.243 | 3.755 | + | P0ABJ9 | P0ABJ9 | Cytochrome bd-I ubiquinol oxidase subunit 1 |
| *rnk* | 2.264 | 4.441 | + | P0AFW4 | P0AFW4 | Regulator of nucleoside diphosphate kinase |
| *metK* | 2.303 | 5.104 | + | P0A817 | P0A817 | S-adenosylmethionine synthase |
| *metB* | 2.312 | 4.036 | + | P00935 | P00935 | Cystathionine gamma-synthase |
| *iscA* | 2.315 | 3.964 | + | P0AAC8 | P0AAC8 | Iron-binding protein IscA |
| *yecD* | 2.345 | 3.152 | + | P0ADI7 | P0ADI7 | Isochorismatase family protein YecD |
| *manX* | 2.354 | 4.528 | + | P69797 | P69797 | PTS system mannose-specific EIIAB component |
| *glpT* | 2.367 | 1.597 | + | P08194 | P08194 | Glycerol-3-phosphate transporter |
| *ompC* | 2.484 | 4.557 | + | P06996;P77747;P77519 | P06996 | Outer membrane protein C |
| *dtpA* | 2.681 | 3.353 | + | P77304 | P77304 | Dipeptide and tripeptide permease A |
| *hha* | 2.725 | 5.151 | + | P0ACE3 | P0ACE3 | Hemolysin expression-modulating protein Hha |
| *manZ* | 2.758 | 4.841 | + | P69805 | P69805 | Mannose permease IID component |
| *glpQ* | 2.869 | 2.361 | + | P09394 | P09394 | Glycerophosphoryl diester phosphodiesterase |
| *fruA* | 2.877 | 4.818 | + | P20966 | P20966 | PTS system fructose-specific EIIBC component |
| *tatE* | 3.003 | 3.283 | + | P0A843 | P0A843 | Sec-independent protein translocase protein TatE |
| *nadA* | 3.042 | 5.158 | + | P11458 | P11458 | Quinolinate synthase A |
| *pyrB* | 3.054 | 5.074 | + | P0A786 | P0A786 | Aspartate carbamoyltransferase catalytic chain |
| *thiC* | 3.152 | 3.991 | + | P30136 | P30136 | Phosphomethylpyrimidine synthase |
| *pyrI* | 3.170 | 5.296 | + | P0A7F3 | P0A7F3 | Aspartate carbamoyltransferase regulatory chain |
| *uxaC* | 3.188 | 5.067 | + | P0A8G3 | P0A8G3 | Uronate isomerase |
| *mnmG* | 3.224 | 3.688 | + | P0A6U3 | P0A6U3 | tRNA uridine 5-carboxymethylaminomethyl modification enzyme MnmG |
| *fruK* | 3.485 | 5.010 | + | P0AEW9 | P0AEW9 | 1-phosphofructokinase |
| *gatA* | 3.578 | 3.732 | + | P69828 | P69828 | Galactitol-specific phosphotransferase enzyme IIA component |
| *fruB* | 3.672 | 6.168 | + | P69811 | P69811 | Multiphosphoryl transfer protein |
| *cspA* | 3.781 | 4.352 | + | P0A9X9 | P0A9X9 | Cold shock protein CspA |
| *nadB* | 3.838 | 5.325 | + | P10902 | P10902 | L-aspartate oxidase |
| *mglB* | 4.407 | 3.887 | + | P0AEE5 | P0AEE5 | D-galactose-binding periplasmic protein |
| *cspD* | 4.725 | 4.827 | + | P0A968 | P0A968 | Cold shock-like protein CspD |
| *hsdR* | 5.055 | 2.292 | + | P08956 | P08956 | Type I restriction enzyme EcoKI R protein |
| *mtr* | 5.071 | 5.675 | + | P0AAD2 | P0AAD2 | Tryptophan-specific transport protein |
| *znuA* | 5.740 | 6.613 | + | P39172 | P39172 | High-affinity zinc uptake system protein ZnuA |
| *sodA* | -0.243 | 0.663 | | P00448 | P00448 | Superoxide dismutase [Mn] |
| *thrB* | 0.734 | 0.998 | | P00547 | P00547 | Homoserine kinase |
| *thrA* | 0.851 | 0.891 | | P00561 | P00561 | Bifunctional aspartokinase/homoserine dehydrogenase 1 |

| Gene names | og2(fold-Change | -Log(p-value) | Significant | Protein IDs | Majority protein IDs | Protein names |
|---|---|---|---|---|---|---|
| *lepB* | -0.168 | 0.887 | | P00803 | P00803 | Signal peptidase I |
| *ppc* | -0.244 | 1.268 | | P00864 | P00864 | Phosphoenolpyruvate carboxylase |
| *ilvI* | 0.460 | 0.519 | | P00893 | P00893 | Acetolactate synthase isozyme 3 large subunit |
| *metG* | -0.055 | 0.404 | | P00959 | P00959 | Methionine--tRNA ligase |
| *glyQ* | -0.109 | 0.941 | | P00960 | P00960 | Glycine--tRNA ligase alpha subunit |
| *glyS* | -0.045 | 0.168 | | P00961 | P00961 | Glycine--tRNA ligase beta subunit |
| *carB* | -0.299 | 0.598 | | P00968 | P00968 | Carbamoyl-phosphate synthase large chain |
| *rpsF* | 0.029 | 0.080 | | P02358 | P02358 | 30S ribosomal protein S6 |
| *rpsG* | -0.082 | 0.434 | | P02359 | P02359 | 30S ribosomal protein S7 |
| *rplO* | 0.039 | 0.084 | | P02413 | P02413 | 50S ribosomal protein L15 |
| *mrcA* | 0.126 | 0.372 | | P02918 | P02918 | Penicillin-binding protein 1A |
| *mrcB* | 0.368 | 1.053 | | P02919 | P02919 | Penicillin-binding protein 1B |
| *tolC* | 0.091 | 0.282 | | P02930 | P02930 | Outer membrane protein TolC |
| *ompF* | 0.217 | 0.254 | | P02931 | P02931 | Outer membrane protein F |
| *lysR* | 0.269 | 0.372 | | P03030 | P03030 | Transcriptional activator protein LysR |
| *ygeA* | 0.083 | 0.629 | | P03813 | P03813 | Uncharacterized protein YgeA |
| *mioC* | 0.436 | 0.741 | | P03817 | P03817 | Protein MioC |
| *frsA* | -0.192 | 0.516 | | P04335 | P04335 | Esterase FrsA |
| *argI* | 0.910 | 0.266 | | P04391 | P04391 | Ornithine carbamoyltransferase chain I |
| *sbcB* | -1.143 | 1.126 | | P04995 | P04995 | Exodeoxyribonuclease I |
| *pnp* | 0.050 | 0.147 | | P05055 | P05055 | Polyribonucleotide nucleotidyltransferase |
| *tsaD* | 0.014 | 0.050 | | P05852 | P05852 | tRNA N6-adenosine threonylcarbamoyltransferase |
| *btuB* | 0.098 | 0.289 | | P06129 | P06129 | Vitamin B12 transporter BtuB |
| *dnaX* | -0.219 | 0.814 | | P06710 | P06710 | DNA polymerase III subunit tau |
| *ebgR* | -0.598 | 1.194 | | P06846 | P06846 | HTH-type transcriptional regulator EbgR |
| *hisC* | -0.228 | 0.322 | | P06986 | P06986 | Histidinol-phosphate aminotransferase |
| *hisB* | -0.497 | 1.190 | | P06987 | P06987 | Histidine biosynthesis bifunctional protein HisB |
| *hisD* | -0.011 | 0.011 | | P06988 | P06988 | Histidinol dehydrogenase |
| *hisI* | -0.825 | 1.152 | | P06989 | P06989 | Histidine biosynthesis bifunctional protein HisIE |
| *pntA* | 0.331 | 1.062 | | P07001 | P07001 | NAD(P) transhydrogenase subunit alpha |
| *prfB* | -0.089 | 0.734 | | P07012 | P07012 | Peptide chain release factor 2 |
| *sdhB* | 0.140 | 1.032 | | P07014 | P07014 | Succinate dehydrogenase iron-sulfur subunit |
| *valS* | -0.172 | 0.586 | | P07118 | P07118 | Valine--tRNA ligase |
| *truA* | -0.384 | 1.086 | | P07649 | P07649 | tRNA pseudouridine synthase A |
| *glgB* | 0.546 | 1.271 | | P07762 | P07762 | 1,4-alpha-glucan branching enzyme GlgB |
| *leuS* | -0.220 | 1.057 | | P07813 | P07813 | Leucine--tRNA ligase |
| *tdh* | 0.369 | 1.264 | | P07913 | P07913 | L-threonine 3-dehydrogenase |
| *usg* | -0.204 | 1.218 | | P08390 | P08390 | USG-1 protein |
| *sppA* | 0.025 | 0.053 | | P08395 | P08395 | Protease 4 |
| *xthA* | -0.159 | 1.282 | | P09030 | P09030 | Exodeoxyribonuclease III |
| *xylB* | -0.108 | 1.233 | | P09099 | P09099 | Xylulose kinase |
| *hemX* | 0.013 | 0.042 | | P09127 | P09127 | Putative uroporphyrinogen-III C-methyltransferase |
| *galE* | -0.105 | 0.602 | | P09147 | P09147 | UDP-glucose 4-epimerase |
| *nagE* | -0.071 | 0.161 | | P09323 | P09323 | PTS system N-acetylglucosamine-specific EIICBA component |
| *dedD* | -0.249 | 0.885 | | P09549 | P09549 | Cell division protein DedD |
| *argT* | -0.732 | 1.161 | | P09551 | P09551 | Lysine/arginine/ornithine-binding periplasmic protein |
| *modC* | 0.011 | 0.039 | | P09833 | P09833 | Molybdenum import ATP-binding protein ModC |
| *rep* | -0.228 | 0.163 | | P09980 | P09980 | ATP-dependent DNA helicase Rep |
| *yidB* | 0.509 | 0.991 | | P09996 | P09996 | Uncharacterized protein YidB |
| *acpP* | 0.102 | 0.168 | | P0A6A8 | P0A6A8 | Acyl carrier protein |
| *argB* | 0.473 | 0.096 | | P0A6C8 | P0A6C8 | Acetylglutamate kinase |
| *argR* | -0.060 | 0.074 | | P0A6D0 | P0A6D0 | Arginine repressor |
| *aroL* | -0.485 | 1.095 | | P0A6E1 | P0A6E1 | Shikimate kinase 2 |

| Gene names | og2(fold-Change | -Log(p-value) | Significant | Protein IDs | Majority protein IDs | Protein names |
|---|---|---|---|---|---|---|
| *argG* | -0.750 | 0.282 | | P0A6E4 | P0A6E4 | Argininosuccinate synthase |
| *carA* | -0.275 | 0.422 | | P0A6F1 | P0A6F1 | Carbamoyl-phosphate synthase small chain |
| *clpX* | -0.028 | 0.163 | | P0A6H1 | P0A6H1 | ATP-dependent Clp protease ATP-binding subunit ClpX |
| *yegH* | -1.467 | 2.088 | | P76389 | P76389 | UPF0053 protein YegH |
| *clsA* | -0.527 | 0.927 | | P0A6H8 | P0A6H8 | Cardiolipin synthase A |
| *coaD* | 0.388 | 0.912 | | P0A6I6 | P0A6I6 | Phosphopantetheine adenylyltransferase |
| *cysZ* | -0.424 | 1.188 | | P0A6J3 | P0A6J3 | Sulfate transporter CysZ |
| *ddlA* | -0.057 | 0.298 | | P0A6J8 | P0A6J8 | D-alanine--D-alanine ligase A |
| *dapF* | -0.117 | 0.312 | | P0A6K1 | P0A6K1 | Diaminopimelate epimerase |
| *dapA* | -0.103 | 0.808 | | P0A6L2 | P0A6L2 | 4-hydroxy-tetrahydrodipicolinate synthase |
| *prmB* | -1.384 | 3.426 | | P39199 | P39199 | 50S ribosomal protein L3 glutamine methyltransferase |
| *iadA* | -1.374 | 2.891 | | P39377 | P39377 | Isoaspartyl dipeptidase |
| *yecS* | -1.368 | 2.442 | | P0AFT2 | P0AFT2 | Inner membrane amino-acid ABC transporter permease protein YecS |
| *qorA* | -1.360 | 3.410 | | P28304 | P28304 | Quinone oxidoreductase 1 |
| *fabZ* | -0.016 | 0.053 | | P0A6Q6 | P0A6Q6 | 3-hydroxyacyl-[acyl-carrier-protein] dehydratase FabZ |
| *tamB* | -1.348 | 3.432 | | P39321 | P39321 | Translocation and assembly module TamB |
| *gpsA* | -0.192 | 0.652 | | P0A6S7 | P0A6S7 | Glycerol-3-phosphate dehydrogenase [NAD(P)+] |
| *sufS* | -1.347 | 1.635 | | P77444 | P77444 | Cysteine desulfurase |
| *ldcA* | -1.345 | 2.672 | | P76008 | P76008 | Murein tetrapeptide carboxypeptidase |
| *sohB* | -1.335 | 3.560 | | P0AG14 | P0AG14 | Probable protease SohB |
| *mug* | -1.320 | 3.024 | | P0A9H1 | P0A9H1 | G/U mismatch-specific DNA glycosylase |
| *lpp* | -1.314 | 3.734 | | P69776 | P69776 | Major outer membrane lipoprotein Lpp |
| *glgA* | 0.403 | 1.026 | | P0A6U8 | P0A6U8 | Glycogen synthase |
| *glgC* | 0.272 | 0.653 | | P0A6V1 | P0A6V1 | Glucose-1-phosphate adenylyltransferase |
| *gnd* | -1.306 | 4.453 | | P00350 | P00350 | 6-phosphogluconate dehydrogenase, decarboxylating |
| *glsA2* | -0.350 | 0.926 | | P0A6W0 | P0A6W0 | Glutaminase 2 |
| *mraY* | -0.449 | 1.298 | | P0A6W3 | P0A6W3 | Phospho-N-acetylmuramoyl-pentapeptide-transferase |
| *greA* | 0.131 | 0.447 | | P0A6W5 | P0A6W5 | Transcription elongation factor GreA |
| *yjgR* | -1.302 | 2.045 | | P39342 | P39342 | Uncharacterized protein YjgR |
| *yggN* | -1.287 | 4.537 | | P0ADS9 | P0ADS9 | Uncharacterized protein YggN |
| *purB* | -1.283 | 3.170 | | P0AB89 | P0AB89 | Adenylosuccinate lyase |
| *rsmF* | -1.279 | 2.217 | | P76273 | P76273 | Ribosomal RNA small subunit methyltransferase F |
| *cyoB* | -1.274 | 2.886 | | P0ABI8 | P0ABI8 | Cytochrome bo(3) ubiquinol oxidase subunit 1 |
| *potA* | -1.242 | 4.370 | | P69874 | P69874 | Spermidine/putrescine import ATP-binding protein PotA |
| *dnaK* | 0.267 | 1.180 | | P0A6Y8 | P0A6Y8 | Chaperone protein DnaK |
| *clpB* | -1.237 | 3.662 | | P63284 | P63284 | Chaperone protein ClpB |
| *yeaJ* | -1.231 | 1.363 | | P76237 | P76237 | Putative diguanylate cyclase YeaJ |
| *ivy* | -1.224 | 2.401 | | P0AD59 | P0AD59 | Inhibitor of vertebrate lysozyme |
| *infB* | 0.011 | 0.035 | | P0A705 | P0A705 | Translation initiation factor IF-2 |
| *infC* | -0.149 | 0.952 | | P0A707 | P0A707 | Translation initiation factor IF-3 |
| *qmcA* | -1.213 | 3.177 | | P0AA53 | P0AA53 | Protein QmcA |
| *yggE* | -1.204 | 3.781 | | P0ADS6 | P0ADS6 | Uncharacterized protein YggE |
| *tmk* | 0.166 | 1.131 | | P0A720 | P0A720 | Thymidylate kinase |
| *lpxA* | 0.016 | 0.091 | | P0A722 | P0A722 | Acyl-[acyl-carrier-protein]--UDP-N-acetylglucosamine O-acyltransferase |
| *cdh* | -1.195 | 2.531 | | P06282 | P06282 | CDP-diacylglycerol pyrophosphatase |
| *mak* | -1.193 | 4.043 | | P23917 | P23917 | Fructokinase |
| *hslJ* | -1.190 | 2.652 | | P52644 | P52644 | Heat shock protein HslJ |
| *minE* | 0.147 | 0.760 | | P0A734 | P0A734 | Cell division topological specificity factor |
| *ompX* | -1.177 | 2.930 | | P0A917 | P0A917 | Outer membrane protein X |
| *tyrS* | -1.169 | 4.722 | | P0AGJ9 | P0AGJ9 | Tyrosine--tRNA ligase |
| *msrA* | -0.020 | 0.090 | | P0A744 | P0A744 | Peptide methionine sulfoxide reductase MsrA |
| *ydbC* | -1.164 | 2.689 | | P25906 | P25906 | Putative oxidoreductase YdbC |
| *yagU* | -1.158 | 3.012 | | P0AAA1 | P0AAA1 | Inner membrane protein YagU |

| Gene names | og2(fold-Change | -Log(p-value) | Significant | Protein IDs | Majority protein IDs | Protein names |
|---|---|---|---|---|---|---|
| *nadD* | -0.141 | 0.736 | | P0A752 | P0A752 | Nicotinate-nucleotide adenylyltransferase |
| *mrr* | -1.149 | 3.566 | | P24202 | P24202 | Mrr restriction system protein |
| *ndk* | -0.315 | 0.950 | | P0A763 | P0A763 | Nucleoside diphosphate kinase |
| *rppH* | -0.024 | 0.146 | | P0A776 | P0A776 | RNA pyrophosphohydrolase |
| *nusB* | 0.112 | 0.420 | | P0A780 | P0A780 | N utilization substance protein B |
| *orn* | 0.151 | 0.755 | | P0A784 | P0A784 | Oligoribonuclease |
| *gdhA* | -1.149 | 2.944 | | P00370 | P00370 | NADP-specific glutamate dehydrogenase |
| *panD* | 0.255 | 0.952 | | P0A790 | P0A790 | Aspartate 1-decarboxylase |
| *msrB* | -1.145 | 2.672 | | P0A746 | P0A746 | Peptide methionine sulfoxide reductase MsrB |
| *lpxC* | -1.144 | 4.460 | | P0A725 | P0A725 | UDP-3-O-[3-hydroxymyristoyl] N-acetylglucosamine deacetylase |
| *uspE* | -1.125 | 4.287 | | P0AAC0 | P0AAC0 | Universal stress protein E |
| *gpmB* | -0.212 | 0.909 | | P0A7A2 | P0A7A2 | Probable phosphoglycerate mutase GpmB |
| *plsB* | 0.249 | 0.665 | | P0A7A7 | P0A7A7 | Glycerol-3-phosphate acyltransferase |
| *fliY* | -1.124 | 2.084 | | P0AEM9 | P0AEM9 | Cystine-binding periplasmic protein |
| *ppk* | 0.414 | 0.837 | | P0A7B1 | P0A7B1 | Polyphosphate kinase |
| *nadK* | 0.161 | 0.513 | | P0A7B3 | P0A7B3 | NAD kinase |
| *gmk* | -1.122 | 3.874 | | P60546 | P60546 | Guanylate kinase |
| *hslV* | -0.117 | 0.547 | | P0A7B8 | P0A7B8 | ATP-dependent protease subunit HslV |
| *lexA* | -0.107 | 0.871 | | P0A7C2 | P0A7C2 | LexA repressor |
| *pth* | -0.114 | 0.643 | | P0A7D1 | P0A7D1 | Peptidyl-tRNA hydrolase |
| *purA* | -0.164 | 1.192 | | P0A7D4 | P0A7D4 | Adenylosuccinate synthetase |
| *tpx* | -1.108 | 2.175 | | P0A862 | P0A862 | Thiol peroxidase |
| *uspF* | -1.108 | 3.006 | | P37903 | P37903 | Universal stress protein F |
| *pyrG* | 0.075 | 0.240 | | P0A7E5 | P0A7E5 | CTP synthase |
| *lysC* | -1.100 | 1.672 | | P08660 | P08660 | Lysine-sensitive aspartokinase 3 |
| *pfkB* | -1.095 | 3.951 | | P06999 | P06999 | ATP-dependent 6-phosphofructokinase isozyme 2 |
| *topA* | -1.092 | 4.722 | | P06612 | P06612 | DNA topoisomerase 1 |
| *queA* | 0.115 | 0.558 | | P0A7F9 | P0A7F9 | S-adenosylmethionine:tRNA ribosyltransferase-isomerase |
| *hdhA* | -1.090 | 6.147 | | P0AET8 | P0AET8 | 7-alpha-hydroxysteroid dehydrogenase |
| *speD* | -1.082 | 3.245 | | P0A7F6 | P0A7F6 | S-adenosylmethionine decarboxylase proenzyme |
| *recR* | -0.061 | 0.700 | | P0A7H6 | P0A7H6 | Recombination protein RecR |
| *yggL* | -1.079 | 2.631 | | P38521 | P38521 | Uncharacterized protein YggL |
| *glnA* | -1.077 | 2.254 | | P0A9C5 | P0A9C5 | Glutamine synthetase |
| *ribA* | -0.117 | 0.646 | | P0A7I7 | P0A7I7 | GTP cyclohydrolase-2 |
| *rpoD* | -1.076 | 2.920 | | P00579 | P00579 | RNA polymerase sigma factor RpoD |
| *rplJ* | -0.153 | 0.654 | | P0A7J3 | P0A7J3 | 50S ribosomal protein L10 |
| *rplK* | 0.123 | 0.541 | | P0A7J7 | P0A7J7 | 50S ribosomal protein L11 |
| *malP* | -1.073 | 3.448 | | P00490 | P00490 | Maltodextrin phosphorylase |
| *rplS* | -0.046 | 0.176 | | P0A7K6 | P0A7K6 | 50S ribosomal protein L19 |
| *tsf* | -1.062 | 4.643 | | P0A6P1 | P0A6P1 | Elongation factor Ts |
| *rplT* | -0.006 | 0.017 | | P0A7L3 | P0A7L3 | 50S ribosomal protein L20 |
| *rpmA* | -0.065 | 0.224 | | P0A7L8 | P0A7L8 | 50S ribosomal protein L27 |
| *rpmB* | -0.131 | 0.440 | | P0A7M2 | P0A7M2 | 50S ribosomal protein L28 |
| *rpmC* | -0.069 | 0.240 | | P0A7M6 | P0A7M6 | 50S ribosomal protein L29 |
| *rpmE* | -0.549 | 1.189 | | P0A7M9 | P0A7M9 | 50S ribosomal protein L31 |
| *rpmF* | -0.059 | 0.194 | | P0A7N4 | P0A7N4 | 50S ribosomal protein L32 |
| *rpmG* | -0.065 | 0.196 | | P0A7N9 | P0A7N9 | 50S ribosomal protein L33 |
| *rpmH* | -0.157 | 0.652 | | P0A7P5 | P0A7P5 | 50S ribosomal protein L34 |
| *ompT* | -1.062 | 1.689 | | P09169 | P09169 | Protease 7 |
| *rplI* | -0.083 | 0.352 | | P0A7R1 | P0A7R1 | 50S ribosomal protein L9 |
| *rpsJ* | 0.238 | 0.910 | | P0A7R5 | P0A7R5 | 30S ribosomal protein S10 |
| *rpsK* | 0.019 | 0.065 | | P0A7R9 | P0A7R9 | 30S ribosomal protein S11 |
| *rpsL* | 0.153 | 0.702 | | P0A7S3 | P0A7S3 | 30S ribosomal protein S12 |

| Gene names | og2(fold-Change | -Log(p-value) | Significant | Protein IDs | Majority protein IDs | Protein names |
|---|---|---|---|---|---|---|
| *rpsM* | 0.061 | 0.160 | | P0A7S9 | P0A7S9 | 30S ribosomal protein S13 |
| *rpsP* | -0.374 | 0.955 | | P0A7T3 | P0A7T3 | 30S ribosomal protein S16 |
| *rpsR* | 0.047 | 0.193 | | P0A7T7 | P0A7T7 | 30S ribosomal protein S18 |
| *rpsS* | -0.144 | 0.540 | | P0A7U3 | P0A7U3 | 30S ribosomal protein S19 |
| *rpsT* | -0.165 | 0.702 | | P0A7U7 | P0A7U7 | 30S ribosomal protein S20 |
| *ybiB* | -1.058 | 4.103 | | P30177 | P30177 | Uncharacterized protein YbiB |
| *rpsC* | -0.326 | 0.853 | | P0A7V3 | P0A7V3 | 30S ribosomal protein S3 |
| *rpsD* | 0.120 | 0.759 | | P0A7V8 | P0A7V8 | 30S ribosomal protein S4 |
| *rpsE* | 0.051 | 0.200 | | P0A7W1 | P0A7W1 | 30S ribosomal protein S5 |
| *rpsH* | -0.076 | 0.330 | | P0A7W7 | P0A7W7 | 30S ribosomal protein S8 |
| *rpsI* | 0.078 | 0.250 | | P0A7X3 | P0A7X3 | 30S ribosomal protein S9 |
| *tamA* | -1.048 | 2.555 | | P0ADE4 | P0ADE4 | Translocation and assembly module TamA |
| *rnhA* | -1.041 | 3.987 | | P0A7Y4 | P0A7Y4 | Ribonuclease HI |
| *zur* | -1.021 | 2.984 | | P0AC51 | P0AC51 | Zinc uptake regulation protein |
| *ymdB* | -1.018 | 2.942 | | P0A8D6 | P0A8D6 | O-acetyl-ADP-ribose deacetylase |
| *rpiA* | 0.265 | 0.975 | | P0A7Z0 | P0A7Z0 | Ribose-5-phosphate isomerase A |
| *rpoA* | -0.028 | 0.197 | | P0A7Z4 | P0A7Z4 | DNA-directed RNA polymerase subunit alpha |
| *rpoZ* | -0.029 | 0.096 | | P0A800 | P0A800 | DNA-directed RNA polymerase subunit omega |
| *frr* | -0.042 | 0.117 | | P0A805 | P0A805 | Ribosome-recycling factor |
| *yeiE* | -1.018 | 2.532 | | P0ACR4 | P0ACR4 | Uncharacterized HTH-type transcriptional regulator YeiE |
| *ppiD* | -1.017 | 4.162 | | P0ADY1 | P0ADY1 | Peptidyl-prolyl cis-trans isomerase D |
| *trmB* | -1.014 | 4.518 | | P0A8I5 | P0A8I5 | tRNA (guanine-N(7)-)-methyltransferase |
| *selA* | 0.107 | 0.416 | | P0A821 | P0A821 | L-seryl-tRNA(Sec) selenium transferase |
| *sfsA* | 0.144 | 0.541 | | P0A823 | P0A823 | Sugar fermentation stimulation protein A |
| *rarA* | -1.010 | 3.235 | | P0AAZ4 | P0AAZ4 | Replication-associated recombination protein A |
| *smpB* | 0.100 | 0.256 | | P0A832 | P0A832 | SsrA-binding protein |
| *cysK* | -1.004 | 2.877 | | P0ABK5 | P0ABK5 | Cysteine synthase A |
| *ftsH* | -0.999 | 3.393 | | P0AAI3 | P0AAI3 | ATP-dependent zinc metalloprotease FtsH |
| *yajL* | -0.994 | 3.212 | | Q46948 | Q46948 | Probable protein deglycase |
| *ybiV* | -0.989 | 2.560 | | P75792 | P75792 | Sugar phosphatase YbiV |
| *argP* | -0.987 | 3.811 | | P0A8S1 | P0A8S1 | HTH-type transcriptional regulator ArgP |
| *csrA* | -0.969 | 3.196 | | P69913 | P69913 | Carbon storage regulator |
| *nagB* | -0.967 | 2.633 | | P0A759 | P0A759 | Glucosamine-6-phosphate deaminase |
| *yeaE* | -0.965 | 3.177 | | P76234 | P76234 | Uncharacterized protein YeaE |
| *cca* | -0.963 | 2.116 | | P06961 | P06961 | Multifunctional CCA protein |
| *queC* | -0.954 | 4.085 | | P77756 | P77756 | 7-cyano-7-deazaguanine synthase |
| *gltD* | -0.946 | 3.388 | | P09832 | P09832 | Glutamate synthase [NADPH] small chain |
| *fabR* | -0.946 | 3.400 | | P0ACU5 | P0ACU5 | HTH-type transcriptional repressor FabR |
| *gltB* | -0.945 | 3.331 | | P09831 | P09831 | Glutamate synthase [NADPH] large chain |
| *maeA* | -0.932 | 3.378 | | P26616 | P26616 | NAD-dependent malic enzyme |
| *ybeZ* | -0.929 | 4.244 | | P0A9K3 | P0A9K3 | PhoH-like protein |
| *rhlE* | -0.927 | 2.604 | | P25888 | P25888 | ATP-dependent RNA helicase RhlE |
| *ybeY* | 0.041 | 0.121 | | P0A898 | P0A898 | Endoribonuclease YbeY |
| *slyB* | -0.926 | 2.845 | | P0A905 | P0A905 | Outer membrane lipoprotein SlyB |
| *dsbB* | -0.924 | 2.065 | | P0A6M2 | P0A6M2 | Disulfide bond formation protein B |
| *aroA* | -0.921 | 3.383 | | P0A6D3 | P0A6D3 | 3-phosphoshikimate 1-carboxyvinyltransferase |
| *rimP* | 0.152 | 0.786 | | P0A8A8 | P0A8A8 | Ribosome maturation factor RimP |
| *dhaL* | -0.914 | 2.828 | | P76014 | P76014 | PTS-dependent dihydroxyacetone kinase, ADP-binding subunit DhaL |
| *ybaB* | -0.120 | 0.387 | | P0A8B5 | P0A8B5 | Nucleoid-associated protein YbaB |
| *ybjQ* | 0.710 | 1.139 | | P0A8C1 | P0A8C1 | UPF0145 protein YbjQ |
| *ygfB* | -0.057 | 0.156 | | P0A8C4 | P0A8C4 | UPF0149 protein YgfB |
| *rpoE* | -0.912 | 4.330 | | P0AGB6 | P0AGB6 | ECF RNA polymerase sigma-E factor |
| *tdk* | -0.911 | 2.207 | | P23331 | P23331 | Thymidine kinase |

| Gene names | og2(fold-Change | -Log(p-value) | Significant | Protein IDs | Majority protein IDs | Protein names |
|---|---|---|---|---|---|---|
| *rpe* | -0.905 | 2.984 | | P0AG07 | P0AG07 | Ribulose-phosphate 3-epimerase |
| *iaaA* | -0.902 | 2.724 | | P37595 | P37595 | Isoaspartyl peptidase |
| *relB* | -0.899 | 2.884 | | P0C079 | P0C079 | Antitoxin RelB |
| *yceH* | -0.896 | 3.625 | | P29217 | P29217 | UPF0502 protein YceH |
| *pdxB* | -0.887 | 4.378 | | P05459 | P05459 | Erythronate-4-phosphate dehydrogenase |
| *slt* | -0.885 | 3.516 | | P0AGC3 | P0AGC3 | Soluble lytic murein transglycosylase |
| *uvrC* | -0.356 | 0.969 | | P0A8G0 | P0A8G0 | UvrABC system protein C |
| *coaE* | -0.884 | 2.141 | | P0A6I9 | P0A6I9 | Dephospho-CoA kinase |
| *yhjJ* | -0.877 | 2.552 | | P37648 | P37648 | Protein YhjJ |
| *zwf* | -0.876 | 3.930 | | P0AC53 | P0AC53 | Glucose-6-phosphate 1-dehydrogenase |
| *ffh* | -0.876 | 2.809 | | P0AGD7 | P0AGD7 | Signal recognition particle protein |
| *yqgF* | 0.191 | 0.872 | | P0A8I1 | P0A8I1 | Putative Holliday junction resolvase |
| *mraZ* | -0.871 | 2.785 | | P22186 | P22186 | Transcriptional regulator MraZ |
| *ydgA* | -0.870 | 3.143 | | P77804 | P77804 | Protein YdgA |
| *ybeD* | -0.117 | 0.290 | | P0A8J4 | P0A8J4 | UPF0250 protein YbeD |
| *adk* | -0.869 | 3.098 | | P69441 | P69441 | Adenylate kinase |
| *gpmA* | -0.858 | 3.608 | | P62707 | P62707 | 2,3-bisphosphoglycerate-dependent phosphoglycerate mutase |
| *amiB* | -0.855 | 1.956 | | P26365 | P26365 | N-acetylmuramoyl-L-alanine amidase AmiB |
| *serS* | 0.084 | 0.857 | | P0A8L1 | P0A8L1 | Serine--tRNA ligase |
| *yeaY* | -0.854 | 2.610 | | P0AA91 | P0AA91 | Uncharacterized lipoprotein YeaY |
| *yfgM* | -0.838 | 2.631 | | P76576 | P76576 | UPF0070 protein YfgM |
| *yeeX* | -0.261 | 0.979 | | P0A8M6 | P0A8M6 | UPF0265 protein YeeX |
| *matP* | 0.194 | 0.637 | | P0A8N0 | P0A8N0 | Macrodomain Ter protein |
| *pgl* | -0.829 | 2.755 | | P52697 | P52697 | 6-phosphogluconolactonase |
| *lysU* | 0.307 | 0.844 | | P0A8N5 | P0A8N5 | Lysine--tRNA ligase, heat inducible |
| *yggX* | -0.323 | 1.184 | | P0A8P3 | P0A8P3 | Probable Fe(2+)-trafficking protein |
| *xerC* | 0.346 | 0.535 | | P0A8P6 | P0A8P6 | Tyrosine recombinase XerC |
| *rnt* | -0.826 | 3.752 | | P30014 | P30014 | Ribonuclease T |
| *ycjF* | -0.171 | 0.066 | | P0A8R7 | P0A8R7 | UPF0283 membrane protein YcjF |
| *hdfR* | -0.016 | 0.039 | | P0A8R9 | P0A8R9 | HTH-type transcriptional regulator HdfR |
| *yidA* | -0.823 | 4.120 | | P0A8Y5 | P0A8Y5 | Sugar phosphatase YidA |
| *prmA* | 0.067 | 0.518 | | P0A8T1 | P0A8T1 | Ribosomal protein L11 methyltransferase |
| *rpoC* | -0.013 | 0.030 | | P0A8T7 | P0A8T7 | DNA-directed RNA polymerase subunit beta |
| *syd* | 0.575 | 1.138 | | P0A8U0 | P0A8U0 | Protein Syd |
| *allR* | -0.820 | 2.515 | | P0ACN4 | P0ACN4 | HTH-type transcriptional repressor AllR |
| *rbn* | 0.189 | 1.192 | | P0A8V0 | P0A8V0 | Ribonuclease BN |
| *rpoB* | -0.193 | 0.707 | | P0A8V2 | P0A8V2 | DNA-directed RNA polymerase subunit beta |
| *fadR* | 0.057 | 0.267 | | P0A8V6 | P0A8V6 | Fatty acid metabolism regulator protein |
| *iraP* | -0.819 | 1.631 | | P0AAN9 | P0AAN9 | Anti-adapter protein IraP |
| *slyA* | -0.585 | 1.275 | | P0A8W2 | P0A8W2 | Transcriptional regulator SlyA |
| *yqgE* | -0.077 | 0.209 | | P0A8W5 | P0A8W5 | UPF0301 protein YqgE |
| *livK* | -0.806 | 2.475 | | P04816 | P04816 | Leucine-specific-binding protein |
| *yjgA* | 0.288 | 1.234 | | P0A8X0 | P0A8X0 | UPF0307 protein YjgA |
| *mdoH* | -0.805 | 1.665 | | P62517 | P62517 | Glucans biosynthesis glucosyltransferase H |
| *yihX* | 0.133 | 0.182 | | P0A8Y3 | P0A8Y3 | Alpha-D-glucose-1-phosphate phosphatase YihX |
| *rstA* | -0.801 | 3.446 | | P52108 | P52108 | Transcriptional regulatory protein RstA |
| *yciA* | 0.372 | 0.884 | | P0A8Z0 | P0A8Z0 | Acyl-CoA thioester hydrolase YciA |
| *ybgC* | -0.131 | 0.330 | | P0A8Z3 | P0A8Z3 | Acyl-CoA thioester hydrolase YbgC |
| *blc* | -0.825 | 0.896 | | P0A901 | P0A901 | Outer membrane lipoprotein Blc |
| *bamC* | -0.234 | 0.662 | | P0A903 | P0A903 | Outer membrane protein assembly factor BamC |
| *glmU* | -0.801 | 2.161 | | P0ACC7 | P0ACC7 | Bifunctional protein GlmU |
| *mipA* | -0.404 | 0.923 | | P0A908 | P0A908 | MltA-interacting protein |
| *cmk* | -0.794 | 4.065 | | P0A6I0 | P0A6I0 | Cytidylate kinase |

| Gene names | og2(fold-Change | -Log(p-value) | Significant | Protein IDs | Majority protein IDs | Protein names |
|---|---|---|---|---|---|---|
| *pal* | 0.415 | 1.216 | | P0A912 | P0A912 | Peptidoglycan-associated lipoprotein |
| *amn* | -0.793 | 2.985 | | P0AE12 | P0AE12 | AMP nucleosidase |
| *sspA* | -0.793 | 3.554 | | P0ACA3 | P0ACA3 | Stringent starvation protein A |
| *bamE* | 0.355 | 0.990 | | P0A937 | P0A937 | Outer membrane protein assembly factor BamE |
| *moeA* | -0.788 | 3.891 | | P12281 | P12281 | Molybdopterin molybdenumtransferase |
| *rimJ* | -0.195 | 0.703 | | P0A948 | P0A948 | Ribosomal-protein-alanine acetyltransferase |
| *speG* | 0.140 | 0.287 | | P0A951 | P0A951 | Spermidine N(1)-acetyltransferase |
| *argS* | -0.787 | 4.186 | | P11875 | P11875 | Arginine--tRNA ligase |
| *ychF* | -0.775 | 3.357 | | P0ABU2 | P0ABU2 | Ribosome-binding ATPase YchF |
| *alaA* | -0.659 | 0.940 | | P0A959 | P0A959 | Glutamate-pyruvate aminotransferase AlaA |
| *ansA* | 0.137 | 0.696 | | P0A962 | P0A962 | L-asparaginase 1 |
| *cbpA* | -0.772 | 2.680 | | P36659 | P36659 | Curved DNA-binding protein |
| *yddE* | -0.769 | 2.283 | | P37757 | P37757 | Uncharacterized isomerase YddE |
| *cyoA* | -0.761 | 2.000 | | P0ABJ1 | P0ABJ1 | Cytochrome bo(3) ubiquinol oxidase subunit 2 |
| *potD* | -0.759 | 1.809 | | P0AFK9 | P0AFK9 | Spermidine/putrescine-binding periplasmic protein |
| *grxC* | -0.759 | 2.197 | | P0AC62 | P0AC62 | Glutaredoxin-3 |
| *gyrB* | -0.756 | 2.739 | | P0AES6 | P0AES6 | DNA gyrase subunit B |
| *speC* | -0.754 | 1.388 | | P21169 | P21169 | Ornithine decarboxylase, constitutive |
| *fur* | -0.152 | 0.394 | | P0A9A9 | P0A9A9 | Ferric uptake regulation protein |
| *gapA* | 0.017 | 0.052 | | P0A9B2;P33898 | P0A9B2 | Glyceraldehyde-3-phosphate dehydrogenase A |
| *ulaR* | -0.753 | 2.011 | | P0A9W0 | P0A9W0 | HTH-type transcriptional regulator UlaR |
| *ptrA* | -0.750 | 1.757 | | P05458 | P05458 | Protease 3 |
| *tyrR* | -0.749 | 2.867 | | P07604 | P07604 | Transcriptional regulatory protein TyrR |
| *trkA* | -0.747 | 2.403 | | P0AGI8 | P0AGI8 | Trk system potassium uptake protein TrkA |
| *mtlD* | -0.743 | 2.236 | | P09424 | P09424 | Mannitol-1-phosphate 5-dehydrogenase |
| *mqsA* | -0.738 | 1.986 | | Q46864 | Q46864 | Antitoxin MqsA |
| *dapD* | -0.067 | 0.275 | | P0A9D8 | P0A9D8 | 2,3,4,5-tetrahydropyridine-2,6-dicarboxylate N-succinyltransferase |
| *yeeZ* | -0.737 | 3.271 | | P0AD12 | P0AD12 | Protein YeeZ |
| *mntR* | -0.147 | 0.377 | | P0A9F1 | P0A9F1 | Transcriptional regulator MntR |
| *rseA* | -0.735 | 2.778 | | P0AFX7 | P0AFX7 | Anti-sigma-E factor RseA |
| *gyrA* | -0.735 | 2.012 | | P0AES4 | P0AES4 | DNA gyrase subunit A |
| *rpsA* | -0.734 | 2.541 | | P0AG67 | P0AG67 | 30S ribosomal protein S1 |
| *rnd* | -0.732 | 4.375 | | P09155 | P09155 | Ribonuclease D |
| *sthA* | -0.729 | 1.809 | | P27306 | P27306 | Soluble pyridine nucleotide transhydrogenase |
| *kdsA* | -0.728 | 4.830 | | P0A715 | P0A715 | 2-dehydro-3-deoxyphosphooctonate aldolase |
| *rbsK* | 0.167 | 1.225 | | P0A9J6 | P0A9J6 | Ribokinase |
| *def* | -0.724 | 2.613 | | P0A6K3 | P0A6K3 | Peptide deformylase |
| *yrdD* | -0.717 | 2.023 | | P45771 | P45771 | Uncharacterized protein YrdD |
| *suhB* | -0.716 | 2.583 | | P0ADG4 | P0ADG4 | Inositol-1-monophosphatase |
| *dapB* | -0.703 | 4.952 | | P04036 | P04036 | 4-hydroxy-tetrahydrodipicolinate reductase |
| *corA* | -0.695 | 2.003 | | P0ABI4 | P0ABI4 | Magnesium transport protein CorA |
| *nrdR* | -0.693 | 2.944 | | P0A8D0 | P0A8D0 | Transcriptional repressor NrdR |
| *pykA* | -0.691 | 2.557 | | P21599 | P21599 | Pyruvate kinase II |
| *yecA* | -0.690 | 1.471 | | P0AD05 | P0AD05 | Uncharacterized protein YecA |
| *dicA* | -0.690 | 2.516 | | P06966 | P06966 | HTH-type transcriptional regulator DicA |
| *der* | -0.687 | 2.499 | | P0A6P5 | P0A6P5 | GTPase Der |
| *moeB* | -0.679 | 1.628 | | P12282 | P12282 | Molybdopterin-synthase adenylyltransferase |
| *fbp* | -0.679 | 2.098 | | P0A993 | P0A993 | Fructose-1,6-bisphosphatase class 1 |
| *nagZ* | -0.675 | 2.244 | | P75949 | P75949 | Beta-hexosaminidase |
| *uvrB* | -0.672 | 2.200 | | P0A8F8 | P0A8F8 | UvrABC system protein B |
| *ycaO* | -0.667 | 4.567 | | P75838 | P75838 | Ribosomal protein S12 methylthiotransferase accessory factor YcaO |
| *yjdC* | -0.667 | 2.467 | | P0ACU7 | P0ACU7 | HTH-type transcriptional regulator YjdC |
| *aroC* | -0.666 | 2.712 | | P12008 | P12008 | Chorismate synthase |

| Gene names | og2(fold-Change | -Log(p-value) | Significant | Protein IDs | Majority protein IDs | Protein names |
|---|---|---|---|---|---|---|
| *wbbI* | -0.664 | 2.897 | | P37749 | P37749 | Beta-1,6-galactofuranosyltransferase WbbI |
| *dcp* | -0.664 | 3.451 | | P24171 | P24171 | Peptidyl-dipeptidase dcp |
| *modE* | -0.656 | 3.710 | | P0A9G8 | P0A9G8 | Transcriptional regulator ModE |
| *pykF* | -0.652 | 2.605 | | P0AD61 | P0AD61 | Pyruvate kinase I |
| *ftsE* | -0.129 | 1.116 | | P0A9R7 | P0A9R7 | Cell division ATP-binding protein FtsE |
| *gldA* | 0.606 | 1.021 | | P0A9S5 | P0A9S5 | Glycerol dehydrogenase |
| *livG* | -0.114 | 0.208 | | P0A9S7 | P0A9S7 | High-affinity branched-chain amino acid transport ATP-binding protein LivG |
| *serA* | 0.069 | 0.280 | | P0A9T0 | P0A9T0 | D-3-phosphoglycerate dehydrogenase |
| *eda* | -0.648 | 2.553 | | P0A955 | P0A955 | KHG/KDPG aldolase |
| *ybbA* | -0.134 | 1.115 | | P0A9T8 | P0A9T8 | Uncharacterized ABC transporter ATP-binding protein YbbA |
| *yeaK* | -0.646 | 2.267 | | P64483 | P64483 | Uncharacterized protein YeaK |
| *grpE* | -0.644 | 2.729 | | P09372 | P09372 | Protein GrpE |
| *ushA* | -0.640 | 3.334 | | P07024 | P07024 | UDP-sugar hydrolase |
| *ettA* | 0.176 | 0.644 | | P0A9W3 | P0A9W3 | Energy-dependent translational throttle protein EttA |
| *pheT* | -0.635 | 2.328 | | P07395 | P07395 | Phenylalanine--tRNA ligase beta subunit |
| *yrdA* | -0.337 | 0.757 | | P0A9W9 | P0A9W9 | Protein YrdA |
| *fumA* | -0.628 | 3.138 | | P0AC33 | P0AC33 | Fumarate hydratase class I, aerobic |
| *pepP* | -0.627 | 2.983 | | P15034 | P15034 | Xaa-Pro aminopeptidase |
| *speE* | -0.627 | 3.523 | | P09158 | P09158 | Polyamine aminopropyltransferase |
| *glnB* | 0.273 | 0.744 | | P0A9Z1 | P0A9Z1 | Nitrogen regulatory protein P-II 1 |
| *eno* | -0.626 | 6.973 | | P0A6P9 | P0A6P9 | Enolase |
| *rplM* | -0.089 | 0.483 | | P0AA10 | P0AA10 | 50S ribosomal protein L13 |
| *ruvA* | -0.624 | 2.589 | | P0A809 | P0A809 | Holliday junction ATP-dependent DNA helicase RuvA |
| *cysS* | -0.620 | 2.773 | | P21888 | P21888 | Cysteine--tRNA ligase |
| *menB* | -0.619 | 1.471 | | P0ABU0 | P0ABU0 | 1,4-dihydroxy-2-naphthoyl-CoA synthase |
| *trmJ* | -0.618 | 2.255 | | P0AE01 | P0AE01 | tRNA (cytidine/uridine-2-O-)-methyltransferase TrmJ |
| *csdE* | -0.617 | 1.896 | | P0AGF2 | P0AGF2 | Sulfur acceptor protein CsdE |
| *pdxK* | -0.615 | 3.168 | | P40191 | P40191 | Pyridoxine kinase |
| *yacC* | -0.229 | 1.163 | | P0AA95 | P0AA95 | Uncharacterized protein YacC |
| *yaeQ* | -0.047 | 0.089 | | P0AA97 | P0AA97 | Uncharacterized protein YaeQ |
| *tolB* | -0.609 | 2.484 | | P0A855 | P0A855 | Protein TolB |
| *ubiD* | 0.017 | 0.047 | | P0AAB4 | P0AAB4 | 3-octaprenyl-4-hydroxybenzoate carboxy-lyase |
| *baeR* | -0.605 | 4.258 | | P69228 | P69228 | Transcriptional regulatory protein BaeR |
| *ygdH* | -0.598 | 2.630 | | P0ADR8 | P0ADR8 | LOG family protein YgdH |
| *pncA* | -0.596 | 2.513 | | P21369 | P21369 | Pyrazinamidase/nicotinamidase |
| *trpA* | -0.595 | 1.733 | | P0A877 | P0A877 | Tryptophan synthase alpha chain |
| *rpmJ* | -0.593 | 1.480 | | P0A7Q6 | P0A7Q6 | 50S ribosomal protein L36 |
| *cycA* | 0.507 | 1.236 | | P0AAE0 | P0AAE0 | D-serine/D-alanine/glycine transporter |
| *artP* | -0.227 | 0.081 | | P0AAF6 | P0AAF6 | Arginine transport ATP-binding protein ArtP |
| *dppD* | 0.261 | 0.834 | | P0AAG0 | P0AAG0 | Dipeptide transport ATP-binding protein DppD |
| *comR* | -0.592 | 3.065 | | P75952 | P75952 | HTH-type transcriptional repressor ComR |
| *hemC* | -0.589 | 2.628 | | P06983 | P06983 | Porphobilinogen deaminase |
| *sapD* | -0.197 | 0.377 | | P0AAH4 | P0AAH4 | Peptide transport system ATP-binding protein SapD |
| *sapF* | 0.324 | 1.111 | | P0AAH8 | P0AAH8 | Peptide transport system ATP-binding protein SapF |
| *ynbE* | -0.587 | 2.666 | | P64448 | P64448 | Uncharacterized protein YnbE |
| *clpP* | -0.586 | 2.717 | | P0A6G7 | P0A6G7 | ATP-dependent Clp protease proteolytic subunit |
| *fabD* | -0.027 | 0.119 | | P0AAI9 | P0AAI9 | Malonyl CoA-acyl carrier protein transacylase |
| *tldD* | -0.572 | 3.541 | | P0AGG8 | P0AGG8 | Metalloprotease TldD |
| *queE* | -0.566 | 1.369 | | P64554 | P64554 | 7-carboxy-7-deazaguanine synthase |
| *ybaK* | 0.123 | 0.498 | | P0AAR3 | P0AAR3 | Cys-tRNA(Pro)/Cys-tRNA(Cys) deacylase YbaK |
| *polA* | -0.564 | 1.807 | | P00582 | P00582 | DNA polymerase I |
| *guaB* | -0.555 | 3.725 | | P0ADG7 | P0ADG7 | Inosine-5-monophosphate dehydrogenase |
| *ompA* | -0.539 | 1.888 | | P0A910 | P0A910 | Outer membrane protein A |

| Gene names | og2(fold-Change | -Log(p-value) | Significant | Protein IDs | Majority protein IDs | Protein names |
|---|---|---|---|---|---|---|
| *guaA* | -0.535 | 2.932 | | P04079 | P04079 | GMP synthase [glutamine-hydrolyzing] |
| *ybiS* | 0.310 | 1.240 | | P0AAX8 | P0AAX8 | Probable L,D-transpeptidase YbiS |
| *rpsB* | -0.531 | 3.596 | | P0A7V0 | P0A7V0 | 30S ribosomal protein S2 |
| *ycaR* | -0.011 | 0.042 | | P0AAZ7 | P0AAZ7 | UPF0434 protein YcaR |
| *ymbA* | 0.153 | 0.271 | | P0AB10 | P0AB10 | Uncharacterized lipoprotein YmbA |
| *yccF* | -0.438 | 0.998 | | P0AB12 | P0AB12 | Inner membrane protein YccF |
| *tusE* | 0.554 | 0.816 | | P0AB18 | P0AB18 | Sulfurtransferase TusE |
| *ybiT* | -0.531 | 3.011 | | P0A9U3 | P0A9U3 | Uncharacterized ABC transporter ATP-binding protein YbiT |
| *pncB* | -0.525 | 2.905 | | P18133 | P18133 | Nicotinate phosphoribosyltransferase |
| *yceD* | -0.040 | 0.333 | | P0AB28 | P0AB28 | Uncharacterized protein YceD |
| *cysM* | -0.525 | 1.357 | | P16703 | P16703 | Cysteine synthase B |
| *ycgL* | 0.397 | 1.173 | | P0AB43 | P0AB43 | Protein YcgL |
| *ychN* | 0.499 | 1.161 | | P0AB52 | P0AB52 | Protein YchN |
| *yciI* | -0.215 | 0.471 | | P0AB55 | P0AB55 | Protein YciI |
| *lapB* | -0.833 | 1.222 | | P0AB58 | P0AB58 | Lipopolysaccharide assembly protein B |
| *thiL* | -0.520 | 2.765 | | P0AGG0 | P0AGG0 | Thiamine-monophosphate kinase |
| *yigL* | -0.514 | 2.168 | | P27848 | P27848 | Pyridoxal phosphate phosphatase YigL |
| *pntB* | 0.229 | 0.796 | | P0AB67 | P0AB67 | NAD(P) transhydrogenase subunit beta |
| *yafC* | -0.512 | 1.445 | | P30864 | P30864 | Uncharacterized HTH-type transcriptional regulator YafC |
| *yafV* | -0.512 | 2.003 | | Q47679 | Q47679 | Hydrolase YafV |
| *yfbR* | -0.505 | 1.412 | | P76491 | P76491 | 5-deoxynucleotidase YfbR |
| *nth* | -0.003 | 0.006 | | P0AB83 | P0AB83 | Endonuclease III |
| *pheS* | -0.504 | 3.136 | | P08312 | P08312 | Phenylalanine--tRNA ligase alpha subunit |
| *ybhK* | -0.503 | 2.347 | | P75767 | P75767 | Putative gluconeogenesis factor |
| *nagC* | -0.502 | 2.113 | | P0AF20 | P0AF20 | N-acetylglucosamine repressor |
| *rplL* | -0.502 | 1.735 | | P0A7K2 | P0A7K2 | 50S ribosomal protein L7/L12 |
| *rob* | -0.501 | 1.828 | | P0ACI0 | P0ACI0 | Right origin-binding protein |
| *rsd* | -0.499 | 2.484 | | P0AFX4 | P0AFX4 | Regulator of sigma D |
| *yfhM* | -0.496 | 1.431 | | P76578 | P76578 | Uncharacterized lipoprotein YfhM |
| *lolA* | -0.495 | 1.335 | | P61316 | P61316 | Outer-membrane lipoprotein carrier protein |
| *hflC* | 0.147 | 1.244 | | P0ABC3 | P0ABC3 | Modulator of FtsH protease HflC |
| *hflK* | 0.150 | 0.667 | | P0ABC7 | P0ABC7 | Modulator of FtsH protease HflK |
| *asnA* | -0.492 | 2.412 | | P00963 | P00963 | Aspartate--ammonia ligase |
| *zapA* | -0.489 | 1.546 | | P0ADS2 | P0ADS2 | Cell division protein ZapA |
| *accB* | 0.453 | 0.560 | | P0ABD8 | P0ABD8 | Biotin carboxyl carrier protein of acetyl-CoA carboxylase |
| *nfsA* | -0.488 | 4.678 | | P17117 | P17117 | Oxygen-insensitive NADPH nitroreductase |
| *pcnB* | 0.095 | 0.334 | | P0ABF1 | P0ABF1 | Poly(A) polymerase I |
| *phoP* | -0.487 | 2.687 | | P23836 | P23836 | Transcriptional regulatory protein PhoP |
| *ftsA* | 0.188 | 0.784 | | P0ABH0 | P0ABH0 | Cell division protein FtsA |
| *sspB* | -0.487 | 2.055 | | P0AFZ3 | P0AFZ3 | Stringent starvation protein B |
| *ispG* | -0.479 | 3.045 | | P62620 | P62620 | 4-hydroxy-3-methylbut-2-en-1-yl diphosphate synthase (flavodoxin) |
| *lplA* | -0.478 | 1.708 | | P32099 | P32099 | Lipoate-protein ligase A |
| *rdoA* | -0.477 | 2.445 | | P0C0K3 | P0C0K3 | Protein RdoA |
| *hslO* | -0.470 | 2.685 | | P0A6Y5 | P0A6Y5 | 33 kDa chaperonin |
| *aspS* | -0.467 | 2.223 | | P21889 | P21889 | Aspartate--tRNA ligase |
| *thiE* | -0.465 | 1.620 | | P30137 | P30137 | Thiamine-phosphate synthase |
| *infA* | -0.462 | 1.662 | | P69222 | P69222 | Translation initiation factor IF-1 |
| *yebC* | -0.461 | 3.569 | | P0A8A0 | P0A8A0 | Probable transcriptional regulatory protein YebC |
| *arcB* | -0.459 | 1.492 | | P0AEC3 | P0AEC3 | Aerobic respiration control sensor protein ArcB |
| *coaBC* | 0.119 | 0.253 | | P0ABQ0 | P0ABQ0 | Coenzyme A biosynthesis bifunctional protein CoaBC |
| *folA* | -0.079 | 0.254 | | P0ABQ4 | P0ABQ4 | Dihydrofolate reductase |
| *dksA* | -0.115 | 0.955 | | P0ABS1 | P0ABS1 | RNA polymerase-binding transcription factor DksA |
| *rsmC* | -0.456 | 2.436 | | P39406 | P39406 | Ribosomal RNA small subunit methyltransferase C |

| Gene names | og2(fold-Change | -Log(p-value) | Significant | Protein IDs | Majority protein IDs | Protein names |
|---|---|---|---|---|---|---|
| *selD* | -0.455 | 5.026 | | P16456 | P16456 | Selenide, water dikinase |
| *mlaE* | -0.448 | 1.512 | | P64606 | P64606 | Probable phospholipid ABC transporter permease protein MlaE |
| *elbB* | 0.070 | 0.183 | | P0ABU5 | P0ABU5 | Enhancing lycopene biosynthesis protein 2 |
| *exbB* | 1.279 | 1.252 | | P0ABU7 | P0ABU7 | Biopolymer transport protein ExbB |
| *tolQ* | 0.162 | 0.824 | | P0ABU9 | P0ABU9 | Protein TolQ |
| *tolR* | 0.139 | 0.855 | | P0ABV6 | P0ABV6 | Protein TolR |
| *hisS* | -0.448 | 2.312 | | P60906 | P60906 | Histidine--tRNA ligase |
| *kdsC* | 0.201 | 0.859 | | P0ABZ4 | P0ABZ4 | 3-deoxy-D-manno-octulosonate 8-phosphate phosphatase KdsC |
| *ftsZ* | -0.445 | 3.601 | | P0A9A6 | P0A9A6 | Cell division protein FtsZ |
| *bamD* | -0.296 | 0.836 | | P0AC02 | P0AC02 | Outer membrane protein assembly factor BamD |
| *aceB* | -0.444 | 2.217 | | P08997 | P08997 | Malate synthase A |
| *folB* | 0.257 | 0.511 | | P0AC16 | P0AC16 | Dihydroneopterin aldolase |
| *yfcE* | -0.440 | 1.438 | | P67095 | P67095 | Phosphodiesterase YfcE |
| *focA* | 0.213 | 0.807 | | P0AC23 | P0AC23 | Probable formate transporter 1 |
| *ftsX* | -0.278 | 0.879 | | P0AC30 | P0AC30 | Cell division protein FtsX |
| *yqhD* | -0.431 | 1.896 | | Q46856 | Q46856 | Alcohol dehydrogenase YqhD |
| *sdhA* | 0.240 | 1.159 | | P0AC41 | P0AC41 | Succinate dehydrogenase flavoprotein subunit |
| *ybbN* | -0.430 | 3.247 | | P77395 | P77395 | Uncharacterized protein YbbN |
| *ydgJ* | -0.427 | 1.425 | | P77376 | P77376 | Uncharacterized oxidoreductase YdgJ |
| *ycgM* | -0.426 | 1.671 | | P76004 | P76004 | Uncharacterized protein YcgM |
| *cutC* | -0.426 | 2.200 | | P67826 | P67826 | Copper homeostasis protein CutC |
| *grxD* | 0.174 | 0.419 | | P0AC69 | P0AC69 | Glutaredoxin-4 |
| *rseB* | -0.425 | 2.830 | | P0AFX9 | P0AFX9 | Sigma-E factor regulatory protein RseB |
| *yfiF* | -0.425 | 2.536 | | P0AGJ5 | P0AGJ5 | Uncharacterized tRNA/rRNA methyltransferase YfiF |
| *glgP* | 0.552 | 1.208 | | P0AC86 | P0AC86 | Glycogen phosphorylase |
| *frmA* | -0.420 | 1.857 | | P25437 | P25437 | S-(hydroxymethyl)glutathione dehydrogenase |
| *glcB* | -0.419 | 2.276 | | P37330 | P37330 | Malate synthase G |
| *gutQ* | -0.415 | 2.320 | | P17115 | P17115 | Arabinose 5-phosphate isomerase GutQ |
| *dnaB* | 0.303 | 1.024 | | P0ACB0 | P0ACB0 | Replicative DNA helicase |
| *hemB* | -0.080 | 1.038 | | P0ACB2 | P0ACB2 | Delta-aminolevulinic acid dehydratase |
| *hemG* | -0.781 | 0.846 | | P0ACB4 | P0ACB4 | Protoporphyrinogen IX dehydrogenase [menaquinone] |
| *hemY* | -0.149 | 0.930 | | P0ACB7 | P0ACB7 | Protein HemY |
| *prmC* | -0.278 | 0.413 | | P0ACC1 | P0ACC1 | Release factor glutamine methyltransferase |
| *mltC* | -0.414 | 1.630 | | P0C066 | P0C066 | Membrane-bound lytic murein transglycosylase C |
| *tehB* | -0.413 | 1.703 | | P25397 | P25397 | Tellurite methyltransferase |
| *ydjA* | -0.413 | 2.113 | | P0ACY1 | P0ACY1 | Putative NAD(P)H nitroreductase YdjA |
| *rplQ* | -0.410 | 1.439 | | P0AG44 | P0AG44 | 50S ribosomal protein L17 |
| *yciK* | -0.409 | 2.187 | | P31808 | P31808 | Uncharacterized oxidoreductase YciK |
| *hupA* | -0.117 | 0.380 | | P0ACF0 | P0ACF0 | DNA-binding protein HU-alpha |
| *galU* | -0.409 | 2.381 | | P0AEP3 | P0AEP3 | UTP--glucose-1-phosphate uridylyltransferase |
| *hns* | -0.137 | 0.304 | | P0ACF8 | P0ACF8 | DNA-binding protein H-NS |
| *agaR* | -0.402 | 1.614 | | P0ACK2 | P0ACK2 | Putative aga operon transcriptional repressor |
| *mtnN* | -0.401 | 1.860 | | P0AF12 | P0AF12 | 5-methylthioadenosine/S-adenosylhomocysteine nucleosidase |
| *lrp* | -0.090 | 0.191 | | P0ACJ0 | P0ACJ0 | Leucine-responsive regulatory protein |
| *pfkA* | -0.397 | 3.316 | | P0A796 | P0A796 | ATP-dependent 6-phosphofructokinase isozyme 1 |
| *gsk* | -0.395 | 2.443 | | P0AEW6 | P0AEW6 | Inosine-guanosine kinase |
| *deoR* | -0.270 | 1.062 | | P0ACK5 | P0ACK5 | Deoxyribose operon repressor |
| *serC* | -0.391 | 2.370 | | P23721 | P23721 | Phosphoserine aminotransferase |
| *ileS* | -0.390 | 2.010 | | P00956 | P00956 | Isoleucine--tRNA ligase |
| *rspR* | -0.389 | 1.182 | | P0ACM2 | P0ACM2 | HTH-type transcriptional repressor RspR |
| *pepN* | -0.390 | 2.234 | | P04825 | P04825 | Aminopeptidase N |
| *rraA* | -0.387 | 1.608 | | P0A8R0 | P0A8R0 | Regulator of ribonuclease activity A |
| *gshA* | -0.385 | 1.772 | | P0A6W9 | P0A6W9 | Glutamate--cysteine ligase |

| Gene names | og2(fold-Change | -Log(p-value) | Significant | Protein IDs | Majority protein IDs | Protein names |
|---|---|---|---|---|---|---|
| *gntR* | -0.016 | 0.038 | | P0ACP5 | P0ACP5 | HTH-type transcriptional regulator GntR |
| *purR* | -0.064 | 0.237 | | P0ACP7 | P0ACP7 | HTH-type transcriptional repressor PurR |
| *hemN* | -0.381 | 1.411 | | P32131 | P32131 | Oxygen-independent coproporphyrinogen-III oxidase |
| *trpB* | -0.381 | 2.611 | | P0A879 | P0A879 | Tryptophan synthase beta chain |
| *yhhW* | -0.380 | 1.609 | | P46852 | P46852 | Quercetin 2,3-dioxygenase |
| *asnB* | -0.380 | 1.913 | | P22106 | P22106 | Asparagine synthetase B [glutamine-hydrolyzing] |
| *uidR* | 0.041 | 0.070 | | P0ACT6 | P0ACT6 | HTH-type transcriptional regulator UidR |
| *rutR* | 0.519 | 0.596 | | P0ACU2 | P0ACU2 | HTH-type transcriptional regulator RutR |
| *tas* | -0.378 | 3.030 | | P0A9T4 | P0A9T4 | Protein tas |
| *zipA* | -0.377 | 2.188 | | P77173 | P77173 | Cell division protein ZipA |
| *lpxL* | -0.864 | 1.093 | | P0ACV0 | P0ACV0 | Lipid A biosynthesis lauroyltransferase |
| *rcsD* | -0.371 | 1.663 | | P39838 | P39838 | Phosphotransferase RcsD |
| *ydcH* | 0.865 | 0.409 | | P0ACW6 | P0ACW6 | Uncharacterized protein YdcH |
| *ydhR* | 0.320 | 0.642 | | P0ACX3 | P0ACX3 | Putative monooxygenase YdhR |
| *ybfF* | -0.370 | 2.132 | | P75736 | P75736 | Esterase YbfF |
| *galF* | -0.367 | 3.794 | | P0AAB6 | P0AAB6 | UTP--glucose-1-phosphate uridylyltransferase |
| *cusR* | -0.023 | 0.068 | | P0ACZ8 | P0ACZ8 | Transcriptional regulatory protein CusR |
| *dcuR* | -0.162 | 0.683 | | P0AD01 | P0AD01 | Transcriptional regulatory protein DcuR |
| *tgt* | -0.361 | 3.295 | | P0A847 | P0A847 | Queuine tRNA-ribosyltransferase |
| *yecJ* | 0.082 | 0.187 | | P0AD10 | P0AD10 | Uncharacterized protein YecJ |
| *hflX* | -0.351 | 1.354 | | P25519 | P25519 | GTPase HflX |
| *yejL* | -0.292 | 0.875 | | P0AD24 | P0AD24 | UPF0352 protein YejL |
| *yejM* | 0.082 | 0.217 | | P0AD27 | P0AD27 | Inner membrane protein YejM |
| *yfcZ* | 0.251 | 1.240 | | P0AD33 | P0AD33 | UPF0381 protein YfcZ |
| *ygaC* | 0.522 | 0.532 | | P0AD53 | P0AD53 | Uncharacterized protein YgaC |
| *ydfG* | -0.346 | 2.232 | | P39831 | P39831 | NADP-dependent 3-hydroxy acid dehydrogenase YdfG |
| *hfq* | -0.344 | 2.012 | | P0A6X3 | P0A6X3 | RNA-binding protein Hfq |
| *dacB* | -0.343 | 3.175 | | P24228 | P24228 | D-alanyl-D-alanine carboxypeptidase DacB |
| *degQ* | -0.340 | 2.009 | | P39099 | P39099 | Periplasmic pH-dependent serine endoprotease DegQ |
| *dld* | -0.340 | 2.026 | | P06149 | P06149 | D-lactate dehydrogenase |
| *nudE* | -0.333 | 3.121 | | P45799 | P45799 | ADP compounds hydrolase NudE |
| *tesA* | -0.197 | 0.375 | | P0ADA1 | P0ADA1 | Acyl-CoA thioesterase I |
| *ispB* | -0.328 | 2.621 | | P0AD57 | P0AD57 | Octaprenyl-diphosphate synthase |
| *kdgK* | -0.328 | 1.517 | | P37647 | P37647 | 2-dehydro-3-deoxygluconokinase |
| *oxyR* | -0.321 | 1.599 | | P0ACQ4 | P0ACQ4 | Hydrogen peroxide-inducible genes activator |
| *pyrD* | -0.316 | 1.371 | | P0A7E1 | P0A7E1 | Dihydroorotate dehydrogenase (quinone) |
| *lptE* | -0.100 | 0.298 | | P0ADC1 | P0ADC1 | LPS-assembly lipoprotein LptE |
| *lolC* | -0.023 | 0.050 | | P0ADC3 | P0ADC3 | Lipoprotein-releasing system transmembrane protein LolC |
| *lasT* | -0.316 | 1.946 | | P37005 | P37005 | Uncharacterized tRNA/rRNA methyltransferase LasT |
| *fusA* | -0.314 | 1.341 | | P0A6M8 | P0A6M8 | Elongation factor G |
| *ybiC* | -0.305 | 1.390 | | P30178 | P30178 | Uncharacterized oxidoreductase YbiC |
| *ygfZ* | 0.218 | 1.261 | | P0ADE8 | P0ADE8 | tRNA-modifying protein YgfZ |
| *pdxJ* | -0.305 | 1.702 | | P0A794 | P0A794 | Pyridoxine 5-phosphate synthase |
| *lon* | -0.299 | 1.841 | | P0A9M0 | P0A9M0 | Lon protease |
| *lysS* | -0.297 | 1.597 | | P0A8N3 | P0A8N3 | Lysine--tRNA ligase |
| *pflA* | -0.291 | 1.472 | | P0A9N4 | P0A9N4 | Pyruvate formate-lyase 1-activating enzyme |
| *yiaF* | -0.028 | 0.083 | | P0ADK0 | P0ADK0 | Uncharacterized protein YiaF |
| *yibL* | -0.235 | 0.957 | | P0ADK8 | P0ADK8 | Uncharacterized protein YibL |
| *nanR* | -0.282 | 1.474 | | P0A8W0 | P0A8W0 | Transcriptional regulator NanR |
| *yifL* | 0.517 | 0.437 | | P0ADN6 | P0ADN6 | Uncharacterized lipoprotein YifL |
| *cysB* | -0.281 | 1.333 | | P0A9F3 | P0A9F3 | HTH-type transcriptional regulator CysB |
| *hpt* | -0.280 | 1.489 | | P0A9M2 | P0A9M2 | Hypoxanthine phosphoribosyltransferase |
| *mlaD* | -0.278 | 5.423 | | P64604 | P64604 | Probable phospholipid ABC transporter-binding protein MlaD |

| Gene names | og2(fold-Change | -Log(p-value) | Significant | Protein IDs | Majority protein IDs | Protein names |
|---|---|---|---|---|---|---|
| *murB* | -0.272 | 1.365 | | P08373 | P08373 | UDP-N-acetylenolpyruvoylglucosamine reductase |
| *rlmM* | -0.439 | 1.156 | | P0ADR6 | P0ADR6 | Ribosomal RNA large subunit methyltransferase M |
| *accD* | -0.271 | 2.770 | | P0A9Q5 | P0A9Q5 | Acetyl-coenzyme A carboxylase carboxyl transferase subunit beta |
| *pdxH* | -0.270 | 1.467 | | P0AFI7 | P0AFI7 | Pyridoxine/pyridoxamine 5-phosphate oxidase |
| *yceI* | -0.266 | 1.313 | | P0A8X2 | P0A8X2 | Protein YceI |
| *accA* | -0.262 | 3.432 | | P0ABD5 | P0ABD5 | Acetyl-coenzyme A carboxylase carboxyl transferase subunit alpha |
| *proC* | -0.257 | 1.405 | | P0A9L8 | P0A9L8 | Pyrroline-5-carboxylate reductase |
| *cpxR* | -0.256 | 2.625 | | P0AE88 | P0AE88 | Transcriptional regulatory protein CpxR |
| *nagK* | -0.253 | 1.428 | | P75959 | P75959 | N-acetyl-D-glucosamine kinase |
| *ygiN* | -0.096 | 0.303 | | P0ADU2 | P0ADU2 | Probable quinol monooxygenase YgiN |
| *ygiW* | 0.128 | 0.120 | | P0ADU5 | P0ADU5 | Protein YgiW |
| *lptA* | 0.032 | 0.102 | | P0ADV1 | P0ADV1 | Lipopolysaccharide export system protein LptA |
| *mlaC* | -0.171 | 0.662 | | P0ADV7 | P0ADV7 | Probable phospholipid-binding protein MlaC |
| *prfA* | -0.252 | 1.405 | | P0A7I0 | P0A7I0 | Peptide chain release factor 1 |
| *wzzB* | -0.251 | 2.571 | | P76372 | P76372 | Chain length determinant protein |
| *pgm* | -0.250 | 3.053 | | P36938 | P36938 | Phosphoglucomutase |
| *ecpC* | -0.243 | 1.600 | | P77802 | P77802 | Probable outer membrane usher protein EcpC |
| *nudC* | -0.237 | 1.634 | | P32664 | P32664 | NADH pyrophosphatase |
| *rplN* | -0.013 | 0.039 | | P0ADY3 | P0ADY3 | 50S ribosomal protein L14 |
| *rplP* | -0.719 | 0.787 | | P0ADY7 | P0ADY7 | 50S ribosomal protein L16 |
| *rplW* | -0.191 | 0.903 | | P0ADZ0 | P0ADZ0 | 50S ribosomal protein L23 |
| *rpsO* | 0.018 | 0.057 | | P0ADZ4 | P0ADZ4 | 30S ribosomal protein S15 |
| *mlaF* | -0.236 | 1.703 | | P63386 | P63386 | Probable phospholipid import ATP-binding protein MlaF |
| *lipB* | -0.235 | 1.412 | | P60720 | P60720 | Octanoyltransferase |
| *acrA* | 0.013 | 0.081 | | P0AE06 | P0AE06 | Multidrug efflux pump subunit AcrA |
| *rplF* | -0.227 | 1.449 | | P0AG55 | P0AG55 | 50S ribosomal protein L6 |
| *lpdA* | -0.225 | 1.500 | | P0A9P0 | P0A9P0 | Dihydrolipoyl dehydrogenase |
| *ydhF* | -0.216 | 1.901 | | P76187 | P76187 | Oxidoreductase YdhF |
| *aroM* | 0.399 | 0.757 | | P0AE28 | P0AE28 | Protein AroM |
| *mrp* | -0.193 | 2.314 | | P0AF08 | P0AF08 | Protein mrp |
| *ytfP* | 0.206 | 0.565 | | P0AE48 | P0AE48 | Gamma-glutamylcyclotransferase family protein YtfP |
| *bcp* | 0.248 | 0.717 | | P0AE52 | P0AE52 | Putative peroxiredoxin bcp |
| *rplA* | -0.190 | 1.516 | | P0A7L0 | P0A7L0 | 50S ribosomal protein L1 |
| *fabI* | -0.158 | 1.438 | | P0AEK4 | P0AEK4 | Enoyl-[acyl-carrier-protein] reductase [NADH] FabI |
| *ruvB* | -0.152 | 1.571 | | P0A812 | P0A812 | Holliday junction ATP-dependent DNA helicase RuvB |
| *rlmI* | -0.149 | 1.472 | | P75876 | P75876 | Ribosomal RNA large subunit methyltransferase I |
| *cysG* | -0.015 | 0.051 | | P0AEA8 | P0AEA8 | Siroheme synthase |
| *crp* | -0.142 | 1.768 | | P0ACJ8 | P0ACJ8 | cAMP-activated global transcriptional regulator CRP |
| *yoaB* | -0.419 | 1.012 | | P0AEB7 | P0AEB7 | RutC family protein YoaB |
| *acnB* | -0.109 | 1.436 | | P36683 | P36683 | Aconitate hydratase B |
| *ttcA* | -0.107 | 1.675 | | P76055 | P76055 | tRNA 2-thiocytidine biosynthesis protein TtcA |
| *barA* | -0.154 | 0.247 | | P0AEC5 | P0AEC5 | Signal transduction histidine-protein kinase BarA |
| *mnmA* | -0.058 | 1.583 | | P25745 | P25745 | tRNA-specific 2-thiouridylase MnmA |
| *dapE* | 0.150 | 0.341 | | P0AED7 | P0AED7 | Succinyl-diaminopimelate desuccinylase |
| *dcrB* | -0.100 | 0.287 | | P0AEE1 | P0AEE1 | Protein DcrB |
| *degS* | -0.098 | 0.410 | | P0AEE3 | P0AEE3 | Serine endoprotease DegS |
| *dnaJ* | 0.069 | 1.550 | | P08622 | P08622 | Chaperone protein DnaJ |
| *dnaC* | 0.259 | 0.910 | | P0AEF0 | P0AEF0 | DNA replication protein DnaC |
| *dppB* | 0.303 | 0.324 | | P0AEF8 | P0AEF8 | Dipeptide transport system permease protein DppB |
| *trpS* | 0.074 | 1.517 | | P00954 | P00954 | Tryptophan--tRNA ligase |
| *sucB* | 0.092 | 1.524 | | P0AFG6 | P0AFG6 | Dihydrolipoyllysine-residue succinyltransferase |
| *rseP* | -0.225 | 0.472 | | P0AEH1 | P0AEH1 | Regulator of sigma-E protease RseP |
| *elaA* | 0.240 | 1.062 | | P0AEH3 | P0AEH3 | Protein ElaA |

| Gene names | og2(fold-Change | -Log(p-value) | Significant | Protein IDs | Majority protein IDs | Protein names |
|---|---|---|---|---|---|---|
| *rsmH* | 0.093 | 1.724 | | P60390 | P60390 | Ribosomal RNA small subunit methyltransferase H |
| *proA* | 0.114 | 1.411 | | P07004 | P07004 | Gamma-glutamyl phosphate reductase |
| *rimO* | -0.067 | 0.378 | | P0AEI4 | P0AEI4 | Ribosomal protein S12 methylthiotransferase RimO |
| *tyrB* | 0.135 | 2.999 | | P04693 | P04693 | Aromatic-amino-acid aminotransferase |
| *fabG* | 0.073 | 0.553 | | P0AEK2 | P0AEK2 | 3-oxoacyl-[acyl-carrier-protein] reductase FabG |
| *rfbD* | 0.139 | 2.691 | | P37760 | P37760 | dTDP-4-dehydrorhamnose reductase |
| *fepB* | -0.857 | 0.758 | | P0AEL6 | P0AEL6 | Ferrienterobactin-binding periplasmic protein |
| *acuI* | 0.146 | 1.657 | | P26646 | P26646 | Probable acrylyl-CoA reductase AcuI |
| *pepQ* | 0.190 | 1.576 | | P21165 | P21165 | Xaa-Pro dipeptidase |
| *purC* | 0.204 | 1.789 | | P0A7D7 | P0A7D7 | Phosphoribosylaminoimidazole-succinocarboxamide synthase |
| *prc* | 0.209 | 1.598 | | P23865 | P23865 | Tail-specific protease |
| *rsmA* | 0.214 | 1.815 | | P06992 | P06992 | Ribosomal RNA small subunit methyltransferase A |
| *tig* | 0.222 | 2.183 | | P0A850 | P0A850 | Trigger factor |
| *rmlA1* | 0.227 | 2.129 | | P37744 | P37744 | Glucose-1-phosphate thymidylyltransferase 1 |
| *lptB* | 0.228 | 1.804 | | P0A9V1 | P0A9V1 | Lipopolysaccharide export system ATP-binding protein LptB |
| *hisJ* | -1.211 | 0.547 | | P0AEU0 | P0AEU0 | Histidine-binding periplasmic protein |
| *skp* | -0.369 | 1.159 | | P0AEU7 | P0AEU7 | Chaperone protein Skp |
| *cpdA* | 0.213 | 0.712 | | P0AEW4 | P0AEW4 | 3,5-cyclic adenosine monophosphate phosphodiesterase CpdA |
| *bepA* | 0.228 | 1.559 | | P66948 | P66948 | Beta-barrel assembly-enhancing protease |
| *thyA* | 0.240 | 1.765 | | P0A884 | P0A884 | Thymidylate synthase |
| *atpG* | 0.247 | 2.215 | | P0ABA6 | P0ABA6 | ATP synthase gamma chain |
| *pyrH* | 0.251 | 2.219 | | P0A7E9 | P0A7E9 | Uridylate kinase |
| *purN* | 0.262 | 2.188 | | P08179 | P08179 | Phosphoribosylglycinamide formyltransferase |
| *nadC* | 0.267 | 1.943 | | P30011 | P30011 | Nicotinate-nucleotide pyrophosphorylase [carboxylating] |
| *minD* | 0.033 | 0.163 | | P0AEZ3 | P0AEZ3 | Septum site-determining protein MinD |
| *rne* | 0.268 | 1.409 | | P21513 | P21513 | Ribonuclease E |
| *rimL* | 0.269 | 1.600 | | P13857 | P13857 | Ribosomal-protein-serine acetyltransferase |
| *ddlB* | 0.270 | 1.324 | | P07862 | P07862 | D-alanine--D-alanine ligase B |
| *surA* | 0.281 | 1.496 | | P0ABZ6 | P0ABZ6 | Chaperone SurA |
| *bglA* | 0.287 | 1.354 | | Q46829 | Q46829 | 6-phospho-beta-glucosidase BglA |
| *prs* | 0.288 | 2.175 | | P0A717 | P0A717 | Ribose-phosphate pyrophosphokinase |
| *nagD* | 0.036 | 0.114 | | P0AF24 | P0AF24 | Ribonucleotide monophosphatase NagD |
| *narL* | 0.043 | 0.180 | | P0AF28 | P0AF28 | Nitrate/nitrite response regulator protein NarL |
| *zapB* | 0.070 | 0.198 | | P0AF36 | P0AF36 | Cell division protein ZapB |
| *yjbR* | 0.013 | 0.028 | | P0AF50 | P0AF50 | Uncharacterized protein YjbR |
| *nsrR* | 0.212 | 0.874 | | P0AF63 | P0AF63 | HTH-type transcriptional repressor NsrR |
| *tsaE* | 0.090 | 0.309 | | P0AF67 | P0AF67 | tRNA threonylcarbamoyladenosine biosynthesis protein TsaE |
| *hemL* | 0.292 | 2.731 | | P23893 | P23893 | Glutamate-1-semialdehyde 2,1-aminomutase |
| *yceF* | 0.304 | 1.753 | | P0A729 | P0A729 | Maf-like protein YceF |
| *rraB* | -0.050 | 0.113 | | P0AF90 | P0AF90 | Regulator of ribonuclease activity B |
| *groL* | 0.306 | 2.284 | | P0A6F5 | P0A6F5 | 60 kDa chaperonin |
| *lptF* | 0.333 | 0.891 | | P0AF98 | P0AF98 | Lipopolysaccharide export system permease protein LptF |
| *nlpI* | 0.244 | 0.862 | | P0AFB1 | P0AFB1 | Lipoprotein NlpI |
| *glnL* | -0.958 | 1.046 | | P0AFB5 | P0AFB5 | Nitrogen regulation protein NR(II) |
| *glnG* | 0.185 | 0.290 | | P0AFB8 | P0AFB8 | Nitrogen regulation protein NR(I) |
| *purM* | 0.313 | 1.709 | | P08178 | P08178 | Phosphoribosylformylglycinamidine cyclo-ligase |
| *gltX* | 0.314 | 2.866 | | P04805 | P04805 | Glutamate--tRNA ligase |
| *uup* | 0.315 | 1.649 | | P43672 | P43672 | ABC transporter ATP-binding protein uup |
| *pssA* | 0.319 | 1.389 | | P23830 | P23830 | CDP-diacylglycerol--serine O-phosphatidyltransferase |
| *fdhE* | 0.320 | 3.223 | | P13024 | P13024 | Protein FdhE |
| *yhhX* | 0.323 | 1.634 | | P46853 | P46853 | Uncharacterized oxidoreductase YhhX |
| *nuoN* | 0.613 | 0.739 | | P0AFF0 | P0AFF0 | NADH-quinone oxidoreductase subunit N |
| *nupC* | -0.086 | 0.229 | | P0AFF2 | P0AFF2 | Nucleoside permease NupC |

| Gene names | og2(fold-Change | -Log(p-value) | Significant | Protein IDs | Majority protein IDs | Protein names |
|---|---|---|---|---|---|---|
| *nusA* | -0.050 | 0.209 | | P0AFF6 | P0AFF6 | Transcription termination/antitermination protein NusA |
| *lipA* | 0.325 | 3.243 | | P60716 | P60716 | Lipoyl synthase |
| *sucA* | 0.124 | 0.999 | | P0AFG3 | P0AFG3 | 2-oxoglutarate dehydrogenase E1 component |
| *udk* | 0.327 | 2.177 | | P0A8F4 | P0A8F4 | Uridine kinase |
| *rapZ* | 0.330 | 1.708 | | P0A894 | P0A894 | RNase adapter protein RapZ |
| *ogt* | 0.248 | 1.156 | | P0AFH0 | P0AFH0 | Methylated-DNA--protein-cysteine methyltransferase |
| *prkB* | 0.330 | 2.799 | | P0AEX5 | P0AEX5 | Probable phosphoribulokinase |
| *rbfA* | 0.336 | 1.661 | | P0A7G2 | P0A7G2 | Ribosome-binding factor A |
| *fabB* | 0.336 | 1.664 | | P0A953 | P0A953 | 3-oxoacyl-[acyl-carrier-protein] synthase 1 |
| *pbpG* | -0.270 | 0.912 | | P0AFI5 | P0AFI5 | D-alanyl-D-alanine endopeptidase |
| *sapA* | 0.337 | 1.476 | | Q47622 | Q47622 | Peptide transport periplasmic protein SapA |
| *phnA* | -0.262 | 0.680 | | P0AFJ1 | P0AFJ1 | Protein PhnA |
| *pitA* | -0.768 | 0.211 | | P0AFJ7 | P0AFJ7 | Low-affinity inorganic phosphate transporter 1 |
| *pmbA* | 0.102 | 0.675 | | P0AFK0 | P0AFK0 | Metalloprotease PmbA |
| *pnuC* | 0.221 | 0.305 | | P0AFK2 | P0AFK2 | Nicotinamide riboside transporter PnuC |
| *yajC* | 0.339 | 1.405 | | P0ADZ7 | P0ADZ7 | UPF0092 membrane protein YajC |
| *dppA* | 0.339 | 2.360 | | P23847 | P23847 | Periplasmic dipeptide transport protein |
| *glmM* | 0.341 | 1.446 | | P31120 | P31120 | Phosphoglucosamine mutase |
| *yeiG* | 0.343 | 1.404 | | P33018 | P33018 | S-formylglutathione hydrolase YeiG |
| *pspA* | 0.389 | 1.229 | | P0AFM6 | P0AFM6 | Phage shock protein A |
| *pspB* | -0.612 | 0.396 | | P0AFM9 | P0AFM9 | Phage shock protein B |
| *ybbO* | -0.295 | 0.731 | | P0AFP4 | P0AFP4 | Uncharacterized oxidoreductase YbbO |
| *xseB* | 0.345 | 2.231 | | P0A8G9 | P0A8G9 | Exodeoxyribonuclease 7 small subunit |
| *ycfH* | 0.125 | 0.602 | | P0AFQ7 | P0AFQ7 | Uncharacterized deoxyribonuclease YcfH |
| *dauA* | 0.033 | 0.019 | | P0AFR2 | P0AFR2 | C4-dicarboxylic acid transporter DauA |
| *mukF* | 0.346 | 1.365 | | P60293 | P60293 | Chromosome partition protein MukF |
| *folM* | -0.609 | 0.903 | | P0AFS3 | P0AFS3 | Dihydrofolate reductase FolM |
| *udp* | 0.347 | 2.485 | | P12758 | P12758 | Uridine phosphorylase |
| *ribC* | -0.045 | 0.105 | | P0AFU8 | P0AFU8 | Riboflavin synthase |
| *ygjR* | 0.348 | 1.519 | | P42599 | P42599 | Uncharacterized oxidoreductase YgjR |
| *yajQ* | 0.349 | 1.930 | | P0A8E7 | P0A8E7 | UPF0234 protein YajQ |
| *hpf* | -0.252 | 0.139 | | P0AFX0 | P0AFX0 | Ribosome hibernation promoting factor |
| *yadG* | 0.351 | 1.980 | | P36879 | P36879 | Uncharacterized ABC transporter ATP-binding protein YadG |
| *alaS* | 0.355 | 2.386 | | P00957 | P00957 | Alanine--tRNA ligase |
| *murD* | 0.356 | 1.574 | | P14900 | P14900 | UDP-N-acetylmuramoylalanine--D-glutamate ligase |
| *sanA* | 0.439 | 0.968 | | P0AFY2 | P0AFY2 | Protein SanA |
| *seqA* | -0.086 | 0.343 | | P0AFY8 | P0AFY8 | Negative modulator of initiation of replication |
| *pqiB* | 0.367 | 2.715 | | P43671 | P43671 | Paraquat-inducible protein B |
| *phoU* | 0.367 | 2.463 | | P0A9K7 | P0A9K7 | Phosphate-specific transport system accessory protein PhoU |
| *wzzE* | -0.236 | 1.161 | | P0AG00 | P0AG00 | Lipopolysaccharide biosynthesis protein WzzE |
| *mrdA* | 0.370 | 1.320 | | P0AD65 | P0AD65 | Penicillin-binding protein 2 |
| *helD* | 0.372 | 1.887 | | P15038 | P15038 | Helicase IV |
| *cmoA* | 0.385 | 2.280 | | P76290 | P76290 | tRNA (cmo5U34)-methyltransferase |
| *purK* | 0.386 | 3.403 | | P09029 | P09029 | N5-carboxyaminoimidazole ribonucleotide synthase |
| *relA* | -0.296 | 1.156 | | P0AG20 | P0AG20 | GTP pyrophosphokinase |
| *spoT* | -0.038 | 0.067 | | P0AG24 | P0AG24 | Bifunctional (p)ppGpp synthase/hydrolase SpoT |
| *ybgI* | 0.386 | 2.085 | | P0AFP6 | P0AFP6 | Putative GTP cyclohydrolase 1 type 2 |
| *amiC* | 0.386 | 1.530 | | P63883 | P63883 | N-acetylmuramoyl-L-alanine amidase AmiC |
| *ribF* | -0.165 | 0.399 | | P0AG40 | P0AG40 | Riboflavin biosynthesis protein RibF |
| *azoR* | 0.390 | 2.135 | | P41407 | P41407 | FMN-dependent NADH-azoreductase |
| *rplU* | 0.000 | 0.000 | | P0AG48 | P0AG48 | 50S ribosomal protein L21 |
| *rpmD* | -0.105 | 0.413 | | P0AG51 | P0AG51 | 50S ribosomal protein L30 |
| *dcd* | 0.391 | 5.092 | | P28248 | P28248 | Deoxycytidine triphosphate deaminase |

| Gene names | og2(fold-Change | -Log(p-value) | Significant | Protein IDs | Majority protein IDs | Protein names |
|---|---|---|---|---|---|---|
| *rpsN* | -0.108 | 0.405 | | P0AG59 | P0AG59 | 30S ribosomal protein S14 |
| *rpsQ* | -0.067 | 0.424 | | P0AG63 | P0AG63 | 30S ribosomal protein S17 |
| *rsmB* | 0.391 | 1.576 | | P36929 | P36929 | Ribosomal RNA small subunit methyltransferase B |
| *wecC* | 0.391 | 1.713 | | P27829 | P27829 | UDP-N-acetyl-D-mannosamine dehydrogenase |
| *kdsB* | 0.392 | 3.343 | | P04951 | P04951 | 3-deoxy-manno-octulosonate cytidylyltransferase |
| *hflD* | 0.393 | 2.002 | | P25746 | P25746 | High frequency lysogenization protein HflD |
| *secD* | 0.287 | 1.035 | | P0AG90 | P0AG90 | Protein translocase subunit SecD |
| *secF* | 0.275 | 0.983 | | P0AG93 | P0AG93 | Protein translocase subunit SecF |
| *secE* | -0.126 | 0.314 | | P0AG96 | P0AG96 | Protein translocase subunit SecE |
| *secG* | 0.133 | 0.814 | | P0AG99 | P0AG99 | Protein-export membrane protein SecG |
| *secY* | -0.097 | 0.239 | | P0AGA2 | P0AGA2 | Protein translocase subunit SecY |
| *sdhC* | 0.394 | 1.395 | | P69054 | P69054 | Succinate dehydrogenase cytochrome b556 subunit |
| *cmoB* | 0.396 | 1.543 | | P76291 | P76291 | tRNA (mo5U34)-methyltransferase |
| *aroH* | 0.399 | 3.751 | | P00887 | P00887 | Phospho-2-dehydro-3-deoxyheptonate aldolase, Trp-sensitive |
| *secB* | 0.399 | 1.799 | | P0AG86 | P0AG86 | Protein-export protein SecB |
| *slmA* | 0.400 | 1.925 | | P0C093 | P0C093 | Nucleoid occlusion factor SlmA |
| *ssb* | -0.588 | 0.769 | | P0AGE0;P18310 | P0AGE0 | Single-stranded DNA-binding protein |
| *murG* | 0.402 | 1.929 | | P17443 | P17443 | N-acetylglucosaminyl transferase |
| *chrR* | 0.015 | 0.044 | | P0AGE6 | P0AGE6 | Chromate reductase |
| *rluC* | 0.405 | 1.724 | | P0AA39 | P0AA39 | Ribosomal large subunit pseudouridine synthase C |
| *asd* | 0.408 | 2.038 | | P0A9Q9 | P0A9Q9 | Aspartate-semialdehyde dehydrogenase |
| *rnc* | 0.410 | 2.268 | | P0A7Y0 | P0A7Y0 | Ribonuclease 3 |
| *bioH* | 0.411 | 2.095 | | P13001 | P13001 | Pimeloyl-[acyl-carrier protein] methyl ester esterase |
| *dsbC* | 0.412 | 1.841 | | P0AEG6 | P0AEG6 | Thiol:disulfide interchange protein DsbC |
| *yfbT* | 0.412 | 2.016 | | P77625 | P77625 | Sugar phosphatase YfbT |
| *trmH* | 0.149 | 0.625 | | P0AGJ2 | P0AGJ2 | tRNA (guanosine(18)-2-O)-methyltransferase |
| *deoD* | 0.414 | 2.047 | | P0ABP8 | P0ABP8 | Purine nucleoside phosphorylase DeoD-type |
| *trmL* | 0.151 | 0.490 | | P0AGJ7 | P0AGJ7 | tRNA (cytidine(34)-2-O)-methyltransferase |
| *atpD* | 0.417 | 2.362 | | P0ABB4 | P0ABB4 | ATP synthase subunit beta |
| *iscS* | 0.417 | 3.183 | | P0A6B7 | P0A6B7 | Cysteine desulfurase IscS |
| *rluB* | 0.421 | 2.585 | | P37765 | P37765 | Ribosomal large subunit pseudouridine synthase B |
| *yejK* | 0.421 | 2.157 | | P33920 | P33920 | Nucleoid-associated protein YejK |
| *ratA* | 0.154 | 0.629 | | P0AGL5 | P0AGL5 | Ribosome association toxin RatA |
| *psd* | 0.423 | 2.578 | | P0A8K1 | P0A8K1 | Phosphatidylserine decarboxylase proenzyme |
| *rplR* | -0.006 | 0.017 | | P0C018 | P0C018 | 50S ribosomal protein L18 |
| *yaiE* | -0.092 | 0.189 | | P0C037 | P0C037 | UPF0345 protein YaiE |
| *diaA* | 0.424 | 1.879 | | P66817 | P66817 | DnaA initiator-associating protein DiaA |
| *hldE* | 0.424 | 2.604 | | P76658 | P76658 | Bifunctional protein HldE |
| *ispA* | 0.424 | 3.652 | | P22939 | P22939 | Farnesyl diphosphate synthase |
| *ydgH* | 0.426 | 1.960 | | P76177 | P76177 | Protein YdgH |
| *yghU* | 0.434 | 2.778 | | Q46845 | Q46845 | Disulfide-bond oxidoreductase YghU |
| *aceE* | 0.435 | 2.018 | | P0AFG8 | P0AFG8 | Pyruvate dehydrogenase E1 component |
| *rlmB* | 0.436 | 2.956 | | P63177 | P63177 | 23S rRNA (guanosine-2-O-)-methyltransferase RlmB |
| *mukE* | 0.436 | 2.629 | | P22524 | P22524 | Chromosome partition protein MukE |
| *degP* | -0.027 | 0.118 | | P0C0V0 | P0C0V0 | Periplasmic serine endoprotease DegP |
| *gatY* | 0.668 | 0.217 | | P0C8J6 | P0C8J6 | D-tagatose-1,6-bisphosphate aldolase subunit GatY |
| *galM* | 0.436 | 1.324 | | P0A9C3 | P0A9C3 | Aldose 1-epimerase |
| *fabF* | 0.437 | 2.034 | | P0AAI5 | P0AAI5 | 3-oxoacyl-[acyl-carrier-protein] synthase 2 |
| *tufB;tufA* | 0.176 | 1.185 | | P0CE48;P0CE47 | P0CE48;P0CE47 | Elongation factor Tu |
| *rcsC* | 0.259 | 0.475 | | P0DMC5 | P0DMC5 | Sensor histidine kinase RcsC |
| *rcsB* | -0.047 | 0.175 | | P0DMC7 | P0DMC7 | Transcriptional regulatory protein RcsB |
| *rlpA* | 0.370 | 1.186 | | P10100 | P10100 | Rare lipoprotein A |
| *ftsY* | -0.190 | 0.892 | | P10121 | P10121 | Signal recognition particle receptor FtsY |

| Gene names | og2(fold-Change | -Log(p-value) | Significant | Protein IDs | Majority protein IDs | Protein names |
|---|---|---|---|---|---|---|
| *tpiA* | 0.437 | 1.991 | | P0A858 | P0A858 | Triosephosphate isomerase |
| *hisA* | -0.163 | 0.266 | | P10371 | P10371 | 1-(5-phosphoribosyl)-5-[(5-phosphoribosylamino)methylideneamino] imidazole-4-carboxamide isomerase |
| *secA* | -0.057 | 0.172 | | P10408 | P10408 | Protein translocase subunit SecA |
| *lpxB* | 0.178 | 0.584 | | P10441 | P10441 | Lipid-A-disaccharide synthase |
| *dnaE* | 1.088 | 0.613 | | P10443 | P10443 | DNA polymerase III subunit alpha |
| *dppF* | 0.438 | 1.540 | | P37313 | P37313 | Dipeptide transport ATP-binding protein DppF |
| *ugpQ* | -0.199 | 0.202 | | P10908 | P10908 | Glycerophosphoryl diester phosphodiesterase |
| *argH* | -0.302 | 0.084 | | P11447 | P11447 | Argininosuccinate lyase |
| *lptC* | 0.439 | 1.446 | | P0ADV9 | P0ADV9 | Lipopolysaccharide export system protein LptC |
| *damX* | 0.199 | 1.166 | | P11557 | P11557 | Cell division protein DamX |
| *trxB* | 0.440 | 2.647 | | P0A9P4 | P0A9P4 | Thioredoxin reductase |
| *murF* | 0.125 | 0.986 | | P11880 | P11880 | UDP-N-acetylmuramoyl-tripeptide--D-alanyl-D-alanine ligase |
| *parC* | 0.442 | 2.351 | | P0AFI2 | P0AFI2 | DNA topoisomerase 4 subunit A |
| *hslU* | 0.442 | 2.187 | | P0A6H5 | P0A6H5 | ATP-dependent protease ATPase subunit HslU |
| *ymjA* | 0.448 | 2.126 | | P0ACV8 | P0ACV8 | Uncharacterized protein YmjA |
| *cydC* | 0.452 | 1.302 | | P23886 | P23886 | ATP-binding/permease protein CydC |
| *adhE* | 0.454 | 2.129 | | P0A9Q7 | P0A9Q7 | Aldehyde-alcohol dehydrogenase |
| *ybhB* | 0.243 | 0.916 | | P12994 | P12994 | UPF0098 protein YbhB |
| *yhfA* | 0.455 | 1.785 | | P0ADX1 | P0ADX1 | Protein YhfA |
| *bioB* | 0.866 | 0.433 | | P12996 | P12996 | Biotin synthase |
| *bioD1* | 0.198 | 1.106 | | P13000 | P13000 | ATP-dependent dethiobiotin synthetase BioD 1 |
| *ilvD* | 0.456 | 1.704 | | P05791 | P05791 | Dihydroxy-acid dehydratase |
| *metH* | 0.263 | 1.040 | | P13009 | P13009 | Methionine synthase |
| *yiiD* | 0.457 | 1.319 | | P0ADQ2 | P0ADQ2 | Uncharacterized protein YiiD |
| *proQ* | 0.458 | 2.673 | | P45577 | P45577 | RNA chaperone ProQ |
| *sucD* | 0.459 | 2.894 | | P0AGE9 | P0AGE9 | Succinyl-CoA ligase [ADP-forming] subunit alpha |
| *folX* | 0.462 | 1.479 | | P0AC19 | P0AC19 | D-erythro-7,8-dihydroneopterin triphosphate epimerase |
| *yfcN* | 0.463 | 1.349 | | P0A8B2 | P0A8B2 | UPF0115 protein YfcN |
| *selB* | -0.081 | 0.172 | | P14081 | P14081 | Selenocysteine-specific elongation factor |
| *frlR* | 0.466 | 2.302 | | P45544 | P45544 | HTH-type transcriptional regulator FrlR |
| *glnS* | 0.467 | 2.513 | | P00962 | P00962 | Glutamine--tRNA ligase |
| *folD* | 0.468 | 2.524 | | P24186 | P24186 | Bifunctional protein FolD |
| *ligA* | -0.042 | 0.141 | | P15042 | P15042 | DNA ligase |
| *ppiC* | 0.470 | 1.440 | | P0A9L5 | P0A9L5 | Peptidyl-prolyl cis-trans isomerase C |
| *srlR* | -0.176 | 0.516 | | P15082 | P15082 | Glucitol operon repressor |
| *purL* | 0.183 | 0.619 | | P15254 | P15254 | Phosphoribosylformylglycinamidine synthase |
| *slyD* | 0.471 | 1.630 | | P0A9K9 | P0A9K9 | FKBP-type peptidyl-prolyl cis-trans isomerase SlyD |
| *prlF* | -0.218 | 0.406 | | P15373 | P15373 | Antitoxin PrlF |
| *asnS* | 0.476 | 2.186 | | P0A8M0 | P0A8M0 | Asparagine--tRNA ligase |
| *truB* | 0.480 | 1.877 | | P60340 | P60340 | tRNA pseudouridine synthase B |
| *aroE* | 0.088 | 0.534 | | P15770 | P15770 | Shikimate dehydrogenase (NADP(+)) |
| *malQ* | 0.453 | 0.433 | | P15977 | P15977 | 4-alpha-glucanotransferase |
| *sdaA* | 0.457 | 0.928 | | P16095 | P16095 | L-serine dehydratase 1 |
| *miaA* | -0.304 | 0.876 | | P16384 | P16384 | tRNA dimethylallyltransferase |
| *yheO* | 0.481 | 1.626 | | P64624 | P64624 | Uncharacterized protein YheO |
| *ubiE* | 0.482 | 2.979 | | P0A887 | P0A887 | Ubiquinone/menaquinone biosynthesis C-methyltransferase UbiE |
| *ompR* | 0.489 | 3.892 | | P0AA16 | P0AA16 | Transcriptional regulatory protein OmpR |
| *fldB* | 0.489 | 1.407 | | P0ABY4 | P0ABY4 | Flavodoxin-2 |
| *dtd* | 0.491 | 2.646 | | P0A6M4 | P0A6M4 | D-aminoacyl-tRNA deacylase |
| *nlpD* | 0.492 | 1.633 | | P0ADA3 | P0ADA3 | Murein hydrolase activator NlpD |
| *glmS* | -0.227 | 0.609 | | P17169 | P17169 | Glutamine--fructose-6-phosphate aminotransferase [isomerizing] |
| *tatA* | 0.493 | 1.814 | | P69428 | P69428 | Sec-independent protein translocase protein TatA |
| *cysI* | 0.445 | 0.341 | | P17846 | P17846 | Sulfite reductase [NADPH] hemoprotein beta-component |

| Gene names | og2(fold-Change | -Log(p-value) | Significant | Protein IDs | Majority protein IDs | Protein names |
|---|---|---|---|---|---|---|
| *murC* | -0.087 | 0.527 | | P17952 | P17952 | UDP-N-acetylmuramate--L-alanine ligase |
| *ispD* | 0.493 | 2.977 | | Q46893 | Q46893 | 2-C-methyl-D-erythritol 4-phosphate cytidylyltransferase |
| *purH* | 0.494 | 1.422 | | P15639 | P15639 | Bifunctional purine biosynthesis protein PurH |
| *minC* | -0.226 | 1.152 | | P18196 | P18196 | Septum site-determining protein MinC |
| *argD* | 0.066 | 0.012 | | P18335 | P18335 | Acetylornithine/succinyldiaminopimelate aminotransferase |
| *yqiC* | 0.494 | 1.503 | | Q46868 | Q46868 | Uncharacterized protein YqiC |
| *aroB* | 0.497 | 3.866 | | P07639 | P07639 | 3-dehydroquinate synthase |
| *ydhQ* | 0.504 | 1.905 | | P77552 | P77552 | Uncharacterized protein YdhQ |
| *tolA* | -0.298 | 1.007 | | P19934 | P19934 | Protein TolA |
| *yfbU* | 0.504 | 3.308 | | P0A8W8 | P0A8W8 | UPF0304 protein YfbU |
| *yjaG* | 0.506 | 3.001 | | P32680 | P32680 | Uncharacterized protein YjaG |
| *clpA* | 0.506 | 2.427 | | P0ABH9 | P0ABH9 | ATP-dependent Clp protease ATP-binding subunit ClpA |
| *panC* | 0.507 | 2.427 | | P31663 | P31663 | Pantothenate synthetase |
| *speA* | -0.415 | 0.114 | | P21170 | P21170 | Biosynthetic arginine decarboxylase |
| *mukB* | 0.507 | 1.780 | | P22523 | P22523 | Chromosome partition protein MukB |
| *nlpA* | 0.512 | 2.502 | | P04846 | P04846 | Lipoprotein 28 |
| *cpxA* | 0.513 | 2.000 | | P0AE82 | P0AE82 | Sensor protein CpxA |
| *rlmE* | 0.513 | 3.347 | | P0C0R7 | P0C0R7 | Ribosomal RNA large subunit methyltransferase E |
| *yhcB* | 0.515 | 3.373 | | P0ADW3 | P0ADW3 | Inner membrane protein YhcB |
| *rnr* | 0.316 | 0.779 | | P21499 | P21499 | Ribonuclease R |
| *srmB* | -0.132 | 0.524 | | P21507 | P21507 | ATP-dependent RNA helicase SrmB |
| *aceF* | 0.516 | 2.287 | | P06959 | P06959 | Pyruvate dehydrogenase, E2 subunit |
| *glpK* | 0.521 | 2.154 | | P0A6F3 | P0A6F3 | Glycerol kinase |
| *lpxD* | -0.013 | 0.041 | | P21645 | P21645 | UDP-3-O-(3-hydroxymyristoyl)glucosamine N-acyltransferase |
| *sucC* | 0.522 | 4.745 | | P0A836 | P0A836 | Succinyl-CoA ligase [ADP-forming] subunit beta |
| *yaiL* | 0.523 | 2.084 | | P51024 | P51024 | Uncharacterized protein YaiL |
| *yhhY* | 0.524 | 2.986 | | P46854 | P46854 | Uncharacterized N-acetyltransferase YhhY |
| *hscA* | 0.525 | 2.268 | | P0A6Z1 | P0A6Z1 | Chaperone protein HscA |
| *yibN* | 0.526 | 2.530 | | P0AG27 | P0AG27 | Uncharacterized protein YibN |
| *murE* | 0.214 | 1.201 | | P22188 | P22188 | UDP-N-acetylmuramoyl-L-alanyl-D-glutamate--2,6-diaminopimelate ligase |
| *cysQ* | 0.055 | 0.363 | | P22255 | P22255 | 3(2),5-bisphosphate nucleotidase CysQ |
| *surE* | 0.526 | 1.932 | | P0A840 | P0A840 | 5/3-nucleotidase SurE |
| *ptsN* | 0.532 | 2.144 | | P69829 | P69829 | Nitrogen regulatory protein |
| *yggS* | 0.532 | 2.457 | | P67080 | P67080 | UPF0001 protein YggS |
| *bamA* | 0.532 | 1.873 | | P0A940 | P0A940 | Outer membrane protein assembly factor BamA |
| *folP* | 0.542 | 3.437 | | P0AC13 | P0AC13 | Dihydropteroate synthase |
| *livF* | 0.762 | 0.321 | | P22731 | P22731 | High-affinity branched-chain amino acid transport ATP-binding protein LivF |
| *rimM* | 0.543 | 2.299 | | P0A7X6 | P0A7X6 | Ribosome maturation factor RimM |
| *katG* | 0.546 | 2.291 | | P13029 | P13029 | Catalase-peroxidase |
| *nfuA* | 0.547 | 2.072 | | P63020 | P63020 | Fe/S biogenesis protein NfuA |
| *ilvA* | 0.550 | 1.722 | | P04968 | P04968 | L-threonine dehydratase biosynthetic IlvA |
| *mutL* | -0.336 | 0.786 | | P23367 | P23367 | DNA mismatch repair protein MutL |
| *dacA* | 0.552 | 4.525 | | P0AEB2 | P0AEB2 | D-alanyl-D-alanine carboxypeptidase DacA |
| *yeeN* | 0.552 | 2.347 | | P0A8A2 | P0A8A2 | Probable transcriptional regulatory protein YeeN |
| *leuB* | 0.555 | 1.679 | | P30125 | P30125 | 3-isopropylmalate dehydrogenase |
| *atpB* | 0.557 | 1.380 | | P0AB98 | P0AB98 | ATP synthase subunit a |
| *mscM* | 0.558 | 3.086 | | P39285 | P39285 | Miniconductance mechanosensitive channel MscM |
| *ybcJ* | 0.558 | 1.747 | | P0AAS7 | P0AAS7 | Uncharacterized protein YbcJ |
| *pdhR* | 0.559 | 3.201 | | P0ACL9 | P0ACL9 | Pyruvate dehydrogenase complex repressor |
| *rlmL* | 0.570 | 1.870 | | P75864 | P75864 | Ribosomal RNA large subunit methyltransferase K/L |
| *ybgJ* | 0.572 | 2.906 | | P0AAV4 | P0AAV4 | Uncharacterized protein YbgJ |
| *ppiB* | -0.234 | 0.777 | | P23869 | P23869 | Peptidyl-prolyl cis-trans isomerase B |
| *hemH* | -0.601 | 0.823 | | P23871 | P23871 | Ferrochelatase |

| Gene names | og2(fold-Change | -Log(p-value) | Significant | Protein IDs | Majority protein IDs | Protein names |
|---|---|---|---|---|---|---|
| *fmt* | -0.179 | 0.538 | | P23882 | P23882 | Methionyl-tRNA formyltransferase |
| *ppa* | 0.574 | 1.942 | | P0A7A9 | P0A7A9 | Inorganic pyrophosphatase |
| *pyrF* | 0.576 | 3.157 | | P08244 | P08244 | Orotidine 5-phosphate decarboxylase |
| *htpX* | -0.165 | 0.676 | | P23894 | P23894 | Protease HtpX |
| *argE* | -0.556 | 0.710 | | P23908 | P23908 | Acetylornithine deacetylase |
| *yihI* | 0.578 | 2.180 | | P0A8H6 | P0A8H6 | Der GTPase-activating protein YihI |
| *lnt* | 0.044 | 0.098 | | P23930 | P23930 | Apolipoprotein N-acyltransferase |
| *kch* | 0.580 | 2.055 | | P31069 | P31069 | Voltage-gated potassium channel Kch |
| *yffB* | 0.819 | 0.952 | | P24178 | P24178 | Protein YffB |
| *accC* | 0.023 | 0.092 | | P24182 | P24182 | Biotin carboxylase |
| *cytR* | 0.580 | 1.792 | | P0ACN7 | P0ACN7 | HTH-type transcriptional repressor CytR |
| *rna* | 0.585 | 3.718 | | P21338 | P21338 | Ribonuclease I |
| *yjiA* | 0.045 | 0.154 | | P24203 | P24203 | Uncharacterized GTP-binding protein YjiA |
| *fabH* | 0.586 | 2.159 | | P0A6R0 | P0A6R0 | 3-oxoacyl-[acyl-carrier-protein] synthase 3 |
| *trmA* | 0.588 | 3.148 | | P23003;REV__P0ABC3 | P23003 | tRNA/tmRNA (uracil-C(5))-methyltransferase |
| *thiM* | 0.589 | 1.939 | | P76423 | P76423 | Hydroxyethylthiazole kinase |
| *rluD* | 0.590 | 1.713 | | P33643 | P33643 | Ribosomal large subunit pseudouridine synthase D |
| *rpoN* | 0.096 | 0.434 | | P24255 | P24255 | RNA polymerase sigma-54 factor |
| *nuoJ* | 0.590 | 1.925 | | P0AFE0 | P0AFE0 | NADH-quinone oxidoreductase subunit J |
| *engB* | 0.595 | 2.877 | | P0A6P7 | P0A6P7 | Probable GTP-binding protein EngB |
| *cspC* | 0.595 | 1.847 | | P0A9Y6 | P0A9Y6 | Cold shock-like protein CspC |
| *hscB* | 0.598 | 1.386 | | P0A6L9 | P0A6L9 | Co-chaperone protein HscB |
| *exuR* | 0.599 | 2.794 | | P0ACL2 | P0ACL2 | Exu regulon transcriptional regulator |
| *mnmE* | 0.165 | 0.364 | | P25522 | P25522 | tRNA modification GTPase MnmE |
| *codA* | 0.279 | 0.339 | | P25524 | P25524 | Cytosine deaminase |
| *deoB* | 0.600 | 1.892 | | P0A6K6 | P0A6K6 | Phosphopentomutase |
| *ubiI* | 0.033 | 0.015 | | P25535 | P25535 | 2-octaprenylphenol hydroxylase |
| *yhdE* | 0.053 | 0.376 | | P25536 | P25536 | Maf-like protein YhdE |
| *ribD* | -0.265 | 0.859 | | P25539 | P25539 | Riboflavin biosynthesis protein RibD |
| *anmK* | 0.605 | 2.737 | | P77570 | P77570 | Anhydro-N-acetylmuramic acid kinase |
| *aldA* | -0.811 | 0.603 | | P25553 | P25553 | Lactaldehyde dehydrogenase |
| *lptG* | 0.608 | 1.323 | | P0ADC6 | P0ADC6 | Lipopolysaccharide export system permease protein LptG |
| *ispF* | 0.610 | 1.617 | | P62617 | P62617 | 2-C-methyl-D-erythritol 2,4-cyclodiphosphate synthase |
| *gcvR* | 0.611 | 2.632 | | P0A9I3 | P0A9I3 | Glycine cleavage system transcriptional repressor |
| *glgX* | 0.615 | 1.417 | | P15067 | P15067 | Glycogen debranching enzyme |
| *corC* | 0.617 | 4.481 | | P0AE78 | P0AE78 | Magnesium and cobalt efflux protein CorC |
| *loiP* | -0.016 | 0.034 | | P25894 | P25894 | Metalloprotease LoiP |
| *ycbL* | 0.617 | 1.317 | | P75849 | P75849 | Uncharacterized protein YcbL |
| *prfC* | 0.617 | 2.662 | | P0A7I4 | P0A7I4 | Peptide chain release factor 3 |
| *rluF* | 0.624 | 2.460 | | P32684 | P32684 | Ribosomal large subunit pseudouridine synthase F |
| *nfo* | 0.626 | 2.671 | | P0A6C1 | P0A6C1 | Endonuclease 4 |
| *pgi* | 0.627 | 2.000 | | P0A6T1 | P0A6T1 | Glucose-6-phosphate isomerase |
| *obgE* | 0.629 | 3.383 | | P42641 | P42641 | GTPase ObgE/CgtA |
| *ftsP* | 0.341 | 1.234 | | P26648 | P26648 | Cell division protein FtsP |
| *ilvH* | 0.630 | 2.863 | | P00894 | P00894 | Acetolactate synthase isozyme 3 small subunit |
| *cydD* | 0.632 | 2.751 | | P29018 | P29018 | ATP-binding/permease protein CydD |
| *moaB* | 0.635 | 2.960 | | P0AEZ9 | P0AEZ9 | Molybdenum cofactor biosynthesis protein B |
| *nadR* | 0.231 | 1.200 | | P27278 | P27278 | Trifunctional NAD biosynthesis/regulator protein NadR |
| *gloA* | 0.638 | 1.622 | | P0AC81 | P0AC81 | Lactoylglutathione lyase |
| *tsaC* | 0.642 | 1.881 | | P45748 | P45748 | Threonylcarbamoyl-AMP synthase |
| *dsdA* | 0.645 | 2.305 | | P00926 | P00926 | D-serine dehydratase |
| *roxA* | 0.126 | 1.143 | | P27431 | P27431 | 50S ribosomal protein L16 arginine hydroxylase |
| *rodZ* | -0.019 | 0.124 | | P27434 | P27434 | Cytoskeleton protein RodZ |

| Gene names | og2(fold-Change | -Log(p-value) | Significant | Protein IDs | Majority protein IDs | Protein names |
|---|---|---|---|---|---|---|
| *wecB* | 0.095 | 0.171 | | P27828 | P27828 | UDP-N-acetylglucosamine 2-epimerase |
| *yniC* | 0.648 | 3.017 | | P77247 | P77247 | 2-deoxyglucose-6-phosphate phosphatase |
| *rffG* | 0.192 | 1.206 | | P27830 | P27830 | dTDP-glucose 4,6-dehydratase 2 |
| *rffA* | 0.222 | 0.775 | | P27833 | P27833 | dTDP-4-amino-4,6-dideoxygalactose transaminase |
| *wecG* | 0.005 | 0.028 | | P27836 | P27836 | UDP-N-acetyl-D-mannosaminuronic acid transferase |
| *rng* | 0.651 | 1.979 | | P0A9J0 | P0A9J0 | Ribonuclease G |
| *rbsB* | 0.653 | 1.642 | | P02925 | P02925 | D-ribose-binding periplasmic protein |
| *asmA* | 0.335 | 1.234 | | P28249 | P28249 | Protein AsmA |
| *pepD* | 0.653 | 3.062 | | P15288 | P15288 | Cytosol non-specific dipeptidase |
| *pabC* | -0.131 | 1.131 | | P28305 | P28305 | Aminodeoxychorismate lyase |
| *yceG* | 0.140 | 0.827 | | P28306 | P28306 | UPF0755 protein YceG |
| *holD* | -0.141 | 0.373 | | P28632 | P28632 | DNA polymerase III subunit psi |
| *pta* | 0.656 | 1.513 | | P0A9M8 | P0A9M8 | Phosphate acetyltransferase |
| *fpr* | 0.122 | 0.361 | | P28861 | P28861 | Ferredoxin--NADP reductase |
| *nusG* | 0.656 | 3.873 | | P0AFG0 | P0AFG0 | Transcription termination/antitermination protein NusG |
| *dadX* | 0.233 | 0.465 | | P29012 | P29012 | Alanine racemase, catabolic |
| *map* | 0.658 | 3.807 | | P0AE18 | P0AE18 | Methionine aminopeptidase |
| *ftsN* | 0.233 | 0.521 | | P29131 | P29131 | Cell division protein FtsN |
| *ung* | 0.660 | 4.158 | | P12295 | P12295 | Uracil-DNA glycosylase |
| *hemE* | -0.072 | 0.375 | | P29680 | P29680 | Uroporphyrinogen decarboxylase |
| *atpF* | 0.673 | 3.221 | | P0ABA0 | P0ABA0 | ATP synthase subunit b |
| *kbl* | 0.673 | 2.095 | | P0AB77 | P0AB77 | 2-amino-3-ketobutyrate coenzyme A ligase |
| *cobB* | 0.674 | 3.957 | | P75960 | P75960 | NAD-dependent protein deacylase |
| *aroG* | 0.674 | 2.862 | | P0AB91 | P0AB91 | Phospho-2-dehydro-3-deoxyheptonate aldolase, Phe-sensitive |
| *groS* | 0.675 | 2.593 | | P0A6F9 | P0A6F9 | 10 kDa chaperonin |
| *queD* | 0.676 | 1.905 | | P65870 | P65870 | 6-carboxy-5,6,7,8-tetrahydropterin synthase |
| *purE* | 0.682 | 2.454 | | P0AG18 | P0AG18 | N5-carboxyaminoimidazole ribonucleotide mutase |
| *atpH* | 0.684 | 3.240 | | P0ABA4 | P0ABA4 | ATP synthase subunit delta |
| *zapE* | 0.684 | 1.938 | | P64612 | P64612 | Cell division protein ZapE |
| *yigB* | 0.688 | 2.492 | | P0ADP0 | P0ADP0 | Flavin mononucleotide phosphatase YigB |
| *yoaF* | 0.688 | 1.448 | | P64493 | P64493 | Uncharacterized protein YoaF |
| *gshB* | 0.689 | 3.428 | | P04425 | P04425 | Glutathione synthetase |
| *moaD* | 0.187 | 0.536 | | P30748 | P30748 | Molybdopterin synthase sulfur carrier subunit |
| *galT* | 0.690 | 1.881 | | P09148 | P09148 | Galactose-1-phosphate uridylyltransferase |
| *mazG* | 0.691 | 2.428 | | P0AEY3 | P0AEY3 | Nucleoside triphosphate pyrophosphohydrolase |
| *rnb* | 0.246 | 1.076 | | P30850 | P30850 | Exoribonuclease 2 |
| *artI* | 0.090 | 0.033 | | P30859 | P30859 | Putative ABC transporter arginine-binding protein 2 |
| *artJ* | 0.395 | 0.062 | | P30860 | P30860 | ABC transporter arginine-binding protein 1 |
| *dkgB* | -0.052 | 0.109 | | P30863 | P30863 | 2,5-diketo-D-gluconic acid reductase B |
| *ybgL* | 0.692 | 1.738 | | P75746 | P75746 | UPF0271 protein YbgL |
| *glnE* | 0.501 | 0.997 | | P30870 | P30870 | Glutamate-ammonia-ligase adenylyltransferase |
| *ygiF* | 0.106 | 0.709 | | P30871 | P30871 | Inorganic triphosphatase |
| *mfd* | -0.206 | 0.563 | | P30958 | P30958 | Transcription-repair-coupling factor |
| *ackA* | 0.694 | 2.281 | | P0A6A3 | P0A6A3 | Acetate kinase |
| *gcvT* | 0.701 | 1.851 | | P27248 | P27248 | Aminomethyltransferase |
| *yedD* | -0.338 | 0.802 | | P31063 | P31063 | Uncharacterized lipoprotein YedD |
| *proS* | 0.702 | 3.097 | | P16659 | P16659 | Proline--tRNA ligase |
| *rnpA* | 0.702 | 1.974 | | P0A7Y8 | P0A7Y8 | Ribonuclease P protein component |
| *ydeJ* | -0.136 | 0.032 | | P31131 | P31131 | Protein YdeJ |
| *potF* | -1.095 | 0.505 | | P31133 | P31133 | Putrescine-binding periplasmic protein |
| *sseA* | -0.078 | 0.557 | | P31142 | P31142 | 3-mercaptopyruvate sulfurtransferase |
| *acrB* | -0.205 | 0.626 | | P31224 | P31224 | Multidrug efflux pump subunit AcrB |
| *yidR* | 0.146 | 0.227 | | P31455 | P31455 | Uncharacterized protein YidR |

| Gene names | og2(fold-Change | -Log(p-value) | Significant | Protein IDs | Majority protein IDs | Protein names |
|---|---|---|---|---|---|---|
| *adeP* | 0.465 | 0.809 | | P31466 | P31466 | Adenine permease AdeP |
| *ravA* | -0.044 | 0.088 | | P31473 | P31473 | ATPase RavA |
| *yieP* | -0.596 | 0.449 | | P31475 | P31475 | Uncharacterized HTH-type transcriptional regulator YieP |
| *thiQ* | 0.223 | 0.493 | | P31548 | P31548 | Thiamine import ATP-binding protein ThiQ |
| *thiB* | -0.032 | 0.102 | | P31550 | P31550 | Thiamine-binding periplasmic protein |
| *lptD* | 0.046 | 0.106 | | P31554 | P31554 | LPS-assembly protein LptD |
| *moaC* | 0.704 | 2.942 | | P0A738 | P0A738 | Cyclic pyranopterin monophosphate synthase accessory protein |
| *atpA* | 0.705 | 3.375 | | P0ABB0 | P0ABB0 | ATP synthase subunit alpha |
| *otsA* | -0.991 | 0.700 | | P31677 | P31677 | Alpha,alpha-trehalose-phosphate synthase [UDP-forming] |
| *djlA* | 0.429 | 0.561 | | P31680 | P31680 | DnaJ-like protein DjlA |
| *ilvE* | 0.707 | 3.526 | | P0AB80 | P0AB80 | Branched-chain-amino-acid aminotransferase |
| *dnaN* | 0.707 | 2.679 | | P0A988 | P0A988 | DNA polymerase III subunit beta |
| *wcaI* | -0.555 | 0.599 | | P32057 | P32057 | Putative colanic acid biosynthesis glycosyl transferase WcaI |
| *ibaG* | 0.714 | 2.204 | | P0A9W6 | P0A9W6 | Acid stress protein IbaG |
| *cueO* | 0.719 | 2.366 | | P36649 | P36649 | Blue copper oxidase CueO |
| *typA* | 0.126 | 0.450 | | P32132 | P32132 | GTP-binding protein TypA/BipA |
| *yiiM* | 0.204 | 0.923 | | P32157 | P32157 | Protein YiiM |
| *proB* | 0.722 | 4.323 | | P0A7B5 | P0A7B5 | Glutamate 5-kinase |
| *mobA* | -0.018 | 0.042 | | P32173 | P32173 | Molybdenum cofactor guanylyltransferase |
| *gph* | -0.033 | 0.121 | | P32662 | P32662 | Phosphoglycolate phosphatase |
| *mltA* | 0.726 | 2.463 | | P0A935 | P0A935 | Membrane-bound lytic murein transglycosylase A |
| *gpmI* | 0.729 | 4.108 | | P37689 | P37689 | 2,3-bisphosphoglycerate-independent phosphoglycerate mutase |
| *xseA* | 0.729 | 2.641 | | P04994 | P04994 | Exodeoxyribonuclease 7 large subunit |
| *yjbD* | -0.407 | 1.116 | | P32685 | P32685 | Uncharacterized protein YjbD |
| *thiI* | 0.732 | 2.801 | | P77718 | P77718 | tRNA sulfurtransferase |
| *purF* | 0.738 | 3.163 | | P0AG16 | P0AG16 | Amidophosphoribosyltransferase |
| *mdoG* | 0.044 | 0.376 | | P33136 | P33136 | Glucans biosynthesis protein G |
| *add* | 0.738 | 2.909 | | P22333 | P22333 | Adenosine deaminase |
| *yebE* | 0.114 | 0.138 | | P33218 | P33218 | Inner membrane protein YebE |
| *purT* | -0.263 | 0.926 | | P33221 | P33221 | Phosphoribosylglycinamide formyltransferase 2 |
| *yehS* | 0.344 | 1.199 | | P33355 | P33355 | Uncharacterized protein YehS |
| *glyA* | 0.743 | 2.492 | | P0A825 | P0A825 | Serine hydroxymethyltransferase |
| *bglX* | -0.173 | 0.485 | | P33363 | P33363 | Periplasmic beta-glucosidase |
| *yaiI* | 0.743 | 2.577 | | P0A8D3 | P0A8D3 | UPF0178 protein YaiI |
| *mreB* | 0.746 | 3.263 | | P0A9X4 | P0A9X4 | Rod shape-determining protein MreB |
| *cysE* | 0.748 | 3.457 | | P0A9D4 | P0A9D4 | Serine acetyltransferase |
| *gcvP* | 0.749 | 2.267 | | P33195 | P33195 | Glycine dehydrogenase (decarboxylating) |
| *maeB* | 0.751 | 2.720 | | P76558 | P76558 | NADP-dependent malic enzyme |
| *moaE* | 0.753 | 2.945 | | P30749 | P30749 | Molybdopterin synthase catalytic subunit |
| *ptsO* | 0.754 | 1.323 | | P0A9N0 | P0A9N0 | Phosphocarrier protein NPr |
| *fkpA* | 0.755 | 2.561 | | P45523 | P45523 | FKBP-type peptidyl-prolyl cis-trans isomerase FkpA |
| *ydcF* | 0.264 | 1.114 | | P34209 | P34209 | Protein YdcF |
| *nuoL* | 0.760 | 1.850 | | P33607 | P33607 | NADH-quinone oxidoreductase subunit L |
| *purD* | 0.760 | 2.434 | | P15640 | P15640 | Phosphoribosylamine--glycine ligase |
| *ppiA* | 0.766 | 1.655 | | P0AFL3 | P0AFL3 | Peptidyl-prolyl cis-trans isomerase A |
| *cobT* | 0.235 | 0.341 | | P36562 | P36562 | Nicotinate-nucleotide--dimethylbenzimidazole phosphoribosyltransferase |
| *smtA* | 0.076 | 0.079 | | P36566;REV__P0A858 | P36566 | Protein SmtA |
| *epd* | 0.767 | 2.673 | | P0A9B6 | P0A9B6 | D-erythrose-4-phosphate dehydrogenase |
| *mreC* | 0.776 | 2.182 | | P16926 | P16926 | Cell shape-determining protein MreC |
| *treR* | 0.142 | 0.409 | | P36673 | P36673 | HTH-type transcriptional regulator TreR |
| *zapD* | -0.096 | 0.464 | | P36680 | P36680 | Cell division protein ZapD |
| *gltL* | 0.778 | 2.531 | | P0AAG3 | P0AAG3 | Glutamate/aspartate transport ATP-binding protein GltL |
| *rdgC* | 0.276 | 1.292 | | P36767 | P36767 | Recombination-associated protein RdgC |

| Gene names | og2(fold-Change | -Log(p-value) | Significant | Protein IDs | Majority protein IDs | Protein names |
|---|---|---|---|---|---|---|
| *rdgB* | 0.781 | 2.747 | | P52061 | P52061 | dITP/XTP pyrophosphatase |
| *atpC* | 0.784 | 2.416 | | P0A6E6 | P0A6E6 | ATP synthase epsilon chain |
| *yidC* | 0.790 | 2.293 | | P25714 | P25714 | Membrane protein insertase YidC |
| *yihD* | 0.794 | 1.912 | | P0ADP9 | P0ADP9 | Protein YihD |
| *purU* | 0.132 | 0.832 | | P37051 | P37051 | Formyltetrahydrofolate deformylase |
| *ychJ* | 0.258 | 0.567 | | P37052 | P37052 | UPF0225 protein YchJ |
| *nuoF* | 0.795 | 3.545 | | P31979 | P31979 | NADH-quinone oxidoreductase subunit F |
| *nudK* | -0.402 | 1.096 | | P37128 | P37128 | GDP-mannose pyrophosphatase NudK |
| *ptsP* | -0.115 | 0.359 | | P37177 | P37177 | Phosphoenolpyruvate-protein phosphotransferase PtsP |
| *plsC* | 0.795 | 2.688 | | P26647 | P26647 | 1-acyl-sn-glycerol-3-phosphate acyltransferase |
| *htpG* | 0.795 | 4.553 | | P0A6Z3 | P0A6Z3 | Chaperone protein HtpG |
| *apt* | 0.795 | 2.281 | | P69503 | P69503 | Adenine phosphoribosyltransferase |
| *dxs* | 0.796 | 2.875 | | P77488 | P77488 | 1-deoxy-D-xylulose-5-phosphate synthase |
| *mdtK* | -0.727 | 0.239 | | P37340 | P37340 | Multidrug resistance protein MdtK |
| *pspF* | -0.997 | 0.675 | | P37344 | P37344 | Psp operon transcriptional activator |
| *ahpF* | 0.798 | 3.888 | | P35340 | P35340 | Alkyl hydroperoxide reductase subunit F |
| *mdh* | 0.803 | 2.257 | | P61889 | P61889 | Malate dehydrogenase |
| *yhbY* | 0.809 | 3.008 | | P0AGK4 | P0AGK4 | RNA-binding protein YhbY |
| *gmhB* | 0.811 | 2.693 | | P63228 | P63228 | D-glycero-beta-D-manno-heptose-1,7-bisphosphate 7-phosphatase |
| *fabA* | 0.812 | 1.850 | | P0A6Q3 | P0A6Q3 | 3-hydroxydecanoyl-[acyl-carrier-protein] dehydratase |
| *avtA* | 0.813 | 3.674 | | P09053 | P09053 | Valine--pyruvate aminotransferase |
| *gltI* | 0.816 | 1.891 | | P37902 | P37902 | Glutamate/aspartate periplasmic-binding protein |
| *yhjK* | -0.930 | 1.053 | | P37649 | P37649 | Protein YhjK |
| *bcsG* | 1.172 | 0.454 | | P37659 | P37659 | Protein BcsG homolog |
| *eptC* | 0.816 | 2.394 | | P0CB39 | P0CB39 | Phosphoethanolamine transferase EptC |
| *ghrB* | -0.076 | 0.267 | | P37666 | P37666 | Glyoxylate/hydroxypyruvate reductase B |
| *yiaJ* | 0.045 | 0.185 | | P37671 | P37671 | HTH-type transcriptional regulator YiaJ |
| *iscR* | 0.825 | 1.904 | | P0AGK8 | P0AGK8 | HTH-type transcriptional regulator IscR |
| *envC* | 0.089 | 0.145 | | P37690 | P37690 | Murein hydrolase activator EnvC |
| *uvrY* | 0.832 | 3.187 | | P0AED5 | P0AED5 | Response regulator UvrY |
| *rfbC* | 0.230 | 0.404 | | P37745 | P37745 | dTDP-4-dehydrorhamnose 3,5-epimerase |
| *pdxA* | 0.833 | 2.525 | | P19624 | P19624 | 4-hydroxythreonine-4-phosphate dehydrogenase |
| *trpR* | 0.833 | 2.621 | | P0A881 | P0A881 | Trp operon repressor |
| *wbbJ* | -0.125 | 0.366 | | P37750 | P37750 | Putative lipopolysaccharide biosynthesis O-acetyl transferase WbbJ |
| *wbbK* | -0.051 | 0.187 | | P37751 | P37751 | Putative glycosyltransferase WbbK |
| *ygiM* | 0.835 | 3.953 | | P0ADT8 | P0ADT8 | Uncharacterized protein YgiM |
| *rfbB* | -0.075 | 0.881 | | P37759 | P37759 | dTDP-glucose 4,6-dehydratase 1 |
| *yjeI* | 0.838 | 2.040 | | P0AF70 | P0AF70 | Uncharacterized protein YjeI |
| *fbaA* | 0.839 | 4.654 | | P0AB71 | P0AB71 | Fructose-bisphosphate aldolase class 2 |
| *mpl* | 0.040 | 0.180 | | P37773 | P37773 | UDP-N-acetylmuramate--L-alanyl-gamma-D-glutamyl-meso--2,6-diaminoheptandioate ligase |
| *gloB* | 0.844 | 3.132 | | P0AC84 | P0AC84 | Hydroxyacylglutathione hydrolase |
| *mprA* | 0.851 | 4.475 | | P0ACR9 | P0ACR9 | Transcriptional repressor MprA |
| *parE* | 0.854 | 2.404 | | P20083 | P20083 | DNA topoisomerase 4 subunit B |
| *yfjD* | -0.826 | 1.246 | | P37908 | P37908 | UPF0053 inner membrane protein YfjD |
| *murA* | 0.858 | 2.837 | | P0A749 | P0A749 | UDP-N-acetylglucosamine 1-carboxyvinyltransferase |
| *modA* | 0.860 | 1.594 | | P37329 | P37329 | Molybdate-binding periplasmic protein |
| *pflB* | 0.861 | 2.290 | | P09373 | P09373 | Formate acetyltransferase 1 |
| *yciH* | 0.862 | 2.512 | | P08245 | P08245 | Uncharacterized protein YciH |
| *fnr* | 0.866 | 2.664 | | P0A9E5 | P0A9E5 | Fumarate and nitrate reduction regulatory protein |
| *thiF* | 0.869 | 4.637 | | P30138 | P30138 | Sulfur carrier protein ThiS adenylyltransferase |
| *mgsA* | 0.870 | 2.908 | | P0A731 | P0A731 | Methylglyoxal synthase |

| Gene names | og2(fold-Change | -Log(p-value) | Significant | Protein IDs | Majority protein IDs | Protein names |
|---|---|---|---|---|---|---|
| *yaaA* | 0.874 | 4.864 | | P0A8I3 | P0A8I3 | UPF0246 protein YaaA |
| *radA* | 0.874 | 2.584 | | P24554 | P24554 | DNA repair protein RadA |
| *nuoG* | 0.878 | 2.953 | | P33602 | P33602 | NADH-quinone oxidoreductase subunit G |
| *rsgA* | -0.107 | 0.730 | | P39286 | P39286 | Putative ribosome biogenesis GTPase RsgA |
| *yjfP* | -0.103 | 0.119 | | P39298 | P39298 | Esterase YjfP |
| *ytfB* | 0.085 | 0.350 | | P39310 | P39310 | Uncharacterized protein YtfB |
| *qorB* | -0.168 | 0.907 | | P39315 | P39315 | Quinone oxidoreductase 2 |
| *ppx* | 0.885 | 2.458 | | P0AFL6 | P0AFL6 | Exopolyphosphatase |
| *bdcR* | 0.120 | 0.424 | | P39334 | P39334 | HTH-type transcriptional repressor BdcR |
| *gltA* | 0.891 | 3.201 | | P0ABH7 | P0ABH7 | Citrate synthase |
| *fdx* | 0.893 | 2.124 | | P0A9R4 | P0A9R4 | 2Fe-2S ferredoxin |
| *yjhU* | 0.014 | 0.029 | | P39356 | P39356 | Uncharacterized transcriptional regulator YjhU |
| *nuoI* | 0.893 | 3.189 | | P0AFD6 | P0AFD6 | NADH-quinone oxidoreductase subunit I |
| *yicC* | 0.895 | 3.007 | | P23839 | P23839 | UPF0701 protein YicC |
| *folC* | 0.897 | 2.512 | | P08192 | P08192 | Bifunctional protein FolC |
| *yjjV* | -0.643 | 1.125 | | P39408 | P39408 | Uncharacterized deoxyribonuclease YjjV |
| *adhP* | -1.311 | 0.751 | | P39451 | P39451 | Alcohol dehydrogenase, propanol-preferring |
| *nrdE* | -0.202 | 0.079 | | P39452 | P39452 | Ribonucleoside-diphosphate reductase 2 subunit alpha |
| *dsbA* | 0.903 | 2.176 | | P0AEG4 | P0AEG4 | Thiol:disulfide interchange protein DsbA |
| *oppD* | 0.906 | 2.503 | | P76027 | P76027 | Oligopeptide transport ATP-binding protein OppD |
| *tatB* | 0.910 | 4.057 | | P69425 | P69425 | Sec-independent protein translocase protein TatB |
| *mdoD* | 0.142 | 0.635 | | P40120 | P40120 | Glucans biosynthesis protein D |
| *thrS* | 0.916 | 3.498 | | P0A8M3 | P0A8M3 | Threonine--tRNA ligase |
| *nlpE* | 0.203 | 0.311 | | P40710 | P40710 | Lipoprotein NlpE |
| *agp* | 0.920 | 1.569 | | P19926 | P19926 | Glucose-1-phosphatase |
| *solA* | 0.140 | 1.028 | | P40874 | P40874 | N-methyl-L-tryptophan oxidase |
| *mltB* | -0.386 | 1.296 | | P41052 | P41052 | Membrane-bound lytic murein transglycosylase B |
| *dusA* | 0.936 | 4.206 | | P32695 | P32695 | tRNA-dihydrouridine synthase A |
| *yqhA* | 0.937 | 1.881 | | P67244 | P67244 | UPF0114 protein YqhA |
| *serB* | 0.938 | 2.227 | | P0AGB0 | P0AGB0 | Phosphoserine phosphatase |
| *rfaH* | 0.944 | 2.907 | | P0AFW0 | P0AFW0 | Transcription antitermination protein RfaH |
| *hrpA* | -0.501 | 1.049 | | P43329 | P43329 | ATP-dependent RNA helicase HrpA |
| *ytfL* | 0.953 | 2.942 | | P0AE45 | P0AE45 | UPF0053 inner membrane protein YtfL |
| *glnD* | 0.955 | 2.086 | | P27249 | P27249 | Bifunctional uridylyltransferase/uridylyl-removing enzyme |
| *kdsD* | -0.132 | 0.468 | | P45395 | P45395 | Arabinose 5-phosphate isomerase KdsD |
| *ybgF* | 0.955 | 2.196 | | P45955 | P45955 | Uncharacterized protein YbgF |
| *nuoA* | 0.957 | 2.793 | | P0AFC3 | P0AFC3 | NADH-quinone oxidoreductase subunit A |
| *ispE* | 0.959 | 3.631 | | P62615 | P62615 | 4-diphosphocytidyl-2-C-methyl-D-erythritol kinase |
| *tusB* | -0.132 | 0.098 | | P45530 | P45530 | Protein TusB |
| *thrC* | 0.966 | 1.388 | | P00934 | P00934 | Threonine synthase |
| *lpoA* | 0.974 | 2.577 | | P45464 | P45464 | Penicillin-binding protein activator LpoA |
| *lpxM* | 0.979 | 3.045 | | P24205 | P24205 | Lipid A biosynthesis myristoyltransferase |
| *yfiH* | 0.989 | 3.268 | | P33644 | P33644 | Laccase domain protein YfiH |
| *pstB* | 0.989 | 3.430 | | P0AAH0 | P0AAH0 | Phosphate import ATP-binding protein PstB |
| *qseC* | 0.994 | 1.400 | | P40719 | P40719 | Sensor protein QseC |
| *yfcD* | 1.004 | 3.317 | | P65556 | P65556 | Uncharacterized Nudix hydrolase YfcD |
| *yrfF* | 0.378 | 0.851 | | P45800 | P45800 | Putative membrane protein IgaA homolog |
| *pepB* | 1.005 | 4.160 | | P37095 | P37095 | Peptidase B |
| *nuoC* | 1.006 | 3.186 | | P33599 | P33599 | NADH-quinone oxidoreductase subunit C/D |
| *galK* | 1.012 | 3.425 | | P0A6T3 | P0A6T3 | Galactokinase |
| *gatZ* | 1.013 | 1.991 | | P0C8J8;P0C8K0 | P0C8J8 | D-tagatose-1,6-bisphosphate aldolase subunit GatZ |
| *trxA* | 1.014 | 2.249 | | P0AA25 | P0AA25 | Thioredoxin-1 |
| *yejF* | 1.020 | 1.861 | | P33916 | P33916 | Uncharacterized ABC transporter ATP-binding protein YejF |

| Gene names | og2(fold-Change | -Log(p-value) | Significant | Protein IDs | Majority protein IDs | Protein names |
|---|---|---|---|---|---|---|
| *ilvB* | 1.020 | 2.446 | | P08142 | P08142 | Acetolactate synthase isozyme 1 large subunit |
| *yiaD* | 1.030 | 2.787 | | P37665 | P37665 | Probable lipoprotein YiaD |
| *nuoB* | 1.032 | 4.049 | | P0AFC7 | P0AFC7 | NADH-quinone oxidoreductase subunit B |
| *ycjG* | 0.210 | 0.579 | | P51981 | P51981 | L-Ala-D/L-Glu epimerase |
| *frdA* | 1.033 | 2.177 | | P00363 | P00363 | Fumarate reductase flavoprotein subunit |
| *yggW* | 0.315 | 0.894 | | P52062 | P52062 | Oxygen-independent coproporphyrinogen-III oxidase-like protein YggW |
| *oppF* | 1.037 | 3.109 | | P77737 | P77737 | Oligopeptide transport ATP-binding protein OppF |
| *ygiB* | 1.038 | 3.666 | | P0ADT2 | P0ADT2 | UPF0441 protein YgiB |
| *iscU* | 1.046 | 2.702 | | P0ACD4 | P0ACD4 | Iron-sulfur cluster assembly scaffold protein IscU |
| *talB* | 1.052 | 4.435 | | P0A870 | P0A870 | Transaldolase B |
| *dnaA* | 1.061 | 2.952 | | P03004 | P03004 | Chromosomal replication initiator protein DnaA |
| *pheA* | 1.065 | 2.262 | | P0A9J8 | P0A9J8 | P-protein |
| *rlmD* | 0.543 | 0.252 | | P55135 | P55135 | 23S rRNA (uracil(1939)-C(5))-methyltransferase RlmD |
| *ptsI* | 1.070 | 4.173 | | P08839 | P08839 | Phosphoenolpyruvate-protein phosphotransferase |
| *rapA* | 0.023 | 0.061 | | P60240 | P60240 | RNA polymerase-associated protein RapA |
| *ahpC* | 1.086 | 2.913 | | P0AE08 | P0AE08 | Alkyl hydroperoxide reductase subunit C |
| *manA* | 1.092 | 3.799 | | P00946 | P00946 | Mannose-6-phosphate isomerase |
| *rhlB* | 1.094 | 3.043 | | P0A8J8 | P0A8J8 | ATP-dependent RNA helicase RhlB |
| *rplB* | -0.323 | 1.066 | | P60422 | P60422 | 50S ribosomal protein L2 |
| *rplC* | 0.137 | 0.790 | | P60438 | P60438 | 50S ribosomal protein L3 |
| *oppA* | 1.098 | 4.441 | | P23843 | P23843 | Periplasmic oligopeptide-binding protein |
| *guaC* | -0.550 | 1.175 | | P60560 | P60560 | GMP reductase |
| *hisH* | -0.324 | 0.487 | | P60595 | P60595 | Imidazole glycerol phosphate synthase subunit HisH |
| *rplX* | -0.026 | 0.102 | | P60624 | P60624 | 50S ribosomal protein L24 |
| *speB* | -0.027 | 0.077 | | P60651 | P60651 | Agmatinase |
| *hisF* | -0.369 | 0.532 | | P60664 | P60664 | Imidazole glycerol phosphate synthase subunit HisF |
| *yoaE* | 1.099 | 2.937 | | P0AEC0 | P0AEC0 | UPF0053 inner membrane protein YoaE |
| *panB* | 1.111 | 4.397 | | P31057 | P31057 | 3-methyl-2-oxobutanoate hydroxymethyltransferase |
| *rplD* | -0.136 | 1.182 | | P60723 | P60723 | 50S ribosomal protein L4 |
| *msbA* | -0.006 | 0.012 | | P60752 | P60752 | Lipid A export ATP-binding/permease protein MsbA |
| *hisG* | 0.011 | 0.012 | | P60757 | P60757 | ATP phosphoribosyltransferase |
| *plsY* | 0.509 | 0.731 | | P60782 | P60782 | Probable glycerol-3-phosphate acyltransferase |
| *lepA* | 0.109 | 0.517 | | P60785 | P60785 | Elongation factor 4 |
| *thiD* | 1.118 | 3.406 | | P76422 | P76422 | Hydroxymethylpyrimidine/phosphomethylpyrimidine kinase |
| *lgt* | -0.754 | 1.056 | | P60955 | P60955 | Prolipoprotein diacylglyceryl transferase |
| *rplV* | -0.185 | 0.822 | | P61175 | P61175 | 50S ribosomal protein L22 |
| *aspC* | 1.119 | 4.731 | | P00509 | P00509 | Aspartate aminotransferase |
| *gppA* | 1.122 | 2.217 | | P25552 | P25552 | Guanosine-5-triphosphate,3-diphosphate pyrophosphatase |
| *oppB* | 1.139 | 3.182 | | P0AFH2 | P0AFH2 | Oligopeptide transport system permease protein OppB |
| *ribE* | 0.239 | 0.812 | | P61714 | P61714 | 6,7-dimethyl-8-ribityllumazine synthase |
| *rmlA2* | 0.446 | 0.668 | | P61887 | P61887 | Glucose-1-phosphate thymidylyltransferase 2 |
| *lolB* | 1.139 | 2.562 | | P61320 | P61320 | Outer-membrane lipoprotein LolB |
| *fldA* | 0.054 | 0.110 | | P61949 | P61949 | Flavodoxin-1 |
| *rplE* | -0.005 | 0.018 | | P62399 | P62399 | 50S ribosomal protein L5 |
| *tktA* | 1.143 | 3.808 | | P27302 | P27302 | Transketolase 1 |
| *cydB* | 1.146 | 2.691 | | P0ABK2 | P0ABK2 | Cytochrome bd-I ubiquinol oxidase subunit 2 |
| *nrdD* | 1.152 | 2.323 | | P28903 | P28903 | Anaerobic ribonucleoside-triphosphate reductase |
| *ilvN* | 1.163 | 1.804 | | P0ADF8 | P0ADF8 | Acetolactate synthase isozyme 1 small subunit |
| *nuoH* | 1.166 | 1.450 | | P0AFD4 | P0AFD4 | NADH-quinone oxidoreductase subunit H |
| *rpiR* | 1.175 | 2.240 | | P0ACS7 | P0ACS7 | HTH-type transcriptional regulator RpiR |
| *nikR* | 1.176 | 3.516 | | P0A6Z6 | P0A6Z6 | Nickel-responsive regulator |
| *hemF* | 1.177 | 4.494 | | P36553 | P36553 | Oxygen-dependent coproporphyrinogen-III oxidase |
| *nudF* | 1.178 | 3.964 | | Q93K97 | Q93K97 | ADP-ribose pyrophosphatase |

| Gene names | og2(fold-Change | -Log(p-value) | Significant | Protein IDs | Majority protein IDs | Protein names |
|---|---|---|---|---|---|---|
| *gmhA* | 0.236 | 1.064 | | P63224 | P63224 | Phosphoheptose isomerase |
| *mog* | 1.178 | 4.224 | | P0AF03 | P0AF03 | Molybdopterin adenylyltransferase |
| *efp* | 1.181 | 4.564 | | P0A6N4 | P0A6N4 | Elongation factor P |
| *ptsH* | 1.183 | 2.675 | | P0AA04 | P0AA04 | Phosphocarrier protein HPr |
| *yheS* | 0.418 | 0.980 | | P63389 | P63389 | Uncharacterized ABC transporter ATP-binding protein YheS |
| *yhbS* | -0.298 | 0.864 | | P63417 | P63417 | Uncharacterized N-acetyltransferase YhbS |
| *folE* | 1.184 | 5.233 | | P0A6T5 | P0A6T5 | GTP cyclohydrolase 1 |
| *zntB* | -1.249 | 1.288 | | P64423 | P64423 | Zinc transport protein ZntB |
| *ypfJ* | 0.107 | 0.363 | | P64429 | P64429 | Uncharacterized protein YpfJ |
| *envZ* | 1.186 | 2.671 | | P0AEJ4 | P0AEJ4 | Osmolarity sensor protein EnvZ |
| *ydcL* | -0.104 | 0.289 | | P64451 | P64451 | Uncharacterized lipoprotein YdcL |
| *hupB* | 1.195 | 3.081 | | P0ACF4 | P0ACF4 | DNA-binding protein HU-beta |
| *cnu* | -0.256 | 1.013 | | P64467 | P64467 | OriC-binding nucleoid-associated protein |
| *ispH* | 1.219 | 3.773 | | P62623 | P62623 | 4-hydroxy-3-methylbut-2-enyl diphosphate reductase |
| *yebY* | 1.222 | 2.316 | | P64506 | P64506 | Uncharacterized protein YebY |
| *ubiG* | 1.224 | 4.150 | | P17993 | P17993 | Ubiquinone biosynthesis O-methyltransferase |
| *yecN* | 0.009 | 0.013 | | P64515 | P64515 | Inner membrane protein YecN |
| *panM* | 1.225 | 2.832 | | P37613 | P37613 | PanD maturation factor |
| *ucpA* | 1.225 | 4.737 | | P37440 | P37440 | Oxidoreductase UcpA |
| *mtlA* | 1.228 | 1.858 | | P00550 | P00550 | PTS system mannitol-specific EIICBA component |
| *yeaD* | 1.230 | 4.142 | | P39173 | P39173 | Putative glucose-6-phosphate 1-epimerase |
| *yqjI* | 0.238 | 1.189 | | P64588 | P64588 | Transcriptional regulator YqjI |
| *yraP* | 0.410 | 1.291 | | P64596 | P64596 | Uncharacterized protein YraP |
| *mlaB* | -0.072 | 0.303 | | P64602 | P64602 | Probable phospholipid ABC transporter-binding protein MlaB |
| *hsdM* | 1.230 | 3.887 | | P08957 | P08957 | Type I restriction enzyme EcoKI M protein |
| *dadA* | 1.235 | 2.874 | | P0A6J5 | P0A6J5 | D-amino acid dehydrogenase |
| *era* | 1.236 | 3.636 | | P06616 | P06616 | GTPase Era |
| *greB* | 1.237 | 3.508 | | P30128 | P30128 | Transcription elongation factor GreB |
| *uxuA* | 1.237 | 4.402 | | P24215 | P24215 | Mannonate dehydratase |
| *rsmE* | 1.238 | 4.909 | | P0AGL7 | P0AGL7 | Ribosomal RNA small subunit methyltransferase E |
| *yqcA* | -0.150 | 0.488 | | P65367 | P65367 | Uncharacterized protein YqcA |
| *fkpB* | 1.251 | 3.192 | | P0AEM0 | P0AEM0 | FKBP-type 16 kDa peptidyl-prolyl cis-trans isomerase |
| *creA* | 1.256 | 3.620 | | P0AE91 | P0AE91 | Protein CreA |
| *uxuB* | 1.258 | 3.695 | | P39160;P77260 | P39160 | D-mannonate oxidoreductase |
| *glpD* | 1.282 | 4.116 | | P13035 | P13035 | Aerobic glycerol-3-phosphate dehydrogenase |
| *cra* | 1.282 | 5.522 | | P0ACP1 | P0ACP1 | Catabolite repressor/activator |
| *rsmI* | 0.102 | 0.814 | | P67087 | P67087 | Ribosomal RNA small subunit methyltransferase I |
| *rsfS* | 1.290 | 2.580 | | P0AAT6 | P0AAT6 | Ribosomal silencing factor RsfS |
| *yajG* | 1.292 | 2.654 | | P0ADA5 | P0ADA5 | Uncharacterized lipoprotein YajG |
| *yqfB* | 0.240 | 0.697 | | P67603 | P67603 | UPF0267 protein YqfB |
| *yhaJ* | 0.202 | 0.897 | | P67660 | P67660 | Uncharacterized HTH-type transcriptional regulator YhaJ |
| *yifE* | 1.292 | 3.322 | | P0ADN2 | P0ADN2 | UPF0438 protein YifE |
| *hldD* | 0.074 | 0.374 | | P67910 | P67910 | ADP-L-glycero-D-manno-heptose-6-epimerase |
| *grcA* | -0.102 | 0.259 | | P68066 | P68066 | Autonomous glycyl radical cofactor |
| *frmB* | 1.295 | 3.418 | | P51025 | P51025 | S-formylglutathione hydrolase FrmB |
| *rpsU* | -0.144 | 0.590 | | P68679 | P68679 | 30S ribosomal protein S21 |
| *aroK* | 1.303 | 4.470 | | P0A6D7 | P0A6D7 | Shikimate kinase 1 |
| *atpE* | 0.498 | 1.086 | | P68699 | P68699 | ATP synthase subunit c |
| *pepA* | -0.238 | 1.115 | | P68767 | P68767 | Cytosol aminopeptidase |
| *rplY* | 0.037 | 0.176 | | P68919 | P68919 | 50S ribosomal protein L25 |
| *hemA* | 1.315 | 2.407 | | P0A6X1 | P0A6X1 | Glutamyl-tRNA reductase |
| *pgk* | 1.326 | 5.174 | | P0A799 | P0A799 | Phosphoglycerate kinase |
| *pncC* | 1.331 | 2.850 | | P0A6G3 | P0A6G3 | Nicotinamide-nucleotide amidohydrolase PncC |

| Gene names | og2(fold-Change | -Log(p-value) | Significant | Protein IDs | Majority protein IDs | Protein names |
|---|---|---|---|---|---|---|
| *rcsF* | 0.251 | 0.828 | | P69411 | P69411 | Outer membrane lipoprotein RcsF |
| *leuC* | 1.335 | 3.561 | | P0A6A6 | P0A6A6 | 3-isopropylmalate dehydratase large subunit |
| *leuA* | 1.337 | 3.505 | | P09151 | P09151 | 2-isopropylmalate synthase |
| *pyrC* | 1.338 | 5.601 | | P05020 | P05020 | Dihydroorotase |
| *alr* | 1.356 | 3.973 | | P0A6B4 | P0A6B4 | Alanine racemase, biosynthetic |
| *crr* | -0.063 | 0.248 | | P69783 | P69783 | Glucose-specific phosphotransferase enzyme IIA component |
| *dut* | 1.372 | 3.395 | | P06968 | P06968 | Deoxyuridine 5-triphosphate nucleotidohydrolase |
| *nei* | 1.375 | 3.472 | | P50465 | P50465 | Endonuclease 8 |
| *erpA* | 1.386 | 1.798 | | P0ACC3 | P0ACC3 | Iron-sulfur cluster insertion protein ErpA |
| *tusD* | 1.447 | 2.342 | | P45532 | P45532 | Sulfurtransferase TusD |
| *ubiF* | -0.338 | 0.551 | | P75728 | P75728 | 2-octaprenyl-3-methyl-6-methoxy-1,4-benzoquinol hydroxylase |
| *ybgK* | 0.305 | 1.057 | | P75745 | P75745 | Uncharacterized protein YbgK |
| *gsiA* | -0.174 | 0.079 | | P75796 | P75796 | Glutathione import ATP-binding protein GsiA |
| *gsiB* | -0.016 | 0.202 | | P75797 | P75797 | Glutathione-binding protein GsiB |
| *ybjI* | 0.045 | 0.178 | | P75809 | P75809 | Flavin mononucleotide phosphatase YbjI |
| *amiD* | -0.026 | 0.079 | | P75820 | P75820 | N-acetylmuramoyl-L-alanine amidase AmiD |
| *ltaE* | 0.031 | 0.088 | | P75823 | P75823 | Low specificity L-threonine aldolase |
| *macA* | -0.262 | 1.088 | | P75830 | P75830 | Macrolide export protein MacA |
| *macB* | -0.475 | 0.464 | | P75831 | P75831 | Macrolide export ATP-binding/permease protein MacB |
| *ycbX* | 0.084 | 0.695 | | P75863 | P75863 | Uncharacterized protein YcbX |
| *ghrA* | 0.104 | 1.153 | | P75913 | P75913 | Glyoxylate/hydroxypyruvate reductase A |
| *ycdX* | -0.149 | 0.313 | | P75914 | P75914 | Probable phosphatase YcdX |
| *ycdY* | -0.172 | 0.500 | | P75915 | P75915 | Chaperone protein YcdY |
| *yceM* | 0.612 | 0.981 | | P75931 | P75931 | Putative oxidoreductase YceM |
| *lolD* | 0.058 | 0.384 | | P75957 | P75957 | Lipoprotein-releasing system ATP-binding protein LolD |
| *lolE* | -0.257 | 0.784 | | P75958 | P75958 | Lipoprotein-releasing system transmembrane protein LolE |
| *jayE* | 0.196 | 0.591 | | P75981 | P75981 | Putative protein JayE from lambdoid prophage e14 region |
| *dhaK* | -0.291 | 0.917 | | P76015 | P76015 | PTS-dependent dihydroxyacetone kinase, dihydroxyacetone-binding subunit DhaK |
| *yciT* | -0.154 | 0.499 | | P76034 | P76034 | Uncharacterized HTH-type transcriptional regulator YciT |
| *ynfB* | 0.190 | 0.427 | | P76170 | P76170 | UPF0482 protein YnfB |
| *yniB* | 0.126 | 0.206 | | P76208 | P76208 | Uncharacterized protein YniB |
| *tsaB* | 0.044 | 0.190 | | P76256 | P76256 | tRNA threonylcarbamoyladenosine biosynthesis protein TsaB |
| *kdgR* | -0.019 | 0.085 | | P76268 | P76268 | Transcriptional regulator KdgR |
| *msrC* | 0.538 | 1.163 | | P76270 | P76270 | Free methionine-R-sulfoxide reductase |
| *yebT* | -0.286 | 1.223 | | P76272 | P76272 | Uncharacterized protein YebT |
| *yedQ* | 0.192 | 0.496 | | P76330 | P76330 | Probable diguanylate cyclase YedQ |
| *yedY* | -0.049 | 0.158 | | P76342 | P76342 | Sulfoxide reductase catalytic subunit YedY |
| *yeeJ* | -0.025 | 0.015 | | P76347 | P76347 | Uncharacterized protein YeeJ |
| *mdtA* | 0.006 | 0.036 | | P76397 | P76397 | Multidrug resistance protein MdtA |
| *yegQ* | 0.249 | 1.115 | | P76403 | P76403 | Uncharacterized protease YegQ |
| *rtn* | 0.365 | 0.323 | | P76446 | P76446 | Protein Rtn |
| *mlaA* | 0.175 | 0.339 | | P76506 | P76506 | Probable phospholipid-binding lipoprotein MlaA |
| *yfeX* | 0.087 | 0.488 | | P76536 | P76536 | Probable deferrochelatase/peroxidase YfeX |
| *yfeY* | 0.032 | 0.062 | | P76537 | P76537 | Uncharacterized protein YfeY |
| *ypeA* | -0.307 | 0.938 | | P76539 | P76539 | Acetyltransferase YpeA |
| *eutL* | -0.317 | 0.533 | | P76541 | P76541 | Ethanolamine utilization protein EutL |
| *yfgD* | -0.109 | 0.275 | | P76569 | P76569 | Uncharacterized protein YfgD |
| *pdxY* | 0.002 | 0.006 | | P77150 | P77150 | Pyridoxamine kinase |
| *mnmC* | 0.485 | 0.886 | | P77182 | P77182 | tRNA 5-methylaminomethyl-2-thiouridine biosynthesis protein MnmC |
| *dsbG* | -0.320 | 0.841 | | P77202 | P77202 | Thiol:disulfide interchange protein DsbG |
| *prpD* | -0.114 | 0.080 | | P77243 | P77243 | 2-methylcitrate dehydratase |
| *rsxG* | 0.122 | 0.248 | | P77285 | P77285 | Electron transport complex subunit RsxG |
| *yfdH* | 0.244 | 0.630 | | P77293 | P77293 | Bactoprenol glucosyl transferase homolog from prophage CPS-53 |

| Gene names | og2(fold-Change | -Log(p-value) | Significant | Protein IDs | Majority protein IDs | Protein names |
|---|---|---|---|---|---|---|
| *fetB* | 0.055 | 0.057 | | P77307 | P77307 | Probable iron export permease protein FetB |
| *mscK* | -0.029 | 0.107 | | P77338 | P77338 | Mechanosensitive channel MscK |
| *mppA* | -0.080 | 0.237 | | P77348 | P77348 | Periplasmic murein peptide-binding protein |
| *alaC* | -0.157 | 0.706 | | P77434 | P77434 | Glutamate-pyruvate aminotransferase AlaC |
| *sufC* | -1.532 | 1.187 | | P77499 | P77499 | Probable ATP-dependent transporter SufC |
| *sufD* | -0.792 | 1.029 | | P77689 | P77689 | FeS cluster assembly protein SufD |
| *gnsB* | 0.304 | 0.648 | | P77695 | P77695 | Protein GnsB |
| *ydiJ* | -0.129 | 0.749 | | P77748 | P77748 | Uncharacterized protein YdiJ |
| *bamB* | -0.351 | 1.025 | | P77774 | P77774 | Outer membrane protein assembly factor BamB |
| *menI* | 0.295 | 0.776 | | P77781 | P77781 | 1,4-dihydroxy-2-naphthoyl-CoA hydrolase |
| *fsaA* | 0.205 | 0.412 | | P78055 | P78055 | Fructose-6-phosphate aldolase 1 |
| *ynjE* | -0.438 | 1.167 | | P78067 | P78067 | Thiosulfate sulfurtransferase YnjE |
| *yobH* | 0.022 | 0.044 | | Q2MB16 | Q2MB16 | Uncharacterized protein YobH |
| *ygeR* | -0.315 | 1.150 | | Q46798 | Q46798 | Uncharacterized lipoprotein YgeR |
| *yqcC* | -0.060 | 0.133 | | Q46919 | Q46919 | Uncharacterized protein YqcC |
| *queF* | 0.238 | 1.080 | | Q46920 | Q46920 | NADPH-dependent 7-cyano-7-deazaguanine reductase |
| *dinJ* | 0.152 | 0.434 | | Q47150 | Q47150 | Antitoxin DinJ |
| *truD* | 0.332 | 1.199 | | Q57261 | Q57261 | tRNA pseudouridine synthase D |
| *copA* | -0.174 | 0.446 | | Q59385 | Q59385 | Copper-exporting P-type ATPase A |
