## Supplementary material for "Laboratory evolution of *Escherichia coli* enables life based on fluorinated amino acids": Auxiliary Table 3_Metabolomics

| Feature identifiers | | | | | |
|---|---|---|---|---|---|
| ClusterIdx | m/z | RT [min] | Subnetwork | Compound name | Adduct |
| 491 | 205.195 | 9.007 | -1 | Cadinene | M+H |
| 24 | 399.252 | 8.753 | 58 | Spectral Match to Tris(2-butoxyethyl) phosphate from NIST14 | M+H |
| 44 | 421.232 | 8.753 | -1 | Spectral Match to Tris(2-butoxyethyl) phosphate from NIST14 | M+Na |
| 130 | 242.100 | 0.564 | 54 | Spectral Match to Pantothenic acid from NIST14 | M+Na |
| 479 | 242.101 | 0.569 | 54 | Spectral Match to Pantothenic acid from NIST14 | M+Na |
| 478 | 204.087 | 0.529 | 51 | Spectral Match to N-Acetyl-D-glucosamine from NIST14 | M+H-H2O |
| 200 | 311.256 | 7.678 | 39 | Spectral Match to Monopalmitolein (9c) from NIST14 | M+H-H2O |
| 461 | 311.256 | 7.795 | 39 | Spectral Match to Monopalmitolein (9c) from NIST14 | M+H-H2O |
| 536 | 339.290 | 8.561 | 39 | Spectral Match to Monoelaidin from NIST14 | M+H-H2O |
| 574 | 339.289 | 8.380 | 39 | Spectral Match to Monoelaidin from NIST14 | M+H-H2O |
| 666 | 188.071 | 1.307 | 27 | Spectral Match to L-Tryptophan from NIST14 | M+H-NH3 |
| 6 | 166.086 | 0.517 | 41 | Spectral Match to DL-Phenylalanine from NIST14 | M+H |
| 441 | 223.090 | 6.761 | -1 | Spectral Match to Diethyl phthalate from NIST14 | M+H |
| 378 | 391.285 | 10.644 | -1 | Spectral Match to Bis(2-ethylhexyl) phthalate from NIST14 | M+H |
| 12 | 245.096 | 3.603 | 62 | Spectral Match to Biotin from NIST14 | M+H |
| 408 | 489.184 | 3.610 | 62 | Spectral Match to Biotin from NIST14 | 2M+H |
| 467 | 250.118 | 7.417 | -1 | Spectral Match to 2,4,6-Triallyloxy-1,3,5-triazine from NIST14 | M+H |
| 462 | 482.321 | 9.064 | 4 | Spectral Match to 1-Stearoyl-2-hydroxy-sn-glycero-3-phosphoethanolamine from NIST14 | M+H |
| 832 | 577.516 | 12.699 | 39 | Spectral Match to 1-Palmitoyl-2-oleoyl-sn-glycerol from NIST14 | M+H-H2O |
| 17 | 454.293 | 8.329 | 4 | Spectral Match to 1-Palmitoyl-2-hydroxy-sn-glycero-3-phosphoethanolamine from NIST14 | M+H |
| 89 | 454.293 | 8.190 | 4 | Spectral Match to 1-Palmitoyl-2-hydroxy-sn-glycero-3-phosphoethanolamine from NIST14 | M+H |
| 211 | 476.267 | 8.328 | 4 | Spectral Match to 1-Hexadecanoyl-sn-glycero-3-phosphoethanolamine from NIST14 | M+Na |
| 743 | 907.578 | 8.331 | -1 | Spectral Match to 1-Hexadecanoyl-sn-glycero-3-phosphoethanolamine from NIST14 | 2M+H |
| 138 | 338.342 | 10.765 | 39 | Spectral Match to 13-Docosenamide, (Z)- from NIST14 | M+H |
| 456 | 338.342 | 9.820 | 39 | Spectral Match to 13-Docosenamide, (Z)- from NIST14 | M+H |
| 767 | 688.493 | 12.701 | 4 | Spectral Match to 1,2-Dipalmitoleoyl-sn-glycero-3-phosphoethanolamine from NIST14 | M+H |
| 30 | 480.309 | 8.382 | 4 | Spectral Match to 1-(9Z-Octadecenoyl)-sn-glycero-3-phosphoethanolamine from NIST14 | M+H |
| 36 | 480.309 | 8.512 | 4 | Spectral Match to 1-(9Z-Octadecenoyl)-sn-glycero-3-phosphoethanolamine from NIST14 | M+H |
| 342 | 377.146 | 3.769 | -1 | Spectral Match to (-)-Riboflavin from NIST14 | M+H |
| 473 | 220.118 | 0.532 | 61 | Pantothenic Acid | M+H |
| 372 | 628.196 | 10.499 | 40 | 2-[3,4-bis[[(2S,3R,4S,5S,6R)-3,4,5-trihydroxy-6-(hydroxymethyl)oxan-2-yl]oxy]phenyl]-5,7-dihydroxychromen-4-one | M+NH4 |
| 3 | 432.238 | 7.526 | 24 | 7b,9-Dihydroxy-3-(hydroxymethyl)-1,1,6,8-tetramethyl-5-oxo-1,1a,1b,4,4a,5,7a,7b,8,9-decahydro-9aH-cyclopropa[3,4]benzo[1,2-e]azulen-9a-yl acetate | M+ACN+H |
| 161 | 432.238 | 7.249 | 24 | 7b,9-Dihydroxy-3-(hydroxymethyl)-1,1,6,8-tetramethyl-5-oxo-1,1a,1b,4,4a,5,7a,7b,8,9-decahydro-9aH-cyclopropa[3,4]benzo[1,2-e]azulen-9a-yl acetate | M+ACN+H |
| 442 | 432.238 | 7.812 | 24 | 7b,9-Dihydroxy-3-(hydroxymethyl)-1,1,6,8-tetramethyl-5-oxo-1,1a,1b,4,4a,5,7a,7b,8,9-decahydro-9aH-cyclopropa[3,4]benzo[1,2-e]azulen-9a-yl acetate | M+ACN+H |
| 2 | 415.212 | 7.526 | 24 | 2H-Oxireno[1,10a]phenanthro[3,2-b]furan-10(11bH)-one, 5,7-bis(acetyloxy)-3,3a,4,5,6,7,7a,7b,8,8a-decahydro-4,4,7a,11-tetramethyl-, (1aS,3aR,5S,7S,7aR,7bS,8aR,11bR) | M-H2O+H |
| 78 | 415.212 | 7.248 | 24 | 2H-Oxireno[1,10a]phenanthro[3,2-b]furan-10(11bH)-one, 5,7-bis(acetyloxy)-3,3a,4,5,6,7,7a,7b,8,8a-decahydro-4,4,7a,11-tetramethyl-, (1aS,3aR,5S,7S,7aR,7bS,8aR,11bR) | M-H2O+H |
| 373 | 415.212 | 7.817 | 24 | 2H-Oxireno[1,10a]phenanthro[3,2-b]furan-10(11bH)-one, 5,7-bis(acetyloxy)-3,3a,4,5,6,7,7a,7b,8,8a-decahydro-4,4,7a,11-tetramethyl-, (1aS,3aR,5S,7S,7aR,7bS,8aR,11bR) | M-H2O+H |
| 40 | 327.078 | 8.430 | -1 | MassbankEU:SM879502 Triphenylphosphate | [M+H]+ |
| 435 | 313.143 | 8.965 | -1 | MassbankEU:SM836901 2-O-benzyl 1-O-butyl benzene-1,2-dicarboxylate | [M+H]+ |
| 80 | 223.089 | 2.607 | 27 | Massbank:BML01533 6-Fluoro-DL-tryptophan | [M+H]+ |
| 697 | 223.089 | 2.398 | 27 | Massbank:BML01533 6-Fluoro-DL-tryptophan | [M+H]+ |
| 798 | 223.089 | 2.491 | 27 | Massbank:BML01533 6-Fluoro-DL-tryptophan | [M+H]+ |
| 665 | 205.097 | 1.310 | 27 | Massbank:BML01185 L-Tryptophan | [M+H]+ |
| 154 | 457.112 | 3.497 | -1 | Flavine mononucleotide | M+H |
| 29 | 220.118 | 0.526 | 61 | D-Pantothenic acid | M+H |
| 681 | 331.165 | 0.498 | 41 | DL-Phenylalanine | 2M+H |
| 192 | 343.296 | 6.715 | 23 | Cocamidopropylbetaine | M+H |
| 589 | 481.400 | 5.241 | 3 | Bis(2,2,6,6-tetramethyl-4-piperidyl) sebacate | M+H |
| 603 | 810.134 | 3.065 | 20 | Acetyl-CoA | M+H |
| 521 | 716.523 | 12.701 | 4 | 2-Linoleoyl-1-palmitoyl-sn-glycero-3-phosphoethanolamine | M+H |
| 572 | 716.525 | 10.404 | 4 | 2-Linoleoyl-1-palmitoyl-sn-glycero-3-phosphoethanolamine | M+H |
| 654 | 716.524 | 10.304 | 4 | 2-Linoleoyl-1-palmitoyl-sn-glycero-3-phosphoethanolamine | M+H |
| 852 | 716.524 | 10.151 | 4 | 2-Linoleoyl-1-palmitoyl-sn-glycero-3-phosphoethanolamine | M+H |
| 524 | 426.262 | 7.583 | 4 | 1-Myristoyl-2-hydroxy-sn-glycero-3-phosphoethanolamine | M+H |
| 552 | 426.262 | 7.443 | 4 | 1-Myristoyl-2-hydroxy-sn-glycero-3-phosphoethanolamine | M+H |
| 507 | 718.539 | 12.704 | 4 | 1-Hexadecanoyl-2-(9Z-octadecenoyl)-sn-glycero-3-phosphoethanolamine | M+H |
| 523 | 718.539 | 10.302 | 4 | 1-Hexadecanoyl-2-(9Z-octadecenoyl)-sn-glycero-3-phosphoethanolamine | M+H |
| 632 | 718.538 | 10.143 | 4 | 1-Hexadecanoyl-2-(9Z-octadecenoyl)-sn-glycero-3-phosphoethanolamine | M+H |
| 823 | 771.514 | 12.696 | 4 | 1-Hexadecanoyl-2-(9Z-octadecenoyl)-sn-glycero-3-phospho-(1'-rac-glycerol) | M+Na |
| 346 | 502.290 | 8.380 | 4 | 1-(9Z-Octadecenoyl)-sn-glycero-3-phosphoethanolamine | M+Na |
| 388 | 502.291 | 8.511 | 4 | 1-(9Z-Octadecenoyl)-sn-glycero-3-phosphoethanolamine | M+Na |
| 476 | 979.349 | 0.522 | 91 | N/A | N/A |
| 488 | 995.318 | 0.510 | 91 | N/A | N/A |
| 52 | 256.264 | 10.439 | 88 | N/A | N/A |
| 427 | 284.295 | 10.783 | 88 | N/A | N/A |
| 62 | 291.096 | 6.457 | 87 | N/A | N/A |
| 153 | 291.096 | 6.559 | 87 | N/A | N/A |
| 374 | 748.356 | 4.106 | 84 | N/A | N/A |
| 384 | 783.875 | 4.128 | 84 | N/A | N/A |
| 284 | 243.196 | 7.929 | 80 | N/A | N/A |

| Feature identifiers | | | | | |
|---|---|---|---|---|---|
| ClusterIdx | m/z | RT [min] | Subnetwork | Compound name | Adduct |
| 415 | 225.185 | 7.934 | 80 | N/A | N/A |
| 667 | 176.074 | 0.736 | 79 | N/A | N/A |
| 668 | 218.121 | 4.865 | 79 | N/A | N/A |
| 670 | 246.152 | 6.015 | 79 | N/A | N/A |
| 674 | 232.137 | 5.471 | 79 | N/A | N/A |
| 675 | 204.105 | 4.140 | 79 | N/A | N/A |
| 772 | 551.192 | 3.315 | 78 | N/A | N/A |
| 774 | 615.714 | 3.354 | 78 | N/A | N/A |
| 776 | 486.670 | 3.240 | 78 | N/A | N/A |
| 777 | 680.235 | 3.406 | 78 | N/A | N/A |
| 780 | 357.628 | 3.065 | 78 | N/A | N/A |
| 46 | 226.158 | 9.291 | 77 | N/A | N/A |
| 201 | 282.221 | 10.073 | 77 | N/A | N/A |
| 634 | 282.221 | 10.084 | 77 | N/A | N/A |
| 150 | 818.299 | 3.185 | 75 | N/A | N/A |
| 323 | 1226.943 | 3.181 | 75 | N/A | N/A |
| 843 | 1226.943 | 3.183 | 75 | N/A | N/A |
| 186 | 135.079 | 7.525 | 74 | N/A | N/A |
| 319 | 135.079 | 4.463 | 74 | N/A | N/A |
| 180 | 529.790 | 4.168 | 73 | N/A | N/A |
| 264 | 465.270 | 4.196 | 73 | N/A | N/A |
| 48 | 431.565 | 4.119 | 72 | N/A | N/A |
| 170 | 417.549 | 3.504 | 72 | N/A | N/A |
| 314 | 396.544 | 3.209 | 72 | N/A | N/A |
| 117 | 832.302 | 3.461 | 70 | N/A | N/A |
| 371 | 1247.949 | 3.461 | 70 | N/A | N/A |
| 512 | 1247.454 | 3.462 | 70 | N/A | N/A |
| 7 | 271.106 | 8.022 | 69 | N/A | N/A |
| 45 | 390.122 | 6.709 | 69 | N/A | N/A |
| 112 | 388.106 | 7.488 | 69 | N/A | N/A |
| 209 | 414.122 | 6.104 | 69 | N/A | N/A |
| 663 | 414.122 | 6.419 | 69 | N/A | N/A |
| 468 | 432.238 | 6.913 | 68 | N/A | N/A |
| 792 | 404.208 | 6.914 | 68 | N/A | N/A |
| 793 | 387.181 | 6.913 | 68 | N/A | N/A |
| 671 | 331.207 | 5.234 | 67 | N/A | N/A |
| 795 | 317.195 | 4.828 | 67 | N/A | N/A |
| 84 | 280.188 | 5.797 | 65 | N/A | N/A |
| 158 | 308.222 | 6.723 | 65 | N/A | N/A |
| 263 | 693.878 | 3.983 | 64 | N/A | N/A |
| 392 | 665.367 | 3.986 | 64 | N/A | N/A |
| 703 | 204.534 | 6.795 | 63 | N/A | N/A |
| 704 | 193.026 | 6.808 | 63 | N/A | N/A |
| 88 | 227.084 | 3.606 | 62 | N/A | N/A |
| 215 | 290.153 | 3.607 | 62 | N/A | N/A |
| 4 | 182.985 | 12.752 | 60 | N/A | N/A |
| 5 | 154.990 | 12.767 | 60 | N/A | N/A |
| 19 | 138.996 | 12.748 | 60 | N/A | N/A |
| 23 | 205.017 | 12.741 | 60 | N/A | N/A |
| 49 | 196.017 | 12.766 | 60 | N/A | N/A |
| 63 | 224.012 | 12.748 | 60 | N/A | N/A |
| 75 | 195.994 | 12.775 | 60 | N/A | N/A |
| 20 | 131.534 | 12.743 | 59 | N/A | N/A |
| 409 | 131.534 | 13.009 | 59 | N/A | N/A |
| 465 | 131.534 | 13.069 | 59 | N/A | N/A |
| 696 | 131.534 | 4.507 | 59 | N/A | N/A |
| 266 | 444.309 | 8.752 | 58 | N/A | N/A |
| 490 | 350.102 | 0.493 | 57 | N/A | N/A |
| 495 | 364.118 | 0.634 | 57 | N/A | N/A |
| 134 | 519.318 | 10.374 | 56 | N/A | N/A |
| 658 | 519.309 | 10.382 | 56 | N/A | N/A |
| 498 | 532.098 | 0.493 | 55 | N/A | N/A |
| 501 | 532.099 | 0.497 | 55 | N/A | N/A |
| 132 | 711.319 | 4.210 | 53 | N/A | N/A |
| 282 | 624.289 | 4.196 | 53 | N/A | N/A |
| 133 | 226.180 | 5.976 | 52 | N/A | N/A |
| 542 | 384.349 | 10.283 | 52 | N/A | N/A |
| 131 | 614.463 | 9.741 | 51 | N/A | N/A |
| 243 | 388.269 | 6.414 | 51 | N/A | N/A |
| 472 | 479.186 | 0.514 | 51 | N/A | N/A |

| Feature identifiers | | | | | |
|---|---|---|---|---|---|
| ClusterIdx | m/z | RT [min] | Subnetwork | Compound name | Adduct |
| 475 | 957.367 | 0.530 | 51 | N/A | N/A |
| 477 | 524.246 | 0.489 | 51 | N/A | N/A |
| 480 | 268.076 | 0.497 | 51 | N/A | N/A |
| 482 | 498.161 | 0.503 | 51 | N/A | N/A |
| 483 | 644.267 | 0.510 | 51 | N/A | N/A |
| 484 | 279.584 | 0.532 | 51 | N/A | N/A |
| 487 | 276.062 | 0.490 | 51 | N/A | N/A |
| 492 | 737.248 | 0.522 | 51 | N/A | N/A |
| 493 | 506.147 | 0.488 | 51 | N/A | N/A |
| 500 | 287.570 | 0.486 | 51 | N/A | N/A |
| 687 | 441.187 | 0.508 | 51 | N/A | N/A |
| 732 | 632.474 | 10.196 | 51 | N/A | N/A |
| 802 | 860.514 | 9.355 | 51 | N/A | N/A |
| 168 | 471.307 | 5.923 | 50 | N/A | N/A |
| 616 | 473.286 | 4.748 | 50 | N/A | N/A |
| 144 | 393.207 | 10.310 | 49 | N/A | N/A |
| 176 | 409.183 | 10.451 | 49 | N/A | N/A |
| 43 | 418.117 | 7.633 | 48 | N/A | N/A |
| 146 | 551.149 | 6.292 | 48 | N/A | N/A |
| 363 | 418.117 | 7.570 | 48 | N/A | N/A |
| 119 | 343.130 | 3.690 | 47 | N/A | N/A |
| 182 | 537.334 | 9.562 | 47 | N/A | N/A |
| 231 | 509.303 | 8.961 | 47 | N/A | N/A |
| 239 | 245.616 | 7.678 | 46 | N/A | N/A |
| 393 | 266.129 | 7.678 | 46 | N/A | N/A |
| 429 | 266.129 | 7.807 | 46 | N/A | N/A |
| 506 | 734.471 | 12.708 | 45 | N/A | N/A |
| 544 | 734.471 | 10.338 | 45 | N/A | N/A |
| 550 | 765.459 | 12.703 | 45 | N/A | N/A |
| 646 | 734.471 | 10.261 | 45 | N/A | N/A |
| 652 | 762.502 | 12.706 | 45 | N/A | N/A |
| 370 | 514.353 | 10.382 | 43 | N/A | N/A |
| 740 | 744.593 | 12.218 | 43 | N/A | N/A |
| 94 | 247.596 | 7.524 | 42 | N/A | N/A |
| 110 | 256.601 | 7.527 | 42 | N/A | N/A |
| 233 | 268.115 | 7.525 | 42 | N/A | N/A |
| 283 | 554.177 | 10.161 | 40 | N/A | N/A |
| 545 | 684.203 | 10.983 | 40 | N/A | N/A |
| 630 | 702.214 | 10.746 | 40 | N/A | N/A |
| 693 | 463.131 | 9.688 | 40 | N/A | N/A |
| 821 | 776.232 | 11.006 | 40 | N/A | N/A |
| 858 | 610.185 | 12.140 | 40 | N/A | N/A |
| 143 | 228.196 | 7.148 | 39 | N/A | N/A |
| 230 | 228.196 | 6.462 | 39 | N/A | N/A |
| 824 | 549.485 | 10.160 | 39 | N/A | N/A |
| 826 | 549.486 | 12.704 | 39 | N/A | N/A |
| 840 | 549.486 | 9.853 | 39 | N/A | N/A |
| 850 | 549.485 | 10.361 | 39 | N/A | N/A |
| 159 | 480.238 | 3.674 | 37 | N/A | N/A |
| 332 | 959.468 | 3.674 | 37 | N/A | N/A |
| 301 | 467.250 | 5.033 | 36 | N/A | N/A |
| 333 | 933.494 | 5.044 | 36 | N/A | N/A |
| 420 | 516.785 | 5.231 | 36 | N/A | N/A |
| 68 | 217.105 | 4.009 | 35 | N/A | N/A |
| 330 | 234.134 | 4.002 | 35 | N/A | N/A |
| 431 | 681.651 | 10.701 | 33 | N/A | N/A |
| 434 | 399.395 | 7.402 | 33 | N/A | N/A |
| 171 | 278.185 | 3.992 | 32 | N/A | N/A |
| 437 | 280.186 | 4.202 | 32 | N/A | N/A |
| 560 | 271.180 | 3.637 | 32 | N/A | N/A |
| 35 | 255.159 | 6.531 | 31 | N/A | N/A |
| 99 | 272.185 | 6.531 | 31 | N/A | N/A |
| 466 | 269.175 | 7.631 | 31 | N/A | N/A |
| 28 | 297.133 | 7.153 | 30 | N/A | N/A |
| 38 | 431.207 | 7.153 | 30 | N/A | N/A |
| 56 | 447.202 | 6.602 | 30 | N/A | N/A |
| 91 | 448.233 | 7.152 | 30 | N/A | N/A |
| 102 | 429.191 | 6.409 | 30 | N/A | N/A |
| 163 | 297.133 | 7.367 | 30 | N/A | N/A |
| 202 | 448.233 | 7.470 | 30 | N/A | N/A |

| Feature identifiers | | | | | |
|---|---|---|---|---|---|
| ClusterIdx | m/z | RT [min] | Subnetwork | Compound name | Adduct |
| 208 | 431.207 | 7.472 | 30 | N/A | N/A |
| 213 | 447.202 | 7.074 | 30 | N/A | N/A |
| 219 | 447.202 | 6.409 | 30 | N/A | N/A |
| 235 | 297.133 | 6.600 | 30 | N/A | N/A |
| 245 | 469.184 | 6.601 | 30 | N/A | N/A |
| 249 | 429.191 | 6.602 | 30 | N/A | N/A |
| 251 | 297.133 | 6.410 | 30 | N/A | N/A |
| 262 | 447.202 | 6.882 | 30 | N/A | N/A |
| 411 | 527.717 | 3.679 | 29 | N/A | N/A |
| 444 | 791.074 | 3.678 | 29 | N/A | N/A |
| 355 | 500.380 | 6.151 | 28 | N/A | N/A |
| 391 | 442.338 | 5.719 | 28 | N/A | N/A |
| 394 | 309.227 | 4.773 | 28 | N/A | N/A |
| 446 | 384.296 | 5.267 | 28 | N/A | N/A |
| 458 | 674.506 | 7.346 | 28 | N/A | N/A |
| 631 | 790.590 | 8.085 | 28 | N/A | N/A |
| 635 | 549.416 | 9.554 | 28 | N/A | N/A |
| 860 | 694.522 | 10.428 | 28 | N/A | N/A |
| 357 | 206.062 | 2.609 | 27 | N/A | N/A |
| 440 | 237.106 | 4.092 | 27 | N/A | N/A |
| 699 | 206.061 | 2.395 | 27 | N/A | N/A |
| 799 | 206.062 | 2.458 | 27 | N/A | N/A |
| 448 | 213.144 | 5.071 | 26 | N/A | N/A |
| 692 | 381.331 | 9.796 | 26 | N/A | N/A |
| 210 | 445.290 | 4.792 | 25 | N/A | N/A |
| 227 | 467.300 | 4.849 | 25 | N/A | N/A |
| 250 | 423.276 | 4.729 | 25 | N/A | N/A |
| 275 | 401.261 | 4.663 | 25 | N/A | N/A |
| 307 | 379.249 | 4.593 | 25 | N/A | N/A |
| 353 | 489.316 | 4.904 | 25 | N/A | N/A |
| 358 | 505.391 | 10.706 | 25 | N/A | N/A |
| 402 | 527.403 | 10.694 | 25 | N/A | N/A |
| 459 | 549.417 | 10.679 | 25 | N/A | N/A |
| 535 | 463.345 | 9.103 | 25 | N/A | N/A |
| 540 | 441.330 | 9.123 | 25 | N/A | N/A |
| 638 | 529.383 | 9.048 | 25 | N/A | N/A |
| 653 | 507.370 | 9.067 | 25 | N/A | N/A |
| 662 | 511.328 | 4.955 | 25 | N/A | N/A |
| 759 | 571.429 | 10.663 | 25 | N/A | N/A |
| 859 | 533.341 | 5.004 | 25 | N/A | N/A |
| 9 | 460.270 | 7.525 | 24 | N/A | N/A |
| 25 | 846.443 | 7.526 | 24 | N/A | N/A |
| 67 | 446.254 | 7.523 | 24 | N/A | N/A |
| 81 | 460.270 | 7.249 | 24 | N/A | N/A |
| 139 | 476.266 | 7.526 | 24 | N/A | N/A |
| 261 | 397.201 | 7.526 | 24 | N/A | N/A |
| 272 | 429.191 | 6.884 | 24 | N/A | N/A |
| 316 | 531.275 | 7.525 | 24 | N/A | N/A |
| 375 | 462.213 | 6.365 | 24 | N/A | N/A |
| 418 | 445.186 | 6.364 | 24 | N/A | N/A |
| 419 | 460.270 | 7.814 | 24 | N/A | N/A |
| 447 | 450.204 | 8.853 | 24 | N/A | N/A |
| 469 | 445.186 | 6.211 | 24 | N/A | N/A |
| 578 | 448.233 | 6.302 | 24 | N/A | N/A |
| 582 | 448.233 | 6.159 | 24 | N/A | N/A |
| 636 | 462.213 | 6.211 | 24 | N/A | N/A |
| 637 | 431.207 | 6.148 | 24 | N/A | N/A |
| 770 | 474.249 | 6.864 | 24 | N/A | N/A |
| 790 | 431.207 | 6.021 | 24 | N/A | N/A |
| 221 | 371.326 | 7.517 | 23 | N/A | N/A |
| 31 | 328.919 | 12.753 | 22 | N/A | N/A |
| 39 | 312.943 | 12.757 | 22 | N/A | N/A |
| 70 | 287.888 | 12.760 | 22 | N/A | N/A |
| 92 | 313.078 | 6.460 | 22 | N/A | N/A |
| 187 | 312.943 | 12.757 | 22 | N/A | N/A |
| 196 | 313.078 | 6.532 | 22 | N/A | N/A |
| 255 | 313.235 | 8.431 | 22 | N/A | N/A |
| 558 | 313.235 | 8.075 | 22 | N/A | N/A |
| 197 | 272.258 | 8.673 | 21 | N/A | N/A |
| 223 | 300.290 | 9.476 | 21 | N/A | N/A |

| Feature identifiers | | | | | |
|---|---|---|---|---|---|
| ClusterIdx | m/z | RT [min] | Subnetwork | Compound name | Adduct |
| 460 | 406.354 | 10.232 | 21 | N/A | N/A |
| 157 | 429.146 | 4.153 | 20 | N/A | N/A |
| 166 | 549.571 | 3.384 | 20 | N/A | N/A |
| 481 | 549.078 | 3.211 | 20 | N/A | N/A |
| 520 | 432.026 | 3.379 | 20 | N/A | N/A |
| 584 | 432.026 | 3.502 | 20 | N/A | N/A |
| 596 | 405.571 | 3.038 | 20 | N/A | N/A |
| 600 | 359.104 | 0.595 | 20 | N/A | N/A |
| 611 | 401.115 | 2.454 | 20 | N/A | N/A |
| 622 | 411.021 | 3.222 | 20 | N/A | N/A |
| 315 | 515.353 | 9.549 | 19 | N/A | N/A |
| 326 | 321.148 | 3.682 | 19 | N/A | N/A |
| 595 | 366.206 | 3.688 | 19 | N/A | N/A |
| 494 | 453.189 | 7.153 | 18 | N/A | N/A |
| 591 | 453.189 | 7.367 | 18 | N/A | N/A |
| 861 | 453.187 | 7.297 | 18 | N/A | N/A |
| 21 | 453.169 | 7.511 | 17 | N/A | N/A |
| 292 | 453.168 | 7.525 | 17 | N/A | N/A |
| 640 | 453.186 | 7.468 | 17 | N/A | N/A |
| 59 | 281.138 | 7.525 | 16 | N/A | N/A |
| 97 | 299.149 | 4.461 | 16 | N/A | N/A |
| 172 | 299.148 | 7.523 | 16 | N/A | N/A |
| 296 | 344.206 | 4.461 | 16 | N/A | N/A |
| 381 | 316.176 | 4.464 | 16 | N/A | N/A |
| 42 | 473.318 | 10.647 | 15 | N/A | N/A |
| 72 | 680.481 | 12.424 | 15 | N/A | N/A |
| 74 | 475.298 | 9.832 | 15 | N/A | N/A |
| 95 | 492.324 | 9.832 | 15 | N/A | N/A |
| 96 | 654.333 | 10.021 | 15 | N/A | N/A |
| 105 | 490.345 | 10.647 | 15 | N/A | N/A |
| 129 | 663.454 | 12.425 | 15 | N/A | N/A |
| 141 | 708.511 | 12.424 | 15 | N/A | N/A |
| 181 | 637.306 | 10.020 | 15 | N/A | N/A |
| 205 | 518.376 | 10.647 | 15 | N/A | N/A |
| 273 | 287.141 | 6.170 | 15 | N/A | N/A |
| 298 | 515.329 | 10.526 | 15 | N/A | N/A |
| 311 | 474.317 | 10.257 | 15 | N/A | N/A |
| 317 | 457.288 | 10.396 | 15 | N/A | N/A |
| 318 | 387.194 | 7.399 | 15 | N/A | N/A |
| 426 | 682.364 | 10.020 | 15 | N/A | N/A |
| 433 | 532.356 | 10.526 | 15 | N/A | N/A |
| 452 | 519.377 | 10.257 | 15 | N/A | N/A |
| 529 | 520.355 | 9.833 | 15 | N/A | N/A |
| 726 | 1342.926 | 12.423 | 15 | N/A | N/A |
| 755 | 949.587 | 9.831 | 15 | N/A | N/A |
| 783 | 491.374 | 10.264 | 15 | N/A | N/A |
| 849 | 1370.957 | 12.421 | 15 | N/A | N/A |
| 585 | 834.093 | 3.504 | 14 | N/A | N/A |
| 619 | 792.082 | 3.197 | 14 | N/A | N/A |
| 714 | 542.297 | 4.609 | 13 | N/A | N/A |
| 717 | 492.763 | 4.165 | 13 | N/A | N/A |
| 51 | 247.034 | 5.943 | 12 | N/A | N/A |
| 698 | 247.034 | 6.796 | 12 | N/A | N/A |
| 718 | 291.023 | 6.253 | 12 | N/A | N/A |
| 16 | 122.547 | 12.762 | 11 | N/A | N/A |
| 60 | 122.547 | 14.272 | 11 | N/A | N/A |
| 103 | 122.547 | 2.944 | 11 | N/A | N/A |
| 113 | 122.547 | 2.472 | 11 | N/A | N/A |
| 115 | 122.547 | 13.880 | 11 | N/A | N/A |
| 120 | 122.547 | 2.832 | 11 | N/A | N/A |
| 123 | 122.547 | 13.067 | 11 | N/A | N/A |
| 125 | 122.547 | 2.720 | 11 | N/A | N/A |
| 127 | 122.547 | 0.660 | 11 | N/A | N/A |
| 135 | 122.547 | 13.843 | 11 | N/A | N/A |
| 140 | 122.547 | 2.622 | 11 | N/A | N/A |
| 145 | 122.547 | 2.598 | 11 | N/A | N/A |
| 147 | 122.547 | 2.749 | 11 | N/A | N/A |
| 151 | 122.547 | 13.172 | 11 | N/A | N/A |
| 156 | 122.547 | 2.528 | 11 | N/A | N/A |
| 160 | 122.547 | 2.262 | 11 | N/A | N/A |

| Feature identifiers | | | | | |
|---|---|---|---|---|---|
| ClusterIdx | m/z | RT [min] | Subnetwork | Compound name | Adduct |
| 169 | 122.547 | 13.121 | 11 | N/A | N/A |
| 174 | 122.547 | 2.436 | 11 | N/A | N/A |
| 177 | 122.547 | 13.217 | 11 | N/A | N/A |
| 178 | 122.547 | 2.294 | 11 | N/A | N/A |
| 183 | 122.547 | 1.983 | 11 | N/A | N/A |
| 185 | 122.547 | 13.651 | 11 | N/A | N/A |
| 189 | 122.547 | 0.617 | 11 | N/A | N/A |
| 206 | 122.547 | 1.515 | 11 | N/A | N/A |
| 207 | 122.547 | 2.189 | 11 | N/A | N/A |
| 216 | 122.547 | 1.421 | 11 | N/A | N/A |
| 220 | 122.547 | 2.347 | 11 | N/A | N/A |
| 248 | 122.547 | 2.158 | 11 | N/A | N/A |
| 587 | 122.547 | 2.681 | 11 | N/A | N/A |
| 644 | 122.547 | 2.919 | 11 | N/A | N/A |
| 645 | 122.547 | 2.389 | 11 | N/A | N/A |
| 656 | 122.547 | 2.584 | 11 | N/A | N/A |
| 657 | 122.547 | 2.792 | 11 | N/A | N/A |
| 710 | 122.547 | 3.154 | 11 | N/A | N/A |
| 715 | 122.547 | 3.094 | 11 | N/A | N/A |
| 719 | 122.547 | 2.222 | 11 | N/A | N/A |
| 773 | 122.547 | 13.630 | 11 | N/A | N/A |
| 61 | 537.536 | 11.439 | 10 | N/A | N/A |
| 109 | 565.567 | 11.404 | 10 | N/A | N/A |
| 295 | 582.594 | 11.445 | 10 | N/A | N/A |
| 816 | 593.598 | 10.672 | 10 | N/A | N/A |
| 191 | 608.393 | 9.449 | 9 | N/A | N/A |
| 313 | 638.404 | 9.642 | 9 | N/A | N/A |
| 348 | 624.388 | 9.076 | 9 | N/A | N/A |
| 15 | 824.976 | 3.461 | 8 | N/A | N/A |
| 18 | 810.973 | 3.183 | 8 | N/A | N/A |
| 26 | 838.980 | 3.758 | 8 | N/A | N/A |
| 65 | 1215.954 | 3.181 | 8 | N/A | N/A |
| 87 | 1236.959 | 3.462 | 8 | N/A | N/A |
| 121 | 837.625 | 3.464 | 8 | N/A | N/A |
| 136 | 823.621 | 3.183 | 8 | N/A | N/A |
| 194 | 618.984 | 3.459 | 8 | N/A | N/A |
| 204 | 1257.965 | 3.758 | 8 | N/A | N/A |
| 240 | 852.983 | 4.057 | 8 | N/A | N/A |
| 244 | 846.306 | 3.758 | 8 | N/A | N/A |
| 253 | 1069.394 | 4.071 | 8 | N/A | N/A |
| 257 | 851.628 | 3.759 | 8 | N/A | N/A |
| 258 | 608.481 | 3.183 | 8 | N/A | N/A |
| 288 | 1083.397 | 4.311 | 8 | N/A | N/A |
| 289 | 629.487 | 3.760 | 8 | N/A | N/A |
| 312 | 1055.391 | 3.820 | 8 | N/A | N/A |
| 352 | 1234.935 | 3.184 | 8 | N/A | N/A |
| 406 | 802.298 | 4.071 | 8 | N/A | N/A |
| 445 | 812.801 | 4.312 | 8 | N/A | N/A |
| 450 | 1041.387 | 3.604 | 8 | N/A | N/A |
| 457 | 1255.942 | 3.463 | 8 | N/A | N/A |
| 497 | 1278.970 | 4.056 | 8 | N/A | N/A |
| 504 | 1268.955 | 3.761 | 8 | N/A | N/A |
| 511 | 865.632 | 4.058 | 8 | N/A | N/A |
| 514 | 639.989 | 4.058 | 8 | N/A | N/A |
| 515 | 860.310 | 4.057 | 8 | N/A | N/A |
| 518 | 1276.945 | 3.762 | 8 | N/A | N/A |
| 519 | 856.952 | 3.759 | 8 | N/A | N/A |
| 522 | 1090.726 | 4.311 | 8 | N/A | N/A |
| 531 | 1076.722 | 4.075 | 8 | N/A | N/A |
| 533 | 1096.048 | 4.311 | 8 | N/A | N/A |
| 534 | 791.796 | 3.823 | 8 | N/A | N/A |
| 537 | 810.973 | 3.375 | 8 | N/A | N/A |
| 559 | 824.976 | 3.689 | 8 | N/A | N/A |
| 561 | 1289.962 | 4.060 | 8 | N/A | N/A |
| 567 | 1082.043 | 4.072 | 8 | N/A | N/A |
| 588 | 1027.384 | 3.427 | 8 | N/A | N/A |
| 590 | 1013.379 | 3.335 | 8 | N/A | N/A |
| 610 | 770.790 | 3.427 | 8 | N/A | N/A |
| 614 | 837.291 | 3.461 | 8 | N/A | N/A |
| 620 | 760.288 | 3.338 | 8 | N/A | N/A |

| Feature identifiers | | | | | |
|---|---|---|---|---|---|
| ClusterIdx | m/z | RT [min] | Subnetwork | Compound name | Adduct |
| 633 | 1062.719 | 3.822 | 8 | N/A | N/A |
| 650 | 629.236 | 3.761 | 8 | N/A | N/A |
| 659 | 813.049 | 4.312 | 8 | N/A | N/A |
| 664 | 866.987 | 4.353 | 8 | N/A | N/A |
| 672 | 1299.976 | 4.354 | 8 | N/A | N/A |
| 676 | 874.315 | 4.356 | 8 | N/A | N/A |
| 677 | 856.617 | 3.758 | 8 | N/A | N/A |
| 678 | 879.636 | 4.362 | 8 | N/A | N/A |
| 680 | 1297.948 | 4.058 | 8 | N/A | N/A |
| 683 | 975.108 | 4.244 | 8 | N/A | N/A |
| 685 | 1101.371 | 4.312 | 8 | N/A | N/A |
| 691 | 822.287 | 4.312 | 8 | N/A | N/A |
| 730 | 796.969 | 3.070 | 8 | N/A | N/A |
| 741 | 1068.040 | 3.824 | 8 | N/A | N/A |
| 746 | 1069.395 | 4.241 | 8 | N/A | N/A |
| 751 | 1221.446 | 3.186 | 8 | N/A | N/A |
| 757 | 1055.390 | 3.992 | 8 | N/A | N/A |
| 760 | 975.359 | 4.241 | 8 | N/A | N/A |
| 761 | 802.298 | 4.239 | 8 | N/A | N/A |
| 766 | 618.733 | 3.459 | 8 | N/A | N/A |
| 768 | 838.979 | 3.988 | 8 | N/A | N/A |
| 778 | 796.636 | 3.069 | 8 | N/A | N/A |
| 785 | 1215.954 | 3.392 | 8 | N/A | N/A |
| 787 | 608.481 | 3.384 | 8 | N/A | N/A |
| 788 | 1236.958 | 3.690 | 8 | N/A | N/A |
| 789 | 618.734 | 3.694 | 8 | N/A | N/A |
| 791 | 792.044 | 3.972 | 8 | N/A | N/A |
| 796 | 1069.727 | 4.238 | 8 | N/A | N/A |
| 800 | 824.643 | 3.674 | 8 | N/A | N/A |
| 801 | 1055.390 | 4.161 | 8 | N/A | N/A |
| 803 | 791.795 | 4.163 | 8 | N/A | N/A |
| 804 | 1300.145 | 4.240 | 8 | N/A | N/A |
| 805 | 810.973 | 3.576 | 8 | N/A | N/A |
| 807 | 1082.043 | 4.246 | 8 | N/A | N/A |
| 809 | 629.487 | 3.997 | 8 | N/A | N/A |
| 812 | 1268.461 | 3.759 | 8 | N/A | N/A |
| 820 | 1300.134 | 4.241 | 8 | N/A | N/A |
| 829 | 1137.663 | 4.195 | 8 | N/A | N/A |
| 830 | 1263.457 | 3.755 | 8 | N/A | N/A |
| 833 | 1076.722 | 4.246 | 8 | N/A | N/A |
| 836 | 1063.052 | 3.825 | 8 | N/A | N/A |
| 838 | 791.796 | 3.996 | 8 | N/A | N/A |
| 842 | 822.537 | 4.313 | 8 | N/A | N/A |
| 845 | 1257.962 | 3.997 | 8 | N/A | N/A |
| 847 | 781.292 | 3.601 | 8 | N/A | N/A |
| 848 | 811.784 | 4.077 | 8 | N/A | N/A |
| 851 | 1276.945 | 3.763 | 8 | N/A | N/A |
| 854 | 1194.947 | 3.069 | 8 | N/A | N/A |
| 862 | 964.605 | 4.154 | 8 | N/A | N/A |
| 354 | 828.611 | 3.182 | 7 | N/A | N/A |
| 360 | 842.615 | 3.460 | 7 | N/A | N/A |
| 649 | 842.948 | 3.459 | 7 | N/A | N/A |
| 811 | 828.945 | 3.185 | 7 | N/A | N/A |
| 22 | 274.274 | 6.821 | 6 | N/A | N/A |
| 104 | 302.305 | 7.664 | 6 | N/A | N/A |
| 277 | 330.337 | 8.495 | 6 | N/A | N/A |
| 341 | 358.368 | 9.237 | 6 | N/A | N/A |
| 321 | 311.185 | 7.063 | 5 | N/A | N/A |
| 347 | 423.310 | 10.200 | 5 | N/A | N/A |
| 661 | 468.369 | 10.166 | 5 | N/A | N/A |
| 11 | 452.277 | 7.678 | 4 | N/A | N/A |
| 155 | 474.259 | 7.679 | 4 | N/A | N/A |
| 238 | 474.258 | 7.808 | 4 | N/A | N/A |
| 350 | 690.508 | 12.705 | 4 | N/A | N/A |
| 424 | 511.304 | 8.562 | 4 | N/A | N/A |
| 486 | 690.508 | 10.348 | 4 | N/A | N/A |
| 489 | 712.490 | 12.701 | 4 | N/A | N/A |
| 505 | 712.490 | 10.356 | 4 | N/A | N/A |
| 510 | 485.288 | 8.510 | 4 | N/A | N/A |
| 513 | 743.484 | 12.700 | 4 | N/A | N/A |

| Feature identifiers | | | | | |
|---|---|---|---|---|---|
| ClusterIdx | m/z | RT [min] | Subnetwork | Compound name | Adduct |
| 516 | 483.272 | 7.795 | 4 | N/A | N/A |
| 517 | 740.522 | 12.702 | 4 | N/A | N/A |
| 532 | 738.526 | 10.287 | 4 | N/A | N/A |
| 554 | 903.547 | 7.681 | 4 | N/A | N/A |
| 555 | 467.277 | 8.509 | 4 | N/A | N/A |
| 562 | 743.484 | 10.306 | 4 | N/A | N/A |
| 565 | 496.304 | 8.835 | 4 | N/A | N/A |
| 577 | 483.272 | 7.944 | 4 | N/A | N/A |
| 581 | 511.304 | 8.710 | 4 | N/A | N/A |
| 583 | 452.277 | 7.806 | 4 | N/A | N/A |
| 639 | 690.508 | 10.220 | 4 | N/A | N/A |
| 643 | 712.490 | 10.268 | 4 | N/A | N/A |
| 669 | 466.293 | 8.072 | 4 | N/A | N/A |
| 679 | 466.293 | 8.206 | 4 | N/A | N/A |
| 701 | 440.278 | 7.956 | 4 | N/A | N/A |
| 713 | 662.477 | 12.666 | 4 | N/A | N/A |
| 721 | 533.285 | 8.561 | 4 | N/A | N/A |
| 722 | 440.278 | 7.818 | 4 | N/A | N/A |
| 724 | 684.458 | 12.702 | 4 | N/A | N/A |
| 727 | 465.262 | 7.796 | 4 | N/A | N/A |
| 728 | 505.254 | 7.795 | 4 | N/A | N/A |
| 729 | 507.269 | 8.510 | 4 | N/A | N/A |
| 731 | 493.293 | 8.564 | 4 | N/A | N/A |
| 736 | 676.492 | 12.707 | 4 | N/A | N/A |
| 745 | 493.293 | 8.710 | 4 | N/A | N/A |
| 749 | 500.300 | 7.793 | 4 | N/A | N/A |
| 756 | 533.285 | 8.715 | 4 | N/A | N/A |
| 775 | 704.523 | 12.703 | 4 | N/A | N/A |
| 779 | 704.523 | 10.367 | 4 | N/A | N/A |
| 781 | 726.506 | 12.701 | 4 | N/A | N/A |
| 813 | 494.288 | 8.224 | 4 | N/A | N/A |
| 825 | 664.490 | 9.773 | 4 | N/A | N/A |
| 837 | 766.559 | 12.703 | 4 | N/A | N/A |
| 841 | 676.491 | 10.424 | 4 | N/A | N/A |
| 844 | 698.475 | 10.420 | 4 | N/A | N/A |
| 853 | 1380.008 | 10.186 | 4 | N/A | N/A |
| 855 | 1380.006 | 9.800 | 4 | N/A | N/A |
| 27 | 316.212 | 3.277 | 3 | N/A | N/A |
| 37 | 430.244 | 4.459 | 3 | N/A | N/A |
| 55 | 300.200 | 3.938 | 3 | N/A | N/A |
| 69 | 416.228 | 3.930 | 3 | N/A | N/A |
| 122 | 243.183 | 3.050 | 3 | N/A | N/A |
| 124 | 334.178 | 3.942 | 3 | N/A | N/A |
| 128 | 357.214 | 4.538 | 3 | N/A | N/A |
| 149 | 350.208 | 4.208 | 3 | N/A | N/A |
| 167 | 599.391 | 3.938 | 3 | N/A | N/A |
| 190 | 294.861 | 4.052 | 3 | N/A | N/A |
| 224 | 302.196 | 0.594 | 3 | N/A | N/A |
| 246 | 309.181 | 4.412 | 3 | N/A | N/A |
| 268 | 328.200 | 4.325 | 3 | N/A | N/A |
| 290 | 272.185 | 3.424 | 3 | N/A | N/A |
| 293 | 293.191 | 3.577 | 3 | N/A | N/A |
| 306 | 300.180 | 5.792 | 3 | N/A | N/A |
| 322 | 713.423 | 4.538 | 3 | N/A | N/A |
| 328 | 252.166 | 3.641 | 3 | N/A | N/A |
| 329 | 332.872 | 4.600 | 3 | N/A | N/A |
| 343 | 471.307 | 6.113 | 3 | N/A | N/A |
| 361 | 441.788 | 4.079 | 3 | N/A | N/A |
| 386 | 241.204 | 5.244 | 3 | N/A | N/A |
| 396 | 469.328 | 4.897 | 3 | N/A | N/A |
| 401 | 256.850 | 3.715 | 3 | N/A | N/A |
| 404 | 699.407 | 4.204 | 3 | N/A | N/A |
| 413 | 292.192 | 4.464 | 3 | N/A | N/A |
| 439 | 585.375 | 3.578 | 3 | N/A | N/A |
| 443 | 384.773 | 3.708 | 3 | N/A | N/A |
| 463 | 293.191 | 3.406 | 3 | N/A | N/A |
| 499 | 599.391 | 3.791 | 3 | N/A | N/A |
| 526 | 485.359 | 3.054 | 3 | N/A | N/A |
| 527 | 441.788 | 4.168 | 3 | N/A | N/A |
| 530 | 290.193 | 3.898 | 3 | N/A | N/A |

| Feature identifiers | | | | | |
|---|---|---|---|---|---|
| ClusterIdx | m/z | RT [min] | Subnetwork | Compound name | Adduct |
| 553 | 332.207 | 4.839 | 3 | N/A | N/A |
| 568 | 379.210 | 4.343 | 3 | N/A | N/A |
| 623 | 343.200 | 3.890 | 3 | N/A | N/A |
| 624 | 263.684 | 4.125 | 3 | N/A | N/A |
| 627 | 230.153 | 3.714 | 3 | N/A | N/A |
| 629 | 347.535 | 4.414 | 3 | N/A | N/A |
| 747 | 540.426 | 9.375 | 3 | N/A | N/A |
| 750 | 568.458 | 9.844 | 3 | N/A | N/A |
| 758 | 263.684 | 4.034 | 3 | N/A | N/A |
| 782 | 220.150 | 0.536 | 3 | N/A | N/A |
| 784 | 498.804 | 4.596 | 3 | N/A | N/A |
| 810 | 294.861 | 4.145 | 3 | N/A | N/A |
| 817 | 384.772 | 3.711 | 3 | N/A | N/A |
| 819 | 292.192 | 4.644 | 3 | N/A | N/A |
| 93 | 256.264 | 9.789 | 2 | N/A | N/A |
| 142 | 284.295 | 10.325 | 2 | N/A | N/A |
| 175 | 308.279 | 8.430 | 2 | N/A | N/A |
| 218 | 482.408 | 9.562 | 2 | N/A | N/A |
| 226 | 526.434 | 9.538 | 2 | N/A | N/A |
| 228 | 336.311 | 9.373 | 2 | N/A | N/A |
| 229 | 291.253 | 8.431 | 2 | N/A | N/A |
| 241 | 424.364 | 9.342 | 2 | N/A | N/A |
| 242 | 380.338 | 9.364 | 2 | N/A | N/A |
| 247 | 352.306 | 8.443 | 2 | N/A | N/A |
| 269 | 438.382 | 9.584 | 2 | N/A | N/A |
| 270 | 408.372 | 10.006 | 2 | N/A | N/A |
| 276 | 570.460 | 9.514 | 2 | N/A | N/A |
| 278 | 512.417 | 9.299 | 2 | N/A | N/A |
| 280 | 468.390 | 9.320 | 2 | N/A | N/A |
| 291 | 366.322 | 8.959 | 2 | N/A | N/A |
| 294 | 452.398 | 9.985 | 2 | N/A | N/A |
| 297 | 556.443 | 9.273 | 2 | N/A | N/A |
| 302 | 247.227 | 8.431 | 2 | N/A | N/A |
| 303 | 392.338 | 10.385 | 2 | N/A | N/A |
| 310 | 496.424 | 9.963 | 2 | N/A | N/A |
| 336 | 364.344 | 10.020 | 2 | N/A | N/A |
| 337 | 540.449 | 9.937 | 2 | N/A | N/A |
| 344 | 476.301 | 4.018 | 2 | N/A | N/A |
| 349 | 440.359 | 8.424 | 2 | N/A | N/A |
| 359 | 600.469 | 9.251 | 2 | N/A | N/A |
| 362 | 322.295 | 8.962 | 2 | N/A | N/A |
| 364 | 614.485 | 9.493 | 2 | N/A | N/A |
| 366 | 396.333 | 8.432 | 2 | N/A | N/A |
| 367 | 422.389 | 10.267 | 2 | N/A | N/A |
| 369 | 436.364 | 10.348 | 2 | N/A | N/A |
| 377 | 422.347 | 10.141 | 2 | N/A | N/A |
| 382 | 466.373 | 10.105 | 2 | N/A | N/A |
| 383 | 319.284 | 9.373 | 2 | N/A | N/A |
| 387 | 466.415 | 10.248 | 2 | N/A | N/A |
| 389 | 526.434 | 9.632 | 2 | N/A | N/A |
| 390 | 468.390 | 9.206 | 2 | N/A | N/A |
| 399 | 520.331 | 4.142 | 2 | N/A | N/A |
| 400 | 484.385 | 8.410 | 2 | N/A | N/A |
| 412 | 512.416 | 9.184 | 2 | N/A | N/A |
| 414 | 528.411 | 8.398 | 2 | N/A | N/A |
| 416 | 564.360 | 4.248 | 2 | N/A | N/A |
| 417 | 644.495 | 9.227 | 2 | N/A | N/A |
| 422 | 480.390 | 10.316 | 2 | N/A | N/A |
| 432 | 556.443 | 9.161 | 2 | N/A | N/A |
| 438 | 584.475 | 9.912 | 2 | N/A | N/A |
| 455 | 628.501 | 9.881 | 2 | N/A | N/A |
| 464 | 608.392 | 4.346 | 2 | N/A | N/A |
| 471 | 352.306 | 8.095 | 2 | N/A | N/A |
| 528 | 308.279 | 8.075 | 2 | N/A | N/A |
| 539 | 688.521 | 9.203 | 2 | N/A | N/A |
| 541 | 363.310 | 9.363 | 2 | N/A | N/A |
| 546 | 291.253 | 8.075 | 2 | N/A | N/A |
| 548 | 396.333 | 8.097 | 2 | N/A | N/A |
| 556 | 732.548 | 9.180 | 2 | N/A | N/A |
| 564 | 440.359 | 8.096 | 2 | N/A | N/A |

| Feature identifiers | | | | | |
|---|---|---|---|---|---|
| ClusterIdx | m/z | RT [min] | Subnetwork | Compound name | Adduct |
| 570 | 484.385 | 8.094 | 2 | N/A | N/A |
| 571 | 308.279 | 8.290 | 2 | N/A | N/A |
| 573 | 528.411 | 8.088 | 2 | N/A | N/A |
| 576 | 247.227 | 8.064 | 2 | N/A | N/A |
| 579 | 335.280 | 8.098 | 2 | N/A | N/A |
| 580 | 572.437 | 8.084 | 2 | N/A | N/A |
| 598 | 396.333 | 8.315 | 2 | N/A | N/A |
| 604 | 440.359 | 8.302 | 2 | N/A | N/A |
| 608 | 672.527 | 9.854 | 2 | N/A | N/A |
| 615 | 350.328 | 9.687 | 2 | N/A | N/A |
| 618 | 658.511 | 9.464 | 2 | N/A | N/A |
| 625 | 702.536 | 9.449 | 2 | N/A | N/A |
| 647 | 394.355 | 9.697 | 2 | N/A | N/A |
| 50 | 701.620 | 10.613 | 1 | N/A | N/A |
| 199 | 419.364 | 7.499 | 1 | N/A | N/A |
| 689 | 685.624 | 12.053 | 1 | N/A | N/A |
| 1 | 110.020 | 12.761 | -1 | N/A | N/A |
| 8 | 437.194 | 7.524 | -1 | N/A | N/A |
| 10 | 120.081 | 0.519 | -1 | N/A | N/A |
| 13 | 242.284 | 8.651 | -1 | N/A | N/A |
| 14 | 851.398 | 7.527 | -1 | N/A | N/A |
| 32 | 437.194 | 7.249 | -1 | N/A | N/A |
| 33 | 559.518 | 11.442 | -1 | N/A | N/A |
| 34 | 867.372 | 7.528 | -1 | N/A | N/A |
| 41 | 587.549 | 11.407 | -1 | N/A | N/A |
| 47 | 163.991 | 12.748 | -1 | N/A | N/A |
| 53 | 243.088 | 4.621 | -1 | N/A | N/A |
| 54 | 399.209 | 10.100 | -1 | N/A | N/A |
| 57 | 685.436 | 12.425 | -1 | N/A | N/A |
| 58 | 250.112 | 5.753 | -1 | N/A | N/A |
| 64 | 255.073 | 4.542 | -1 | N/A | N/A |
| 66 | 261.109 | 5.951 | -1 | N/A | N/A |
| 71 | 267.077 | 3.607 | -1 | N/A | N/A |
| 73 | 453.168 | 7.250 | -1 | N/A | N/A |
| 76 | 324.142 | 7.584 | -1 | N/A | N/A |
| 77 | 372.882 | 12.723 | -1 | N/A | N/A |
| 79 | 321.131 | 4.460 | -1 | N/A | N/A |
| 82 | 269.015 | 5.941 | -1 | N/A | N/A |
| 83 | 230.141 | 12.721 | -1 | N/A | N/A |
| 85 | 233.079 | 4.825 | -1 | N/A | N/A |
| 86 | 239.089 | 4.007 | -1 | N/A | N/A |
| 90 | 332.913 | 12.737 | -1 | N/A | N/A |
| 98 | 195.102 | 7.496 | -1 | N/A | N/A |
| 100 | 440.099 | 7.644 | -1 | N/A | N/A |
| 101 | 235.169 | 8.489 | -1 | N/A | N/A |
| 106 | 670.153 | 5.750 | -1 | N/A | N/A |
| 107 | 183.150 | 4.641 | -1 | N/A | N/A |
| 108 | 281.088 | 8.127 | -1 | N/A | N/A |
| 111 | 262.165 | 4.000 | -1 | N/A | N/A |
| 114 | 244.191 | 5.975 | -1 | N/A | N/A |
| 116 | 228.232 | 9.984 | -1 | N/A | N/A |
| 118 | 219.175 | 8.820 | -1 | N/A | N/A |
| 126 | 296.219 | 7.149 | -1 | N/A | N/A |
| 137 | 431.183 | 10.115 | -1 | N/A | N/A |
| 148 | 319.187 | 7.588 | -1 | N/A | N/A |
| 152 | 707.875 | 4.385 | -1 | N/A | N/A |
| 162 | 174.999 | 12.765 | -1 | N/A | N/A |
| 164 | 338.147 | 5.707 | -1 | N/A | N/A |
| 165 | 269.210 | 8.432 | -1 | N/A | N/A |
| 173 | 451.246 | 9.774 | -1 | N/A | N/A |
| 179 | 723.608 | 10.615 | -1 | N/A | N/A |
| 184 | 434.186 | 7.527 | -1 | N/A | N/A |
| 188 | 270.279 | 10.247 | -1 | N/A | N/A |
| 193 | 242.284 | 7.258 | -1 | N/A | N/A |
| 195 | 312.326 | 10.525 | -1 | N/A | N/A |
| 198 | 499.165 | 7.526 | -1 | N/A | N/A |
| 203 | 122.547 | 1.761 | -1 | N/A | N/A |
| 212 | 409.201 | 10.114 | -1 | N/A | N/A |
| 214 | 270.279 | 9.930 | -1 | N/A | N/A |
| 217 | 266.248 | 6.949 | -1 | N/A | N/A |

| Feature identifiers | | | | | |
|---|---|---|---|---|---|
| ClusterIdx | m/z | RT [min] | Subnetwork | Compound name | Adduct |
| 222 | 495.300 | 10.647 | -1 | N/A | N/A |
| 225 | 330.300 | 9.151 | -1 | N/A | N/A |
| 232 | 250.178 | 6.460 | -1 | N/A | N/A |
| 234 | 555.250 | 4.014 | -1 | N/A | N/A |
| 236 | 149.988 | 12.784 | -1 | N/A | N/A |
| 237 | 640.348 | 4.374 | -1 | N/A | N/A |
| 252 | 157.084 | 0.657 | -1 | N/A | N/A |
| 254 | 111.039 | 13.492 | -1 | N/A | N/A |
| 256 | 297.241 | 9.395 | -1 | N/A | N/A |
| 259 | 249.185 | 8.504 | -1 | N/A | N/A |
| 260 | 290.269 | 6.924 | -1 | N/A | N/A |
| 265 | 535.153 | 8.517 | -1 | N/A | N/A |
| 267 | 453.168 | 7.817 | -1 | N/A | N/A |
| 271 | 283.104 | 5.343 | -1 | N/A | N/A |
| 274 | 537.311 | 10.526 | -1 | N/A | N/A |
| 279 | 149.024 | 6.761 | -1 | N/A | N/A |
| 281 | 467.147 | 6.891 | -1 | N/A | N/A |
| 285 | 408.309 | 10.166 | -1 | N/A | N/A |
| 286 | 291.195 | 8.162 | -1 | N/A | N/A |
| 287 | 573.110 | 8.520 | -1 | N/A | N/A |
| 299 | 437.194 | 8.754 | -1 | N/A | N/A |
| 300 | 604.812 | 4.484 | -1 | N/A | N/A |
| 304 | 365.137 | 8.808 | -1 | N/A | N/A |
| 305 | 480.273 | 3.605 | -1 | N/A | N/A |
| 308 | 659.288 | 10.020 | -1 | N/A | N/A |
| 309 | 165.076 | 4.458 | -1 | N/A | N/A |
| 320 | 341.267 | 9.373 | -1 | N/A | N/A |
| 324 | 270.315 | 9.426 | -1 | N/A | N/A |
| 325 | 326.378 | 9.409 | -1 | N/A | N/A |
| 327 | 131.049 | 0.513 | -1 | N/A | N/A |
| 331 | 260.165 | 6.499 | -1 | N/A | N/A |
| 334 | 469.184 | 6.409 | -1 | N/A | N/A |
| 335 | 469.163 | 6.314 | -1 | N/A | N/A |
| 338 | 499.780 | 3.714 | -1 | N/A | N/A |
| 339 | 615.581 | 10.604 | -1 | N/A | N/A |
| 340 | 354.553 | 5.749 | -1 | N/A | N/A |
| 345 | 236.142 | 5.355 | -1 | N/A | N/A |
| 351 | 451.173 | 6.881 | -1 | N/A | N/A |
| 356 | 310.173 | 6.844 | -1 | N/A | N/A |
| 365 | 519.261 | 9.835 | -1 | N/A | N/A |
| 368 | 273.167 | 4.221 | -1 | N/A | N/A |
| 376 | 288.181 | 4.136 | -1 | N/A | N/A |
| 379 | 496.301 | 10.257 | -1 | N/A | N/A |
| 380 | 497.280 | 9.835 | -1 | N/A | N/A |
| 385 | 188.069 | 0.519 | -1 | N/A | N/A |
| 395 | 721.506 | 12.424 | -1 | N/A | N/A |
| 397 | 459.263 | 10.315 | -1 | N/A | N/A |
| 398 | 445.291 | 10.157 | -1 | N/A | N/A |
| 403 | 763.517 | 12.425 | -1 | N/A | N/A |
| 405 | 528.788 | 4.417 | -1 | N/A | N/A |
| 407 | 455.335 | 10.125 | -1 | N/A | N/A |
| 410 | 830.949 | 3.185 | -1 | N/A | N/A |
| 421 | 514.951 | 4.135 | -1 | N/A | N/A |
| 423 | 409.180 | 7.399 | -1 | N/A | N/A |
| 425 | 707.607 | 12.056 | -1 | N/A | N/A |
| 428 | 366.633 | 3.310 | -1 | N/A | N/A |
| 430 | 379.862 | 4.127 | -1 | N/A | N/A |
| 436 | 267.137 | 8.329 | -1 | N/A | N/A |
| 449 | 658.341 | 3.958 | -1 | N/A | N/A |
| 451 | 254.660 | 3.065 | -1 | N/A | N/A |
| 453 | 441.346 | 7.492 | -1 | N/A | N/A |
| 454 | 409.163 | 6.914 | -1 | N/A | N/A |
| 470 | 469.163 | 6.021 | -1 | N/A | N/A |
| 474 | 501.170 | 0.519 | -1 | N/A | N/A |
| 485 | 517.142 | 0.507 | -1 | N/A | N/A |
| 496 | 226.012 | 12.739 | -1 | N/A | N/A |
| 502 | 102.034 | 0.568 | -1 | N/A | N/A |
| 503 | 228.207 | 4.644 | -1 | N/A | N/A |
| 508 | 1011.286 | 0.501 | -1 | N/A | N/A |
| 509 | 385.095 | 3.893 | -1 | N/A | N/A |

| Feature identifiers | | | | | |
|---|---|---|---|---|---|
| ClusterIdx | m/z | RT [min] | Subnetwork | Compound name | Adduct |
| 525 | 269.210 | 8.072 | -1 | N/A | N/A |
| 538 | 844.952 | 3.465 | -1 | N/A | N/A |
| 543 | 453.188 | 6.313 | -1 | N/A | N/A |
| 547 | 372.717 | 12.740 | -1 | N/A | N/A |
| 549 | 287.114 | 4.597 | -1 | N/A | N/A |
| 551 | 385.294 | 9.364 | -1 | N/A | N/A |
| 557 | 858.957 | 3.760 | -1 | N/A | N/A |
| 563 | 429.319 | 9.342 | -1 | N/A | N/A |
| 566 | 472.175 | 4.688 | -1 | N/A | N/A |
| 569 | 294.074 | 0.519 | -1 | N/A | N/A |
| 575 | 453.189 | 6.022 | -1 | N/A | N/A |
| 586 | 191.086 | 3.681 | -1 | N/A | N/A |
| 592 | 359.104 | 3.680 | -1 | N/A | N/A |
| 593 | 511.165 | 3.612 | -1 | N/A | N/A |
| 594 | 700.315 | 10.020 | -1 | N/A | N/A |
| 597 | 225.149 | 6.492 | -1 | N/A | N/A |
| 599 | 131.049 | 0.515 | -1 | N/A | N/A |
| 601 | 357.262 | 8.098 | -1 | N/A | N/A |
| 602 | 303.138 | 3.688 | -1 | N/A | N/A |
| 605 | 347.222 | 7.583 | -1 | N/A | N/A |
| 606 | 438.143 | 7.111 | -1 | N/A | N/A |
| 607 | 851.398 | 7.250 | -1 | N/A | N/A |
| 609 | 830.614 | 3.188 | -1 | N/A | N/A |
| 612 | 863.046 | 3.487 | -1 | N/A | N/A |
| 613 | 619.272 | 4.460 | -1 | N/A | N/A |
| 617 | 454.172 | 7.812 | -1 | N/A | N/A |
| 621 | 815.644 | 3.349 | -1 | N/A | N/A |
| 626 | 331.209 | 4.774 | -1 | N/A | N/A |
| 628 | 103.054 | 0.515 | -1 | N/A | N/A |
| 641 | 158.154 | 6.250 | -1 | N/A | N/A |
| 642 | 165.114 | 12.714 | -1 | N/A | N/A |
| 648 | 425.137 | 6.913 | -1 | N/A | N/A |
| 651 | 267.157 | 4.332 | -1 | N/A | N/A |
| 655 | 290.232 | 4.304 | -1 | N/A | N/A |
| 660 | 391.205 | 10.042 | -1 | N/A | N/A |
| 673 | 870.956 | 4.058 | -1 | N/A | N/A |
| 682 | 425.215 | 9.512 | -1 | N/A | N/A |
| 684 | 612.425 | 0.524 | -1 | N/A | N/A |
| 686 | 231.081 | 0.527 | -1 | N/A | N/A |
| 688 | 726.463 | 12.425 | -1 | N/A | N/A |
| 690 | 403.233 | 9.514 | -1 | N/A | N/A |
| 694 | 111.020 | 12.539 | -1 | N/A | N/A |
| 695 | 351.102 | 8.964 | -1 | N/A | N/A |
| 700 | 164.051 | 4.799 | -1 | N/A | N/A |
| 702 | 111.039 | 2.302 | -1 | N/A | N/A |
| 705 | 271.031 | 5.195 | -1 | N/A | N/A |
| 706 | 249.050 | 5.192 | -1 | N/A | N/A |
| 707 | 279.060 | 6.671 | -1 | N/A | N/A |
| 708 | 162.035 | 6.813 | -1 | N/A | N/A |
| 709 | 177.083 | 2.375 | -1 | N/A | N/A |
| 711 | 164.051 | 2.397 | -1 | N/A | N/A |
| 712 | 245.070 | 2.402 | -1 | N/A | N/A |
| 716 | 111.039 | 2.354 | -1 | N/A | N/A |
| 720 | 495.251 | 4.182 | -1 | N/A | N/A |
| 723 | 162.073 | 2.320 | -1 | N/A | N/A |
| 725 | 1347.881 | 12.426 | -1 | N/A | N/A |
| 733 | 297.083 | 6.817 | -1 | N/A | N/A |
| 734 | 471.252 | 7.681 | -1 | N/A | N/A |
| 735 | 597.678 | 0.494 | -1 | N/A | N/A |
| 737 | 701.411 | 12.427 | -1 | N/A | N/A |
| 738 | 253.602 | 7.678 | -1 | N/A | N/A |
| 739 | 749.548 | 12.237 | -1 | N/A | N/A |
| 742 | 287.650 | 8.328 | -1 | N/A | N/A |
| 744 | 247.129 | 0.492 | -1 | N/A | N/A |
| 748 | 654.456 | 10.203 | -1 | N/A | N/A |
| 752 | 286.520 | 6.796 | -1 | N/A | N/A |
| 753 | 335.125 | 8.963 | -1 | N/A | N/A |
| 754 | 533.934 | 3.609 | -1 | N/A | N/A |
| 762 | 533.282 | 9.675 | -1 | N/A | N/A |
| 763 | 1194.349 | 0.496 | -1 | N/A | N/A |

| Feature identifiers | | | | | |
|---|---|---|---|---|---|
| ClusterIdx | m/z | RT [min] | Subnetwork | Compound name | Adduct |
| 764 | 616.651 | 0.503 | -1 | N/A | N/A |
| 765 | 223.064 | 6.361 | -1 | N/A | N/A |
| 769 | 498.173 | 7.669 | -1 | N/A | N/A |
| 771 | 143.107 | 1.088 | -1 | N/A | N/A |
| 786 | 819.476 | 8.754 | -1 | N/A | N/A |
| 794 | 811.308 | 6.909 | -1 | N/A | N/A |
| 797 | 818.299 | 3.374 | -1 | N/A | N/A |
| 806 | 618.984 | 3.690 | -1 | N/A | N/A |
| 808 | 832.302 | 3.693 | -1 | N/A | N/A |
| 814 | 513.189 | 7.230 | -1 | N/A | N/A |
| 815 | 432.241 | 7.420 | -1 | N/A | N/A |
| 818 | 123.079 | 0.380 | -1 | N/A | N/A |
| 822 | 376.087 | 6.788 | -1 | N/A | N/A |
| 827 | 326.112 | 5.859 | -1 | N/A | N/A |
| 828 | 319.064 | 6.836 | -1 | N/A | N/A |
| 831 | 1295.589 | 10.022 | -1 | N/A | N/A |
| 834 | 628.471 | 3.460 | -1 | N/A | N/A |
| 835 | 850.609 | 3.462 | -1 | N/A | N/A |
| 839 | 372.717 | 10.091 | -1 | N/A | N/A |
| 846 | 681.132 | 5.759 | -1 | N/A | N/A |
| 856 | 1194.817 | 11.488 | -1 | N/A | N/A |
| 857 | 804.296 | 3.067 | -1 | N/A | N/A |
| 863 | 353.267 | 9.157 | -1 | N/A | N/A |

| Feature identifiers | | | | Library Hits | | | | | | | | | | |
|---|---|---|---|---|---|---|---|---|---|---|---|---|---|---|
| ClusterIdx | m/z | RT [min] | Subnetwork | Library | Instrument | Ion source | Ion mode | Library m/z | MasDiff | Cosine | PPM | Parent mass | Precursor charge | Precursor mass |
| 491 | 205.195 | 9.007 | -1 | METLIN | LC-ESI-QTOF | LC-ESI | Positive-0eV | 205.195 | 0.000 | 0.963 | 1.264 | 205.195 | 1.000 | 205.195 |
| 24 | 399.252 | 8.753 | 58 | GNPS-NIST14-MATCHES | HCD | ESI | Positive | 399.251 | 0.001 | 0.961 | 2.522 | 399.252 | 1.000 | 399.252 |
| 44 | 421.232 | 8.753 | -1 | GNPS-NIST14-MATCHES | HCD | ESI | Positive | 421.233 | 0.001 | 0.987 | 1.811 | 421.232 | 1.000 | 421.232 |
| 130 | 242.100 | 0.564 | 54 | GNPS-NIST14-MATCHES | IT/ion trap | ESI | Positive | 242.100 | 0.000 | 0.930 | 0.378 | 242.100 | 1.000 | 242.100 |
| 479 | 242.101 | 0.569 | 54 | GNPS-NIST14-MATCHES | IT/ion trap | ESI | Positive | 242.100 | 0.001 | 0.946 | 4.916 | 242.101 | 1.000 | 242.101 |
| 478 | 204.087 | 0.529 | 51 | GNPS-NIST14-MATCHES | Q-TOF | ESI | Positive | 204.084 | 0.003 | 0.999 | 12.710 | 204.087 | 1.000 | 204.087 |
| 200 | 311.256 | 7.678 | 39 | GNPS-NIST14-MATCHES | HCD | ESI | Positive | 311.257 | 0.001 | 0.954 | 2.941 | 311.256 | 1.000 | 311.256 |
| 461 | 311.256 | 7.795 | 39 | GNPS-NIST14-MATCHES | HCD | ESI | Positive | 311.257 | 0.001 | 0.952 | 2.157 | 311.256 | 1.000 | 311.256 |
| 536 | 339.290 | 8.561 | 39 | GNPS-NIST14-MATCHES | HCD | ESI | Positive | 339.289 | 0.001 | 0.980 | 1.979 | 339.290 | 1.000 | 339.290 |
| 574 | 339.289 | 8.380 | 39 | GNPS-NIST14-MATCHES | HCD | ESI | Positive | 339.289 | 0.000 | 0.984 | 1.349 | 339.289 | 1.000 | 339.289 |
| 666 | 188.071 | 1.307 | 27 | GNPS-NIST14-MATCHES | HCD | ESI | Positive | 188.070 | 0.001 | 0.988 | 4.057 | 188.071 | 1.000 | 188.071 |
| 6 | 166.086 | 0.517 | 41 | GNPS-NIST14-MATCHES | QqQ | ESI | Positive | 166.083 | 0.003 | 0.998 | 20.304 | 166.086 | 1.000 | 166.086 |
| 441 | 223.090 | 6.761 | -1 | GNPS-NIST14-MATCHES | QqQ | ESI | Positive | 223.097 | 0.007 | 0.995 | 32.898 | 223.090 | 1.000 | 223.090 |
| 378 | 391.285 | 10.644 | -1 | GNPS-NIST14-MATCHES | IT/ion trap | ESI | Positive | 391.284 | 0.001 | 0.971 | 2.262 | 391.285 | 1.000 | 391.285 |
| 12 | 245.096 | 3.603 | 62 | GNPS-NIST14-MATCHES | Q-TOF | ESI | Positive | 245.095 | 0.001 | 0.999 | 2.366 | 245.096 | 1.000 | 245.096 |
| 408 | 489.184 | 3.610 | 62 | GNPS-NIST14-MATCHES | Q-TOF | ESI | Positive | 489.180 | 0.004 | 0.983 | 7.798 | 489.184 | 1.000 | 489.184 |
| 467 | 250.118 | 7.417 | -1 | GNPS-NIST14-MATCHES | HCD | ESI | Positive | 250.119 | 0.001 | 0.959 | 5.918 | 250.118 | 1.000 | 250.118 |
| 462 | 482.321 | 9.064 | 4 | GNPS-NIST14-MATCHES | HCD | ESI | Positive | 482.320 | 0.001 | 0.860 | 1.645 | 482.321 | 1.000 | 482.321 |
| 832 | 577.516 | 12.699 | 39 | GNPS-NIST14-MATCHES | HCD | ESI | Positive | 577.518 | 0.002 | 0.938 | 4.016 | 577.516 | 1.000 | 577.516 |
| 17 | 454.293 | 8.329 | 4 | GNPS-NIST14-MATCHES | HCD | ESI | Positive | 454.291 | 0.002 | 0.876 | 4.568 | 454.293 | 1.000 | 454.293 |
| 89 | 454.293 | 8.190 | 4 | GNPS-NIST14-MATCHES | HCD | ESI | Positive | 454.291 | 0.002 | 0.891 | 4.904 | 454.293 | 1.000 | 454.293 |
| 211 | 476.267 | 8.328 | 4 | GNPS-NIST14-MATCHES | Q-TOF | ESI | Positive | 476.275 | 0.008 | 0.972 | 15.955 | 476.267 | 1.000 | 476.267 |
| 743 | 907.578 | 8.331 | -1 | GNPS-NIST14-MATCHES | Q-TOF | ESI | Positive | 907.579 | 0.001 | 0.820 | 0.942 | 907.578 | 1.000 | 907.578 |
| 138 | 338.342 | 10.765 | 39 | GNPS-NIST14-MATCHES | QqQ | ESI | Positive | 338.342 | 0.000 | 0.809 | 0.631 | 338.342 | 1.000 | 338.342 |
| 456 | 338.342 | 9.820 | 39 | GNPS-NIST14-MATCHES | QqQ | ESI | Positive | 338.341 | 0.001 | 0.900 | 2.345 | 338.342 | 1.000 | 338.342 |
| 767 | 688.493 | 12.701 | 4 | GNPS-NIST14-MATCHES | HCD | ESI | Positive | 688.486 | 0.007 | 0.806 | 10.372 | 688.493 | 1.000 | 688.493 |
| 30 | 480.309 | 8.382 | 4 | GNPS-NIST14-MATCHES | QqIT | ESI | Positive | 480.308 | 0.001 | 0.846 | 1.652 | 480.309 | 1.000 | 480.309 |
| 36 | 480.309 | 8.512 | 4 | GNPS-NIST14-MATCHES | QqIT | ESI | Positive | 480.308 | 0.001 | 0.948 | 1.716 | 480.309 | 1.000 | 480.309 |
| 342 | 377.146 | 3.769 | -1 | GNPS-NIST14-MATCHES | IT/ion trap | ESI | Positive | 377.145 | 0.001 | 0.903 | 2.266 | 377.146 | 1.000 | 377.146 |
| 473 | 220.118 | 0.532 | 61 | METLIN | LC-ESI-QTOF | LC-ESI | Positive-10eV | 220.118 | 0.000 | 0.956 | 1.109 | 220.118 | 1.000 | 220.118 |
| 372 | 628.196 | 10.499 | 40 | GNPS-NIH-NATURALPRODUCTSLIBRARY | Maxis II HD Q-TOF Bruker | LC-ESI | positive | 628.187 | 0.009 | 0.756 | 13.603 | 628.196 | 1.000 | 628.196 |
| 3 | 432.238 | 7.526 | 24 | GNPS-NIH-NATURALPRODUCTSLIBRARY | Maxis II HD Q-TOF Bruker | LC-ESI | positive | 432.238 | 0.000 | 0.896 | 0.847 | 432.238 | 1.000 | 432.238 |
| 161 | 432.238 | 7.249 | 24 | GNPS-NIH-NATURALPRODUCTSLIBRARY | Maxis II HD Q-TOF Bruker | LC-ESI | positive | 432.238 | 0.000 | 0.897 | 0.777 | 432.238 | 1.000 | 432.238 |
| 442 | 432.238 | 7.812 | 24 | GNPS-NIH-NATURALPRODUCTSLIBRARY | Maxis II HD Q-TOF Bruker | LC-ESI | positive | 432.238 | 0.000 | 0.882 | 0.918 | 432.238 | 1.000 | 432.238 |
| 2 | 415.212 | 7.526 | 24 | GNPS-NIH-NATURALPRODUCTSLIBRARY | Maxis II HD Q-TOF Bruker | LC-ESI | positive | 415.211 | 0.001 | 0.931 | 1.837 | 415.212 | 1.000 | 415.212 |
| 78 | 415.212 | 7.248 | 24 | GNPS-NIH-NATURALPRODUCTSLIBRARY | Maxis II HD Q-TOF Bruker | LC-ESI | positive | 415.211 | 0.001 | 0.936 | 1.837 | 415.212 | 1.000 | 415.212 |
| 373 | 415.212 | 7.817 | 24 | GNPS-NIH-NATURALPRODUCTSLIBRARY | Maxis II HD Q-TOF Bruker | LC-ESI | positive | 415.211 | 0.001 | 0.931 | 1.984 | 415.212 | 1.000 | 415.212 |
| 40 | 327.078 | 8.430 | -1 | MASSBANKEU | LC-ESI-QFT | ESI | Positive | 327.078 | 0.000 | 0.967 | 0.373 | 327.078 | 1.000 | 327.078 |
| 435 | 313.143 | 8.965 | -1 | MASSBANKEU | LC-ESI-QFT | ESI | Positive | 313.143 | 0.000 | 0.982 | 0.487 | 313.143 | 1.000 | 313.143 |
| 80 | 223.089 | 2.607 | 27 | MASSBANK | LC-ESI-QTOF | ESI | Positive | 223.088 | 0.001 | 0.972 | 5.882 | 223.089 | 1.000 | 223.089 |
| 697 | 223.089 | 2.398 | 27 | MASSBANK | LC-ESI-QTOF | ESI | Positive | 223.088 | 0.001 | 0.999 | 4.309 | 223.089 | 1.000 | 223.089 |
| 798 | 223.089 | 2.491 | 27 | MASSBANK | LC-ESI-QTOF | ESI | Positive | 223.088 | 0.001 | 0.998 | 4.172 | 223.089 | 1.000 | 223.089 |
| 665 | 205.097 | 1.310 | 27 | MASSBANK | LC-ESI-QTOF | ESI | Positive | 205.097 | 0.000 | 0.997 | 1.414 | 205.097 | 1.000 | 205.097 |
| 154 | 457.112 | 3.497 | -1 | NIST14 | HCD | ESI | Positive | 457.112 | 0.000 | 0.924 | 0.334 | 457.112 | 1.000 | 457.112 |
| 29 | 220.118 | 0.526 | 61 | GNPS-EMBL-MCF | Q-Exactive Plus | LC-ESI | Positive | 220.120 | 0.002 | 0.982 | 8.041 | 220.118 | 1.000 | 220.118 |
| 681 | 331.165 | 0.498 | 41 | NIST14 | Q-TOF | ESI | Positive | 331.165 | 0.000 | 0.993 | 0.461 | 331.165 | 1.000 | 331.165 |
| 192 | 343.296 | 6.715 | 23 | GNPS-LIBRARY | qTof | LC-ESI | Positive | 343.300 | 0.004 | 0.971 | 11.112 | 343.296 | 1.000 | 343.296 |
| 589 | 481.400 | 5.241 | 3 | NIST14 | HCD | ESI | Positive | 481.400 | 0.000 | 0.807 | 0.951 | 481.400 | 1.000 | 481.400 |
| 603 | 810.134 | 3.065 | 20 | METLIN | LC-ESI-QTOF | LC-ESI | Positive-40eV | 810.133 | 0.001 | 0.939 | 1.582 | 810.134 | 1.000 | 810.134 |
| 521 | 716.523 | 12.701 | 4 | NIST14 | HCD | ESI | Positive | 716.523 | 0.000 | 0.769 | 0.085 | 716.523 | 1.000 | 716.523 |
| 572 | 716.525 | 10.404 | 4 | NIST14 | HCD | ESI | Positive | 716.523 | 0.002 | 0.756 | 2.896 | 716.525 | 1.000 | 716.525 |
| 654 | 716.524 | 10.304 | 4 | NIST14 | HCD | ESI | Positive | 716.523 | 0.001 | 0.767 | 1.874 | 716.524 | 1.000 | 716.524 |
| 852 | 716.524 | 10.151 | 4 | NIST14 | HCD | ESI | Positive | 716.523 | 0.001 | 0.769 | 1.448 | 716.524 | 1.000 | 716.524 |
| 524 | 426.262 | 7.583 | 4 | NIST14 | HCD | ESI | Positive | 426.262 | 0.001 | 0.841 | 1.503 | 426.262 | 1.000 | 426.262 |
| 552 | 426.262 | 7.443 | 4 | NIST14 | HCD | ESI | Positive | 426.262 | 0.001 | 0.826 | 1.432 | 426.262 | 1.000 | 426.262 |
| 507 | 718.539 | 12.704 | 4 | NIST14 | HCD | ESI | Positive | 718.538 | 0.001 | 0.894 | 1.189 | 718.539 | 1.000 | 718.539 |
| 523 | 718.539 | 10.302 | 4 | NIST14 | HCD | ESI | Positive | 718.538 | 0.001 | 0.898 | 0.934 | 718.539 | 1.000 | 718.539 |
| 632 | 718.538 | 10.143 | 4 | NIST14 | HCD | ESI | Positive | 718.538 | 0.000 | 0.898 | 0.255 | 718.538 | 1.000 | 718.538 |
| 823 | 771.514 | 12.696 | 4 | NIST14 | Q-TOF | ESI | Positive | 771.515 | 0.001 | 0.885 | 1.424 | 771.514 | 1.000 | 771.514 |
| 346 | 502.290 | 8.380 | 4 | NIST14 | HCD | ESI | Positive | 502.290 | 0.000 | 0.820 | 0.911 | 502.290 | 1.000 | 502.290 |
| 388 | 502.291 | 8.511 | 4 | NIST14 | HCD | ESI | Positive | 502.290 | 0.000 | 0.865 | 0.425 | 502.291 | 1.000 | 502.291 |
| 476 | 979.349 | 0.522 | 91 | N/A | N/A | N/A | N/A | N/A | N/A | N/A | N/A | 979.349 | 1.000 | 979.349 |
| 488 | 995.318 | 0.510 | 91 | N/A | N/A | N/A | N/A | N/A | N/A | N/A | N/A | 995.318 | 1.000 | 995.318 |
| 52 | 256.264 | 10.439 | 88 | N/A | N/A | N/A | N/A | N/A | N/A | N/A | N/A | 256.264 | 1.000 | 256.264 |
| 427 | 284.295 | 10.783 | 88 | N/A | N/A | N/A | N/A | N/A | N/A | N/A | N/A | 284.295 | 1.000 | 284.295 |
| 62 | 291.096 | 6.457 | 87 | N/A | N/A | N/A | N/A | N/A | N/A | N/A | N/A | 291.096 | 1.000 | 291.096 |
| 153 | 291.096 | 6.559 | 87 | N/A | N/A | N/A | N/A | N/A | N/A | N/A | N/A | 291.096 | 1.000 | 291.096 |
| 374 | 748.356 | 4.106 | 84 | N/A | N/A | N/A | N/A | N/A | N/A | N/A | N/A | 1495.700 | 2.000 | 748.352 |
| 384 | 783.875 | 4.128 | 84 | N/A | N/A | N/A | N/A | N/A | N/A | N/A | N/A | 1566.740 | 2.000 | 783.871 |
| 284 | 243.196 | 7.929 | 80 | N/A | N/A | N/A | N/A | N/A | N/A | N/A | N/A | 243.196 | 1.000 | 243.196 |

| Feature identifiers | | | | Library Hits | | | | | | | | | | |
|---|---|---|---|---|---|---|---|---|---|---|---|---|---|---|
| ClusterIdx | m/z | RT [min] | Subnetwork | Library | Instrument | Ion source | Ion mode | Library m/z | MasDiff | Cosine | PPM | Parent mass | Precursor charge | Precursor mass |
| 415 | 225.185 | 7.934 | 80 | N/A | N/A | N/A | N/A | N/A | N/A | N/A | N/A | 225.185 | 1.000 | 225.185 |
| 667 | 176.074 | 0.736 | 79 | N/A | N/A | N/A | N/A | N/A | N/A | N/A | N/A | 176.074 | 1.000 | 176.074 |
| 668 | 218.121 | 4.865 | 79 | N/A | N/A | N/A | N/A | N/A | N/A | N/A | N/A | 218.121 | 1.000 | 218.121 |
| 670 | 246.152 | 6.015 | 79 | N/A | N/A | N/A | N/A | N/A | N/A | N/A | N/A | 246.152 | 1.000 | 246.152 |
| 674 | 232.137 | 5.471 | 79 | N/A | N/A | N/A | N/A | N/A | N/A | N/A | N/A | 232.137 | 1.000 | 232.137 |
| 675 | 204.105 | 4.140 | 79 | N/A | N/A | N/A | N/A | N/A | N/A | N/A | N/A | 204.105 | 1.000 | 204.105 |
| 772 | 551.192 | 3.315 | 78 | N/A | N/A | N/A | N/A | N/A | N/A | N/A | N/A | 1101.380 | 2.000 | 551.188 |
| 774 | 615.714 | 3.354 | 78 | N/A | N/A | N/A | N/A | N/A | N/A | N/A | N/A | 1230.420 | 2.000 | 615.710 |
| 776 | 486.670 | 3.240 | 78 | N/A | N/A | N/A | N/A | N/A | N/A | N/A | N/A | 972.333 | 2.000 | 486.666 |
| 777 | 680.235 | 3.406 | 78 | N/A | N/A | N/A | N/A | N/A | N/A | N/A | N/A | 1359.460 | 2.000 | 680.231 |
| 780 | 357.628 | 3.065 | 78 | N/A | N/A | N/A | N/A | N/A | N/A | N/A | N/A | 714.249 | 2.000 | 357.624 |
| 46 | 226.158 | 9.291 | 77 | N/A | N/A | N/A | N/A | N/A | N/A | N/A | N/A | 226.158 | 1.000 | 226.158 |
| 201 | 282.221 | 10.073 | 77 | N/A | N/A | N/A | N/A | N/A | N/A | N/A | N/A | 282.221 | 1.000 | 282.221 |
| 634 | 282.221 | 10.084 | 77 | N/A | N/A | N/A | N/A | N/A | N/A | N/A | N/A | 282.221 | 1.000 | 282.221 |
| 150 | 818.299 | 3.185 | 75 | N/A | N/A | N/A | N/A | N/A | N/A | N/A | N/A | 2452.880 | 3.000 | 818.294 |
| 323 | 1226.943 | 3.181 | 75 | N/A | N/A | N/A | N/A | N/A | N/A | N/A | N/A | 2452.880 | 2.000 | 1226.940 |
| 843 | 1226.943 | 3.183 | 75 | N/A | N/A | N/A | N/A | N/A | N/A | N/A | N/A | 2452.880 | 2.000 | 1226.940 |
| 186 | 135.079 | 7.525 | 74 | N/A | N/A | N/A | N/A | N/A | N/A | N/A | N/A | 135.079 | 1.000 | 135.079 |
| 319 | 135.079 | 4.463 | 74 | N/A | N/A | N/A | N/A | N/A | N/A | N/A | N/A | 135.079 | 1.000 | 135.079 |
| 180 | 529.790 | 4.168 | 73 | N/A | N/A | N/A | N/A | N/A | N/A | N/A | N/A | 1058.570 | 2.000 | 529.786 |
| 264 | 465.270 | 4.196 | 73 | N/A | N/A | N/A | N/A | N/A | N/A | N/A | N/A | 929.533 | 2.000 | 465.266 |
| 48 | 431.565 | 4.119 | 72 | N/A | N/A | N/A | N/A | N/A | N/A | N/A | N/A | 862.123 | 2.000 | 431.562 |
| 170 | 417.549 | 3.504 | 72 | N/A | N/A | N/A | N/A | N/A | N/A | N/A | N/A | 834.091 | 2.000 | 417.546 |
| 314 | 396.544 | 3.209 | 72 | N/A | N/A | N/A | N/A | N/A | N/A | N/A | N/A | 792.081 | 2.000 | 396.541 |
| 117 | 832.302 | 3.461 | 70 | N/A | N/A | N/A | N/A | N/A | N/A | N/A | N/A | 2494.890 | 3.000 | 832.297 |
| 371 | 1247.949 | 3.461 | 70 | N/A | N/A | N/A | N/A | N/A | N/A | N/A | N/A | 2494.890 | 2.000 | 1247.950 |
| 512 | 1247.454 | 3.462 | 70 | N/A | N/A | N/A | N/A | N/A | N/A | N/A | N/A | 2493.900 | 2.000 | 1247.450 |
| 7 | 271.106 | 8.022 | 69 | N/A | N/A | N/A | N/A | N/A | N/A | N/A | N/A | 271.106 | 1.000 | 271.106 |
| 45 | 390.122 | 6.709 | 69 | N/A | N/A | N/A | N/A | N/A | N/A | N/A | N/A | 390.122 | 1.000 | 390.122 |
| 112 | 388.106 | 7.488 | 69 | N/A | N/A | N/A | N/A | N/A | N/A | N/A | N/A | 388.106 | 1.000 | 388.106 |
| 209 | 414.122 | 6.104 | 69 | N/A | N/A | N/A | N/A | N/A | N/A | N/A | N/A | 414.122 | 1.000 | 414.122 |
| 663 | 414.122 | 6.419 | 69 | N/A | N/A | N/A | N/A | N/A | N/A | N/A | N/A | 414.122 | 1.000 | 414.122 |
| 468 | 432.238 | 6.913 | 68 | N/A | N/A | N/A | N/A | N/A | N/A | N/A | N/A | 432.238 | 1.000 | 432.238 |
| 792 | 404.208 | 6.914 | 68 | N/A | N/A | N/A | N/A | N/A | N/A | N/A | N/A | 404.208 | 1.000 | 404.208 |
| 793 | 387.181 | 6.913 | 68 | N/A | N/A | N/A | N/A | N/A | N/A | N/A | N/A | 387.181 | 1.000 | 387.181 |
| 671 | 331.207 | 5.234 | 67 | N/A | N/A | N/A | N/A | N/A | N/A | N/A | N/A | 331.207 | 1.000 | 331.207 |
| 795 | 317.195 | 4.828 | 67 | N/A | N/A | N/A | N/A | N/A | N/A | N/A | N/A | 317.195 | 1.000 | 317.195 |
| 84 | 280.188 | 5.797 | 65 | N/A | N/A | N/A | N/A | N/A | N/A | N/A | N/A | 280.188 | 1.000 | 280.188 |
| 158 | 308.222 | 6.723 | 65 | N/A | N/A | N/A | N/A | N/A | N/A | N/A | N/A | 308.222 | 1.000 | 308.222 |
| 263 | 693.878 | 3.983 | 64 | N/A | N/A | N/A | N/A | N/A | N/A | N/A | N/A | 1386.750 | 2.000 | 693.874 |
| 392 | 665.367 | 3.986 | 64 | N/A | N/A | N/A | N/A | N/A | N/A | N/A | N/A | 1329.730 | 2.000 | 665.363 |
| 703 | 204.534 | 6.795 | 63 | N/A | N/A | N/A | N/A | N/A | N/A | N/A | N/A | 408.060 | 2.000 | 204.530 |
| 704 | 193.026 | 6.808 | 63 | N/A | N/A | N/A | N/A | N/A | N/A | N/A | N/A | 385.044 | 2.000 | 193.022 |
| 88 | 227.084 | 3.606 | 62 | N/A | N/A | N/A | N/A | N/A | N/A | N/A | N/A | 227.084 | 1.000 | 227.084 |
| 215 | 290.153 | 3.607 | 62 | N/A | N/A | N/A | N/A | N/A | N/A | N/A | N/A | 290.153 | 1.000 | 290.153 |
| 4 | 182.985 | 12.752 | 60 | N/A | N/A | N/A | N/A | N/A | N/A | N/A | N/A | 182.985 | 1.000 | 182.985 |
| 5 | 154.990 | 12.767 | 60 | N/A | N/A | N/A | N/A | N/A | N/A | N/A | N/A | 154.990 | 1.000 | 154.990 |
| 19 | 138.996 | 12.748 | 60 | N/A | N/A | N/A | N/A | N/A | N/A | N/A | N/A | 138.996 | 1.000 | 138.996 |
| 23 | 205.017 | 12.741 | 60 | N/A | N/A | N/A | N/A | N/A | N/A | N/A | N/A | 205.017 | 1.000 | 205.017 |
| 49 | 196.017 | 12.766 | 60 | N/A | N/A | N/A | N/A | N/A | N/A | N/A | N/A | 196.017 | 1.000 | 196.017 |
| 63 | 224.012 | 12.748 | 60 | N/A | N/A | N/A | N/A | N/A | N/A | N/A | N/A | 224.012 | 1.000 | 224.012 |
| 75 | 195.994 | 12.775 | 60 | N/A | N/A | N/A | N/A | N/A | N/A | N/A | N/A | 195.994 | 1.000 | 195.994 |
| 20 | 131.534 | 12.743 | 59 | N/A | N/A | N/A | N/A | N/A | N/A | N/A | N/A | 262.059 | 2.000 | 131.530 |
| 409 | 131.534 | 13.009 | 59 | N/A | N/A | N/A | N/A | N/A | N/A | N/A | N/A | 262.059 | 2.000 | 131.530 |
| 465 | 131.534 | 13.069 | 59 | N/A | N/A | N/A | N/A | N/A | N/A | N/A | N/A | 262.059 | 2.000 | 131.530 |
| 696 | 131.534 | 4.507 | 59 | N/A | N/A | N/A | N/A | N/A | N/A | N/A | N/A | 262.059 | 2.000 | 131.530 |
| 266 | 444.309 | 8.752 | 58 | N/A | N/A | N/A | N/A | N/A | N/A | N/A | N/A | 444.309 | 1.000 | 444.309 |
| 490 | 350.102 | 0.493 | 57 | N/A | N/A | N/A | N/A | N/A | N/A | N/A | N/A | 350.102 | 1.000 | 350.102 |
| 495 | 364.118 | 0.634 | 57 | N/A | N/A | N/A | N/A | N/A | N/A | N/A | N/A | 364.118 | 1.000 | 364.118 |
| 134 | 519.318 | 10.374 | 56 | N/A | N/A | N/A | N/A | N/A | N/A | N/A | N/A | 519.318 | 1.000 | 519.318 |
| 658 | 519.309 | 10.382 | 56 | N/A | N/A | N/A | N/A | N/A | N/A | N/A | N/A | 519.309 | 1.000 | 519.309 |
| 498 | 532.098 | 0.493 | 55 | N/A | N/A | N/A | N/A | N/A | N/A | N/A | N/A | 532.098 | 1.000 | 532.098 |
| 501 | 532.099 | 0.497 | 55 | N/A | N/A | N/A | N/A | N/A | N/A | N/A | N/A | 532.099 | 1.000 | 532.099 |
| 132 | 711.319 | 4.210 | 53 | N/A | N/A | N/A | N/A | N/A | N/A | N/A | N/A | 711.319 | 1.000 | 711.319 |
| 282 | 624.289 | 4.196 | 53 | N/A | N/A | N/A | N/A | N/A | N/A | N/A | N/A | 624.289 | 1.000 | 624.289 |
| 133 | 226.180 | 5.976 | 52 | N/A | N/A | N/A | N/A | N/A | N/A | N/A | N/A | 226.180 | 1.000 | 226.180 |
| 542 | 384.349 | 10.283 | 52 | N/A | N/A | N/A | N/A | N/A | N/A | N/A | N/A | 384.349 | 1.000 | 384.349 |
| 131 | 614.463 | 9.741 | 51 | N/A | N/A | N/A | N/A | N/A | N/A | N/A | N/A | 614.463 | 1.000 | 614.463 |
| 243 | 388.269 | 6.414 | 51 | N/A | N/A | N/A | N/A | N/A | N/A | N/A | N/A | 388.269 | 1.000 | 388.269 |
| 472 | 479.186 | 0.514 | 51 | N/A | N/A | N/A | N/A | N/A | N/A | N/A | N/A | 479.186 | 1.000 | 479.186 |

| Feature identifiers | | | | Library Hits | | | | | | | | | | |
|---|---|---|---|---|---|---|---|---|---|---|---|---|---|---|
| ClusterIdx | m/z | RT [min] | Subnetwork | Library | Instrument | Ion source | Ion mode | Library m/z | MasDiff | Cosine | PPM | Parent mass | Precursor charge | Precursor mass |
| 475 | 957.367 | 0.530 | 51 | N/A | N/A | N/A | N/A | N/A | N/A | N/A | N/A | 957.367 | 1.000 | 957.367 |
| 477 | 524.246 | 0.489 | 51 | N/A | N/A | N/A | N/A | N/A | N/A | N/A | N/A | 524.246 | 1.000 | 524.246 |
| 480 | 268.076 | 0.497 | 51 | N/A | N/A | N/A | N/A | N/A | N/A | N/A | N/A | 535.145 | 2.000 | 268.072 |
| 482 | 498.161 | 0.503 | 51 | N/A | N/A | N/A | N/A | N/A | N/A | N/A | N/A | 995.314 | 2.000 | 498.157 |
| 483 | 644.267 | 0.510 | 51 | N/A | N/A | N/A | N/A | N/A | N/A | N/A | N/A | 644.267 | 1.000 | 644.267 |
| 484 | 279.584 | 0.532 | 51 | N/A | N/A | N/A | N/A | N/A | N/A | N/A | N/A | 558.160 | 2.000 | 279.580 |
| 487 | 276.062 | 0.490 | 51 | N/A | N/A | N/A | N/A | N/A | N/A | N/A | N/A | 276.062 | 1.000 | 276.062 |
| 492 | 737.248 | 0.522 | 51 | N/A | N/A | N/A | N/A | N/A | N/A | N/A | N/A | 1473.490 | 2.000 | 737.244 |
| 493 | 506.147 | 0.488 | 51 | N/A | N/A | N/A | N/A | N/A | N/A | N/A | N/A | 1011.290 | 2.000 | 506.143 |
| 500 | 287.570 | 0.486 | 51 | N/A | N/A | N/A | N/A | N/A | N/A | N/A | N/A | 574.131 | 2.000 | 287.566 |
| 687 | 441.187 | 0.508 | 51 | N/A | N/A | N/A | N/A | N/A | N/A | N/A | N/A | 441.187 | 1.000 | 441.187 |
| 732 | 632.474 | 10.196 | 51 | N/A | N/A | N/A | N/A | N/A | N/A | N/A | N/A | 632.474 | 1.000 | 632.474 |
| 802 | 860.514 | 9.355 | 51 | N/A | N/A | N/A | N/A | N/A | N/A | N/A | N/A | 860.514 | 1.000 | 860.514 |
| 168 | 471.307 | 5.923 | 50 | N/A | N/A | N/A | N/A | N/A | N/A | N/A | N/A | 471.307 | 1.000 | 471.307 |
| 616 | 473.286 | 4.748 | 50 | N/A | N/A | N/A | N/A | N/A | N/A | N/A | N/A | 473.286 | 1.000 | 473.286 |
| 144 | 393.207 | 10.310 | 49 | N/A | N/A | N/A | N/A | N/A | N/A | N/A | N/A | 393.207 | 1.000 | 393.207 |
| 176 | 409.183 | 10.451 | 49 | N/A | N/A | N/A | N/A | N/A | N/A | N/A | N/A | 409.183 | 1.000 | 409.183 |
| 43 | 418.117 | 7.633 | 48 | N/A | N/A | N/A | N/A | N/A | N/A | N/A | N/A | 418.117 | 1.000 | 418.117 |
| 146 | 551.149 | 6.292 | 48 | N/A | N/A | N/A | N/A | N/A | N/A | N/A | N/A | 551.149 | 1.000 | 551.149 |
| 363 | 418.117 | 7.570 | 48 | N/A | N/A | N/A | N/A | N/A | N/A | N/A | N/A | 418.117 | 1.000 | 418.117 |
| 119 | 343.130 | 3.690 | 47 | N/A | N/A | N/A | N/A | N/A | N/A | N/A | N/A | 343.130 | 1.000 | 343.130 |
| 182 | 537.334 | 9.562 | 47 | N/A | N/A | N/A | N/A | N/A | N/A | N/A | N/A | 537.334 | 1.000 | 537.334 |
| 231 | 509.303 | 8.961 | 47 | N/A | N/A | N/A | N/A | N/A | N/A | N/A | N/A | 509.303 | 1.000 | 509.303 |
| 239 | 245.616 | 7.678 | 46 | N/A | N/A | N/A | N/A | N/A | N/A | N/A | N/A | 490.224 | 2.000 | 245.612 |
| 393 | 266.129 | 7.678 | 46 | N/A | N/A | N/A | N/A | N/A | N/A | N/A | N/A | 531.250 | 2.000 | 266.125 |
| 429 | 266.129 | 7.807 | 46 | N/A | N/A | N/A | N/A | N/A | N/A | N/A | N/A | 531.250 | 2.000 | 266.125 |
| 506 | 734.471 | 12.708 | 45 | N/A | N/A | N/A | N/A | N/A | N/A | N/A | N/A | 734.471 | 1.000 | 734.471 |
| 544 | 734.471 | 10.338 | 45 | N/A | N/A | N/A | N/A | N/A | N/A | N/A | N/A | 734.471 | 1.000 | 734.471 |
| 550 | 765.459 | 12.703 | 45 | N/A | N/A | N/A | N/A | N/A | N/A | N/A | N/A | 765.459 | 1.000 | 765.459 |
| 646 | 734.471 | 10.261 | 45 | N/A | N/A | N/A | N/A | N/A | N/A | N/A | N/A | 734.471 | 1.000 | 734.471 |
| 652 | 762.502 | 12.706 | 45 | N/A | N/A | N/A | N/A | N/A | N/A | N/A | N/A | 762.502 | 1.000 | 762.502 |
| 370 | 514.353 | 10.382 | 43 | N/A | N/A | N/A | N/A | N/A | N/A | N/A | N/A | 514.353 | 1.000 | 514.353 |
| 740 | 744.593 | 12.218 | 43 | N/A | N/A | N/A | N/A | N/A | N/A | N/A | N/A | 744.593 | 1.000 | 744.593 |
| 94 | 247.596 | 7.524 | 42 | N/A | N/A | N/A | N/A | N/A | N/A | N/A | N/A | 494.184 | 2.000 | 247.592 |
| 110 | 256.601 | 7.527 | 42 | N/A | N/A | N/A | N/A | N/A | N/A | N/A | N/A | 512.195 | 2.000 | 256.598 |
| 233 | 268.115 | 7.525 | 42 | N/A | N/A | N/A | N/A | N/A | N/A | N/A | N/A | 535.223 | 2.000 | 268.111 |
| 283 | 554.177 | 10.161 | 40 | N/A | N/A | N/A | N/A | N/A | N/A | N/A | N/A | 554.177 | 1.000 | 554.177 |
| 545 | 684.203 | 10.983 | 40 | N/A | N/A | N/A | N/A | N/A | N/A | N/A | N/A | 684.203 | 1.000 | 684.203 |
| 630 | 702.214 | 10.746 | 40 | N/A | N/A | N/A | N/A | N/A | N/A | N/A | N/A | 702.214 | 1.000 | 702.214 |
| 693 | 463.131 | 9.688 | 40 | N/A | N/A | N/A | N/A | N/A | N/A | N/A | N/A | 463.131 | 1.000 | 463.131 |
| 821 | 776.232 | 11.006 | 40 | N/A | N/A | N/A | N/A | N/A | N/A | N/A | N/A | 776.232 | 1.000 | 776.232 |
| 858 | 610.185 | 12.140 | 40 | N/A | N/A | N/A | N/A | N/A | N/A | N/A | N/A | 610.185 | 1.000 | 610.185 |
| 143 | 228.196 | 7.148 | 39 | N/A | N/A | N/A | N/A | N/A | N/A | N/A | N/A | 228.196 | 1.000 | 228.196 |
| 230 | 228.196 | 6.462 | 39 | N/A | N/A | N/A | N/A | N/A | N/A | N/A | N/A | 228.196 | 1.000 | 228.196 |
| 824 | 549.485 | 10.160 | 39 | N/A | N/A | N/A | N/A | N/A | N/A | N/A | N/A | 549.485 | 1.000 | 549.485 |
| 826 | 549.486 | 12.704 | 39 | N/A | N/A | N/A | N/A | N/A | N/A | N/A | N/A | 549.486 | 1.000 | 549.486 |
| 840 | 549.486 | 9.853 | 39 | N/A | N/A | N/A | N/A | N/A | N/A | N/A | N/A | 549.486 | 1.000 | 549.486 |
| 850 | 549.485 | 10.361 | 39 | N/A | N/A | N/A | N/A | N/A | N/A | N/A | N/A | 549.485 | 1.000 | 549.485 |
| 159 | 480.238 | 3.674 | 37 | N/A | N/A | N/A | N/A | N/A | N/A | N/A | N/A | 959.468 | 2.000 | 480.234 |
| 332 | 959.468 | 3.674 | 37 | N/A | N/A | N/A | N/A | N/A | N/A | N/A | N/A | 959.468 | 1.000 | 959.468 |
| 301 | 467.250 | 5.033 | 36 | N/A | N/A | N/A | N/A | N/A | N/A | N/A | N/A | 933.493 | 2.000 | 467.247 |
| 333 | 933.494 | 5.044 | 36 | N/A | N/A | N/A | N/A | N/A | N/A | N/A | N/A | 933.494 | 1.000 | 933.494 |
| 420 | 516.785 | 5.231 | 36 | N/A | N/A | N/A | N/A | N/A | N/A | N/A | N/A | 1032.560 | 2.000 | 516.781 |
| 68 | 217.105 | 4.009 | 35 | N/A | N/A | N/A | N/A | N/A | N/A | N/A | N/A | 217.105 | 1.000 | 217.105 |
| 330 | 234.134 | 4.002 | 35 | N/A | N/A | N/A | N/A | N/A | N/A | N/A | N/A | 234.134 | 1.000 | 234.134 |
| 431 | 681.651 | 10.701 | 33 | N/A | N/A | N/A | N/A | N/A | N/A | N/A | N/A | 681.651 | 1.000 | 681.651 |
| 434 | 399.395 | 7.402 | 33 | N/A | N/A | N/A | N/A | N/A | N/A | N/A | N/A | 399.395 | 1.000 | 399.395 |
| 171 | 278.185 | 3.992 | 32 | N/A | N/A | N/A | N/A | N/A | N/A | N/A | N/A | 555.363 | 2.000 | 278.182 |
| 437 | 280.186 | 4.202 | 32 | N/A | N/A | N/A | N/A | N/A | N/A | N/A | N/A | 838.542 | 3.000 | 280.181 |
| 560 | 271.180 | 3.637 | 32 | N/A | N/A | N/A | N/A | N/A | N/A | N/A | N/A | 541.352 | 2.000 | 271.176 |
| 35 | 255.159 | 6.531 | 31 | N/A | N/A | N/A | N/A | N/A | N/A | N/A | N/A | 255.159 | 1.000 | 255.159 |
| 99 | 272.185 | 6.531 | 31 | N/A | N/A | N/A | N/A | N/A | N/A | N/A | N/A | 272.185 | 1.000 | 272.185 |
| 466 | 269.175 | 7.631 | 31 | N/A | N/A | N/A | N/A | N/A | N/A | N/A | N/A | 269.175 | 1.000 | 269.175 |
| 28 | 297.133 | 7.153 | 30 | N/A | N/A | N/A | N/A | N/A | N/A | N/A | N/A | 297.133 | 1.000 | 297.133 |
| 38 | 431.207 | 7.153 | 30 | N/A | N/A | N/A | N/A | N/A | N/A | N/A | N/A | 431.207 | 1.000 | 431.207 |
| 56 | 447.202 | 6.602 | 30 | N/A | N/A | N/A | N/A | N/A | N/A | N/A | N/A | 447.202 | 1.000 | 447.202 |
| 91 | 448.233 | 7.152 | 30 | N/A | N/A | N/A | N/A | N/A | N/A | N/A | N/A | 448.233 | 1.000 | 448.233 |
| 102 | 429.191 | 6.409 | 30 | N/A | N/A | N/A | N/A | N/A | N/A | N/A | N/A | 429.191 | 1.000 | 429.191 |
| 163 | 297.133 | 7.367 | 30 | N/A | N/A | N/A | N/A | N/A | N/A | N/A | N/A | 297.133 | 1.000 | 297.133 |
| 202 | 448.233 | 7.470 | 30 | N/A | N/A | N/A | N/A | N/A | N/A | N/A | N/A | 448.233 | 1.000 | 448.233 |

| Feature identifiers | | | | Library Hits | | | | | | | | | | |
|---|---|---|---|---|---|---|---|---|---|---|---|---|---|---|
| ClusterIdx | m/z | RT [min] | Subnetwork | Library | Instrument | Ion source | Ion mode | Library m/z | MasDiff | Cosine | PPM | Parent mass | Precursor charge | Precursor mass |
| 208 | 431.207 | 7.472 | 30 | N/A | N/A | N/A | N/A | N/A | N/A | N/A | N/A | 431.207 | 1.000 | 431.207 |
| 213 | 447.202 | 7.074 | 30 | N/A | N/A | N/A | N/A | N/A | N/A | N/A | N/A | 447.202 | 1.000 | 447.202 |
| 219 | 447.202 | 6.409 | 30 | N/A | N/A | N/A | N/A | N/A | N/A | N/A | N/A | 447.202 | 1.000 | 447.202 |
| 235 | 297.133 | 6.600 | 30 | N/A | N/A | N/A | N/A | N/A | N/A | N/A | N/A | 297.133 | 1.000 | 297.133 |
| 245 | 469.184 | 6.601 | 30 | N/A | N/A | N/A | N/A | N/A | N/A | N/A | N/A | 469.184 | 1.000 | 469.184 |
| 249 | 429.191 | 6.602 | 30 | N/A | N/A | N/A | N/A | N/A | N/A | N/A | N/A | 429.191 | 1.000 | 429.191 |
| 251 | 297.133 | 6.410 | 30 | N/A | N/A | N/A | N/A | N/A | N/A | N/A | N/A | 297.133 | 1.000 | 297.133 |
| 262 | 447.202 | 6.882 | 30 | N/A | N/A | N/A | N/A | N/A | N/A | N/A | N/A | 447.202 | 1.000 | 447.202 |
| 411 | 527.717 | 3.679 | 29 | N/A | N/A | N/A | N/A | N/A | N/A | N/A | N/A | 1581.140 | 3.000 | 527.712 |
| 444 | 791.074 | 3.678 | 29 | N/A | N/A | N/A | N/A | N/A | N/A | N/A | N/A | 1581.140 | 2.000 | 791.070 |
| 355 | 500.380 | 6.151 | 28 | N/A | N/A | N/A | N/A | N/A | N/A | N/A | N/A | 500.380 | 1.000 | 500.380 |
| 391 | 442.338 | 5.719 | 28 | N/A | N/A | N/A | N/A | N/A | N/A | N/A | N/A | 442.338 | 1.000 | 442.338 |
| 394 | 309.227 | 4.773 | 28 | N/A | N/A | N/A | N/A | N/A | N/A | N/A | N/A | 309.227 | 1.000 | 309.227 |
| 446 | 384.296 | 5.267 | 28 | N/A | N/A | N/A | N/A | N/A | N/A | N/A | N/A | 384.296 | 1.000 | 384.296 |
| 458 | 674.506 | 7.346 | 28 | N/A | N/A | N/A | N/A | N/A | N/A | N/A | N/A | 674.506 | 1.000 | 674.506 |
| 631 | 790.590 | 8.085 | 28 | N/A | N/A | N/A | N/A | N/A | N/A | N/A | N/A | 790.590 | 1.000 | 790.590 |
| 635 | 549.416 | 9.554 | 28 | N/A | N/A | N/A | N/A | N/A | N/A | N/A | N/A | 1097.820 | 2.000 | 549.412 |
| 860 | 694.522 | 10.428 | 28 | N/A | N/A | N/A | N/A | N/A | N/A | N/A | N/A | 1388.040 | 2.000 | 694.518 |
| 357 | 206.062 | 2.609 | 27 | N/A | N/A | N/A | N/A | N/A | N/A | N/A | N/A | 206.062 | 1.000 | 206.062 |
| 440 | 237.106 | 4.092 | 27 | N/A | N/A | N/A | N/A | N/A | N/A | N/A | N/A | 237.106 | 1.000 | 237.106 |
| 699 | 206.061 | 2.395 | 27 | N/A | N/A | N/A | N/A | N/A | N/A | N/A | N/A | 206.061 | 1.000 | 206.061 |
| 799 | 206.062 | 2.458 | 27 | N/A | N/A | N/A | N/A | N/A | N/A | N/A | N/A | 206.062 | 1.000 | 206.062 |
| 448 | 213.144 | 5.071 | 26 | N/A | N/A | N/A | N/A | N/A | N/A | N/A | N/A | 213.144 | 1.000 | 213.144 |
| 692 | 381.331 | 9.796 | 26 | N/A | N/A | N/A | N/A | N/A | N/A | N/A | N/A | 381.331 | 1.000 | 381.331 |
| 210 | 445.290 | 4.792 | 25 | N/A | N/A | N/A | N/A | N/A | N/A | N/A | N/A | 889.573 | 2.000 | 445.286 |
| 227 | 467.300 | 4.849 | 25 | N/A | N/A | N/A | N/A | N/A | N/A | N/A | N/A | 933.592 | 2.000 | 467.296 |
| 250 | 423.276 | 4.729 | 25 | N/A | N/A | N/A | N/A | N/A | N/A | N/A | N/A | 845.543 | 2.000 | 423.272 |
| 275 | 401.261 | 4.663 | 25 | N/A | N/A | N/A | N/A | N/A | N/A | N/A | N/A | 801.514 | 2.000 | 401.257 |
| 307 | 379.249 | 4.593 | 25 | N/A | N/A | N/A | N/A | N/A | N/A | N/A | N/A | 757.491 | 2.000 | 379.246 |
| 353 | 489.316 | 4.904 | 25 | N/A | N/A | N/A | N/A | N/A | N/A | N/A | N/A | 977.624 | 2.000 | 489.312 |
| 358 | 505.391 | 10.706 | 25 | N/A | N/A | N/A | N/A | N/A | N/A | N/A | N/A | 1009.770 | 2.000 | 505.387 |
| 402 | 527.403 | 10.694 | 25 | N/A | N/A | N/A | N/A | N/A | N/A | N/A | N/A | 1053.800 | 2.000 | 527.399 |
| 459 | 549.417 | 10.679 | 25 | N/A | N/A | N/A | N/A | N/A | N/A | N/A | N/A | 1097.830 | 2.000 | 549.413 |
| 535 | 463.345 | 9.103 | 25 | N/A | N/A | N/A | N/A | N/A | N/A | N/A | N/A | 925.681 | 2.000 | 463.341 |
| 540 | 441.330 | 9.123 | 25 | N/A | N/A | N/A | N/A | N/A | N/A | N/A | N/A | 881.652 | 2.000 | 441.326 |
| 638 | 529.383 | 9.048 | 25 | N/A | N/A | N/A | N/A | N/A | N/A | N/A | N/A | 1057.760 | 2.000 | 529.379 |
| 653 | 507.370 | 9.067 | 25 | N/A | N/A | N/A | N/A | N/A | N/A | N/A | N/A | 1013.730 | 2.000 | 507.366 |
| 662 | 511.328 | 4.955 | 25 | N/A | N/A | N/A | N/A | N/A | N/A | N/A | N/A | 1021.650 | 2.000 | 511.324 |
| 759 | 571.429 | 10.663 | 25 | N/A | N/A | N/A | N/A | N/A | N/A | N/A | N/A | 1141.850 | 2.000 | 571.425 |
| 859 | 533.341 | 5.004 | 25 | N/A | N/A | N/A | N/A | N/A | N/A | N/A | N/A | 1065.670 | 2.000 | 533.337 |
| 9 | 460.270 | 7.525 | 24 | N/A | N/A | N/A | N/A | N/A | N/A | N/A | N/A | 460.270 | 1.000 | 460.270 |
| 25 | 846.443 | 7.526 | 24 | N/A | N/A | N/A | N/A | N/A | N/A | N/A | N/A | 846.443 | 1.000 | 846.443 |
| 67 | 446.254 | 7.523 | 24 | N/A | N/A | N/A | N/A | N/A | N/A | N/A | N/A | 446.254 | 1.000 | 446.254 |
| 81 | 460.270 | 7.249 | 24 | N/A | N/A | N/A | N/A | N/A | N/A | N/A | N/A | 460.270 | 1.000 | 460.270 |
| 139 | 476.266 | 7.526 | 24 | N/A | N/A | N/A | N/A | N/A | N/A | N/A | N/A | 476.266 | 1.000 | 476.266 |
| 261 | 397.201 | 7.526 | 24 | N/A | N/A | N/A | N/A | N/A | N/A | N/A | N/A | 397.201 | 1.000 | 397.201 |
| 272 | 429.191 | 6.884 | 24 | N/A | N/A | N/A | N/A | N/A | N/A | N/A | N/A | 429.191 | 1.000 | 429.191 |
| 316 | 531.275 | 7.525 | 24 | N/A | N/A | N/A | N/A | N/A | N/A | N/A | N/A | 531.275 | 1.000 | 531.275 |
| 375 | 462.213 | 6.365 | 24 | N/A | N/A | N/A | N/A | N/A | N/A | N/A | N/A | 462.213 | 1.000 | 462.213 |
| 418 | 445.186 | 6.364 | 24 | N/A | N/A | N/A | N/A | N/A | N/A | N/A | N/A | 445.186 | 1.000 | 445.186 |
| 419 | 460.270 | 7.814 | 24 | N/A | N/A | N/A | N/A | N/A | N/A | N/A | N/A | 460.270 | 1.000 | 460.270 |
| 447 | 450.204 | 8.853 | 24 | N/A | N/A | N/A | N/A | N/A | N/A | N/A | N/A | 450.204 | 1.000 | 450.204 |
| 469 | 445.186 | 6.211 | 24 | N/A | N/A | N/A | N/A | N/A | N/A | N/A | N/A | 445.186 | 1.000 | 445.186 |
| 578 | 448.233 | 6.302 | 24 | N/A | N/A | N/A | N/A | N/A | N/A | N/A | N/A | 448.233 | 1.000 | 448.233 |
| 582 | 448.233 | 6.159 | 24 | N/A | N/A | N/A | N/A | N/A | N/A | N/A | N/A | 448.233 | 1.000 | 448.233 |
| 636 | 462.213 | 6.211 | 24 | N/A | N/A | N/A | N/A | N/A | N/A | N/A | N/A | 462.213 | 1.000 | 462.213 |
| 637 | 431.207 | 6.148 | 24 | N/A | N/A | N/A | N/A | N/A | N/A | N/A | N/A | 431.207 | 1.000 | 431.207 |
| 770 | 474.249 | 6.864 | 24 | N/A | N/A | N/A | N/A | N/A | N/A | N/A | N/A | 474.249 | 1.000 | 474.249 |
| 790 | 431.207 | 6.021 | 24 | N/A | N/A | N/A | N/A | N/A | N/A | N/A | N/A | 431.207 | 1.000 | 431.207 |
| 221 | 371.326 | 7.517 | 23 | N/A | N/A | N/A | N/A | N/A | N/A | N/A | N/A | 371.326 | 1.000 | 371.326 |
| 31 | 328.919 | 12.753 | 22 | N/A | N/A | N/A | N/A | N/A | N/A | N/A | N/A | 328.919 | 1.000 | 328.919 |
| 39 | 312.943 | 12.757 | 22 | N/A | N/A | N/A | N/A | N/A | N/A | N/A | N/A | 312.943 | 1.000 | 312.943 |
| 70 | 287.888 | 12.760 | 22 | N/A | N/A | N/A | N/A | N/A | N/A | N/A | N/A | 287.888 | 1.000 | 287.888 |
| 92 | 313.078 | 6.460 | 22 | N/A | N/A | N/A | N/A | N/A | N/A | N/A | N/A | 313.078 | 1.000 | 313.078 |
| 187 | 312.943 | 12.757 | 22 | N/A | N/A | N/A | N/A | N/A | N/A | N/A | N/A | 312.943 | 1.000 | 312.943 |
| 196 | 313.078 | 6.532 | 22 | N/A | N/A | N/A | N/A | N/A | N/A | N/A | N/A | 313.078 | 1.000 | 313.078 |
| 255 | 313.235 | 8.431 | 22 | N/A | N/A | N/A | N/A | N/A | N/A | N/A | N/A | 313.235 | 1.000 | 313.235 |
| 558 | 313.235 | 8.075 | 22 | N/A | N/A | N/A | N/A | N/A | N/A | N/A | N/A | 313.235 | 1.000 | 313.235 |
| 197 | 272.258 | 8.673 | 21 | N/A | N/A | N/A | N/A | N/A | N/A | N/A | N/A | 272.258 | 1.000 | 272.258 |
| 223 | 300.290 | 9.476 | 21 | N/A | N/A | N/A | N/A | N/A | N/A | N/A | N/A | 300.290 | 1.000 | 300.290 |

| Feature identifiers | | | | Library Hits | | | | | | | | | | |
|---|---|---|---|---|---|---|---|---|---|---|---|---|---|---|
| ClusterIdx | m/z | RT [min] | Subnetwork | Library | Instrument | Ion source | Ion mode | Library m/z | MasDiff | Cosine | PPM | Parent mass | Precursor charge | Precursor mass |
| 460 | 406.354 | 10.232 | 21 | N/A | N/A | N/A | N/A | N/A | N/A | N/A | N/A | 406.354 | 1.000 | 406.354 |
| 157 | 429.146 | 4.153 | 20 | N/A | N/A | N/A | N/A | N/A | N/A | N/A | N/A | 429.146 | 1.000 | 429.146 |
| 166 | 549.571 | 3.384 | 20 | N/A | N/A | N/A | N/A | N/A | N/A | N/A | N/A | 1098.130 | 2.000 | 549.567 |
| 481 | 549.078 | 3.211 | 20 | N/A | N/A | N/A | N/A | N/A | N/A | N/A | N/A | 1097.150 | 2.000 | 549.075 |
| 520 | 432.026 | 3.379 | 20 | N/A | N/A | N/A | N/A | N/A | N/A | N/A | N/A | 863.044 | 2.000 | 432.022 |
| 584 | 432.026 | 3.502 | 20 | N/A | N/A | N/A | N/A | N/A | N/A | N/A | N/A | 863.044 | 2.000 | 432.022 |
| 596 | 405.571 | 3.038 | 20 | N/A | N/A | N/A | N/A | N/A | N/A | N/A | N/A | 810.133 | 2.000 | 405.567 |
| 600 | 359.104 | 0.595 | 20 | N/A | N/A | N/A | N/A | N/A | N/A | N/A | N/A | 359.104 | 1.000 | 359.104 |
| 611 | 401.115 | 2.454 | 20 | N/A | N/A | N/A | N/A | N/A | N/A | N/A | N/A | 401.115 | 1.000 | 401.115 |
| 622 | 411.021 | 3.222 | 20 | N/A | N/A | N/A | N/A | N/A | N/A | N/A | N/A | 821.034 | 2.000 | 411.017 |
| 315 | 515.353 | 9.549 | 19 | N/A | N/A | N/A | N/A | N/A | N/A | N/A | N/A | 515.353 | 1.000 | 515.353 |
| 326 | 321.148 | 3.682 | 19 | N/A | N/A | N/A | N/A | N/A | N/A | N/A | N/A | 321.148 | 1.000 | 321.148 |
| 595 | 366.206 | 3.688 | 19 | N/A | N/A | N/A | N/A | N/A | N/A | N/A | N/A | 366.206 | 1.000 | 366.206 |
| 494 | 453.189 | 7.153 | 18 | N/A | N/A | N/A | N/A | N/A | N/A | N/A | N/A | 453.189 | 1.000 | 453.189 |
| 591 | 453.189 | 7.367 | 18 | N/A | N/A | N/A | N/A | N/A | N/A | N/A | N/A | 453.189 | 1.000 | 453.189 |
| 861 | 453.187 | 7.297 | 18 | N/A | N/A | N/A | N/A | N/A | N/A | N/A | N/A | 453.187 | 1.000 | 453.187 |
| 21 | 453.169 | 7.511 | 17 | N/A | N/A | N/A | N/A | N/A | N/A | N/A | N/A | 453.169 | 1.000 | 453.169 |
| 292 | 453.168 | 7.525 | 17 | N/A | N/A | N/A | N/A | N/A | N/A | N/A | N/A | 453.168 | 1.000 | 453.168 |
| 640 | 453.186 | 7.468 | 17 | N/A | N/A | N/A | N/A | N/A | N/A | N/A | N/A | 453.186 | 1.000 | 453.186 |
| 59 | 281.138 | 7.525 | 16 | N/A | N/A | N/A | N/A | N/A | N/A | N/A | N/A | 281.138 | 1.000 | 281.138 |
| 97 | 299.149 | 4.461 | 16 | N/A | N/A | N/A | N/A | N/A | N/A | N/A | N/A | 299.149 | 1.000 | 299.149 |
| 172 | 299.148 | 7.523 | 16 | N/A | N/A | N/A | N/A | N/A | N/A | N/A | N/A | 299.148 | 1.000 | 299.148 |
| 296 | 344.206 | 4.461 | 16 | N/A | N/A | N/A | N/A | N/A | N/A | N/A | N/A | 344.206 | 1.000 | 344.206 |
| 381 | 316.176 | 4.464 | 16 | N/A | N/A | N/A | N/A | N/A | N/A | N/A | N/A | 316.176 | 1.000 | 316.176 |
| 42 | 473.318 | 10.647 | 15 | N/A | N/A | N/A | N/A | N/A | N/A | N/A | N/A | 473.318 | 1.000 | 473.318 |
| 72 | 680.481 | 12.424 | 15 | N/A | N/A | N/A | N/A | N/A | N/A | N/A | N/A | 680.481 | 1.000 | 680.481 |
| 74 | 475.298 | 9.832 | 15 | N/A | N/A | N/A | N/A | N/A | N/A | N/A | N/A | 475.298 | 1.000 | 475.298 |
| 95 | 492.324 | 9.832 | 15 | N/A | N/A | N/A | N/A | N/A | N/A | N/A | N/A | 492.324 | 1.000 | 492.324 |
| 96 | 654.333 | 10.021 | 15 | N/A | N/A | N/A | N/A | N/A | N/A | N/A | N/A | 654.333 | 1.000 | 654.333 |
| 105 | 490.345 | 10.647 | 15 | N/A | N/A | N/A | N/A | N/A | N/A | N/A | N/A | 490.345 | 1.000 | 490.345 |
| 129 | 663.454 | 12.425 | 15 | N/A | N/A | N/A | N/A | N/A | N/A | N/A | N/A | 663.454 | 1.000 | 663.454 |
| 141 | 708.511 | 12.424 | 15 | N/A | N/A | N/A | N/A | N/A | N/A | N/A | N/A | 708.511 | 1.000 | 708.511 |
| 181 | 637.306 | 10.020 | 15 | N/A | N/A | N/A | N/A | N/A | N/A | N/A | N/A | 637.306 | 1.000 | 637.306 |
| 205 | 518.376 | 10.647 | 15 | N/A | N/A | N/A | N/A | N/A | N/A | N/A | N/A | 518.376 | 1.000 | 518.376 |
| 273 | 287.141 | 6.170 | 15 | N/A | N/A | N/A | N/A | N/A | N/A | N/A | N/A | 287.141 | 1.000 | 287.141 |
| 298 | 515.329 | 10.526 | 15 | N/A | N/A | N/A | N/A | N/A | N/A | N/A | N/A | 515.329 | 1.000 | 515.329 |
| 311 | 474.317 | 10.257 | 15 | N/A | N/A | N/A | N/A | N/A | N/A | N/A | N/A | 474.317 | 1.000 | 474.317 |
| 317 | 457.288 | 10.396 | 15 | N/A | N/A | N/A | N/A | N/A | N/A | N/A | N/A | 457.288 | 1.000 | 457.288 |
| 318 | 387.194 | 7.399 | 15 | N/A | N/A | N/A | N/A | N/A | N/A | N/A | N/A | 387.194 | 1.000 | 387.194 |
| 426 | 682.364 | 10.020 | 15 | N/A | N/A | N/A | N/A | N/A | N/A | N/A | N/A | 682.364 | 1.000 | 682.364 |
| 433 | 532.356 | 10.526 | 15 | N/A | N/A | N/A | N/A | N/A | N/A | N/A | N/A | 532.356 | 1.000 | 532.356 |
| 452 | 519.377 | 10.257 | 15 | N/A | N/A | N/A | N/A | N/A | N/A | N/A | N/A | 519.377 | 1.000 | 519.377 |
| 529 | 520.355 | 9.833 | 15 | N/A | N/A | N/A | N/A | N/A | N/A | N/A | N/A | 520.355 | 1.000 | 520.355 |
| 726 | 1342.926 | 12.423 | 15 | N/A | N/A | N/A | N/A | N/A | N/A | N/A | N/A | 1342.930 | 1.000 | 1342.930 |
| 755 | 949.587 | 9.831 | 15 | N/A | N/A | N/A | N/A | N/A | N/A | N/A | N/A | 949.587 | 1.000 | 949.587 |
| 783 | 491.374 | 10.264 | 15 | N/A | N/A | N/A | N/A | N/A | N/A | N/A | N/A | 981.740 | 2.000 | 491.370 |
| 849 | 1370.957 | 12.421 | 15 | N/A | N/A | N/A | N/A | N/A | N/A | N/A | N/A | 1370.960 | 1.000 | 1370.960 |
| 585 | 834.093 | 3.504 | 14 | N/A | N/A | N/A | N/A | N/A | N/A | N/A | N/A | 834.093 | 1.000 | 834.093 |
| 619 | 792.082 | 3.197 | 14 | N/A | N/A | N/A | N/A | N/A | N/A | N/A | N/A | 792.082 | 1.000 | 792.082 |
| 714 | 542.297 | 4.609 | 13 | N/A | N/A | N/A | N/A | N/A | N/A | N/A | N/A | 1083.590 | 2.000 | 542.293 |
| 717 | 492.763 | 4.165 | 13 | N/A | N/A | N/A | N/A | N/A | N/A | N/A | N/A | 984.519 | 2.000 | 492.759 |
| 51 | 247.034 | 5.943 | 12 | N/A | N/A | N/A | N/A | N/A | N/A | N/A | N/A | 247.034 | 1.000 | 247.034 |
| 698 | 247.034 | 6.796 | 12 | N/A | N/A | N/A | N/A | N/A | N/A | N/A | N/A | 247.034 | 1.000 | 247.034 |
| 718 | 291.023 | 6.253 | 12 | N/A | N/A | N/A | N/A | N/A | N/A | N/A | N/A | 291.023 | 1.000 | 291.023 |
| 16 | 122.547 | 12.762 | 11 | N/A | N/A | N/A | N/A | N/A | N/A | N/A | N/A | 244.087 | 2.000 | 122.543 |
| 60 | 122.547 | 14.272 | 11 | N/A | N/A | N/A | N/A | N/A | N/A | N/A | N/A | 244.087 | 2.000 | 122.543 |
| 103 | 122.547 | 2.944 | 11 | N/A | N/A | N/A | N/A | N/A | N/A | N/A | N/A | 244.087 | 2.000 | 122.543 |
| 113 | 122.547 | 2.472 | 11 | N/A | N/A | N/A | N/A | N/A | N/A | N/A | N/A | 244.087 | 2.000 | 122.543 |
| 115 | 122.547 | 13.880 | 11 | N/A | N/A | N/A | N/A | N/A | N/A | N/A | N/A | 244.087 | 2.000 | 122.543 |
| 120 | 122.547 | 2.832 | 11 | N/A | N/A | N/A | N/A | N/A | N/A | N/A | N/A | 244.087 | 2.000 | 122.543 |
| 123 | 122.547 | 13.067 | 11 | N/A | N/A | N/A | N/A | N/A | N/A | N/A | N/A | 244.087 | 2.000 | 122.543 |
| 125 | 122.547 | 2.720 | 11 | N/A | N/A | N/A | N/A | N/A | N/A | N/A | N/A | 244.087 | 2.000 | 122.543 |
| 127 | 122.547 | 0.660 | 11 | N/A | N/A | N/A | N/A | N/A | N/A | N/A | N/A | 244.087 | 2.000 | 122.543 |
| 135 | 122.547 | 13.843 | 11 | N/A | N/A | N/A | N/A | N/A | N/A | N/A | N/A | 244.087 | 2.000 | 122.543 |
| 140 | 122.547 | 2.622 | 11 | N/A | N/A | N/A | N/A | N/A | N/A | N/A | N/A | 244.087 | 2.000 | 122.543 |
| 145 | 122.547 | 2.598 | 11 | N/A | N/A | N/A | N/A | N/A | N/A | N/A | N/A | 244.087 | 2.000 | 122.543 |
| 147 | 122.547 | 2.749 | 11 | N/A | N/A | N/A | N/A | N/A | N/A | N/A | N/A | 244.087 | 2.000 | 122.543 |
| 151 | 122.547 | 13.172 | 11 | N/A | N/A | N/A | N/A | N/A | N/A | N/A | N/A | 244.087 | 2.000 | 122.543 |
| 156 | 122.547 | 2.528 | 11 | N/A | N/A | N/A | N/A | N/A | N/A | N/A | N/A | 244.087 | 2.000 | 122.543 |
| 160 | 122.547 | 2.262 | 11 | N/A | N/A | N/A | N/A | N/A | N/A | N/A | N/A | 244.087 | 2.000 | 122.543 |

| Feature identifiers | | | | Library Hits | | | | | | | | | | |
|---|---|---|---|---|---|---|---|---|---|---|---|---|---|---|
| ClusterIdx | m/z | RT [min] | Subnetwork | Library | Instrument | Ion source | Ion mode | Library m/z | MasDiff | Cosine | PPM | Parent mass | Precursor charge | Precursor mass |
| 169 | 122.547 | 13.121 | 11 | N/A | N/A | N/A | N/A | N/A | N/A | N/A | N/A | 244.087 | 2.000 | 122.543 |
| 174 | 122.547 | 2.436 | 11 | N/A | N/A | N/A | N/A | N/A | N/A | N/A | N/A | 244.087 | 2.000 | 122.543 |
| 177 | 122.547 | 13.217 | 11 | N/A | N/A | N/A | N/A | N/A | N/A | N/A | N/A | 244.087 | 2.000 | 122.543 |
| 178 | 122.547 | 2.294 | 11 | N/A | N/A | N/A | N/A | N/A | N/A | N/A | N/A | 244.087 | 2.000 | 122.543 |
| 183 | 122.547 | 1.983 | 11 | N/A | N/A | N/A | N/A | N/A | N/A | N/A | N/A | 244.087 | 2.000 | 122.543 |
| 185 | 122.547 | 13.651 | 11 | N/A | N/A | N/A | N/A | N/A | N/A | N/A | N/A | 244.087 | 2.000 | 122.543 |
| 189 | 122.547 | 0.617 | 11 | N/A | N/A | N/A | N/A | N/A | N/A | N/A | N/A | 244.087 | 2.000 | 122.543 |
| 206 | 122.547 | 1.515 | 11 | N/A | N/A | N/A | N/A | N/A | N/A | N/A | N/A | 244.087 | 2.000 | 122.543 |
| 207 | 122.547 | 2.189 | 11 | N/A | N/A | N/A | N/A | N/A | N/A | N/A | N/A | 244.087 | 2.000 | 122.543 |
| 216 | 122.547 | 1.421 | 11 | N/A | N/A | N/A | N/A | N/A | N/A | N/A | N/A | 244.087 | 2.000 | 122.543 |
| 220 | 122.547 | 2.347 | 11 | N/A | N/A | N/A | N/A | N/A | N/A | N/A | N/A | 244.087 | 2.000 | 122.543 |
| 248 | 122.547 | 2.158 | 11 | N/A | N/A | N/A | N/A | N/A | N/A | N/A | N/A | 244.087 | 2.000 | 122.543 |
| 587 | 122.547 | 2.681 | 11 | N/A | N/A | N/A | N/A | N/A | N/A | N/A | N/A | 244.087 | 2.000 | 122.543 |
| 644 | 122.547 | 2.919 | 11 | N/A | N/A | N/A | N/A | N/A | N/A | N/A | N/A | 244.087 | 2.000 | 122.543 |
| 645 | 122.547 | 2.389 | 11 | N/A | N/A | N/A | N/A | N/A | N/A | N/A | N/A | 244.087 | 2.000 | 122.543 |
| 656 | 122.547 | 2.584 | 11 | N/A | N/A | N/A | N/A | N/A | N/A | N/A | N/A | 244.087 | 2.000 | 122.543 |
| 657 | 122.547 | 2.792 | 11 | N/A | N/A | N/A | N/A | N/A | N/A | N/A | N/A | 244.087 | 2.000 | 122.543 |
| 710 | 122.547 | 3.154 | 11 | N/A | N/A | N/A | N/A | N/A | N/A | N/A | N/A | 244.086 | 2.000 | 122.543 |
| 715 | 122.547 | 3.094 | 11 | N/A | N/A | N/A | N/A | N/A | N/A | N/A | N/A | 244.086 | 2.000 | 122.543 |
| 719 | 122.547 | 2.222 | 11 | N/A | N/A | N/A | N/A | N/A | N/A | N/A | N/A | 244.087 | 2.000 | 122.543 |
| 773 | 122.547 | 13.630 | 11 | N/A | N/A | N/A | N/A | N/A | N/A | N/A | N/A | 244.087 | 2.000 | 122.543 |
| 61 | 537.536 | 11.439 | 10 | N/A | N/A | N/A | N/A | N/A | N/A | N/A | N/A | 537.536 | 1.000 | 537.536 |
| 109 | 565.567 | 11.404 | 10 | N/A | N/A | N/A | N/A | N/A | N/A | N/A | N/A | 565.567 | 1.000 | 565.567 |
| 295 | 582.594 | 11.445 | 10 | N/A | N/A | N/A | N/A | N/A | N/A | N/A | N/A | 582.594 | 1.000 | 582.594 |
| 816 | 593.598 | 10.672 | 10 | N/A | N/A | N/A | N/A | N/A | N/A | N/A | N/A | 593.598 | 1.000 | 593.598 |
| 191 | 608.393 | 9.449 | 9 | N/A | N/A | N/A | N/A | N/A | N/A | N/A | N/A | 608.393 | 1.000 | 608.393 |
| 313 | 638.404 | 9.642 | 9 | N/A | N/A | N/A | N/A | N/A | N/A | N/A | N/A | 638.404 | 1.000 | 638.404 |
| 348 | 624.388 | 9.076 | 9 | N/A | N/A | N/A | N/A | N/A | N/A | N/A | N/A | 624.388 | 1.000 | 624.388 |
| 15 | 824.976 | 3.461 | 8 | N/A | N/A | N/A | N/A | N/A | N/A | N/A | N/A | 2472.910 | 3.000 | 824.971 |
| 18 | 810.973 | 3.183 | 8 | N/A | N/A | N/A | N/A | N/A | N/A | N/A | N/A | 2430.900 | 3.000 | 810.968 |
| 26 | 838.980 | 3.758 | 8 | N/A | N/A | N/A | N/A | N/A | N/A | N/A | N/A | 2514.920 | 3.000 | 838.974 |
| 65 | 1215.954 | 3.181 | 8 | N/A | N/A | N/A | N/A | N/A | N/A | N/A | N/A | 2430.900 | 2.000 | 1215.950 |
| 87 | 1236.959 | 3.462 | 8 | N/A | N/A | N/A | N/A | N/A | N/A | N/A | N/A | 2472.910 | 2.000 | 1236.960 |
| 121 | 837.625 | 3.464 | 8 | N/A | N/A | N/A | N/A | N/A | N/A | N/A | N/A | 2510.860 | 3.000 | 837.620 |
| 136 | 823.621 | 3.183 | 8 | N/A | N/A | N/A | N/A | N/A | N/A | N/A | N/A | 2468.850 | 3.000 | 823.616 |
| 194 | 618.984 | 3.459 | 8 | N/A | N/A | N/A | N/A | N/A | N/A | N/A | N/A | 2472.910 | 4.000 | 618.978 |
| 204 | 1257.965 | 3.758 | 8 | N/A | N/A | N/A | N/A | N/A | N/A | N/A | N/A | 2514.920 | 2.000 | 1257.960 |
| 240 | 852.983 | 4.057 | 8 | N/A | N/A | N/A | N/A | N/A | N/A | N/A | N/A | 2556.930 | 3.000 | 852.978 |
| 244 | 846.306 | 3.758 | 8 | N/A | N/A | N/A | N/A | N/A | N/A | N/A | N/A | 2536.900 | 3.000 | 846.301 |
| 253 | 1069.394 | 4.071 | 8 | N/A | N/A | N/A | N/A | N/A | N/A | N/A | N/A | 3206.170 | 3.000 | 1069.390 |
| 257 | 851.628 | 3.759 | 8 | N/A | N/A | N/A | N/A | N/A | N/A | N/A | N/A | 2552.870 | 3.000 | 851.623 |
| 258 | 608.481 | 3.183 | 8 | N/A | N/A | N/A | N/A | N/A | N/A | N/A | N/A | 2430.900 | 4.000 | 608.475 |
| 288 | 1083.397 | 4.311 | 8 | N/A | N/A | N/A | N/A | N/A | N/A | N/A | N/A | 3248.180 | 3.000 | 1083.390 |
| 289 | 629.487 | 3.760 | 8 | N/A | N/A | N/A | N/A | N/A | N/A | N/A | N/A | 2514.920 | 4.000 | 629.481 |
| 312 | 1055.391 | 3.820 | 8 | N/A | N/A | N/A | N/A | N/A | N/A | N/A | N/A | 3164.160 | 3.000 | 1055.390 |
| 352 | 1234.935 | 3.184 | 8 | N/A | N/A | N/A | N/A | N/A | N/A | N/A | N/A | 2468.860 | 2.000 | 1234.930 |
| 406 | 802.298 | 4.071 | 8 | N/A | N/A | N/A | N/A | N/A | N/A | N/A | N/A | 3206.170 | 4.000 | 802.293 |
| 445 | 812.801 | 4.312 | 8 | N/A | N/A | N/A | N/A | N/A | N/A | N/A | N/A | 3248.180 | 4.000 | 812.795 |
| 450 | 1041.387 | 3.604 | 8 | N/A | N/A | N/A | N/A | N/A | N/A | N/A | N/A | 3122.150 | 3.000 | 1041.380 |
| 457 | 1255.942 | 3.463 | 8 | N/A | N/A | N/A | N/A | N/A | N/A | N/A | N/A | 2510.880 | 2.000 | 1255.940 |
| 497 | 1278.970 | 4.056 | 8 | N/A | N/A | N/A | N/A | N/A | N/A | N/A | N/A | 2556.930 | 2.000 | 1278.970 |
| 504 | 1268.955 | 3.761 | 8 | N/A | N/A | N/A | N/A | N/A | N/A | N/A | N/A | 2536.900 | 2.000 | 1268.950 |
| 511 | 865.632 | 4.058 | 8 | N/A | N/A | N/A | N/A | N/A | N/A | N/A | N/A | 2594.880 | 3.000 | 865.627 |
| 514 | 639.989 | 4.058 | 8 | N/A | N/A | N/A | N/A | N/A | N/A | N/A | N/A | 2556.930 | 4.000 | 639.983 |
| 515 | 860.310 | 4.057 | 8 | N/A | N/A | N/A | N/A | N/A | N/A | N/A | N/A | 2578.920 | 3.000 | 860.305 |
| 518 | 1276.945 | 3.762 | 8 | N/A | N/A | N/A | N/A | N/A | N/A | N/A | N/A | 2552.880 | 2.000 | 1276.940 |
| 519 | 856.952 | 3.759 | 8 | N/A | N/A | N/A | N/A | N/A | N/A | N/A | N/A | 2568.840 | 3.000 | 856.947 |
| 522 | 1090.726 | 4.311 | 8 | N/A | N/A | N/A | N/A | N/A | N/A | N/A | N/A | 3270.160 | 3.000 | 1090.720 |
| 531 | 1076.722 | 4.075 | 8 | N/A | N/A | N/A | N/A | N/A | N/A | N/A | N/A | 3228.150 | 3.000 | 1076.720 |
| 533 | 1096.048 | 4.311 | 8 | N/A | N/A | N/A | N/A | N/A | N/A | N/A | N/A | 3286.130 | 3.000 | 1096.040 |
| 534 | 791.796 | 3.823 | 8 | N/A | N/A | N/A | N/A | N/A | N/A | N/A | N/A | 3164.160 | 4.000 | 791.790 |
| 537 | 810.973 | 3.375 | 8 | N/A | N/A | N/A | N/A | N/A | N/A | N/A | N/A | 2430.900 | 3.000 | 810.968 |
| 559 | 824.976 | 3.689 | 8 | N/A | N/A | N/A | N/A | N/A | N/A | N/A | N/A | 2472.910 | 3.000 | 824.971 |
| 561 | 1289.962 | 4.060 | 8 | N/A | N/A | N/A | N/A | N/A | N/A | N/A | N/A | 2578.920 | 2.000 | 1289.960 |
| 567 | 1082.043 | 4.072 | 8 | N/A | N/A | N/A | N/A | N/A | N/A | N/A | N/A | 3244.110 | 3.000 | 1082.040 |
| 588 | 1027.384 | 3.427 | 8 | N/A | N/A | N/A | N/A | N/A | N/A | N/A | N/A | 3080.140 | 3.000 | 1027.380 |
| 590 | 1013.379 | 3.335 | 8 | N/A | N/A | N/A | N/A | N/A | N/A | N/A | N/A | 3038.120 | 3.000 | 1013.370 |
| 610 | 770.790 | 3.427 | 8 | N/A | N/A | N/A | N/A | N/A | N/A | N/A | N/A | 3080.140 | 4.000 | 770.784 |
| 614 | 837.291 | 3.461 | 8 | N/A | N/A | N/A | N/A | N/A | N/A | N/A | N/A | 2509.860 | 3.000 | 837.285 |
| 620 | 760.288 | 3.338 | 8 | N/A | N/A | N/A | N/A | N/A | N/A | N/A | N/A | 3038.130 | 4.000 | 760.282 |

| Feature identifiers | | | | Library Hits | | | | | | | | | | |
|---|---|---|---|---|---|---|---|---|---|---|---|---|---|---|
| ClusterIdx | m/z | RT [min] | Subnetwork | Library | Instrument | Ion source | Ion mode | Library m/z | MasDiff | Cosine | PPM | Parent mass | Precursor charge | Precursor mass |
| 633 | 1062.719 | 3.822 | 8 | N/A | N/A | N/A | N/A | N/A | N/A | N/A | N/A | 3186.140 | 3.000 | 1062.710 |
| 650 | 629.236 | 3.761 | 8 | N/A | N/A | N/A | N/A | N/A | N/A | N/A | N/A | 2513.920 | 4.000 | 629.230 |
| 659 | 813.049 | 4.312 | 8 | N/A | N/A | N/A | N/A | N/A | N/A | N/A | N/A | 3249.170 | 4.000 | 813.043 |
| 664 | 866.987 | 4.353 | 8 | N/A | N/A | N/A | N/A | N/A | N/A | N/A | N/A | 2598.940 | 3.000 | 866.981 |
| 672 | 1299.976 | 4.354 | 8 | N/A | N/A | N/A | N/A | N/A | N/A | N/A | N/A | 2598.940 | 2.000 | 1299.970 |
| 676 | 874.315 | 4.356 | 8 | N/A | N/A | N/A | N/A | N/A | N/A | N/A | N/A | 2620.930 | 3.000 | 874.310 |
| 677 | 856.617 | 3.758 | 8 | N/A | N/A | N/A | N/A | N/A | N/A | N/A | N/A | 2567.840 | 3.000 | 856.612 |
| 678 | 879.636 | 4.362 | 8 | N/A | N/A | N/A | N/A | N/A | N/A | N/A | N/A | 2636.890 | 3.000 | 879.631 |
| 680 | 1297.948 | 4.058 | 8 | N/A | N/A | N/A | N/A | N/A | N/A | N/A | N/A | 2594.890 | 2.000 | 1297.940 |
| 683 | 975.108 | 4.244 | 8 | N/A | N/A | N/A | N/A | N/A | N/A | N/A | N/A | 3897.410 | 4.000 | 975.102 |
| 685 | 1101.371 | 4.312 | 8 | N/A | N/A | N/A | N/A | N/A | N/A | N/A | N/A | 3302.100 | 3.000 | 1101.370 |
| 691 | 822.287 | 4.312 | 8 | N/A | N/A | N/A | N/A | N/A | N/A | N/A | N/A | 3286.130 | 4.000 | 822.282 |
| 730 | 796.969 | 3.070 | 8 | N/A | N/A | N/A | N/A | N/A | N/A | N/A | N/A | 2388.890 | 3.000 | 796.964 |
| 741 | 1068.040 | 3.824 | 8 | N/A | N/A | N/A | N/A | N/A | N/A | N/A | N/A | 3202.100 | 3.000 | 1068.030 |
| 746 | 1069.395 | 4.241 | 8 | N/A | N/A | N/A | N/A | N/A | N/A | N/A | N/A | 3206.170 | 3.000 | 1069.390 |
| 751 | 1221.446 | 3.186 | 8 | N/A | N/A | N/A | N/A | N/A | N/A | N/A | N/A | 4882.760 | 4.000 | 1221.440 |
| 757 | 1055.390 | 3.992 | 8 | N/A | N/A | N/A | N/A | N/A | N/A | N/A | N/A | 3164.160 | 3.000 | 1055.390 |
| 760 | 975.359 | 4.241 | 8 | N/A | N/A | N/A | N/A | N/A | N/A | N/A | N/A | 3898.410 | 4.000 | 975.353 |
| 761 | 802.298 | 4.239 | 8 | N/A | N/A | N/A | N/A | N/A | N/A | N/A | N/A | 3206.170 | 4.000 | 802.292 |
| 766 | 618.733 | 3.459 | 8 | N/A | N/A | N/A | N/A | N/A | N/A | N/A | N/A | 2471.910 | 4.000 | 618.727 |
| 768 | 838.979 | 3.988 | 8 | N/A | N/A | N/A | N/A | N/A | N/A | N/A | N/A | 2514.920 | 3.000 | 838.974 |
| 778 | 796.636 | 3.069 | 8 | N/A | N/A | N/A | N/A | N/A | N/A | N/A | N/A | 2387.890 | 3.000 | 796.630 |
| 785 | 1215.954 | 3.392 | 8 | N/A | N/A | N/A | N/A | N/A | N/A | N/A | N/A | 2430.900 | 2.000 | 1215.950 |
| 787 | 608.481 | 3.384 | 8 | N/A | N/A | N/A | N/A | N/A | N/A | N/A | N/A | 2430.900 | 4.000 | 608.475 |
| 788 | 1236.958 | 3.690 | 8 | N/A | N/A | N/A | N/A | N/A | N/A | N/A | N/A | 2472.910 | 2.000 | 1236.950 |
| 789 | 618.734 | 3.694 | 8 | N/A | N/A | N/A | N/A | N/A | N/A | N/A | N/A | 2471.910 | 4.000 | 618.728 |
| 791 | 792.044 | 3.972 | 8 | N/A | N/A | N/A | N/A | N/A | N/A | N/A | N/A | 3165.150 | 4.000 | 792.038 |
| 796 | 1069.727 | 4.238 | 8 | N/A | N/A | N/A | N/A | N/A | N/A | N/A | N/A | 3207.170 | 3.000 | 1069.720 |
| 800 | 824.643 | 3.674 | 8 | N/A | N/A | N/A | N/A | N/A | N/A | N/A | N/A | 2471.910 | 3.000 | 824.638 |
| 801 | 1055.390 | 4.161 | 8 | N/A | N/A | N/A | N/A | N/A | N/A | N/A | N/A | 3164.150 | 3.000 | 1055.380 |
| 803 | 791.795 | 4.163 | 8 | N/A | N/A | N/A | N/A | N/A | N/A | N/A | N/A | 3164.160 | 4.000 | 791.789 |
| 804 | 1300.145 | 4.240 | 8 | N/A | N/A | N/A | N/A | N/A | N/A | N/A | N/A | 3898.420 | 3.000 | 1300.140 |
| 805 | 810.973 | 3.576 | 8 | N/A | N/A | N/A | N/A | N/A | N/A | N/A | N/A | 2430.900 | 3.000 | 810.968 |
| 807 | 1082.043 | 4.246 | 8 | N/A | N/A | N/A | N/A | N/A | N/A | N/A | N/A | 3244.110 | 3.000 | 1082.040 |
| 809 | 629.487 | 3.997 | 8 | N/A | N/A | N/A | N/A | N/A | N/A | N/A | N/A | 2514.920 | 4.000 | 629.481 |
| 812 | 1268.461 | 3.759 | 8 | N/A | N/A | N/A | N/A | N/A | N/A | N/A | N/A | 2535.910 | 2.000 | 1268.460 |
| 820 | 1300.134 | 4.241 | 8 | N/A | N/A | N/A | N/A | N/A | N/A | N/A | N/A | 3898.390 | 3.000 | 1300.130 |
| 829 | 1137.663 | 4.195 | 8 | N/A | N/A | N/A | N/A | N/A | N/A | N/A | N/A | 4547.630 | 4.000 | 1137.660 |
| 830 | 1263.457 | 3.755 | 8 | N/A | N/A | N/A | N/A | N/A | N/A | N/A | N/A | 5050.810 | 4.000 | 1263.450 |
| 833 | 1076.722 | 4.246 | 8 | N/A | N/A | N/A | N/A | N/A | N/A | N/A | N/A | 3228.150 | 3.000 | 1076.720 |
| 836 | 1063.052 | 3.825 | 8 | N/A | N/A | N/A | N/A | N/A | N/A | N/A | N/A | 3187.140 | 3.000 | 1063.050 |
| 838 | 791.796 | 3.996 | 8 | N/A | N/A | N/A | N/A | N/A | N/A | N/A | N/A | 3164.160 | 4.000 | 791.790 |
| 842 | 822.537 | 4.313 | 8 | N/A | N/A | N/A | N/A | N/A | N/A | N/A | N/A | 3287.120 | 4.000 | 822.531 |
| 845 | 1257.962 | 3.997 | 8 | N/A | N/A | N/A | N/A | N/A | N/A | N/A | N/A | 2514.920 | 2.000 | 1257.960 |
| 847 | 781.292 | 3.601 | 8 | N/A | N/A | N/A | N/A | N/A | N/A | N/A | N/A | 3122.140 | 4.000 | 781.286 |
| 848 | 811.784 | 4.077 | 8 | N/A | N/A | N/A | N/A | N/A | N/A | N/A | N/A | 3244.110 | 4.000 | 811.778 |
| 851 | 1276.945 | 3.763 | 8 | N/A | N/A | N/A | N/A | N/A | N/A | N/A | N/A | 2552.880 | 2.000 | 1276.940 |
| 854 | 1194.947 | 3.069 | 8 | N/A | N/A | N/A | N/A | N/A | N/A | N/A | N/A | 2388.890 | 2.000 | 1194.940 |
| 862 | 964.605 | 4.154 | 8 | N/A | N/A | N/A | N/A | N/A | N/A | N/A | N/A | 3855.400 | 4.000 | 964.599 |
| 354 | 828.611 | 3.182 | 7 | N/A | N/A | N/A | N/A | N/A | N/A | N/A | N/A | 2483.820 | 3.000 | 828.605 |
| 360 | 842.615 | 3.460 | 7 | N/A | N/A | N/A | N/A | N/A | N/A | N/A | N/A | 2525.830 | 3.000 | 842.609 |
| 649 | 842.948 | 3.459 | 7 | N/A | N/A | N/A | N/A | N/A | N/A | N/A | N/A | 2526.830 | 3.000 | 842.943 |
| 811 | 828.945 | 3.185 | 7 | N/A | N/A | N/A | N/A | N/A | N/A | N/A | N/A | 2484.820 | 3.000 | 828.940 |
| 22 | 274.274 | 6.821 | 6 | N/A | N/A | N/A | N/A | N/A | N/A | N/A | N/A | 274.274 | 1.000 | 274.274 |
| 104 | 302.305 | 7.664 | 6 | N/A | N/A | N/A | N/A | N/A | N/A | N/A | N/A | 302.305 | 1.000 | 302.305 |
| 277 | 330.337 | 8.495 | 6 | N/A | N/A | N/A | N/A | N/A | N/A | N/A | N/A | 330.337 | 1.000 | 330.337 |
| 341 | 358.368 | 9.237 | 6 | N/A | N/A | N/A | N/A | N/A | N/A | N/A | N/A | 358.368 | 1.000 | 358.368 |
| 321 | 311.185 | 7.063 | 5 | N/A | N/A | N/A | N/A | N/A | N/A | N/A | N/A | 311.185 | 1.000 | 311.185 |
| 347 | 423.310 | 10.200 | 5 | N/A | N/A | N/A | N/A | N/A | N/A | N/A | N/A | 423.310 | 1.000 | 423.310 |
| 661 | 468.369 | 10.166 | 5 | N/A | N/A | N/A | N/A | N/A | N/A | N/A | N/A | 468.369 | 1.000 | 468.369 |
| 11 | 452.277 | 7.678 | 4 | N/A | N/A | N/A | N/A | N/A | N/A | N/A | N/A | 452.277 | 1.000 | 452.277 |
| 155 | 474.259 | 7.679 | 4 | N/A | N/A | N/A | N/A | N/A | N/A | N/A | N/A | 474.259 | 1.000 | 474.259 |
| 238 | 474.258 | 7.808 | 4 | N/A | N/A | N/A | N/A | N/A | N/A | N/A | N/A | 474.258 | 1.000 | 474.258 |
| 350 | 690.508 | 12.705 | 4 | N/A | N/A | N/A | N/A | N/A | N/A | N/A | N/A | 690.508 | 1.000 | 690.508 |
| 424 | 511.304 | 8.562 | 4 | N/A | N/A | N/A | N/A | N/A | N/A | N/A | N/A | 511.304 | 1.000 | 511.304 |
| 486 | 690.508 | 10.348 | 4 | N/A | N/A | N/A | N/A | N/A | N/A | N/A | N/A | 690.508 | 1.000 | 690.508 |
| 489 | 712.490 | 12.701 | 4 | N/A | N/A | N/A | N/A | N/A | N/A | N/A | N/A | 712.490 | 1.000 | 712.490 |
| 505 | 712.490 | 10.356 | 4 | N/A | N/A | N/A | N/A | N/A | N/A | N/A | N/A | 712.490 | 1.000 | 712.490 |
| 510 | 485.288 | 8.510 | 4 | N/A | N/A | N/A | N/A | N/A | N/A | N/A | N/A | 485.288 | 1.000 | 485.288 |
| 513 | 743.484 | 12.700 | 4 | N/A | N/A | N/A | N/A | N/A | N/A | N/A | N/A | 743.484 | 1.000 | 743.484 |

| Feature identifiers | | | | Library Hits | | | | | | | | | | |
|---|---|---|---|---|---|---|---|---|---|---|---|---|---|---|
| ClusterIdx | m/z | RT [min] | Subnetwork | Library | Instrument | Ion source | Ion mode | Library m/z | MasDiff | Cosine | PPM | Parent mass | Precursor charge | Precursor mass |
| 516 | 483.272 | 7.795 | 4 | N/A | N/A | N/A | N/A | N/A | N/A | N/A | N/A | 483.272 | 1.000 | 483.272 |
| 517 | 740.522 | 12.702 | 4 | N/A | N/A | N/A | N/A | N/A | N/A | N/A | N/A | 740.522 | 1.000 | 740.522 |
| 532 | 738.526 | 10.287 | 4 | N/A | N/A | N/A | N/A | N/A | N/A | N/A | N/A | 738.526 | 1.000 | 738.526 |
| 554 | 903.547 | 7.681 | 4 | N/A | N/A | N/A | N/A | N/A | N/A | N/A | N/A | 903.547 | 1.000 | 903.547 |
| 555 | 467.277 | 8.509 | 4 | N/A | N/A | N/A | N/A | N/A | N/A | N/A | N/A | 467.277 | 1.000 | 467.277 |
| 562 | 743.484 | 10.306 | 4 | N/A | N/A | N/A | N/A | N/A | N/A | N/A | N/A | 743.484 | 1.000 | 743.484 |
| 565 | 496.304 | 8.835 | 4 | N/A | N/A | N/A | N/A | N/A | N/A | N/A | N/A | 496.304 | 1.000 | 496.304 |
| 577 | 483.272 | 7.944 | 4 | N/A | N/A | N/A | N/A | N/A | N/A | N/A | N/A | 483.272 | 1.000 | 483.272 |
| 581 | 511.304 | 8.710 | 4 | N/A | N/A | N/A | N/A | N/A | N/A | N/A | N/A | 511.304 | 1.000 | 511.304 |
| 583 | 452.277 | 7.806 | 4 | N/A | N/A | N/A | N/A | N/A | N/A | N/A | N/A | 452.277 | 1.000 | 452.277 |
| 639 | 690.508 | 10.220 | 4 | N/A | N/A | N/A | N/A | N/A | N/A | N/A | N/A | 690.508 | 1.000 | 690.508 |
| 643 | 712.490 | 10.268 | 4 | N/A | N/A | N/A | N/A | N/A | N/A | N/A | N/A | 712.490 | 1.000 | 712.490 |
| 669 | 466.293 | 8.072 | 4 | N/A | N/A | N/A | N/A | N/A | N/A | N/A | N/A | 466.293 | 1.000 | 466.293 |
| 679 | 466.293 | 8.206 | 4 | N/A | N/A | N/A | N/A | N/A | N/A | N/A | N/A | 466.293 | 1.000 | 466.293 |
| 701 | 440.278 | 7.956 | 4 | N/A | N/A | N/A | N/A | N/A | N/A | N/A | N/A | 440.278 | 1.000 | 440.278 |
| 713 | 662.477 | 12.666 | 4 | N/A | N/A | N/A | N/A | N/A | N/A | N/A | N/A | 662.477 | 1.000 | 662.477 |
| 721 | 533.285 | 8.561 | 4 | N/A | N/A | N/A | N/A | N/A | N/A | N/A | N/A | 533.285 | 1.000 | 533.285 |
| 722 | 440.278 | 7.818 | 4 | N/A | N/A | N/A | N/A | N/A | N/A | N/A | N/A | 440.278 | 1.000 | 440.278 |
| 724 | 684.458 | 12.702 | 4 | N/A | N/A | N/A | N/A | N/A | N/A | N/A | N/A | 684.458 | 1.000 | 684.458 |
| 727 | 465.262 | 7.796 | 4 | N/A | N/A | N/A | N/A | N/A | N/A | N/A | N/A | 465.262 | 1.000 | 465.262 |
| 728 | 505.254 | 7.795 | 4 | N/A | N/A | N/A | N/A | N/A | N/A | N/A | N/A | 505.254 | 1.000 | 505.254 |
| 729 | 507.269 | 8.510 | 4 | N/A | N/A | N/A | N/A | N/A | N/A | N/A | N/A | 507.269 | 1.000 | 507.269 |
| 731 | 493.293 | 8.564 | 4 | N/A | N/A | N/A | N/A | N/A | N/A | N/A | N/A | 493.293 | 1.000 | 493.293 |
| 736 | 676.492 | 12.707 | 4 | N/A | N/A | N/A | N/A | N/A | N/A | N/A | N/A | 676.492 | 1.000 | 676.492 |
| 745 | 493.293 | 8.710 | 4 | N/A | N/A | N/A | N/A | N/A | N/A | N/A | N/A | 493.293 | 1.000 | 493.293 |
| 749 | 500.300 | 7.793 | 4 | N/A | N/A | N/A | N/A | N/A | N/A | N/A | N/A | 500.300 | 1.000 | 500.300 |
| 756 | 533.285 | 8.715 | 4 | N/A | N/A | N/A | N/A | N/A | N/A | N/A | N/A | 533.285 | 1.000 | 533.285 |
| 775 | 704.523 | 12.703 | 4 | N/A | N/A | N/A | N/A | N/A | N/A | N/A | N/A | 704.523 | 1.000 | 704.523 |
| 779 | 704.523 | 10.367 | 4 | N/A | N/A | N/A | N/A | N/A | N/A | N/A | N/A | 704.523 | 1.000 | 704.523 |
| 781 | 726.506 | 12.701 | 4 | N/A | N/A | N/A | N/A | N/A | N/A | N/A | N/A | 726.505 | 1.000 | 726.505 |
| 813 | 494.288 | 8.224 | 4 | N/A | N/A | N/A | N/A | N/A | N/A | N/A | N/A | 494.288 | 1.000 | 494.288 |
| 825 | 664.490 | 9.773 | 4 | N/A | N/A | N/A | N/A | N/A | N/A | N/A | N/A | 664.490 | 1.000 | 664.490 |
| 837 | 766.559 | 12.703 | 4 | N/A | N/A | N/A | N/A | N/A | N/A | N/A | N/A | 766.559 | 1.000 | 766.559 |
| 841 | 676.491 | 10.424 | 4 | N/A | N/A | N/A | N/A | N/A | N/A | N/A | N/A | 676.491 | 1.000 | 676.491 |
| 844 | 698.475 | 10.420 | 4 | N/A | N/A | N/A | N/A | N/A | N/A | N/A | N/A | 698.475 | 1.000 | 698.475 |
| 853 | 1380.008 | 10.186 | 4 | N/A | N/A | N/A | N/A | N/A | N/A | N/A | N/A | 1380.010 | 1.000 | 1380.010 |
| 855 | 1380.006 | 9.800 | 4 | N/A | N/A | N/A | N/A | N/A | N/A | N/A | N/A | 1380.010 | 1.000 | 1380.010 |
| 27 | 316.212 | 3.277 | 3 | N/A | N/A | N/A | N/A | N/A | N/A | N/A | N/A | 316.212 | 1.000 | 316.212 |
| 37 | 430.244 | 4.459 | 3 | N/A | N/A | N/A | N/A | N/A | N/A | N/A | N/A | 430.244 | 1.000 | 430.244 |
| 55 | 300.200 | 3.938 | 3 | N/A | N/A | N/A | N/A | N/A | N/A | N/A | N/A | 599.393 | 2.000 | 300.197 |
| 69 | 416.228 | 3.930 | 3 | N/A | N/A | N/A | N/A | N/A | N/A | N/A | N/A | 416.228 | 1.000 | 416.228 |
| 122 | 243.183 | 3.050 | 3 | N/A | N/A | N/A | N/A | N/A | N/A | N/A | N/A | 485.358 | 2.000 | 243.179 |
| 124 | 334.178 | 3.942 | 3 | N/A | N/A | N/A | N/A | N/A | N/A | N/A | N/A | 334.178 | 1.000 | 334.178 |
| 128 | 357.214 | 4.538 | 3 | N/A | N/A | N/A | N/A | N/A | N/A | N/A | N/A | 713.421 | 2.000 | 357.211 |
| 149 | 350.208 | 4.208 | 3 | N/A | N/A | N/A | N/A | N/A | N/A | N/A | N/A | 699.407 | 2.000 | 350.204 |
| 167 | 599.391 | 3.938 | 3 | N/A | N/A | N/A | N/A | N/A | N/A | N/A | N/A | 599.391 | 1.000 | 599.391 |
| 190 | 294.861 | 4.052 | 3 | N/A | N/A | N/A | N/A | N/A | N/A | N/A | N/A | 882.567 | 3.000 | 294.856 |
| 224 | 302.196 | 0.594 | 3 | N/A | N/A | N/A | N/A | N/A | N/A | N/A | N/A | 302.196 | 1.000 | 302.196 |
| 246 | 309.181 | 4.412 | 3 | N/A | N/A | N/A | N/A | N/A | N/A | N/A | N/A | 617.355 | 2.000 | 309.178 |
| 268 | 328.200 | 4.325 | 3 | N/A | N/A | N/A | N/A | N/A | N/A | N/A | N/A | 982.583 | 3.000 | 328.194 |
| 290 | 272.185 | 3.424 | 3 | N/A | N/A | N/A | N/A | N/A | N/A | N/A | N/A | 272.185 | 1.000 | 272.185 |
| 293 | 293.191 | 3.577 | 3 | N/A | N/A | N/A | N/A | N/A | N/A | N/A | N/A | 585.373 | 2.000 | 293.187 |
| 306 | 300.180 | 5.792 | 3 | N/A | N/A | N/A | N/A | N/A | N/A | N/A | N/A | 300.180 | 1.000 | 300.180 |
| 322 | 713.423 | 4.538 | 3 | N/A | N/A | N/A | N/A | N/A | N/A | N/A | N/A | 713.423 | 1.000 | 713.423 |
| 328 | 252.166 | 3.641 | 3 | N/A | N/A | N/A | N/A | N/A | N/A | N/A | N/A | 503.324 | 2.000 | 252.162 |
| 329 | 332.872 | 4.600 | 3 | N/A | N/A | N/A | N/A | N/A | N/A | N/A | N/A | 996.599 | 3.000 | 332.866 |
| 343 | 471.307 | 6.113 | 3 | N/A | N/A | N/A | N/A | N/A | N/A | N/A | N/A | 471.307 | 1.000 | 471.307 |
| 361 | 441.788 | 4.079 | 3 | N/A | N/A | N/A | N/A | N/A | N/A | N/A | N/A | 882.568 | 2.000 | 441.784 |
| 386 | 241.204 | 5.244 | 3 | N/A | N/A | N/A | N/A | N/A | N/A | N/A | N/A | 481.400 | 2.000 | 241.200 |
| 396 | 469.328 | 4.897 | 3 | N/A | N/A | N/A | N/A | N/A | N/A | N/A | N/A | 469.328 | 1.000 | 469.328 |
| 401 | 256.850 | 3.715 | 3 | N/A | N/A | N/A | N/A | N/A | N/A | N/A | N/A | 768.536 | 3.000 | 256.845 |
| 404 | 699.407 | 4.204 | 3 | N/A | N/A | N/A | N/A | N/A | N/A | N/A | N/A | 699.407 | 1.000 | 699.407 |
| 413 | 292.192 | 4.464 | 3 | N/A | N/A | N/A | N/A | N/A | N/A | N/A | N/A | 1165.740 | 4.000 | 292.186 |
| 439 | 585.375 | 3.578 | 3 | N/A | N/A | N/A | N/A | N/A | N/A | N/A | N/A | 585.375 | 1.000 | 585.375 |
| 443 | 384.773 | 3.708 | 3 | N/A | N/A | N/A | N/A | N/A | N/A | N/A | N/A | 768.537 | 2.000 | 384.769 |
| 463 | 293.191 | 3.406 | 3 | N/A | N/A | N/A | N/A | N/A | N/A | N/A | N/A | 585.373 | 2.000 | 293.187 |
| 499 | 599.391 | 3.791 | 3 | N/A | N/A | N/A | N/A | N/A | N/A | N/A | N/A | 599.391 | 1.000 | 599.391 |
| 526 | 485.359 | 3.054 | 3 | N/A | N/A | N/A | N/A | N/A | N/A | N/A | N/A | 485.359 | 1.000 | 485.359 |
| 527 | 441.788 | 4.168 | 3 | N/A | N/A | N/A | N/A | N/A | N/A | N/A | N/A | 882.568 | 2.000 | 441.784 |
| 530 | 290.193 | 3.898 | 3 | N/A | N/A | N/A | N/A | N/A | N/A | N/A | N/A | 868.563 | 3.000 | 290.188 |

| Feature identifiers | | | | Library Hits | | | | | | | | | | |
|---|---|---|---|---|---|---|---|---|---|---|---|---|---|---|
| ClusterIdx | m/z | RT [min] | Subnetwork | Library | Instrument | Ion source | Ion mode | Library m/z | MasDiff | Cosine | PPM | Parent mass | Precursor charge | Precursor mass |
| 553 | 332.207 | 4.839 | 3 | N/A | N/A | N/A | N/A | N/A | N/A | N/A | N/A | 332.207 | 1.000 | 332.207 |
| 568 | 379.210 | 4.343 | 3 | N/A | N/A | N/A | N/A | N/A | N/A | N/A | N/A | 757.412 | 2.000 | 379.206 |
| 623 | 343.200 | 3.890 | 3 | N/A | N/A | N/A | N/A | N/A | N/A | N/A | N/A | 685.391 | 2.000 | 343.196 |
| 624 | 263.684 | 4.125 | 3 | N/A | N/A | N/A | N/A | N/A | N/A | N/A | N/A | 1051.710 | 4.000 | 263.678 |
| 627 | 230.153 | 3.714 | 3 | N/A | N/A | N/A | N/A | N/A | N/A | N/A | N/A | 459.299 | 2.000 | 230.150 |
| 629 | 347.535 | 4.414 | 3 | N/A | N/A | N/A | N/A | N/A | N/A | N/A | N/A | 1040.590 | 3.000 | 347.530 |
| 747 | 540.426 | 9.375 | 3 | N/A | N/A | N/A | N/A | N/A | N/A | N/A | N/A | 540.426 | 1.000 | 540.426 |
| 750 | 568.458 | 9.844 | 3 | N/A | N/A | N/A | N/A | N/A | N/A | N/A | N/A | 568.458 | 1.000 | 568.458 |
| 758 | 263.684 | 4.034 | 3 | N/A | N/A | N/A | N/A | N/A | N/A | N/A | N/A | 1051.710 | 4.000 | 263.678 |
| 782 | 220.150 | 0.536 | 3 | N/A | N/A | N/A | N/A | N/A | N/A | N/A | N/A | 220.150 | 1.000 | 220.150 |
| 784 | 498.804 | 4.596 | 3 | N/A | N/A | N/A | N/A | N/A | N/A | N/A | N/A | 996.601 | 2.000 | 498.800 |
| 810 | 294.861 | 4.145 | 3 | N/A | N/A | N/A | N/A | N/A | N/A | N/A | N/A | 882.566 | 3.000 | 294.855 |
| 817 | 384.772 | 3.711 | 3 | N/A | N/A | N/A | N/A | N/A | N/A | N/A | N/A | 768.537 | 2.000 | 384.769 |
| 819 | 292.192 | 4.644 | 3 | N/A | N/A | N/A | N/A | N/A | N/A | N/A | N/A | 1165.740 | 4.000 | 292.186 |
| 93 | 256.264 | 9.789 | 2 | N/A | N/A | N/A | N/A | N/A | N/A | N/A | N/A | 256.264 | 1.000 | 256.264 |
| 142 | 284.295 | 10.325 | 2 | N/A | N/A | N/A | N/A | N/A | N/A | N/A | N/A | 284.295 | 1.000 | 284.295 |
| 175 | 308.279 | 8.430 | 2 | N/A | N/A | N/A | N/A | N/A | N/A | N/A | N/A | 308.279 | 1.000 | 308.279 |
| 218 | 482.408 | 9.562 | 2 | N/A | N/A | N/A | N/A | N/A | N/A | N/A | N/A | 482.408 | 1.000 | 482.408 |
| 226 | 526.434 | 9.538 | 2 | N/A | N/A | N/A | N/A | N/A | N/A | N/A | N/A | 526.434 | 1.000 | 526.434 |
| 228 | 336.311 | 9.373 | 2 | N/A | N/A | N/A | N/A | N/A | N/A | N/A | N/A | 336.311 | 1.000 | 336.311 |
| 229 | 291.253 | 8.431 | 2 | N/A | N/A | N/A | N/A | N/A | N/A | N/A | N/A | 291.253 | 1.000 | 291.253 |
| 241 | 424.364 | 9.342 | 2 | N/A | N/A | N/A | N/A | N/A | N/A | N/A | N/A | 424.364 | 1.000 | 424.364 |
| 242 | 380.338 | 9.364 | 2 | N/A | N/A | N/A | N/A | N/A | N/A | N/A | N/A | 380.338 | 1.000 | 380.338 |
| 247 | 352.306 | 8.443 | 2 | N/A | N/A | N/A | N/A | N/A | N/A | N/A | N/A | 352.306 | 1.000 | 352.306 |
| 269 | 438.382 | 9.584 | 2 | N/A | N/A | N/A | N/A | N/A | N/A | N/A | N/A | 438.382 | 1.000 | 438.382 |
| 270 | 408.372 | 10.006 | 2 | N/A | N/A | N/A | N/A | N/A | N/A | N/A | N/A | 408.372 | 1.000 | 408.372 |
| 276 | 570.460 | 9.514 | 2 | N/A | N/A | N/A | N/A | N/A | N/A | N/A | N/A | 570.460 | 1.000 | 570.460 |
| 278 | 512.417 | 9.299 | 2 | N/A | N/A | N/A | N/A | N/A | N/A | N/A | N/A | 512.417 | 1.000 | 512.417 |
| 280 | 468.390 | 9.320 | 2 | N/A | N/A | N/A | N/A | N/A | N/A | N/A | N/A | 468.390 | 1.000 | 468.390 |
| 291 | 366.322 | 8.959 | 2 | N/A | N/A | N/A | N/A | N/A | N/A | N/A | N/A | 366.322 | 1.000 | 366.322 |
| 294 | 452.398 | 9.985 | 2 | N/A | N/A | N/A | N/A | N/A | N/A | N/A | N/A | 452.397 | 1.000 | 452.397 |
| 297 | 556.443 | 9.273 | 2 | N/A | N/A | N/A | N/A | N/A | N/A | N/A | N/A | 556.443 | 1.000 | 556.443 |
| 302 | 247.227 | 8.431 | 2 | N/A | N/A | N/A | N/A | N/A | N/A | N/A | N/A | 247.227 | 1.000 | 247.227 |
| 303 | 392.338 | 10.385 | 2 | N/A | N/A | N/A | N/A | N/A | N/A | N/A | N/A | 392.338 | 1.000 | 392.338 |
| 310 | 496.424 | 9.963 | 2 | N/A | N/A | N/A | N/A | N/A | N/A | N/A | N/A | 496.424 | 1.000 | 496.424 |
| 336 | 364.344 | 10.020 | 2 | N/A | N/A | N/A | N/A | N/A | N/A | N/A | N/A | 364.344 | 1.000 | 364.344 |
| 337 | 540.449 | 9.937 | 2 | N/A | N/A | N/A | N/A | N/A | N/A | N/A | N/A | 540.449 | 1.000 | 540.449 |
| 344 | 476.301 | 4.018 | 2 | N/A | N/A | N/A | N/A | N/A | N/A | N/A | N/A | 476.301 | 1.000 | 476.301 |
| 349 | 440.359 | 8.424 | 2 | N/A | N/A | N/A | N/A | N/A | N/A | N/A | N/A | 440.359 | 1.000 | 440.359 |
| 359 | 600.469 | 9.251 | 2 | N/A | N/A | N/A | N/A | N/A | N/A | N/A | N/A | 600.469 | 1.000 | 600.469 |
| 362 | 322.295 | 8.962 | 2 | N/A | N/A | N/A | N/A | N/A | N/A | N/A | N/A | 322.295 | 1.000 | 322.295 |
| 364 | 614.485 | 9.493 | 2 | N/A | N/A | N/A | N/A | N/A | N/A | N/A | N/A | 614.485 | 1.000 | 614.485 |
| 366 | 396.333 | 8.432 | 2 | N/A | N/A | N/A | N/A | N/A | N/A | N/A | N/A | 396.333 | 1.000 | 396.333 |
| 367 | 422.389 | 10.267 | 2 | N/A | N/A | N/A | N/A | N/A | N/A | N/A | N/A | 422.389 | 1.000 | 422.389 |
| 369 | 436.364 | 10.348 | 2 | N/A | N/A | N/A | N/A | N/A | N/A | N/A | N/A | 436.364 | 1.000 | 436.364 |
| 377 | 422.347 | 10.141 | 2 | N/A | N/A | N/A | N/A | N/A | N/A | N/A | N/A | 422.347 | 1.000 | 422.347 |
| 382 | 466.373 | 10.105 | 2 | N/A | N/A | N/A | N/A | N/A | N/A | N/A | N/A | 466.373 | 1.000 | 466.373 |
| 383 | 319.284 | 9.373 | 2 | N/A | N/A | N/A | N/A | N/A | N/A | N/A | N/A | 319.284 | 1.000 | 319.284 |
| 387 | 466.415 | 10.248 | 2 | N/A | N/A | N/A | N/A | N/A | N/A | N/A | N/A | 466.415 | 1.000 | 466.415 |
| 389 | 526.434 | 9.632 | 2 | N/A | N/A | N/A | N/A | N/A | N/A | N/A | N/A | 526.434 | 1.000 | 526.434 |
| 390 | 468.390 | 9.206 | 2 | N/A | N/A | N/A | N/A | N/A | N/A | N/A | N/A | 468.390 | 1.000 | 468.390 |
| 399 | 520.331 | 4.142 | 2 | N/A | N/A | N/A | N/A | N/A | N/A | N/A | N/A | 520.331 | 1.000 | 520.331 |
| 400 | 484.385 | 8.410 | 2 | N/A | N/A | N/A | N/A | N/A | N/A | N/A | N/A | 484.385 | 1.000 | 484.385 |
| 412 | 512.416 | 9.184 | 2 | N/A | N/A | N/A | N/A | N/A | N/A | N/A | N/A | 512.417 | 1.000 | 512.417 |
| 414 | 528.411 | 8.398 | 2 | N/A | N/A | N/A | N/A | N/A | N/A | N/A | N/A | 528.411 | 1.000 | 528.411 |
| 416 | 564.360 | 4.248 | 2 | N/A | N/A | N/A | N/A | N/A | N/A | N/A | N/A | 564.359 | 1.000 | 564.359 |
| 417 | 644.495 | 9.227 | 2 | N/A | N/A | N/A | N/A | N/A | N/A | N/A | N/A | 644.495 | 1.000 | 644.495 |
| 422 | 480.390 | 10.316 | 2 | N/A | N/A | N/A | N/A | N/A | N/A | N/A | N/A | 480.390 | 1.000 | 480.390 |
| 432 | 556.443 | 9.161 | 2 | N/A | N/A | N/A | N/A | N/A | N/A | N/A | N/A | 556.443 | 1.000 | 556.443 |
| 438 | 584.475 | 9.912 | 2 | N/A | N/A | N/A | N/A | N/A | N/A | N/A | N/A | 584.475 | 1.000 | 584.475 |
| 455 | 628.501 | 9.881 | 2 | N/A | N/A | N/A | N/A | N/A | N/A | N/A | N/A | 628.501 | 1.000 | 628.501 |
| 464 | 608.392 | 4.346 | 2 | N/A | N/A | N/A | N/A | N/A | N/A | N/A | N/A | 608.392 | 1.000 | 608.392 |
| 471 | 352.306 | 8.095 | 2 | N/A | N/A | N/A | N/A | N/A | N/A | N/A | N/A | 352.306 | 1.000 | 352.306 |
| 528 | 308.279 | 8.075 | 2 | N/A | N/A | N/A | N/A | N/A | N/A | N/A | N/A | 308.279 | 1.000 | 308.279 |
| 539 | 688.521 | 9.203 | 2 | N/A | N/A | N/A | N/A | N/A | N/A | N/A | N/A | 688.521 | 1.000 | 688.521 |
| 541 | 363.310 | 9.363 | 2 | N/A | N/A | N/A | N/A | N/A | N/A | N/A | N/A | 363.310 | 1.000 | 363.310 |
| 546 | 291.253 | 8.075 | 2 | N/A | N/A | N/A | N/A | N/A | N/A | N/A | N/A | 291.253 | 1.000 | 291.253 |
| 548 | 396.333 | 8.097 | 2 | N/A | N/A | N/A | N/A | N/A | N/A | N/A | N/A | 396.333 | 1.000 | 396.333 |
| 556 | 732.548 | 9.180 | 2 | N/A | N/A | N/A | N/A | N/A | N/A | N/A | N/A | 732.548 | 1.000 | 732.548 |
| 564 | 440.359 | 8.096 | 2 | N/A | N/A | N/A | N/A | N/A | N/A | N/A | N/A | 440.359 | 1.000 | 440.359 |

| Feature identifiers | | | | Library Hits | | | | | | | | | | |
|---|---|---|---|---|---|---|---|---|---|---|---|---|---|---|
| ClusterIdx | m/z | RT [min] | Subnetwork | Library | Instrument | Ion source | Ion mode | Library m/z | MasDiff | Cosine | PPM | Parent mass | Precursor charge | Precursor mass |
| 570 | 484.385 | 8.094 | 2 | N/A | N/A | N/A | N/A | N/A | N/A | N/A | N/A | 484.385 | 1.000 | 484.385 |
| 571 | 308.279 | 8.290 | 2 | N/A | N/A | N/A | N/A | N/A | N/A | N/A | N/A | 308.279 | 1.000 | 308.279 |
| 573 | 528.411 | 8.088 | 2 | N/A | N/A | N/A | N/A | N/A | N/A | N/A | N/A | 528.411 | 1.000 | 528.411 |
| 576 | 247.227 | 8.064 | 2 | N/A | N/A | N/A | N/A | N/A | N/A | N/A | N/A | 247.227 | 1.000 | 247.227 |
| 579 | 335.280 | 8.098 | 2 | N/A | N/A | N/A | N/A | N/A | N/A | N/A | N/A | 335.280 | 1.000 | 335.280 |
| 580 | 572.437 | 8.084 | 2 | N/A | N/A | N/A | N/A | N/A | N/A | N/A | N/A | 572.437 | 1.000 | 572.437 |
| 598 | 396.333 | 8.315 | 2 | N/A | N/A | N/A | N/A | N/A | N/A | N/A | N/A | 396.333 | 1.000 | 396.333 |
| 604 | 440.359 | 8.302 | 2 | N/A | N/A | N/A | N/A | N/A | N/A | N/A | N/A | 440.359 | 1.000 | 440.359 |
| 608 | 672.527 | 9.854 | 2 | N/A | N/A | N/A | N/A | N/A | N/A | N/A | N/A | 672.527 | 1.000 | 672.527 |
| 615 | 350.328 | 9.687 | 2 | N/A | N/A | N/A | N/A | N/A | N/A | N/A | N/A | 350.328 | 1.000 | 350.328 |
| 618 | 658.511 | 9.464 | 2 | N/A | N/A | N/A | N/A | N/A | N/A | N/A | N/A | 658.511 | 1.000 | 658.511 |
| 625 | 702.536 | 9.449 | 2 | N/A | N/A | N/A | N/A | N/A | N/A | N/A | N/A | 702.536 | 1.000 | 702.536 |
| 647 | 394.355 | 9.697 | 2 | N/A | N/A | N/A | N/A | N/A | N/A | N/A | N/A | 394.355 | 1.000 | 394.355 |
| 50 | 701.620 | 10.613 | 1 | N/A | N/A | N/A | N/A | N/A | N/A | N/A | N/A | 701.620 | 1.000 | 701.620 |
| 199 | 419.364 | 7.499 | 1 | N/A | N/A | N/A | N/A | N/A | N/A | N/A | N/A | 419.364 | 1.000 | 419.364 |
| 689 | 685.624 | 12.053 | 1 | N/A | N/A | N/A | N/A | N/A | N/A | N/A | N/A | 685.624 | 1.000 | 685.624 |
| 1 | 110.020 | 12.761 | -1 | N/A | N/A | N/A | N/A | N/A | N/A | N/A | N/A | 219.032 | 2.000 | 110.016 |
| 8 | 437.194 | 7.524 | -1 | N/A | N/A | N/A | N/A | N/A | N/A | N/A | N/A | 437.194 | 1.000 | 437.194 |
| 10 | 120.081 | 0.519 | -1 | N/A | N/A | N/A | N/A | N/A | N/A | N/A | N/A | 120.081 | 1.000 | 120.081 |
| 13 | 242.284 | 8.651 | -1 | N/A | N/A | N/A | N/A | N/A | N/A | N/A | N/A | 242.284 | 1.000 | 242.284 |
| 14 | 851.398 | 7.527 | -1 | N/A | N/A | N/A | N/A | N/A | N/A | N/A | N/A | 851.398 | 1.000 | 851.398 |
| 32 | 437.194 | 7.249 | -1 | N/A | N/A | N/A | N/A | N/A | N/A | N/A | N/A | 437.194 | 1.000 | 437.194 |
| 33 | 559.518 | 11.442 | -1 | N/A | N/A | N/A | N/A | N/A | N/A | N/A | N/A | 559.518 | 1.000 | 559.518 |
| 34 | 867.372 | 7.528 | -1 | N/A | N/A | N/A | N/A | N/A | N/A | N/A | N/A | 867.372 | 1.000 | 867.372 |
| 41 | 587.549 | 11.407 | -1 | N/A | N/A | N/A | N/A | N/A | N/A | N/A | N/A | 587.549 | 1.000 | 587.549 |
| 47 | 163.991 | 12.748 | -1 | N/A | N/A | N/A | N/A | N/A | N/A | N/A | N/A | 163.991 | 1.000 | 163.991 |
| 53 | 243.088 | 4.621 | -1 | N/A | N/A | N/A | N/A | N/A | N/A | N/A | N/A | 243.088 | 1.000 | 243.088 |
| 54 | 399.209 | 10.100 | -1 | N/A | N/A | N/A | N/A | N/A | N/A | N/A | N/A | 399.209 | 1.000 | 399.209 |
| 57 | 685.436 | 12.425 | -1 | N/A | N/A | N/A | N/A | N/A | N/A | N/A | N/A | 685.436 | 1.000 | 685.436 |
| 58 | 250.112 | 5.753 | -1 | N/A | N/A | N/A | N/A | N/A | N/A | N/A | N/A | 250.112 | 1.000 | 250.112 |
| 64 | 255.073 | 4.542 | -1 | N/A | N/A | N/A | N/A | N/A | N/A | N/A | N/A | 255.073 | 1.000 | 255.073 |
| 66 | 261.109 | 5.951 | -1 | N/A | N/A | N/A | N/A | N/A | N/A | N/A | N/A | 261.109 | 1.000 | 261.109 |
| 71 | 267.077 | 3.607 | -1 | N/A | N/A | N/A | N/A | N/A | N/A | N/A | N/A | 267.077 | 1.000 | 267.077 |
| 73 | 453.168 | 7.250 | -1 | N/A | N/A | N/A | N/A | N/A | N/A | N/A | N/A | 453.168 | 1.000 | 453.168 |
| 76 | 324.142 | 7.584 | -1 | N/A | N/A | N/A | N/A | N/A | N/A | N/A | N/A | 324.142 | 1.000 | 324.142 |
| 77 | 372.882 | 12.723 | -1 | N/A | N/A | N/A | N/A | N/A | N/A | N/A | N/A | 372.882 | 1.000 | 372.882 |
| 79 | 321.131 | 4.460 | -1 | N/A | N/A | N/A | N/A | N/A | N/A | N/A | N/A | 321.131 | 1.000 | 321.131 |
| 82 | 269.015 | 5.941 | -1 | N/A | N/A | N/A | N/A | N/A | N/A | N/A | N/A | 269.015 | 1.000 | 269.015 |
| 83 | 230.141 | 12.721 | -1 | N/A | N/A | N/A | N/A | N/A | N/A | N/A | N/A | 230.141 | 1.000 | 230.141 |
| 85 | 233.079 | 4.825 | -1 | N/A | N/A | N/A | N/A | N/A | N/A | N/A | N/A | 233.079 | 1.000 | 233.079 |
| 86 | 239.089 | 4.007 | -1 | N/A | N/A | N/A | N/A | N/A | N/A | N/A | N/A | 239.089 | 1.000 | 239.089 |
| 90 | 332.913 | 12.737 | -1 | N/A | N/A | N/A | N/A | N/A | N/A | N/A | N/A | 332.913 | 1.000 | 332.913 |
| 98 | 195.102 | 7.496 | -1 | N/A | N/A | N/A | N/A | N/A | N/A | N/A | N/A | 195.102 | 1.000 | 195.102 |
| 100 | 440.099 | 7.644 | -1 | N/A | N/A | N/A | N/A | N/A | N/A | N/A | N/A | 440.099 | 1.000 | 440.099 |
| 101 | 235.169 | 8.489 | -1 | N/A | N/A | N/A | N/A | N/A | N/A | N/A | N/A | 235.169 | 1.000 | 235.169 |
| 106 | 670.153 | 5.750 | -1 | N/A | N/A | N/A | N/A | N/A | N/A | N/A | N/A | 670.153 | 1.000 | 670.153 |
| 107 | 183.150 | 4.641 | -1 | N/A | N/A | N/A | N/A | N/A | N/A | N/A | N/A | 183.150 | 1.000 | 183.150 |
| 108 | 281.088 | 8.127 | -1 | N/A | N/A | N/A | N/A | N/A | N/A | N/A | N/A | 281.088 | 1.000 | 281.088 |
| 111 | 262.165 | 4.000 | -1 | N/A | N/A | N/A | N/A | N/A | N/A | N/A | N/A | 262.165 | 1.000 | 262.165 |
| 114 | 244.191 | 5.975 | -1 | N/A | N/A | N/A | N/A | N/A | N/A | N/A | N/A | 244.191 | 1.000 | 244.191 |
| 116 | 228.232 | 9.984 | -1 | N/A | N/A | N/A | N/A | N/A | N/A | N/A | N/A | 228.232 | 1.000 | 228.232 |
| 118 | 219.175 | 8.820 | -1 | N/A | N/A | N/A | N/A | N/A | N/A | N/A | N/A | 219.175 | 1.000 | 219.175 |
| 126 | 296.219 | 7.149 | -1 | N/A | N/A | N/A | N/A | N/A | N/A | N/A | N/A | 296.219 | 1.000 | 296.219 |
| 137 | 431.183 | 10.115 | -1 | N/A | N/A | N/A | N/A | N/A | N/A | N/A | N/A | 431.184 | 1.000 | 431.184 |
| 148 | 319.187 | 7.588 | -1 | N/A | N/A | N/A | N/A | N/A | N/A | N/A | N/A | 319.187 | 1.000 | 319.187 |
| 152 | 707.875 | 4.385 | -1 | N/A | N/A | N/A | N/A | N/A | N/A | N/A | N/A | 1414.740 | 2.000 | 707.871 |
| 162 | 174.999 | 12.765 | -1 | N/A | N/A | N/A | N/A | N/A | N/A | N/A | N/A | 348.991 | 2.000 | 174.995 |
| 164 | 338.147 | 5.707 | -1 | N/A | N/A | N/A | N/A | N/A | N/A | N/A | N/A | 338.147 | 1.000 | 338.147 |
| 165 | 269.210 | 8.432 | -1 | N/A | N/A | N/A | N/A | N/A | N/A | N/A | N/A | 269.210 | 1.000 | 269.210 |
| 173 | 451.246 | 9.774 | -1 | N/A | N/A | N/A | N/A | N/A | N/A | N/A | N/A | 451.246 | 1.000 | 451.246 |
| 179 | 723.608 | 10.615 | -1 | N/A | N/A | N/A | N/A | N/A | N/A | N/A | N/A | 723.608 | 1.000 | 723.608 |
| 184 | 434.186 | 7.527 | -1 | N/A | N/A | N/A | N/A | N/A | N/A | N/A | N/A | 867.364 | 2.000 | 434.182 |
| 188 | 270.279 | 10.247 | -1 | N/A | N/A | N/A | N/A | N/A | N/A | N/A | N/A | 270.279 | 1.000 | 270.279 |
| 193 | 242.284 | 7.258 | -1 | N/A | N/A | N/A | N/A | N/A | N/A | N/A | N/A | 242.284 | 1.000 | 242.284 |
| 195 | 312.326 | 10.525 | -1 | N/A | N/A | N/A | N/A | N/A | N/A | N/A | N/A | 312.326 | 1.000 | 312.326 |
| 198 | 499.165 | 7.526 | -1 | N/A | N/A | N/A | N/A | N/A | N/A | N/A | N/A | 499.165 | 1.000 | 499.165 |
| 203 | 122.547 | 1.761 | -1 | N/A | N/A | N/A | N/A | N/A | N/A | N/A | N/A | 244.087 | 2.000 | 122.543 |
| 212 | 409.201 | 10.114 | -1 | N/A | N/A | N/A | N/A | N/A | N/A | N/A | N/A | 409.202 | 1.000 | 409.202 |
| 214 | 270.279 | 9.930 | -1 | N/A | N/A | N/A | N/A | N/A | N/A | N/A | N/A | 270.279 | 1.000 | 270.279 |
| 217 | 266.248 | 6.949 | -1 | N/A | N/A | N/A | N/A | N/A | N/A | N/A | N/A | 266.248 | 1.000 | 266.248 |

| Feature identifiers | | | | Library Hits | | | | | | | | | | |
|---|---|---|---|---|---|---|---|---|---|---|---|---|---|---|
| ClusterIdx | m/z | RT [min] | Subnetwork | Library | Instrument | Ion source | Ion mode | Library m/z | MasDiff | Cosine | PPM | Parent mass | Precursor charge | Precursor mass |
| 222 | 495.300 | 10.647 | -1 | N/A | N/A | N/A | N/A | N/A | N/A | N/A | N/A | 495.300 | 1.000 | 495.300 |
| 225 | 330.300 | 9.151 | -1 | N/A | N/A | N/A | N/A | N/A | N/A | N/A | N/A | 330.300 | 1.000 | 330.300 |
| 232 | 250.178 | 6.460 | -1 | N/A | N/A | N/A | N/A | N/A | N/A | N/A | N/A | 250.178 | 1.000 | 250.178 |
| 234 | 555.250 | 4.014 | -1 | N/A | N/A | N/A | N/A | N/A | N/A | N/A | N/A | 1109.490 | 2.000 | 555.246 |
| 236 | 149.988 | 12.784 | -1 | N/A | N/A | N/A | N/A | N/A | N/A | N/A | N/A | 149.988 | 1.000 | 149.988 |
| 237 | 640.348 | 4.374 | -1 | N/A | N/A | N/A | N/A | N/A | N/A | N/A | N/A | 1279.690 | 2.000 | 640.344 |
| 252 | 157.084 | 0.657 | -1 | N/A | N/A | N/A | N/A | N/A | N/A | N/A | N/A | 157.084 | 1.000 | 157.084 |
| 254 | 111.039 | 13.492 | -1 | N/A | N/A | N/A | N/A | N/A | N/A | N/A | N/A | 221.070 | 2.000 | 111.035 |
| 256 | 297.241 | 9.395 | -1 | N/A | N/A | N/A | N/A | N/A | N/A | N/A | N/A | 297.241 | 1.000 | 297.241 |
| 259 | 249.185 | 8.504 | -1 | N/A | N/A | N/A | N/A | N/A | N/A | N/A | N/A | 249.185 | 1.000 | 249.185 |
| 260 | 290.269 | 6.924 | -1 | N/A | N/A | N/A | N/A | N/A | N/A | N/A | N/A | 290.269 | 1.000 | 290.269 |
| 265 | 535.153 | 8.517 | -1 | N/A | N/A | N/A | N/A | N/A | N/A | N/A | N/A | 535.153 | 1.000 | 535.153 |
| 267 | 453.168 | 7.817 | -1 | N/A | N/A | N/A | N/A | N/A | N/A | N/A | N/A | 453.168 | 1.000 | 453.168 |
| 271 | 283.104 | 5.343 | -1 | N/A | N/A | N/A | N/A | N/A | N/A | N/A | N/A | 283.104 | 1.000 | 283.104 |
| 274 | 537.311 | 10.526 | -1 | N/A | N/A | N/A | N/A | N/A | N/A | N/A | N/A | 537.311 | 1.000 | 537.311 |
| 279 | 149.024 | 6.761 | -1 | N/A | N/A | N/A | N/A | N/A | N/A | N/A | N/A | 149.024 | 1.000 | 149.024 |
| 281 | 467.147 | 6.891 | -1 | N/A | N/A | N/A | N/A | N/A | N/A | N/A | N/A | 467.147 | 1.000 | 467.147 |
| 285 | 408.309 | 10.166 | -1 | N/A | N/A | N/A | N/A | N/A | N/A | N/A | N/A | 408.309 | 1.000 | 408.309 |
| 286 | 291.195 | 8.162 | -1 | N/A | N/A | N/A | N/A | N/A | N/A | N/A | N/A | 291.195 | 1.000 | 291.195 |
| 287 | 573.110 | 8.520 | -1 | N/A | N/A | N/A | N/A | N/A | N/A | N/A | N/A | 573.110 | 1.000 | 573.110 |
| 299 | 437.194 | 8.754 | -1 | N/A | N/A | N/A | N/A | N/A | N/A | N/A | N/A | 437.194 | 1.000 | 437.194 |
| 300 | 604.812 | 4.484 | -1 | N/A | N/A | N/A | N/A | N/A | N/A | N/A | N/A | 1208.620 | 2.000 | 604.808 |
| 304 | 365.137 | 8.808 | -1 | N/A | N/A | N/A | N/A | N/A | N/A | N/A | N/A | 365.137 | 1.000 | 365.137 |
| 305 | 480.273 | 3.605 | -1 | N/A | N/A | N/A | N/A | N/A | N/A | N/A | N/A | 959.538 | 2.000 | 480.269 |
| 308 | 659.288 | 10.020 | -1 | N/A | N/A | N/A | N/A | N/A | N/A | N/A | N/A | 659.288 | 1.000 | 659.288 |
| 309 | 165.076 | 4.458 | -1 | N/A | N/A | N/A | N/A | N/A | N/A | N/A | N/A | 165.076 | 1.000 | 165.076 |
| 320 | 341.267 | 9.373 | -1 | N/A | N/A | N/A | N/A | N/A | N/A | N/A | N/A | 341.267 | 1.000 | 341.267 |
| 324 | 270.315 | 9.426 | -1 | N/A | N/A | N/A | N/A | N/A | N/A | N/A | N/A | 270.315 | 1.000 | 270.315 |
| 325 | 326.378 | 9.409 | -1 | N/A | N/A | N/A | N/A | N/A | N/A | N/A | N/A | 326.378 | 1.000 | 326.378 |
| 327 | 131.049 | 0.513 | -1 | N/A | N/A | N/A | N/A | N/A | N/A | N/A | N/A | 131.049 | 1.000 | 131.049 |
| 331 | 260.165 | 6.499 | -1 | N/A | N/A | N/A | N/A | N/A | N/A | N/A | N/A | 519.322 | 2.000 | 260.161 |
| 334 | 469.184 | 6.409 | -1 | N/A | N/A | N/A | N/A | N/A | N/A | N/A | N/A | 469.184 | 1.000 | 469.184 |
| 335 | 469.163 | 6.314 | -1 | N/A | N/A | N/A | N/A | N/A | N/A | N/A | N/A | 469.163 | 1.000 | 469.163 |
| 338 | 499.780 | 3.714 | -1 | N/A | N/A | N/A | N/A | N/A | N/A | N/A | N/A | 998.551 | 2.000 | 499.776 |
| 339 | 615.581 | 10.604 | -1 | N/A | N/A | N/A | N/A | N/A | N/A | N/A | N/A | 615.581 | 1.000 | 615.581 |
| 340 | 354.553 | 5.749 | -1 | N/A | N/A | N/A | N/A | N/A | N/A | N/A | N/A | 708.099 | 2.000 | 354.550 |
| 345 | 236.142 | 5.355 | -1 | N/A | N/A | N/A | N/A | N/A | N/A | N/A | N/A | 236.142 | 1.000 | 236.142 |
| 351 | 451.173 | 6.881 | -1 | N/A | N/A | N/A | N/A | N/A | N/A | N/A | N/A | 451.173 | 1.000 | 451.173 |
| 356 | 310.173 | 6.844 | -1 | N/A | N/A | N/A | N/A | N/A | N/A | N/A | N/A | 619.339 | 2.000 | 310.170 |
| 365 | 519.261 | 9.835 | -1 | N/A | N/A | N/A | N/A | N/A | N/A | N/A | N/A | 519.261 | 1.000 | 519.261 |
| 368 | 273.167 | 4.221 | -1 | N/A | N/A | N/A | N/A | N/A | N/A | N/A | N/A | 273.167 | 1.000 | 273.167 |
| 376 | 288.181 | 4.136 | -1 | N/A | N/A | N/A | N/A | N/A | N/A | N/A | N/A | 575.353 | 2.000 | 288.177 |
| 379 | 496.301 | 10.257 | -1 | N/A | N/A | N/A | N/A | N/A | N/A | N/A | N/A | 496.301 | 1.000 | 496.301 |
| 380 | 497.280 | 9.835 | -1 | N/A | N/A | N/A | N/A | N/A | N/A | N/A | N/A | 497.280 | 1.000 | 497.280 |
| 385 | 188.069 | 0.519 | -1 | N/A | N/A | N/A | N/A | N/A | N/A | N/A | N/A | 188.069 | 1.000 | 188.069 |
| 395 | 721.506 | 12.424 | -1 | N/A | N/A | N/A | N/A | N/A | N/A | N/A | N/A | 721.506 | 1.000 | 721.506 |
| 397 | 459.263 | 10.315 | -1 | N/A | N/A | N/A | N/A | N/A | N/A | N/A | N/A | 459.263 | 1.000 | 459.263 |
| 398 | 445.291 | 10.157 | -1 | N/A | N/A | N/A | N/A | N/A | N/A | N/A | N/A | 445.291 | 1.000 | 445.291 |
| 403 | 763.517 | 12.425 | -1 | N/A | N/A | N/A | N/A | N/A | N/A | N/A | N/A | 763.517 | 1.000 | 763.517 |
| 405 | 528.788 | 4.417 | -1 | N/A | N/A | N/A | N/A | N/A | N/A | N/A | N/A | 1056.570 | 2.000 | 528.784 |
| 407 | 455.335 | 10.125 | -1 | N/A | N/A | N/A | N/A | N/A | N/A | N/A | N/A | 455.335 | 1.000 | 455.335 |
| 410 | 830.949 | 3.185 | -1 | N/A | N/A | N/A | N/A | N/A | N/A | N/A | N/A | 2490.830 | 3.000 | 830.944 |
| 421 | 514.951 | 4.135 | -1 | N/A | N/A | N/A | N/A | N/A | N/A | N/A | N/A | 1542.840 | 3.000 | 514.946 |
| 423 | 409.180 | 7.399 | -1 | N/A | N/A | N/A | N/A | N/A | N/A | N/A | N/A | 409.180 | 1.000 | 409.180 |
| 425 | 707.607 | 12.056 | -1 | N/A | N/A | N/A | N/A | N/A | N/A | N/A | N/A | 707.607 | 1.000 | 707.607 |
| 428 | 366.633 | 3.310 | -1 | N/A | N/A | N/A | N/A | N/A | N/A | N/A | N/A | 732.259 | 2.000 | 366.629 |
| 430 | 379.862 | 4.127 | -1 | N/A | N/A | N/A | N/A | N/A | N/A | N/A | N/A | 1137.570 | 3.000 | 379.857 |
| 436 | 267.137 | 8.329 | -1 | N/A | N/A | N/A | N/A | N/A | N/A | N/A | N/A | 533.267 | 2.000 | 267.133 |
| 449 | 658.341 | 3.958 | -1 | N/A | N/A | N/A | N/A | N/A | N/A | N/A | N/A | 1315.670 | 2.000 | 658.337 |
| 451 | 254.660 | 3.065 | -1 | N/A | N/A | N/A | N/A | N/A | N/A | N/A | N/A | 508.312 | 2.000 | 254.656 |
| 453 | 441.346 | 7.492 | -1 | N/A | N/A | N/A | N/A | N/A | N/A | N/A | N/A | 441.346 | 1.000 | 441.346 |
| 454 | 409.163 | 6.914 | -1 | N/A | N/A | N/A | N/A | N/A | N/A | N/A | N/A | 409.163 | 1.000 | 409.163 |
| 470 | 469.163 | 6.021 | -1 | N/A | N/A | N/A | N/A | N/A | N/A | N/A | N/A | 469.163 | 1.000 | 469.163 |
| 474 | 501.170 | 0.519 | -1 | N/A | N/A | N/A | N/A | N/A | N/A | N/A | N/A | 501.170 | 1.000 | 501.170 |
| 485 | 517.142 | 0.507 | -1 | N/A | N/A | N/A | N/A | N/A | N/A | N/A | N/A | 517.142 | 1.000 | 517.142 |
| 496 | 226.012 | 12.739 | -1 | N/A | N/A | N/A | N/A | N/A | N/A | N/A | N/A | 226.012 | 1.000 | 226.012 |
| 502 | 102.034 | 0.568 | -1 | N/A | N/A | N/A | N/A | N/A | N/A | N/A | N/A | 203.060 | 2.000 | 102.030 |
| 503 | 228.207 | 4.644 | -1 | N/A | N/A | N/A | N/A | N/A | N/A | N/A | N/A | 228.207 | 1.000 | 228.207 |
| 508 | 1011.286 | 0.501 | -1 | N/A | N/A | N/A | N/A | N/A | N/A | N/A | N/A | 1011.290 | 1.000 | 1011.290 |
| 509 | 385.095 | 3.893 | -1 | N/A | N/A | N/A | N/A | N/A | N/A | N/A | N/A | 769.181 | 2.000 | 385.091 |

| Feature identifiers | | | | Library Hits | | | | | | | | | | |
|---|---|---|---|---|---|---|---|---|---|---|---|---|---|---|
| ClusterIdx | m/z | RT [min] | Subnetwork | Library | Instrument | Ion source | Ion mode | Library m/z | MasDiff | Cosine | PPM | Parent mass | Precursor charge | Precursor mass |
| 525 | 269.210 | 8.072 | -1 | N/A | N/A | N/A | N/A | N/A | N/A | N/A | N/A | 269.210 | 1.000 | 269.210 |
| 538 | 844.952 | 3.465 | -1 | N/A | N/A | N/A | N/A | N/A | N/A | N/A | N/A | 2532.840 | 3.000 | 844.947 |
| 543 | 453.188 | 6.313 | -1 | N/A | N/A | N/A | N/A | N/A | N/A | N/A | N/A | 453.188 | 1.000 | 453.188 |
| 547 | 372.717 | 12.740 | -1 | N/A | N/A | N/A | N/A | N/A | N/A | N/A | N/A | 744.427 | 2.000 | 372.714 |
| 549 | 287.114 | 4.597 | -1 | N/A | N/A | N/A | N/A | N/A | N/A | N/A | N/A | 287.114 | 1.000 | 287.114 |
| 551 | 385.294 | 9.364 | -1 | N/A | N/A | N/A | N/A | N/A | N/A | N/A | N/A | 385.294 | 1.000 | 385.294 |
| 557 | 858.957 | 3.760 | -1 | N/A | N/A | N/A | N/A | N/A | N/A | N/A | N/A | 2574.850 | 3.000 | 858.951 |
| 563 | 429.319 | 9.342 | -1 | N/A | N/A | N/A | N/A | N/A | N/A | N/A | N/A | 429.319 | 1.000 | 429.319 |
| 566 | 472.175 | 4.688 | -1 | N/A | N/A | N/A | N/A | N/A | N/A | N/A | N/A | 472.175 | 1.000 | 472.175 |
| 569 | 294.074 | 0.519 | -1 | N/A | N/A | N/A | N/A | N/A | N/A | N/A | N/A | 294.074 | 1.000 | 294.074 |
| 575 | 453.189 | 6.022 | -1 | N/A | N/A | N/A | N/A | N/A | N/A | N/A | N/A | 453.189 | 1.000 | 453.189 |
| 586 | 191.086 | 3.681 | -1 | N/A | N/A | N/A | N/A | N/A | N/A | N/A | N/A | 191.086 | 1.000 | 191.086 |
| 592 | 359.104 | 3.680 | -1 | N/A | N/A | N/A | N/A | N/A | N/A | N/A | N/A | 359.104 | 1.000 | 359.104 |
| 593 | 511.165 | 3.612 | -1 | N/A | N/A | N/A | N/A | N/A | N/A | N/A | N/A | 511.165 | 1.000 | 511.165 |
| 594 | 700.315 | 10.020 | -1 | N/A | N/A | N/A | N/A | N/A | N/A | N/A | N/A | 700.315 | 1.000 | 700.315 |
| 597 | 225.149 | 6.492 | -1 | N/A | N/A | N/A | N/A | N/A | N/A | N/A | N/A | 225.149 | 1.000 | 225.149 |
| 599 | 131.049 | 0.515 | -1 | N/A | N/A | N/A | N/A | N/A | N/A | N/A | N/A | 131.049 | 1.000 | 131.049 |
| 601 | 357.262 | 8.098 | -1 | N/A | N/A | N/A | N/A | N/A | N/A | N/A | N/A | 357.262 | 1.000 | 357.262 |
| 602 | 303.138 | 3.688 | -1 | N/A | N/A | N/A | N/A | N/A | N/A | N/A | N/A | 303.138 | 1.000 | 303.138 |
| 605 | 347.222 | 7.583 | -1 | N/A | N/A | N/A | N/A | N/A | N/A | N/A | N/A | 347.222 | 1.000 | 347.222 |
| 606 | 438.143 | 7.111 | -1 | N/A | N/A | N/A | N/A | N/A | N/A | N/A | N/A | 438.143 | 1.000 | 438.143 |
| 607 | 851.398 | 7.250 | -1 | N/A | N/A | N/A | N/A | N/A | N/A | N/A | N/A | 851.398 | 1.000 | 851.398 |
| 609 | 830.614 | 3.188 | -1 | N/A | N/A | N/A | N/A | N/A | N/A | N/A | N/A | 2489.830 | 3.000 | 830.608 |
| 612 | 863.046 | 3.487 | -1 | N/A | N/A | N/A | N/A | N/A | N/A | N/A | N/A | 863.046 | 1.000 | 863.046 |
| 613 | 619.272 | 4.460 | -1 | N/A | N/A | N/A | N/A | N/A | N/A | N/A | N/A | 619.272 | 1.000 | 619.272 |
| 617 | 454.172 | 7.812 | -1 | N/A | N/A | N/A | N/A | N/A | N/A | N/A | N/A | 454.172 | 1.000 | 454.172 |
| 621 | 815.644 | 3.349 | -1 | N/A | N/A | N/A | N/A | N/A | N/A | N/A | N/A | 2444.920 | 3.000 | 815.639 |
| 626 | 331.209 | 4.774 | -1 | N/A | N/A | N/A | N/A | N/A | N/A | N/A | N/A | 331.209 | 1.000 | 331.209 |
| 628 | 103.054 | 0.515 | -1 | N/A | N/A | N/A | N/A | N/A | N/A | N/A | N/A | 103.054 | 1.000 | 103.054 |
| 641 | 158.154 | 6.250 | -1 | N/A | N/A | N/A | N/A | N/A | N/A | N/A | N/A | 158.154 | 1.000 | 158.154 |
| 642 | 165.114 | 12.714 | -1 | N/A | N/A | N/A | N/A | N/A | N/A | N/A | N/A | 165.114 | 1.000 | 165.114 |
| 648 | 425.137 | 6.913 | -1 | N/A | N/A | N/A | N/A | N/A | N/A | N/A | N/A | 425.137 | 1.000 | 425.137 |
| 651 | 267.157 | 4.332 | -1 | N/A | N/A | N/A | N/A | N/A | N/A | N/A | N/A | 267.157 | 1.000 | 267.157 |
| 655 | 290.232 | 4.304 | -1 | N/A | N/A | N/A | N/A | N/A | N/A | N/A | N/A | 290.232 | 1.000 | 290.232 |
| 660 | 391.205 | 10.042 | -1 | N/A | N/A | N/A | N/A | N/A | N/A | N/A | N/A | 391.205 | 1.000 | 391.205 |
| 673 | 870.956 | 4.058 | -1 | N/A | N/A | N/A | N/A | N/A | N/A | N/A | N/A | 2610.850 | 3.000 | 870.951 |
| 682 | 425.215 | 9.512 | -1 | N/A | N/A | N/A | N/A | N/A | N/A | N/A | N/A | 425.215 | 1.000 | 425.215 |
| 684 | 612.425 | 0.524 | -1 | N/A | N/A | N/A | N/A | N/A | N/A | N/A | N/A | 612.425 | 1.000 | 612.425 |
| 686 | 231.081 | 0.527 | -1 | N/A | N/A | N/A | N/A | N/A | N/A | N/A | N/A | 231.081 | 1.000 | 231.081 |
| 688 | 726.463 | 12.425 | -1 | N/A | N/A | N/A | N/A | N/A | N/A | N/A | N/A | 726.463 | 1.000 | 726.463 |
| 690 | 403.233 | 9.514 | -1 | N/A | N/A | N/A | N/A | N/A | N/A | N/A | N/A | 403.233 | 1.000 | 403.233 |
| 694 | 111.020 | 12.539 | -1 | N/A | N/A | N/A | N/A | N/A | N/A | N/A | N/A | 111.020 | 1.000 | 111.020 |
| 695 | 351.102 | 8.964 | -1 | N/A | N/A | N/A | N/A | N/A | N/A | N/A | N/A | 351.102 | 1.000 | 351.102 |
| 700 | 164.051 | 4.799 | -1 | N/A | N/A | N/A | N/A | N/A | N/A | N/A | N/A | 164.051 | 1.000 | 164.051 |
| 702 | 111.039 | 2.302 | -1 | N/A | N/A | N/A | N/A | N/A | N/A | N/A | N/A | 221.070 | 2.000 | 111.035 |
| 705 | 271.031 | 5.195 | -1 | N/A | N/A | N/A | N/A | N/A | N/A | N/A | N/A | 271.031 | 1.000 | 271.031 |
| 706 | 249.050 | 5.192 | -1 | N/A | N/A | N/A | N/A | N/A | N/A | N/A | N/A | 249.050 | 1.000 | 249.050 |
| 707 | 279.060 | 6.671 | -1 | N/A | N/A | N/A | N/A | N/A | N/A | N/A | N/A | 279.060 | 1.000 | 279.060 |
| 708 | 162.035 | 6.813 | -1 | N/A | N/A | N/A | N/A | N/A | N/A | N/A | N/A | 162.035 | 1.000 | 162.035 |
| 709 | 177.083 | 2.375 | -1 | N/A | N/A | N/A | N/A | N/A | N/A | N/A | N/A | 177.083 | 1.000 | 177.083 |
| 711 | 164.051 | 2.397 | -1 | N/A | N/A | N/A | N/A | N/A | N/A | N/A | N/A | 164.051 | 1.000 | 164.051 |
| 712 | 245.070 | 2.402 | -1 | N/A | N/A | N/A | N/A | N/A | N/A | N/A | N/A | 245.070 | 1.000 | 245.070 |
| 716 | 111.039 | 2.354 | -1 | N/A | N/A | N/A | N/A | N/A | N/A | N/A | N/A | 221.070 | 2.000 | 111.035 |
| 720 | 495.251 | 4.182 | -1 | N/A | N/A | N/A | N/A | N/A | N/A | N/A | N/A | 989.494 | 2.000 | 495.247 |
| 723 | 162.073 | 2.320 | -1 | N/A | N/A | N/A | N/A | N/A | N/A | N/A | N/A | 162.073 | 1.000 | 162.073 |
| 725 | 1347.881 | 12.426 | -1 | N/A | N/A | N/A | N/A | N/A | N/A | N/A | N/A | 1347.880 | 1.000 | 1347.880 |
| 733 | 297.083 | 6.817 | -1 | N/A | N/A | N/A | N/A | N/A | N/A | N/A | N/A | 297.083 | 1.000 | 297.083 |
| 734 | 471.252 | 7.681 | -1 | N/A | N/A | N/A | N/A | N/A | N/A | N/A | N/A | 941.495 | 2.000 | 471.248 |
| 735 | 597.678 | 0.494 | -1 | N/A | N/A | N/A | N/A | N/A | N/A | N/A | N/A | 1194.350 | 2.000 | 597.674 |
| 737 | 701.411 | 12.427 | -1 | N/A | N/A | N/A | N/A | N/A | N/A | N/A | N/A | 701.411 | 1.000 | 701.411 |
| 738 | 253.602 | 7.678 | -1 | N/A | N/A | N/A | N/A | N/A | N/A | N/A | N/A | 506.196 | 2.000 | 253.598 |
| 739 | 749.548 | 12.237 | -1 | N/A | N/A | N/A | N/A | N/A | N/A | N/A | N/A | 749.548 | 1.000 | 749.548 |
| 742 | 287.650 | 8.328 | -1 | N/A | N/A | N/A | N/A | N/A | N/A | N/A | N/A | 574.292 | 2.000 | 287.646 |
| 744 | 247.129 | 0.492 | -1 | N/A | N/A | N/A | N/A | N/A | N/A | N/A | N/A | 247.129 | 1.000 | 247.129 |
| 748 | 654.456 | 10.203 | -1 | N/A | N/A | N/A | N/A | N/A | N/A | N/A | N/A | 654.456 | 1.000 | 654.456 |
| 752 | 286.520 | 6.796 | -1 | N/A | N/A | N/A | N/A | N/A | N/A | N/A | N/A | 572.032 | 2.000 | 286.516 |
| 753 | 335.125 | 8.963 | -1 | N/A | N/A | N/A | N/A | N/A | N/A | N/A | N/A | 335.125 | 1.000 | 335.125 |
| 754 | 533.934 | 3.609 | -1 | N/A | N/A | N/A | N/A | N/A | N/A | N/A | N/A | 1599.790 | 3.000 | 533.929 |
| 762 | 533.282 | 9.675 | -1 | N/A | N/A | N/A | N/A | N/A | N/A | N/A | N/A | 533.282 | 1.000 | 533.282 |
| 763 | 1194.349 | 0.496 | -1 | N/A | N/A | N/A | N/A | N/A | N/A | N/A | N/A | 1194.350 | 1.000 | 1194.350 |

| Feature identifiers | | | | Library Hits | | | | | | | | | | |
|---|---|---|---|---|---|---|---|---|---|---|---|---|---|---|
| **ClusterIdx** | **m/z** | **RT [min]** | **Subnetwork** | **Library** | **Instrument** | **Ion source** | **Ion mode** | **Library m/z** | **MasDiff** | **Cosine** | **PPM** | **Parent mass** | **Precursor charge** | **Precursor mass** |
| 764 | 616.651 | 0.503 | -1 | N/A | N/A | N/A | N/A | N/A | N/A | N/A | N/A | 1232.300 | 2.000 | 616.648 |
| 765 | 223.064 | 6.361 | -1 | N/A | N/A | N/A | N/A | N/A | N/A | N/A | N/A | 223.064 | 1.000 | 223.064 |
| 769 | 498.173 | 7.669 | -1 | N/A | N/A | N/A | N/A | N/A | N/A | N/A | N/A | 498.173 | 1.000 | 498.173 |
| 771 | 143.107 | 1.088 | -1 | N/A | N/A | N/A | N/A | N/A | N/A | N/A | N/A | 143.107 | 1.000 | 143.107 |
| 786 | 819.476 | 8.754 | -1 | N/A | N/A | N/A | N/A | N/A | N/A | N/A | N/A | 819.476 | 1.000 | 819.476 |
| 794 | 811.308 | 6.909 | -1 | N/A | N/A | N/A | N/A | N/A | N/A | N/A | N/A | 811.308 | 1.000 | 811.308 |
| 797 | 818.299 | 3.374 | -1 | N/A | N/A | N/A | N/A | N/A | N/A | N/A | N/A | 2452.880 | 3.000 | 818.294 |
| 806 | 618.984 | 3.690 | -1 | N/A | N/A | N/A | N/A | N/A | N/A | N/A | N/A | 2472.910 | 4.000 | 618.978 |
| 808 | 832.302 | 3.693 | -1 | N/A | N/A | N/A | N/A | N/A | N/A | N/A | N/A | 2494.890 | 3.000 | 832.297 |
| 814 | 513.189 | 7.230 | -1 | N/A | N/A | N/A | N/A | N/A | N/A | N/A | N/A | 513.189 | 1.000 | 513.189 |
| 815 | 432.241 | 7.420 | -1 | N/A | N/A | N/A | N/A | N/A | N/A | N/A | N/A | 432.241 | 1.000 | 432.241 |
| 818 | 123.079 | 0.380 | -1 | N/A | N/A | N/A | N/A | N/A | N/A | N/A | N/A | 123.079 | 1.000 | 123.079 |
| 822 | 376.087 | 6.788 | -1 | N/A | N/A | N/A | N/A | N/A | N/A | N/A | N/A | 376.087 | 1.000 | 376.087 |
| 827 | 326.112 | 5.859 | -1 | N/A | N/A | N/A | N/A | N/A | N/A | N/A | N/A | 326.112 | 1.000 | 326.112 |
| 828 | 319.064 | 6.836 | -1 | N/A | N/A | N/A | N/A | N/A | N/A | N/A | N/A | 319.064 | 1.000 | 319.064 |
| 831 | 1295.589 | 10.022 | -1 | N/A | N/A | N/A | N/A | N/A | N/A | N/A | N/A | 2590.170 | 2.000 | 1295.580 |
| 834 | 628.471 | 3.460 | -1 | N/A | N/A | N/A | N/A | N/A | N/A | N/A | N/A | 2510.860 | 4.000 | 628.465 |
| 835 | 850.609 | 3.462 | -1 | N/A | N/A | N/A | N/A | N/A | N/A | N/A | N/A | 2549.810 | 3.000 | 850.604 |
| 839 | 372.717 | 10.091 | -1 | N/A | N/A | N/A | N/A | N/A | N/A | N/A | N/A | 744.426 | 2.000 | 372.713 |
| 846 | 681.132 | 5.759 | -1 | N/A | N/A | N/A | N/A | N/A | N/A | N/A | N/A | 681.132 | 1.000 | 681.132 |
| 856 | 1194.817 | 11.488 | -1 | N/A | N/A | N/A | N/A | N/A | N/A | N/A | N/A | 1194.820 | 1.000 | 1194.820 |
| 857 | 804.296 | 3.067 | -1 | N/A | N/A | N/A | N/A | N/A | N/A | N/A | N/A | 2410.870 | 3.000 | 804.291 |
| 863 | 353.267 | 9.157 | -1 | N/A | N/A | N/A | N/A | N/A | N/A | N/A | N/A | 353.267 | 1.000 | 353.267 |

| Feature identifiers | | | | | Analog Hits | |
|---|---|---|---|---|---|---|
| ClusterIdx | m/z | RT [min] | Subnetwork | Shared Peaks | Analog name | Cosine |
| 491 | 205.195 | 9.007 | -1 | 7 | Cadinene | 0.963 |
| 24 | 399.252 | 8.753 | 58 | 10 | Spectral Match to Tris(2-butoxyethyl) phosphate from NIST14 | 0.961 |
| 44 | 421.232 | 8.753 | -1 | 8 | Spectral Match to Tris(2-butoxyethyl) phosphate from NIST14 | 0.987 |
| 130 | 242.100 | 0.564 | 54 | 6 | Spectral Match to Pantothenic acid from NIST14 | 0.930 |
| 479 | 242.101 | 0.569 | 54 | 6 | Spectral Match to Pantothenic acid from NIST14 | 0.946 |
| 478 | 204.087 | 0.529 | 51 | 8 | Spectral Match to N-Acetyl-D-glucosamine from NIST14 | 0.999 |
| 200 | 311.256 | 7.678 | 39 | 10 | Spectral Match to Monopalmitolein (9c) from NIST14 | 0.954 |
| 461 | 311.256 | 7.795 | 39 | 10 | Spectral Match to Monopalmitolein (9c) from NIST14 | 0.952 |
| 536 | 339.290 | 8.561 | 39 | 11 | 1-Oleoyl-2-acetyl-sn-glycerol | 0.983 |
| 574 | 339.289 | 8.380 | 39 | 11 | 1-Oleoyl-2-acetyl-sn-glycerol | 0.989 |
| 666 | 188.071 | 1.307 | 27 | 7 | Spectral Match to L-Tryptophan from NIST14 | 0.988 |
| 6 | 166.086 | 0.517 | 41 | 7 | Spectral Match to DL-Phenylalanine from NIST14 | 0.998 |
| 441 | 223.090 | 6.761 | -1 | 8 | Spectral Match to Diethyl phthalate from NIST14 | 0.995 |
| 378 | 391.285 | 10.644 | -1 | 4 | Spectral Match to Bis(2-ethylhexyl) phthalate from NIST14 | 0.971 |
| 12 | 245.096 | 3.603 | 62 | 11 | Spectral Match to Biotin from NIST14 | 0.999 |
| 408 | 489.184 | 3.610 | 62 | 7 | Spectral Match to Biotin from NIST14 | 0.983 |
| 467 | 250.118 | 7.417 | -1 | 6 | Spectral Match to 2,4,6-Triallyloxy-1,3,5-triazine from NIST14 | 0.959 |
| 462 | 482.321 | 9.064 | 4 | 9 | Spectral Match to 1-(9Z-Octadecenoyl)-sn-glycero-3-phosphoethanolamine from NIST14 | 0.959 |
| 832 | 577.516 | 12.699 | 39 | 11 | Spectral Match to 1-Palmitoyl-2-oleoyl-sn-glycerol from NIST14 | 0.938 |
| 17 | 454.293 | 8.329 | 4 | 10 | Spectral Match to 1-(9Z-Octadecenoyl)-sn-glycero-3-phosphoethanolamine from NIST14 | 0.961 |
| 89 | 454.293 | 8.190 | 4 | 10 | PE(16:0/0:0); [M+H]+ C21H45N1O7P1 | 0.922 |
| 211 | 476.267 | 8.328 | 4 | 12 | Spectral Match to 1-Hexadecanoyl-sn-glycero-3-phosphoethanolamine from NIST14 | 0.972 |
| 743 | 907.578 | 8.331 | -1 | 6 | Spectral Match to 1-Hexadecanoyl-sn-glycero-3-phosphoethanolamine from NIST14 | 0.820 |
| 138 | 338.342 | 10.765 | 39 | 11 | Monoerucin | 0.811 |
| 456 | 338.342 | 9.820 | 39 | 10 | Spectral Match to 13-Docosenamide, (Z)- from NIST14 | 0.900 |
| 767 | 688.493 | 12.701 | 4 | 11 | PE(16:1/16:1); [M+H]+ C37H71N1O8P1 | 0.839 |
| 30 | 480.309 | 8.382 | 4 | 12 | PE(18:1/0:0); [M+H]+ C23H47N1O7P1 | 0.924 |
| 36 | 480.309 | 8.512 | 4 | 14 | Spectral Match to 1-(9Z-Octadecenoyl)-sn-glycero-3-phosphoethanolamine from NIST14 | 0.948 |
| 342 | 377.146 | 3.769 | -1 | 9 | Spectral Match to (-)-Riboflavin from NIST14 | 0.903 |
| 473 | 220.118 | 0.532 | 61 | 9 | Pantothenic Acid | 0.956 |
| 372 | 628.196 | 10.499 | 40 | 14 | 2-[3,4-bis[[(2S,3R,4S,5S,6R)-3,4,5-trihydroxy-6-(hydroxymethyl)oxan-2-yl]oxy]phenyl]-5,7-dihydroxychromen-4-one | 0.756 |
| 3 | 432.238 | 7.526 | 24 | 8 | 7b,9-Dihydroxy-3-(hydroxymethyl)-1,1,6,8-tetramethyl-5-oxo-1,1a,1b,4,4a,5,7a,7b,8,9-decahydro-9aH-cyclopropa[3,4]benzo[1,2-e]azulen-9a-yl acetate | 0.896 |
| 161 | 432.238 | 7.249 | 24 | 7 | 7b,9-Dihydroxy-3-(hydroxymethyl)-1,1,6,8-tetramethyl-5-oxo-1,1a,1b,4,4a,5,7a,7b,8,9-decahydro-9aH-cyclopropa[3,4]benzo[1,2-e]azulen-9a-yl acetate | 0.897 |
| 442 | 432.238 | 7.812 | 24 | 6 | 6-[3-[(3,4-dimethoxyphenyl)methyl]-4-methoxy-2-(methoxymethyl)butyl]-4-methoxy-1,3-benzodioxole | 0.887 |
| 2 | 415.212 | 7.526 | 24 | 6 | 2H-Oxireno[1,10a]phenanthro[3,2-b]furan-10(11bH)-one, 5,7-bis(acetyloxy)-3,3a,4,5,6,7,7a,7b,8,8a-decahydro-4,4,7a,11-tetramethyl-, (1aS,3aR,5S,7S,7aR,7bS,8aR,11bR)- | 0.931 |
| 78 | 415.212 | 7.248 | 24 | 6 | 2H-Oxireno[1,10a]phenanthro[3,2-b]furan-10(11bH)-one, 5,7-bis(acetyloxy)-3,3a,4,5,6,7,7a,7b,8,8a-decahydro-4,4,7a,11-tetramethyl-, (1aS,3aR,5S,7S,7aR,7bS,8aR,11bR)- | 0.936 |
| 373 | 415.212 | 7.817 | 24 | 6 | 2H-Oxireno[1,10a]phenanthro[3,2-b]furan-10(11bH)-one, 5,7-bis(acetyloxy)-3,3a,4,5,6,7,7a,7b,8,8a-decahydro-4,4,7a,11-tetramethyl-, (1aS,3aR,5S,7S,7aR,7bS,8aR,11bR)- | 0.931 |
| 40 | 327.078 | 8.430 | -1 | 16 | MassbankEU:SM879502 Triphenylphosphate\|Triphenyl phosphate | 0.967 |
| 435 | 313.143 | 8.965 | -1 | 4 | MassbankEU:SM836901 Benzyl butyl phthalate\|2-O-benzyl 1-O-butyl benzene-1,2-dicarboxylate\|2-O-benzyl 1-O-butyl benzene-1,2-dicarboxylate | 0.982 |
| 80 | 223.089 | 2.607 | 27 | 7 | L-Tryptophan | 0.975 |
| 697 | 223.089 | 2.398 | 27 | 8 | Massbank:BML01533 6-Fluoro-DL-tryptophan | 0.999 |
| 798 | 223.089 | 2.491 | 27 | 8 | Massbank:BML01533 6-Fluoro-DL-tryptophan | 0.998 |
| 665 | 205.097 | 1.310 | 27 | 7 | Massbank:BML01533 6-Fluoro-DL-tryptophan | 0.998 |
| 154 | 457.112 | 3.497 | -1 | 19 | Flavine mononucleotide | 0.924 |
| 29 | 220.118 | 0.526 | 61 | 11 | D-PANTOTHENIC ACID | 0.982 |
| 681 | 331.165 | 0.498 | 41 | 5 | DL-Phenylalanine | 0.993 |
| 192 | 343.296 | 6.715 | 23 | 5 | CocamidoprpylBetaine | 0.971 |
| 589 | 481.400 | 5.241 | 3 | 4 | Bis(2,2,6,6-tetramethyl-4-piperidyl) sebacate | 0.807 |
| 603 | 810.134 | 3.065 | 20 | 14 | Acetyl-CoA | 0.939 |
| 521 | 716.523 | 12.701 | 4 | 6 | 1-Stearoyl-2-linoleoyl-sn-glycero-3-phosphoethanolamine | 0.902 |
| 572 | 716.525 | 10.404 | 4 | 6 | 1-Stearoyl-2-linoleoyl-sn-glycero-3-phosphoethanolamine | 0.898 |
| 654 | 716.524 | 10.304 | 4 | 6 | PE(16:1/18:1); [M+H]+ C39H75N1O8P1 | 0.910 |
| 852 | 716.524 | 10.151 | 4 | 5 | 1-Stearoyl-2-linoleoyl-sn-glycero-3-phosphoethanolamine | 0.892 |
| 524 | 426.262 | 7.583 | 4 | 7 | Spectral Match to 1-(9Z-Octadecenoyl)-sn-glycero-3-phosphoethanolamine from NIST14 | 0.950 |
| 552 | 426.262 | 7.443 | 4 | 7 | Spectral Match to 1-Palmitoyl-2-hydroxy-sn-glycero-3-phosphoethanolamine from NIST14 | 0.894 |
| 507 | 718.539 | 12.704 | 4 | 13 | 1-Hexadecanoyl-2-(9Z-octadecenoyl)-sn-glycero-3-phosphoethanolamine | 0.894 |
| 523 | 718.539 | 10.302 | 4 | 13 | 1-Hexadecanoyl-2-(9Z-octadecenoyl)-sn-glycero-3-phosphoethanolamine | 0.898 |
| 632 | 718.538 | 10.143 | 4 | 13 | 1-Hexadecanoyl-2-(9Z-octadecenoyl)-sn-glycero-3-phosphoethanolamine | 0.898 |
| 823 | 771.514 | 12.696 | 4 | 5 | 1,2-Dioleoyl-sn-glycero-3-phospho-rac-1-glycerol | 0.920 |
| 346 | 502.290 | 8.380 | 4 | 14 | 1-Palmitoyl-2-hydroxy-sn-glycero-3-phosphoethanolamine | 0.839 |
| 388 | 502.291 | 8.511 | 4 | 14 | Spectral Match to 1-Palmitoyl-2-hydroxy-sn-glycero-3-phosphoethanolamine from NIST14 | 0.893 |
| 476 | 979.349 | 0.522 | 91 | N/A | N/A | N/A |
| 488 | 995.318 | 0.510 | 91 | N/A | N/A | N/A |
| 52 | 256.264 | 10.439 | 88 | N/A | N/A | N/A |
| 427 | 284.295 | 10.783 | 88 | N/A | N/A | N/A |
| 62 | 291.096 | 6.457 | 87 | N/A | N/A | N/A |
| 153 | 291.096 | 6.559 | 87 | N/A | N/A | N/A |
| 374 | 748.356 | 4.106 | 84 | N/A | N/A | N/A |
| 384 | 783.875 | 4.128 | 84 | N/A | N/A | N/A |
| 284 | 243.196 | 7.929 | 80 | N/A | 2,4-dihydroxyheptadecyl acetate | 0.814 |

| Feature identifiers | | | | | Analog Hits | |
|---|---|---|---|---|---|---|
| ClusterIdx | m/z | RT [min] | Subnetwork | Shared Peaks | Analog name | Cosine |
| 415 | 225.185 | 7.934 | 80 | N/A | Conjugated linoleic Acid (10E,12Z) | 0.799 |
| 667 | 176.074 | 0.736 | 79 | N/A | Cysteine | 0.715 |
| 668 | 218.121 | 4.865 | 79 | N/A | CAY10444 | 0.864 |
| 670 | 246.152 | 6.015 | 79 | N/A | CAY10444 | 0.870 |
| 674 | 232.137 | 5.471 | 79 | N/A | CAY10444 | 0.851 |
| 675 | 204.105 | 4.140 | 79 | N/A | CAY10444 | 0.836 |
| 772 | 551.192 | 3.315 | 78 | N/A | N/A | N/A |
| 774 | 615.714 | 3.354 | 78 | N/A | N/A | N/A |
| 776 | 486.670 | 3.240 | 78 | N/A | N/A | N/A |
| 777 | 680.235 | 3.406 | 78 | N/A | N/A | N/A |
| 780 | 357.628 | 3.065 | 78 | N/A | N/A | N/A |
| 46 | 226.158 | 9.291 | 77 | N/A | Phenyltriethylammonium cation | 0.777 |
| 201 | 282.221 | 10.073 | 77 | N/A | N/A | N/A |
| 634 | 282.221 | 10.084 | 77 | N/A | N/A | N/A |
| 150 | 818.299 | 3.185 | 75 | N/A | N/A | N/A |
| 323 | 1226.943 | 3.181 | 75 | N/A | N/A | N/A |
| 843 | 1226.943 | 3.183 | 75 | N/A | N/A | N/A |
| 186 | 135.079 | 7.525 | 74 | N/A | MoNA:805057 3,5-dimethyladamantan-1-amine | 0.863 |
| 319 | 135.079 | 4.463 | 74 | N/A | MoNA:805057 3,5-dimethyladamantan-1-amine | 0.875 |
| 180 | 529.790 | 4.168 | 73 | N/A | N/A | N/A |
| 264 | 465.270 | 4.196 | 73 | N/A | N/A | N/A |
| 48 | 431.565 | 4.119 | 72 | N/A | N/A | N/A |
| 170 | 417.549 | 3.504 | 72 | N/A | N/A | N/A |
| 314 | 396.544 | 3.209 | 72 | N/A | N/A | N/A |
| 117 | 832.302 | 3.461 | 70 | N/A | N/A | N/A |
| 371 | 1247.949 | 3.461 | 70 | N/A | N/A | N/A |
| 512 | 1247.454 | 3.462 | 70 | N/A | N/A | N/A |
| 7 | 271.106 | 8.022 | 69 | N/A | N/A | N/A |
| 45 | 390.122 | 6.709 | 69 | N/A | N/A | N/A |
| 112 | 388.106 | 7.488 | 69 | N/A | N/A | N/A |
| 209 | 414.122 | 6.104 | 69 | N/A | N/A | N/A |
| 663 | 414.122 | 6.419 | 69 | N/A | N/A | N/A |
| 468 | 432.238 | 6.913 | 68 | N/A | N/A | N/A |
| 792 | 404.208 | 6.914 | 68 | N/A | N/A | N/A |
| 793 | 387.181 | 6.913 | 68 | N/A | N/A | N/A |
| 671 | 331.207 | 5.234 | 67 | N/A | N/A | N/A |
| 795 | 317.195 | 4.828 | 67 | N/A | N/A | N/A |
| 84 | 280.188 | 5.797 | 65 | N/A | (.+/-.)-Baclofen | 0.735 |
| 158 | 308.222 | 6.723 | 65 | N/A | N/A | N/A |
| 263 | 693.878 | 3.983 | 64 | N/A | N/A | N/A |
| 392 | 665.367 | 3.986 | 64 | N/A | N/A | N/A |
| 703 | 204.534 | 6.795 | 63 | N/A | N/A | N/A |
| 704 | 193.026 | 6.808 | 63 | N/A | N/A | N/A |
| 88 | 227.084 | 3.606 | 62 | N/A | Biotin | 0.886 |
| 215 | 290.153 | 3.607 | 62 | N/A | Massbank:EA290101 Citalopram\|1-[3-(dimethylamino)propyl]-1-(4-fluorophenyl)-3H-2-benzofuran-5-carbonitrile | 0.776 |
| 4 | 182.985 | 12.752 | 60 | N/A | N/A | N/A |
| 5 | 154.990 | 12.767 | 60 | N/A | N/A | N/A |
| 19 | 138.996 | 12.748 | 60 | N/A | N/A | N/A |
| 23 | 205.017 | 12.741 | 60 | N/A | N/A | N/A |
| 49 | 196.017 | 12.766 | 60 | N/A | N/A | N/A |
| 63 | 224.012 | 12.748 | 60 | N/A | N/A | N/A |
| 75 | 195.994 | 12.775 | 60 | N/A | N/A | N/A |
| 20 | 131.534 | 12.743 | 59 | N/A | N/A | N/A |
| 409 | 131.534 | 13.009 | 59 | N/A | N/A | N/A |
| 465 | 131.534 | 13.069 | 59 | N/A | N/A | N/A |
| 696 | 131.534 | 4.507 | 59 | N/A | N/A | N/A |
| 266 | 444.309 | 8.752 | 58 | N/A | Spectral Match to Tris(2-butoxyethyl) phosphate from NIST14 | 0.741 |
| 490 | 350.102 | 0.493 | 57 | N/A | S-Lactoylglutathione | 0.878 |
| 495 | 364.118 | 0.634 | 57 | N/A | S-Lactoylglutathione | 0.863 |
| 134 | 519.318 | 10.374 | 56 | N/A | N/A | N/A |
| 658 | 519.309 | 10.382 | 56 | N/A | N/A | N/A |
| 498 | 532.098 | 0.493 | 55 | N/A | N/A | N/A |
| 501 | 532.099 | 0.497 | 55 | N/A | N/A | N/A |
| 132 | 711.319 | 4.210 | 53 | N/A | N/A | N/A |
| 282 | 624.289 | 4.196 | 53 | N/A | N/A | N/A |
| 133 | 226.180 | 5.976 | 52 | N/A | 1,11-Undecanedicarboxylic acid | 0.798 |
| 542 | 384.349 | 10.283 | 52 | N/A | N/A | N/A |
| 131 | 614.463 | 9.741 | 51 | N/A | N/A | N/A |
| 243 | 388.269 | 6.414 | 51 | N/A | N/A | N/A |
| 472 | 479.186 | 0.514 | 51 | N/A | Spectral Match to Glycan Le-X Trisaccharide from NIST14 | 0.964 |

| Feature identifiers | | | | | Analog Hits | |
|---|---|---|---|---|---|---|
| ClusterIdx | m/z | RT [min] | Subnetwork | Shared Peaks | Analog name | Cosine |
| 475 | 957.367 | 0.530 | 51 | N/A | Glycan Man3a | 0.902 |
| 477 | 524.246 | 0.489 | 51 | N/A | Spectral Match to Glycan Le-A Trisaccharide from NIST14 | 0.937 |
| 480 | 268.076 | 0.497 | 51 | N/A | GalNAc1-4GlcNAc-Sp | 0.749 |
| 482 | 498.161 | 0.503 | 51 | N/A | Glycan Man3b | 0.703 |
| 483 | 644.267 | 0.510 | 51 | N/A | Glycan Man2a | 0.918 |
| 484 | 279.584 | 0.532 | 51 | N/A | N/A | N/A |
| 487 | 276.062 | 0.490 | 51 | N/A | N-Acetyl-D-glucosamine | 0.977 |
| 492 | 737.248 | 0.522 | 51 | N/A | N/A | N/A |
| 493 | 506.147 | 0.488 | 51 | N/A | Glycan Man3b | 0.741 |
| 500 | 287.570 | 0.486 | 51 | N/A | GlcNAc1-3Gal1-4Glc-Sp | 0.867 |
| 687 | 441.187 | 0.508 | 51 | N/A | Glycan Lacto-N-biose | 0.738 |
| 732 | 632.474 | 10.196 | 51 | N/A | N/A | N/A |
| 802 | 860.514 | 9.355 | 51 | N/A | N/A | N/A |
| 168 | 471.307 | 5.923 | 50 | N/A | N/A | N/A |
| 616 | 473.286 | 4.748 | 50 | N/A | N/A | N/A |
| 144 | 393.207 | 10.310 | 49 | N/A | 2-(3,4-dihydroxyphenyl)-5,7-dihydroxy-6-[3,4,5-trihydroxy-6-(hydroxymethyl)oxan-2-yl]chromen-4-one | 0.838 |
| 176 | 409.183 | 10.451 | 49 | N/A | 3,5,7,8-tetramethoxy-2-(3,4,5-trimethoxyphenyl)chromen-4-one | 0.833 |
| 43 | 418.117 | 7.633 | 48 | N/A | N/A | N/A |
| 146 | 551.149 | 6.292 | 48 | N/A | N/A | N/A |
| 363 | 418.117 | 7.570 | 48 | N/A | N/A | N/A |
| 119 | 343.130 | 3.690 | 47 | N/A | N/A | N/A |
| 182 | 537.334 | 9.562 | 47 | N/A | N/A | N/A |
| 231 | 509.303 | 8.961 | 47 | N/A | N/A | N/A |
| 239 | 245.616 | 7.678 | 46 | N/A | N/A | N/A |
| 393 | 266.129 | 7.678 | 46 | N/A | N/A | N/A |
| 429 | 266.129 | 7.807 | 46 | N/A | N/A | N/A |
| 506 | 734.471 | 12.708 | 45 | N/A | N/A | N/A |
| 544 | 734.471 | 10.338 | 45 | N/A | N/A | N/A |
| 550 | 765.459 | 12.703 | 45 | N/A | 1-Hexadecanoyl-2-(5Z,8Z,11Z,14Z-eicosatetraenoyl)-sn-glycero-3-phospho-(1'-sn-glycerol) | 0.845 |
| 646 | 734.471 | 10.261 | 45 | N/A | N/A | N/A |
| 652 | 762.502 | 12.706 | 45 | N/A | N/A | N/A |
| 370 | 514.353 | 10.382 | 43 | N/A | N/A | N/A |
| 740 | 744.593 | 12.218 | 43 | N/A | Coenzyme Q9 | 0.837 |
| 94 | 247.596 | 7.524 | 42 | N/A | N/A | N/A |
| 110 | 256.601 | 7.527 | 42 | N/A | N/A | N/A |
| 233 | 268.115 | 7.525 | 42 | N/A | N/A | N/A |
| 283 | 554.177 | 10.161 | 40 | N/A | 2-[3,4-bis[[(2S,3R,4S,5S,6R)-3,4,5-trihydroxy-6-(hydroxymethyl)oxan-2-yl]oxy]phenyl]-5,7-dihydroxychromen-4-one | 0.757 |
| 545 | 684.203 | 10.983 | 40 | N/A | N/A | N/A |
| 630 | 702.214 | 10.746 | 40 | N/A | 2-[3,4-bis[[(2S,3R,4S,5S,6R)-3,4,5-trihydroxy-6-(hydroxymethyl)oxan-2-yl]oxy]phenyl]-5,7-dihydroxychromen-4-one | 0.723 |
| 693 | 463.131 | 9.688 | 40 | N/A | N/A | N/A |
| 821 | 776.232 | 11.006 | 40 | N/A | N/A | N/A |
| 858 | 610.185 | 12.140 | 40 | N/A | N/A | N/A |
| 143 | 228.196 | 7.148 | 39 | N/A | Linoleic acid ethyl ester | 0.818 |
| 230 | 228.196 | 6.462 | 39 | N/A | Linoleic acid ethyl ester | 0.809 |
| 824 | 549.485 | 10.160 | 39 | N/A | Spectral Match to 1-Palmitoyl-2-oleoyl-sn-glycerol from NIST14 | 0.968 |
| 826 | 549.486 | 12.704 | 39 | N/A | Spectral Match to 1-Palmitoyl-2-oleoyl-sn-glycerol from NIST14 | 0.963 |
| 840 | 549.486 | 9.853 | 39 | N/A | 1-Palmitoyl-2-oleoyl-sn-glycerol | 0.880 |
| 850 | 549.485 | 10.361 | 39 | N/A | 1-Palmitoyl-2-oleoyl-sn-glycerol | 0.904 |
| 159 | 480.238 | 3.674 | 37 | N/A | N/A | N/A |
| 332 | 959.468 | 3.674 | 37 | N/A | N/A | N/A |
| 301 | 467.250 | 5.033 | 36 | N/A | N/A | N/A |
| 333 | 933.494 | 5.044 | 36 | N/A | N/A | N/A |
| 420 | 516.785 | 5.231 | 36 | N/A | N/A | N/A |
| 68 | 217.105 | 4.009 | 35 | N/A | Triethylene glycol | 0.816 |
| 330 | 234.134 | 4.002 | 35 | N/A | N/A | N/A |
| 431 | 681.651 | 10.701 | 33 | N/A | N/A | N/A |
| 434 | 399.395 | 7.402 | 33 | N/A | N/A | N/A |
| 171 | 278.185 | 3.992 | 32 | N/A | N/A | N/A |
| 437 | 280.186 | 4.202 | 32 | N/A | N/A | N/A |
| 560 | 271.180 | 3.637 | 32 | N/A | N/A | N/A |
| 35 | 255.159 | 6.531 | 31 | N/A | N-Acetyl-L-glutamic acid | 0.781 |
| 99 | 272.185 | 6.531 | 31 | N/A | N/A | N/A |
| 466 | 269.175 | 7.631 | 31 | N/A | N/A | N/A |
| 28 | 297.133 | 7.153 | 30 | N/A | N/A | N/A |
| 38 | 431.207 | 7.153 | 30 | N/A | Spectral Match to Tebufenozide from NIST14 | 0.812 |
| 56 | 447.202 | 6.602 | 30 | N/A | Spectral Match to Tebufenozide from NIST14 | 0.867 |
| 91 | 448.233 | 7.152 | 30 | N/A | N/A | N/A |
| 102 | 429.191 | 6.409 | 30 | N/A | Spectral Match to Tebufenozide from NIST14 | 0.903 |
| 163 | 297.133 | 7.367 | 30 | N/A | N/A | N/A |
| 202 | 448.233 | 7.470 | 30 | N/A | N/A | N/A |

| Feature identifiers | | | | | Analog Hits | |
|---|---|---|---|---|---|---|
| ClusterIdx | m/z | RT [min] | Subnetwork | Shared Peaks | Analog name | Cosine |
| 208 | 431.207 | 7.472 | 30 | N/A | N/A | N/A |
| 213 | 447.202 | 7.074 | 30 | N/A | Spectral Match to Tebufenozide from NIST14 | 0.878 |
| 219 | 447.202 | 6.409 | 30 | N/A | Spectral Match to Tebufenozide from NIST14 | 0.857 |
| 235 | 297.133 | 6.600 | 30 | N/A | N/A | N/A |
| 245 | 469.184 | 6.601 | 30 | N/A | N/A | N/A |
| 249 | 429.191 | 6.602 | 30 | N/A | Spectral Match to Tebufenozide from NIST14 | 0.901 |
| 251 | 297.133 | 6.410 | 30 | N/A | N/A | N/A |
| 262 | 447.202 | 6.882 | 30 | N/A | Spectral Match to Tebufenozide from NIST14 | 0.857 |
| 411 | 527.717 | 3.679 | 29 | N/A | N/A | N/A |
| 444 | 791.074 | 3.678 | 29 | N/A | N/A | N/A |
| 355 | 500.380 | 6.151 | 28 | N/A | Octapropylene glycol | 0.701 |
| 391 | 442.338 | 5.719 | 28 | N/A | Octapropylene glycol | 0.703 |
| 394 | 309.227 | 4.773 | 28 | N/A | Hexapropylene glycol | 0.739 |
| 446 | 384.296 | 5.267 | 28 | N/A | N/A | N/A |
| 458 | 674.506 | 7.346 | 28 | N/A | N/A | N/A |
| 631 | 790.590 | 8.085 | 28 | N/A | N/A | N/A |
| 635 | 549.416 | 9.554 | 28 | N/A | N/A | N/A |
| 860 | 694.522 | 10.428 | 28 | N/A | N/A | N/A |
| 357 | 206.062 | 2.609 | 27 | N/A | *5-Hydroxy-L-tryptophan* | 0.980 |
| 440 | 237.106 | 4.092 | 27 | N/A | Massbank:BML01533 6-Fluoro-DL-tryptophan | 0.918 |
| 699 | 206.061 | 2.395 | 27 | N/A | *5-Hydroxy-L-tryptophan* | 0.978 |
| 799 | 206.062 | 2.458 | 27 | N/A | *5-Hydroxy-L-tryptophan* | 0.978 |
| 448 | 213.144 | 5.071 | 26 | N/A | N/A | N/A |
| 692 | 381.331 | 9.796 | 26 | N/A | N/A | N/A |
| 210 | 445.290 | 4.792 | 25 | N/A | N/A | N/A |
| 227 | 467.300 | 4.849 | 25 | N/A | N/A | N/A |
| 250 | 423.276 | 4.729 | 25 | N/A | N/A | N/A |
| 275 | 401.261 | 4.663 | 25 | N/A | N/A | N/A |
| 307 | 379.249 | 4.593 | 25 | N/A | N/A | N/A |
| 353 | 489.316 | 4.904 | 25 | N/A | N/A | N/A |
| 358 | 505.391 | 10.706 | 25 | N/A | N/A | N/A |
| 402 | 527.403 | 10.694 | 25 | N/A | N/A | N/A |
| 459 | 549.417 | 10.679 | 25 | N/A | N/A | N/A |
| 535 | 463.345 | 9.103 | 25 | N/A | N/A | N/A |
| 540 | 441.330 | 9.123 | 25 | N/A | N/A | N/A |
| 638 | 529.383 | 9.048 | 25 | N/A | N/A | N/A |
| 653 | 507.370 | 9.067 | 25 | N/A | N/A | N/A |
| 662 | 511.328 | 4.955 | 25 | N/A | N/A | N/A |
| 759 | 571.429 | 10.663 | 25 | N/A | N/A | N/A |
| 859 | 533.341 | 5.004 | 25 | N/A | N/A | N/A |
| 9 | 460.270 | 7.525 | 24 | N/A | 6-[3-[(3,4-dimethoxyphenyl)methyl]-4-methoxy-2-(methoxymethyl)butyl]-4-methoxy-1,3-benzodioxole | 0.845 |
| 25 | 846.443 | 7.526 | 24 | N/A | N/A | N/A |
| 67 | 446.254 | 7.523 | 24 | N/A | 7b,9-Dihydroxy-3-(hydroxymethyl)-1,1,6,8-tetramethyl-5-oxo-1,1a,1b,4,4a,5,7a,7b,8,9-decahydro-9aH-cyclopropa[3,4]benzo[1,2-e]azulen-9a-yl acetate | 0.866 |
| 81 | 460.270 | 7.249 | 24 | N/A | 7b,9-Dihydroxy-3-(hydroxymethyl)-1,1,6,8-tetramethyl-5-oxo-1,1a,1b,4,4a,5,7a,7b,8,9-decahydro-9aH-cyclopropa[3,4]benzo[1,2-e]azulen-9a-yl acetate | 0.870 |
| 139 | 476.266 | 7.526 | 24 | N/A | 7b,9-Dihydroxy-3-(hydroxymethyl)-1,1,6,8-tetramethyl-5-oxo-1,1a,1b,4,4a,5,7a,7b,8,9-decahydro-9aH-cyclopropa[3,4]benzo[1,2-e]azulen-9a-yl acetate | 0.868 |
| 261 | 397.201 | 7.526 | 24 | N/A | 2H-Oxireno[1,10a]phenanthro[3,2-b]furan-10(11bH)-one, 5,7-bis(acetyloxy)-3,3a,4,5,6,7,7a,7b,8,8a-decahydro-4,4,7a,11-tetramethyl-, (1aS,3aR,5S,7S,7aR,7bS,8aR,11bR)- | 0.865 |
| 272 | 429.191 | 6.884 | 24 | N/A | 2H-Oxireno[1,10a]phenanthro[3,2-b]furan-10(11bH)-one, 5,7-bis(acetyloxy)-3,3a,4,5,6,7,7a,7b,8,8a-decahydro-4,4,7a,11-tetramethyl-, (1aS,3aR,5S,7S,7aR,7bS,8aR,11bR)- | 0.819 |
| 316 | 531.275 | 7.525 | 24 | N/A | N/A | N/A |
| 375 | 462.213 | 6.365 | 24 | N/A | N/A | N/A |
| 418 | 445.186 | 6.364 | 24 | N/A | 2H-Oxireno[1,10a]phenanthro[3,2-b]furan-10(11bH)-one, 5,7-bis(acetyloxy)-3,3a,4,5,6,7,7a,7b,8,8a-decahydro-4,4,7a,11-tetramethyl-, (1aS,3aR,5S,7S,7aR,7bS,8aR,11bR)- | 0.812 |
| 419 | 460.270 | 7.814 | 24 | N/A | 6-[3-[(3,4-dimethoxyphenyl)methyl]-4-methoxy-2-(methoxymethyl)butyl]-4-methoxy-1,3-benzodioxole | 0.861 |
| 447 | 450.204 | 8.853 | 24 | N/A | 2H-Oxireno[1,10a]phenanthro[3,2-b]furan-10(11bH)-one, 5,7-bis(acetyloxy)-3,3a,4,5,6,7,7a,7b,8,8a-decahydro-4,4,7a,11-tetramethyl-, (1aS,3aR,5S,7S,7aR,7bS,8aR,11bR)- | 0.884 |
| 469 | 445.186 | 6.211 | 24 | N/A | 2H-Oxireno[1,10a]phenanthro[3,2-b]furan-10(11bH)-one, 5,7-bis(acetyloxy)-3,3a,4,5,6,7,7a,7b,8,8a-decahydro-4,4,7a,11-tetramethyl-, (1aS,3aR,5S,7S,7aR,7bS,8aR,11bR)- | 0.811 |
| 578 | 448.233 | 6.302 | 24 | N/A | N/A | N/A |
| 582 | 448.233 | 6.159 | 24 | N/A | 7b,9-Dihydroxy-3-(hydroxymethyl)-1,1,6,8-tetramethyl-5-oxo-1,1a,1b,4,4a,5,7a,7b,8,9-decahydro-9aH-cyclopropa[3,4]benzo[1,2-e]azulen-9a-yl acetate | 0.796 |
| 636 | 462.213 | 6.211 | 24 | N/A | 7b,9-Dihydroxy-3-(hydroxymethyl)-1,1,6,8-tetramethyl-5-oxo-1,1a,1b,4,4a,5,7a,7b,8,9-decahydro-9aH-cyclopropa[3,4]benzo[1,2-e]azulen-9a-yl acetate | 0.700 |
| 637 | 431.207 | 6.148 | 24 | N/A | 6-[3-[(3,4-dimethoxyphenyl)methyl]-4-methoxy-2-(methoxymethyl)butyl]-4-methoxy-1,3-benzodioxole | 0.932 |
| 770 | 474.249 | 6.864 | 24 | N/A | N/A | N/A |
| 790 | 431.207 | 6.021 | 24 | N/A | 6-[3-[(3,4-dimethoxyphenyl)methyl]-4-methoxy-2-(methoxymethyl)butyl]-4-methoxy-1,3-benzodioxole | 0.935 |
| 221 | 371.326 | 7.517 | 23 | N/A | CocamidopropylBetaine | 0.974 |
| 31 | 328.919 | 12.753 | 22 | N/A | ethyl 2,3-dioxo-1,4-dihydroquinoline-4-carboxylate | 0.709 |
| 39 | 312.943 | 12.757 | 22 | N/A | N/A | N/A |
| 70 | 287.888 | 12.760 | 22 | N/A | N/A | N/A |
| 92 | 313.078 | 6.460 | 22 | N/A | N/A | N/A |
| 187 | 312.943 | 12.757 | 22 | N/A | N/A | N/A |
| 196 | 313.078 | 6.532 | 22 | N/A | N/A | N/A |
| 255 | 313.235 | 8.431 | 22 | N/A | N/A | N/A |
| 558 | 313.235 | 8.075 | 22 | N/A | N/A | N/A |
| 197 | 272.258 | 8.673 | 21 | N/A | Heptadecanoyl ethanolamide | 0.951 |
| 223 | 300.290 | 9.476 | 21 | N/A | Spectral Match to Heptadecanoyl ethanolamide from NIST14 | 0.908 |

| Feature identifiers | | | | | Analog Hits | |
|---|---|---|---|---|---|---|
| ClusterIdx | m/z | RT [min] | Subnetwork | Shared Peaks | Analog name | Cosine |
| 460 | 406.354 | 10.232 | 21 | N/A | N/A | N/A |
| 157 | 429.146 | 4.153 | 20 | N/A | N/A | N/A |
| 166 | 549.571 | 3.384 | 20 | N/A | N/A | N/A |
| 481 | 549.078 | 3.211 | 20 | N/A | N/A | N/A |
| 520 | 432.026 | 3.379 | 20 | N/A | N/A | N/A |
| 584 | 432.026 | 3.502 | 20 | N/A | N/A | N/A |
| 596 | 405.571 | 3.038 | 20 | N/A | Octanoyl coenzyme A | 0.772 |
| 600 | 359.104 | 0.595 | 20 | N/A | N/A | N/A |
| 611 | 401.115 | 2.454 | 20 | N/A | N/A | N/A |
| 622 | 411.021 | 3.222 | 20 | N/A | Adenosine 5'-diphosphate | 0.781 |
| 315 | 515.353 | 9.549 | 19 | N/A | N/A | N/A |
| 326 | 321.148 | 3.682 | 19 | N/A | N/A | N/A |
| 595 | 366.206 | 3.688 | 19 | N/A | N/A | N/A |
| 494 | 453.189 | 7.153 | 18 | N/A | N/A | N/A |
| 591 | 453.189 | 7.367 | 18 | N/A | N/A | N/A |
| 861 | 453.187 | 7.297 | 18 | N/A | N/A | N/A |
| 21 | 453.169 | 7.511 | 17 | N/A | N/A | N/A |
| 292 | 453.168 | 7.525 | 17 | N/A | N/A | N/A |
| 640 | 453.186 | 7.468 | 17 | N/A | N/A | N/A |
| 59 | 281.138 | 7.525 | 16 | N/A | N/A | N/A |
| 97 | 299.149 | 4.461 | 16 | N/A | (+-)-2-cis-4-trans-Abscisic acid | 0.701 |
| 172 | 299.148 | 7.523 | 16 | N/A | (+-)-2-cis-4-trans-Abscisic acid | 0.703 |
| 296 | 344.206 | 4.461 | 16 | N/A | N/A | N/A |
| 381 | 316.176 | 4.464 | 16 | N/A | N/A | N/A |
| 42 | 473.318 | 10.647 | 15 | N/A | Tri-2-ethylhexyl trimellitate | 0.720 |
| 72 | 680.481 | 12.424 | 15 | N/A | N/A | N/A |
| 74 | 475.298 | 9.832 | 15 | N/A | PC(18:2/0:0); [M+H]+ C26H51N1O7P1 | 0.727 |
| 95 | 492.324 | 9.832 | 15 | N/A | PC(18:2/0:0); [M+H]+ C26H51N1O7P1 | 0.767 |
| 96 | 654.333 | 10.021 | 15 | N/A | N/A | N/A |
| 105 | 490.345 | 10.647 | 15 | N/A | N/A | N/A |
| 129 | 663.454 | 12.425 | 15 | N/A | N/A | N/A |
| 141 | 708.511 | 12.424 | 15 | N/A | N/A | N/A |
| 181 | 637.306 | 10.020 | 15 | N/A | N/A | N/A |
| 205 | 518.376 | 10.647 | 15 | N/A | N/A | N/A |
| 273 | 287.141 | 6.170 | 15 | N/A | N/A | N/A |
| 298 | 515.329 | 10.526 | 15 | N/A | N/A | N/A |
| 311 | 474.317 | 10.257 | 15 | N/A | N/A | N/A |
| 317 | 457.288 | 10.396 | 15 | N/A | N/A | N/A |
| 318 | 387.194 | 7.399 | 15 | N/A | Massbank: Cholesterol | 0.863 |
| 426 | 682.364 | 10.020 | 15 | N/A | N/A | N/A |
| 433 | 532.356 | 10.526 | 15 | N/A | N/A | N/A |
| 452 | 519.377 | 10.257 | 15 | N/A | N/A | N/A |
| 529 | 520.355 | 9.833 | 15 | N/A | PC(18:2/0:0); [M+H]+ C26H51N1O7P1 | 0.761 |
| 726 | 1342.926 | 12.423 | 15 | N/A | N/A | N/A |
| 755 | 949.587 | 9.831 | 15 | N/A | N/A | N/A |
| 783 | 491.374 | 10.264 | 15 | N/A | N/A | N/A |
| 849 | 1370.957 | 12.421 | 15 | N/A | N/A | N/A |
| 585 | 834.093 | 3.504 | 14 | N/A | N/A | N/A |
| 619 | 792.082 | 3.197 | 14 | N/A | N/A | N/A |
| 714 | 542.297 | 4.609 | 13 | N/A | N/A | N/A |
| 717 | 492.763 | 4.165 | 13 | N/A | N/A | N/A |
| 51 | 247.034 | 5.943 | 12 | N/A | N/A | N/A |
| 698 | 247.034 | 6.796 | 12 | N/A | N/A | N/A |
| 718 | 291.023 | 6.253 | 12 | N/A | N/A | N/A |
| 16 | 122.547 | 12.762 | 11 | N/A | N/A | N/A |
| 60 | 122.547 | 14.272 | 11 | N/A | N/A | N/A |
| 103 | 122.547 | 2.944 | 11 | N/A | N/A | N/A |
| 113 | 122.547 | 2.472 | 11 | N/A | N/A | N/A |
| 115 | 122.547 | 13.880 | 11 | N/A | N/A | N/A |
| 120 | 122.547 | 2.832 | 11 | N/A | N/A | N/A |
| 123 | 122.547 | 13.067 | 11 | N/A | N/A | N/A |
| 125 | 122.547 | 2.720 | 11 | N/A | N/A | N/A |
| 127 | 122.547 | 0.660 | 11 | N/A | N/A | N/A |
| 135 | 122.547 | 13.843 | 11 | N/A | N/A | N/A |
| 140 | 122.547 | 2.622 | 11 | N/A | N/A | N/A |
| 145 | 122.547 | 2.598 | 11 | N/A | N/A | N/A |
| 147 | 122.547 | 2.749 | 11 | N/A | N/A | N/A |
| 151 | 122.547 | 13.172 | 11 | N/A | N/A | N/A |
| 156 | 122.547 | 2.528 | 11 | N/A | N/A | N/A |
| 160 | 122.547 | 2.262 | 11 | N/A | N/A | N/A |

| Feature identifiers | | | | | Analog Hits | |
|---|---|---|---|---|---|---|
| ClusterIdx | m/z | RT [min] | Subnetwork | Shared Peaks | Analog name | Cosine |
| 169 | 122.547 | 13.121 | 11 | N/A | N/A | N/A |
| 174 | 122.547 | 2.436 | 11 | N/A | N/A | N/A |
| 177 | 122.547 | 13.217 | 11 | N/A | N/A | N/A |
| 178 | 122.547 | 2.294 | 11 | N/A | N/A | N/A |
| 183 | 122.547 | 1.983 | 11 | N/A | N/A | N/A |
| 185 | 122.547 | 13.651 | 11 | N/A | N/A | N/A |
| 189 | 122.547 | 0.617 | 11 | N/A | N/A | N/A |
| 206 | 122.547 | 1.515 | 11 | N/A | N/A | N/A |
| 207 | 122.547 | 2.189 | 11 | N/A | N/A | N/A |
| 216 | 122.547 | 1.421 | 11 | N/A | N/A | N/A |
| 220 | 122.547 | 2.347 | 11 | N/A | N/A | N/A |
| 248 | 122.547 | 2.158 | 11 | N/A | N/A | N/A |
| 587 | 122.547 | 2.681 | 11 | N/A | N/A | N/A |
| 644 | 122.547 | 2.919 | 11 | N/A | N/A | N/A |
| 645 | 122.547 | 2.389 | 11 | N/A | N/A | N/A |
| 656 | 122.547 | 2.584 | 11 | N/A | N/A | N/A |
| 657 | 122.547 | 2.792 | 11 | N/A | N/A | N/A |
| 710 | 122.547 | 3.154 | 11 | N/A | N/A | N/A |
| 715 | 122.547 | 3.094 | 11 | N/A | N/A | N/A |
| 719 | 122.547 | 2.222 | 11 | N/A | N/A | N/A |
| 773 | 122.547 | 13.630 | 11 | N/A | N/A | N/A |
| 61 | 537.536 | 11.439 | 10 | N/A | N/A | N/A |
| 109 | 565.567 | 11.404 | 10 | N/A | N/A | N/A |
| 295 | 582.594 | 11.445 | 10 | N/A | N/A | N/A |
| 816 | 593.598 | 10.672 | 10 | N/A | N/A | N/A |
| 191 | 608.393 | 9.449 | 9 | N/A | N/A | N/A |
| 313 | 638.404 | 9.642 | 9 | N/A | 2-Linoleoyl-1-palmitoyl-sn-glycero-3-phosphoethanolamine | 0.770 |
| 348 | 624.388 | 9.076 | 9 | N/A | 2-Linoleoyl-1-palmitoyl-sn-glycero-3-phosphoethanolamine | 0.797 |
| 15 | 824.976 | 3.461 | 8 | N/A | N/A | N/A |
| 18 | 810.973 | 3.183 | 8 | N/A | N/A | N/A |
| 26 | 838.980 | 3.758 | 8 | N/A | N/A | N/A |
| 65 | 1215.954 | 3.181 | 8 | N/A | N/A | N/A |
| 87 | 1236.959 | 3.462 | 8 | N/A | N/A | N/A |
| 121 | 837.625 | 3.464 | 8 | N/A | N/A | N/A |
| 136 | 823.621 | 3.183 | 8 | N/A | N/A | N/A |
| 194 | 618.984 | 3.459 | 8 | N/A | N/A | N/A |
| 204 | 1257.965 | 3.758 | 8 | N/A | N/A | N/A |
| 240 | 852.983 | 4.057 | 8 | N/A | N/A | N/A |
| 244 | 846.306 | 3.758 | 8 | N/A | N/A | N/A |
| 253 | 1069.394 | 4.071 | 8 | N/A | N/A | N/A |
| 257 | 851.628 | 3.759 | 8 | N/A | N/A | N/A |
| 258 | 608.481 | 3.183 | 8 | N/A | N/A | N/A |
| 288 | 1083.397 | 4.311 | 8 | N/A | N/A | N/A |
| 289 | 629.487 | 3.760 | 8 | N/A | N/A | N/A |
| 312 | 1055.391 | 3.820 | 8 | N/A | N/A | N/A |
| 352 | 1234.935 | 3.184 | 8 | N/A | N/A | N/A |
| 406 | 802.298 | 4.071 | 8 | N/A | N/A | N/A |
| 445 | 812.801 | 4.312 | 8 | N/A | N/A | N/A |
| 450 | 1041.387 | 3.604 | 8 | N/A | N/A | N/A |
| 457 | 1255.942 | 3.463 | 8 | N/A | N/A | N/A |
| 497 | 1278.970 | 4.056 | 8 | N/A | N/A | N/A |
| 504 | 1268.955 | 3.761 | 8 | N/A | N/A | N/A |
| 511 | 865.632 | 4.058 | 8 | N/A | N/A | N/A |
| 514 | 639.989 | 4.058 | 8 | N/A | N/A | N/A |
| 515 | 860.310 | 4.057 | 8 | N/A | N/A | N/A |
| 518 | 1276.945 | 3.762 | 8 | N/A | N/A | N/A |
| 519 | 856.952 | 3.759 | 8 | N/A | N/A | N/A |
| 522 | 1090.726 | 4.311 | 8 | N/A | N/A | N/A |
| 531 | 1076.722 | 4.075 | 8 | N/A | N/A | N/A |
| 533 | 1096.048 | 4.311 | 8 | N/A | N/A | N/A |
| 534 | 791.796 | 3.823 | 8 | N/A | N/A | N/A |
| 537 | 810.973 | 3.375 | 8 | N/A | N/A | N/A |
| 559 | 824.976 | 3.689 | 8 | N/A | N/A | N/A |
| 561 | 1289.962 | 4.060 | 8 | N/A | N/A | N/A |
| 567 | 1082.043 | 4.072 | 8 | N/A | N/A | N/A |
| 588 | 1027.384 | 3.427 | 8 | N/A | N/A | N/A |
| 590 | 1013.379 | 3.335 | 8 | N/A | N/A | N/A |
| 610 | 770.790 | 3.427 | 8 | N/A | N/A | N/A |
| 614 | 837.291 | 3.461 | 8 | N/A | N/A | N/A |
| 620 | 760.288 | 3.338 | 8 | N/A | N/A | N/A |

| Feature identifiers | | | | | Analog Hits | |
|---|---|---|---|---|---|---|
| ClusterIdx | m/z | RT [min] | Subnetwork | Shared Peaks | Analog name | Cosine |
| 633 | 1062.719 | 3.822 | 8 | N/A | N/A | N/A |
| 650 | 629.236 | 3.761 | 8 | N/A | N/A | N/A |
| 659 | 813.049 | 4.312 | 8 | N/A | N/A | N/A |
| 664 | 866.987 | 4.353 | 8 | N/A | N/A | N/A |
| 672 | 1299.976 | 4.354 | 8 | N/A | N/A | N/A |
| 676 | 874.315 | 4.356 | 8 | N/A | N/A | N/A |
| 677 | 856.617 | 3.758 | 8 | N/A | N/A | N/A |
| 678 | 879.636 | 4.362 | 8 | N/A | N/A | N/A |
| 680 | 1297.948 | 4.058 | 8 | N/A | N/A | N/A |
| 683 | 975.108 | 4.244 | 8 | N/A | N/A | N/A |
| 685 | 1101.371 | 4.312 | 8 | N/A | N/A | N/A |
| 691 | 822.287 | 4.312 | 8 | N/A | N/A | N/A |
| 730 | 796.969 | 3.070 | 8 | N/A | N/A | N/A |
| 741 | 1068.040 | 3.824 | 8 | N/A | N/A | N/A |
| 746 | 1069.395 | 4.241 | 8 | N/A | N/A | N/A |
| 751 | 1221.446 | 3.186 | 8 | N/A | N/A | N/A |
| 757 | 1055.390 | 3.992 | 8 | N/A | N/A | N/A |
| 760 | 975.359 | 4.241 | 8 | N/A | N/A | N/A |
| 761 | 802.298 | 4.239 | 8 | N/A | N/A | N/A |
| 766 | 618.733 | 3.459 | 8 | N/A | N/A | N/A |
| 768 | 838.979 | 3.988 | 8 | N/A | N/A | N/A |
| 778 | 796.636 | 3.069 | 8 | N/A | N/A | N/A |
| 785 | 1215.954 | 3.392 | 8 | N/A | N/A | N/A |
| 787 | 608.481 | 3.384 | 8 | N/A | N/A | N/A |
| 788 | 1236.958 | 3.690 | 8 | N/A | N/A | N/A |
| 789 | 618.734 | 3.694 | 8 | N/A | N/A | N/A |
| 791 | 792.044 | 3.972 | 8 | N/A | N/A | N/A |
| 796 | 1069.727 | 4.238 | 8 | N/A | N/A | N/A |
| 800 | 824.643 | 3.674 | 8 | N/A | N/A | N/A |
| 801 | 1055.390 | 4.161 | 8 | N/A | N/A | N/A |
| 803 | 791.795 | 4.163 | 8 | N/A | N/A | N/A |
| 804 | 1300.145 | 4.240 | 8 | N/A | N/A | N/A |
| 805 | 810.973 | 3.576 | 8 | N/A | N/A | N/A |
| 807 | 1082.043 | 4.246 | 8 | N/A | N/A | N/A |
| 809 | 629.487 | 3.997 | 8 | N/A | N/A | N/A |
| 812 | 1268.461 | 3.759 | 8 | N/A | N/A | N/A |
| 820 | 1300.134 | 4.241 | 8 | N/A | N/A | N/A |
| 829 | 1137.663 | 4.195 | 8 | N/A | N/A | N/A |
| 830 | 1263.457 | 3.755 | 8 | N/A | N/A | N/A |
| 833 | 1076.722 | 4.246 | 8 | N/A | N/A | N/A |
| 836 | 1063.052 | 3.825 | 8 | N/A | N/A | N/A |
| 838 | 791.796 | 3.996 | 8 | N/A | N/A | N/A |
| 842 | 822.537 | 4.313 | 8 | N/A | N/A | N/A |
| 845 | 1257.962 | 3.997 | 8 | N/A | N/A | N/A |
| 847 | 781.292 | 3.601 | 8 | N/A | N/A | N/A |
| 848 | 811.784 | 4.077 | 8 | N/A | N/A | N/A |
| 851 | 1276.945 | 3.763 | 8 | N/A | N/A | N/A |
| 854 | 1194.947 | 3.069 | 8 | N/A | N/A | N/A |
| 862 | 964.605 | 4.154 | 8 | N/A | N/A | N/A |
| 354 | 828.611 | 3.182 | 7 | N/A | N/A | N/A |
| 360 | 842.615 | 3.460 | 7 | N/A | N/A | N/A |
| 649 | 842.948 | 3.459 | 7 | N/A | N/A | N/A |
| 811 | 828.945 | 3.185 | 7 | N/A | N/A | N/A |
| 22 | 274.274 | 6.821 | 6 | N/A | C17-Sphinganine | 0.702 |
| 104 | 302.305 | 7.664 | 6 | N/A | C17-Sphinganine | 0.780 |
| 277 | 330.337 | 8.495 | 6 | N/A | C17-Sphinganine | 0.774 |
| 341 | 358.368 | 9.237 | 6 | N/A | C17-Sphinganine | 0.756 |
| 321 | 311.185 | 7.063 | 5 | N/A | Spectral Match to Diethyl phthalate from NIST14 | 0.747 |
| 347 | 423.310 | 10.200 | 5 | N/A | N/A | N/A |
| 661 | 468.369 | 10.166 | 5 | N/A | N/A | N/A |
| 11 | 452.277 | 7.678 | 4 | N/A | 4-[5-[[4-[5-[acetyl(hydroxy)amino]pentylamino]-4-oxobutanoyl]-hydroxyamino]pentylamino]-4-oxobutanoic acid | 0.894 |
| 155 | 474.259 | 7.679 | 4 | N/A | 1-(9Z-Octadecenoyl)-sn-glycero-3-phosphoethanolamine | 0.825 |
| 238 | 474.258 | 7.808 | 4 | N/A | Spectral Match to 1-Hexadecanoyl-sn-glycero-3-phosphoethanolamine from NIST14 | 0.969 |
| 350 | 690.508 | 12.705 | 4 | N/A | 2-Linoleoyl-1-palmitoyl-sn-glycero-3-phosphoethanolamine | 0.911 |
| 424 | 511.304 | 8.562 | 4 | N/A | PE(18:1/0:0); [M+H]+ C23H47N1O7P1 | 0.880 |
| 486 | 690.508 | 10.348 | 4 | N/A | 2-Linoleoyl-1-palmitoyl-sn-glycero-3-phosphoethanolamine | 0.909 |
| 489 | 712.490 | 12.701 | 4 | N/A | 1,2-Di-(9Z-octadecenoyl)-sn-glycero-3-phosphoethanolamine | 0.885 |
| 505 | 712.490 | 10.356 | 4 | N/A | 1,2-Di-(9Z-octadecenoyl)-sn-glycero-3-phosphoethanolamine | 0.881 |
| 510 | 485.288 | 8.510 | 4 | N/A | Glycerophospho-N-oleoyl ethanolamine | 0.840 |
| 513 | 743.484 | 12.700 | 4 | N/A | 1,2-Dioleoyl-sn-glycero-3-phospho-rac-1-glycerol | 0.926 |

| Feature identifiers | | | | | Analog Hits | |
|---|---|---|---|---|---|---|
| ClusterIdx | m/z | RT [min] | Subnetwork | Shared Peaks | Analog name | Cosine |
| 516 | 483.272 | 7.795 | 4 | N/A | PE(16:1/0:0); [M+H]+ C21H43N1O7P1 | 0.787 |
| 517 | 740.522 | 12.702 | 4 | N/A | 1,2-Di-(9Z-octadecenoyl)-sn-glycero-3-phosphoethanolamine | 0.917 |
| 532 | 738.526 | 10.287 | 4 | N/A | PE(16:0/16:1); [M+H]+ C37H73N1O8P1 | 0.872 |
| 554 | 903.547 | 7.681 | 4 | N/A | N/A | N/A |
| 555 | 467.277 | 8.509 | 4 | N/A | 1-Hexadecanoyl-2-sn-glycero-3-phosphate | 0.812 |
| 562 | 743.484 | 10.306 | 4 | N/A | 1,2-Dioleoyl-sn-glycero-3-phospho-rac-1-glycerol | 0.919 |
| 565 | 496.304 | 8.835 | 4 | N/A | Spectral Match to 1-Palmitoyl-2-hydroxy-sn-glycero-3-phosphoethanolamine from NIST14 | 0.841 |
| 577 | 483.272 | 7.944 | 4 | N/A | Glycerophospho-N-oleoyl ethanolamine | 0.817 |
| 581 | 511.304 | 8.710 | 4 | N/A | 1-Oleoyl-L-.alpha.-lysophosphatidic acid | 0.859 |
| 583 | 452.277 | 7.806 | 4 | N/A | Spectral Match to 1-(9Z-Octadecenoyl)-sn-glycero-3-phosphoethanolamine from NIST14 | 0.966 |
| 639 | 690.508 | 10.220 | 4 | N/A | 2-Linoleoyl-1-palmitoyl-sn-glycero-3-phosphoethanolamine | 0.912 |
| 643 | 712.490 | 10.268 | 4 | N/A | 1,2-Di-(9Z-octadecenoyl)-sn-glycero-3-phosphoethanolamine | 0.892 |
| 669 | 466.293 | 8.072 | 4 | N/A | 4-[5-[[4-[5-[acetyl(hydroxy)amino]pentylamino]-4-oxobutanoyl]-hydroxyamino]pentylamino]-4-oxobutanoic acid | 0.866 |
| 679 | 466.293 | 8.206 | 4 | N/A | Spectral Match to 1-(9Z-Octadecenoyl)-sn-glycero-3-phosphoethanolamine from NIST14 | 0.967 |
| 701 | 440.278 | 7.956 | 4 | N/A | Spectral Match to 1-(9Z-Octadecenoyl)-sn-glycero-3-phosphoethanolamine from NIST14 | 0.950 |
| 713 | 662.477 | 12.666 | 4 | N/A | 2-Linoleoyl-1-palmitoyl-sn-glycero-3-phosphoethanolamine | 0.807 |
| 721 | 533.285 | 8.561 | 4 | N/A | 1-Oleoyl-L-.alpha.-lysophosphatidic acid | 0.887 |
| 722 | 440.278 | 7.818 | 4 | N/A | Spectral Match to 1-Palmitoyl-2-hydroxy-sn-glycero-3-phosphoethanolamine from NIST14 | 0.892 |
| 724 | 684.458 | 12.702 | 4 | N/A | 1,2-Di-(9Z-octadecenoyl)-sn-glycero-3-phosphoethanolamine | 0.860 |
| 727 | 465.262 | 7.796 | 4 | N/A | 1-Oleoyl-L-.alpha.-lysophosphatidic acid | 0.783 |
| 728 | 505.254 | 7.795 | 4 | N/A | Dipalmitolein (9c) | 0.863 |
| 729 | 507.269 | 8.510 | 4 | N/A | 1-Hexadecanoyl-2-sn-glycero-3-phosphate | 0.792 |
| 731 | 493.293 | 8.564 | 4 | N/A | 1-Oleoyl-L-.alpha.-lysophosphatidic acid | 0.861 |
| 736 | 676.492 | 12.707 | 4 | N/A | Spectral Match to 2-Linoleoyl-1-palmitoyl-sn-glycero-3-phosphoethanolamine from NIST14 | 0.825 |
| 745 | 493.293 | 8.710 | 4 | N/A | 1-Oleoyl-L-.alpha.-lysophosphatidic acid | 0.862 |
| 749 | 500.300 | 7.793 | 4 | N/A | PE(16:1/0:0); [M+H]+ C21H43N1O7P1 | 0.798 |
| 756 | 533.285 | 8.715 | 4 | N/A | 1-Oleoyl-L-.alpha.-lysophosphatidic acid | 0.819 |
| 775 | 704.523 | 12.703 | 4 | N/A | 1-Hexadecanoyl-2-(9Z-octadecenoyl)-sn-glycero-3-phosphoethanolamine | 0.888 |
| 779 | 704.523 | 10.367 | 4 | N/A | 1-Hexadecanoyl-2-(9Z-octadecenoyl)-sn-glycero-3-phosphoethanolamine | 0.890 |
| 781 | 726.506 | 12.701 | 4 | N/A | 1,2-Di-(9Z-octadecenoyl)-sn-glycero-3-phosphoethanolamine | 0.887 |
| 813 | 494.288 | 8.224 | 4 | N/A | 1-(9Z-Octadecenoyl)-sn-glycero-3-phosphoethanolamine | 0.763 |
| 825 | 664.490 | 9.773 | 4 | N/A | 2-Linoleoyl-1-palmitoyl-sn-glycero-3-phosphoethanolamine | 0.918 |
| 837 | 766.559 | 12.703 | 4 | N/A | N/A | N/A |
| 841 | 676.491 | 10.424 | 4 | N/A | 2-Linoleoyl-1-palmitoyl-sn-glycero-3-phosphoethanolamine | 0.817 |
| 844 | 698.475 | 10.420 | 4 | N/A | 1,2-Di-(9Z-octadecenoyl)-sn-glycero-3-phosphoethanolamine | 0.883 |
| 853 | 1380.008 | 10.186 | 4 | N/A | N/A | N/A |
| 855 | 1380.006 | 9.800 | 4 | N/A | N/A | N/A |
| 27 | 316.212 | 3.277 | 3 | N/A | N/A | N/A |
| 37 | 430.244 | 4.459 | 3 | N/A | N/A | N/A |
| 55 | 300.200 | 3.938 | 3 | N/A | N/A | N/A |
| 69 | 416.228 | 3.930 | 3 | N/A | N/A | N/A |
| 122 | 243.183 | 3.050 | 3 | N/A | N/A | N/A |
| 124 | 334.178 | 3.942 | 3 | N/A | N/A | N/A |
| 128 | 357.214 | 4.538 | 3 | N/A | N/A | N/A |
| 149 | 350.208 | 4.208 | 3 | N/A | N/A | N/A |
| 167 | 599.391 | 3.938 | 3 | N/A | N/A | N/A |
| 190 | 294.861 | 4.052 | 3 | N/A | N/A | N/A |
| 224 | 302.196 | 0.594 | 3 | N/A | N/A | N/A |
| 246 | 309.181 | 4.412 | 3 | N/A | N/A | N/A |
| 268 | 328.200 | 4.325 | 3 | N/A | N/A | N/A |
| 290 | 272.185 | 3.424 | 3 | N/A | N/A | N/A |
| 293 | 293.191 | 3.577 | 3 | N/A | N/A | N/A |
| 306 | 300.180 | 5.792 | 3 | N/A | N/A | N/A |
| 322 | 713.423 | 4.538 | 3 | N/A | N/A | N/A |
| 328 | 252.166 | 3.641 | 3 | N/A | N/A | N/A |
| 329 | 332.872 | 4.600 | 3 | N/A | N/A | N/A |
| 343 | 471.307 | 6.113 | 3 | N/A | N/A | N/A |
| 361 | 441.788 | 4.079 | 3 | N/A | N/A | N/A |
| 386 | 241.204 | 5.244 | 3 | N/A | N/A | N/A |
| 396 | 469.328 | 4.897 | 3 | N/A | N/A | N/A |
| 401 | 256.850 | 3.715 | 3 | N/A | N/A | N/A |
| 404 | 699.407 | 4.204 | 3 | N/A | N/A | N/A |
| 413 | 292.192 | 4.464 | 3 | N/A | N/A | N/A |
| 439 | 585.375 | 3.578 | 3 | N/A | N/A | N/A |
| 443 | 384.773 | 3.708 | 3 | N/A | N/A | N/A |
| 463 | 293.191 | 3.406 | 3 | N/A | N/A | N/A |
| 499 | 599.391 | 3.791 | 3 | N/A | N/A | N/A |
| 526 | 485.359 | 3.054 | 3 | N/A | N/A | N/A |
| 527 | 441.788 | 4.168 | 3 | N/A | N/A | N/A |
| 530 | 290.193 | 3.898 | 3 | N/A | N/A | N/A |

| Feature identifiers | | | | | Analog Hits | |
|---|---|---|---|---|---|---|
| ClusterIdx | m/z | RT [min] | Subnetwork | Shared Peaks | Analog name | Cosine |
| 553 | 332.207 | 4.839 | 3 | N/A | N/A | N/A |
| 568 | 379.210 | 4.343 | 3 | N/A | N/A | N/A |
| 623 | 343.200 | 3.890 | 3 | N/A | N/A | N/A |
| 624 | 263.684 | 4.125 | 3 | N/A | N/A | N/A |
| 627 | 230.153 | 3.714 | 3 | N/A | N/A | N/A |
| 629 | 347.535 | 4.414 | 3 | N/A | N/A | N/A |
| 747 | 540.426 | 9.375 | 3 | N/A | N/A | N/A |
| 750 | 568.458 | 9.844 | 3 | N/A | N/A | N/A |
| 758 | 263.684 | 4.034 | 3 | N/A | N/A | N/A |
| 782 | 220.150 | 0.536 | 3 | N/A | N/A | N/A |
| 784 | 498.804 | 4.596 | 3 | N/A | N/A | N/A |
| 810 | 294.861 | 4.145 | 3 | N/A | N/A | N/A |
| 817 | 384.772 | 3.711 | 3 | N/A | N/A | N/A |
| 819 | 292.192 | 4.644 | 3 | N/A | N/A | N/A |
| 93 | 256.264 | 9.789 | 2 | N/A | N/A | N/A |
| 142 | 284.295 | 10.325 | 2 | N/A | Spectral Match to Palmitamide from NIST14 | 0.789 |
| 175 | 308.279 | 8.430 | 2 | N/A | N/A | N/A |
| 218 | 482.408 | 9.562 | 2 | N/A | Spectral Match to Nonaethylene glycol from NIST14 | 0.780 |
| 226 | 526.434 | 9.538 | 2 | N/A | Spectral Match to Decaethylene glycol from NIST14 | 0.789 |
| 228 | 336.311 | 9.373 | 2 | N/A | N/A | N/A |
| 229 | 291.253 | 8.431 | 2 | N/A | Tetraethylene glycol | 0.801 |
| 241 | 424.364 | 9.342 | 2 | N/A | N/A | N/A |
| 242 | 380.338 | 9.364 | 2 | N/A | N/A | N/A |
| 247 | 352.306 | 8.443 | 2 | N/A | N/A | N/A |
| 269 | 438.382 | 9.584 | 2 | N/A | Spectral Match to Nonaethylene glycol from NIST14 | 0.726 |
| 270 | 408.372 | 10.006 | 2 | N/A | N/A | N/A |
| 276 | 570.460 | 9.514 | 2 | N/A | Spectral Match to Undecaethylene glycol from NIST14 | 0.763 |
| 278 | 512.417 | 9.299 | 2 | N/A | Spectral Match to Nonaethylene glycol from NIST14 | 0.738 |
| 280 | 468.390 | 9.320 | 2 | N/A | N/A | N/A |
| 291 | 366.322 | 8.959 | 2 | N/A | N/A | N/A |
| 294 | 452.398 | 9.985 | 2 | N/A | N/A | N/A |
| 297 | 556.443 | 9.273 | 2 | N/A | Spectral Match to Decaethylene glycol from NIST14 | 0.739 |
| 302 | 247.227 | 8.431 | 2 | N/A | N/A | N/A |
| 303 | 392.338 | 10.385 | 2 | N/A | N/A | N/A |
| 310 | 496.424 | 9.963 | 2 | N/A | N/A | N/A |
| 336 | 364.344 | 10.020 | 2 | N/A | N/A | N/A |
| 337 | 540.449 | 9.937 | 2 | N/A | Spectral Match to Decaethylene glycol from NIST14 | 0.702 |
| 344 | 476.301 | 4.018 | 2 | N/A | Spectral Match to Undecaethylene glycol from NIST14 | 0.857 |
| 349 | 440.359 | 8.424 | 2 | N/A | N/A | N/A |
| 359 | 600.469 | 9.251 | 2 | N/A | Spectral Match to Undecaethylene glycol from NIST14 | 0.738 |
| 362 | 322.295 | 8.962 | 2 | N/A | N/A | N/A |
| 364 | 614.485 | 9.493 | 2 | N/A | Spectral Match to Decaethylene glycol tridecyl ether from NIST14 | 0.728 |
| 366 | 396.333 | 8.432 | 2 | N/A | N/A | N/A |
| 367 | 422.389 | 10.267 | 2 | N/A | N/A | N/A |
| 369 | 436.364 | 10.348 | 2 | N/A | N/A | N/A |
| 377 | 422.347 | 10.141 | 2 | N/A | N/A | N/A |
| 382 | 466.373 | 10.105 | 2 | N/A | N/A | N/A |
| 383 | 319.284 | 9.373 | 2 | N/A | Triethylene glycol | 0.828 |
| 387 | 466.415 | 10.248 | 2 | N/A | N/A | N/A |
| 389 | 526.434 | 9.632 | 2 | N/A | Spectral Match to Decaethylene glycol from NIST14 | 0.787 |
| 390 | 468.390 | 9.206 | 2 | N/A | Spectral Match to Nonaethylene glycol from NIST14 | 0.752 |
| 399 | 520.331 | 4.142 | 2 | N/A | Spectral Match to Decaethylene glycol from NIST14 | 0.838 |
| 400 | 484.385 | 8.410 | 2 | N/A | Spectral Match to Nonaethylene glycol from NIST14 | 0.711 |
| 412 | 512.416 | 9.184 | 2 | N/A | Spectral Match to Nonaethylene glycol from NIST14 | 0.822 |
| 414 | 528.411 | 8.398 | 2 | N/A | Spectral Match to Decaethylene glycol from NIST14 | 0.710 |
| 416 | 564.360 | 4.248 | 2 | N/A | Spectral Match to Undecaethylene glycol from NIST14 | 0.834 |
| 417 | 644.495 | 9.227 | 2 | N/A | Spectral Match to Decaethylene glycol tridecyl ether from NIST14 | 0.701 |
| 422 | 480.390 | 10.316 | 2 | N/A | Spectral Match to Nonaethylene glycol from NIST14 | 0.701 |
| 432 | 556.443 | 9.161 | 2 | N/A | Spectral Match to Decaethylene glycol from NIST14 | 0.835 |
| 438 | 584.475 | 9.912 | 2 | N/A | Spectral Match to Undecaethylene glycol from NIST14 | 0.725 |
| 455 | 628.501 | 9.881 | 2 | N/A | N/A | N/A |
| 464 | 608.392 | 4.346 | 2 | N/A | N/A | N/A |
| 471 | 352.306 | 8.095 | 2 | N/A | N/A | N/A |
| 528 | 308.279 | 8.075 | 2 | N/A | N/A | N/A |
| 539 | 688.521 | 9.203 | 2 | N/A | N/A | N/A |
| 541 | 363.310 | 9.363 | 2 | N/A | Spectral Match to Hexaethylene glycol from NIST14 | 0.826 |
| 546 | 291.253 | 8.075 | 2 | N/A | N/A | N/A |
| 548 | 396.333 | 8.097 | 2 | N/A | N/A | N/A |
| 556 | 732.548 | 9.180 | 2 | N/A | N/A | N/A |
| 564 | 440.359 | 8.096 | 2 | N/A | N/A | N/A |

| Feature identifiers | | | | | Analog Hits | |
|---|---|---|---|---|---|---|
| ClusterIdx | m/z | RT [min] | Subnetwork | Shared Peaks | Analog name | Cosine |
| 570 | 484.385 | 8.094 | 2 | N/A | N/A | N/A |
| 571 | 308.279 | 8.290 | 2 | N/A | N/A | N/A |
| 573 | 528.411 | 8.088 | 2 | N/A | N/A | N/A |
| 576 | 247.227 | 8.064 | 2 | N/A | Spectral Match to Ethanol, 2-(2-butoxyethoxy)- from NIST14 | 0.827 |
| 579 | 335.280 | 8.098 | 2 | N/A | Spectral Match to Hexaethylene glycol from NIST14 | 0.774 |
| 580 | 572.437 | 8.084 | 2 | N/A | Spectral Match to Undecaethylene glycol from NIST14 | 0.703 |
| 598 | 396.333 | 8.315 | 2 | N/A | N/A | N/A |
| 604 | 440.359 | 8.302 | 2 | N/A | N/A | N/A |
| 608 | 672.527 | 9.854 | 2 | N/A | N/A | N/A |
| 615 | 350.328 | 9.687 | 2 | N/A | N/A | N/A |
| 618 | 658.511 | 9.464 | 2 | N/A | Spectral Match to Decaethylene glycol tridecyl ether from NIST14 | 0.737 |
| 625 | 702.536 | 9.449 | 2 | N/A | N/A | N/A |
| 647 | 394.355 | 9.697 | 2 | N/A | N/A | N/A |
| 50 | 701.620 | 10.613 | 1 | N/A | N/A | N/A |
| 199 | 419.364 | 7.499 | 1 | N/A | N/A | N/A |
| 689 | 685.624 | 12.053 | 1 | N/A | N/A | N/A |
| 1 | 110.020 | 12.761 | -1 | N/A | N/A | N/A |
| 8 | 437.194 | 7.524 | -1 | N/A | N/A | N/A |
| 10 | 120.081 | 0.519 | -1 | N/A | Massbank:PB000474 Tyramine\|4-(2-aminoethyl)phenol | 0.946 |
| 13 | 242.284 | 8.651 | -1 | N/A | Tetradecylamine | 0.958 |
| 14 | 851.398 | 7.527 | -1 | N/A | N/A | N/A |
| 32 | 437.194 | 7.249 | -1 | N/A | N/A | N/A |
| 33 | 559.518 | 11.442 | -1 | N/A | N/A | N/A |
| 34 | 867.372 | 7.528 | -1 | N/A | N/A | N/A |
| 41 | 587.549 | 11.407 | -1 | N/A | N/A | N/A |
| 47 | 163.991 | 12.748 | -1 | N/A | N/A | N/A |
| 53 | 243.088 | 4.621 | -1 | N/A | N/A | N/A |
| 54 | 399.209 | 10.100 | -1 | N/A | N/A | N/A |
| 57 | 685.436 | 12.425 | -1 | N/A | N/A | N/A |
| 58 | 250.112 | 5.753 | -1 | N/A | 3-(3-methylbut-2-enyl)purin-6-amine | 0.781 |
| 64 | 255.073 | 4.542 | -1 | N/A | N/A | N/A |
| 66 | 261.109 | 5.951 | -1 | N/A | N/A | N/A |
| 71 | 267.077 | 3.607 | -1 | N/A | N/A | N/A |
| 73 | 453.168 | 7.250 | -1 | N/A | N/A | N/A |
| 76 | 324.142 | 7.584 | -1 | N/A | N/A | N/A |
| 77 | 372.882 | 12.723 | -1 | N/A | N/A | N/A |
| 79 | 321.131 | 4.460 | -1 | N/A | N/A | N/A |
| 82 | 269.015 | 5.941 | -1 | N/A | N/A | N/A |
| 83 | 230.141 | 12.721 | -1 | N/A | NCGC00385962-01_C16H24O3 | 0.730 |
| 85 | 233.079 | 4.825 | -1 | N/A | N/A | N/A |
| 86 | 239.089 | 4.007 | -1 | N/A | N/A | N/A |
| 90 | 332.913 | 12.737 | -1 | N/A | N/A | N/A |
| 98 | 195.102 | 7.496 | -1 | N/A | MassbankEU:SM862301 Benzocaine\|Benzocain\|ethyl 4-aminobenzoate | 0.771 |
| 100 | 440.099 | 7.644 | -1 | N/A | N/A | N/A |
| 101 | 235.169 | 8.489 | -1 | N/A | N/A | N/A |
| 106 | 670.153 | 5.750 | -1 | N/A | N/A | N/A |
| 107 | 183.150 | 4.641 | -1 | N/A | N/A | N/A |
| 108 | 281.088 | 8.127 | -1 | N/A | N/A | N/A |
| 111 | 262.165 | 4.000 | -1 | N/A | N/A | N/A |
| 114 | 244.191 | 5.975 | -1 | N/A | 1,11-Undecanedicarboxylic acid | 0.941 |
| 116 | 228.232 | 9.984 | -1 | N/A | N/A | N/A |
| 118 | 219.175 | 8.820 | -1 | N/A | Spectral Match to 7.alpha.,24(S)-Dihydroxy-4-cholesten-3-one from NIST14 | 0.760 |
| 126 | 296.219 | 7.149 | -1 | N/A | N/A | N/A |
| 137 | 431.183 | 10.115 | -1 | N/A | N/A | N/A |
| 148 | 319.187 | 7.588 | -1 | N/A | N/A | N/A |
| 152 | 707.875 | 4.385 | -1 | N/A | N/A | N/A |
| 162 | 174.999 | 12.765 | -1 | N/A | N/A | N/A |
| 164 | 338.147 | 5.707 | -1 | N/A | Huperzine A | 0.791 |
| 165 | 269.210 | 8.432 | -1 | N/A | N/A | N/A |
| 173 | 451.246 | 9.774 | -1 | N/A | N/A | N/A |
| 179 | 723.608 | 10.615 | -1 | N/A | N/A | N/A |
| 184 | 434.186 | 7.527 | -1 | N/A | N/A | N/A |
| 188 | 270.279 | 10.247 | -1 | N/A | N/A | N/A |
| 193 | 242.284 | 7.258 | -1 | N/A | N/A | N/A |
| 195 | 312.326 | 10.525 | -1 | N/A | N/A | N/A |
| 198 | 499.165 | 7.526 | -1 | N/A | N/A | N/A |
| 203 | 122.547 | 1.761 | -1 | N/A | N/A | N/A |
| 212 | 409.201 | 10.114 | -1 | N/A | N/A | N/A |
| 214 | 270.279 | 9.930 | -1 | N/A | N/A | N/A |
| 217 | 266.248 | 6.949 | -1 | N/A | N/A | N/A |

| Feature identifiers | | | | | Analog Hits | |
|---|---|---|---|---|---|---|
| ClusterIdx | m/z | RT [min] | Subnetwork | Shared Peaks | Analog name | Cosine |
| 222 | 495.300 | 10.647 | -1 | N/A | N/A | N/A |
| 225 | 330.300 | 9.151 | -1 | N/A | Spectral Match to R-Palmitoyl-(1-methyl) ethanolamide from NIST14 | 0.971 |
| 232 | 250.178 | 6.460 | -1 | N/A | N/A | N/A |
| 234 | 555.250 | 4.014 | -1 | N/A | N/A | N/A |
| 236 | 149.988 | 12.784 | -1 | N/A | N/A | N/A |
| 237 | 640.348 | 4.374 | -1 | N/A | N/A | N/A |
| 252 | 157.084 | 0.657 | -1 | N/A | N/A | N/A |
| 254 | 111.039 | 13.492 | -1 | N/A | N/A | N/A |
| 256 | 297.241 | 9.395 | -1 | N/A | N/A | N/A |
| 259 | 249.185 | 8.504 | -1 | N/A | N/A | N/A |
| 260 | 290.269 | 6.924 | -1 | N/A | N/A | N/A |
| 265 | 535.153 | 8.517 | -1 | N/A | N/A | N/A |
| 267 | 453.168 | 7.817 | -1 | N/A | N/A | N/A |
| 271 | 283.104 | 5.343 | -1 | N/A | N/A | N/A |
| 274 | 537.311 | 10.526 | -1 | N/A | N/A | N/A |
| 279 | 149.024 | 6.761 | -1 | N/A | N/A | N/A |
| 281 | 467.147 | 6.891 | -1 | N/A | N/A | N/A |
| 285 | 408.309 | 10.166 | -1 | N/A | N/A | N/A |
| 286 | 291.195 | 8.162 | -1 | N/A | N/A | N/A |
| 287 | 573.110 | 8.520 | -1 | N/A | N/A | N/A |
| 299 | 437.194 | 8.754 | -1 | N/A | N/A | N/A |
| 300 | 604.812 | 4.484 | -1 | N/A | N/A | N/A |
| 304 | 365.137 | 8.808 | -1 | N/A | N/A | N/A |
| 305 | 480.273 | 3.605 | -1 | N/A | N/A | N/A |
| 308 | 659.288 | 10.020 | -1 | N/A | N/A | N/A |
| 309 | 165.076 | 4.458 | -1 | N/A | Erythritol | 0.862 |
| 320 | 341.267 | 9.373 | -1 | N/A | N/A | N/A |
| 324 | 270.315 | 9.426 | -1 | N/A | N/A | N/A |
| 325 | 326.378 | 9.409 | -1 | N/A | N/A | N/A |
| 327 | 131.049 | 0.513 | -1 | N/A | N/A | N/A |
| 331 | 260.165 | 6.499 | -1 | N/A | N/A | N/A |
| 334 | 469.184 | 6.409 | -1 | N/A | N/A | N/A |
| 335 | 469.163 | 6.314 | -1 | N/A | N/A | N/A |
| 338 | 499.780 | 3.714 | -1 | N/A | N/A | N/A |
| 339 | 615.581 | 10.604 | -1 | N/A | N/A | N/A |
| 340 | 354.553 | 5.749 | -1 | N/A | N/A | N/A |
| 345 | 236.142 | 5.355 | -1 | N/A | N/A | N/A |
| 351 | 451.173 | 6.881 | -1 | N/A | N/A | N/A |
| 356 | 310.173 | 6.844 | -1 | N/A | N/A | N/A |
| 365 | 519.261 | 9.835 | -1 | N/A | N/A | N/A |
| 368 | 273.167 | 4.221 | -1 | N/A | N/A | N/A |
| 376 | 288.181 | 4.136 | -1 | N/A | N/A | N/A |
| 379 | 496.301 | 10.257 | -1 | N/A | N/A | N/A |
| 380 | 497.280 | 9.835 | -1 | N/A | N/A | N/A |
| 385 | 188.069 | 0.519 | -1 | N/A | N/A | N/A |
| 395 | 721.506 | 12.424 | -1 | N/A | N/A | N/A |
| 397 | 459.263 | 10.315 | -1 | N/A | N/A | N/A |
| 398 | 445.291 | 10.157 | -1 | N/A | N/A | N/A |
| 403 | 763.517 | 12.425 | -1 | N/A | N/A | N/A |
| 405 | 528.788 | 4.417 | -1 | N/A | N/A | N/A |
| 407 | 455.335 | 10.125 | -1 | N/A | N/A | N/A |
| 410 | 830.949 | 3.185 | -1 | N/A | N/A | N/A |
| 421 | 514.951 | 4.135 | -1 | N/A | N/A | N/A |
| 423 | 409.180 | 7.399 | -1 | N/A | N/A | N/A |
| 425 | 707.607 | 12.056 | -1 | N/A | N/A | N/A |
| 428 | 366.633 | 3.310 | -1 | N/A | N/A | N/A |
| 430 | 379.862 | 4.127 | -1 | N/A | N/A | N/A |
| 436 | 267.137 | 8.329 | -1 | N/A | Tributyl phosphate | 0.914 |
| 449 | 658.341 | 3.958 | -1 | N/A | N/A | N/A |
| 451 | 254.660 | 3.065 | -1 | N/A | N/A | N/A |
| 453 | 441.346 | 7.492 | -1 | N/A | N/A | N/A |
| 454 | 409.163 | 6.914 | -1 | N/A | N/A | N/A |
| 470 | 469.163 | 6.021 | -1 | N/A | N/A | N/A |
| 474 | 501.170 | 0.519 | -1 | N/A | N/A | N/A |
| 485 | 517.142 | 0.507 | -1 | N/A | N/A | N/A |
| 496 | 226.012 | 12.739 | -1 | N/A | N/A | N/A |
| 502 | 102.034 | 0.568 | -1 | N/A | N/A | N/A |
| 503 | 228.207 | 4.644 | -1 | N/A | N/A | N/A |
| 508 | 1011.286 | 0.501 | -1 | N/A | N/A | N/A |
| 509 | 385.095 | 3.893 | -1 | N/A | N/A | N/A |

| Feature identifiers | | | | | Analog Hits | |
|---|---|---|---|---|---|---|
| ClusterIdx | m/z | RT [min] | Subnetwork | Shared Peaks | Analog name | Cosine |
| 525 | 269.210 | 8.072 | -1 | N/A | N/A | N/A |
| 538 | 844.952 | 3.465 | -1 | N/A | N/A | N/A |
| 543 | 453.188 | 6.313 | -1 | N/A | N/A | N/A |
| 547 | 372.717 | 12.740 | -1 | N/A | N/A | N/A |
| 549 | 287.114 | 4.597 | -1 | N/A | N/A | N/A |
| 551 | 385.294 | 9.364 | -1 | N/A | N/A | N/A |
| 557 | 858.957 | 3.760 | -1 | N/A | N/A | N/A |
| 563 | 429.319 | 9.342 | -1 | N/A | N/A | N/A |
| 566 | 472.175 | 4.688 | -1 | N/A | N/A | N/A |
| 569 | 294.074 | 0.519 | -1 | N/A | N/A | N/A |
| 575 | 453.189 | 6.022 | -1 | N/A | N/A | N/A |
| 586 | 191.086 | 3.681 | -1 | N/A | N/A | N/A |
| 592 | 359.104 | 3.680 | -1 | N/A | N/A | N/A |
| 593 | 511.165 | 3.612 | -1 | N/A | N/A | N/A |
| 594 | 700.315 | 10.020 | -1 | N/A | N/A | N/A |
| 597 | 225.149 | 6.492 | -1 | N/A | N/A | N/A |
| 599 | 131.049 | 0.515 | -1 | N/A | N/A | N/A |
| 601 | 357.262 | 8.098 | -1 | N/A | N/A | N/A |
| 602 | 303.138 | 3.688 | -1 | N/A | N/A | N/A |
| 605 | 347.222 | 7.583 | -1 | N/A | N/A | N/A |
| 606 | 438.143 | 7.111 | -1 | N/A | 2'-Deoxyguanosine 5'-diphosphate | 0.716 |
| 607 | 851.398 | 7.250 | -1 | N/A | N/A | N/A |
| 609 | 830.614 | 3.188 | -1 | N/A | N/A | N/A |
| 612 | 863.046 | 3.487 | -1 | N/A | N/A | N/A |
| 613 | 619.272 | 4.460 | -1 | N/A | N/A | N/A |
| 617 | 454.172 | 7.812 | -1 | N/A | N/A | N/A |
| 621 | 815.644 | 3.349 | -1 | N/A | N/A | N/A |
| 626 | 331.209 | 4.774 | -1 | N/A | N/A | N/A |
| 628 | 103.054 | 0.515 | -1 | N/A | N/A | N/A |
| 641 | 158.154 | 6.250 | -1 | N/A | N/A | N/A |
| 642 | 165.114 | 12.714 | -1 | N/A | 2Aminomethylpyrimidinehydrochloride | 0.905 |
| 648 | 425.137 | 6.913 | -1 | N/A | N/A | N/A |
| 651 | 267.157 | 4.332 | -1 | N/A | N/A | N/A |
| 655 | 290.232 | 4.304 | -1 | N/A | N/A | N/A |
| 660 | 391.205 | 10.042 | -1 | N/A | N/A | N/A |
| 673 | 870.956 | 4.058 | -1 | N/A | N/A | N/A |
| 682 | 425.215 | 9.512 | -1 | N/A | N/A | N/A |
| 684 | 612.425 | 0.524 | -1 | N/A | N/A | N/A |
| 686 | 231.081 | 0.527 | -1 | N/A | Diaminopimelic acid | 0.891 |
| 688 | 726.463 | 12.425 | -1 | N/A | N/A | N/A |
| 690 | 403.233 | 9.514 | -1 | N/A | N/A | N/A |
| 694 | 111.020 | 12.539 | -1 | N/A | N/A | N/A |
| 695 | 351.102 | 8.964 | -1 | N/A | N/A | N/A |
| 700 | 164.051 | 4.799 | -1 | N/A | 4-formyl Indole | 0.991 |
| 702 | 111.039 | 2.302 | -1 | N/A | N/A | N/A |
| 705 | 271.031 | 5.195 | -1 | N/A | N/A | N/A |
| 706 | 249.050 | 5.192 | -1 | N/A | N/A | N/A |
| 707 | 279.060 | 6.671 | -1 | N/A | N/A | N/A |
| 708 | 162.035 | 6.813 | -1 | N/A | N/A | N/A |
| 709 | 177.083 | 2.375 | -1 | N/A | N/A | N/A |
| 711 | 164.051 | 2.397 | -1 | N/A | N/A | N/A |
| 712 | 245.070 | 2.402 | -1 | N/A | N/A | N/A |
| 716 | 111.039 | 2.354 | -1 | N/A | N/A | N/A |
| 720 | 495.251 | 4.182 | -1 | N/A | N/A | N/A |
| 723 | 162.073 | 2.320 | -1 | N/A | N/A | N/A |
| 725 | 1347.881 | 12.426 | -1 | N/A | N/A | N/A |
| 733 | 297.083 | 6.817 | -1 | N/A | N/A | N/A |
| 734 | 471.252 | 7.681 | -1 | N/A | N/A | N/A |
| 735 | 597.678 | 0.494 | -1 | N/A | N/A | N/A |
| 737 | 701.411 | 12.427 | -1 | N/A | N/A | N/A |
| 738 | 253.602 | 7.678 | -1 | N/A | N/A | N/A |
| 739 | 749.548 | 12.237 | -1 | N/A | N/A | N/A |
| 742 | 287.650 | 8.328 | -1 | N/A | N/A | N/A |
| 744 | 247.129 | 0.492 | -1 | N/A | 5-L-Glutamyl-L-alanine | 0.917 |
| 748 | 654.456 | 10.203 | -1 | N/A | N/A | N/A |
| 752 | 286.520 | 6.796 | -1 | N/A | N/A | N/A |
| 753 | 335.125 | 8.963 | -1 | N/A | N/A | N/A |
| 754 | 533.934 | 3.609 | -1 | N/A | N/A | N/A |
| 762 | 533.282 | 9.675 | -1 | N/A | N/A | N/A |
| 763 | 1194.349 | 0.496 | -1 | N/A | N/A | N/A |

| Feature identifiers | | | | | Analog Hits | |
|---|---|---|---|---|---|---|
| ClusterIdx | m/z | RT [min] | Subnetwork | Shared Peaks | Analog name | Cosine |
| 764 | 616.651 | 0.503 | -1 | N/A | N/A | N/A |
| 765 | 223.064 | 6.361 | -1 | N/A | N/A | N/A |
| 769 | 498.173 | 7.669 | -1 | N/A | N/A | N/A |
| 771 | 143.107 | 1.088 | -1 | N/A | N/A | N/A |
| 786 | 819.476 | 8.754 | -1 | N/A | Spectral Match to Tris(2-butoxyethyl) phosphate from NIST14 | 0.822 |
| 794 | 811.308 | 6.909 | -1 | N/A | N/A | N/A |
| 797 | 818.299 | 3.374 | -1 | N/A | N/A | N/A |
| 806 | 618.984 | 3.690 | -1 | N/A | N/A | N/A |
| 808 | 832.302 | 3.693 | -1 | N/A | N/A | N/A |
| 814 | 513.189 | 7.230 | -1 | N/A | N/A | N/A |
| 815 | 432.241 | 7.420 | -1 | N/A | N/A | N/A |
| 818 | 123.079 | 0.380 | -1 | N/A | N/A | N/A |
| 822 | 376.087 | 6.788 | -1 | N/A | N/A | N/A |
| 827 | 326.112 | 5.859 | -1 | N/A | N/A | N/A |
| 828 | 319.064 | 6.836 | -1 | N/A | N/A | N/A |
| 831 | 1295.589 | 10.022 | -1 | N/A | N/A | N/A |
| 834 | 628.471 | 3.460 | -1 | N/A | N/A | N/A |
| 835 | 850.609 | 3.462 | -1 | N/A | N/A | N/A |
| 839 | 372.717 | 10.091 | -1 | N/A | N/A | N/A |
| 846 | 681.132 | 5.759 | -1 | N/A | N/A | N/A |
| 856 | 1194.817 | 11.488 | -1 | N/A | N/A | N/A |
| 857 | 804.296 | 3.067 | -1 | N/A | N/A | N/A |
| 863 | 353.267 | 9.157 | -1 | N/A | N/A | N/A |

| Feature identifiers | | | | | | Average XIC | | | | | | | Standard Err of Average XIC | | | | | | |
|---|---|---|---|---|---|---|---|---|---|---|---|---|---|---|---|---|---|---|---|
| ClusterIdx | m/z | RT [min] | Subnetwork | PPM | MassDiff | TUB00 | 4TUB34 | 4TUB81 | 4TUB93 | 5TUB23 | 5TUB48 | 5TUB83 | TUB00 | 4TUB34 | 4TUB81 | 4TUB93 | 5TUB23 | 5TUB48 | 5TUB83 |
| 491 | 205.195 | 9.007 | -1 | 1.264E+00 | 0.000 | 1.909E+04 | 2.137E+06 | 3.903E+05 | 6.898E+06 | 4.447E+03 | 5.214E+03 | 2.757E+03 | 8.503E+03 | 8.003E+05 | 1.416E+05 | 3.158E+06 | 8.547E+02 | 3.885E+03 | 1.728E+03 |
| 24 | 399.252 | 8.753 | 58 | 2.522E+00 | 0.001 | 4.232E+07 | 6.262E+07 | 4.260E+07 | 9.864E+07 | 4.716E+07 | 5.358E+07 | 5.810E+07 | 2.333E+06 | 7.571E+06 | 4.285E+05 | 5.530E+07 | 4.138E+06 | 5.506E+06 | 1.063E+07 |
| 44 | 421.232 | 8.753 | -1 | 1.811E+00 | 0.001 | 1.393E+07 | 2.184E+07 | 1.364E+07 | 3.428E+07 | 1.565E+07 | 1.656E+07 | 1.470E+07 | 5.378E+05 | 2.805E+06 | 3.351E+05 | 1.970E+07 | 9.327E+05 | 2.145E+06 | 7.994E+05 |
| 130 | 242.100 | 0.564 | 54 | 3.782E-01 | 0.000 | 1.380E+07 | 1.469E+07 | 1.231E+07 | 3.362E+07 | 2.003E+05 | 9.131E+04 | 1.302E+04 | 1.489E+06 | 7.124E+06 | 1.218E+06 | 2.573E+07 | 1.377E+05 | 8.852E+04 | 1.104E+04 |
| 479 | 242.101 | 0.569 | 54 | 4.916E+00 | 0.001 | 9.614E+06 | 4.750E+06 | 5.060E+06 | 3.605E+07 | 2.003E+05 | 9.337E+04 | 1.302E+04 | 8.951E+05 | 3.787E+06 | 1.589E+06 | 2.741E+07 | 1.377E+05 | 8.747E+04 | 1.104E+04 |
| 478 | 204.087 | 0.529 | 51 | 1.271E+01 | 0.003 | 3.619E+04 | 6.603E+04 | 6.239E+03 | 8.886E+07 | 7.285E+04 | 2.449E+04 | 3.685E+04 | 1.270E+04 | 2.224E+04 | 1.455E+03 | 1.084E+07 | 6.805E+03 | 1.756E+04 | 1.193E+04 |
| 200 | 311.256 | 7.678 | 39 | 2.941E+00 | 0.001 | 7.956E+05 | 5.712E+06 | 3.886E+06 | 8.176E+06 | 1.302E+07 | 1.121E+07 | 1.686E+07 | 1.009E+05 | 1.782E+06 | 3.299E+05 | 3.700E+06 | 2.928E+06 | 7.673E+05 | 2.983E+06 |
| 461 | 311.256 | 7.795 | 39 | 2.157E+00 | 0.001 | 2.272E+05 | 8.595E+05 | 1.162E+06 | 1.563E+06 | 1.148E+07 | 8.557E+06 | 3.336E+07 | 2.600E+04 | 2.888E+05 | 2.361E+05 | 4.758E+05 | 2.985E+06 | 1.857E+06 | 1.334E+07 |
| 536 | 339.290 | 8.561 | 39 | 1.102E+05 | 42.010 | 2.312E+05 | 9.624E+04 | 1.029E+06 | 7.860E+05 | 1.055E+07 | 4.169E+06 | 1.534E+07 | 9.868E+04 | 9.078E+03 | 2.094E+05 | 4.383E+05 | 9.035E+05 | 2.004E+06 | 4.849E+06 |
| 574 | 339.289 | 8.380 | 39 | 1.102E+05 | 42.010 | 4.818E+05 | 2.773E+05 | 7.918E+05 | 7.085E+05 | 2.632E+06 | 1.139E+06 | 2.332E+06 | 8.029E+03 | 9.016E+04 | 5.959E+04 | 2.903E+05 | 1.235E+05 | 2.735E+05 | 4.608E+05 |
| 666 | 188.071 | 1.307 | 27 | 4.057E+00 | 0.001 | 2.027E+08 | 5.777E+03 | 0.000E+00 | 5.199E+02 | 3.323E+03 | 0.000E+00 | 2.558E+03 | 2.112E+07 | 1.177E+03 | 0.000E+00 | 5.199E+02 | 7.320E+02 | 0.000E+00 | 2.558E+03 |
| 6 | 166.086 | 0.517 | 41 | 2.030E+01 | 0.003 | 1.415E+09 | 3.286E+09 | 2.841E+08 | 9.561E+08 | 1.768E+09 | 1.835E+09 | 2.766E+08 | 1.525E+08 | 7.083E+08 | 4.621E+07 | 2.216E+08 | 1.264E+08 | 1.447E+09 | 7.073E+07 |
| 441 | 223.090 | 6.761 | -1 | 3.290E+01 | 0.007 | 1.875E+06 | 4.945E+06 | 1.652E+06 | 2.901E+06 | 1.694E+06 | 2.292E+06 | 2.128E+06 | 5.186E+05 | 1.396E+06 | 2.672E+05 | 1.488E+06 | 1.125E+05 | 6.836E+05 | 6.258E+05 |
| 378 | 391.285 | 10.644 | -1 | 2.262E+00 | 0.001 | 5.357E+05 | 6.080E+05 | 4.799E+05 | 5.667E+05 | 6.006E+05 | 5.690E+05 | 5.934E+05 | 1.491E+04 | 7.693E+04 | 1.067E+05 | 3.261E+04 | 1.825E+04 | 1.210E+05 | 9.409E+04 |
| 12 | 245.096 | 3.603 | 62 | 2.366E+00 | 0.001 | 2.741E+07 | 2.175E+08 | 1.620E+08 | 2.371E+08 | 1.421E+08 | 1.961E+08 | 1.608E+08 | 1.813E+06 | 2.858E+07 | 8.840E+06 | 4.908E+07 | 3.110E+07 | 2.534E+07 | 2.374E+07 |
| 408 | 489.184 | 3.610 | 62 | 7.798E+00 | 0.004 | 0.000E+00 | 3.287E+06 | 1.696E+06 | 4.055E+06 | 1.380E+06 | 2.706E+06 | 1.916E+06 | 0.000E+00 | 9.199E+05 | 1.422E+05 | 1.640E+06 | 6.625E+05 | 7.380E+05 | 8.256E+05 |
| 467 | 250.118 | 7.417 | -1 | 5.918E+00 | 0.001 | 6.814E+05 | 6.177E+05 | 1.277E+06 | 6.091E+05 | 6.628E+05 | 5.176E+05 | 4.397E+05 | 1.293E+05 | 1.859E+05 | 2.844E+04 | 7.791E+04 | 8.078E+04 | 8.899E+04 | 1.057E+05 |
| 462 | 482.321 | 9.064 | 4 | 4.191E+03 | 2.013 | 7.453E+04 | 9.074E+04 | 6.668E+05 | 5.523E+05 | 1.945E+06 | 1.375E+06 | 2.494E+06 | 7.333E+03 | 3.013E+04 | 7.907E+04 | 2.836E+05 | 4.827E+05 | 6.841E+05 | 1.276E+06 |
| 832 | 577.516 | 12.699 | 39 | 4.016E+00 | 0.002 | 0.000E+00 | 2.582E+03 | 2.263E+04 | 7.216E+04 | 1.652E+05 | 5.256E+05 | 1.923E+06 | 0.000E+00 | 2.582E+03 | 2.263E+04 | 7.041E+03 | 5.271E+04 | 2.665E+05 | 8.736E+05 |
| 17 | 454.293 | 8.329 | 4 | 5.416E+04 | 26.015 | 2.290E+07 | 3.916E+07 | 5.484E+07 | 7.319E+07 | 1.363E+08 | 1.006E+08 | 1.521E+08 | 1.490E+06 | 1.271E+07 | 4.841E+06 | 3.058E+07 | 2.711E+07 | 9.389E+06 | 3.868E+07 |
| 89 | 454.293 | 8.190 | 4 | 2.222E+03 | 1.007 | 1.734E+06 | 5.405E+06 | 8.597E+06 | 1.055E+07 | 1.898E+07 | 1.491E+07 | 1.293E+07 | 2.039E+05 | 1.867E+06 | 6.256E+05 | 5.197E+06 | 3.924E+06 | 2.823E+06 | 2.526E+06 |
| 211 | 476.267 | 8.328 | 4 | 1.595E+01 | 0.008 | 1.484E+06 | 2.622E+06 | 3.725E+06 | 5.062E+06 | 8.685E+06 | 6.827E+06 | 9.610E+06 | 5.356E+04 | 8.837E+05 | 3.519E+05 | 2.080E+06 | 1.483E+06 | 5.178E+05 | 2.162E+06 |
| 743 | 907.578 | 8.331 | -1 | 9.415E-01 | 0.001 | 3.871E+03 | 9.447E+04 | 1.423E+05 | 3.329E+05 | 7.701E+05 | 5.513E+05 | 1.374E+06 | 3.871E+03 | 7.147E+04 | 4.850E+04 | 2.337E+05 | 3.160E+05 | 3.209E+04 | 6.224E+05 |
| 138 | 338.342 | 10.765 | 39 | 1.442E+05 | 57.010 | 4.678E+06 | 5.636E+06 | 4.718E+06 | 4.099E+06 | 4.304E+06 | 4.795E+06 | 5.349E+06 | 1.656E+05 | 7.485E+05 | 1.702E+05 | 1.610E+05 | 3.991E+05 | 5.102E+05 | 1.400E+06 |
| 456 | 338.342 | 9.820 | 39 | 2.345E+00 | 0.001 | 1.499E+06 | 1.428E+06 | 1.501E+06 | 1.470E+06 | 1.618E+06 | 1.739E+06 | 1.851E+06 | 3.872E+04 | 1.913E+04 | 3.645E+04 | 2.590E+04 | 1.469E+04 | 2.086E+05 | 1.026E+05 |
| 767 | 688.493 | 12.701 | 4 | 1.468E+03 | 1.009 | 0.000E+00 | 0.000E+00 | 0.000E+00 | 0.000E+00 | 1.391E+06 | 5.369E+06 | 9.426E+06 | 0.000E+00 | 0.000E+00 | 0.000E+00 | 0.000E+00 | 9.688E+05 | 3.405E+06 | 6.283E+06 |
| 30 | 480.309 | 8.382 | 4 | 2.103E+03 | 1.008 | 1.217E+07 | 7.529E+06 | 2.092E+07 | 1.948E+07 | 1.111E+08 | 2.770E+07 | 8.067E+07 | 9.869E+05 | 2.059E+06 | 1.707E+06 | 7.138E+06 | 4.396E+06 | 7.075E+06 | 2.501E+07 |
| 36 | 480.309 | 8.512 | 4 | 1.716E+00 | 0.001 | 3.880E+06 | 2.317E+06 | 1.740E+07 | 8.643E+06 | 4.681E+07 | 1.408E+07 | 3.120E+07 | 4.668E+05 | 6.112E+05 | 1.730E+06 | 2.536E+06 | 1.935E+06 | 3.272E+06 | 7.315E+06 |
| 342 | 377.146 | 3.769 | -1 | 2.266E+00 | 0.001 | 7.366E+05 | 6.381E+05 | 3.117E+06 | 4.419E+06 | 1.307E+06 | 9.266E+05 | 8.651E+05 | 2.128E+05 | 3.319E+05 | 2.704E+05 | 2.739E+06 | 2.713E+05 | 3.659E+05 | 9.453E+04 |
| 473 | 220.118 | 0.532 | 61 | 1.109E+00 | 0.000 | 2.155E+07 | 6.867E+06 | 3.836E+07 | 2.615E+08 | 4.199E+05 | 6.647E+05 | 2.006E+05 | 1.173E+07 | 2.836E+06 | 3.530E+06 | 1.785E+08 | 2.876E+05 | 6.590E+05 | 1.289E+05 |
| 372 | 628.196 | 10.499 | 40 | 1.360E+01 | 0.009 | 2.107E+06 | 1.599E+06 | 1.397E+06 | 1.083E+06 | 1.121E+06 | 1.177E+06 | 1.148E+06 | 2.096E+05 | 1.434E+05 | 1.812E+05 | 4.373E+04 | 2.067E+04 | 7.934E+04 | 1.588E+04 |
| 3 | 432.238 | 7.526 | 24 | 8.472E-01 | 0.000 | 2.282E+08 | 4.128E+08 | 3.904E+08 | 3.950E+08 | 4.468E+08 | 4.810E+08 | 5.690E+08 | 1.742E+07 | 4.662E+07 | 4.298E+07 | 1.695E+07 | 7.967E+07 | 4.136E+07 | 2.462E+08 |
| 161 | 432.238 | 7.249 | 24 | 7.766E-01 | 0.000 | 1.307E+06 | 3.194E+06 | 4.140E+06 | 5.161E+06 | 4.796E+06 | 4.913E+06 | 4.494E+06 | 5.798E+04 | 1.329E+06 | 4.636E+05 | 2.487E+06 | 1.516E+06 | 1.978E+06 | 1.854E+06 |
| 442 | 432.238 | 7.812 | 24 | 4.101E+04 | 17.027 | 5.174E+05 | 2.234E+06 | 1.168E+06 | 1.512E+06 | 1.874E+06 | 2.176E+06 | 1.995E+06 | 8.656E+04 | 4.263E+05 | 8.751E+04 | 6.190E+05 | 1.236E+05 | 4.267E+05 | 8.218E+05 |
| 2 | 415.212 | 7.526 | 24 | 1.837E+00 | 0.001 | 2.127E+08 | 4.157E+08 | 3.885E+08 | 4.058E+08 | 4.720E+08 | 5.114E+08 | 6.189E+08 | 1.728E+07 | 5.610E+07 | 4.865E+07 | 2.578E+07 | 9.226E+07 | 5.553E+07 | 2.965E+08 |
| 78 | 415.212 | 7.248 | 24 | 1.837E+00 | 0.001 | 2.428E+06 | 5.973E+06 | 7.704E+06 | 9.834E+06 | 9.006E+06 | 9.121E+06 | 8.755E+06 | 1.648E+05 | 2.423E+06 | 8.674E+05 | 4.762E+06 | 2.668E+06 | 3.635E+06 | 3.484E+06 |
| 373 | 415.212 | 7.817 | 24 | 1.984E+00 | 0.001 | 1.067E+06 | 5.933E+06 | 2.075E+06 | 3.099E+06 | 3.836E+06 | 4.756E+06 | 3.285E+06 | 1.008E+05 | 1.681E+06 | 1.714E+05 | 1.303E+06 | 2.643E+05 | 8.560E+05 | 1.191E+06 |
| 40 | 327.078 | 8.430 | -1 | 3.732E-01 | 0.000 | 7.004E+05 | 1.062E+06 | 1.921E+07 | 6.872E+06 | 1.911E+06 | 9.002E+05 | 8.452E+05 | 5.002E+04 | 1.215E+05 | 2.260E+06 | 3.813E+06 | 5.042E+05 | 1.443E+05 | 1.175E+05 |
| 435 | 313.143 | 8.965 | -1 | 4.873E-01 | 0.000 | 1.260E+06 | 2.870E+06 | 8.395E+05 | 9.517E+05 | 4.911E+06 | 1.455E+06 | 9.595E+05 | 2.296E+05 | 1.144E+06 | 3.582E+04 | 6.157E+04 | 3.524E+06 | 3.926E+05 | 1.769E+05 |
| 80 | 223.089 | 2.607 | 27 | 8.772E+04 | 17.992 | 1.184E+04 | 3.156E+08 | 5.788E+07 | 2.102E+08 | 9.644E+05 | 2.778E+07 | 2.064E+05 | 6.901E+02 | 6.782E+07 | 3.161E+06 | 6.894E+07 | 4.398E+05 | 2.537E+07 | 1.972E+05 |
| 697 | 223.089 | 2.398 | 27 | 4.309E+00 | 0.001 | 3.144E+04 | 4.816E+07 | 7.745E+06 | 4.220E+07 | 1.092E+09 | 2.011E+09 | 5.093E+06 | 9.586E+03 | 1.032E+07 | 1.932E+06 | 2.023E+07 | 8.571E+08 | 4.964E+08 | 2.098E+06 |
| 798 | 223.089 | 2.491 | 27 | 4.172E+00 | 0.001 | 2.934E+04 | 3.205E+07 | 5.218E+06 | 5.537E+07 | 1.969E+08 | 9.110E+08 | 4.138E+06 | 9.852E+03 | 1.121E+07 | 2.816E+06 | 2.067E+07 | 8.604E+07 | 5.299E+08 | 3.821E+06 |
| 665 | 205.097 | 1.310 | 27 | 8.065E+04 | 17.991 | 2.521E+08 | 6.214E+03 | 2.726E+03 | 4.876E+02 | 1.481E+03 | 0.000E+00 | 1.341E+03 | 2.593E+07 | 2.492E+03 | 1.372E+03 | 4.876E+02 | 1.481E+03 | 0.000E+00 | 1.341E+03 |
| 154 | 457.112 | 3.497 | -1 | 3.338E-01 | 0.000 | 8.205E+06 | 3.071E+07 | 8.126E+06 | 2.094E+07 | 1.406E+07 | 2.028E+07 | 1.148E+07 | 4.302E+05 | 7.904E+06 | 1.047E+06 | 4.024E+06 | 5.820E+05 | 7.586E+05 | 9.403E+05 |
| 29 | 220.118 | 0.526 | 61 | 8.041E+00 | 0.002 | 3.994E+07 | 4.367E+07 | 7.338E+07 | 1.416E+08 | 1.332E+06 | 3.253E+05 | 2.006E+05 | 5.487E+06 | 2.093E+07 | 4.533E+06 | 6.964E+07 | 1.198E+06 | 3.196E+05 | 1.289E+05 |
| 681 | 331.165 | 0.498 | 41 | 4.608E-01 | 0.000 | 3.156E+06 | 1.084E+07 | 2.898E+03 | 8.293E+03 | 4.533E+06 | 5.988E+06 | 1.208E+04 | 4.045E+05 | 3.834E+06 | 1.587E+03 | 8.293E+03 | 5.755E+05 | 5.790E+06 | 8.672E+03 |
| 192 | 343.296 | 6.715 | 23 | 1.111E+01 | 0.004 | 4.940E+06 | 8.454E+06 | 3.651E+06 | 3.440E+06 | 6.753E+06 | 3.939E+06 | 3.867E+06 | 1.120E+06 | 6.860E+05 | 3.773E+04 | 3.026E+04 | 1.180E+06 | 5.605E+05 | 6.690E+05 |
| 589 | 481.400 | 5.241 | 3 | 9.509E-01 | 0.000 | 4.406E+04 | 3.133E+05 | 5.387E+03 | 2.247E+06 | 4.265E+04 | 9.467E+04 | 6.043E+05 | 3.001E+04 | 6.986E+04 | 4.400E+02 | 2.179E+06 | 1.603E+04 | 1.692E+04 | 5.864E+05 |
| 603 | 810.134 | 3.065 | 20 | 1.582E+00 | 0.001 | 1.926E+05 | 1.777E+07 | 3.536E+04 | 1.931E+07 | 1.509E+07 | 1.963E+07 | 5.197E+07 | 1.085E+05 | 7.396E+06 | 2.226E+04 | 1.921E+07 | 9.860E+06 | 1.277E+07 | 1.404E+07 |
| 521 | 716.523 | 12.701 | 4 | 3.765E+04 | 28.031 | 0.000E+00 | 9.809E+04 | 7.184E+05 | 9.279E+05 | 7.533E+05 | 7.704E+06 | 1.090E+07 | 0.000E+00 | 5.509E+04 | 3.714E+05 | 7.717E+05 | 3.413E+05 | 3.575E+06 | 4.222E+06 |
| 572 | 716.525 | 10.404 | 4 | 3.765E+04 | 28.029 | 0.000E+00 | 9.098E+03 | 2.223E+06 | 1.716E+06 | 1.588E+05 | 1.482E+07 | 1.100E+07 | 0.000E+00 | 2.284E+03 | 1.601E+06 | 1.693E+06 | 1.066E+05 | 8.907E+06 | 7.791E+06 |
| 654 | 716.524 | 10.304 | 4 | 1.410E+03 | 1.009 | 0.000E+00 | 2.035E+04 | 3.130E+06 | 4.972E+05 | 2.011E+06 | 9.939E+07 | 4.815E+07 | 0.000E+00 | 6.980E+03 | 2.742E+06 | 4.670E+05 | 1.944E+06 | 8.546E+07 | 2.409E+07 |
| 852 | 716.524 | 10.151 | 4 | 3.765E+04 | 28.030 | 2.973E+03 | 3.061E+04 | 4.454E+05 | 9.272E+04 | 2.423E+05 | 1.286E+08 | 1.352E+08 | 2.973E+03 | 1.476E+04 | 2.965E+05 | 3.792E+04 | 5.088E+04 | 1.175E+08 | 1.273E+08 |
| 524 | 426.262 | 7.583 | 4 | 1.125E+05 | 54.046 | 8.384E+05 | 2.880E+06 | 8.364E+05 | 1.986E+06 | 8.465E+06 | 8.655E+06 | 8.775E+06 | 7.851E+04 | 9.550E+05 | 7.324E+04 | 7.824E+05 | 1.882E+06 | 2.919E+06 | 1.813E+06 |
| 552 | 426.262 | 7.443 | 4 | 6.170E+04 | 28.029 | 3.055E+06 | 4.437E+06 | 6.123E+05 | 2.416E+06 | 9.472E+06 | 1.526E+07 | 1.581E+07 | 1.270E+05 | 1.743E+06 | 9.931E+04 | 1.187E+06 | 2.186E+06 | 3.596E+06 | 2.473E+06 |
| 507 | 718.539 | 12.704 | 4 | 1.189E+00 | 0.001 | 0.000E+00 | 1.556E+06 | 2.204E+06 | 2.607E+06 | 3.917E+06 | 4.698E+07 | 5.483E+07 | 0.000E+00 | 1.556E+06 | 1.732E+06 | 1.061E+06 | 2.322E+06 | 2.185E+07 | 3.466E+07 |
| 523 | 718.539 | 10.302 | 4 | 9.344E-01 | 0.001 | 0.000E+00 | 4.079E+05 | 1.411E+06 | 1.549E+07 | 3.306E+07 | 5.628E+06 | 1.221E+07 | 0.000E+00 | 4.079E+05 | 1.044E+06 | 1.137E+07 | 1.793E+07 | 2.154E+06 | 4.437E+06 |
| 632 | 718.538 | 10.143 | 4 | 2.548E-01 | 0.000 | 0.000E+00 | 1.122E+05 | 8.844E+06 | 8.043E+06 | 2.056E+06 | 2.522E+07 | 4.675E+07 | 0.000E+00 | 1.122E+05 | 8.438E+06 | 7.163E+06 | 1.290E+06 | 1.818E+07 | 2.536E+07 |
| 823 | 771.514 | 12.696 | 4 | 3.262E+04 | 26.017 | 0.000E+00 | 0.000E+00 | 0.000E+00 | 1.644E+05 | 3.651E+05 | 1.405E+05 | 5.176E+06 | 0.000E+00 | 0.000E+00 | 0.000E+00 | 3.274E+04 | 1.321E+05 | 4.612E+04 | 2.464E+06 |
| 346 | 502.290 | 8.380 | 4 | 5.462E+04 | 26.015 | 7.953E+05 | 4.936E+05 | 1.479E+06 | 1.404E+06 | 4.733E+06 | 2.033E+06 | 3.994E+06 | 5.363E+04 | 1.124E+05 | 1.228E+05 | 5.706E+05 | 1.316E+05 | 5.407E+05 | 7.662E+05 |
| 388 | 502.291 | 8.511 | 4 | 5.463E+04 | 26.018 | 2.150E+05 | 9.234E+04 | 1.211E+06 | 5.629E+05 | 3.135E+06 | 8.916E+05 | 1.387E+06 | 4.605E+04 | 3.095E+04 | 1.495E+05 | 1.848E+05 | 1.380E+05 | 2.391E+05 | 1.737E+05 |
| 476 | 979.349 | 0.522 | 91 | N/A | N/A | 0.000E+00 | 0.000E+00 | 0.000E+00 | 1.694E+08 | 0.000E+00 | 0.000E+00 | 0.000E+00 | 0.000E+00 | 0.000E+00 | 0.000E+00 | 1.388E+07 | 0.000E+00 | 0.000E+00 | 0.000E+00 |
| 488 | 995.318 | 0.510 | 91 | N/A | N/A | 0.000E+00 | 0.000E+00 | 0.000E+00 | 2.068E+07 | 0.000E+00 | 0.000E+00 | 0.000E+00 | 0.000E+00 | 0.000E+00 | 0.000E+00 | 1.685E+06 | 0.000E+00 | 0.000E+00 | 0.000E+00 |
| 52 | 256.264 | 10.439 | 88 | N/A | N/A | 1.580E+07 | 1.358E+07 | 1.543E+07 | 1.242E+07 | 1.044E+07 | 1.138E+07 | 9.037E+06 | 2.979E+05 | 4.033E+06 | 1.298E+06 | 2.449E+06 | 1.400E+06 | 2.731E+06 | 1.285E+06 |
| 427 | 284.295 | 10.783 | 88 | N/A | N/A | 1.285E+06 | 1.112E+06 | 1.188E+06 | 1.224E+06 | 2.232E+06 | 2.298E+06 | 2.094E+06 | 7.020E+04 | 2.689E+05 | 2.339E+05 | 3.483E+05 | 2.620E+05 | 5.415E+05 | 6.624E+05 |
| 62 | 291.096 | 6.457 | 87 | N/A | N/A | 0.000E+00 | 0.000E+00 | 9.898E+06 | 4.719E+05 | 0.000E+00 | 0.000E+00 | 0.000E+00 | 0.000E+00 | 0.000E+00 | 1.412E+06 | 4.719E+05 | 0.000E+00 | 0.000E+00 | 0.000E+00 |
| 153 | 291.096 | 6.559 | 87 | N/A | N/A | 0.000E+00 | 0.000E+00 | 5.014E+06 | 2.051E+05 | 2.532E+03 | 0.000E+00 | 0.000E+00 | 0.000E+00 | 0.000E+00 | 7.494E+05 | 2.051E+05 | 2.532E+03 | 0.000E+00 | 0.000E+00 |
| 374 | 748.356 | 4.106 | 84 | N/A | N/A | 0.000E+00 | 0.000E+00 | 1.322E+06 | 2.184E+05 | 0.000E+00 | 0.000E+00 | 0.000E+00 | 0.000E+00 | 0.000E+00 | 2.590E+04 | 2.184E+05 | 0.000E+00 | 0.000E+00 | 0.000E+00 |
| 384 | 783.875 | 4.128 | 84 | N/A | N/A | 0.000E+00 | 0.000E+00 | 1.600E+06 | 3.041E+05 | 0.000E+00 | 0.000E+00 | 0.000E+00 | 0.000E+00 | 0.000E+00 | 5.230E+04 | 3.041E+05 | 0.000E+00 | 0.000E+00 | 0.000E+00 |
| 284 | 243.196 | 7.929 | 80 | 2.237E+05 | 70.078 | 2.242E+04 | 5.767E+04 | 2.719E+06 | 3.469E+06 | 3.956E+04 | 1.278E+04 | 2.033E+03 | 5.636E+03 | 2.340E+04 | 3.122E+05 | 2.221E+06 | 2.947E+04 | 3.086E+03 | 2.033E+03 |

| Feature identifiers | | | | | | Average XIC | | | | | | | Standard Err of Average XIC | | | | | | |
|---|---|---|---|---|---|---|---|---|---|---|---|---|---|---|---|---|---|---|---|
| ClusterIdx | m/z | RT [min] | Subnetwork | PPM | MassDiff | TUB00 | 4TUB34 | 4TUB81 | 4TUB93 | 5TUB23 | 5TUB48 | 5TUB83 | TUB00 | 4TUB34 | 4TUB81 | 4TUB93 | 5TUB23 | 5TUB48 | 5TUB83 |
| 415 | 225.185 | 7.934 | 80 | 1.993E+05 | 56.063 | 2.298E+04 | 3.921E+04 | 1.409E+06 | 1.825E+06 | 2.450E+04 | 1.253E+04 | 7.395E+03 | 7.142E+03 | 1.429E+04 | 1.664E+05 | 1.170E+06 | 1.556E+04 | 5.302E+03 | 1.446E+03 |
| 667 | 176.074 | 0.736 | 79 | 4.429E+05 | 54.044 | 9.158E+07 | 1.451E+08 | 6.134E+04 | 4.477E+05 | 5.988E+07 | 3.046E+07 | 1.864E+05 | 1.188E+07 | 4.828E+07 | 2.150E+04 | 1.691E+05 | 7.355E+06 | 2.839E+07 | 1.020E+05 |
| 668 | 218.121 | 4.865 | 79 | 2.432E+05 | 70.078 | 1.517E+07 | 3.044E+07 | 5.726E+04 | 1.993E+05 | 1.214E+07 | 4.775E+06 | 1.058E+05 | 1.052E+06 | 7.638E+06 | 2.949E+04 | 6.613E+04 | 7.026E+05 | 4.582E+06 | 6.987E+04 |
| 670 | 246.152 | 6.015 | 79 | 1.459E+05 | 42.047 | 1.999E+07 | 2.641E+07 | 1.426E+05 | 1.640E+05 | 1.498E+07 | 5.263E+06 | 2.320E+04 | 1.284E+05 | 5.558E+06 | 6.919E+04 | 1.130E+05 | 1.458E+06 | 5.168E+06 | 8.452E+03 |
| 674 | 232.137 | 5.471 | 79 | 1.945E+05 | 56.062 | 4.557E+06 | 8.336E+06 | 1.080E+04 | 1.485E+04 | 3.808E+06 | 1.303E+06 | 1.934E+03 | 1.223E+05 | 1.417E+06 | 1.080E+04 | 1.224E+04 | 4.635E+05 | 1.281E+06 | 1.934E+03 |
| 675 | 204.105 | 4.140 | 79 | 2.918E+05 | 84.094 | 8.099E+06 | 1.868E+07 | 0.000E+00 | 5.546E+04 | 7.407E+06 | 1.666E+06 | 2.015E+04 | 3.436E+05 | 2.969E+06 | 0.000E+00 | 3.562E+04 | 7.960E+05 | 1.666E+06 | 2.015E+04 |
| 772 | 551.192 | 3.315 | 78 | N/A | N/A | 2.191E+05 | 4.496E+06 | 0.000E+00 | 8.216E+04 | 2.808E+03 | 5.721E+05 | 1.404E+05 | 1.098E+05 | 2.249E+06 | 0.000E+00 | 7.026E+04 | 1.596E+03 | 5.667E+05 | 1.404E+05 |
| 774 | 615.714 | 3.354 | 78 | N/A | N/A | 5.927E+04 | 3.689E+06 | 0.000E+00 | 3.845E+03 | 0.000E+00 | 6.022E+05 | 2.114E+05 | 2.972E+04 | 1.844E+06 | 0.000E+00 | 3.845E+03 | 0.000E+00 | 5.871E+05 | 2.114E+05 |
| 776 | 486.670 | 3.240 | 78 | N/A | N/A | 7.745E+04 | 2.192E+06 | 0.000E+00 | 8.458E+04 | 0.000E+00 | 2.600E+05 | 4.543E+04 | 3.875E+04 | 1.096E+06 | 0.000E+00 | 8.458E+04 | 0.000E+00 | 2.600E+05 | 4.543E+04 |
| 777 | 680.235 | 3.406 | 78 | N/A | N/A | 3.079E+04 | 1.789E+06 | 0.000E+00 | 5.307E+03 | 0.000E+00 | 5.047E+05 | 2.135E+05 | 1.888E+04 | 9.364E+05 | 0.000E+00 | 5.307E+03 | 0.000E+00 | 4.961E+05 | 2.135E+05 |
| 780 | 357.628 | 3.065 | 78 | N/A | N/A | 1.032E+05 | 4.166E+06 | 1.610E+04 | 2.074E+05 | 1.495E+05 | 3.012E+05 | 5.679E+04 | 4.852E+04 | 2.106E+06 | 3.964E+03 | 5.719E+04 | 1.223E+05 | 2.681E+05 | 5.679E+04 |
| 46 | 226.158 | 9.291 | 77 | 2.694E+05 | 47.999 | 6.041E+05 | 1.746E+06 | 9.603E+06 | 1.538E+07 | 2.365E+06 | 4.147E+05 | 4.093E+05 | 1.024E+05 | 5.207E+05 | 1.835E+06 | 8.938E+06 | 6.162E+05 | 3.165E+04 | 2.148E+05 |
| 201 | 282.221 | 10.073 | 77 | N/A | N/A | 1.018E+06 | 1.396E+06 | 1.732E+06 | 2.503E+06 | 3.773E+06 | 3.953E+05 | 1.766E+05 | 3.429E+05 | 8.638E+05 | 1.140E+06 | 1.709E+06 | 2.256E+06 | 6.959E+04 | 7.700E+04 |
| 634 | 282.221 | 10.084 | 77 | N/A | N/A | 7.502E+05 | 9.664E+05 | 1.152E+06 | 1.732E+06 | 3.524E+06 | 3.953E+05 | 1.595E+05 | 1.738E+05 | 5.369E+05 | 7.580E+05 | 1.371E+06 | 1.802E+06 | 6.959E+04 | 6.066E+04 |
| 150 | 818.299 | 3.185 | 75 | N/A | N/A | 1.890E+06 | 2.122E+07 | 9.529E+06 | 1.581E+07 | 1.803E+07 | 4.416E+07 | 2.194E+07 | 7.105E+05 | 4.415E+06 | 1.003E+06 | 1.857E+06 | 7.802E+06 | 7.008E+06 | 4.387E+06 |
| 323 | 1226.943 | 3.181 | 75 | N/A | N/A | 4.911E+05 | 5.718E+06 | 2.652E+06 | 4.618E+06 | 3.838E+06 | 1.109E+07 | 4.645E+06 | 6.558E+04 | 1.339E+06 | 1.946E+05 | 6.308E+05 | 1.508E+06 | 6.077E+05 | 2.038E+06 |
| 843 | 1226.943 | 3.183 | 75 | N/A | N/A | 4.177E+05 | 4.397E+06 | 1.864E+06 | 3.384E+06 | 2.387E+06 | 9.866E+06 | 4.559E+06 | 1.478E+05 | 7.446E+05 | 4.497E+05 | 4.855E+05 | 3.535E+05 | 3.619E+06 | 9.189E+05 |
| 186 | 135.079 | 7.525 | 74 | 2.503E+05 | 45.096 | 3.177E+06 | 6.618E+06 | 6.102E+06 | 6.632E+06 | 8.043E+06 | 8.439E+06 | 1.088E+07 | 2.622E+05 | 8.343E+05 | 5.885E+05 | 2.893E+05 | 1.745E+06 | 8.902E+05 | 5.547E+06 |
| 319 | 135.079 | 4.463 | 74 | 2.503E+05 | 45.096 | 9.112E+05 | 2.161E+06 | 1.959E+06 | 2.311E+06 | 3.082E+06 | 3.696E+06 | 5.214E+06 | 5.235E+04 | 4.743E+05 | 2.715E+05 | 6.431E+05 | 5.304E+05 | 5.402E+05 | 2.425E+06 |
| 180 | 529.790 | 4.168 | 73 | N/A | N/A | 0.000E+00 | 0.000E+00 | 5.694E+06 | 1.983E+06 | 3.825E+03 | 0.000E+00 | 0.000E+00 | 0.000E+00 | 0.000E+00 | 5.704E+05 | 1.983E+06 | 3.825E+03 | 0.000E+00 | 0.000E+00 |
| 264 | 465.270 | 4.196 | 73 | N/A | N/A | 0.000E+00 | 0.000E+00 | 3.650E+06 | 1.281E+06 | 5.743E+03 | 0.000E+00 | 0.000E+00 | 0.000E+00 | 0.000E+00 | 4.530E+05 | 1.281E+06 | 2.931E+03 | 0.000E+00 | 0.000E+00 |
| 48 | 431.565 | 4.119 | 72 | N/A | N/A | 2.161E+04 | 1.717E+04 | 1.369E+07 | 2.106E+05 | 0.000E+00 | 0.000E+00 | 0.000E+00 | 1.675E+04 | 1.717E+04 | 1.737E+06 | 4.307E+04 | 0.000E+00 | 0.000E+00 | 0.000E+00 |
| 170 | 417.549 | 3.504 | 72 | N/A | N/A | 4.190E+07 | 5.204E+07 | 7.068E+06 | 6.493E+07 | 6.974E+06 | 1.020E+07 | 1.516E+07 | 1.244E+07 | 2.368E+07 | 1.631E+06 | 5.365E+07 | 4.819E+06 | 4.701E+06 | 9.846E+05 |
| 314 | 396.544 | 3.209 | 72 | N/A | N/A | 6.715E+07 | 4.419E+06 | 5.259E+06 | 1.705E+07 | 1.244E+06 | 7.634E+05 | 9.883E+05 | 8.326E+06 | 2.496E+06 | 5.761E+05 | 9.088E+06 | 9.228E+05 | 3.715E+05 | 9.844E+05 |
| 117 | 832.302 | 3.461 | 70 | N/A | N/A | 3.211E+06 | 1.270E+07 | 8.253E+06 | 1.296E+07 | 9.164E+06 | 2.487E+07 | 3.156E+07 | 8.228E+05 | 2.889E+06 | 5.411E+05 | 5.232E+06 | 1.535E+06 | 7.834E+06 | 5.165E+06 |
| 371 | 1247.949 | 3.461 | 70 | N/A | N/A | 1.224E+06 | 3.849E+06 | 2.474E+06 | 4.329E+06 | 2.845E+06 | 8.094E+06 | 8.847E+06 | 1.663E+05 | 8.419E+05 | 2.722E+05 | 1.786E+06 | 4.848E+05 | 2.900E+06 | 9.722E+05 |
| 512 | 1247.454 | 3.462 | 70 | N/A | N/A | 5.948E+05 | 3.544E+06 | 1.732E+06 | 3.436E+06 | 2.228E+06 | 5.189E+06 | 8.185E+06 | 8.601E+04 | 8.658E+05 | 2.049E+05 | 1.483E+06 | 6.003E+05 | 7.769E+05 | 1.069E+06 |
| 7 | 271.106 | 8.022 | 69 | N/A | N/A | 8.953E+03 | 3.153E+07 | 1.232E+08 | 5.492E+07 | 2.901E+05 | 7.422E+04 | 1.912E+04 | 4.477E+03 | 3.634E+06 | 1.146E+07 | 2.283E+07 | 1.479E+05 | 3.442E+04 | 2.077E+03 |
| 45 | 390.122 | 6.709 | 69 | N/A | N/A | 0.000E+00 | 1.529E+06 | 1.276E+07 | 5.182E+06 | 0.000E+00 | 0.000E+00 | 0.000E+00 | 0.000E+00 | 1.129E+05 | 8.927E+05 | 2.394E+06 | 0.000E+00 | 0.000E+00 | 0.000E+00 |
| 112 | 388.106 | 7.488 | 69 | N/A | N/A | 0.000E+00 | 1.400E+06 | 5.143E+06 | 3.447E+06 | 0.000E+00 | 0.000E+00 | 0.000E+00 | 0.000E+00 | 3.235E+05 | 3.785E+05 | 1.547E+06 | 0.000E+00 | 0.000E+00 | 0.000E+00 |
| 209 | 414.122 | 6.104 | 69 | N/A | N/A | 0.000E+00 | 4.453E+06 | 2.502E+06 | 7.308E+06 | 1.612E+03 | 0.000E+00 | 0.000E+00 | 0.000E+00 | 9.417E+05 | 3.329E+05 | 4.363E+06 | 1.612E+03 | 0.000E+00 | 0.000E+00 |
| 663 | 414.122 | 6.419 | 69 | N/A | N/A | 7.841E+04 | 4.417E+04 | 1.543E+05 | 7.747E+05 | 1.728E+06 | 3.735E+06 | 1.325E+06 | 7.841E+04 | 2.468E+04 | 1.381E+05 | 7.250E+05 | 1.206E+06 | 8.326E+05 | 6.233E+05 |
| 468 | 432.238 | 6.913 | 68 | N/A | N/A | 9.316E+05 | 1.395E+07 | 9.898E+05 | 3.732E+05 | 4.407E+05 | 1.037E+07 | 1.711E+05 | 1.297E+05 | 7.612E+06 | 2.577E+05 | 3.205E+04 | 9.223E+04 | 9.831E+06 | 2.072E+04 |
| 792 | 404.208 | 6.914 | 68 | N/A | N/A | 6.144E+05 | 1.033E+07 | 5.795E+05 | 1.594E+05 | 2.574E+05 | 7.637E+06 | 7.469E+04 | 8.607E+04 | 5.744E+06 | 1.288E+05 | 1.575E+04 | 6.875E+04 | 7.306E+06 | 1.531E+04 |
| 793 | 387.181 | 6.913 | 68 | N/A | N/A | 6.030E+05 | 9.983E+06 | 5.851E+05 | 1.889E+05 | 2.556E+05 | 7.493E+06 | 6.517E+04 | 7.564E+04 | 5.462E+06 | 1.190E+05 | 4.933E+04 | 6.784E+04 | 7.167E+06 | 6.269E+03 |
| 671 | 331.207 | 5.234 | 67 | N/A | N/A | 1.290E+07 | 9.695E+06 | 1.323E+05 | 5.809E+04 | 4.049E+06 | 7.946E+06 | 2.180E+04 | 1.588E+06 | 3.406E+06 | 7.827E+04 | 3.566E+04 | 8.758E+05 | 7.799E+06 | 1.150E+04 |
| 795 | 317.195 | 4.828 | 67 | N/A | N/A | 8.310E+05 | 9.431E+05 | 4.933E+04 | 1.226E+05 | 5.115E+05 | 1.102E+06 | 9.712E+04 | 1.206E+05 | 3.098E+05 | 1.525E+04 | 3.140E+04 | 1.260E+05 | 8.021E+05 | 3.366E+04 |
| 84 | 280.188 | 5.797 | 65 | 4.291E+05 | 84.135 | 1.203E+04 | 5.142E+03 | 6.768E+06 | 5.432E+06 | 1.143E+03 | 0.000E+00 | 1.822E+03 | 1.512E+03 | 1.991E+03 | 4.623E+05 | 5.429E+06 | 1.143E+03 | 0.000E+00 | 1.822E+03 |
| 158 | 308.222 | 6.723 | 65 | N/A | N/A | 9.850E+03 | 0.000E+00 | 4.477E+06 | 2.603E+06 | 0.000E+00 | 0.000E+00 | 0.000E+00 | 7.547E+03 | 0.000E+00 | 4.152E+05 | 2.601E+06 | 0.000E+00 | 0.000E+00 | 0.000E+00 |
| 263 | 693.878 | 3.983 | 64 | N/A | N/A | 0.000E+00 | 0.000E+00 | 2.570E+06 | 2.728E+05 | 0.000E+00 | 0.000E+00 | 0.000E+00 | 0.000E+00 | 0.000E+00 | 8.661E+04 | 2.728E+05 | 0.000E+00 | 0.000E+00 | 0.000E+00 |
| 392 | 665.367 | 3.986 | 64 | N/A | N/A | 0.000E+00 | 0.000E+00 | 1.452E+06 | 3.505E+05 | 0.000E+00 | 0.000E+00 | 0.000E+00 | 0.000E+00 | 0.000E+00 | 1.222E+05 | 3.505E+05 | 0.000E+00 | 0.000E+00 | 0.000E+00 |
| 703 | 204.534 | 6.795 | 63 | N/A | N/A | 0.000E+00 | 1.696E+03 | 0.000E+00 | 0.000E+00 | 1.265E+07 | 0.000E+00 | 1.504E+05 | 0.000E+00 | 1.696E+03 | 0.000E+00 | 0.000E+00 | 2.742E+06 | 0.000E+00 | 8.635E+04 |
| 704 | 193.026 | 6.808 | 63 | N/A | N/A | 0.000E+00 | 0.000E+00 | 0.000E+00 | 0.000E+00 | 1.379E+07 | 0.000E+00 | 2.042E+05 | 0.000E+00 | 0.000E+00 | 0.000E+00 | 0.000E+00 | 3.059E+06 | 0.000E+00 | 1.010E+05 |
| 88 | 227.084 | 3.606 | 62 | 7.349E+04 | 18.011 | 2.728E+06 | 2.387E+07 | 1.773E+07 | 2.612E+07 | 1.558E+07 | 2.176E+07 | 1.791E+07 | 1.250E+05 | 3.208E+06 | 9.281E+05 | 5.537E+06 | 3.418E+06 | 2.857E+06 | 2.626E+06 |
| 215 | 290.153 | 3.607 | 62 | 1.077E+05 | 35.018 | 4.388E+05 | 8.919E+06 | 6.702E+06 | 1.095E+07 | 5.351E+06 | 8.491E+06 | 6.890E+06 | 3.721E+04 | 1.632E+06 | 4.536E+05 | 2.894E+06 | 1.213E+06 | 1.076E+06 | 1.389E+06 |
| 4 | 182.985 | 12.752 | 60 | N/A | N/A | 5.116E+08 | 5.361E+08 | 8.037E+08 | 7.867E+08 | 7.552E+08 | 7.261E+08 | 7.010E+08 | 1.593E+08 | 5.970E+07 | 1.042E+07 | 6.643E+06 | 7.803E+06 | 6.831E+06 | 4.161E+06 |
| 5 | 154.990 | 12.767 | 60 | N/A | N/A | 8.835E+08 | 7.944E+08 | 7.553E+08 | 7.516E+08 | 7.253E+08 | 7.001E+08 | 6.745E+08 | 6.694E+06 | 2.442E+07 | 1.753E+07 | 8.684E+06 | 1.102E+07 | 2.093E+07 | 1.256E+07 |
| 19 | 138.996 | 12.748 | 60 | N/A | N/A | 1.356E+08 | 1.191E+08 | 1.114E+08 | 1.131E+08 | 1.090E+08 | 1.018E+08 | 9.963E+07 | 1.407E+06 | 4.923E+06 | 2.497E+06 | 2.510E+06 | 2.248E+06 | 9.701E+05 | 2.553E+06 |
| 23 | 205.017 | 12.741 | 60 | N/A | N/A | 5.733E+07 | 5.114E+07 | 4.994E+07 | 4.897E+07 | 4.652E+07 | 4.359E+07 | 4.163E+07 | 2.511E+05 | 2.976E+06 | 1.952E+06 | 1.903E+06 | 1.356E+06 | 1.758E+06 | 8.678E+05 |
| 49 | 196.017 | 12.766 | 60 | N/A | N/A | 4.261E+07 | 3.618E+07 | 3.219E+07 | 3.217E+07 | 3.007E+07 | 2.789E+07 | 2.569E+07 | 1.431E+06 | 2.445E+06 | 2.032E+06 | 6.412E+05 | 1.409E+06 | 1.196E+06 | 8.923E+05 |
| 63 | 224.012 | 12.748 | 60 | N/A | N/A | 3.457E+07 | 2.918E+07 | 2.577E+07 | 2.536E+07 | 2.315E+07 | 2.089E+07 | 1.887E+07 | 9.195E+05 | 1.718E+06 | 1.680E+06 | 1.575E+05 | 1.078E+06 | 1.221E+06 | 3.404E+05 |
| 75 | 195.994 | 12.775 | 60 | N/A | N/A | 3.380E+07 | 3.215E+07 | 3.114E+07 | 3.195E+07 | 3.088E+07 | 2.961E+07 | 2.992E+07 | 6.013E+05 | 8.298E+04 | 4.220E+05 | 3.796E+05 | 1.706E+05 | 6.312E+05 | 9.722E+05 |
| 20 | 131.534 | 12.743 | 59 | N/A | N/A | 8.479E+07 | 7.549E+07 | 7.205E+07 | 6.839E+07 | 6.489E+07 | 5.969E+07 | 5.648E+07 | 4.544E+04 | 2.501E+06 | 1.838E+06 | 1.208E+06 | 1.579E+06 | 2.468E+06 | 4.864E+05 |
| 409 | 131.534 | 13.009 | 59 | N/A | N/A | 8.964E+05 | 1.438E+06 | 6.609E+05 | 6.915E+05 | 7.453E+05 | 3.919E+05 | 6.800E+05 | 2.423E+05 | 3.143E+05 | 2.482E+05 | 3.813E+05 | 3.158E+05 | 9.790E+04 | 1.868E+05 |
| 465 | 131.534 | 13.069 | 59 | N/A | N/A | 6.260E+05 | 1.009E+06 | 3.605E+05 | 3.739E+05 | 4.458E+05 | 5.653E+05 | 3.712E+05 | 1.760E+05 | 3.164E+05 | 8.460E+04 | 1.364E+05 | 1.812E+05 | 1.528E+05 | 1.314E+05 |
| 696 | 131.534 | 4.507 | 59 | N/A | N/A | 2.432E+05 | 1.507E+05 | 2.283E+05 | 1.445E+05 | 8.517E+04 | 1.290E+05 | 1.290E+05 | 3.927E+04 | 2.438E+04 | 7.184E+04 | 4.888E+04 | 2.179E+04 | 3.136E+04 | 1.672E+04 |
| 266 | 444.309 | 8.752 | 58 | 1.129E+05 | 45.058 | 2.348E+06 | 4.084E+06 | 2.633E+06 | 6.605E+06 | 3.178E+06 | 3.543E+06 | 3.105E+06 | 1.562E+05 | 7.021E+05 | 7.297E+04 | 3.831E+06 | 2.414E+05 | 4.683E+05 | 2.563E+05 |
| 490 | 350.102 | 0.493 | 57 | 7.895E+04 | 30.011 | 5.129E+06 | 8.618E+06 | 2.726E+05 | 5.066E+06 | 1.345E+07 | 1.493E+07 | 1.439E+07 | 6.958E+05 | 4.896E+06 | 1.910E+05 | 3.236E+06 | 5.235E+06 | 1.166E+07 | 1.348E+07 |
| 495 | 364.118 | 0.634 | 57 | 4.208E+04 | 15.995 | 2.703E+05 | 4.944E+06 | 0.000E+00 | 2.673E+07 | 2.210E+07 | 7.315E+04 | 6.361E+03 | 7.399E+04 | 2.592E+06 | 0.000E+00 | 2.628E+07 | 2.192E+07 | 4.086E+04 | 6.361E+03 |
| 134 | 519.318 | 10.374 | 56 | N/A | N/A | 1.525E+06 | 5.359E+05 | 3.446E+06 | 2.565E+06 | 1.342E+06 | 1.593E+06 | 1.209E+06 | 1.369E+06 | 4.987E+05 | 1.727E+06 | 8.852E+05 | 6.393E+05 | 6.992E+05 | 4.902E+05 |
| 658 | 519.309 | 10.382 | 56 | N/A | N/A | 3.425E+06 | 1.827E+06 | 4.540E+06 | 2.788E+06 | 2.126E+06 | 2.499E+06 | 1.906E+06 | 4.729E+05 | 1.001E+05 | 6.493E+05 | 7.744E+05 | 3.815E+05 | 3.364E+04 | 9.998E+04 |
| 498 | 532.098 | 0.493 | 55 | N/A | N/A | 0.000E+00 | 0.000E+00 | 0.000E+00 | 5.368E+06 | 0.000E+00 | 0.000E+00 | 0.000E+00 | 0.000E+00 | 0.000E+00 | 0.000E+00 | 2.530E+06 | 0.000E+00 | 0.000E+00 | 0.000E+00 |
| 501 | 532.099 | 0.497 | 55 | N/A | N/A | 0.000E+00 | 0.000E+00 | 0.000E+00 | 4.490E+06 | 0.000E+00 | 0.000E+00 | 0.000E+00 | 0.000E+00 | 0.000E+00 | 0.000E+00 | 2.850E+06 | 0.000E+00 | 0.000E+00 | 0.000E+00 |
| 132 | 711.319 | 4.210 | 53 | N/A | N/A | 0.000E+00 | 3.155E+04 | 5.754E+06 | 9.097E+05 | 0.000E+00 | 0.000E+00 | 0.000E+00 | 0.000E+00 | 3.155E+04 | 6.612E+05 | 9.097E+05 | 0.000E+00 | 0.000E+00 | 0.000E+00 |
| 282 | 624.289 | 4.196 | 53 | N/A | N/A | 0.000E+00 | 0.000E+00 | 1.819E+06 | 2.519E+05 | 0.000E+00 | 0.000E+00 | 0.000E+00 | 0.000E+00 | 0.000E+00 | 3.871E+05 | 2.519E+05 | 0.000E+00 | 0.000E+00 | 0.000E+00 |
| 133 | 226.180 | 5.976 | 52 | 4.330E+03 | 0.984 | 6.080E+06 | 6.010E+06 | 5.153E+06 | 5.763E+06 | 5.679E+06 | 6.245E+06 | 5.714E+06 | 5.748E+05 | 1.775E+05 | 2.464E+05 | 3.087E+05 | 1.729E+05 | 3.427E+05 | 4.364E+05 |
| 542 | 384.349 | 10.283 | 52 | N/A | N/A | 1.433E+06 | 1.477E+06 | 1.688E+06 | 1.578E+06 | 1.497E+06 | 2.552E+06 | 2.570E+06 | 3.258E+04 | 3.264E+04 | 1.497E+05 | 6.888E+04 | 4.446E+04 | 2.811E+04 | 2.299E+05 |
| 131 | 614.463 | 9.741 | 51 | N/A | N/A | 1.053E+05 | 1.619E+06 | 1.095E+07 | 7.460E+06 | 1.394E+06 | 4.696E+05 | 8.730E+05 | 3.247E+04 | 7.066E+05 | 2.958E+05 | 1.813E+06 | 6.020E+05 | 5.543E+04 | 2.534E+05 |
| 243 | 388.269 | 6.414 | 51 | N/A | N/A | 1.031E+04 | 5.405E+05 | 4.794E+06 | 2.414E+06 | 2.101E+05 | 9.054E+04 | 6.744E+04 | 4.607E+02 | 7.374E+04 | 8.484E+05 | 6.189E+05 | 7.798E+04 | 3.783E+04 | 2.652E+04 |
| 472 | 479.186 | 0.514 | 51 | 6.445E+04 | 33.011 | 5.898E+05 | 1.164E+05 | 0.000E+00 | 3.371E+09 | 2.987E+06 | 2.775E+05 | 7.668E+04 | 5.898E+05 | 9.490E+04 | 0.000E+00 | 3.906E+08 | 9.211E+05 | 2.000E+05 | 4.636E+04 |

| Feature identifiers | | | | | | Average XIC | | | | | | | Standard Err of Average XIC | | | | | | |
|---|---|---|---|---|---|---|---|---|---|---|---|---|---|---|---|---|---|---|---|
| ClusterIdx | m/z | RT [min] | Subnetwork | PPM | MassDiff | TUB00 | 4TUB34 | 4TUB81 | 4TUB93 | 5TUB23 | 5TUB48 | 5TUB83 | TUB00 | 4TUB34 | 4TUB81 | 4TUB93 | 5TUB23 | 5TUB48 | 5TUB83 |
| 475 | 957.367 | 0.530 | 51 | 5.051E+04 | 46.032 | 0.000E+00 | 0.000E+00 | 0.000E+00 | 1.833E+08 | 0.000E+00 | 0.000E+00 | 0.000E+00 | 0.000E+00 | 0.000E+00 | 0.000E+00 | 3.389E+07 | 0.000E+00 | 0.000E+00 | 0.000E+00 |
| 477 | 524.246 | 0.489 | 51 | 2.352E+04 | 12.049 | 0.000E+00 | 0.000E+00 | 0.000E+00 | 7.175E+07 | 0.000E+00 | 0.000E+00 | 1.810E+03 | 0.000E+00 | 0.000E+00 | 0.000E+00 | 6.796E+06 | 0.000E+00 | 0.000E+00 | 1.810E+03 |
| 480 | 268.076 | 0.497 | 51 | 4.576E+05 | 40.936 | 0.000E+00 | 0.000E+00 | 0.000E+00 | 5.224E+07 | 0.000E+00 | 0.000E+00 | 0.000E+00 | 0.000E+00 | 0.000E+00 | 0.000E+00 | 3.788E+06 | 0.000E+00 | 0.000E+00 | 0.000E+00 |
| 482 | 498.161 | 0.503 | 51 | 4.534E+05 | 83.979 | 8.311E+03 | 0.000E+00 | 0.000E+00 | 5.093E+07 | 0.000E+00 | 0.000E+00 | 0.000E+00 | 7.395E+03 | 0.000E+00 | 0.000E+00 | 6.075E+06 | 0.000E+00 | 0.000E+00 | 0.000E+00 |
| 483 | 644.267 | 0.510 | 51 | 1.190E+05 | 87.005 | 0.000E+00 | 0.000E+00 | 0.000E+00 | 4.430E+07 | 5.480E+03 | 0.000E+00 | 0.000E+00 | 0.000E+00 | 0.000E+00 | 0.000E+00 | 1.025E+07 | 5.122E+03 | 0.000E+00 | 0.000E+00 |
| 484 | 279.584 | 0.532 | 51 | N/A | N/A | 0.000E+00 | 0.000E+00 | 0.000E+00 | 5.076E+07 | 0.000E+00 | 0.000E+00 | 0.000E+00 | 0.000E+00 | 0.000E+00 | 0.000E+00 | 3.928E+06 | 0.000E+00 | 0.000E+00 | 0.000E+00 |
| 487 | 276.062 | 0.490 | 51 | 3.527E+05 | 71.975 | 0.000E+00 | 0.000E+00 | 0.000E+00 | 1.432E+07 | 0.000E+00 | 0.000E+00 | 0.000E+00 | 0.000E+00 | 0.000E+00 | 0.000E+00 | 1.647E+06 | 0.000E+00 | 0.000E+00 | 0.000E+00 |
| 492 | 737.248 | 0.522 | 51 | N/A | N/A | 0.000E+00 | 0.000E+00 | 0.000E+00 | 1.525E+07 | 0.000E+00 | 0.000E+00 | 0.000E+00 | 0.000E+00 | 0.000E+00 | 0.000E+00 | 2.473E+06 | 0.000E+00 | 0.000E+00 | 0.000E+00 |
| 493 | 506.147 | 0.488 | 51 | 4.446E+05 | 99.950 | 0.000E+00 | 0.000E+00 | 0.000E+00 | 1.004E+07 | 0.000E+00 | 0.000E+00 | 0.000E+00 | 0.000E+00 | 0.000E+00 | 0.000E+00 | 9.238E+05 | 0.000E+00 | 0.000E+00 | 0.000E+00 |
| 500 | 287.570 | 0.486 | 51 | 5.326E+05 | 41.104 | 0.000E+00 | 0.000E+00 | 0.000E+00 | 6.915E+06 | 0.000E+00 | 0.000E+00 | 0.000E+00 | 0.000E+00 | 0.000E+00 | 0.000E+00 | 1.872E+06 | 0.000E+00 | 0.000E+00 | 0.000E+00 |
| 687 | 441.187 | 0.508 | 51 | 2.050E+05 | 75.047 | 7.266E+05 | 6.885E+06 | 1.373E+03 | 0.000E+00 | 1.112E+07 | 3.975E+05 | 5.707E+05 | 2.219E+05 | 3.631E+06 | 1.373E+03 | 0.000E+00 | 3.323E+06 | 3.233E+05 | 4.451E+05 |
| 732 | 632.474 | 10.196 | 51 | N/A | N/A | 2.491E+03 | 1.337E+05 | 4.601E+04 | 7.001E+04 | 2.100E+06 | 2.336E+06 | 1.860E+06 | 2.491E+03 | 6.124E+04 | 1.589E+04 | 3.874E+04 | 8.076E+05 | 7.682E+05 | 6.616E+05 |
| 802 | 860.514 | 9.355 | 51 | N/A | N/A | 0.000E+00 | 6.117E+04 | 1.118E+04 | 8.915E+04 | 1.327E+06 | 6.512E+05 | 2.130E+06 | 0.000E+00 | 1.689E+04 | 1.118E+04 | 6.314E+04 | 3.776E+05 | 1.566E+05 | 8.240E+05 |
| 168 | 471.307 | 5.923 | 50 | N/A | N/A | 3.287E+06 | 4.663E+06 | 3.614E+06 | 4.087E+06 | 5.424E+06 | 4.657E+06 | 3.367E+06 | 5.827E+05 | 1.085E+06 | 2.291E+05 | 2.361E+05 | 1.349E+06 | 1.802E+05 | 2.301E+05 |
| 616 | 473.286 | 4.748 | 50 | N/A | N/A | 1.160E+06 | 3.338E+06 | 5.739E+05 | 1.790E+06 | 8.821E+05 | 2.243E+06 | 1.049E+06 | 5.595E+05 | 2.837E+05 | 1.951E+05 | 8.773E+05 | 4.020E+05 | 4.659E+05 | 5.823E+05 |
| 144 | 393.207 | 10.310 | 49 | 1.653E+05 | 77.883 | 5.332E+06 | 5.275E+06 | 5.701E+06 | 5.653E+06 | 5.768E+06 | 5.996E+06 | 5.973E+06 | 1.221E+05 | 1.272E+05 | 3.454E+05 | 3.756E+04 | 8.323E+04 | 7.088E+04 | 2.024E+05 |
| 176 | 409.183 | 10.451 | 49 | 1.314E+05 | 61.922 | 3.870E+06 | 3.807E+06 | 4.206E+06 | 3.966E+06 | 3.811E+06 | 4.301E+06 | 4.350E+06 | 1.302E+05 | 1.538E+05 | 8.476E+04 | 2.202E+04 | 1.121E+05 | 4.583E+04 | 1.674E+05 |
| 43 | 418.117 | 7.633 | 48 | N/A | N/A | 0.000E+00 | 1.930E+06 | 1.357E+07 | 1.158E+07 | 0.000E+00 | 0.000E+00 | 0.000E+00 | 0.000E+00 | 1.567E+05 | 7.129E+05 | 5.882E+06 | 0.000E+00 | 0.000E+00 | 0.000E+00 |
| 146 | 551.149 | 6.292 | 48 | N/A | N/A | 0.000E+00 | 3.162E+06 | 4.749E+06 | 4.326E+06 | 0.000E+00 | 0.000E+00 | 0.000E+00 | 0.000E+00 | 1.478E+06 | 1.556E+06 | 2.985E+06 | 0.000E+00 | 0.000E+00 | 0.000E+00 |
| 363 | 418.117 | 7.570 | 48 | N/A | N/A | 0.000E+00 | 1.400E+06 | 2.879E+06 | 1.791E+06 | 0.000E+00 | 0.000E+00 | 0.000E+00 | 0.000E+00 | 6.288E+05 | 1.406E+06 | 7.219E+05 | 0.000E+00 | 0.000E+00 | 0.000E+00 |
| 119 | 343.130 | 3.690 | 47 | N/A | N/A | 5.601E+05 | 1.333E+07 | 9.907E+06 | 5.993E+07 | 1.421E+05 | 2.155E+05 | 2.736E+04 | 3.449E+05 | 2.579E+06 | 3.533E+05 | 2.828E+07 | 1.128E+05 | 1.270E+05 | 8.638E+03 |
| 182 | 537.334 | 9.562 | 47 | N/A | N/A | 6.605E+03 | 4.457E+04 | 6.444E+06 | 1.876E+06 | 1.630E+04 | 1.133E+04 | 0.000E+00 | 1.483E+03 | 1.959E+04 | 2.316E+06 | 1.759E+06 | 8.168E+03 | 7.448E+03 | 0.000E+00 |
| 231 | 509.303 | 8.961 | 47 | N/A | N/A | 0.000E+00 | 0.000E+00 | 3.088E+06 | 9.030E+05 | 0.000E+00 | 0.000E+00 | 0.000E+00 | 0.000E+00 | 0.000E+00 | 4.796E+05 | 8.925E+05 | 0.000E+00 | 0.000E+00 | 0.000E+00 |
| 239 | 245.616 | 7.678 | 46 | N/A | N/A | 4.563E+05 | 3.631E+06 | 2.284E+06 | 4.822E+06 | 7.721E+06 | 6.844E+06 | 1.023E+07 | 1.115E+05 | 1.157E+06 | 9.235E+04 | 2.163E+06 | 1.541E+06 | 2.787E+05 | 1.760E+06 |
| 393 | 266.129 | 7.678 | 46 | N/A | N/A | 1.878E+05 | 2.353E+06 | 1.552E+06 | 3.292E+06 | 4.669E+06 | 4.224E+06 | 5.731E+06 | 4.697E+03 | 8.126E+05 | 1.201E+05 | 1.448E+06 | 9.707E+05 | 2.177E+05 | 8.712E+05 |
| 429 | 266.129 | 7.807 | 46 | N/A | N/A | 3.680E+04 | 4.175E+05 | 1.124E+06 | 1.110E+06 | 2.393E+06 | 1.011E+06 | 1.234E+06 | 7.298E+03 | 1.164E+05 | 2.479E+05 | 3.339E+05 | 4.407E+05 | 2.622E+05 | 2.411E+05 |
| 506 | 734.471 | 12.708 | 45 | N/A | N/A | 0.000E+00 | 3.956E+05 | 3.555E+06 | 1.316E+06 | 2.223E+06 | 3.487E+06 | 1.150E+07 | 0.000E+00 | 2.048E+05 | 1.706E+06 | 8.762E+05 | 9.726E+05 | 1.384E+06 | 5.139E+06 |
| 544 | 734.471 | 10.338 | 45 | N/A | N/A | 0.000E+00 | 4.638E+04 | 8.616E+06 | 3.125E+06 | 7.151E+06 | 3.468E+06 | 6.019E+06 | 0.000E+00 | 2.778E+04 | 7.690E+06 | 2.973E+06 | 6.217E+06 | 1.743E+06 | 2.852E+06 |
| 550 | 765.459 | 12.703 | 45 | 6.134E+04 | 50.022 | 0.000E+00 | 0.000E+00 | 0.000E+00 | 4.863E+05 | 2.883E+05 | 3.730E+06 | 3.153E+06 | 0.000E+00 | 0.000E+00 | 0.000E+00 | 4.485E+05 | 2.883E+05 | 1.799E+06 | 2.014E+06 |
| 646 | 734.471 | 10.261 | 45 | N/A | N/A | 0.000E+00 | 1.910E+04 | 1.299E+07 | 9.888E+05 | 5.012E+05 | 5.770E+07 | 6.629E+07 | 0.000E+00 | 1.086E+04 | 1.063E+07 | 9.334E+05 | 3.034E+05 | 3.196E+07 | 3.867E+07 |
| 652 | 762.502 | 12.706 | 45 | N/A | N/A | 0.000E+00 | 1.522E+05 | 2.449E+05 | 5.253E+05 | 5.188E+05 | 5.621E+06 | 4.210E+06 | 0.000E+00 | 1.522E+05 | 2.258E+05 | 2.924E+05 | 4.034E+05 | 2.527E+06 | 1.791E+06 |
| 370 | 514.353 | 10.382 | 43 | N/A | N/A | 9.242E+05 | 5.311E+05 | 1.432E+06 | 7.181E+05 | 5.974E+05 | 1.072E+06 | 9.373E+05 | 9.859E+04 | 5.817E+04 | 2.505E+05 | 1.774E+05 | 7.535E+04 | 3.771E+04 | 2.193E+05 |
| 740 | 744.593 | 12.218 | 43 | 6.414E+04 | 51.035 | 0.000E+00 | 2.801E+03 | 1.612E+04 | 5.005E+03 | 1.792E+06 | 4.301E+06 | 8.976E+06 | 0.000E+00 | 2.801E+03 | 1.270E+04 | 5.005E+03 | 1.721E+06 | 2.649E+06 | 6.811E+06 |
| 94 | 247.596 | 7.524 | 42 | N/A | N/A | 4.874E+06 | 9.447E+06 | 8.849E+06 | 1.026E+07 | 1.057E+07 | 1.170E+07 | 1.338E+07 | 1.532E+05 | 1.422E+06 | 1.409E+06 | 4.262E+05 | 2.074E+06 | 1.136E+06 | 5.451E+06 |
| 110 | 256.601 | 7.527 | 42 | N/A | N/A | 3.965E+06 | 8.110E+06 | 7.870E+06 | 8.120E+06 | 9.860E+06 | 1.077E+07 | 1.198E+07 | 5.389E+05 | 1.139E+06 | 1.035E+06 | 2.988E+05 | 2.562E+06 | 1.223E+06 | 4.956E+06 |
| 233 | 268.115 | 7.525 | 42 | N/A | N/A | 2.502E+06 | 4.145E+06 | 3.899E+06 | 3.487E+06 | 3.730E+06 | 4.133E+06 | 3.630E+06 | 2.128E+05 | 1.692E+05 | 5.200E+05 | 3.667E+05 | 8.507E+05 | 3.436E+05 | 1.225E+06 |
| 283 | 554.177 | 10.161 | 40 | 1.178E+05 | 74.011 | 2.355E+06 | 1.917E+06 | 1.870E+06 | 1.409E+06 | 1.682E+06 | 1.801E+06 | 1.948E+06 | 1.006E+05 | 1.474E+05 | 1.299E+05 | 8.217E+04 | 1.469E+05 | 2.289E+05 | 2.218E+05 |
| 545 | 684.203 | 10.983 | 40 | N/A | N/A | 1.704E+05 | 8.710E+05 | 9.520E+05 | 1.017E+06 | 1.288E+06 | 1.708E+06 | 1.856E+06 | 6.135E+04 | 7.454E+05 | 7.606E+05 | 9.007E+05 | 1.122E+06 | 1.631E+06 | 1.654E+06 |
| 630 | 702.214 | 10.746 | 40 | 1.178E+05 | 74.027 | 1.073E+06 | 8.043E+05 | 6.706E+05 | 5.738E+05 | 5.135E+05 | 5.478E+05 | 5.196E+05 | 9.875E+04 | 1.362E+05 | 4.574E+04 | 1.605E+04 | 6.061E+04 | 4.398E+04 | 9.628E+03 |
| 693 | 463.131 | 9.688 | 40 | N/A | N/A | 6.290E+05 | 7.028E+05 | 4.567E+05 | 4.876E+05 | 5.552E+05 | 4.819E+05 | 4.951E+05 | 8.135E+03 | 5.785E+04 | 5.767E+04 | 2.188E+04 | 9.686E+04 | 1.940E+04 | 1.102E+05 |
| 821 | 776.232 | 11.006 | 40 | N/A | N/A | 6.206E+05 | 2.898E+05 | 3.427E+05 | 3.080E+05 | 2.389E+05 | 2.612E+05 | 2.608E+05 | 2.064E+04 | 4.093E+04 | 1.315E+04 | 2.418E+04 | 4.074E+04 | 2.168E+04 | 4.500E+04 |
| 858 | 610.185 | 12.140 | 40 | N/A | N/A | 7.022E+05 | 7.875E+05 | 7.365E+05 | 7.704E+05 | 7.722E+05 | 9.314E+05 | 1.037E+06 | 1.050E+05 | 1.234E+05 | 1.488E+05 | 2.823E+04 | 1.941E+05 | 2.092E+05 | 7.781E+04 |
| 143 | 228.196 | 7.148 | 39 | 2.622E+05 | 81.083 | 1.706E+06 | 2.649E+06 | 3.107E+06 | 3.989E+06 | 2.844E+06 | 4.165E+06 | 5.921E+06 | 1.476E+05 | 6.702E+05 | 7.035E+05 | 7.066E+05 | 3.976E+05 | 1.126E+06 | 1.807E+06 |
| 230 | 228.196 | 6.462 | 39 | 2.622E+05 | 81.083 | 8.176E+06 | 1.352E+07 | 1.122E+07 | 1.460E+07 | 9.723E+06 | 1.375E+07 | 1.605E+07 | 7.682E+05 | 3.536E+06 | 2.080E+06 | 2.361E+06 | 1.041E+06 | 3.453E+06 | 4.707E+06 |
| 824 | 549.485 | 10.160 | 39 | 4.854E+04 | 28.033 | 0.000E+00 | 0.000E+00 | 1.309E+05 | 2.537E+05 | 1.303E+05 | 2.536E+06 | 2.920E+07 | 0.000E+00 | 0.000E+00 | 6.566E+04 | 1.272E+05 | 6.461E+04 | 1.125E+06 | 2.085E+07 |
| 826 | 549.486 | 12.704 | 39 | 4.854E+04 | 28.032 | 0.000E+00 | 0.000E+00 | 2.222E+05 | 4.772E+05 | 3.402E+05 | 3.550E+06 | 5.145E+06 | 0.000E+00 | 0.000E+00 | 1.349E+05 | 4.011E+05 | 1.842E+05 | 1.659E+06 | 1.989E+06 |
| 840 | 549.486 | 9.853 | 39 | 7.732E+04 | 46.044 | 0.000E+00 | 1.307E+03 | 2.713E+04 | 3.368E+05 | 2.831E+04 | 2.809E+07 | 6.682E+07 | 0.000E+00 | 1.307E+03 | 4.861E+03 | 2.082E+05 | 1.432E+04 | 2.340E+07 | 6.360E+07 |
| 850 | 549.485 | 10.361 | 39 | 7.732E+04 | 46.045 | 0.000E+00 | 0.000E+00 | 5.180E+05 | 2.191E+05 | 8.865E+04 | 2.037E+07 | 4.634E+06 | 0.000E+00 | 0.000E+00 | 2.684E+05 | 1.922E+05 | 2.800E+04 | 1.909E+07 | 1.313E+06 |
| 159 | 480.238 | 3.674 | 37 | N/A | N/A | 0.000E+00 | 0.000E+00 | 8.413E+06 | 2.843E+06 | 0.000E+00 | 0.000E+00 | 0.000E+00 | 0.000E+00 | 0.000E+00 | 9.895E+05 | 2.843E+06 | 0.000E+00 | 0.000E+00 | 0.000E+00 |
| 332 | 959.468 | 3.674 | 37 | N/A | N/A | 0.000E+00 | 0.000E+00 | 4.008E+06 | 1.298E+06 | 0.000E+00 | 0.000E+00 | 0.000E+00 | 0.000E+00 | 0.000E+00 | 4.387E+05 | 1.298E+06 | 0.000E+00 | 0.000E+00 | 0.000E+00 |
| 301 | 467.250 | 5.033 | 36 | N/A | N/A | 0.000E+00 | 0.000E+00 | 3.403E+06 | 1.227E+06 | 0.000E+00 | 0.000E+00 | 0.000E+00 | 0.000E+00 | 0.000E+00 | 8.924E+05 | 1.227E+06 | 0.000E+00 | 0.000E+00 | 0.000E+00 |
| 333 | 933.494 | 5.044 | 36 | N/A | N/A | 0.000E+00 | 0.000E+00 | 3.665E+06 | 1.266E+06 | 0.000E+00 | 0.000E+00 | 0.000E+00 | 0.000E+00 | 0.000E+00 | 1.077E+06 | 1.265E+06 | 0.000E+00 | 0.000E+00 | 0.000E+00 |
| 420 | 516.785 | 5.231 | 36 | N/A | N/A | 0.000E+00 | 0.000E+00 | 1.881E+06 | 6.738E+05 | 0.000E+00 | 0.000E+00 | 0.000E+00 | 0.000E+00 | 0.000E+00 | 4.239E+05 | 6.738E+05 | 0.000E+00 | 0.000E+00 | 0.000E+00 |
| 68 | 217.105 | 4.009 | 35 | 4.369E+05 | 66.009 | 3.769E+07 | 5.986E+07 | 3.088E+07 | 4.026E+07 | 2.718E+07 | 3.296E+07 | 3.107E+07 | 1.368E+07 | 7.852E+06 | 6.794E+06 | 7.703E+06 | 3.518E+06 | 5.238E+06 | 8.888E+06 |
| 330 | 234.134 | 4.002 | 35 | N/A | N/A | 7.160E+06 | 1.165E+07 | 5.826E+06 | 7.551E+06 | 4.860E+06 | 5.794E+06 | 5.593E+06 | 2.598E+06 | 1.394E+06 | 1.432E+06 | 1.525E+06 | 7.153E+05 | 7.801E+05 | 1.752E+06 |
| 431 | 681.651 | 10.701 | 33 | N/A | N/A | 1.227E+06 | 1.147E+06 | 1.076E+06 | 8.436E+05 | 8.101E+05 | 8.114E+05 | 8.192E+05 | 5.794E+04 | 4.921E+04 | 7.394E+04 | 2.316E+04 | 6.406E+03 | 3.081E+04 | 4.256E+04 |
| 434 | 399.395 | 7.402 | 33 | N/A | N/A | 1.083E+06 | 1.083E+06 | 1.033E+06 | 9.992E+05 | 1.026E+06 | 9.297E+05 | 9.574E+05 | 2.023E+04 | 2.076E+04 | 4.006E+04 | 1.579E+04 | 3.633E+03 | 3.764E+04 | 1.504E+04 |
| 171 | 278.185 | 3.992 | 32 | N/A | N/A | 8.619E+06 | 1.715E+07 | 7.861E+06 | 1.336E+07 | 1.152E+07 | 1.531E+07 | 1.106E+07 | 8.142E+05 | 2.112E+06 | 2.661E+05 | 5.421E+05 | 6.278E+05 | 7.546E+05 | 9.483E+05 |
| 437 | 280.186 | 4.202 | 32 | N/A | N/A | 2.780E+06 | 6.166E+06 | 2.557E+06 | 5.021E+06 | 4.268E+06 | 6.147E+06 | 4.424E+06 | 4.340E+05 | 1.210E+06 | 1.587E+05 | 7.754E+05 | 1.006E+05 | 1.140E+05 | 1.995E+05 |
| 560 | 271.180 | 3.637 | 32 | N/A | N/A | 2.499E+06 | 3.214E+06 | 1.025E+06 | 2.602E+06 | 3.116E+06 | 3.580E+06 | 3.109E+06 | 9.707E+05 | 4.360E+05 | 4.838E+04 | 6.731E+05 | 3.945E+04 | 1.541E+05 | 4.753E+05 |
| 35 | 255.159 | 6.531 | 31 | 3.424E+05 | 65.088 | 1.848E+07 | 2.161E+07 | 2.162E+07 | 2.155E+07 | 2.201E+07 | 2.493E+07 | 2.648E+07 | 8.529E+05 | 4.139E+05 | 9.607E+05 | 9.413E+05 | 1.101E+06 | 9.873E+05 | 4.642E+06 |
| 99 | 272.185 | 6.531 | 31 | N/A | N/A | 6.254E+06 | 7.230E+06 | 7.188E+06 | 7.067E+06 | 7.257E+06 | 8.086E+06 | 8.894E+06 | 2.071E+05 | 3.852E+05 | 3.264E+05 | 2.503E+05 | 3.614E+05 | 1.662E+05 | 1.779E+06 |
| 466 | 269.175 | 7.631 | 31 | N/A | N/A | 1.171E+06 | 3.590E+06 | 2.232E+06 | 3.593E+06 | 9.607E+05 | 3.215E+06 | 4.208E+06 | 5.759E+05 | 1.203E+06 | 8.545E+05 | 2.140E+06 | 2.364E+05 | 1.608E+06 | 2.435E+06 |
| 28 | 297.133 | 7.153 | 30 | N/A | N/A | 1.838E+07 | 2.170E+07 | 1.857E+07 | 2.548E+07 | 2.021E+07 | 2.387E+07 | 1.517E+07 | 1.340E+06 | 1.345E+06 | 2.412E+06 | 1.546E+06 | 3.050E+06 | 2.182E+06 | 4.695E+06 |
| 38 | 431.207 | 7.153 | 30 | 2.208E+05 | 77.984 | 1.259E+07 | 1.452E+07 | 1.227E+07 | 1.679E+07 | 1.282E+07 | 1.551E+07 | 9.588E+06 | 7.452E+05 | 1.005E+06 | 1.386E+06 | 1.028E+06 | 2.141E+06 | 1.468E+06 | 2.878E+06 |
| 56 | 447.202 | 6.602 | 30 | 2.661E+05 | 93.979 | 1.100E+07 | 1.273E+07 | 7.708E+06 | 1.265E+07 | 8.894E+06 | 1.289E+07 | 8.126E+06 | 1.234E+06 | 1.085E+06 | 1.074E+06 | 1.448E+06 | 6.154E+05 | 1.346E+06 | 2.832E+06 |
| 91 | 448.233 | 7.152 | 30 | N/A | N/A | 6.006E+06 | 7.478E+06 | 5.588E+06 | 8.253E+06 | 6.495E+06 | 7.666E+06 | 4.715E+06 | 4.880E+05 | 5.696E+05 | 5.705E+05 | 6.630E+05 | 1.152E+06 | 6.889E+05 | 1.148E+06 |
| 102 | 429.191 | 6.409 | 30 | 2.151E+05 | 75.968 | 6.579E+06 | 7.526E+06 | 5.065E+06 | 9.123E+06 | 6.106E+06 | 9.669E+06 | 6.074E+06 | 6.462E+05 | 5.350E+05 | 5.954E+05 | 1.186E+06 | 4.646E+05 | 1.423E+06 | 2.117E+06 |
| 163 | 297.133 | 7.367 | 30 | N/A | N/A | 3.929E+06 | 5.624E+06 | 4.870E+06 | 5.895E+06 | 5.221E+06 | 6.165E+06 | 3.918E+06 | 5.080E+05 | 8.562E+05 | 5.263E+05 | 5.352E+05 | 1.042E+06 | 4.966E+05 | 5.156E+05 |
| 202 | 448.233 | 7.470 | 30 | N/A | N/A | 2.897E+06 | 3.825E+06 | 2.831E+06 | 3.784E+06 | 4.413E+06 | 4.994E+06 | 3.180E+06 | 4.264E+05 | 4.244E+05 | 2.766E+05 | 3.696E+05 | 1.016E+06 | 6.175E+05 | 4.395E+05 |

| Feature identifiers | | | | | | Average XIC | | | | | | | Standard Err of Average XIC | | | | | | |
|---|---|---|---|---|---|---|---|---|---|---|---|---|---|---|---|---|---|---|---|
| ClusterIdx | m/z | RT [min] | Subnetwork | PPM | MassDiff | TUB00 | 4TUB34 | 4TUB81 | 4TUB93 | 5TUB23 | 5TUB48 | 5TUB83 | TUB00 | 4TUB34 | 4TUB81 | 4TUB93 | 5TUB23 | 5TUB48 | 5TUB83 |
| 208 | 431.207 | 7.472 | 30 | N/A | N/A | 2.929E+06 | 5.647E+06 | 3.732E+06 | 6.379E+06 | 5.968E+06 | 6.839E+06 | 3.056E+06 | 5.521E+05 | 1.480E+06 | 9.658E+05 | 5.763E+05 | 1.138E+06 | 5.610E+05 | 1.202E+06 |
| 213 | 447.202 | 7.074 | 30 | 2.661E+05 | 93.979 | 1.601E+06 | 1.770E+06 | 2.263E+06 | 2.793E+06 | 2.248E+06 | 2.753E+06 | 2.003E+06 | 1.038E+05 | 1.074E+05 | 3.444E+05 | 3.972E+04 | 9.518E+04 | 5.202E+05 | 5.423E+05 |
| 219 | 447.202 | 6.409 | 30 | 2.661E+05 | 93.979 | 3.090E+06 | 3.365E+06 | 2.221E+06 | 4.001E+06 | 2.656E+06 | 4.221E+06 | 2.696E+06 | 2.904E+05 | 2.820E+05 | 2.535E+05 | 5.611E+05 | 1.968E+05 | 6.669E+05 | 8.850E+05 |
| 235 | 297.133 | 6.600 | 30 | N/A | N/A | 2.414E+06 | 2.976E+06 | 1.880E+06 | 3.271E+06 | 2.251E+06 | 3.365E+06 | 2.068E+06 | 3.460E+05 | 2.730E+05 | 2.576E+05 | 3.606E+05 | 1.362E+05 | 4.353E+05 | 7.348E+05 |
| 245 | 469.184 | 6.601 | 30 | N/A | N/A | 2.400E+06 | 3.056E+06 | 1.750E+06 | 3.048E+06 | 2.136E+06 | 3.215E+06 | 2.007E+06 | 3.251E+05 | 3.119E+05 | 2.942E+05 | 3.294E+05 | 1.534E+05 | 3.076E+05 | 7.040E+05 |
| 249 | 429.191 | 6.602 | 30 | 2.151E+05 | 75.968 | 2.547E+06 | 3.062E+06 | 1.751E+06 | 3.036E+06 | 2.102E+06 | 3.133E+06 | 1.955E+06 | 3.050E+05 | 3.271E+05 | 2.830E+05 | 3.866E+05 | 1.449E+05 | 3.399E+05 | 6.780E+05 |
| 251 | 297.133 | 6.410 | 30 | N/A | N/A | 2.240E+06 | 2.641E+06 | 1.739E+06 | 3.181E+06 | 2.096E+06 | 3.465E+06 | 2.168E+06 | 2.212E+05 | 1.780E+05 | 2.956E+05 | 4.282E+05 | 1.197E+05 | 5.176E+05 | 7.862E+05 |
| 262 | 447.202 | 6.882 | 30 | 2.661E+05 | 93.979 | 4.814E+06 | 5.283E+06 | 4.441E+06 | 6.322E+06 | 4.666E+06 | 6.785E+06 | 6.457E+06 | 2.810E+05 | 1.843E+05 | 5.448E+05 | 6.029E+05 | 3.163E+05 | 9.497E+05 | 1.201E+06 |
| 411 | 527.717 | 3.679 | 29 | N/A | N/A | 3.396E+04 | 0.000E+00 | 7.903E+05 | 9.698E+05 | 1.818E+03 | 1.217E+03 | 0.000E+00 | 3.291E+04 | 0.000E+00 | 4.005E+05 | 9.375E+05 | 1.818E+03 | 1.217E+03 | 0.000E+00 |
| 444 | 791.074 | 3.678 | 29 | N/A | N/A | 2.058E+04 | 0.000E+00 | 5.906E+05 | 8.470E+05 | 1.727E+03 | 0.000E+00 | 0.000E+00 | 2.058E+04 | 0.000E+00 | 2.793E+05 | 8.331E+05 | 1.727E+03 | 0.000E+00 | 0.000E+00 |
| 355 | 500.380 | 6.151 | 28 | 3.523E+04 | 17.027 | 4.001E+06 | 3.464E+06 | 3.534E+06 | 3.242E+06 | 3.459E+06 | 3.210E+06 | 3.229E+06 | 1.329E+05 | 1.749E+05 | 6.390E+04 | 7.839E+04 | 5.911E+04 | 3.701E+04 | 4.950E+04 |
| 391 | 442.338 | 5.719 | 28 | 8.485E+04 | 41.015 | 2.875E+06 | 2.870E+06 | 2.739E+06 | 2.754E+06 | 2.776E+06 | 2.779E+06 | 2.663E+06 | 7.906E+04 | 7.724E+04 | 5.602E+04 | 5.229E+04 | 6.273E+04 | 8.519E+04 | 2.589E+04 |
| 394 | 309.227 | 4.773 | 28 | 1.580E+05 | 58.042 | 3.342E+06 | 4.762E+06 | 3.113E+06 | 3.596E+06 | 3.701E+06 | 3.452E+06 | 2.764E+06 | 3.806E+05 | 2.643E+05 | 2.643E+05 | 1.929E+05 | 3.313E+05 | 1.465E+05 | 1.145E+05 |
| 446 | 384.296 | 5.267 | 28 | N/A | N/A | 2.614E+06 | 3.080E+06 | 2.275E+06 | 2.555E+06 | 2.483E+06 | 2.304E+06 | 2.095E+06 | 2.845E+05 | 1.008E+05 | 1.501E+05 | 4.790E+04 | 8.673E+04 | 9.957E+04 | 6.335E+04 |
| 458 | 674.506 | 7.346 | 28 | N/A | N/A | 2.203E+06 | 1.401E+06 | 2.201E+06 | 1.832E+06 | 1.022E+06 | 1.449E+06 | 7.556E+05 | 1.077E+05 | 2.967E+05 | 5.953E+04 | 2.965E+05 | 4.434E+05 | 3.437E+05 | 1.983E+05 |
| 631 | 790.590 | 8.085 | 28 | N/A | N/A | 2.213E+06 | 1.510E+06 | 1.133E+06 | 1.671E+06 | 1.321E+06 | 1.460E+06 | 1.186E+06 | 3.810E+04 | 3.601E+05 | 2.185E+05 | 3.959E+05 | 1.568E+05 | 3.345E+05 | 1.325E+05 |
| 635 | 549.416 | 9.554 | 28 | N/A | N/A | 8.709E+05 | 3.379E+05 | 6.074E+05 | 9.627E+05 | 5.056E+05 | 4.742E+05 | 4.587E+05 | 1.376E+05 | 9.541E+04 | 5.188E+04 | 4.070E+05 | 1.795E+05 | 8.068E+04 | 4.452E+04 |
| 860 | 694.522 | 10.428 | 28 | N/A | N/A | 2.658E+05 | 3.636E+05 | 5.944E+05 | 4.529E+05 | 3.158E+05 | 9.781E+05 | 8.360E+05 | 6.048E+04 | 3.615E+04 | 2.386E+05 | 5.061E+04 | 1.912E+04 | 5.142E+05 | 2.972E+05 |
| 357 | 206.062 | 2.609 | 27 | 9.781E+03 | 1.996 | 0.000E+00 | 5.742E+07 | 1.035E+07 | 3.881E+07 | 5.807E+05 | 1.367E+07 | 3.672E+05 | 0.000E+00 | 1.231E+07 | 5.349E+05 | 1.289E+07 | 3.077E+05 | 1.231E+07 | 3.660E+05 |
| 440 | 237.106 | 4.092 | 27 | 6.284E+04 | 14.019 | 4.233E+04 | 1.516E+07 | 1.014E+06 | 8.669E+06 | 1.276E+06 | 2.305E+06 | 3.812E+04 | 1.280E+04 | 1.718E+06 | 9.772E+04 | 5.384E+05 | 4.683E+05 | 7.110E+05 | 6.280E+03 |
| 699 | 206.061 | 2.395 | 27 | 9.781E+03 | 1.996 | 0.000E+00 | 6.921E+06 | 5.948E+05 | 1.418E+06 | 5.262E+08 | 8.512E+08 | 2.820E+05 | 0.000E+00 | 2.075E+06 | 1.963E+05 | 2.201E+05 | 4.226E+08 | 3.641E+08 | 7.414E+04 |
| 799 | 206.062 | 2.458 | 27 | 9.781E+03 | 1.996 | 5.878E+02 | 2.851E+06 | 5.930E+05 | 1.913E+06 | 8.242E+07 | 3.889E+08 | 4.465E+05 | 5.878E+02 | 5.943E+05 | 1.437E+05 | 7.249E+05 | 3.866E+07 | 2.852E+08 | 1.942E+05 |
| 448 | 213.144 | 5.071 | 26 | N/A | N/A | 2.424E+06 | 1.818E+06 | 1.615E+06 | 1.153E+06 | 9.274E+05 | 9.363E+05 | 8.895E+05 | 1.681E+05 | 7.962E+04 | 6.252E+04 | 2.373E+05 | 2.416E+05 | 2.929E+05 | 1.583E+05 |
| 692 | 381.331 | 9.796 | 26 | N/A | N/A | 1.824E+06 | 8.956E+05 | 5.340E+05 | 4.691E+05 | 5.591E+05 | 5.514E+05 | 6.001E+05 | 7.039E+05 | 2.645E+05 | 8.424E+04 | 5.775E+03 | 2.498E+05 | 6.343E+04 | 1.855E+05 |
| 210 | 445.290 | 4.792 | 25 | N/A | N/A | 2.883E+06 | 2.727E+06 | 2.670E+06 | 2.593E+06 | 2.761E+06 | 3.621E+06 | 2.676E+06 | 1.228E+05 | 1.395E+05 | 4.852E+04 | 3.975E+04 | 3.237E+04 | 3.434E+05 | 2.446E+05 |
| 227 | 467.300 | 4.849 | 25 | N/A | N/A | 2.251E+06 | 2.338E+06 | 2.231E+06 | 2.254E+06 | 2.133E+06 | 3.027E+06 | 2.312E+06 | 8.113E+04 | 2.645E+04 | 4.797E+04 | 6.939E+04 | 1.103E+05 | 2.797E+05 | 5.618E+04 |
| 250 | 423.276 | 4.729 | 25 | N/A | N/A | 2.311E+06 | 2.197E+06 | 2.117E+06 | 2.062E+06 | 2.066E+06 | 2.616E+06 | 2.105E+06 | 1.155E+05 | 1.446E+05 | 5.322E+04 | 8.121E+04 | 4.690E+04 | 2.836E+05 | 7.933E+04 |
| 275 | 401.261 | 4.663 | 25 | N/A | N/A | 1.870E+06 | 1.614E+06 | 1.744E+06 | 1.715E+06 | 1.644E+06 | 2.077E+06 | 1.711E+06 | 2.073E+04 | 1.691E+05 | 9.437E+04 | 6.435E+04 | 4.159E+04 | 1.703E+05 | 3.467E+04 |
| 307 | 379.249 | 4.593 | 25 | N/A | N/A | 1.642E+06 | 1.613E+06 | 1.401E+06 | 1.414E+06 | 1.355E+06 | 1.785E+06 | 1.394E+06 | 1.068E+05 | 9.425E+04 | 4.886E+04 | 8.199E+04 | 3.098E+04 | 1.648E+05 | 4.087E+04 |
| 353 | 489.316 | 4.904 | 25 | N/A | N/A | 1.343E+06 | 1.299E+06 | 1.208E+06 | 1.214E+06 | 1.175E+06 | 1.944E+06 | 1.299E+06 | 3.750E+04 | 7.335E+04 | 6.080E+04 | 1.234E+04 | 4.610E+04 | 2.840E+05 | 6.898E+04 |
| 358 | 505.391 | 10.706 | 25 | N/A | N/A | 1.282E+06 | 1.864E+06 | 1.387E+06 | 1.358E+06 | 2.031E+06 | 1.500E+06 | 1.304E+06 | 9.085E+04 | 2.861E+04 | 2.755E+04 | 7.457E+04 | 2.792E+05 | 6.328E+04 | 1.421E+05 |
| 402 | 527.403 | 10.694 | 25 | N/A | N/A | 1.120E+06 | 1.819E+06 | 1.163E+06 | 1.169E+06 | 1.899E+06 | 1.357E+06 | 1.102E+06 | 6.629E+04 | 6.033E+04 | 2.917E+04 | 1.022E+05 | 3.450E+05 | 8.288E+04 | 9.874E+04 |
| 459 | 549.417 | 10.679 | 25 | N/A | N/A | 7.928E+05 | 1.435E+06 | 6.039E+05 | 6.563E+05 | 1.512E+06 | 9.891E+05 | 7.501E+05 | 1.303E+05 | 7.798E+04 | 5.057E+04 | 1.214E+05 | 3.339E+05 | 1.015E+05 | 4.545E+04 |
| 535 | 463.345 | 9.103 | 25 | N/A | N/A | 2.187E+05 | 1.168E+06 | 1.096E+06 | 1.246E+06 | 4.140E+05 | 4.258E+05 | 3.385E+05 | 3.696E+04 | 4.565E+05 | 4.731E+05 | 1.878E+05 | 8.351E+04 | 1.196E+05 | 1.142E+04 |
| 540 | 441.330 | 9.123 | 25 | N/A | N/A | 2.534E+05 | 1.118E+06 | 1.177E+06 | 1.372E+06 | 5.462E+05 | 4.356E+05 | 2.941E+05 | 3.037E+04 | 3.508E+05 | 6.112E+05 | 2.710E+05 | 7.479E+04 | 2.057E+05 | 3.554E+04 |
| 638 | 529.383 | 9.048 | 25 | N/A | N/A | 2.762E+05 | 1.237E+06 | 4.703E+05 | 5.071E+05 | 3.217E+05 | 4.470E+05 | 2.509E+05 | 8.008E+04 | 1.714E+05 | 2.213E+05 | 1.924E+05 | 6.984E+04 | 1.380E+05 | 5.784E+04 |
| 653 | 507.370 | 9.067 | 25 | N/A | N/A | 2.821E+05 | 1.155E+06 | 5.655E+05 | 7.542E+05 | 3.979E+05 | 4.464E+05 | 2.714E+05 | 8.100E+04 | 3.248E+05 | 2.276E+05 | 8.761E+04 | 7.872E+04 | 1.180E+05 | 2.006E+04 |
| 662 | 511.328 | 4.955 | 25 | N/A | N/A | 5.506E+05 | 4.642E+05 | 5.206E+05 | 4.367E+05 | 4.585E+05 | 8.921E+05 | 4.895E+05 | 7.535E+04 | 1.148E+05 | 4.964E+04 | 9.592E+04 | 4.625E+04 | 1.247E+05 | 7.310E+04 |
| 759 | 571.429 | 10.663 | 25 | N/A | N/A | 5.374E+05 | 1.084E+06 | 4.139E+05 | 3.149E+05 | 1.156E+06 | 5.428E+05 | 4.728E+05 | 1.257E+05 | 5.171E+04 | 4.557E+04 | 6.674E+04 | 2.849E+05 | 1.763E+05 | 3.166E+04 |
| 859 | 533.341 | 5.004 | 25 | N/A | N/A | 3.480E+05 | 2.121E+05 | 1.938E+05 | 1.779E+05 | 1.835E+05 | 5.038E+05 | 2.625E+05 | 2.902E+04 | 3.004E+04 | 6.185E+04 | 2.096E+04 | 4.144E+04 | 1.017E+05 | 2.825E+04 |
| 9 | 460.270 | 7.525 | 24 | 1.085E+05 | 45.059 | 1.202E+08 | 1.734E+08 | 1.742E+08 | 1.789E+08 | 2.047E+08 | 2.135E+08 | 2.271E+08 | 3.481E+06 | 1.147E+07 | 1.291E+07 | 7.359E+06 | 2.207E+07 | 1.142E+07 | 5.234E+07 |
| 25 | 846.443 | 7.526 | 24 | N/A | N/A | 7.909E+06 | 3.361E+07 | 2.856E+07 | 2.902E+07 | 4.403E+07 | 4.936E+07 | 9.007E+07 | 1.449E+06 | 1.044E+07 | 7.916E+06 | 4.113E+06 | 1.533E+07 | 1.075E+07 | 7.064E+07 |
| 67 | 446.254 | 7.523 | 24 | 3.243E+04 | 14.016 | 7.765E+06 | 1.205E+07 | 1.250E+07 | 1.290E+07 | 1.457E+07 | 1.457E+07 | 1.386E+07 | 8.455E+05 | 6.721E+05 | 1.058E+06 | 4.291E+05 | 2.012E+06 | 9.630E+05 | 2.845E+06 |
| 81 | 460.270 | 7.249 | 24 | 6.485E+04 | 28.032 | 2.770E+06 | 5.877E+06 | 8.034E+06 | 9.891E+06 | 9.850E+06 | 9.607E+06 | 9.589E+06 | 1.417E+05 | 2.175E+06 | 6.829E+05 | 4.347E+06 | 2.921E+06 | 3.205E+06 | 3.550E+06 |
| 139 | 476.266 | 7.526 | 24 | 1.019E+05 | 44.028 | 5.064E+06 | 7.067E+06 | 6.394E+06 | 5.893E+06 | 6.322E+06 | 6.620E+06 | 6.868E+06 | 1.836E+05 | 3.616E+05 | 5.595E+05 | 1.720E+05 | 8.120E+05 | 6.981E+05 | 1.753E+06 |
| 261 | 397.201 | 7.526 | 24 | 4.338E+04 | 18.010 | 1.257E+06 | 2.735E+06 | 2.520E+06 | 2.489E+06 | 3.266E+06 | 3.593E+06 | 4.475E+06 | 3.170E+04 | 5.330E+05 | 4.186E+05 | 9.259E+04 | 6.651E+05 | 2.181E+05 | 2.419E+06 |
| 272 | 429.191 | 6.884 | 24 | 3.367E+04 | 13.980 | 5.455E+06 | 7.995E+06 | 5.962E+06 | 7.409E+06 | 7.916E+06 | 8.904E+06 | 1.502E+07 | 4.726E+05 | 3.737E+05 | 5.026E+05 | 4.266E+05 | 4.659E+05 | 7.917E+05 | 8.469E+06 |
| 316 | 531.275 | 7.525 | 24 | N/A | N/A | 3.198E+05 | 2.288E+06 | 1.813E+06 | 1.951E+06 | 2.854E+06 | 3.359E+06 | 9.869E+06 | 3.211E+04 | 8.605E+05 | 7.094E+05 | 4.176E+05 | 1.232E+06 | 9.520E+05 | 8.925E+06 |
| 375 | 462.213 | 6.365 | 24 | N/A | N/A | 1.024E+06 | 2.550E+06 | 1.215E+06 | 1.742E+06 | 2.179E+06 | 2.259E+06 | 1.111E+07 | 2.156E+05 | 9.461E+04 | 4.079E+04 | 1.942E+05 | 4.018E+05 | 1.529E+05 | 9.997E+06 |
| 418 | 445.186 | 6.364 | 24 | 7.219E+04 | 29.975 | 9.228E+05 | 2.141E+06 | 9.861E+05 | 2.042E+06 | 1.865E+06 | 2.535E+06 | 9.346E+06 | 1.697E+05 | 4.023E+04 | 6.318E+03 | 6.420E+05 | 3.480E+05 | 5.344E+05 | 8.399E+06 |
| 419 | 460.270 | 7.814 | 24 | 1.085E+05 | 45.059 | 5.929E+05 | 2.570E+06 | 1.718E+06 | 2.159E+06 | 3.147E+06 | 3.193E+06 | 3.325E+06 | 3.555E+04 | 5.414E+05 | 9.894E+04 | 6.824E+05 | 1.062E+05 | 6.383E+05 | 1.162E+06 |
| 447 | 450.204 | 8.853 | 24 | 8.428E+04 | 34.994 | 6.354E+05 | 1.017E+06 | 8.416E+05 | 8.056E+05 | 1.310E+06 | 1.372E+06 | 1.309E+06 | 5.216E+04 | 1.580E+05 | 1.644E+05 | 5.817E+04 | 1.686E+05 | 1.146E+05 | 4.946E+05 |
| 469 | 445.186 | 6.211 | 24 | 7.219E+04 | 29.975 | 4.454E+05 | 1.018E+06 | 4.829E+05 | 6.754E+05 | 9.415E+05 | 9.164E+05 | 3.574E+06 | 9.695E+04 | 1.075E+04 | 1.686E+04 | 1.415E+05 | 1.521E+05 | 1.218E+05 | 3.202E+06 |
| 578 | 448.233 | 6.302 | 24 | N/A | N/A | 2.689E+05 | 6.491E+05 | 3.707E+05 | 4.786E+05 | 5.198E+05 | 5.650E+05 | 1.394E+06 | 6.908E+04 | 2.803E+04 | 7.356E+04 | 1.549E+04 | 2.170E+04 | 9.659E+04 | 1.161E+06 |
| 582 | 448.233 | 6.159 | 24 | 3.701E+04 | 15.996 | 3.360E+05 | 6.553E+05 | 4.423E+05 | 4.357E+05 | 5.500E+05 | 5.578E+05 | 1.842E+06 | 5.995E+04 | 4.654E+04 | 4.287E+04 | 3.536E+04 | 8.827E+04 | 7.548E+04 | 1.516E+06 |
| 636 | 462.213 | 6.211 | 24 | 6.935E+04 | 29.975 | 2.338E+05 | 5.939E+05 | 2.667E+05 | 4.180E+05 | 5.900E+05 | 5.999E+05 | 2.321E+06 | 1.359E+04 | 2.016E+04 | 6.307E+04 | 1.216E+04 | 1.153E+05 | 5.922E+04 | 2.090E+06 |
| 637 | 431.207 | 6.148 | 24 | 3.852E+04 | 15.996 | 2.454E+05 | 6.370E+05 | 3.606E+05 | 4.335E+05 | 5.391E+05 | 5.017E+05 | 1.670E+06 | 4.133E+04 | 3.104E+04 | 3.875E+04 | 4.016E+04 | 3.940E+04 | 3.825E+04 | 1.369E+06 |
| 770 | 474.249 | 6.864 | 24 | N/A | N/A | 3.452E+05 | 1.012E+06 | 5.129E+05 | 4.669E+05 | 9.234E+05 | 8.177E+05 | 3.685E+06 | 7.185E+04 | 2.374E+05 | 8.860E+04 | 7.385E+04 | 2.288E+05 | 3.072E+04 | 3.161E+06 |
| 790 | 431.207 | 6.021 | 24 | 3.853E+04 | 15.996 | 2.143E+05 | 4.128E+05 | 2.420E+05 | 3.004E+05 | 3.478E+05 | 4.340E+05 | 1.154E+06 | 4.856E+04 | 3.099E+04 | 3.319E+04 | 5.062E+04 | 2.493E+04 | 3.205E+04 | 9.457E+05 |
| 221 | 371.326 | 7.517 | 23 | 8.164E+04 | 28.026 | 2.903E+06 | 4.509E+06 | 2.153E+06 | 2.328E+06 | 4.296E+06 | 2.045E+06 | 2.095E+06 | 4.852E+05 | 2.295E+05 | 1.976E+05 | 1.887E+05 | 5.782E+05 | 1.610E+05 | 3.314E+05 |
| 31 | 328.919 | 12.753 | 22 | 4.052E+05 | 94.843 | 7.685E+07 | 6.089E+07 | 5.516E+07 | 5.098E+07 | 4.899E+07 | 4.454E+07 | 4.360E+07 | 2.522E+06 | 2.804E+06 | 4.007E+05 | 9.499E+05 | 2.064E+06 | 8.317E+05 | 1.241E+06 |
| 39 | 312.943 | 12.757 | 22 | N/A | N/A | 3.513E+07 | 2.637E+07 | 2.879E+07 | 3.895E+07 | 2.035E+07 | 3.682E+07 | 2.931E+07 | 1.182E+07 | 8.725E+06 | 7.202E+06 | 1.373E+06 | 9.893E+06 | 4.371E+05 | 3.620E+06 |
| 70 | 287.888 | 12.760 | 22 | N/A | N/A | 3.715E+07 | 2.871E+07 | 2.642E+07 | 2.472E+07 | 2.355E+07 | 2.269E+07 | 2.272E+07 | 1.790E+06 | 1.501E+06 | 2.332E+05 | 7.278E+05 | 2.404E+05 | 2.117E+05 | 1.146E+06 |
| 92 | 313.078 | 6.460 | 22 | N/A | N/A | 0.000E+00 | 0.000E+00 | 7.353E+06 | 3.646E+05 | 0.000E+00 | 0.000E+00 | 0.000E+00 | 0.000E+00 | 0.000E+00 | 9.358E+05 | 3.646E+05 | 0.000E+00 | 0.000E+00 | 0.000E+00 |
| 187 | 312.943 | 12.757 | 22 | N/A | N/A | 4.669E+07 | 4.215E+07 | 4.048E+07 | 4.021E+07 | 3.844E+07 | 2.924E+07 | 3.225E+07 | 7.658E+05 | 1.034E+06 | 1.134E+06 | 1.564E+05 | 1.110E+06 | 8.014E+06 | 2.142E+06 |
| 196 | 313.078 | 6.532 | 22 | N/A | N/A | 0.000E+00 | 0.000E+00 | 3.834E+06 | 2.645E+05 | 0.000E+00 | 0.000E+00 | 0.000E+00 | 0.000E+00 | 0.000E+00 | 5.630E+05 | 2.645E+05 | 0.000E+00 | 0.000E+00 | 0.000E+00 |
| 255 | 313.235 | 8.431 | 22 | N/A | N/A | 2.048E+06 | 2.745E+06 | 3.328E+06 | 3.598E+06 | 2.852E+06 | 3.535E+06 | 3.956E+06 | 2.512E+05 | 3.345E+05 | 1.456E+05 | 1.732E+05 | 2.804E+05 | 1.864E+05 | 1.260E+06 |
| 558 | 313.235 | 8.075 | 22 | N/A | N/A | 7.649E+04 | 1.101E+05 | 1.010E+06 | 2.788E+07 | 3.823E+06 | 1.082E+07 | 6.278E+06 | 8.227E+03 | 2.880E+04 | 6.738E+05 | 1.600E+07 | 1.579E+06 | 4.062E+06 | 1.366E+06 |
| 197 | 272.258 | 8.673 | 21 | 1.338E+05 | 42.047 | 1.033E+05 | 6.107E+05 | 4.363E+06 | 4.095E+06 | 1.526E+05 | 1.558E+05 | 1.704E+05 | 2.775E+04 | 1.438E+05 | 2.951E+05 | 2.945E+06 | 2.404E+04 | 2.121E+04 | 2.104E+04 |
| 223 | 300.290 | 9.476 | 21 | 4.459E+04 | 14.015 | 2.015E+05 | 4.418E+05 | 2.546E+06 | 2.027E+06 | 8.093E+05 | 5.848E+05 | 6.277E+05 | 4.566E+04 | 8.837E+04 | 1.308E+05 | 1.440E+06 | 1.559E+05 | 7.660E+03 | 3.215E+04 |

| Feature identifiers | | | | | | Average XIC | | | | | | | Standard Err of Average XIC | | | | | | |
|---|---|---|---|---|---|---|---|---|---|---|---|---|---|---|---|---|---|---|---|
| ClusterIdx | m/z | RT [min] | Subnetwork | PPM | MassDiff | TUB00 | 4TUB34 | 4TUB81 | 4TUB93 | 5TUB23 | 5TUB48 | 5TUB83 | TUB00 | 4TUB34 | 4TUB81 | 4TUB93 | 5TUB23 | 5TUB48 | 5TUB83 |
| 460 | 406.354 | 10.232 | 21 | N/A | N/A | 4.444E+05 | 6.283E+05 | 4.396E+05 | 4.828E+05 | 8.136E+05 | 1.008E+06 | 1.414E+06 | 7.826E+04 | 1.921E+04 | 7.717E+04 | 1.452E+05 | 2.021E+05 | 5.248E+04 | 7.470E+05 |
| 157 | 429.146 | 4.153 | 20 | N/A | N/A | 0.000E+00 | 7.626E+04 | 7.731E+06 | 6.248E+05 | 7.143E+03 | 0.000E+00 | 0.000E+00 | 0.000E+00 | 3.769E+04 | 6.317E+05 | 6.248E+05 | 7.143E+03 | 0.000E+00 | 0.000E+00 |
| 166 | 549.571 | 3.384 | 20 | N/A | N/A | 0.000E+00 | 5.080E+03 | 4.785E+06 | 8.924E+04 | 0.000E+00 | 0.000E+00 | 0.000E+00 | 0.000E+00 | 5.080E+03 | 1.883E+06 | 5.631E+04 | 0.000E+00 | 0.000E+00 | 0.000E+00 |
| 481 | 549.078 | 3.211 | 20 | N/A | N/A | 4.069E+06 | 5.491E+05 | 2.841E+05 | 1.655E+07 | 5.657E+05 | 5.089E+05 | 9.129E+04 | 8.931E+05 | 5.137E+05 | 1.018E+05 | 1.518E+07 | 8.666E+04 | 2.700E+05 | 7.984E+04 |
| 520 | 432.026 | 3.379 | 20 | N/A | N/A | 1.335E+07 | 6.422E+07 | 7.106E+05 | 4.881E+07 | 5.176E+07 | 3.550E+07 | 3.891E+07 | 8.732E+06 | 4.426E+06 | 1.998E+05 | 2.619E+07 | 2.687E+07 | 1.974E+07 | 2.080E+07 |
| 584 | 432.026 | 3.502 | 20 | N/A | N/A | 2.547E+07 | 1.516E+08 | 8.797E+05 | 6.915E+07 | 9.375E+06 | 5.108E+07 | 8.535E+07 | 1.172E+07 | 2.255E+07 | 1.088E+05 | 6.592E+07 | 1.911E+06 | 4.368E+07 | 3.824E+07 |
| 596 | 405.571 | 3.038 | 20 | 5.465E+05 | 84.094 | 1.087E+05 | 2.200E+07 | 3.988E+04 | 3.398E+07 | 2.163E+07 | 2.548E+07 | 4.810E+07 | 6.825E+04 | 2.008E+07 | 2.962E+03 | 3.377E+07 | 1.242E+07 | 2.471E+07 | 2.605E+07 |
| 600 | 359.104 | 0.595 | 20 | N/A | N/A | 2.559E+06 | 2.169E+06 | 5.851E+06 | 9.560E+06 | 4.345E+06 | 5.751E+05 | 1.867E+05 | 9.865E+05 | 1.053E+06 | 2.765E+06 | 6.644E+06 | 4.095E+06 | 4.060E+05 | 1.749E+05 |
| 611 | 401.115 | 2.454 | 20 | N/A | N/A | 4.551E+04 | 2.500E+07 | 1.617E+05 | 5.646E+07 | 1.537E+05 | 1.783E+06 | 3.860E+05 | 1.233E+04 | 8.365E+06 | 5.166E+04 | 4.669E+07 | 6.885E+04 | 1.654E+06 | 4.170E+04 |
| 622 | 411.021 | 3.222 | 20 | 5.193E+05 | 34.032 | 2.385E+07 | 1.227E+07 | 8.151E+04 | 5.291E+06 | 4.382E+06 | 1.387E+06 | 4.649E+05 | 2.035E+07 | 6.569E+06 | 3.571E+04 | 4.667E+06 | 3.469E+06 | 1.134E+06 | 2.113E+05 |
| 315 | 515.353 | 9.549 | 19 | N/A | N/A | 0.000E+00 | 1.207E+04 | 3.088E+06 | 8.669E+05 | 1.704E+03 | 0.000E+00 | 0.000E+00 | 0.000E+00 | 2.801E+03 | 1.162E+06 | 8.402E+05 | 1.704E+03 | 0.000E+00 | 0.000E+00 |
| 326 | 321.148 | 3.682 | 19 | N/A | N/A | 7.965E+04 | 4.038E+06 | 2.977E+06 | 2.500E+07 | 3.099E+04 | 1.905E+04 | 2.172E+03 | 2.620E+04 | 8.951E+05 | 6.554E+04 | 1.334E+07 | 3.099E+04 | 1.506E+04 | 2.172E+03 |
| 595 | 366.206 | 3.688 | 19 | N/A | N/A | 1.046E+04 | 1.295E+06 | 8.050E+05 | 7.942E+06 | 8.459E+03 | 1.330E+04 | 0.000E+00 | 1.046E+04 | 2.264E+05 | 1.826E+05 | 4.381E+06 | 4.437E+03 | 1.330E+04 | 0.000E+00 |
| 494 | 453.189 | 7.153 | 18 | N/A | N/A | 7.015E+06 | 8.391E+06 | 7.052E+06 | 9.800E+06 | 7.736E+06 | 9.119E+06 | 5.768E+06 | 3.677E+05 | 3.803E+05 | 9.898E+05 | 7.365E+05 | 1.304E+06 | 7.685E+05 | 1.624E+06 |
| 591 | 453.189 | 7.367 | 18 | N/A | N/A | 3.132E+06 | 4.469E+06 | 3.181E+06 | 4.259E+06 | 3.783E+06 | 5.178E+06 | 3.568E+06 | 3.510E+05 | 4.969E+05 | 1.698E+05 | 2.885E+05 | 6.629E+05 | 3.155E+05 | 9.790E+05 |
| 861 | 453.187 | 7.297 | 18 | N/A | N/A | 2.820E+06 | 4.035E+06 | 1.474E+05 | 2.240E+05 | 2.748E+05 | 7.057E+06 | 8.155E+06 | 7.307E+05 | 7.456E+05 | 3.568E+04 | 6.748E+04 | 6.817E+04 | 6.822E+06 | 2.545E+06 |
| 21 | 453.169 | 7.511 | 17 | N/A | N/A | 3.610E+07 | 1.234E+06 | 7.535E+07 | 7.344E+07 | 5.188E+07 | 4.624E+07 | 6.821E+07 | 2.011E+07 | 1.643E+05 | 8.653E+06 | 1.133E+07 | 2.744E+07 | 2.427E+07 | 5.106E+06 |
| 292 | 453.168 | 7.525 | 17 | N/A | N/A | 7.393E+07 | 8.842E+07 | 5.853E+07 | 9.166E+07 | 6.073E+07 | 7.880E+07 | 8.321E+07 | 1.455E+06 | 1.354E+06 | 2.901E+07 | 5.241E+06 | 2.995E+07 | 1.348E+07 | 1.254E+07 |
| 640 | 453.186 | 7.468 | 17 | N/A | N/A | 8.478E+05 | 1.134E+06 | 2.909E+07 | 1.089E+06 | 7.807E+05 | 9.444E+05 | 2.949E+07 | 2.508E+05 | 1.232E+05 | 2.824E+07 | 7.576E+04 | 1.159E+05 | 2.338E+05 | 2.906E+07 |
| 59 | 281.138 | 7.525 | 16 | N/A | N/A | 6.478E+06 | 1.315E+07 | 1.235E+07 | 1.217E+07 | 1.531E+07 | 1.613E+07 | 2.058E+07 | 5.843E+05 | 1.256E+06 | 1.522E+06 | 7.013E+05 | 3.300E+06 | 1.724E+06 | 9.831E+06 |
| 97 | 299.149 | 4.461 | 16 | 1.283E+05 | 34.005 | 4.127E+06 | 1.012E+07 | 8.741E+06 | 1.046E+07 | 1.381E+07 | 1.662E+07 | 2.293E+07 | 2.789E+05 | 1.853E+06 | 8.456E+05 | 3.097E+06 | 2.335E+06 | 2.280E+06 | 1.059E+07 |
| 172 | 299.148 | 7.523 | 16 | 1.283E+05 | 34.005 | 2.761E+06 | 6.479E+06 | 5.940E+06 | 5.815E+06 | 7.302E+06 | 7.982E+06 | 9.999E+06 | 3.366E+05 | 1.036E+06 | 7.555E+05 | 4.151E+05 | 1.654E+06 | 1.253E+06 | 5.164E+06 |
| 296 | 344.206 | 4.461 | 16 | N/A | N/A | 1.147E+06 | 2.065E+06 | 2.263E+06 | 2.461E+06 | 3.688E+06 | 4.223E+06 | 6.664E+06 | 1.446E+05 | 2.791E+05 | 2.139E+05 | 5.171E+05 | 6.458E+05 | 6.326E+05 | 3.375E+06 |
| 381 | 316.176 | 4.464 | 16 | N/A | N/A | 7.462E+05 | 1.416E+06 | 1.343E+06 | 1.530E+06 | 2.191E+06 | 2.425E+06 | 3.544E+06 | 3.974E+04 | 2.639E+05 | 1.280E+05 | 4.327E+05 | 4.771E+05 | 3.446E+05 | 1.644E+06 |
| 42 | 473.318 | 10.647 | 15 | 1.353E+05 | 74.081 | 6.110E+06 | 4.766E+06 | 7.071E+06 | 5.941E+06 | 6.044E+06 | 6.179E+06 | 5.778E+06 | 6.310E+05 | 3.113E+05 | 5.063E+05 | 1.244E+06 | 5.438E+05 | 1.023E+06 | 6.831E+05 |
| 72 | 680.481 | 12.424 | 15 | N/A | N/A | 2.125E+07 | 2.100E+07 | 1.615E+07 | 1.711E+07 | 4.777E+07 | 5.635E+07 | 5.015E+07 | 1.572E+06 | 1.317E+06 | 2.329E+05 | 1.319E+06 | 5.661E+06 | 1.585E+07 | 1.987E+07 |
| 74 | 475.298 | 9.832 | 15 | 8.479E+04 | 44.036 | 1.726E+07 | 1.493E+07 | 1.539E+07 | 1.798E+07 | 6.023E+07 | 4.798E+07 | 5.408E+07 | 9.766E+05 | 3.867E+06 | 8.904E+05 | 4.486E+06 | 3.031E+06 | 1.585E+07 | 1.759E+07 |
| 95 | 492.324 | 9.832 | 15 | 5.201E+04 | 27.009 | 1.614E+07 | 1.342E+07 | 1.387E+07 | 1.582E+07 | 5.366E+07 | 4.470E+07 | 5.149E+07 | 8.509E+05 | 3.362E+06 | 6.694E+05 | 3.880E+06 | 2.929E+06 | 1.551E+07 | 1.855E+07 |
| 96 | 654.333 | 10.021 | 15 | N/A | N/A | 1.229E+06 | 2.203E+07 | 6.087E+06 | 1.629E+07 | 2.197E+07 | 3.374E+07 | 2.376E+07 | 2.770E+05 | 1.125E+07 | 5.946E+05 | 1.340E+07 | 8.721E+06 | 1.233E+07 | 7.795E+06 |
| 105 | 490.345 | 10.647 | 15 | N/A | N/A | 2.817E+06 | 2.290E+06 | 3.334E+06 | 2.701E+06 | 2.621E+06 | 2.692E+06 | 2.526E+06 | 3.710E+05 | 1.953E+05 | 1.383E+05 | 5.661E+05 | 3.639E+05 | 4.229E+05 | 4.825E+05 |
| 129 | 663.454 | 12.425 | 15 | N/A | N/A | 1.268E+07 | 1.302E+07 | 1.025E+07 | 1.101E+07 | 3.007E+07 | 3.616E+07 | 3.340E+07 | 7.570E+05 | 8.645E+05 | 6.876E+04 | 7.597E+05 | 3.983E+06 | 1.091E+07 | 1.338E+07 |
| 141 | 708.511 | 12.424 | 15 | N/A | N/A | 9.786E+06 | 1.167E+07 | 9.979E+06 | 1.187E+07 | 2.625E+07 | 3.034E+07 | 2.968E+07 | 5.703E+05 | 7.819E+05 | 3.084E+05 | 9.599E+05 | 1.999E+06 | 4.747E+06 | 7.159E+06 |
| 181 | 637.306 | 10.020 | 15 | N/A | N/A | 6.458E+05 | 1.250E+07 | 3.566E+06 | 9.492E+06 | 1.273E+07 | 1.752E+07 | 1.042E+07 | 1.217E+05 | 6.372E+06 | 3.560E+05 | 7.871E+06 | 4.932E+06 | 7.127E+06 | 2.484E+06 |
| 205 | 518.376 | 10.647 | 15 | N/A | N/A | 1.432E+06 | 1.072E+06 | 1.795E+06 | 1.596E+06 | 1.549E+06 | 1.588E+06 | 1.571E+06 | 1.674E+05 | 8.448E+04 | 1.342E+05 | 4.125E+05 | 2.013E+05 | 2.047E+05 | 2.864E+05 |
| 273 | 287.141 | 6.170 | 15 | N/A | N/A | 2.575E+06 | 2.266E+06 | 2.633E+06 | 3.390E+06 | 4.215E+06 | 4.959E+06 | 6.272E+06 | 1.057E+05 | 2.972E+05 | 2.788E+05 | 5.494E+05 | 1.983E+05 | 2.955E+05 | 1.882E+06 |
| 298 | 515.329 | 10.526 | 15 | N/A | N/A | 9.877E+05 | 1.841E+06 | 1.105E+06 | 1.389E+06 | 2.967E+06 | 4.185E+06 | 3.715E+06 | 1.870E+05 | 8.543E+04 | 1.025E+04 | 2.542E+05 | 2.664E+05 | 6.721E+05 | 1.290E+06 |
| 311 | 474.317 | 10.257 | 15 | N/A | N/A | 1.496E+06 | 1.915E+06 | 1.498E+06 | 1.522E+06 | 3.419E+06 | 4.688E+06 | 3.278E+06 | 1.568E+05 | 5.362E+05 | 2.080E+05 | 9.465E+04 | 3.536E+05 | 7.640E+05 | 1.048E+06 |
| 317 | 457.288 | 10.396 | 15 | N/A | N/A | 8.649E+05 | 1.288E+06 | 1.274E+06 | 1.215E+06 | 1.485E+06 | 2.114E+06 | 1.834E+06 | 1.316E+05 | 1.436E+05 | 4.242E+04 | 9.402E+04 | 2.174E+05 | 2.480E+05 | 3.507E+05 |
| 318 | 387.194 | 7.399 | 15 | 4.372E+02 | 0.169 | 5.271E+05 | 6.484E+06 | 1.821E+06 | 2.087E+06 | 3.785E+06 | 7.181E+06 | 3.767E+06 | 1.393E+05 | 3.239E+06 | 1.021E+05 | 1.080E+06 | 1.456E+06 | 4.195E+06 | 1.264E+06 |
| 426 | 682.364 | 10.020 | 15 | N/A | N/A | 1.597E+05 | 3.424E+06 | 9.843E+05 | 2.829E+06 | 3.703E+06 | 8.707E+06 | 7.494E+06 | 4.667E+04 | 1.692E+06 | 1.615E+05 | 2.252E+06 | 1.638E+06 | 2.749E+06 | 1.378E+06 |
| 433 | 532.356 | 10.526 | 15 | N/A | N/A | 4.655E+05 | 7.701E+05 | 5.326E+05 | 6.552E+05 | 1.355E+06 | 1.996E+06 | 1.523E+06 | 6.419E+04 | 1.080E+05 | 5.945E+03 | 1.018E+05 | 1.687E+05 | 3.141E+05 | 6.339E+05 |
| 452 | 519.377 | 10.257 | 15 | N/A | N/A | 7.202E+05 | 8.287E+05 | 5.836E+05 | 6.868E+05 | 1.624E+06 | 1.780E+06 | 1.044E+06 | 3.608E+04 | 1.663E+05 | 5.420E+04 | 7.417E+04 | 1.769E+05 | 6.409E+04 | 2.164E+05 |
| 529 | 520.355 | 9.833 | 15 | 1.967E+03 | 1.022 | 7.162E+05 | 9.772E+05 | 4.691E+05 | 1.823E+06 | 1.200E+07 | 1.957E+07 | 2.496E+07 | 4.649E+04 | 5.027E+05 | 1.639E+05 | 7.164E+05 | 1.329E+06 | 8.098E+06 | 9.380E+06 |
| 726 | 1342.926 | 12.423 | 15 | N/A | N/A | 8.097E+04 | 1.886E+05 | 3.845E+04 | 8.454E+04 | 2.059E+06 | 3.638E+06 | 3.151E+06 | 4.992E+04 | 8.465E+04 | 1.552E+04 | 2.171E+04 | 6.515E+05 | 2.083E+06 | 2.608E+06 |
| 755 | 949.587 | 9.831 | 15 | N/A | N/A | 8.292E+03 | 1.229E+04 | 8.933E+03 | 4.405E+04 | 9.932E+05 | 8.330E+05 | 1.333E+06 | 2.165E+03 | 1.100E+04 | 6.895E+03 | 3.121E+04 | 9.695E+04 | 4.468E+05 | 1.049E+06 |
| 783 | 491.374 | 10.264 | 15 | N/A | N/A | 4.807E+05 | 8.625E+05 | 4.594E+05 | 7.199E+05 | 5.413E+05 | 2.042E+05 | 3.146E+05 | 1.051E+05 | 1.849E+05 | 7.253E+04 | 1.546E+05 | 2.523E+05 | 7.525E+04 | 4.845E+04 |
| 849 | 1370.957 | 12.421 | 15 | N/A | N/A | 4.816E+03 | 1.520E+04 | 6.255E+03 | 1.850E+04 | 7.307E+05 | 7.545E+05 | 9.558E+05 | 4.816E+03 | 1.235E+04 | 6.255E+03 | 4.474E+03 | 1.075E+05 | 2.740E+05 | 8.079E+05 |
| 585 | 834.093 | 3.504 | 14 | N/A | N/A | 5.068E+06 | 5.613E+06 | 2.874E+05 | 7.341E+06 | 3.288E+05 | 6.677E+05 | 4.314E+05 | 1.834E+06 | 3.148E+06 | 1.533E+05 | 6.550E+06 | 2.418E+05 | 3.578E+05 | 2.458E+05 |
| 619 | 792.082 | 3.197 | 14 | N/A | N/A | 9.169E+06 | 1.262E+05 | 2.999E+05 | 1.162E+06 | 3.797E+04 | 4.959E+03 | 1.556E+04 | 1.372E+06 | 9.273E+04 | 1.107E+05 | 6.036E+05 | 3.470E+04 | 4.959E+03 | 1.556E+04 |
| 714 | 542.297 | 4.609 | 13 | N/A | N/A | 0.000E+00 | 0.000E+00 | 0.000E+00 | 0.000E+00 | 4.065E+06 | 2.338E+06 | 4.313E+04 | 0.000E+00 | 0.000E+00 | 0.000E+00 | 0.000E+00 | 1.780E+06 | 7.752E+05 | 1.241E+04 |
| 717 | 492.763 | 4.165 | 13 | N/A | N/A | 0.000E+00 | 0.000E+00 | 0.000E+00 | 0.000E+00 | 4.393E+06 | 2.597E+06 | 4.557E+04 | 0.000E+00 | 0.000E+00 | 0.000E+00 | 0.000E+00 | 1.639E+06 | 9.026E+05 | 1.656E+04 |
| 51 | 247.034 | 5.943 | 12 | N/A | N/A | 0.000E+00 | 1.036E+07 | 1.559E+07 | 2.887E+06 | 2.783E+04 | 0.000E+00 | 0.000E+00 | 0.000E+00 | 3.767E+06 | 1.148E+06 | 2.384E+06 | 2.783E+04 | 0.000E+00 | 0.000E+00 |
| 698 | 247.034 | 6.796 | 12 | N/A | N/A | 0.000E+00 | 4.279E+05 | 3.429E+03 | 3.079E+04 | 4.223E+08 | 9.661E+04 | 7.062E+06 | 0.000E+00 | 2.140E+05 | 1.868E+03 | 3.079E+04 | 1.130E+08 | 4.851E+04 | 3.675E+06 |
| 718 | 291.023 | 6.253 | 12 | N/A | N/A | 0.000E+00 | 0.000E+00 | 0.000E+00 | 0.000E+00 | 7.833E+06 | 0.000E+00 | 6.208E+04 | 0.000E+00 | 0.000E+00 | 0.000E+00 | 0.000E+00 | 1.843E+06 | 0.000E+00 | 3.572E+04 |
| 16 | 122.547 | 12.762 | 11 | N/A | N/A | 2.121E+08 | 2.005E+08 | 1.846E+08 | 1.975E+08 | 1.859E+08 | 1.781E+08 | 1.665E+08 | 5.081E+06 | 3.953E+06 | 2.531E+06 | 3.355E+06 | 1.101E+07 | 6.142E+06 | 1.072E+05 |
| 60 | 122.547 | 14.272 | 11 | N/A | N/A | 4.025E+06 | 7.503E+06 | 1.083E+07 | 1.738E+06 | 4.344E+06 | 2.779E+06 | 7.722E+06 | 1.426E+06 | 3.017E+06 | 3.456E+06 | 1.074E+05 | 2.812E+06 | 9.504E+05 | 4.711E+06 |
| 103 | 122.547 | 2.944 | 11 | N/A | N/A | 4.061E+06 | 3.021E+06 | 4.485E+06 | 7.735E+06 | 6.403E+06 | 2.839E+06 | 6.698E+06 | 1.534E+06 | 1.190E+06 | 2.157E+06 | 5.832E+06 | 1.550E+06 | 4.836E+04 | 4.761E+06 |
| 113 | 122.547 | 2.472 | 11 | N/A | N/A | 6.307E+06 | 3.894E+06 | 4.548E+06 | 1.439E+06 | 3.129E+06 | 3.079E+06 | 1.538E+06 | 8.888E+05 | 1.218E+06 | 6.334E+05 | 5.285E+05 | 5.437E+05 | 1.552E+06 | 2.784E+05 |
| 115 | 122.547 | 13.880 | 11 | N/A | N/A | 1.178E+07 | 3.683E+06 | 7.174E+06 | 5.545E+06 | 2.164E+06 | 6.823E+06 | 3.840E+06 | 1.638E+06 | 2.024E+06 | 1.241E+06 | 2.413E+06 | 1.836E+05 | 3.160E+06 | 2.767E+06 |
| 120 | 122.547 | 2.832 | 11 | N/A | N/A | 3.288E+06 | 5.354E+06 | 4.011E+06 | 2.754E+06 | 2.064E+06 | 1.841E+06 | 2.002E+06 | 5.177E+05 | 2.528E+06 | 4.400E+05 | 1.186E+06 | 3.782E+05 | 5.106E+05 | 4.204E+05 |
| 123 | 122.547 | 13.067 | 11 | N/A | N/A | 3.894E+06 | 5.264E+06 | 3.050E+06 | 2.152E+06 | 3.174E+06 | 3.075E+06 | 1.899E+06 | 8.752E+05 | 2.621E+06 | 8.267E+05 | 6.670E+05 | 7.352E+05 | 1.550E+06 | 4.757E+05 |
| 125 | 122.547 | 2.720 | 11 | N/A | N/A | 5.944E+06 | 5.756E+06 | 3.348E+06 | 6.601E+06 | 2.995E+06 | 2.383E+06 | 4.832E+06 | 1.223E+06 | 5.981E+05 | 1.530E+06 | 3.781E+06 | 4.301E+05 | 1.723E+05 | 4.142E+05 |
| 127 | 122.547 | 0.660 | 11 | N/A | N/A | 7.957E+06 | 2.419E+06 | 8.241E+06 | 1.691E+07 | 6.761E+06 | 6.802E+06 | 2.359E+06 | 3.910E+06 | 1.272E+06 | 9.856E+05 | 5.158E+06 | 3.112E+06 | 2.249E+06 | 1.140E+06 |
| 135 | 122.547 | 13.843 | 11 | N/A | N/A | 6.079E+06 | 8.103E+06 | 5.007E+06 | 8.642E+06 | 2.556E+06 | 4.527E+06 | 5.815E+06 | 2.306E+06 | 2.671E+06 | 1.858E+06 | 3.802E+06 | 2.215E+05 | 1.861E+06 | 4.812E+06 |
| 140 | 122.547 | 2.622 | 11 | N/A | N/A | 6.558E+06 | 5.119E+06 | 2.065E+06 | 1.548E+06 | 3.234E+06 | 1.372E+06 | 3.528E+06 | 1.478E+06 | 2.618E+06 | 1.342E+05 | 6.309E+05 | 2.014E+06 | 4.031E+05 | 1.303E+06 |
| 145 | 122.547 | 2.598 | 11 | N/A | N/A | 3.174E+06 | 5.396E+06 | 3.238E+06 | 3.227E+06 | 1.881E+06 | 5.977E+06 | 2.858E+06 | 8.777E+05 | 1.264E+06 | 9.054E+05 | 7.801E+05 | 1.045E+06 | 7.623E+05 | 1.865E+06 |
| 147 | 122.547 | 2.749 | 11 | N/A | N/A | 4.912E+06 | 2.544E+06 | 1.442E+06 | 1.612E+06 | 3.087E+06 | 5.653E+06 | 2.571E+06 | 3.195E+06 | 6.726E+05 | 1.354E+05 | 3.934E+05 | 5.467E+05 | 3.110E+06 | 1.907E+05 |
| 151 | 122.547 | 13.172 | 11 | N/A | N/A | 4.230E+06 | 6.619E+06 | 3.276E+06 | 5.783E+06 | 3.516E+06 | 2.564E+06 | 3.014E+06 | 1.645E+06 | 1.405E+06 | 4.224E+05 | 1.447E+06 | 4.924E+05 | 2.181E+05 | 1.577E+06 |
| 156 | 122.547 | 2.528 | 11 | N/A | N/A | 3.715E+06 | 3.748E+06 | 1.628E+06 | 6.507E+06 | 8.412E+05 | 2.248E+06 | 2.618E+06 | 1.234E+06 | 1.628E+05 | 3.660E+05 | 5.464E+05 | 4.382E+04 | 3.349E+05 | 6.411E+05 |
| 160 | 122.547 | 2.262 | 11 | N/A | N/A | 2.211E+06 | 3.539E+06 | 2.312E+06 | 3.707E+06 | 3.288E+06 | 2.711E+06 | 1.744E+06 | 2.279E+05 | 1.284E+06 | 7.019E+05 | 1.857E+06 | 5.901E+05 | 4.853E+05 | 9.951E+04 |

| Feature identifiers | | | | | | Average XIC | | | | | | | Standard Err of Average XIC | | | | | | |
|---|---|---|---|---|---|---|---|---|---|---|---|---|---|---|---|---|---|---|---|
| ClusterIdx | m/z | RT [min] | Subnetwork | PPM | MassDiff | TUB00 | 4TUB34 | 4TUB81 | 4TUB93 | 5TUB23 | 5TUB48 | 5TUB83 | TUB00 | 4TUB34 | 4TUB81 | 4TUB93 | 5TUB23 | 5TUB48 | 5TUB83 |
| 169 | 122.547 | 13.121 | 11 | N/A | N/A | 1.305E+06 | 7.297E+06 | 2.931E+06 | 2.917E+06 | 3.385E+06 | 3.597E+06 | 3.289E+06 | 6.093E+05 | 2.313E+06 | 8.992E+05 | 1.039E+06 | 8.052E+05 | 1.242E+06 | 1.801E+06 |
| 174 | 122.547 | 2.436 | 11 | N/A | N/A | 1.460E+06 | 2.895E+06 | 2.951E+06 | 3.981E+06 | 2.984E+06 | 5.518E+06 | 1.683E+06 | 7.168E+05 | 4.867E+05 | 5.371E+05 | 1.456E+06 | 1.088E+06 | 1.084E+06 | 2.213E+05 |
| 177 | 122.547 | 13.217 | 11 | N/A | N/A | 1.791E+06 | 4.083E+06 | 2.128E+06 | 1.399E+06 | 3.235E+06 | 1.871E+06 | 1.596E+06 | 7.203E+05 | 1.097E+06 | 1.067E+06 | 1.305E+05 | 2.656E+05 | 9.702E+05 | 8.506E+05 |
| 178 | 122.547 | 2.294 | 11 | N/A | N/A | 3.700E+06 | 4.677E+06 | 2.043E+06 | 2.825E+06 | 3.909E+06 | 1.916E+06 | 2.445E+06 | 1.644E+06 | 1.726E+06 | 9.251E+05 | 1.461E+06 | 1.134E+06 | 4.654E+05 | 3.086E+05 |
| 183 | 122.547 | 1.983 | 11 | N/A | N/A | 2.292E+06 | 1.069E+06 | 1.787E+06 | 2.430E+06 | 1.690E+06 | 3.441E+06 | 8.247E+05 | 8.856E+05 | 4.075E+05 | 8.709E+05 | 8.543E+05 | 5.645E+05 | 1.578E+06 | 1.017E+05 |
| 185 | 122.547 | 13.651 | 11 | N/A | N/A | 6.348E+06 | 1.590E+06 | 2.606E+06 | 3.698E+06 | 1.598E+06 | 3.890E+06 | 1.817E+06 | 7.661E+05 | 2.223E+05 | 9.985E+05 | 1.058E+06 | 7.225E+05 | 2.068E+05 | 1.075E+06 |
| 189 | 122.547 | 0.617 | 11 | N/A | N/A | 6.365E+06 | 1.820E+07 | 5.262E+06 | 1.129E+07 | 7.211E+06 | 2.986E+06 | 5.753E+06 | 3.300E+06 | 1.927E+06 | 1.739E+06 | 4.421E+06 | 3.848E+06 | 1.519E+06 | 2.064E+06 |
| 206 | 122.547 | 1.515 | 11 | N/A | N/A | 2.070E+06 | 7.843E+05 | 2.907E+06 | 1.604E+06 | 1.607E+06 | 9.856E+05 | 1.118E+06 | 4.755E+05 | 1.485E+05 | 7.934E+05 | 6.531E+05 | 6.674E+05 | 1.494E+05 | 1.711E+05 |
| 207 | 122.547 | 2.189 | 11 | N/A | N/A | 3.448E+06 | 3.357E+06 | 3.282E+06 | 1.695E+06 | 2.887E+06 | 3.014E+06 | 1.466E+06 | 4.707E+05 | 4.034E+05 | 1.038E+06 | 6.740E+05 | 1.298E+06 | 9.361E+05 | 3.520E+05 |
| 216 | 122.547 | 1.421 | 11 | N/A | N/A | 1.128E+06 | 2.337E+06 | 1.367E+06 | 1.421E+06 | 1.368E+06 | 1.466E+06 | 7.479E+05 | 4.095E+05 | 1.274E+06 | 3.572E+05 | 7.861E+05 | 2.731E+05 | 5.312E+05 | 1.955E+05 |
| 220 | 122.547 | 2.347 | 11 | N/A | N/A | 3.412E+06 | 4.018E+06 | 2.712E+06 | 1.985E+06 | 2.842E+06 | 6.962E+06 | 1.723E+06 | 6.917E+05 | 2.291E+06 | 8.553E+05 | 7.175E+05 | 1.042E+06 | 3.451E+06 | 6.019E+05 |
| 248 | 122.547 | 2.158 | 11 | N/A | N/A | 2.790E+06 | 9.101E+05 | 1.951E+06 | 3.281E+06 | 3.184E+06 | 3.619E+06 | 1.694E+06 | 2.107E+05 | 7.770E+04 | 9.184E+05 | 1.436E+06 | 6.868E+05 | 8.695E+05 | 2.710E+05 |
| 587 | 122.547 | 2.681 | 11 | N/A | N/A | 2.633E+06 | 6.575E+06 | 2.745E+06 | 1.926E+06 | 1.394E+06 | 1.889E+06 | 3.203E+06 | 1.156E+06 | 2.711E+06 | 1.574E+06 | 4.972E+05 | 5.335E+05 | 5.756E+05 | 1.695E+06 |
| 644 | 122.547 | 2.919 | 11 | N/A | N/A | 7.744E+06 | 6.851E+06 | 3.952E+06 | 2.304E+06 | 3.918E+06 | 1.829E+07 | 3.296E+06 | 3.503E+06 | 3.155E+06 | 2.253E+06 | 7.077E+05 | 9.441E+05 | 1.625E+07 | 1.647E+06 |
| 645 | 122.547 | 2.389 | 11 | N/A | N/A | 3.264E+06 | 1.439E+06 | 9.525E+05 | 2.836E+06 | 2.445E+06 | 3.810E+06 | 2.283E+06 | 9.011E+05 | 4.066E+05 | 2.489E+05 | 9.333E+05 | 6.032E+05 | 1.386E+06 | 7.423E+05 |
| 656 | 122.547 | 2.584 | 11 | N/A | N/A | 3.646E+06 | 3.188E+06 | 1.456E+06 | 1.516E+06 | 4.816E+06 | 1.921E+06 | 9.632E+05 | 8.193E+05 | 1.206E+06 | 7.047E+05 | 5.309E+05 | 1.915E+06 | 8.853E+05 | 2.581E+05 |
| 657 | 122.547 | 2.792 | 11 | N/A | N/A | 2.241E+06 | 3.669E+06 | 1.921E+06 | 4.172E+06 | 2.578E+06 | 4.912E+06 | 3.004E+06 | 6.796E+05 | 1.338E+05 | 1.001E+06 | 2.211E+06 | 1.138E+06 | 2.270E+06 | 9.643E+05 |
| 710 | 122.547 | 3.154 | 11 | N/A | N/A | 1.375E+07 | 7.218E+06 | 9.135E+06 | 5.399E+06 | 1.029E+07 | 1.072E+07 | 1.728E+07 | 3.322E+06 | 2.478E+06 | 1.770E+06 | 8.137E+05 | 2.749E+06 | 3.231E+06 | 8.334E+06 |
| 715 | 122.547 | 3.094 | 11 | N/A | N/A | 3.625E+06 | 6.910E+06 | 9.050E+06 | 6.981E+06 | 9.867E+06 | 1.814E+07 | 3.414E+06 | 1.289E+06 | 2.501E+06 | 5.649E+06 | 1.612E+06 | 2.771E+06 | 1.426E+07 | 8.541E+05 |
| 719 | 122.547 | 2.222 | 11 | N/A | N/A | 6.302E+06 | 1.600E+06 | 1.161E+06 | 1.762E+06 | 2.191E+06 | 2.838E+06 | 1.887E+06 | 3.252E+06 | 4.287E+05 | 3.699E+05 | 5.213E+05 | 3.941E+05 | 7.150E+05 | 7.109E+05 |
| 773 | 122.547 | 13.630 | 11 | N/A | N/A | 1.622E+06 | 1.982E+06 | 1.847E+06 | 1.962E+06 | 2.536E+06 | 1.469E+06 | 3.068E+06 | 7.210E+05 | 1.155E+06 | 6.415E+05 | 4.244E+05 | 1.525E+06 | 2.762E+05 | 1.573E+06 |
| 61 | 537.536 | 11.439 | 10 | N/A | N/A | 6.097E+07 | 5.067E+07 | 5.227E+07 | 4.519E+07 | 4.362E+07 | 4.310E+07 | 5.038E+07 | 3.156E+06 | 3.585E+06 | 4.103E+05 | 3.202E+06 | 2.713E+06 | 3.733E+06 | 1.457E+06 |
| 109 | 565.567 | 11.404 | 10 | N/A | N/A | 6.398E+07 | 5.316E+07 | 4.691E+07 | 4.507E+07 | 4.618E+07 | 4.803E+07 | 4.769E+07 | 2.970E+06 | 1.651E+06 | 3.020E+06 | 2.771E+06 | 5.547E+05 | 9.619E+05 | 2.466E+06 |
| 295 | 582.594 | 11.445 | 10 | N/A | N/A | 6.175E+06 | 6.142E+06 | 6.268E+06 | 6.797E+06 | 7.147E+06 | 7.104E+06 | 8.106E+06 | 5.440E+05 | 8.299E+04 | 5.847E+04 | 8.956E+04 | 7.425E+04 | 2.256E+05 | 1.584E+05 |
| 816 | 593.598 | 10.672 | 10 | N/A | N/A | 2.069E+07 | 1.161E+06 | 8.983E+05 | 8.259E+05 | 9.239E+05 | 1.083E+06 | 7.651E+05 | 1.487E+07 | 3.466E+05 | 3.084E+05 | 2.346E+05 | 1.690E+05 | 3.266E+05 | 2.190E+05 |
| 191 | 608.393 | 9.449 | 9 | N/A | N/A | 3.933E+06 | 9.549E+06 | 9.190E+06 | 1.079E+07 | 4.388E+06 | 5.789E+06 | 4.101E+06 | 7.586E+05 | 9.986E+05 | 8.528E+05 | 1.242E+06 | 3.327E+05 | 2.442E+05 | 4.573E+05 |
| 313 | 638.404 | 9.642 | 9 | 1.090E+05 | 78.119 | 8.859E+05 | 1.986E+06 | 1.687E+06 | 1.811E+06 | 7.665E+05 | 8.701E+05 | 7.860E+05 | 1.147E+05 | 8.519E+04 | 1.535E+05 | 2.572E+05 | 1.314E+05 | 8.594E+04 | 1.059E+05 |
| 348 | 624.388 | 9.076 | 9 | 1.286E+05 | 92.135 | 7.878E+05 | 1.620E+06 | 1.590E+06 | 1.872E+06 | 8.506E+05 | 1.028E+06 | 7.244E+05 | 1.591E+05 | 1.775E+04 | 2.052E+05 | 5.500E+05 | 4.847E+04 | 1.437E+05 | 4.465E+04 |
| 15 | 824.976 | 3.461 | 8 | N/A | N/A | 3.946E+07 | 1.500E+08 | 8.406E+07 | 1.449E+08 | 1.064E+08 | 3.207E+08 | 3.907E+08 | 5.355E+06 | 3.256E+07 | 7.432E+06 | 5.895E+07 | 2.105E+07 | 1.282E+08 | 7.728E+07 |
| 18 | 810.973 | 3.183 | 8 | N/A | N/A | 2.600E+07 | 2.294E+08 | 1.011E+08 | 1.759E+08 | 1.875E+08 | 5.539E+08 | 2.517E+08 | 5.215E+06 | 4.829E+07 | 9.463E+06 | 2.525E+07 | 7.938E+07 | 1.073E+08 | 5.989E+07 |
| 26 | 838.980 | 3.758 | 8 | N/A | N/A | 2.295E+08 | 8.539E+07 | 3.901E+07 | 1.288E+08 | 7.599E+07 | 1.766E+08 | 5.385E+08 | 2.670E+07 | 2.361E+07 | 4.377E+06 | 6.323E+07 | 3.328E+07 | 1.005E+08 | 9.941E+07 |
| 65 | 1215.954 | 3.181 | 8 | N/A | N/A | 4.621E+06 | 3.779E+07 | 1.715E+07 | 3.053E+07 | 2.938E+07 | 9.541E+07 | 3.780E+07 | 9.346E+05 | 7.643E+06 | 1.274E+06 | 4.367E+06 | 9.597E+06 | 1.534E+07 | 7.971E+06 |
| 87 | 1236.959 | 3.462 | 8 | N/A | N/A | 8.253E+06 | 2.942E+07 | 1.599E+07 | 2.797E+07 | 1.921E+07 | 5.760E+07 | 6.831E+07 | 1.179E+06 | 6.443E+06 | 1.475E+06 | 1.207E+07 | 3.828E+06 | 2.193E+07 | 1.034E+07 |
| 121 | 837.625 | 3.464 | 8 | N/A | N/A | 3.875E+06 | 1.490E+07 | 9.271E+06 | 1.440E+07 | 1.100E+07 | 2.828E+07 | 3.519E+07 | 4.583E+05 | 3.697E+06 | 7.287E+05 | 5.923E+06 | 1.776E+06 | 8.582E+06 | 5.847E+06 |
| 136 | 823.621 | 3.183 | 8 | N/A | N/A | 2.531E+06 | 2.359E+07 | 1.142E+07 | 1.888E+07 | 1.996E+07 | 5.000E+07 | 2.558E+07 | 5.299E+05 | 4.781E+06 | 1.156E+06 | 2.484E+06 | 7.575E+06 | 6.558E+06 | 4.823E+06 |
| 194 | 618.984 | 3.459 | 8 | N/A | N/A | 2.093E+06 | 1.010E+07 | 5.321E+06 | 9.888E+06 | 6.210E+06 | 2.125E+07 | 2.640E+07 | 3.524E+05 | 2.398E+06 | 5.695E+05 | 4.296E+06 | 1.091E+06 | 8.893E+06 | 5.488E+06 |
| 204 | 1257.965 | 3.758 | 8 | N/A | N/A | 4.698E+07 | 1.668E+07 | 7.520E+06 | 2.524E+07 | 1.438E+07 | 3.501E+07 | 1.043E+08 | 4.259E+06 | 4.572E+06 | 5.617E+05 | 1.254E+07 | 6.819E+06 | 2.059E+07 | 1.651E+07 |
| 240 | 852.983 | 4.057 | 8 | N/A | N/A | 2.770E+08 | 1.672E+07 | 4.789E+06 | 3.285E+07 | 2.328E+07 | 4.013E+07 | 1.983E+08 | 3.853E+07 | 5.438E+06 | 4.655E+05 | 1.632E+07 | 1.348E+07 | 2.730E+07 | 3.195E+07 |
| 244 | 846.306 | 3.758 | 8 | N/A | N/A | 1.862E+07 | 7.997E+06 | 3.753E+06 | 1.253E+07 | 7.680E+06 | 1.504E+07 | 4.533E+07 | 5.065E+06 | 2.067E+06 | 4.729E+05 | 6.202E+06 | 3.347E+06 | 7.352E+06 | 5.930E+06 |
| 253 | 1069.394 | 4.071 | 8 | N/A | N/A | 2.252E+06 | 1.441E+07 | 6.164E+06 | 1.041E+07 | 1.816E+07 | 7.020E+07 | 3.147E+07 | 5.151E+05 | 3.750E+06 | 9.576E+05 | 4.118E+06 | 5.276E+06 | 1.182E+07 | 6.306E+06 |
| 257 | 851.628 | 3.759 | 8 | N/A | N/A | 2.103E+07 | 8.521E+06 | 4.071E+06 | 1.173E+07 | 7.584E+06 | 1.560E+07 | 4.439E+07 | 1.990E+06 | 2.348E+06 | 4.779E+05 | 5.801E+06 | 3.337E+06 | 8.004E+06 | 6.427E+06 |
| 258 | 608.481 | 3.183 | 8 | N/A | N/A | 7.817E+05 | 1.076E+07 | 4.122E+06 | 7.782E+06 | 9.121E+06 | 2.634E+07 | 1.162E+07 | 1.805E+05 | 2.632E+06 | 4.243E+05 | 1.278E+06 | 4.715E+06 | 3.951E+06 | 2.815E+06 |
| 288 | 1083.397 | 4.311 | 8 | N/A | N/A | 5.521E+07 | 8.230E+06 | 3.057E+06 | 7.504E+06 | 3.064E+07 | 1.163E+08 | 1.776E+08 | 6.815E+06 | 2.524E+06 | 6.759E+05 | 4.571E+06 | 1.357E+07 | 5.120E+07 | 2.823E+07 |
| 289 | 629.487 | 3.760 | 8 | N/A | N/A | 1.614E+07 | 5.701E+06 | 2.599E+06 | 1.010E+07 | 4.894E+06 | 1.293E+07 | 4.411E+07 | 2.497E+06 | 1.800E+06 | 4.326E+05 | 5.025E+06 | 2.362E+06 | 7.656E+06 | 9.173E+06 |
| 312 | 1055.391 | 3.820 | 8 | N/A | N/A | 3.354E+05 | 1.250E+07 | 4.729E+06 | 8.051E+06 | 1.272E+07 | 6.150E+07 | 9.379E+06 | 2.210E+05 | 2.709E+06 | 6.482E+05 | 8.463E+05 | 5.909E+06 | 8.405E+06 | 1.964E+06 |
| 352 | 1234.935 | 3.184 | 8 | N/A | N/A | 3.372E+05 | 2.212E+06 | 1.764E+06 | 3.208E+06 | 2.739E+06 | 8.430E+06 | 3.921E+06 | 1.107E+05 | 1.072E+06 | 2.669E+05 | 5.561E+05 | 5.510E+05 | 9.681E+05 | 5.392E+05 |
| 406 | 802.298 | 4.071 | 8 | N/A | N/A | 3.379E+05 | 5.218E+06 | 2.192E+06 | 3.673E+06 | 6.897E+06 | 2.781E+07 | 1.414E+07 | 4.919E+04 | 1.497E+06 | 2.660E+05 | 1.499E+06 | 2.160E+06 | 4.232E+06 | 2.946E+06 |
| 445 | 812.801 | 4.312 | 8 | N/A | N/A | 2.468E+07 | 3.706E+06 | 1.145E+06 | 4.263E+06 | 1.409E+07 | 5.877E+07 | 9.810E+07 | 2.704E+06 | 1.154E+06 | 1.502E+05 | 2.158E+06 | 4.965E+06 | 2.599E+07 | 1.962E+07 |
| 450 | 1041.387 | 3.604 | 8 | N/A | N/A | 3.158E+04 | 2.691E+06 | 2.048E+06 | 4.102E+06 | 2.141E+06 | 1.019E+07 | 6.810E+05 | 4.025E+03 | 6.509E+05 | 2.732E+05 | 3.126E+06 | 1.670E+06 | 1.247E+06 | 3.972E+05 |
| 457 | 1255.942 | 3.463 | 8 | N/A | N/A | 9.566E+05 | 2.353E+06 | 1.579E+06 | 3.032E+06 | 2.098E+06 | 4.476E+06 | 6.104E+06 | 2.061E+05 | 5.176E+05 | 5.892E+05 | 1.322E+06 | 4.688E+05 | 8.377E+05 | 5.762E+05 |
| 497 | 1278.970 | 4.056 | 8 | N/A | N/A | 6.195E+07 | 2.925E+06 | 5.701E+05 | 6.960E+06 | 4.669E+06 | 7.900E+06 | 4.004E+07 | 7.621E+06 | 1.004E+06 | 1.116E+05 | 3.494E+06 | 2.727E+06 | 5.745E+06 | 7.091E+06 |
| 504 | 1268.955 | 3.761 | 8 | N/A | N/A | 7.772E+06 | 2.195E+06 | 1.258E+06 | 3.681E+06 | 2.471E+06 | 5.351E+06 | 1.209E+07 | 7.870E+05 | 5.654E+05 | 9.464E+04 | 1.855E+06 | 9.912E+05 | 2.737E+06 | 1.055E+06 |
| 511 | 865.632 | 4.058 | 8 | N/A | N/A | 2.166E+07 | 1.367E+06 | 3.654E+05 | 3.142E+06 | 2.068E+06 | 3.317E+06 | 1.701E+07 | 2.261E+06 | 4.603E+05 | 2.813E+04 | 1.573E+06 | 1.301E+06 | 2.423E+06 | 2.518E+06 |
| 514 | 639.989 | 4.058 | 8 | N/A | N/A | 2.806E+07 | 1.337E+06 | 3.291E+05 | 3.015E+06 | 2.158E+06 | 3.763E+06 | 2.239E+07 | 3.974E+06 | 4.355E+05 | 3.758E+04 | 1.508E+06 | 1.355E+06 | 2.583E+06 | 3.449E+06 |
| 515 | 860.310 | 4.057 | 8 | N/A | N/A | 2.257E+07 | 1.325E+06 | 2.264E+05 | 3.148E+06 | 2.242E+06 | 3.389E+06 | 1.746E+07 | 2.519E+06 | 5.920E+05 | 8.330E+04 | 1.577E+06 | 1.453E+06 | 2.409E+06 | 2.667E+06 |
| 518 | 1276.945 | 3.762 | 8 | N/A | N/A | 5.047E+06 | 2.145E+06 | 9.848E+05 | 2.750E+06 | 1.831E+06 | 3.686E+06 | 9.105E+06 | 2.531E+05 | 6.026E+05 | 2.074E+05 | 1.380E+06 | 8.568E+05 | 2.220E+06 | 1.351E+06 |
| 519 | 856.952 | 3.759 | 8 | N/A | N/A | 5.059E+06 | 1.899E+06 | 7.281E+05 | 2.761E+06 | 1.336E+06 | 3.072E+06 | 6.531E+06 | 8.939E+05 | 4.999E+05 | 1.769E+05 | 1.374E+06 | 7.887E+05 | 1.711E+06 | 2.346E+05 |
| 522 | 1090.726 | 4.311 | 8 | N/A | N/A | 7.374E+06 | 1.099E+06 | 3.536E+05 | 1.328E+06 | 4.480E+06 | 1.418E+07 | 2.281E+07 | 1.004E+06 | 3.942E+05 | 1.677E+05 | 6.715E+05 | 1.701E+06 | 5.402E+06 | 3.650E+06 |
| 531 | 1076.722 | 4.075 | 8 | N/A | N/A | 8.618E+04 | 1.944E+06 | 8.719E+05 | 1.387E+06 | 2.861E+06 | 1.098E+07 | 3.252E+06 | 3.114E+04 | 7.700E+05 | 1.277E+05 | 5.495E+05 | 9.153E+05 | 1.726E+06 | 7.695E+05 |
| 533 | 1096.048 | 4.311 | 8 | N/A | N/A | 5.114E+06 | 4.848E+05 | 1.978E+05 | 8.676E+05 | 3.226E+06 | 1.011E+07 | 1.522E+07 | 5.752E+05 | 1.860E+05 | 1.089E+05 | 4.338E+05 | 1.223E+06 | 3.754E+06 | 1.948E+06 |
| 534 | 791.796 | 3.823 | 8 | N/A | N/A | 6.712E+04 | 3.745E+06 | 9.047E+05 | 2.322E+06 | 3.986E+06 | 1.914E+07 | 3.199E+06 | 4.362E+04 | 1.061E+06 | 2.917E+05 | 2.852E+05 | 2.208E+06 | 2.403E+06 | 7.426E+05 |
| 537 | 810.973 | 3.375 | 8 | N/A | N/A | 2.385E+04 | 7.127E+06 | 6.350E+04 | 1.422E+06 | 6.074E+06 | 2.150E+07 | 1.698E+07 | 1.271E+04 | 1.687E+06 | 5.831E+03 | 6.996E+05 | 3.637E+06 | 1.501E+07 | 7.759E+06 |
| 559 | 824.976 | 3.689 | 8 | N/A | N/A | 2.481E+05 | 5.910E+06 | 6.720E+04 | 1.259E+06 | 6.576E+06 | 1.841E+07 | 2.481E+07 | 1.108E+05 | 1.027E+06 | 3.267E+04 | 7.090E+05 | 4.976E+06 | 1.499E+07 | 1.065E+07 |
| 561 | 1289.962 | 4.060 | 8 | N/A | N/A | 9.176E+06 | 3.966E+05 | 5.858E+04 | 1.096E+06 | 7.686E+05 | 9.392E+05 | 6.357E+06 | 9.611E+05 | 1.484E+05 | 2.509E+04 | 5.503E+05 | 5.520E+05 | 6.974E+05 | 8.653E+05 |
| 567 | 1082.043 | 4.072 | 8 | N/A | N/A | 0.000E+00 | 1.072E+06 | 4.218E+05 | 9.482E+05 | 1.822E+06 | 7.212E+06 | 1.552E+06 | 0.000E+00 | 3.052E+05 | 9.329E+04 | 4.670E+05 | 5.678E+05 | 8.996E+05 | 8.263E+05 |
| 588 | 1027.384 | 3.427 | 8 | N/A | N/A | 0.000E+00 | 8.020E+05 | 4.857E+05 | 3.019E+06 | 6.780E+05 | 3.316E+06 | 1.699E+05 | 0.000E+00 | 2.059E+05 | 6.533E+04 | 2.748E+06 | 5.471E+05 | 1.023E+06 | 3.547E+04 |
| 590 | 1013.379 | 3.335 | 8 | N/A | N/A | 2.913E+05 | 6.534E+05 | 4.029E+04 | 2.478E+06 | 4.280E+05 | 5.023E+06 | 3.873E+05 | 3.874E+04 | 1.990E+05 | 1.770E+04 | 2.457E+06 | 4.015E+05 | 6.226E+05 | 1.978E+05 |
| 610 | 770.790 | 3.427 | 8 | N/A | N/A | 5.768E+03 | 2.088E+05 | 5.107E+04 | 8.545E+05 | 1.816E+05 | 1.021E+06 | 2.719E+04 | 5.768E+03 | 9.331E+04 | 9.443E+02 | 8.161E+05 | 1.623E+05 | 3.687E+05 | 6.265E+03 |
| 614 | 837.291 | 3.461 | 8 | N/A | N/A | 2.807E+06 | 1.076E+07 | 5.807E+06 | 1.140E+07 | 7.211E+06 | 2.328E+07 | 2.454E+07 | 5.692E+05 | 3.534E+06 | 1.292E+06 | 4.681E+06 | 1.833E+06 | 7.720E+06 | 6.003E+06 |
| 620 | 760.288 | 3.338 | 8 | N/A | N/A | 1.752E+04 | 8.498E+04 | 1.034E+04 | 6.194E+05 | 1.140E+05 | 1.071E+06 | 4.525E+04 | 8.667E+03 | 5.356E+04 | 7.473E+03 | 6.194E+05 | 1.120E+05 | 2.369E+05 | 5.743E+03 |

| Feature identifiers | | | | | | Average XIC | | | | | | | Standard Err of Average XIC | | | | | | |
|---|---|---|---|---|---|---|---|---|---|---|---|---|---|---|---|---|---|---|---|
| ClusterIdx | m/z | RT [min] | Subnetwork | PPM | MassDiff | TUB00 | 4TUB34 | 4TUB81 | 4TUB93 | 5TUB23 | 5TUB48 | 5TUB83 | TUB00 | 4TUB34 | 4TUB81 | 4TUB93 | 5TUB23 | 5TUB48 | 5TUB83 |
| 633 | 1062.719 | 3.822 | 8 | N/A | N/A | 2.000E+04 | 1.789E+06 | 3.345E+05 | 9.283E+05 | 1.900E+06 | 7.962E+06 | 1.316E+06 | 2.000E+04 | 6.968E+05 | 1.103E+05 | 2.642E+05 | 1.082E+06 | 3.969E+05 | 3.884E+05 |
| 650 | 629.236 | 3.761 | 8 | N/A | N/A | 8.365E+06 | 4.160E+06 | 1.979E+06 | 7.153E+06 | 4.053E+06 | 9.631E+06 | 3.416E+07 | 1.679E+06 | 1.934E+06 | 2.137E+05 | 3.622E+06 | 1.570E+06 | 5.880E+06 | 5.775E+06 |
| 659 | 813.049 | 4.312 | 8 | N/A | N/A | 2.042E+07 | 2.576E+06 | 7.508E+05 | 3.789E+06 | 1.232E+07 | 4.742E+07 | 8.514E+07 | 1.904E+06 | 8.072E+05 | 3.321E+05 | 1.883E+06 | 4.609E+06 | 2.569E+07 | 1.770E+07 |
| 664 | 866.987 | 4.353 | 8 | N/A | N/A | 5.290E+07 | 1.634E+05 | 0.000E+00 | 4.413E+05 | 5.754E+05 | 6.036E+05 | 6.355E+06 | 6.634E+06 | 1.223E+05 | 0.000E+00 | 2.472E+05 | 3.312E+05 | 5.401E+05 | 1.602E+06 |
| 672 | 1299.976 | 4.354 | 8 | N/A | N/A | 9.462E+06 | 4.053E+03 | 1.581E+03 | 2.985E+04 | 2.503E+04 | 0.000E+00 | 5.457E+05 | 1.264E+06 | 4.053E+03 | 1.581E+03 | 1.492E+04 | 1.295E+04 | 0.000E+00 | 1.960E+05 |
| 676 | 874.315 | 4.356 | 8 | N/A | N/A | 4.599E+06 | 0.000E+00 | 0.000E+00 | 1.005E+04 | 1.081E+04 | 3.941E+04 | 3.490E+05 | 6.971E+05 | 0.000E+00 | 0.000E+00 | 1.005E+04 | 1.081E+04 | 3.941E+04 | 1.772E+05 |
| 677 | 856.617 | 3.758 | 8 | N/A | N/A | 7.516E+06 | 2.033E+06 | 8.351E+05 | 1.930E+06 | 1.295E+06 | 3.227E+06 | 9.224E+06 | 1.137E+06 | 5.632E+05 | 8.880E+04 | 1.172E+06 | 7.123E+05 | 2.002E+06 | 1.020E+06 |
| 678 | 879.636 | 4.362 | 8 | N/A | N/A | 3.742E+06 | 0.000E+00 | 0.000E+00 | 2.546E+03 | 9.712E+03 | 0.000E+00 | 2.020E+05 | 4.975E+05 | 0.000E+00 | 0.000E+00 | 2.546E+03 | 9.712E+03 | 0.000E+00 | 1.027E+05 |
| 680 | 1297.948 | 4.058 | 8 | N/A | N/A | 5.513E+06 | 3.323E+05 | 5.690E+04 | 8.280E+05 | 3.310E+05 | 2.376E+05 | 3.908E+06 | 4.351E+05 | 1.597E+05 | 9.902E+03 | 4.156E+05 | 1.772E+05 | 9.234E+04 | 4.990E+05 |
| 683 | 975.108 | 4.244 | 8 | N/A | N/A | 1.061E+06 | 2.014E+05 | 5.661E+04 | 1.364E+05 | 4.927E+05 | 5.681E+06 | 9.280E+06 | 5.431E+05 | 9.016E+04 | 1.177E+04 | 6.689E+04 | 1.292E+05 | 2.365E+06 | 2.359E+06 |
| 685 | 1101.371 | 4.312 | 8 | N/A | N/A | 1.547E+06 | 6.572E+04 | 1.067E+04 | 1.187E+05 | 4.842E+05 | 1.189E+06 | 2.617E+06 | 2.225E+05 | 4.168E+04 | 4.825E+03 | 8.288E+04 | 2.625E+05 | 2.125E+05 | 4.873E+05 |
| 691 | 822.287 | 4.312 | 8 | N/A | N/A | 1.155E+06 | 5.378E+04 | 2.051E+04 | 9.214E+04 | 6.308E+05 | 1.822E+06 | 4.176E+06 | 9.301E+04 | 2.401E+04 | 6.566E+03 | 5.773E+04 | 2.787E+05 | 7.485E+05 | 1.006E+06 |
| 730 | 796.969 | 3.070 | 8 | N/A | N/A | 0.000E+00 | 2.799E+06 | 2.672E+04 | 2.082E+05 | 3.640E+06 | 2.169E+07 | 6.232E+06 | 0.000E+00 | 8.284E+05 | 5.234E+03 | 6.491E+04 | 1.759E+06 | 1.168E+07 | 3.908E+06 |
| 741 | 1068.040 | 3.824 | 8 | N/A | N/A | 0.000E+00 | 8.973E+05 | 2.819E+05 | 3.651E+05 | 1.089E+06 | 5.110E+06 | 7.880E+05 | 0.000E+00 | 2.579E+05 | 8.297E+04 | 1.763E+05 | 5.881E+05 | 4.182E+05 | 1.843E+05 |
| 746 | 1069.395 | 4.241 | 8 | N/A | N/A | 5.417E+04 | 1.283E+06 | 1.255E+04 | 1.425E+05 | 5.881E+06 | 2.494E+07 | 2.568E+07 | 3.964E+04 | 2.515E+05 | 6.353E+03 | 7.171E+04 | 4.444E+06 | 1.869E+07 | 1.090E+07 |
| 751 | 1221.446 | 3.186 | 8 | N/A | N/A | 0.000E+00 | 5.273E+05 | 1.405E+05 | 4.362E+05 | 4.169E+05 | 1.416E+06 | 6.253E+05 | 0.000E+00 | 2.303E+05 | 4.264E+04 | 1.623E+05 | 2.405E+05 | 5.610E+05 | 2.373E+05 |
| 757 | 1055.390 | 3.992 | 8 | N/A | N/A | 6.976E+03 | 1.052E+06 | 8.637E+03 | 1.062E+05 | 1.134E+06 | 7.588E+06 | 3.235E+06 | 4.451E+03 | 1.379E+05 | 8.637E+03 | 5.325E+04 | 5.689E+05 | 4.356E+06 | 1.416E+06 |
| 760 | 975.359 | 4.241 | 8 | N/A | N/A | 1.487E+06 | 2.226E+05 | 3.555E+04 | 3.161E+05 | 9.837E+05 | 5.834E+06 | 9.348E+06 | 2.452E+05 | 9.389E+04 | 6.268E+03 | 2.465E+05 | 4.204E+05 | 2.456E+06 | 2.405E+06 |
| 761 | 802.298 | 4.239 | 8 | N/A | N/A | 7.645E+04 | 4.181E+05 | 2.282E+03 | 6.999E+04 | 3.184E+06 | 1.484E+07 | 1.669E+07 | 4.911E+04 | 1.334E+05 | 2.282E+03 | 5.033E+04 | 2.460E+06 | 1.147E+07 | 7.628E+06 |
| 766 | 618.733 | 3.459 | 8 | N/A | N/A | 1.020E+06 | 8.923E+06 | 2.797E+06 | 7.508E+06 | 5.764E+06 | 1.739E+07 | 2.032E+07 | 3.185E+05 | 2.211E+06 | 4.736E+05 | 3.133E+06 | 9.512E+05 | 7.139E+06 | 6.269E+06 |
| 768 | 838.979 | 3.988 | 8 | N/A | N/A | 2.066E+05 | 1.746E+06 | 3.570E+03 | 2.383E+05 | 3.900E+06 | 6.741E+06 | 1.508E+07 | 1.209E+05 | 2.139E+05 | 3.570E+03 | 1.237E+05 | 3.241E+06 | 6.034E+06 | 5.978E+06 |
| 778 | 796.636 | 3.069 | 8 | N/A | N/A | 0.000E+00 | 2.940E+06 | 1.570E+04 | 1.664E+05 | 1.626E+06 | 1.554E+07 | 5.162E+06 | 0.000E+00 | 7.466E+05 | 4.672E+03 | 5.698E+04 | 7.575E+05 | 7.304E+06 | 3.532E+06 |
| 785 | 1215.954 | 3.392 | 8 | N/A | N/A | 0.000E+00 | 7.851E+05 | 8.660E+03 | 9.774E+04 | 5.606E+05 | 2.771E+06 | 1.789E+06 | 0.000E+00 | 2.814E+05 | 5.299E+03 | 5.528E+04 | 4.225E+05 | 2.154E+06 | 9.634E+05 |
| 787 | 608.481 | 3.384 | 8 | N/A | N/A | 0.000E+00 | 4.329E+05 | 2.677E+03 | 4.212E+04 | 3.644E+05 | 1.959E+06 | 1.690E+06 | 0.000E+00 | 1.733E+05 | 2.677E+03 | 2.522E+04 | 2.614E+05 | 1.411E+06 | 9.328E+05 |
| 788 | 1236.958 | 3.690 | 8 | N/A | N/A | 4.864E+03 | 4.723E+05 | 5.709E+03 | 1.054E+05 | 8.178E+05 | 2.216E+06 | 3.422E+06 | 4.864E+03 | 6.843E+04 | 5.709E+03 | 5.306E+04 | 7.172E+05 | 1.998E+06 | 1.421E+06 |
| 789 | 618.734 | 3.694 | 8 | N/A | N/A | 7.388E+03 | 3.582E+05 | 0.000E+00 | 3.705E+04 | 4.395E+05 | 1.785E+06 | 2.592E+06 | 3.807E+03 | 9.578E+04 | 0.000E+00 | 2.026E+04 | 3.700E+05 | 1.615E+06 | 1.174E+06 |
| 791 | 792.044 | 3.972 | 8 | N/A | N/A | 0.000E+00 | 2.314E+05 | 1.107E+04 | 5.791E+04 | 2.712E+05 | 2.659E+06 | 1.258E+06 | 0.000E+00 | 9.915E+04 | 7.077E+03 | 2.290E+04 | 1.216E+05 | 1.624E+06 | 7.189E+05 |
| 796 | 1069.727 | 4.238 | 8 | N/A | N/A | 1.636E+05 | 9.568E+05 | 7.482E+03 | 2.071E+05 | 5.019E+06 | 2.158E+07 | 2.209E+07 | 1.070E+05 | 1.845E+05 | 3.791E+03 | 1.071E+05 | 3.706E+06 | 1.613E+07 | 9.022E+06 |
| 800 | 824.643 | 3.674 | 8 | N/A | N/A | 1.810E+05 | 5.315E+06 | 3.576E+04 | 1.013E+06 | 5.675E+06 | 1.520E+07 | 2.306E+07 | 1.235E+05 | 8.597E+05 | 1.439E+04 | 5.653E+05 | 4.705E+06 | 1.235E+07 | 9.111E+06 |
| 801 | 1055.390 | 4.161 | 8 | N/A | N/A | 0.000E+00 | 4.477E+04 | 0.000E+00 | 1.866E+04 | 1.144E+06 | 3.570E+06 | 2.251E+06 | 0.000E+00 | 1.516E+04 | 0.000E+00 | 9.369E+03 | 1.083E+06 | 3.455E+06 | 1.687E+06 |
| 803 | 791.795 | 4.163 | 8 | N/A | N/A | 0.000E+00 | 9.255E+04 | 0.000E+00 | 6.539E+03 | 8.411E+05 | 2.876E+06 | 2.014E+06 | 0.000E+00 | 1.921E+04 | 0.000E+00 | 6.539E+03 | 7.952E+05 | 2.594E+06 | 1.117E+06 |
| 804 | 1300.145 | 4.240 | 8 | N/A | N/A | 4.656E+04 | 5.775E+04 | 2.126E+04 | 2.995E+04 | 5.275E+05 | 3.151E+06 | 7.012E+06 | 1.034E+04 | 3.316E+04 | 8.607E+03 | 1.632E+04 | 4.479E+05 | 1.953E+06 | 1.526E+06 |
| 805 | 810.973 | 3.576 | 8 | N/A | N/A | 0.000E+00 | 2.153E+05 | 6.370E+03 | 1.041E+04 | 5.599E+05 | 1.097E+06 | 6.805E+05 | 0.000E+00 | 1.207E+05 | 6.370E+03 | 1.041E+04 | 5.431E+05 | 1.061E+06 | 5.381E+05 |
| 807 | 1082.043 | 4.246 | 8 | N/A | N/A | 1.310E+03 | 2.746E+04 | 0.000E+00 | 0.000E+00 | 5.028E+05 | 1.800E+06 | 2.179E+06 | 1.310E+03 | 1.237E+04 | 0.000E+00 | 0.000E+00 | 4.468E+05 | 1.425E+06 | 9.214E+05 |
| 809 | 629.487 | 3.997 | 8 | N/A | N/A | 6.181E+03 | 1.434E+05 | 0.000E+00 | 2.126E+04 | 3.183E+05 | 7.632E+05 | 2.029E+06 | 6.181E+03 | 7.366E+04 | 0.000E+00 | 1.065E+04 | 2.791E+05 | 7.431E+05 | 9.997E+05 |
| 812 | 1268.461 | 3.759 | 8 | N/A | N/A | 6.316E+06 | 2.361E+06 | 1.010E+06 | 3.585E+06 | 1.407E+06 | 4.237E+06 | 1.206E+07 | 6.074E+05 | 6.179E+05 | 3.192E+04 | 1.790E+06 | 2.976E+05 | 2.738E+06 | 1.851E+06 |
| 820 | 1300.134 | 4.241 | 8 | N/A | N/A | 1.054E+06 | 3.374E+05 | 2.197E+04 | 1.466E+05 | 4.283E+05 | 2.743E+06 | 1.707E+06 | 1.548E+05 | 1.645E+05 | 1.105E+04 | 8.650E+04 | 6.234E+04 | 1.131E+06 | 3.112E+05 |
| 829 | 1137.663 | 4.195 | 8 | N/A | N/A | 9.515E+04 | 6.464E+03 | 0.000E+00 | 8.220E+03 | 9.146E+04 | 1.098E+06 | 2.454E+06 | 5.186E+04 | 6.464E+03 | 0.000E+00 | 8.220E+03 | 5.037E+04 | 7.110E+05 | 6.582E+05 |
| 830 | 1263.457 | 3.755 | 8 | N/A | N/A | 6.790E+05 | 1.029E+05 | 1.030E+04 | 1.746E+05 | 7.392E+04 | 1.981E+05 | 1.948E+06 | 1.172E+05 | 3.208E+04 | 1.030E+04 | 8.793E+04 | 5.120E+04 | 1.109E+05 | 6.708E+05 |
| 833 | 1076.722 | 4.246 | 8 | N/A | N/A | 5.027E+03 | 5.010E+04 | 0.000E+00 | 3.785E+03 | 5.777E+05 | 2.325E+06 | 2.705E+06 | 5.027E+03 | 1.680E+04 | 0.000E+00 | 3.785E+03 | 5.288E+05 | 1.725E+06 | 1.199E+06 |
| 836 | 1063.052 | 3.825 | 8 | N/A | N/A | 1.153E+04 | 1.869E+06 | 4.605E+05 | 6.469E+05 | 1.510E+06 | 6.753E+06 | 8.276E+05 | 1.153E+04 | 4.410E+05 | 1.164E+05 | 1.462E+05 | 1.067E+06 | 2.151E+05 | 3.097E+05 |
| 838 | 791.796 | 3.996 | 8 | N/A | N/A | 3.417E+03 | 4.047E+05 | 0.000E+00 | 2.318E+04 | 1.650E+05 | 1.806E+06 | 1.247E+06 | 3.417E+03 | 1.779E+05 | 0.000E+00 | 1.240E+04 | 8.054E+04 | 5.687E+05 | 6.415E+05 |
| 842 | 822.537 | 4.313 | 8 | N/A | N/A | 7.758E+05 | 1.015E+05 | 1.025E+04 | 6.661E+04 | 3.801E+05 | 2.356E+06 | 3.234E+06 | 9.372E+04 | 5.118E+04 | 5.484E+02 | 4.476E+04 | 1.091E+05 | 1.169E+06 | 2.528E+05 |
| 845 | 1257.962 | 3.997 | 8 | N/A | N/A | 8.223E+03 | 6.125E+04 | 0.000E+00 | 6.502E+03 | 1.928E+05 | 6.239E+05 | 7.941E+05 | 8.223E+03 | 1.561E+04 | 0.000E+00 | 6.502E+03 | 1.842E+05 | 6.007E+05 | 1.848E+05 |
| 847 | 781.292 | 3.601 | 8 | N/A | N/A | 5.044E+03 | 3.991E+05 | 2.623E+05 | 1.217E+06 | 2.478E+05 | 2.383E+06 | 1.396E+05 | 2.697E+03 | 1.720E+05 | 3.201E+04 | 9.203E+05 | 1.937E+05 | 3.484E+05 | 7.181E+04 |
| 848 | 811.784 | 4.077 | 8 | N/A | N/A | 0.000E+00 | 1.508E+05 | 2.367E+04 | 1.113E+05 | 1.028E+05 | 1.232E+06 | 4.181E+05 | 0.000E+00 | 6.648E+04 | 1.223E+04 | 5.640E+04 | 2.535E+04 | 1.509E+05 | 1.426E+05 |
| 851 | 1276.945 | 3.763 | 8 | N/A | N/A | 4.346E+06 | 1.341E+06 | 9.848E+05 | 2.437E+06 | 1.794E+06 | 3.120E+06 | 5.772E+06 | 7.047E+05 | 5.466E+05 | 2.074E+05 | 1.262E+06 | 8.215E+05 | 2.470E+06 | 1.229E+06 |
| 854 | 1194.947 | 3.069 | 8 | N/A | N/A | 0.000E+00 | 1.074E+05 | 0.000E+00 | 8.264E+03 | 1.629E+05 | 2.100E+06 | 4.125E+05 | 0.000E+00 | 4.003E+04 | 0.000E+00 | 4.978E+03 | 7.593E+04 | 1.215E+06 | 3.108E+05 |
| 862 | 964.605 | 4.154 | 8 | N/A | N/A | 0.000E+00 | 5.402E+03 | 0.000E+00 | 4.739E+03 | 1.247E+05 | 1.141E+06 | 1.135E+06 | 0.000E+00 | 5.402E+03 | 0.000E+00 | 4.739E+03 | 1.169E+05 | 8.021E+05 | 6.266E+05 |
| 354 | 828.611 | 3.182 | 7 | N/A | N/A | 9.432E+05 | 6.092E+06 | 3.048E+06 | 4.949E+06 | 4.864E+06 | 1.299E+07 | 5.433E+06 | 2.245E+05 | 1.580E+06 | 3.087E+05 | 6.674E+05 | 1.807E+06 | 1.568E+06 | 1.866E+06 |
| 360 | 842.615 | 3.460 | 7 | N/A | N/A | 1.199E+06 | 4.078E+06 | 2.182E+06 | 3.566E+06 | 2.017E+06 | 6.351E+06 | 7.308E+06 | 3.379E+05 | 6.827E+05 | 2.851E+05 | 1.489E+06 | 8.905E+05 | 1.698E+06 | 7.157E+05 |
| 649 | 842.948 | 3.459 | 7 | N/A | N/A | 7.443E+05 | 3.975E+06 | 1.640E+06 | 3.341E+06 | 2.237E+06 | 6.214E+06 | 6.636E+06 | 2.937E+05 | 8.893E+05 | 1.713E+05 | 1.350E+06 | 5.149E+05 | 1.928E+06 | 2.534E+06 |
| 811 | 828.945 | 3.185 | 7 | N/A | N/A | 9.093E+05 | 6.771E+06 | 1.974E+06 | 4.081E+06 | 3.875E+06 | 1.048E+07 | 4.816E+06 | 1.574E+05 | 1.382E+06 | 4.411E+05 | 6.303E+05 | 2.348E+06 | 2.242E+06 | 7.123E+05 |
| 22 | 274.274 | 6.821 | 6 | 4.862E+04 | 14.017 | 4.419E+07 | 4.232E+07 | 4.304E+07 | 4.328E+07 | 4.289E+07 | 4.167E+07 | 4.232E+07 | 6.880E+05 | 3.186E+05 | 3.848E+05 | 8.827E+05 | 5.450E+05 | 2.085E+05 | 2.959E+05 |
| 104 | 302.305 | 7.664 | 6 | 4.861E+04 | 14.014 | 8.766E+06 | 9.126E+06 | 9.231E+06 | 9.631E+06 | 1.009E+07 | 1.000E+07 | 1.048E+07 | 1.517E+05 | 2.938E+05 | 1.049E+05 | 1.740E+05 | 1.421E+05 | 2.237E+05 | 2.215E+05 |
| 277 | 330.337 | 8.495 | 6 | 1.458E+05 | 42.046 | 3.903E+06 | 3.942E+06 | 4.025E+06 | 4.209E+06 | 4.245E+06 | 4.192E+06 | 4.384E+06 | 3.766E+04 | 4.659E+04 | 1.558E+05 | 7.542E+04 | 7.558E+04 | 2.166E+04 | 1.740E+05 |
| 341 | 358.368 | 9.237 | 6 | 2.431E+05 | 70.078 | 3.472E+06 | 3.312E+06 | 3.490E+06 | 3.869E+06 | 3.468E+06 | 3.614E+06 | 3.659E+06 | 1.289E+05 | 6.570E+04 | 4.543E+04 | 2.191E+05 | 1.552E+05 | 1.087E+05 | 3.379E+05 |
| 321 | 311.185 | 7.063 | 5 | 3.948E+05 | 88.087 | 1.928E+06 | 2.005E+06 | 2.006E+06 | 1.958E+06 | 2.014E+06 | 2.016E+06 | 2.037E+06 | 5.599E+04 | 5.151E+04 | 4.146E+04 | 6.467E+04 | 3.451E+04 | 2.500E+04 | 2.090E+04 |
| 347 | 423.310 | 10.200 | 5 | N/A | N/A | 1.304E+06 | 2.376E+06 | 4.093E+06 | 3.249E+06 | 1.376E+07 | 3.202E+05 | 8.267E+05 | 3.345E+05 | 2.659E+05 | 2.552E+06 | 1.674E+06 | 9.623E+06 | 8.023E+04 | 7.688E+05 |
| 661 | 468.369 | 10.166 | 5 | N/A | N/A | 1.111E+05 | 2.008E+05 | 5.619E+05 | 4.785E+05 | 2.055E+06 | 4.440E+04 | 4.271E+04 | 1.995E+04 | 3.309E+04 | 4.229E+05 | 2.441E+05 | 1.422E+06 | 2.946E+04 | 4.271E+04 |
| 11 | 452.277 | 7.678 | 4 | 5.438E+04 | 26.010 | 1.811E+07 | 1.272E+08 | 1.238E+08 | 1.954E+08 | 3.481E+08 | 2.566E+08 | 3.795E+08 | 1.945E+06 | 4.174E+07 | 1.109E+07 | 8.618E+07 | 7.482E+07 | 1.648E+07 | 6.809E+07 |
| 155 | 474.259 | 7.679 | 4 | 5.581E+04 | 28.031 | 1.092E+06 | 7.281E+06 | 5.266E+06 | 9.793E+06 | 1.491E+07 | 1.314E+07 | 1.823E+07 | 1.070E+05 | 2.146E+06 | 3.734E+05 | 3.840E+06 | 2.904E+06 | 6.861E+05 | 2.524E+06 |
| 238 | 474.258 | 7.808 | 4 | 4.234E+03 | 2.017 | 1.128E+05 | 1.165E+06 | 3.026E+06 | 2.930E+06 | 5.888E+06 | 2.730E+06 | 3.636E+06 | 3.591E+04 | 3.159E+05 | 3.506E+05 | 9.134E+05 | 1.160E+06 | 7.341E+05 | 7.016E+05 |
| 350 | 690.508 | 12.705 | 4 | 3.631E+04 | 26.015 | 0.000E+00 | 3.233E+06 | 2.285E+07 | 9.209E+06 | 1.387E+07 | 2.936E+07 | 1.234E+08 | 0.000E+00 | 1.530E+06 | 1.115E+07 | 5.746E+06 | 6.738E+06 | 1.202E+07 | 7.017E+07 |
| 424 | 511.304 | 8.562 | 4 | 6.677E+04 | 32.003 | 4.943E+05 | 1.115E+05 | 1.229E+06 | 1.173E+06 | 1.489E+07 | 6.115E+06 | 2.848E+07 | 5.302E+04 | 3.170E+04 | 3.712E+05 | 6.268E+05 | 1.342E+06 | 2.868E+06 | 1.040E+07 |
| 486 | 690.508 | 10.348 | 4 | 3.631E+04 | 26.015 | 0.000E+00 | 1.753E+07 | 3.285E+07 | 5.869E+07 | 1.161E+08 | 9.975E+07 | 1.077E+08 | 0.000E+00 | 9.101E+06 | 1.595E+07 | 3.337E+07 | 6.859E+07 | 4.656E+07 | 1.949E+07 |
| 489 | 712.490 | 12.701 | 4 | 7.051E+04 | 54.045 | 0.000E+00 | 1.293E+06 | 1.086E+07 | 4.152E+06 | 6.166E+06 | 1.126E+07 | 4.434E+07 | 0.000E+00 | 6.755E+05 | 5.154E+06 | 2.833E+06 | 2.764E+06 | 4.485E+06 | 2.284E+07 |
| 505 | 712.490 | 10.356 | 4 | 7.051E+04 | 54.046 | 0.000E+00 | 1.289E+06 | 2.635E+07 | 1.022E+07 | 2.356E+07 | 2.880E+07 | 2.720E+07 | 0.000E+00 | 8.740E+05 | 2.197E+07 | 8.166E+06 | 1.698E+07 | 1.114E+07 | 7.418E+06 |
| 510 | 485.288 | 8.510 | 4 | 1.037E+04 | 4.979 | 7.504E+05 | 4.631E+05 | 1.119E+06 | 1.789E+06 | 1.779E+07 | 1.103E+07 | 2.940E+07 | 9.156E+04 | 9.880E+04 | 2.886E+05 | 9.666E+05 | 4.634E+06 | 3.308E+06 | 1.027E+07 |
| 513 | 743.484 | 12.700 | 4 | 6.777E+04 | 54.046 | 0.000E+00 | 4.713E+04 | 1.750E+04 | 1.263E+06 | 7.794E+05 | 1.151E+07 | 1.374E+07 | 0.000E+00 | 4.713E+04 | 1.750E+04 | 9.401E+05 | 7.597E+05 | 5.366E+06 | 8.340E+06 |

| Feature identifiers | | | | | | Average XIC | | | | | | | Standard Err of Average XIC | | | | | | |
|---|---|---|---|---|---|---|---|---|---|---|---|---|---|---|---|---|---|---|---|
| ClusterIdx | m/z | RT [min] | Subnetwork | PPM | MassDiff | TUB00 | 4TUB34 | 4TUB81 | 4TUB93 | 5TUB23 | 5TUB48 | 5TUB83 | TUB00 | 4TUB34 | 4TUB81 | 4TUB93 | 5TUB23 | 5TUB48 | 5TUB83 |
| 516 | 483.272 | 7.795 | 4 | 7.092E+04 | 32.002 | 2.794E+05 | 8.229E+05 | 8.924E+05 | 1.528E+06 | 1.297E+07 | 9.536E+06 | 2.438E+07 | 9.651E+04 | 1.839E+05 | 2.750E+05 | 5.412E+05 | 3.471E+06 | 2.324E+06 | 8.430E+06 |
| 517 | 740.522 | 12.702 | 4 | 3.394E+04 | 26.014 | 0.000E+00 | 6.745E+05 | 8.932E+05 | 1.316E+06 | 2.062E+06 | 1.929E+07 | 1.848E+07 | 0.000E+00 | 6.745E+05 | 7.233E+05 | 5.463E+05 | 1.220E+06 | 8.661E+06 | 9.043E+06 |
| 532 | 738.526 | 10.287 | 4 | 7.110E+04 | 49.026 | 2.154E+03 | 9.099E+02 | 1.206E+04 | 9.133E+06 | 1.558E+05 | 2.954E+07 | 6.357E+07 | 2.154E+03 | 9.099E+02 | 7.322E+03 | 8.925E+06 | 8.080E+04 | 2.580E+07 | 2.906E+07 |
| 554 | 903.547 | 7.681 | 4 | N/A | N/A | 0.000E+00 | 1.062E+06 | 4.549E+05 | 1.825E+06 | 3.971E+06 | 2.857E+06 | 5.550E+06 | 0.000E+00 | 5.230E+05 | 9.956E+04 | 1.272E+06 | 1.509E+06 | 4.508E+05 | 1.704E+06 |
| 555 | 467.277 | 8.509 | 4 | 7.858E+04 | 34.045 | 2.928E+05 | 7.405E+04 | 4.170E+05 | 6.177E+05 | 7.451E+06 | 4.483E+06 | 1.257E+07 | 7.614E+04 | 9.869E+03 | 1.400E+05 | 3.616E+05 | 1.933E+06 | 1.354E+06 | 4.504E+06 |
| 562 | 743.484 | 10.306 | 4 | 6.777E+04 | 54.046 | 0.000E+00 | 0.000E+00 | 0.000E+00 | 5.075E+06 | 1.305E+05 | 2.581E+07 | 6.116E+07 | 0.000E+00 | 0.000E+00 | 0.000E+00 | 4.979E+06 | 1.202E+05 | 1.429E+07 | 3.551E+07 |
| 565 | 496.304 | 8.835 | 4 | 9.248E+04 | 42.013 | 7.705E+05 | 7.580E+06 | 0.000E+00 | 1.639E+06 | 5.471E+05 | 5.808E+05 | 2.506E+05 | 1.656E+05 | 3.312E+06 | 0.000E+00 | 1.169E+06 | 1.575E+05 | 3.774E+05 | 3.308E+04 |
| 577 | 483.272 | 7.944 | 4 | 6.170E+03 | 2.964 | 2.528E+04 | 3.928E+04 | 4.129E+05 | 2.533E+05 | 3.081E+06 | 1.464E+06 | 2.305E+06 | 9.595E+02 | 1.383E+04 | 1.018E+05 | 1.978E+05 | 9.208E+05 | 3.363E+05 | 8.438E+05 |
| 581 | 511.304 | 8.710 | 4 | 1.693E+05 | 74.038 | 5.469E+04 | 2.280E+03 | 3.860E+05 | 1.681E+05 | 5.903E+06 | 2.033E+06 | 7.593E+06 | 1.504E+04 | 2.280E+03 | 1.509E+05 | 1.468E+05 | 3.438E+05 | 9.598E+05 | 2.780E+06 |
| 583 | 452.277 | 7.806 | 4 | 5.836E+04 | 28.031 | 3.419E+06 | 1.787E+07 | 4.595E+07 | 4.393E+07 | 9.716E+07 | 4.508E+07 | 5.987E+07 | 5.633E+05 | 4.795E+06 | 5.306E+06 | 1.358E+07 | 2.093E+07 | 1.121E+07 | 1.476E+07 |
| 639 | 690.508 | 10.220 | 4 | 3.631E+04 | 26.015 | 2.857E+03 | 9.189E+05 | 2.614E+08 | 1.101E+07 | 7.519E+06 | 2.903E+08 | 7.193E+07 | 2.857E+03 | 2.367E+05 | 1.326E+08 | 9.612E+06 | 4.096E+06 | 1.732E+08 | 3.285E+07 |
| 643 | 712.490 | 10.268 | 4 | 7.051E+04 | 54.046 | 0.000E+00 | 1.310E+05 | 3.698E+07 | 2.686E+06 | 4.026E+06 | 1.975E+08 | 2.386E+08 | 0.000E+00 | 5.367E+04 | 3.124E+07 | 2.431E+06 | 3.468E+06 | 1.106E+08 | 1.468E+08 |
| 669 | 466.293 | 8.072 | 4 | 2.508E+04 | 11.994 | 1.508E+07 | 4.243E+04 | 4.317E+04 | 3.646E+05 | 2.902E+07 | 3.262E+06 | 2.412E+07 | 4.594E+05 | 1.843E+04 | 2.674E+04 | 8.962E+04 | 1.188E+07 | 5.260E+05 | 5.326E+06 |
| 679 | 466.293 | 8.206 | 4 | 2.918E+04 | 14.015 | 2.400E+06 | 8.737E+03 | 2.008E+04 | 4.045E+04 | 9.713E+06 | 5.382E+05 | 3.005E+06 | 3.679E+05 | 1.989E+03 | 9.167E+03 | 6.739E+03 | 3.263E+06 | 2.028E+05 | 5.316E+05 |
| 701 | 440.278 | 7.956 | 4 | 8.334E+04 | 40.030 | 0.000E+00 | 0.000E+00 | 1.611E+05 | 6.646E+05 | 3.571E+06 | 2.161E+06 | 1.741E+06 | 0.000E+00 | 0.000E+00 | 5.698E+04 | 2.379E+05 | 3.216E+06 | 3.996E+05 | 3.992E+05 |
| 713 | 662.477 | 12.666 | 4 | 7.543E+04 | 54.046 | 3.994E+03 | 2.301E+05 | 1.779E+04 | 1.061E+04 | 4.859E+06 | 9.625E+06 | 1.313E+07 | 3.994E+03 | 2.301E+05 | 3.463E+03 | 8.581E+03 | 2.576E+06 | 5.010E+06 | 1.048E+07 |
| 721 | 533.285 | 8.561 | 4 | 2.196E+05 | 96.018 | 6.137E+04 | 1.196E+04 | 2.491E+05 | 2.991E+05 | 4.430E+06 | 1.796E+06 | 6.254E+06 | 5.307E+03 | 8.401E+03 | 3.885E+04 | 1.557E+05 | 4.463E+05 | 8.236E+05 | 1.968E+06 |
| 722 | 440.278 | 7.818 | 4 | 3.085E+04 | 14.014 | 0.000E+00 | 0.000E+00 | 4.457E+03 | 2.750E+05 | 1.214E+06 | 9.425E+05 | 5.603E+05 | 0.000E+00 | 0.000E+00 | 4.457E+03 | 1.037E+05 | 1.112E+06 | 1.863E+05 | 1.145E+05 |
| 724 | 684.458 | 12.702 | 4 | 1.071E+05 | 82.077 | 0.000E+00 | 8.187E+04 | 0.000E+00 | 0.000E+00 | 2.199E+06 | 3.755E+06 | 5.237E+06 | 0.000E+00 | 8.187E+04 | 0.000E+00 | 0.000E+00 | 1.203E+06 | 2.110E+06 | 4.367E+06 |
| 727 | 465.262 | 7.796 | 4 | 1.097E+05 | 46.006 | 4.999E+04 | 1.303E+05 | 1.902E+05 | 3.543E+05 | 3.575E+06 | 2.630E+06 | 6.646E+06 | 2.444E+04 | 3.337E+04 | 8.716E+04 | 1.741E+05 | 1.022E+06 | 6.252E+05 | 2.393E+06 |
| 728 | 505.254 | 7.795 | 4 | 7.711E+04 | 42.218 | 2.434E+04 | 2.206E+05 | 2.294E+05 | 3.816E+05 | 3.704E+06 | 2.804E+06 | 6.944E+06 | 7.982E+03 | 7.262E+04 | 1.057E+05 | 1.977E+05 | 8.945E+05 | 7.031E+05 | 2.374E+06 |
| 729 | 507.269 | 8.510 | 4 | 1.709E+05 | 74.037 | 8.403E+04 | 4.212E+04 | 2.588E+05 | 2.708E+05 | 4.148E+06 | 2.523E+06 | 6.576E+06 | 1.081E+04 | 1.385E+04 | 9.288E+04 | 1.826E+05 | 1.063E+06 | 7.904E+05 | 2.163E+06 |
| 731 | 493.293 | 8.564 | 4 | 1.281E+05 | 56.027 | 5.062E+04 | 9.606E+03 | 2.197E+05 | 2.032E+05 | 3.150E+06 | 1.285E+06 | 4.801E+06 | 2.223E+04 | 5.398E+03 | 8.029E+04 | 1.098E+05 | 3.797E+05 | 6.380E+05 | 1.561E+06 |
| 736 | 676.492 | 12.707 | 4 | 5.586E+04 | 40.028 | 0.000E+00 | 0.000E+00 | 0.000E+00 | 1.661E+05 | 5.431E+05 | 3.609E+06 | 1.386E+06 | 0.000E+00 | 0.000E+00 | 0.000E+00 | 1.661E+05 | 4.923E+05 | 2.316E+06 | 1.132E+06 |
| 745 | 493.293 | 8.710 | 4 | 1.281E+05 | 56.027 | 6.295E+03 | 0.000E+00 | 6.892E+04 | 4.369E+04 | 2.055E+06 | 6.793E+05 | 2.681E+06 | 4.210E+03 | 0.000E+00 | 3.078E+04 | 3.899E+04 | 1.562E+05 | 3.121E+05 | 1.050E+06 |
| 749 | 500.300 | 7.793 | 4 | 1.087E+05 | 49.030 | 1.130E+04 | 5.350E+04 | 5.548E+04 | 1.397E+05 | 1.615E+06 | 1.112E+06 | 2.862E+06 | 7.662E+03 | 2.398E+04 | 2.552E+04 | 6.323E+04 | 5.197E+05 | 3.583E+05 | 9.675E+05 |
| 756 | 533.285 | 8.715 | 4 | 1.612E+05 | 74.037 | 4.364E+03 | 0.000E+00 | 3.546E+04 | 3.667E+04 | 1.377E+06 | 4.500E+05 | 1.756E+06 | 4.364E+03 | 0.000E+00 | 1.913E+04 | 3.439E+04 | 1.164E+05 | 2.100E+05 | 6.389E+05 |
| 775 | 704.523 | 12.703 | 4 | 1.950E+04 | 14.015 | 0.000E+00 | 2.308E+06 | 5.683E+04 | 0.000E+00 | 9.462E+04 | 2.230E+07 | 6.743E+06 | 0.000E+00 | 1.156E+06 | 5.683E+04 | 0.000E+00 | 5.177E+04 | 3.199E+06 | 3.009E+06 |
| 779 | 704.523 | 10.367 | 4 | 1.951E+04 | 14.015 | 0.000E+00 | 2.149E+07 | 1.800E+05 | 1.055E+04 | 4.920E+04 | 6.089E+07 | 3.294E+07 | 0.000E+00 | 1.111E+07 | 1.607E+05 | 5.274E+03 | 4.124E+04 | 5.607E+07 | 2.285E+07 |
| 781 | 726.506 | 12.701 | 4 | 5.222E+04 | 40.030 | 0.000E+00 | 8.729E+05 | 2.533E+04 | 0.000E+00 | 8.842E+03 | 9.034E+06 | 2.734E+06 | 0.000E+00 | 4.371E+05 | 2.533E+04 | 0.000E+00 | 8.842E+03 | 1.564E+06 | 1.407E+06 |
| 813 | 494.288 | 8.224 | 4 | 2.911E+04 | 13.980 | 7.435E+02 | 5.677E+05 | 3.659E+05 | 9.477E+05 | 1.157E+05 | 3.664E+04 | 1.748E+04 | 7.435E+02 | 2.219E+05 | 7.801E+04 | 6.765E+05 | 1.020E+05 | 1.296E+04 | 9.147E+03 |
| 825 | 664.490 | 9.773 | 4 | 7.262E+04 | 52.033 | 2.518E+03 | 2.466E+04 | 5.940E+04 | 0.000E+00 | 4.367E+04 | 3.326E+07 | 1.213E+08 | 2.518E+03 | 9.651E+03 | 2.266E+04 | 0.000E+00 | 4.367E+04 | 2.928E+07 | 8.012E+07 |
| 837 | 766.559 | 12.703 | 4 | N/A | N/A | 0.000E+00 | 0.000E+00 | 0.000E+00 | 2.424E+04 | 9.091E+04 | 6.759E+05 | 3.177E+06 | 0.000E+00 | 0.000E+00 | 0.000E+00 | 1.251E+04 | 3.517E+04 | 3.325E+05 | 2.698E+06 |
| 841 | 676.491 | 10.424 | 4 | 5.587E+04 | 40.032 | 0.000E+00 | 0.000E+00 | 0.000E+00 | 9.383E+03 | 1.980E+04 | 4.262E+07 | 1.285E+07 | 0.000E+00 | 0.000E+00 | 0.000E+00 | 9.383E+03 | 1.980E+04 | 3.061E+07 | 9.373E+06 |
| 844 | 698.475 | 10.420 | 4 | 8.879E+04 | 68.061 | 0.000E+00 | 0.000E+00 | 0.000E+00 | 0.000E+00 | 6.130E+03 | 9.856E+06 | 2.237E+06 | 0.000E+00 | 0.000E+00 | 0.000E+00 | 0.000E+00 | 6.130E+03 | 7.126E+06 | 2.091E+06 |
| 853 | 1380.008 | 10.186 | 4 | N/A | N/A | 0.000E+00 | 0.000E+00 | 1.022E+05 | 0.000E+00 | 0.000E+00 | 5.648E+06 | 1.091E+07 | 0.000E+00 | 0.000E+00 | 7.099E+04 | 0.000E+00 | 0.000E+00 | 5.355E+06 | 1.059E+07 |
| 855 | 1380.006 | 9.800 | 4 | N/A | N/A | 0.000E+00 | 0.000E+00 | 0.000E+00 | 0.000E+00 | 0.000E+00 | 1.150E+06 | 2.147E+06 | 0.000E+00 | 0.000E+00 | 0.000E+00 | 0.000E+00 | 0.000E+00 | 1.071E+06 | 2.057E+06 |
| 27 | 316.212 | 3.277 | 3 | N/A | N/A | 1.738E+08 | 3.185E+08 | 1.582E+08 | 2.112E+08 | 2.306E+08 | 2.582E+08 | 1.876E+08 | 2.999E+07 | 3.464E+07 | 1.112E+07 | 7.802E+06 | 2.602E+07 | 1.777E+07 | 2.767E+07 |
| 37 | 430.244 | 4.459 | 3 | N/A | N/A | 2.627E+07 | 3.812E+07 | 2.613E+07 | 3.187E+07 | 2.876E+07 | 3.574E+07 | 3.281E+07 | 1.301E+06 | 2.765E+06 | 1.397E+06 | 8.336E+05 | 2.266E+06 | 2.254E+06 | 5.201E+06 |
| 55 | 300.200 | 3.938 | 3 | N/A | N/A | 4.197E+07 | 9.306E+07 | 3.636E+07 | 5.680E+07 | 6.641E+07 | 7.203E+07 | 4.674E+07 | 5.164E+06 | 9.406E+06 | 2.512E+05 | 1.016E+06 | 5.833E+06 | 1.591E+06 | 4.500E+06 |
| 69 | 416.228 | 3.930 | 3 | N/A | N/A | 2.528E+07 | 3.590E+07 | 1.945E+07 | 2.899E+07 | 3.246E+07 | 3.556E+07 | 2.840E+07 | 2.428E+06 | 2.382E+06 | 3.742E+05 | 1.044E+06 | 2.239E+06 | 3.705E+06 | 4.900E+06 |
| 122 | 243.183 | 3.050 | 3 | N/A | N/A | 1.934E+07 | 2.515E+07 | 1.161E+07 | 2.349E+07 | 1.778E+07 | 2.670E+07 | 2.404E+07 | 3.609E+06 | 3.168E+06 | 9.633E+05 | 4.172E+05 | 1.778E+06 | 9.930E+05 | 5.493E+06 |
| 124 | 334.178 | 3.942 | 3 | N/A | N/A | 1.336E+07 | 2.119E+07 | 1.723E+07 | 2.133E+07 | 1.493E+07 | 2.381E+07 | 1.785E+07 | 3.159E+06 | 5.902E+06 | 2.119E+06 | 1.612E+06 | 3.369E+06 | 1.520E+06 | 2.350E+06 |
| 128 | 357.214 | 4.538 | 3 | N/A | N/A | 8.203E+06 | 1.465E+07 | 9.366E+06 | 1.180E+07 | 1.235E+07 | 1.354E+07 | 1.122E+07 | 5.315E+05 | 8.923E+05 | 5.488E+05 | 2.260E+05 | 1.560E+06 | 1.076E+06 | 1.576E+06 |
| 149 | 350.208 | 4.208 | 3 | N/A | N/A | 1.246E+07 | 2.101E+07 | 1.243E+07 | 1.774E+07 | 1.792E+07 | 2.183E+07 | 1.602E+07 | 1.671E+06 | 2.475E+06 | 4.696E+05 | 1.817E+06 | 8.113E+05 | 1.475E+06 | 1.016E+06 |
| 167 | 599.391 | 3.938 | 3 | N/A | N/A | 5.862E+06 | 2.714E+07 | 8.790E+06 | 1.330E+07 | 1.570E+07 | 1.884E+07 | 1.299E+07 | 1.016E+06 | 5.708E+06 | 3.149E+06 | 5.424E+06 | 4.899E+06 | 5.670E+06 | 5.495E+06 |
| 190 | 294.861 | 4.052 | 3 | N/A | N/A | 1.534E+07 | 3.171E+07 | 1.848E+07 | 3.010E+07 | 3.126E+07 | 4.078E+07 | 1.593E+07 | 3.357E+06 | 1.038E+07 | 8.479E+05 | 1.981E+06 | 5.478E+05 | 1.157E+06 | 5.129E+06 |
| 224 | 302.196 | 0.594 | 3 | N/A | N/A | 3.804E+07 | 5.043E+07 | 1.968E+07 | 4.189E+07 | 4.990E+07 | 6.193E+07 | 4.118E+07 | 6.450E+06 | 5.623E+06 | 1.861E+06 | 4.788E+06 | 3.186E+06 | 2.821E+06 | 3.844E+06 |
| 246 | 309.181 | 4.412 | 3 | N/A | N/A | 5.862E+06 | 7.331E+06 | 5.709E+06 | 7.531E+06 | 5.960E+06 | 7.954E+06 | 7.416E+06 | 8.454E+05 | 1.153E+06 | 3.191E+05 | 1.496E+05 | 6.077E+05 | 3.319E+05 | 1.311E+06 |
| 268 | 328.200 | 4.325 | 3 | N/A | N/A | 4.992E+06 | 9.387E+06 | 5.206E+06 | 8.493E+06 | 8.100E+06 | 1.096E+07 | 7.780E+06 | 6.129E+05 | 1.567E+06 | 2.975E+05 | 7.936E+05 | 2.054E+05 | 8.552E+05 | 4.591E+05 |
| 290 | 272.185 | 3.424 | 3 | N/A | N/A | 1.223E+07 | 3.465E+07 | 1.023E+07 | 1.695E+07 | 1.522E+07 | 2.294E+07 | 1.257E+07 | 3.438E+06 | 5.409E+06 | 1.893E+06 | 4.482E+06 | 4.274E+06 | 2.892E+06 | 2.599E+06 |
| 293 | 293.191 | 3.577 | 3 | N/A | N/A | 6.414E+06 | 1.146E+07 | 2.419E+06 | 6.989E+06 | 1.055E+07 | 1.266E+07 | 6.761E+06 | 2.382E+06 | 1.654E+06 | 1.219E+05 | 1.842E+06 | 1.340E+06 | 1.819E+06 | 8.654E+05 |
| 306 | 300.180 | 5.792 | 3 | N/A | N/A | 1.695E+06 | 3.622E+06 | 2.146E+06 | 2.353E+06 | 2.383E+06 | 2.928E+06 | 1.868E+06 | 2.244E+05 | 1.936E+05 | 2.187E+05 | 1.117E+05 | 3.594E+05 | 2.410E+05 | 3.814E+05 |
| 322 | 713.423 | 4.538 | 3 | N/A | N/A | 2.818E+06 | 4.276E+06 | 2.806E+06 | 4.006E+06 | 4.251E+06 | 4.612E+06 | 3.884E+06 | 2.025E+05 | 7.583E+05 | 4.797E+05 | 1.836E+05 | 5.241E+05 | 3.759E+05 | 5.329E+05 |
| 328 | 252.166 | 3.641 | 3 | N/A | N/A | 5.367E+06 | 1.419E+07 | 3.674E+06 | 9.648E+06 | 1.047E+07 | 1.177E+07 | 5.602E+06 | 1.020E+06 | 2.423E+06 | 4.704E+05 | 8.429E+05 | 1.194E+06 | 3.796E+05 | 1.442E+06 |
| 329 | 332.872 | 4.600 | 3 | N/A | N/A | 5.175E+06 | 1.089E+07 | 5.239E+06 | 8.485E+06 | 8.658E+06 | 1.183E+07 | 7.372E+06 | 6.857E+05 | 1.614E+06 | 2.899E+05 | 7.737E+05 | 3.196E+05 | 1.250E+06 | 7.522E+05 |
| 343 | 471.307 | 6.113 | 3 | N/A | N/A | 1.012E+06 | 1.556E+06 | 1.213E+06 | 1.224E+06 | 1.618E+06 | 1.485E+06 | 1.203E+06 | 9.909E+04 | 2.324E+05 | 6.410E+04 | 8.446E+04 | 2.761E+05 | 3.644E+04 | 7.060E+04 |
| 361 | 441.788 | 4.079 | 3 | N/A | N/A | 8.352E+05 | 7.964E+06 | 3.386E+06 | 4.736E+06 | 2.938E+06 | 4.622E+06 | 4.245E+06 | 2.761E+05 | 2.354E+06 | 2.164E+06 | 3.092E+06 | 7.809E+05 | 8.174E+05 | 2.041E+06 |
| 386 | 241.204 | 5.244 | 3 | N/A | N/A | 2.931E+06 | 9.403E+06 | 1.437E+06 | 5.288E+07 | 2.856E+06 | 4.786E+06 | 1.616E+07 | 6.866E+05 | 2.137E+06 | 4.638E+04 | 4.990E+07 | 4.308E+05 | 2.975E+05 | 1.451E+07 |
| 396 | 469.328 | 4.897 | 3 | N/A | N/A | 1.700E+06 | 2.813E+06 | 1.255E+06 | 1.945E+06 | 2.168E+06 | 2.281E+06 | 1.778E+06 | 3.132E+05 | 1.989E+05 | 7.173E+04 | 7.174E+04 | 1.350E+05 | 9.121E+04 | 2.557E+05 |
| 401 | 256.850 | 3.715 | 3 | N/A | N/A | 1.106E+07 | 1.554E+07 | 6.852E+06 | 1.705E+07 | 1.393E+07 | 2.121E+07 | 1.541E+07 | 1.258E+06 | 2.977E+06 | 4.877E+05 | 1.854E+06 | 1.329E+06 | 1.263E+06 | 1.005E+06 |
| 404 | 699.407 | 4.204 | 3 | N/A | N/A | 2.878E+06 | 4.958E+06 | 2.537E+06 | 4.162E+06 | 4.388E+06 | 5.650E+06 | 3.887E+06 | 6.598E+05 | 4.794E+05 | 1.810E+05 | 1.762E+05 | 1.945E+06 | 6.568E+05 | 2.967E+05 |
| 413 | 292.192 | 4.464 | 3 | N/A | N/A | 1.514E+06 | 2.556E+07 | 7.331E+06 | 2.047E+07 | 1.704E+07 | 2.718E+07 | 1.213E+07 | 8.324E+05 | 4.104E+06 | 3.218E+06 | 2.904E+06 | 1.875E+05 | 3.241E+06 | 5.935E+06 |
| 439 | 585.375 | 3.578 | 3 | N/A | N/A | 2.419E+06 | 3.371E+06 | 1.251E+06 | 2.851E+06 | 3.952E+06 | 4.493E+06 | 3.626E+06 | 6.457E+05 | 4.227E+05 | 1.690E+05 | 4.296E+05 | 2.161E+05 | 1.612E+05 | 3.593E+05 |
| 443 | 384.773 | 3.708 | 3 | N/A | N/A | 6.809E+06 | 9.034E+06 | 4.949E+06 | 1.063E+07 | 9.122E+06 | 6.509E+06 | 3.920E+06 | 1.935E+05 | 1.563E+06 | 2.611E+05 | 1.007E+06 | 1.250E+06 | 1.464E+06 | 2.490E+06 |
| 463 | 293.191 | 3.406 | 3 | N/A | N/A | 1.667E+06 | 3.540E+06 | 1.587E+06 | 3.175E+06 | 2.578E+06 | 3.613E+06 | 3.696E+06 | 2.617E+05 | 5.261E+05 | 9.455E+04 | 5.557E+05 | 4.343E+05 | 7.377E+05 | 3.227E+05 |
| 499 | 599.391 | 3.791 | 3 | N/A | N/A | 1.714E+07 | 2.516E+07 | 1.270E+07 | 1.969E+07 | 2.393E+07 | 2.524E+07 | 1.583E+07 | 2.242E+06 | 1.045E+07 | 2.692E+06 | 4.927E+06 | 6.330E+06 | 7.598E+06 | 3.279E+06 |
| 526 | 485.359 | 3.054 | 3 | N/A | N/A | 3.439E+06 | 5.376E+06 | 1.518E+06 | 5.173E+06 | 4.237E+06 | 7.023E+06 | 5.583E+06 | 8.017E+05 | 9.230E+05 | 4.173E+05 | 2.664E+05 | 5.240E+05 | 3.707E+05 | 9.927E+05 |
| 527 | 441.788 | 4.168 | 3 | N/A | N/A | 7.023E+06 | 1.249E+07 | 4.660E+06 | 8.073E+06 | 1.136E+07 | 1.271E+07 | 6.341E+06 | 4.069E+05 | 3.099E+06 | 1.805E+06 | 2.470E+06 | 2.432E+05 | 1.824E+05 | 1.534E+06 |
| 530 | 290.193 | 3.898 | 3 | N/A | N/A | 4.917E+06 | 7.530E+06 | 2.888E+06 | 7.732E+06 | 8.828E+06 | 1.099E+07 | 7.945E+06 | 5.852E+05 | 1.188E+06 | 9.619E+05 | 7.215E+05 | 9.516E+04 | 6.863E+05 | 1.005E+06 |

| Feature identifiers | | | | | | Average XIC | | | | | | | Standard Err of Average XIC | | | | | | |
|---|---|---|---|---|---|---|---|---|---|---|---|---|---|---|---|---|---|---|---|
| ClusterIdx | m/z | RT [min] | Subnetwork | PPM | MassDiff | TUB00 | 4TUB34 | 4TUB81 | 4TUB93 | 5TUB23 | 5TUB48 | 5TUB83 | TUB00 | 4TUB34 | 4TUB81 | 4TUB93 | 5TUB23 | 5TUB48 | 5TUB83 |
| 553 | 332.207 | 4.839 | 3 | N/A | N/A | 1.166E+06 | 8.212E+05 | 9.252E+05 | 1.613E+06 | 9.635E+05 | 1.666E+06 | 2.015E+06 | 9.531E+04 | 2.804E+05 | 2.596E+05 | 3.004E+04 | 1.054E+05 | 2.238E+05 | 3.104E+05 |
| 568 | 379.210 | 4.343 | 3 | N/A | N/A | 1.155E+06 | 3.634E+06 | 6.307E+05 | 3.044E+06 | 2.930E+06 | 4.095E+06 | 2.964E+06 | 5.872E+05 | 7.783E+05 | 7.339E+04 | 5.189E+05 | 4.025E+05 | 5.269E+05 | 2.943E+05 |
| 623 | 343.200 | 3.890 | 3 | N/A | N/A | 1.375E+06 | 2.014E+06 | 4.682E+05 | 1.791E+06 | 2.186E+06 | 2.584E+06 | 2.067E+06 | 4.097E+05 | 3.599E+05 | 3.711E+04 | 4.241E+05 | 1.578E+05 | 1.454E+05 | 3.935E+05 |
| 624 | 263.684 | 4.125 | 3 | N/A | N/A | 6.373E+05 | 3.866E+06 | 3.089E+05 | 7.976E+06 | 9.421E+05 | 1.046E+07 | 7.278E+06 | 1.150E+05 | 3.107E+06 | 5.655E+04 | 3.976E+06 | 1.587E+05 | 4.134E+06 | 3.450E+06 |
| 627 | 230.153 | 3.714 | 3 | N/A | N/A | 6.655E+05 | 3.998E+06 | 3.631E+05 | 1.807E+06 | 1.088E+06 | 3.092E+06 | 9.979E+05 | 1.128E+05 | 9.682E+05 | 8.292E+04 | 2.698E+05 | 5.091E+05 | 1.462E+05 | 4.154E+05 |
| 629 | 347.535 | 4.414 | 3 | N/A | N/A | 5.695E+05 | 1.863E+06 | 2.956E+05 | 1.769E+06 | 1.677E+06 | 2.342E+06 | 1.882E+06 | 2.543E+05 | 5.628E+05 | 1.175E+05 | 4.909E+05 | 1.956E+05 | 7.236E+05 | 1.325E+05 |
| 747 | 540.426 | 9.375 | 3 | N/A | N/A | 7.940E+05 | 2.151E+06 | 7.420E+05 | 1.134E+06 | 1.752E+06 | 2.172E+06 | 2.218E+06 | 1.554E+05 | 6.662E+05 | 1.147E+04 | 4.328E+05 | 2.287E+05 | 3.109E+05 | 1.348E+06 |
| 750 | 568.458 | 9.844 | 3 | N/A | N/A | 7.564E+04 | 6.689E+05 | 9.585E+04 | 2.932E+05 | 1.127E+06 | 1.512E+06 | 1.997E+06 | 3.117E+03 | 2.748E+05 | 2.200E+04 | 1.584E+05 | 7.883E+04 | 6.505E+05 | 1.436E+06 |
| 758 | 263.684 | 4.034 | 3 | N/A | N/A | 5.530E+05 | 8.417E+06 | 4.158E+05 | 1.975E+06 | 4.297E+06 | 6.202E+06 | 7.397E+05 | 1.971E+05 | 3.932E+06 | 1.443E+05 | 6.633E+05 | 3.549E+06 | 5.347E+06 | 1.993E+05 |
| 782 | 220.150 | 0.536 | 3 | N/A | N/A | 3.900E+04 | 6.938E+05 | 1.927E+05 | 7.637E+03 | 3.815E+04 | 1.295E+06 | 1.612E+06 | 3.010E+04 | 5.166E+05 | 1.277E+05 | 7.637E+03 | 2.173E+04 | 7.110E+05 | 1.017E+06 |
| 784 | 498.804 | 4.596 | 3 | N/A | N/A | 6.851E+05 | 2.217E+06 | 7.216E+05 | 1.972E+06 | 8.160E+05 | 2.623E+06 | 1.231E+06 | 2.040E+05 | 5.979E+05 | 8.629E+04 | 4.385E+05 | 2.779E+05 | 2.895E+05 | 4.225E+05 |
| 810 | 294.861 | 4.145 | 3 | N/A | N/A | 1.022E+07 | 2.392E+07 | 4.159E+06 | 9.764E+06 | 7.310E+06 | 1.682E+07 | 1.967E+07 | 6.085E+06 | 8.974E+06 | 5.539E+05 | 3.898E+06 | 1.389E+06 | 2.451E+06 | 5.909E+06 |
| 817 | 384.772 | 3.711 | 3 | N/A | N/A | 1.923E+06 | 4.616E+06 | 2.094E+06 | 5.103E+06 | 4.727E+06 | 9.515E+06 | 5.404E+06 | 1.796E+05 | 1.277E+06 | 4.662E+05 | 9.757E+05 | 1.056E+06 | 3.276E+06 | 1.371E+06 |
| 819 | 292.192 | 4.644 | 3 | N/A | N/A | 5.176E+06 | 1.081E+06 | 4.987E+05 | 7.071E+05 | 7.633E+05 | 2.059E+06 | 2.925E+06 | 4.363E+06 | 2.634E+05 | 1.791E+05 | 1.482E+05 | 1.180E+05 | 4.601E+05 | 2.306E+06 |
| 93 | 256.264 | 9.789 | 2 | N/A | N/A | 7.134E+06 | 7.143E+06 | 7.736E+06 | 7.679E+06 | 7.452E+06 | 7.832E+06 | 7.785E+06 | 1.636E+05 | 2.225E+05 | 1.553E+05 | 2.987E+05 | 2.121E+05 | 4.849E+05 | 3.898E+05 |
| 142 | 284.295 | 10.325 | 2 | 1.094E+05 | 28.032 | 4.586E+06 | 4.499E+06 | 5.576E+06 | 4.549E+06 | 4.737E+06 | 4.831E+06 | 3.931E+06 | 2.428E+05 | 2.950E+05 | 3.208E+05 | 3.572E+05 | 2.147E+05 | 4.356E+05 | 1.106E+05 |
| 175 | 308.279 | 8.430 | 2 | N/A | N/A | 3.064E+06 | 3.932E+06 | 4.505E+06 | 4.856E+06 | 3.855E+06 | 4.797E+06 | 5.236E+06 | 3.002E+05 | 4.329E+05 | 1.839E+05 | 3.056E+05 | 3.024E+05 | 2.378E+05 | 1.674E+06 |
| 218 | 482.408 | 9.562 | 2 | 1.617E+05 | 67.155 | 2.895E+06 | 3.829E+06 | 3.124E+06 | 3.484E+06 | 5.351E+06 | 4.212E+06 | 3.717E+06 | 3.128E+04 | 2.090E+05 | 7.341E+04 | 4.406E+05 | 1.836E+06 | 3.180E+05 | 3.394E+05 |
| 226 | 526.434 | 9.538 | 2 | 1.462E+05 | 67.154 | 2.961E+06 | 4.332E+06 | 3.291E+06 | 3.329E+06 | 4.302E+06 | 5.332E+06 | 4.102E+06 | 1.293E+05 | 3.518E+05 | 8.780E+04 | 5.854E+05 | 6.130E+05 | 1.174E+06 | 4.442E+05 |
| 228 | 336.311 | 9.373 | 2 | N/A | N/A | 3.439E+06 | 3.578E+06 | 3.743E+06 | 9.585E+06 | 4.449E+06 | 3.975E+06 | 3.805E+06 | 2.583E+04 | 7.989E+04 | 2.671E+05 | 2.880E+06 | 3.421E+05 | 1.691E+05 | 1.946E+05 |
| 229 | 291.253 | 8.431 | 2 | 4.927E+05 | 96.130 | 2.157E+06 | 2.845E+06 | 3.283E+06 | 3.604E+06 | 2.829E+06 | 3.608E+06 | 3.923E+06 | 2.804E+05 | 3.433E+05 | 8.759E+04 | 1.865E+05 | 2.503E+05 | 2.292E+05 | 1.348E+06 |
| 241 | 424.364 | 9.342 | 2 | N/A | N/A | 2.403E+06 | 3.287E+06 | 3.739E+06 | 1.401E+07 | 4.837E+06 | 3.991E+06 | 3.390E+06 | 3.356E+04 | 3.537E+05 | 8.417E+05 | 4.722E+06 | 1.013E+06 | 2.240E+05 | 1.863E+05 |
| 242 | 380.338 | 9.364 | 2 | N/A | N/A | 3.231E+06 | 3.833E+06 | 4.315E+06 | 1.462E+07 | 5.471E+06 | 4.580E+06 | 4.075E+06 | 5.111E+04 | 1.700E+05 | 7.391E+05 | 4.757E+06 | 8.687E+05 | 2.406E+05 | 1.684E+05 |
| 247 | 352.306 | 8.443 | 2 | N/A | N/A | 1.986E+06 | 2.682E+06 | 2.902E+06 | 3.566E+06 | 2.808E+06 | 3.476E+06 | 3.664E+06 | 3.463E+05 | 4.985E+05 | 1.352E+05 | 3.037E+05 | 3.480E+05 | 3.136E+05 | 1.159E+06 |
| 269 | 438.382 | 9.584 | 2 | 5.570E+04 | 23.128 | 4.913E+06 | 5.396E+06 | 5.506E+06 | 6.055E+06 | 5.843E+06 | 7.540E+06 | 6.127E+06 | 1.817E+05 | 1.260E+05 | 2.037E+05 | 2.119E+05 | 1.918E+05 | 1.015E+06 | 4.003E+05 |
| 270 | 408.372 | 10.006 | 2 | N/A | N/A | 3.284E+06 | 4.130E+06 | 3.813E+06 | 4.533E+06 | 4.151E+06 | 4.732E+06 | 3.838E+06 | 9.630E+04 | 1.628E+05 | 1.242E+05 | 3.503E+05 | 8.992E+04 | 3.103E+05 | 1.481E+05 |
| 276 | 570.460 | 9.514 | 2 | 1.334E+05 | 67.149 | 2.409E+06 | 4.313E+06 | 3.042E+06 | 2.716E+06 | 3.732E+06 | 4.663E+06 | 4.191E+06 | 8.591E+04 | 7.093E+05 | 5.507E+05 | 5.103E+05 | 8.111E+05 | 1.249E+06 | 5.032E+05 |
| 278 | 512.417 | 9.299 | 2 | 2.340E+05 | 97.163 | 1.556E+06 | 2.500E+06 | 3.009E+06 | 1.176E+07 | 3.869E+06 | 2.828E+06 | 2.118E+06 | 2.371E+04 | 5.328E+05 | 8.468E+05 | 3.956E+06 | 8.783E+05 | 3.413E+05 | 1.939E+05 |
| 280 | 468.390 | 9.320 | 2 | N/A | N/A | 1.821E+06 | 2.810E+06 | 3.321E+06 | 1.342E+07 | 4.557E+06 | 3.351E+06 | 2.554E+06 | 2.879E+04 | 4.790E+05 | 8.804E+05 | 4.418E+06 | 1.150E+06 | 3.265E+05 | 1.399E+05 |
| 291 | 366.322 | 8.959 | 2 | N/A | N/A | 3.592E+06 | 3.602E+06 | 3.420E+06 | 3.457E+06 | 3.395E+06 | 3.459E+06 | 3.344E+06 | 5.593E+04 | 6.947E+04 | 4.067E+04 | 1.025E+05 | 5.205E+04 | 9.667E+03 | 9.081E+04 |
| 294 | 452.398 | 9.985 | 2 | N/A | N/A | 2.220E+06 | 3.024E+06 | 2.412E+06 | 3.984E+06 | 3.156E+06 | 3.484E+06 | 2.453E+06 | 3.567E+04 | 2.459E+05 | 1.655E+05 | 1.150E+06 | 5.153E+05 | 7.312E+05 | 1.795E+05 |
| 297 | 556.443 | 9.273 | 2 | 2.116E+05 | 97.163 | 1.258E+06 | 2.139E+06 | 2.832E+06 | 9.213E+06 | 3.188E+06 | 2.439E+06 | 1.929E+06 | 2.718E+04 | 4.434E+05 | 9.059E+05 | 2.986E+06 | 6.712E+05 | 2.660E+05 | 2.149E+05 |
| 302 | 247.227 | 8.431 | 2 | N/A | N/A | 1.164E+06 | 1.786E+06 | 1.992E+06 | 1.933E+06 | 1.718E+06 | 2.203E+06 | 2.392E+06 | 4.268E+04 | 2.833E+05 | 9.352E+04 | 2.239E+05 | 1.606E+05 | 1.142E+05 | 8.595E+05 |
| 303 | 392.338 | 10.385 | 2 | N/A | N/A | 2.109E+06 | 2.144E+06 | 2.169E+06 | 1.966E+06 | 1.990E+06 | 2.193E+06 | 2.270E+06 | 6.667E+04 | 1.692E+04 | 1.491E+04 | 4.512E+04 | 1.090E+04 | 1.016E+05 | 3.788E+04 |
| 310 | 496.424 | 9.963 | 2 | N/A | N/A | 1.943E+06 | 4.848E+06 | 2.698E+06 | 4.364E+06 | 3.115E+06 | 4.384E+06 | 2.217E+06 | 3.205E+04 | 5.033E+05 | 5.098E+05 | 1.498E+06 | 5.103E+05 | 1.027E+06 | 5.414E+04 |
| 336 | 364.344 | 10.020 | 2 | N/A | N/A | 2.193E+06 | 2.929E+06 | 2.392E+06 | 2.836E+06 | 2.965E+06 | 3.756E+06 | 3.726E+06 | 4.353E+04 | 2.404E+05 | 6.145E+04 | 3.658E+05 | 3.692E+05 | 5.379E+05 | 9.126E+04 |
| 337 | 540.449 | 9.937 | 2 | 1.767E+05 | 81.169 | 1.528E+06 | 3.970E+06 | 1.874E+06 | 4.031E+06 | 2.086E+06 | 3.945E+06 | 2.446E+06 | 8.673E+03 | 1.169E+06 | 9.078E+04 | 1.440E+06 | 2.765E+05 | 1.060E+06 | 6.732E+05 |
| 344 | 476.301 | 4.018 | 2 | 5.366E+04 | 27.006 | 2.855E+06 | 3.152E+06 | 2.672E+06 | 3.014E+06 | 3.021E+06 | 2.804E+06 | 2.601E+06 | 9.745E+04 | 3.774E+04 | 4.688E+04 | 1.322E+05 | 2.184E+05 | 8.415E+04 | 6.645E+04 |
| 349 | 440.359 | 8.424 | 2 | N/A | N/A | 1.361E+06 | 2.024E+06 | 1.612E+06 | 2.103E+06 | 1.854E+06 | 2.072E+06 | 2.038E+06 | 2.455E+05 | 2.868E+05 | 3.803E+04 | 1.621E+05 | 3.193E+05 | 1.268E+05 | 5.564E+05 |
| 359 | 600.469 | 9.251 | 2 | 1.931E+05 | 97.163 | 9.333E+05 | 1.591E+06 | 2.385E+06 | 6.713E+06 | 2.407E+06 | 1.847E+06 | 1.454E+06 | 4.446E+04 | 2.942E+05 | 7.975E+05 | 2.162E+06 | 4.654E+05 | 2.079E+05 | 1.403E+05 |
| 362 | 322.295 | 8.962 | 2 | N/A | N/A | 2.838E+06 | 2.648E+06 | 2.648E+06 | 2.673E+06 | 2.483E+06 | 2.511E+06 | 2.528E+06 | 8.007E+04 | 2.882E+04 | 3.009E+04 | 4.557E+04 | 4.113E+04 | 5.069E+04 | 1.990E+04 |
| 364 | 614.485 | 9.493 | 2 | 4.209E+04 | 27.000 | 1.555E+06 | 3.647E+06 | 1.683E+06 | 1.891E+06 | 2.983E+06 | 3.216E+06 | 2.747E+06 | 4.517E+04 | 9.199E+05 | 1.280E+05 | 4.550E+05 | 8.413E+05 | 6.873E+05 | 4.152E+05 |
| 366 | 396.333 | 8.432 | 2 | N/A | N/A | 1.445E+06 | 2.030E+06 | 1.752E+06 | 2.347E+06 | 1.823E+06 | 2.225E+06 | 2.370E+06 | 2.130E+05 | 2.698E+05 | 2.433E+04 | 2.238E+05 | 2.857E+05 | 1.674E+05 | 7.098E+05 |
| 367 | 422.389 | 10.267 | 2 | N/A | N/A | 1.794E+06 | 2.141E+06 | 1.902E+06 | 1.821E+06 | 2.033E+06 | 2.249E+06 | 1.744E+06 | 2.964E+04 | 1.241E+05 | 1.218E+04 | 9.288E+04 | 1.489E+05 | 6.775E+04 | 1.256E+05 |
| 369 | 436.364 | 10.348 | 2 | N/A | N/A | 2.259E+06 | 2.247E+06 | 2.302E+06 | 2.229E+06 | 2.210E+06 | 2.321E+06 | 2.299E+06 | 1.349E+04 | 9.753E+03 | 6.776E+04 | 4.537E+04 | 4.512E+04 | 9.578E+03 | 5.997E+04 |
| 377 | 422.347 | 10.141 | 2 | N/A | N/A | 3.927E+06 | 4.085E+06 | 4.130E+06 | 4.051E+06 | 4.125E+06 | 3.963E+06 | 3.790E+06 | 3.775E+04 | 2.104E+04 | 8.086E+04 | 1.051E+05 | 8.752E+04 | 2.408E+05 | 6.761E+05 |
| 382 | 466.373 | 10.105 | 2 | N/A | N/A | 2.940E+06 | 3.094E+06 | 3.221E+06 | 3.330E+06 | 3.406E+06 | 3.037E+06 | 1.949E+06 | 5.098E+05 | 6.106E+05 | 6.049E+05 | 6.445E+05 | 7.342E+05 | 5.205E+05 | 3.952E+04 |
| 383 | 319.284 | 9.373 | 2 | 6.009E+04 | 18.099 | 1.474E+06 | 1.644E+06 | 1.508E+06 | 4.692E+06 | 1.983E+06 | 1.730E+06 | 1.633E+06 | 5.922E+04 | 3.290E+04 | 1.385E+05 | 1.539E+06 | 1.749E+05 | 7.457E+04 | 2.240E+04 |
| 387 | 466.415 | 10.248 | 2 | N/A | N/A | 1.442E+06 | 1.806E+06 | 1.571E+06 | 1.557E+06 | 1.665E+06 | 1.924E+06 | 1.410E+06 | 4.591E+04 | 1.164E+05 | 2.370E+04 | 1.136E+05 | 6.378E+04 | 1.863E+05 | 9.739E+04 |
| 389 | 526.434 | 9.632 | 2 | 1.462E+05 | 67.154 | 1.707E+06 | 2.443E+06 | 1.748E+06 | 2.058E+06 | 2.158E+06 | 2.546E+06 | 1.774E+06 | 1.262E+04 | 3.240E+05 | 6.647E+03 | 2.444E+05 | 2.265E+05 | 3.956E+05 | 6.019E+04 |
| 390 | 468.390 | 9.206 | 2 | 1.280E+05 | 53.137 | 1.054E+06 | 1.562E+06 | 1.191E+06 | 1.261E+06 | 1.606E+06 | 1.719E+06 | 1.500E+06 | 3.164E+04 | 1.311E+05 | 2.849E+04 | 2.053E+05 | 1.986E+05 | 1.658E+05 | 1.665E+05 |
| 399 | 520.331 | 4.142 | 2 | 1.329E+05 | 61.051 | 1.909E+06 | 2.236E+06 | 1.908E+06 | 2.106E+06 | 2.086E+06 | 2.182E+06 | 1.854E+06 | 6.571E+03 | 1.408E+04 | 5.053E+04 | 8.312E+04 | 1.942E+05 | 1.382E+05 | 8.211E+04 |
| 400 | 484.385 | 8.410 | 2 | 1.665E+05 | 69.131 | 1.052E+06 | 1.821E+06 | 1.443E+06 | 1.760E+06 | 1.751E+06 | 1.745E+06 | 1.621E+06 | 3.715E+05 | 3.233E+05 | 6.910E+04 | 1.293E+05 | 3.169E+05 | 3.142E+04 | 4.210E+05 |
| 412 | 512.416 | 9.184 | 2 | 2.340E+05 | 97.164 | 1.029E+06 | 1.847E+06 | 1.148E+06 | 1.249E+06 | 1.757E+06 | 1.809E+06 | 1.617E+06 | 1.991E+04 | 3.040E+05 | 5.792E+04 | 3.145E+05 | 3.438E+05 | 2.437E+05 | 1.888E+05 |
| 414 | 528.411 | 8.398 | 2 | 1.505E+05 | 69.131 | 9.031E+05 | 1.596E+06 | 1.336E+06 | 1.500E+06 | 1.551E+06 | 1.592E+06 | 1.415E+06 | 2.579E+05 | 3.226E+05 | 5.479E+04 | 2.017E+04 | 2.824E+05 | 4.755E+04 | 3.726E+05 |
| 416 | 564.360 | 4.248 | 2 | 1.213E+05 | 61.054 | 1.497E+06 | 1.488E+06 | 1.297E+06 | 1.440E+06 | 1.426E+06 | 1.327E+06 | 1.213E+06 | 1.959E+04 | 1.014E+05 | 6.131E+04 | 1.707E+04 | 7.526E+04 | 8.430E+03 | 8.745E+04 |
| 417 | 644.495 | 9.227 | 2 | 4.693E+03 | 3.010 | 5.738E+05 | 1.048E+06 | 1.852E+06 | 4.242E+06 | 1.461E+06 | 1.177E+06 | 9.320E+05 | 6.359E+04 | 2.204E+05 | 6.302E+05 | 1.407E+06 | 3.989E+05 | 1.119E+05 | 5.339E+04 |
| 422 | 480.390 | 10.316 | 2 | 1.569E+05 | 65.136 | 1.766E+06 | 1.636E+06 | 1.817E+06 | 1.669E+06 | 1.767E+06 | 1.771E+06 | 1.648E+06 | 5.493E+04 | 4.178E+04 | 6.343E+04 | 4.965E+03 | 4.601E+04 | 5.762E+04 | 5.186E+04 |
| 432 | 556.443 | 9.161 | 2 | 2.116E+05 | 97.163 | 8.511E+05 | 1.875E+06 | 9.392E+05 | 1.033E+06 | 1.651E+06 | 1.772E+06 | 1.514E+06 | 2.104E+04 | 4.681E+05 | 7.894E+04 | 3.099E+05 | 4.749E+05 | 4.116E+05 | 2.845E+05 |
| 438 | 584.475 | 9.912 | 2 | 1.613E+05 | 81.169 | 9.982E+05 | 2.033E+06 | 1.315E+06 | 2.770E+06 | 1.430E+06 | 2.171E+06 | 1.003E+06 | 4.333E+04 | 6.141E+05 | 1.772E+05 | 8.040E+05 | 2.330E+05 | 7.217E+05 | 1.003E+05 |
| 455 | 628.501 | 9.881 | 2 | N/A | N/A | 7.090E+05 | 1.677E+06 | 1.090E+06 | 2.145E+06 | 1.065E+06 | 1.275E+06 | 1.057E+06 | 1.497E+05 | 5.519E+05 | 2.277E+05 | 6.309E+05 | 1.972E+05 | 2.661E+05 | 2.307E+05 |
| 464 | 608.392 | 4.346 | 2 | N/A | N/A | 1.010E+06 | 1.077E+06 | 9.295E+05 | 9.838E+05 | 9.700E+05 | 8.868E+05 | 8.106E+05 | 8.400E+04 | 4.662E+04 | 6.475E+04 | 3.444E+04 | 6.412E+04 | 6.909E+04 | 1.362E+05 |
| 471 | 352.306 | 8.095 | 2 | N/A | N/A | 9.663E+04 | 3.429E+05 | 2.563E+06 | 4.436E+07 | 7.768E+06 | 2.845E+07 | 1.581E+07 | 1.242E+04 | 1.385E+05 | 1.319E+06 | 2.227E+07 | 2.335E+06 | 1.240E+07 | 2.591E+06 |
| 528 | 308.279 | 8.075 | 2 | N/A | N/A | 3.662E+04 | 1.244E+05 | 1.808E+06 | 4.188E+07 | 5.134E+06 | 1.600E+07 | 9.170E+06 | 8.053E+03 | 1.810E+04 | 1.194E+06 | 2.381E+07 | 2.472E+06 | 6.378E+06 | 2.086E+06 |
| 539 | 688.521 | 9.203 | 2 | N/A | N/A | 2.185E+05 | 7.508E+05 | 1.401E+06 | 2.726E+06 | 7.574E+05 | 2.003E+05 | 4.393E+05 | 7.243E+04 | 1.379E+05 | 5.057E+05 | 8.887E+05 | 3.938E+05 | 2.003E+05 | 2.198E+05 |
| 541 | 363.310 | 9.363 | 2 | 2.830E+05 | 80.135 | 8.787E+05 | 9.901E+05 | 1.117E+06 | 4.099E+06 | 1.370E+06 | 1.064E+06 | 8.625E+05 | 1.111E+04 | 7.309E+04 | 1.420E+05 | 1.422E+06 | 2.147E+05 | 3.545E+04 | 4.680E+04 |
| 546 | 291.253 | 8.075 | 2 | N/A | N/A | 1.612E+04 | 9.534E+04 | 8.998E+05 | 2.856E+07 | 3.744E+06 | 1.141E+07 | 6.539E+06 | 2.684E+03 | 1.829E+04 | 7.054E+05 | 1.576E+07 | 1.617E+06 | 4.620E+06 | 1.546E+06 |
| 548 | 396.333 | 8.097 | 2 | N/A | N/A | 1.189E+05 | 6.343E+05 | 1.738E+06 | 3.006E+07 | 7.459E+06 | 2.921E+07 | 1.651E+07 | 4.626E+04 | 2.865E+05 | 1.264E+06 | 1.439E+07 | 1.858E+06 | 1.274E+07 | 2.065E+06 |
| 556 | 732.548 | 9.180 | 2 | N/A | N/A | 1.408E+05 | 6.343E+05 | 1.065E+06 | 1.742E+06 | 6.138E+05 | 4.904E+05 | 2.727E+05 | 6.157E+03 | 1.257E+05 | 3.925E+05 | 5.232E+05 | 1.353E+05 | 1.106E+05 | 1.330E+05 |
| 564 | 440.359 | 8.096 | 2 | N/A | N/A | 1.998E+05 | 1.110E+06 | 1.470E+06 | 2.111E+07 | 7.512E+06 | 2.599E+07 | 1.633E+07 | 2.265E+04 | 4.796E+05 | 1.111E+06 | 9.061E+06 | 1.567E+06 | 9.658E+06 | 2.192E+06 |

| Feature identifiers | | | | | | Average XIC | | | | | | | Standard Err of Average XIC | | | | | | |
|---|---|---|---|---|---|---|---|---|---|---|---|---|---|---|---|---|---|---|---|
| ClusterIdx | m/z | RT [min] | Subnetwork | PPM | MassDiff | TUB00 | 4TUB34 | 4TUB81 | 4TUB93 | 5TUB23 | 5TUB48 | 5TUB83 | TUB00 | 4TUB34 | 4TUB81 | 4TUB93 | 5TUB23 | 5TUB48 | 5TUB83 |
| 570 | 484.385 | 8.094 | 2 | N/A | N/A | 7.377E+04 | 7.930E+05 | 1.148E+06 | 1.518E+07 | 6.330E+06 | 1.908E+07 | 1.319E+07 | 2.363E+04 | 4.791E+05 | 7.309E+05 | 6.307E+06 | 1.261E+06 | 5.854E+06 | 2.111E+06 |
| 571 | 308.279 | 8.290 | 2 | N/A | N/A | 1.559E+05 | 1.689E+05 | 2.015E+05 | 5.150E+06 | 5.633E+05 | 1.605E+06 | 8.266E+05 | 4.457E+04 | 8.201E+04 | 1.819E+05 | 2.319E+06 | 2.455E+05 | 8.497E+05 | 1.695E+05 |
| 573 | 528.411 | 8.088 | 2 | N/A | N/A | 4.510E+04 | 4.590E+05 | 8.091E+05 | 1.051E+07 | 5.095E+06 | 1.266E+07 | 9.473E+06 | 9.075E+02 | 3.491E+05 | 6.237E+05 | 4.370E+06 | 1.051E+06 | 3.176E+06 | 1.947E+06 |
| 576 | 247.227 | 8.064 | 2 | 2.399E+05 | 78.031 | 3.665E+03 | 1.100E+04 | 9.534E+04 | 1.066E+07 | 4.030E+05 | 1.605E+06 | 5.588E+05 | 2.082E+03 | 8.238E+03 | 6.243E+04 | 6.926E+06 | 1.965E+05 | 7.755E+05 | 2.363E+05 |
| 579 | 335.280 | 8.098 | 2 | 1.840E+05 | 52.105 | 1.379E+04 | 6.170E+04 | 6.514E+05 | 1.415E+07 | 2.656E+06 | 9.854E+06 | 5.941E+06 | 5.179E+03 | 1.661E+04 | 4.964E+05 | 7.013E+06 | 1.109E+06 | 4.804E+06 | 9.639E+05 |
| 580 | 572.437 | 8.084 | 2 | 1.374E+05 | 69.131 | 2.069E+04 | 2.209E+05 | 2.791E+05 | 6.582E+06 | 3.887E+06 | 8.007E+06 | 6.484E+06 | 5.470E+03 | 9.443E+04 | 1.781E+05 | 2.707E+06 | 8.625E+05 | 1.711E+06 | 1.619E+06 |
| 598 | 396.333 | 8.315 | 2 | N/A | N/A | 6.661E+04 | 9.600E+04 | 2.587E+05 | 2.365E+06 | 6.084E+05 | 2.171E+06 | 1.457E+06 | 1.220E+04 | 3.220E+04 | 1.616E+05 | 1.077E+06 | 2.375E+05 | 1.072E+06 | 2.283E+05 |
| 604 | 440.359 | 8.302 | 2 | N/A | N/A | 1.412E+05 | 1.312E+05 | 1.219E+05 | 2.661E+06 | 5.741E+05 | 2.118E+06 | 1.253E+06 | 4.755E+04 | 3.670E+04 | 5.990E+04 | 1.318E+06 | 1.731E+05 | 8.723E+05 | 2.293E+05 |
| 608 | 672.527 | 9.854 | 2 | N/A | N/A | 6.019E+05 | 1.325E+06 | 9.538E+05 | 1.570E+06 | 7.548E+05 | 8.892E+05 | 7.278E+05 | 1.033E+05 | 4.521E+05 | 1.863E+05 | 3.528E+05 | 1.633E+05 | 2.095E+05 | 1.629E+05 |
| 615 | 350.328 | 9.687 | 2 | N/A | N/A | 4.714E+06 | 5.118E+06 | 2.681E+06 | 4.431E+06 | 2.464E+06 | 4.705E+06 | 1.901E+06 | 2.364E+06 | 2.321E+06 | 1.526E+05 | 2.553E+06 | 3.628E+05 | 2.682E+06 | 1.055E+05 |
| 618 | 658.511 | 9.464 | 2 | 2.654E+04 | 17.026 | 7.715E+05 | 2.696E+06 | 7.566E+05 | 1.155E+06 | 1.928E+06 | 2.228E+06 | 1.734E+06 | 8.563E+04 | 8.783E+05 | 1.328E+05 | 3.710E+05 | 6.379E+05 | 6.525E+05 | 3.065E+05 |
| 625 | 702.536 | 9.449 | 2 | N/A | N/A | 5.349E+05 | 1.751E+06 | 5.821E+05 | 7.554E+05 | 1.058E+06 | 1.474E+06 | 9.679E+05 | 1.170E+04 | 6.158E+05 | 7.901E+04 | 1.632E+05 | 4.407E+05 | 5.159E+05 | 1.137E+05 |
| 647 | 394.355 | 9.697 | 2 | N/A | N/A | 2.288E+06 | 4.987E+06 | 2.346E+06 | 2.437E+06 | 4.844E+06 | 5.350E+06 | 2.157E+06 | 6.731E+04 | 2.588E+06 | 2.387E+05 | 2.016E+05 | 2.629E+06 | 2.968E+06 | 3.313E+05 |
| 50 | 701.620 | 10.613 | 1 | N/A | N/A | 1.081E+07 | 9.633E+06 | 8.856E+06 | 7.113E+06 | 7.042E+06 | 6.557E+06 | 6.903E+06 | 6.501E+05 | 9.567E+04 | 1.380E+05 | 2.535E+05 | 5.305E+05 | 1.980E+05 | 1.060E+05 |
| 199 | 419.364 | 7.499 | 1 | N/A | N/A | 3.314E+06 | 3.047E+06 | 3.196E+06 | 3.005E+06 | 3.048E+06 | 2.852E+06 | 2.668E+06 | 8.700E+04 | 3.579E+04 | 1.621E+05 | 8.024E+04 | 1.257E+05 | 1.734E+05 | 7.405E+04 |
| 689 | 685.624 | 12.053 | 1 | N/A | N/A | 9.169E+05 | 6.524E+05 | 5.499E+05 | 3.508E+05 | 3.018E+05 | 3.384E+05 | 4.653E+05 | 3.047E+05 | 1.222E+05 | 3.557E+04 | 8.554E+04 | 9.666E+04 | 5.241E+04 | 6.884E+04 |
| 1 | 110.020 | 12.761 | -1 | N/A | N/A | 1.076E+09 | 1.379E+09 | 1.471E+09 | 1.442E+09 | 1.361E+09 | 1.299E+09 | 1.247E+09 | 1.348E+08 | 1.893E+08 | 2.030E+07 | 1.577E+07 | 2.251E+07 | 3.267E+07 | 1.979E+07 |
| 8 | 437.194 | 7.524 | -1 | N/A | N/A | 1.965E+08 | 2.672E+08 | 2.636E+08 | 2.708E+08 | 2.748E+08 | 2.837E+08 | 2.810E+08 | 7.268E+06 | 1.113E+07 | 1.047E+07 | 7.799E+06 | 1.732E+07 | 1.082E+07 | 4.254E+07 |
| 10 | 120.081 | 0.519 | -1 | 1.304E+05 | 18.011 | 6.687E+08 | 1.612E+09 | 1.324E+08 | 4.576E+08 | 8.551E+08 | 9.000E+08 | 1.304E+08 | 7.426E+07 | 3.555E+08 | 2.185E+07 | 1.062E+08 | 5.878E+07 | 7.166E+08 | 3.250E+07 |
| 13 | 242.284 | 8.651 | -1 | 1.308E+05 | 28.032 | 1.391E+08 | 1.331E+08 | 2.357E+08 | 2.369E+08 | 2.270E+08 | 2.303E+08 | 2.325E+08 | 4.738E+07 | 5.489E+07 | 1.933E+06 | 6.034E+06 | 8.666E+05 | 3.807E+06 | 2.904E+06 |
| 14 | 851.398 | 7.527 | -1 | N/A | N/A | 3.395E+07 | 7.098E+07 | 6.717E+07 | 6.866E+07 | 8.061E+07 | 8.710E+07 | 9.611E+07 | 3.321E+06 | 1.054E+07 | 8.608E+06 | 3.852E+06 | 1.602E+07 | 8.214E+06 | 4.129E+07 |
| 32 | 437.194 | 7.249 | -1 | N/A | N/A | 6.594E+06 | 1.504E+07 | 1.995E+07 | 2.383E+07 | 2.234E+07 | 2.199E+07 | 2.094E+07 | 1.625E+05 | 5.708E+06 | 2.016E+06 | 1.022E+07 | 6.548E+06 | 7.377E+06 | 7.868E+06 |
| 33 | 559.518 | 11.442 | -1 | N/A | N/A | 1.226E+08 | 1.251E+08 | 1.265E+08 | 1.214E+08 | 1.201E+08 | 1.144E+08 | 1.146E+08 | 9.425E+05 | 8.459E+05 | 1.677E+06 | 1.125E+06 | 3.043E+05 | 4.119E+06 | 2.754E+06 |
| 34 | 867.372 | 7.528 | -1 | N/A | N/A | 1.938E+07 | 2.879E+07 | 2.892E+07 | 2.892E+07 | 3.172E+07 | 3.255E+07 | 3.164E+07 | 1.485E+06 | 1.725E+06 | 2.017E+06 | 1.108E+06 | 3.908E+06 | 1.268E+06 | 6.290E+06 |
| 41 | 587.549 | 11.407 | -1 | N/A | N/A | 1.178E+08 | 1.224E+08 | 1.229E+08 | 1.182E+08 | 1.154E+08 | 1.110E+08 | 1.112E+08 | 2.971E+06 | 2.055E+05 | 2.400E+06 | 1.202E+06 | 2.467E+06 | 3.733E+06 | 1.583E+06 |
| 47 | 163.991 | 12.748 | -1 | N/A | N/A | 1.733E+07 | 1.740E+07 | 1.659E+07 | 1.510E+07 | 1.459E+07 | 1.409E+07 | 1.423E+07 | 4.141E+05 | 1.101E+06 | 1.604E+05 | 3.449E+05 | 5.151E+05 | 2.532E+05 | 2.287E+05 |
| 53 | 243.088 | 4.621 | -1 | N/A | N/A | 4.661E+06 | 1.356E+07 | 9.875E+06 | 1.740E+07 | 9.455E+06 | 1.424E+07 | 1.112E+07 | 3.585E+05 | 2.853E+06 | 1.432E+06 | 3.350E+06 | 4.750E+05 | 3.592E+05 | 1.102E+06 |
| 54 | 399.209 | 10.100 | -1 | N/A | N/A | 1.105E+07 | 1.213E+07 | 1.205E+07 | 1.138E+07 | 1.184E+07 | 8.545E+06 | 7.642E+06 | 4.637E+05 | 2.982E+05 | 6.086E+05 | 4.861E+05 | 3.598E+05 | 8.896E+05 | 8.813E+05 |
| 57 | 685.436 | 12.425 | -1 | N/A | N/A | 2.162E+07 | 2.369E+07 | 1.921E+07 | 2.007E+07 | 5.178E+07 | 5.778E+07 | 5.202E+07 | 1.411E+06 | 1.928E+06 | 4.795E+05 | 1.388E+06 | 5.895E+06 | 1.435E+07 | 1.819E+07 |
| 58 | 250.112 | 5.753 | -1 | 2.253E+05 | 45.988 | 1.614E+05 | 6.302E+03 | 1.270E+07 | 6.322E+06 | 1.281E+05 | 0.000E+00 | 3.261E+04 | 6.516E+04 | 4.594E+03 | 1.393E+06 | 6.147E+06 | 6.400E+04 | 0.000E+00 | 1.007E+04 |
| 64 | 255.073 | 4.542 | -1 | N/A | N/A | 0.000E+00 | 1.035E+07 | 1.870E+07 | 2.041E+07 | 0.000E+00 | 0.000E+00 | 0.000E+00 | 0.000E+00 | 8.140E+05 | 3.479E+06 | 2.581E+06 | 0.000E+00 | 0.000E+00 | 0.000E+00 |
| 66 | 261.109 | 5.951 | -1 | N/A | N/A | 2.855E+07 | 1.132E+07 | 1.622E+07 | 1.971E+07 | 3.066E+07 | 1.593E+07 | 2.562E+07 | 4.379E+05 | 3.544E+06 | 3.287E+06 | 3.160E+06 | 4.155E+06 | 6.082E+05 | 5.609E+06 |
| 71 | 267.077 | 3.607 | -1 | N/A | N/A | 2.451E+06 | 2.448E+07 | 1.778E+07 | 2.906E+07 | 1.436E+07 | 2.402E+07 | 1.924E+07 | 1.566E+05 | 3.409E+06 | 1.342E+06 | 7.162E+06 | 3.169E+06 | 2.733E+06 | 4.404E+06 |
| 73 | 453.168 | 7.250 | -1 | N/A | N/A | 2.966E+06 | 6.822E+06 | 9.043E+06 | 1.060E+07 | 1.017E+07 | 9.560E+06 | 8.990E+06 | 2.415E+04 | 2.548E+06 | 7.808E+05 | 4.249E+06 | 2.931E+06 | 3.205E+06 | 3.262E+06 |
| 76 | 324.142 | 7.584 | -1 | N/A | N/A | 5.357E+06 | 1.144E+07 | 9.861E+06 | 9.276E+06 | 8.785E+06 | 1.272E+07 | 8.609E+06 | 1.464E+06 | 2.519E+06 | 1.678E+06 | 1.347E+06 | 2.149E+06 | 8.964E+05 | 2.412E+06 |
| 77 | 372.882 | 12.723 | -1 | N/A | N/A | 2.497E+06 | 1.755E+06 | 1.480E+06 | 1.557E+06 | 1.231E+06 | 1.360E+06 | 1.427E+06 | 4.769E+05 | 4.743E+05 | 2.989E+04 | 1.417E+05 | 1.275E+05 | 3.069E+05 | 4.282E+05 |
| 79 | 321.131 | 4.460 | -1 | N/A | N/A | 4.688E+06 | 1.182E+07 | 1.078E+07 | 1.273E+07 | 1.789E+07 | 2.273E+07 | 2.847E+07 | 3.780E+05 | 2.033E+06 | 1.067E+06 | 3.725E+06 | 2.450E+06 | 2.746E+06 | 1.184E+07 |
| 82 | 269.015 | 5.941 | -1 | N/A | N/A | 0.000E+00 | 7.679E+06 | 1.156E+07 | 2.136E+06 | 1.077E+04 | 0.000E+00 | 0.000E+00 | 0.000E+00 | 2.865E+06 | 9.798E+05 | 1.760E+06 | 1.077E+04 | 0.000E+00 | 0.000E+00 |
| 83 | 230.141 | 12.721 | -1 | 6.889E+04 | 17.028 | 5.854E+06 | 6.220E+06 | 6.912E+06 | 7.058E+06 | 6.990E+06 | 6.339E+06 | 5.963E+06 | 1.188E+05 | 3.529E+05 | 3.684E+05 | 1.077E+05 | 3.503E+05 | 3.303E+05 | 7.043E+05 |
| 85 | 233.079 | 4.825 | -1 | N/A | N/A | 1.441E+07 | 6.477E+06 | 6.243E+06 | 7.146E+06 | 1.460E+07 | 8.822E+06 | 1.201E+07 | 1.412E+06 | 1.082E+06 | 1.285E+06 | 1.570E+06 | 3.026E+06 | 1.227E+06 | 1.992E+06 |
| 86 | 239.089 | 4.007 | -1 | N/A | N/A | 3.175E+07 | 4.932E+07 | 2.712E+07 | 3.296E+07 | 2.213E+07 | 2.482E+07 | 2.325E+07 | 1.186E+07 | 4.929E+06 | 6.325E+06 | 3.973E+06 | 3.769E+06 | 3.286E+06 | 7.503E+06 |
| 90 | 332.913 | 12.737 | -1 | N/A | N/A | 9.737E+06 | 8.091E+06 | 7.651E+06 | 7.970E+06 | 6.734E+06 | 6.072E+06 | 6.091E+06 | 2.215E+05 | 2.231E+05 | 2.192E+05 | 7.752E+05 | 6.928E+05 | 5.717E+05 | 2.094E+05 |
| 98 | 195.102 | 7.496 | -1 | 1.747E+05 | 29.016 | 7.719E+06 | 8.005E+06 | 8.718E+06 | 8.988E+06 | 8.984E+06 | 8.817E+06 | 8.573E+06 | 3.594E+05 | 4.206E+05 | 8.337E+05 | 5.414E+05 | 5.967E+05 | 7.388E+05 | 1.126E+06 |
| 100 | 440.099 | 7.644 | -1 | N/A | N/A | 0.000E+00 | 8.657E+05 | 6.570E+06 | 5.606E+06 | 0.000E+00 | 1.464E+03 | 0.000E+00 | 0.000E+00 | 8.432E+04 | 2.700E+05 | 2.690E+06 | 0.000E+00 | 1.464E+03 | 0.000E+00 |
| 101 | 235.169 | 8.489 | -1 | N/A | N/A | 1.564E+07 | 3.835E+07 | 1.317E+07 | 2.276E+07 | 1.787E+07 | 2.597E+07 | 1.810E+07 | 3.050E+06 | 1.158E+07 | 1.300E+06 | 8.058E+06 | 2.420E+06 | 4.550E+06 | 2.882E+06 |
| 106 | 670.153 | 5.750 | -1 | N/A | N/A | 1.072E+06 | 2.235E+04 | 4.670E+06 | 3.497E+06 | 6.443E+05 | 5.525E+07 | 3.498E+07 | 5.444E+05 | 2.235E+04 | 2.016E+06 | 9.766E+05 | 4.117E+05 | 5.083E+06 | 8.828E+06 |
| 107 | 183.150 | 4.641 | -1 | N/A | N/A | 1.345E+07 | 7.951E+06 | 1.083E+07 | 1.372E+07 | 9.392E+06 | 1.617E+07 | 1.717E+07 | 6.937E+06 | 1.224E+06 | 1.015E+06 | 4.009E+05 | 5.192E+05 | 1.279E+06 | 8.474E+05 |
| 108 | 281.088 | 8.127 | -1 | N/A | N/A | 0.000E+00 | 2.072E+06 | 5.678E+06 | 5.922E+06 | 0.000E+00 | 0.000E+00 | 0.000E+00 | 0.000E+00 | 4.250E+05 | 6.616E+05 | 4.011E+06 | 0.000E+00 | 0.000E+00 | 0.000E+00 |
| 111 | 262.165 | 4.000 | -1 | N/A | N/A | 1.773E+07 | 2.996E+07 | 1.689E+07 | 2.162E+07 | 1.548E+07 | 1.585E+07 | 1.719E+07 | 6.548E+06 | 3.359E+06 | 3.842E+06 | 3.769E+06 | 2.121E+06 | 1.101E+06 | 6.020E+06 |
| 114 | 244.191 | 5.975 | -1 | 4.014E+03 | 0.984 | 6.298E+06 | 6.067E+06 | 5.183E+06 | 5.736E+06 | 5.629E+06 | 6.240E+06 | 5.665E+06 | 5.433E+05 | 2.281E+05 | 3.150E+05 | 3.124E+05 | 1.663E+05 | 2.145E+05 | 3.717E+05 |
| 116 | 228.232 | 9.984 | -1 | N/A | N/A | 8.642E+06 | 7.196E+06 | 8.920E+06 | 7.965E+06 | 5.962E+06 | 6.554E+06 | 4.800E+06 | 4.549E+05 | 1.409E+06 | 1.325E+06 | 1.333E+06 | 1.163E+06 | 1.379E+06 | 3.676E+05 |
| 118 | 219.175 | 8.820 | -1 | 6.814E+04 | 13.982 | 5.796E+06 | 5.789E+06 | 6.122E+06 | 6.175E+06 | 6.279E+06 | 6.406E+06 | 6.422E+06 | 3.909E+04 | 1.825E+05 | 1.772E+05 | 9.883E+04 | 1.147E+05 | 7.469E+04 | 3.683E+05 |
| 126 | 296.219 | 7.149 | -1 | N/A | N/A | 1.855E+06 | 2.854E+06 | 3.456E+06 | 4.354E+06 | 3.230E+06 | 4.574E+06 | 6.499E+06 | 1.679E+05 | 7.237E+05 | 8.895E+05 | 6.947E+05 | 4.076E+05 | 1.273E+06 | 1.729E+06 |
| 137 | 431.183 | 10.115 | -1 | N/A | N/A | 5.075E+06 | 5.412E+06 | 5.822E+06 | 5.501E+06 | 5.309E+06 | 4.762E+06 | 4.337E+06 | 3.853E+04 | 1.849E+05 | 5.076E+04 | 9.593E+04 | 2.209E+05 | 5.743E+05 | 3.028E+05 |
| 148 | 319.187 | 7.588 | -1 | N/A | N/A | 3.159E+06 | 6.763E+06 | 5.609E+06 | 5.149E+06 | 4.812E+06 | 7.278E+06 | 5.201E+06 | 8.703E+05 | 1.471E+06 | 9.211E+05 | 8.879E+05 | 1.215E+06 | 6.698E+05 | 1.632E+06 |
| 152 | 707.875 | 4.385 | -1 | N/A | N/A | 0.000E+00 | 1.608E+04 | 4.879E+06 | 1.628E+06 | 0.000E+00 | 0.000E+00 | 0.000E+00 | 0.000E+00 | 1.215E+04 | 4.366E+05 | 1.624E+06 | 0.000E+00 | 0.000E+00 | 0.000E+00 |
| 162 | 174.999 | 12.765 | -1 | N/A | N/A | 1.171E+07 | 1.091E+07 | 9.187E+06 | 1.079E+07 | 9.625E+06 | 8.982E+06 | 8.297E+06 | 4.234E+05 | 2.801E+05 | 3.591E+05 | 2.394E+05 | 6.390E+05 | 3.451E+05 | 4.729E+05 |
| 164 | 338.147 | 5.707 | -1 | 3.907E+05 | 94.998 | 0.000E+00 | 1.543E+04 | 2.395E+06 | 4.217E+05 | 0.000E+00 | 0.000E+00 | 0.000E+00 | 0.000E+00 | 9.693E+03 | 1.119E+06 | 3.890E+05 | 0.000E+00 | 0.000E+00 | 0.000E+00 |
| 165 | 269.210 | 8.432 | -1 | N/A | N/A | 4.005E+06 | 5.525E+06 | 7.015E+06 | 7.340E+06 | 4.980E+06 | 6.553E+06 | 6.611E+06 | 1.674E+05 | 7.256E+05 | 1.896E+05 | 8.612E+05 | 3.731E+05 | 4.541E+05 | 1.923E+06 |
| 173 | 451.246 | 9.774 | -1 | N/A | N/A | 3.404E+06 | 2.730E+06 | 6.663E+06 | 4.795E+06 | 2.192E+06 | 3.460E+06 | 2.345E+06 | 4.538E+05 | 7.167E+05 | 1.282E+06 | 2.038E+06 | 4.209E+05 | 3.484E+05 | 1.980E+05 |
| 179 | 723.608 | 10.615 | -1 | N/A | N/A | 4.477E+06 | 4.043E+06 | 3.316E+06 | 2.608E+06 | 2.649E+06 | 2.138E+06 | 2.300E+06 | 2.133E+05 | 2.425E+05 | 1.474E+05 | 3.749E+04 | 2.402E+05 | 6.582E+04 | 1.258E+05 |
| 184 | 434.186 | 7.527 | -1 | N/A | N/A | 7.408E+05 | 2.776E+06 | 2.170E+06 | 2.549E+06 | 3.883E+06 | 3.751E+06 | 7.048E+06 | 1.862E+05 | 8.536E+05 | 7.766E+05 | 3.779E+05 | 1.255E+06 | 6.941E+05 | 4.819E+06 |
| 188 | 270.279 | 10.247 | -1 | N/A | N/A | 3.975E+06 | 3.137E+06 | 3.826E+06 | 3.273E+06 | 2.833E+06 | 3.030E+06 | 2.953E+06 | 1.422E+05 | 9.065E+04 | 1.071E+05 | 4.271E+05 | 2.827E+05 | 3.451E+05 | 2.319E+05 |
| 193 | 242.284 | 7.258 | -1 | N/A | N/A | 1.460E+07 | 8.583E+06 | 5.862E+06 | 4.170E+06 | 3.177E+06 | 2.657E+06 | 2.281E+06 | 1.427E+06 | 5.689E+05 | 5.176E+05 | 2.362E+05 | 1.833E+05 | 1.529E+05 | 3.986E+04 |
| 195 | 312.326 | 10.525 | -1 | N/A | N/A | 2.375E+06 | 1.895E+06 | 2.795E+06 | 2.321E+06 | 2.008E+06 | 2.572E+06 | 2.062E+06 | 1.048E+05 | 3.485E+05 | 2.340E+05 | 4.768E+05 | 3.339E+05 | 5.978E+05 | 2.705E+05 |
| 198 | 499.165 | 7.526 | -1 | N/A | N/A | 2.760E+06 | 5.194E+06 | 4.754E+06 | 5.439E+06 | 5.863E+06 | 6.225E+06 | 6.980E+06 | 2.482E+05 | 6.796E+05 | 3.449E+05 | 2.935E+05 | 9.678E+05 | 6.284E+05 | 2.514E+06 |
| 203 | 122.547 | 1.761 | -1 | N/A | N/A | 3.237E+06 | 3.508E+06 | 1.938E+06 | 1.919E+06 | 9.047E+05 | 1.582E+06 | 1.213E+06 | 7.656E+05 | 1.044E+06 | 4.367E+05 | 3.007E+05 | 8.290E+04 | 6.090E+05 | 6.137E+05 |
| 212 | 409.201 | 10.114 | -1 | N/A | N/A | 4.090E+06 | 3.463E+06 | 4.507E+06 | 4.116E+06 | 4.214E+06 | 5.467E+06 | 5.201E+06 | 6.499E+04 | 2.792E+05 | 1.179E+05 | 6.069E+04 | 9.377E+04 | 2.155E+05 | 3.407E+05 |
| 214 | 270.279 | 9.930 | -1 | N/A | N/A | 7.978E+06 | 6.460E+06 | 5.943E+06 | 5.339E+06 | 4.712E+06 | 5.210E+06 | 5.261E+06 | 4.062E+05 | 3.653E+05 | 2.174E+05 | 1.292E+05 | 9.395E+04 | 4.561E+05 | 1.906E+05 |
| 217 | 266.248 | 6.949 | -1 | N/A | N/A | 1.786E+04 | 0.000E+00 | 5.676E+06 | 6.862E+05 | 0.000E+00 | 0.000E+00 | 0.000E+00 | 3.563E+03 | 0.000E+00 | 1.781E+06 | 6.862E+05 | 0.000E+00 | 0.000E+00 | 0.000E+00 |

| Feature identifiers | | | | | | Average XIC | | | | | | | Standard Err of Average XIC | | | | | | |
|---|---|---|---|---|---|---|---|---|---|---|---|---|---|---|---|---|---|---|---|
| ClusterIdx | m/z | RT [min] | Subnetwork | PPM | MassDiff | TUB00 | 4TUB34 | 4TUB81 | 4TUB93 | 5TUB23 | 5TUB48 | 5TUB83 | TUB00 | 4TUB34 | 4TUB81 | 4TUB93 | 5TUB23 | 5TUB48 | 5TUB83 |
| 222 | 495.300 | 10.647 | -1 | N/A | N/A | 1.120E+06 | 8.418E+05 | 1.452E+06 | 1.201E+06 | 1.161E+06 | 1.109E+06 | 1.093E+06 | 1.178E+05 | 9.668E+04 | 1.380E+05 | 2.144E+05 | 1.622E+05 | 2.036E+05 | 1.721E+05 |
| 225 | 330.300 | 9.151 | -1 | 5.089E+04 | 15.996 | 8.859E+05 | 1.175E+06 | 2.530E+06 | 4.575E+06 | 9.814E+04 | 9.655E+03 | 6.696E+02 | 5.778E+04 | 4.222E+05 | 1.150E+05 | 2.423E+06 | 3.380E+04 | 5.556E+03 | 6.696E+02 |
| 232 | 250.178 | 6.460 | -1 | N/A | N/A | 7.678E+06 | 1.315E+07 | 1.089E+07 | 1.445E+07 | 9.460E+06 | 1.366E+07 | 1.571E+07 | 6.043E+05 | 3.626E+06 | 2.151E+06 | 2.423E+06 | 9.762E+05 | 3.416E+06 | 4.668E+06 |
| 234 | 555.250 | 4.014 | -1 | N/A | N/A | 0.000E+00 | 0.000E+00 | 2.326E+06 | 2.731E+05 | 0.000E+00 | 0.000E+00 | 0.000E+00 | 0.000E+00 | 0.000E+00 | 5.758E+05 | 2.731E+05 | 0.000E+00 | 0.000E+00 | 0.000E+00 |
| 236 | 149.988 | 12.784 | -1 | N/A | N/A | 9.453E+06 | 9.053E+06 | 8.089E+06 | 8.133E+06 | 8.167E+06 | 7.740E+06 | 7.446E+06 | 4.333E+05 | 9.986E+05 | 1.379E+05 | 8.334E+04 | 2.711E+05 | 2.652E+05 | 3.500E+05 |
| 237 | 640.348 | 4.374 | -1 | N/A | N/A | 0.000E+00 | 2.091E+04 | 2.514E+06 | 1.199E+06 | 1.012E+04 | 2.541E+04 | 3.015E+04 | 0.000E+00 | 1.372E+04 | 2.120E+05 | 1.177E+06 | 6.285E+03 | 1.271E+04 | 2.445E+04 |
| 252 | 157.084 | 0.657 | -1 | N/A | N/A | 2.161E+07 | 3.424E+07 | 1.351E+07 | 1.137E+07 | 2.832E+07 | 1.668E+07 | 1.542E+07 | 3.802E+06 | 4.520E+06 | 1.395E+06 | 1.983E+06 | 6.525E+06 | 1.367E+06 | 5.095E+05 |
| 254 | 111.039 | 13.492 | -1 | N/A | N/A | 2.465E+06 | 1.625E+06 | 1.603E+06 | 2.744E+06 | 2.051E+06 | 1.528E+06 | 1.744E+06 | 1.062E+06 | 9.326E+05 | 2.370E+05 | 7.863E+05 | 1.289E+05 | 1.551E+05 | 2.988E+05 |
| 256 | 297.241 | 9.395 | -1 | N/A | N/A | 2.999E+06 | 3.708E+06 | 2.955E+06 | 4.803E+06 | 3.240E+06 | 2.962E+06 | 2.814E+06 | 2.524E+05 | 6.318E+05 | 9.898E+04 | 1.043E+06 | 5.604E+04 | 3.996E+04 | 1.522E+05 |
| 259 | 249.185 | 8.504 | -1 | N/A | N/A | 2.607E+06 | 4.371E+06 | 3.338E+06 | 3.746E+06 | 3.242E+06 | 3.691E+06 | 3.388E+06 | 1.402E+05 | 7.377E+05 | 1.683E+05 | 5.376E+05 | 2.950E+05 | 1.616E+05 | 2.430E+05 |
| 260 | 290.269 | 6.924 | -1 | N/A | N/A | 2.498E+06 | 2.418E+06 | 2.277E+06 | 2.232E+06 | 2.264E+06 | 2.104E+06 | 2.213E+06 | 6.161E+04 | 3.790E+04 | 1.432E+05 | 5.458E+04 | 5.872E+04 | 1.440E+05 | 4.967E+04 |
| 265 | 535.153 | 8.517 | -1 | N/A | N/A | 0.000E+00 | 1.793E+05 | 2.121E+06 | 2.473E+06 | 0.000E+00 | 0.000E+00 | 0.000E+00 | 0.000E+00 | 8.965E+04 | 7.055E+05 | 2.272E+06 | 0.000E+00 | 0.000E+00 | 0.000E+00 |
| 267 | 453.168 | 7.817 | -1 | N/A | N/A | 1.563E+06 | 6.423E+06 | 2.374E+06 | 3.695E+06 | 4.369E+06 | 5.101E+06 | 3.886E+06 | 7.252E+04 | 1.016E+06 | 2.973E+05 | 1.691E+06 | 3.786E+05 | 1.082E+06 | 1.055E+06 |
| 271 | 283.104 | 5.343 | -1 | N/A | N/A | 0.000E+00 | 1.828E+06 | 2.927E+06 | 4.842E+06 | 0.000E+00 | 0.000E+00 | 0.000E+00 | 0.000E+00 | 3.917E+05 | 8.944E+05 | 1.781E+06 | 0.000E+00 | 0.000E+00 | 0.000E+00 |
| 274 | 537.311 | 10.526 | -1 | N/A | N/A | 1.103E+06 | 2.100E+06 | 1.310E+06 | 1.602E+06 | 3.317E+06 | 3.686E+06 | 2.798E+06 | 2.614E+05 | 1.009E+05 | 6.375E+04 | 3.099E+05 | 3.619E+05 | 4.019E+05 | 1.058E+06 |
| 279 | 149.024 | 6.761 | -1 | N/A | N/A | 4.232E+06 | 1.109E+07 | 4.035E+06 | 6.679E+06 | 4.174E+06 | 5.740E+06 | 5.345E+06 | 1.081E+06 | 3.036E+06 | 5.701E+05 | 3.309E+06 | 1.114E+05 | 1.724E+06 | 1.545E+06 |
| 281 | 467.147 | 6.891 | -1 | N/A | N/A | 4.391E+06 | 7.261E+06 | 5.412E+06 | 5.841E+06 | 7.382E+06 | 7.247E+06 | 1.476E+07 | 6.619E+05 | 9.816E+05 | 4.308E+05 | 3.835E+05 | 4.294E+05 | 9.007E+05 | 9.937E+06 |
| 285 | 408.309 | 10.166 | -1 | N/A | N/A | 1.087E+06 | 2.600E+06 | 3.095E+06 | 8.418E+05 | 1.961E+06 | 2.714E+06 | 1.258E+06 | 1.902E+05 | 6.881E+05 | 8.825E+05 | 2.755E+05 | 2.576E+04 | 6.186E+05 | 5.052E+05 |
| 286 | 291.195 | 8.162 | -1 | N/A | N/A | 9.403E+05 | 1.754E+06 | 2.638E+06 | 2.710E+06 | 2.330E+06 | 2.903E+06 | 2.238E+06 | 3.114E+05 | 2.260E+05 | 1.860E+05 | 5.292E+04 | 2.210E+05 | 3.319E+05 | 2.496E+05 |
| 287 | 573.110 | 8.520 | -1 | N/A | N/A | 0.000E+00 | 7.133E+04 | 1.517E+06 | 1.498E+06 | 0.000E+00 | 0.000E+00 | 0.000E+00 | 0.000E+00 | 3.577E+04 | 5.452E+05 | 1.394E+06 | 0.000E+00 | 0.000E+00 | 0.000E+00 |
| 299 | 437.194 | 8.754 | -1 | N/A | N/A | 2.586E+06 | 4.849E+06 | 2.846E+06 | 5.225E+06 | 3.302E+06 | 3.385E+06 | 3.366E+06 | 8.280E+04 | 5.073E+05 | 9.568E+04 | 2.624E+06 | 3.097E+05 | 3.218E+05 | 5.635E+05 |
| 300 | 604.812 | 4.484 | -1 | N/A | N/A | 0.000E+00 | 7.211E+03 | 1.261E+06 | 2.041E+05 | 0.000E+00 | 0.000E+00 | 0.000E+00 | 0.000E+00 | 7.211E+03 | 2.131E+05 | 2.041E+05 | 0.000E+00 | 0.000E+00 | 0.000E+00 |
| 304 | 365.137 | 8.808 | -1 | N/A | N/A | 1.117E+07 | 9.532E+06 | 8.426E+06 | 8.811E+06 | 8.258E+06 | 7.964E+06 | 7.325E+06 | 1.233E+06 | 7.373E+05 | 6.726E+05 | 6.706E+05 | 4.765E+05 | 6.154E+05 | 4.042E+05 |
| 305 | 480.273 | 3.605 | -1 | N/A | N/A | 0.000E+00 | 4.950E+03 | 1.761E+06 | 1.238E+05 | 0.000E+00 | 1.097E+04 | 0.000E+00 | 0.000E+00 | 4.950E+03 | 1.121E+06 | 1.238E+05 | 0.000E+00 | 1.097E+04 | 0.000E+00 |
| 308 | 659.288 | 10.020 | -1 | N/A | N/A | 2.319E+05 | 5.587E+06 | 1.607E+06 | 4.231E+06 | 6.189E+06 | 1.284E+07 | 1.157E+07 | 4.064E+04 | 2.711E+06 | 2.280E+05 | 3.559E+06 | 2.644E+06 | 3.595E+06 | 3.733E+06 |
| 309 | 165.076 | 4.458 | -1 | 3.414E+05 | 42.011 | 1.012E+06 | 2.137E+06 | 2.162E+06 | 2.366E+06 | 3.229E+06 | 3.689E+06 | 5.428E+06 | 4.962E+04 | 3.913E+05 | 1.970E+05 | 7.320E+05 | 5.537E+05 | 4.293E+05 | 2.504E+06 |
| 320 | 341.267 | 9.373 | -1 | N/A | N/A | 2.005E+06 | 2.241E+06 | 2.356E+06 | 6.460E+06 | 2.797E+06 | 2.375E+06 | 2.073E+06 | 6.008E+04 | 3.996E+04 | 1.694E+05 | 2.028E+06 | 3.310E+05 | 9.439E+04 | 9.644E+04 |
| 324 | 270.315 | 9.426 | -1 | N/A | N/A | 1.359E+07 | 8.913E+06 | 9.096E+06 | 7.112E+06 | 1.418E+07 | 1.509E+07 | 1.629E+07 | 1.281E+06 | 1.962E+06 | 1.612E+06 | 1.667E+06 | 3.011E+06 | 5.037E+06 | 3.394E+06 |
| 325 | 326.378 | 9.409 | -1 | N/A | N/A | 4.139E+06 | 6.779E+06 | 3.711E+06 | 4.167E+06 | 6.422E+06 | 4.581E+06 | 3.633E+06 | 3.311E+05 | 4.576E+05 | 2.327E+04 | 5.065E+05 | 9.240E+05 | 3.823E+05 | 9.770E+04 |
| 327 | 131.049 | 0.513 | -1 | N/A | N/A | 1.238E+07 | 2.959E+07 | 1.964E+06 | 6.349E+06 | 1.529E+07 | 1.221E+07 | 2.134E+06 | 1.440E+06 | 6.529E+06 | 9.183E+05 | 2.960E+06 | 1.891E+06 | 9.286E+06 | 9.134E+05 |
| 331 | 260.165 | 6.499 | -1 | N/A | N/A | 9.113E+06 | 1.072E+07 | 1.416E+06 | 1.667E+06 | 7.178E+06 | 3.647E+06 | 1.556E+06 | 2.044E+05 | 1.871E+06 | 8.773E+04 | 1.542E+05 | 1.120E+06 | 1.890E+06 | 1.679E+05 |
| 334 | 469.184 | 6.409 | -1 | N/A | N/A | 1.414E+06 | 1.623E+06 | 1.141E+06 | 2.033E+06 | 1.266E+06 | 2.121E+06 | 1.345E+06 | 1.945E+05 | 1.131E+05 | 1.491E+05 | 2.597E+05 | 7.131E+04 | 3.122E+05 | 4.052E+05 |
| 335 | 469.163 | 6.314 | -1 | N/A | N/A | 9.713E+05 | 2.092E+06 | 1.479E+06 | 1.638E+06 | 1.834E+06 | 2.096E+06 | 9.316E+06 | 2.201E+05 | 9.555E+04 | 1.214E+05 | 1.279E+05 | 1.228E+05 | 2.492E+05 | 8.259E+06 |
| 338 | 499.780 | 3.714 | -1 | N/A | N/A | 0.000E+00 | 0.000E+00 | 3.258E+06 | 1.432E+06 | 0.000E+00 | 0.000E+00 | 0.000E+00 | 0.000E+00 | 0.000E+00 | 3.239E+05 | 1.432E+06 | 0.000E+00 | 0.000E+00 | 0.000E+00 |
| 339 | 615.581 | 10.604 | -1 | N/A | N/A | 5.889E+06 | 1.489E+07 | 1.569E+07 | 1.551E+07 | 1.520E+07 | 1.388E+07 | 1.325E+07 | 5.025E+06 | 1.629E+05 | 2.090E+05 | 1.060E+05 | 3.634E+05 | 1.753E+05 | 1.438E+05 |
| 340 | 354.553 | 5.749 | -1 | N/A | N/A | 1.918E+05 | 2.259E+03 | 9.969E+05 | 8.094E+05 | 1.160E+05 | 1.215E+07 | 7.596E+06 | 1.063E+05 | 2.259E+03 | 4.865E+05 | 2.066E+05 | 8.819E+04 | 7.041E+05 | 1.750E+06 |
| 345 | 236.142 | 5.355 | -1 | N/A | N/A | 8.979E+05 | 3.381E+06 | 2.435E+06 | 2.233E+06 | 2.130E+06 | 3.183E+06 | 2.058E+06 | 2.076E+05 | 6.122E+05 | 3.828E+05 | 2.889E+05 | 4.178E+05 | 2.240E+05 | 8.301E+05 |
| 351 | 451.173 | 6.881 | -1 | N/A | N/A | 3.699E+06 | 5.397E+06 | 4.531E+06 | 5.149E+06 | 7.442E+06 | 5.203E+06 | 1.698E+07 | 6.251E+05 | 1.983E+06 | 3.863E+05 | 4.767E+05 | 5.567E+05 | 2.000E+06 | 1.254E+07 |
| 356 | 310.173 | 6.844 | -1 | N/A | N/A | 1.332E+06 | 1.949E+06 | 1.273E+06 | 1.484E+06 | 1.982E+06 | 2.065E+06 | 1.830E+06 | 8.888E+04 | 5.679E+05 | 1.352E+05 | 1.072E+05 | 6.825E+05 | 5.250E+05 | 5.107E+05 |
| 365 | 519.261 | 9.835 | -1 | N/A | N/A | 3.053E+06 | 2.665E+06 | 2.776E+06 | 3.137E+06 | 1.019E+07 | 5.870E+06 | 6.436E+06 | 2.578E+05 | 7.530E+05 | 1.634E+05 | 7.599E+05 | 5.597E+05 | 1.511E+06 | 1.556E+06 |
| 368 | 273.167 | 4.221 | -1 | N/A | N/A | 5.092E+06 | 7.378E+06 | 5.053E+06 | 5.530E+06 | 4.990E+06 | 5.349E+06 | 4.562E+06 | 1.018E+06 | 7.402E+05 | 3.509E+05 | 1.560E+05 | 2.526E+05 | 4.591E+05 | 3.471E+05 |
| 376 | 288.181 | 4.136 | -1 | N/A | N/A | 1.813E+06 | 5.415E+06 | 2.778E+06 | 3.478E+06 | 4.062E+06 | 5.188E+06 | 3.323E+06 | 3.597E+05 | 9.605E+05 | 5.839E+05 | 1.685E+05 | 5.804E+05 | 5.118E+05 | 8.105E+05 |
| 379 | 496.301 | 10.257 | -1 | N/A | N/A | 9.582E+05 | 1.267E+06 | 8.014E+05 | 8.605E+05 | 1.989E+06 | 1.800E+06 | 1.128E+06 | 6.157E+04 | 4.201E+05 | 4.438E+04 | 1.337E+05 | 1.900E+05 | 1.419E+05 | 2.056E+05 |
| 380 | 497.280 | 9.835 | -1 | N/A | N/A | 2.765E+06 | 2.214E+06 | 2.390E+06 | 2.645E+06 | 1.103E+07 | 9.165E+06 | 1.115E+07 | 1.350E+05 | 6.995E+05 | 1.363E+05 | 8.420E+05 | 8.335E+05 | 3.199E+06 | 3.544E+06 |
| 385 | 188.069 | 0.519 | -1 | N/A | N/A | 1.171E+07 | 2.882E+07 | 1.438E+06 | 5.796E+04 | 1.211E+07 | 1.402E+07 | 4.871E+05 | 1.158E+06 | 9.924E+06 | 6.619E+05 | 2.075E+04 | 1.384E+06 | 1.288E+07 | 3.395E+05 |
| 395 | 721.506 | 12.424 | -1 | N/A | N/A | 2.538E+06 | 3.068E+06 | 2.387E+06 | 2.673E+06 | 6.277E+06 | 7.505E+06 | 6.726E+06 | 4.475E+04 | 2.728E+05 | 1.100E+05 | 2.426E+05 | 5.435E+05 | 1.461E+06 | 1.608E+06 |
| 397 | 459.263 | 10.315 | -1 | N/A | N/A | 6.012E+05 | 5.330E+05 | 6.462E+05 | 6.592E+05 | 7.136E+05 | 9.687E+05 | 8.757E+05 | 3.683E+04 | 1.113E+05 | 7.530E+04 | 7.310E+04 | 8.954E+04 | 1.294E+05 | 2.962E+05 |
| 398 | 445.291 | 10.157 | -1 | N/A | N/A | 7.228E+05 | 1.342E+06 | 2.543E+06 | 2.010E+06 | 7.918E+06 | 1.937E+05 | 8.978E+04 | 1.411E+05 | 1.405E+05 | 1.480E+06 | 1.039E+06 | 5.642E+06 | 8.834E+04 | 7.749E+04 |
| 403 | 763.517 | 12.425 | -1 | N/A | N/A | 2.650E+06 | 2.770E+06 | 2.205E+06 | 2.341E+06 | 5.905E+06 | 7.161E+06 | 6.634E+06 | 1.128E+05 | 2.538E+05 | 5.905E+04 | 1.981E+05 | 5.356E+05 | 1.526E+06 | 1.772E+06 |
| 405 | 528.788 | 4.417 | -1 | N/A | N/A | 0.000E+00 | 0.000E+00 | 1.106E+06 | 2.511E+05 | 1.721E+04 | 0.000E+00 | 0.000E+00 | 0.000E+00 | 0.000E+00 | 2.392E+05 | 2.511E+05 | 1.721E+04 | 0.000E+00 | 0.000E+00 |
| 407 | 455.335 | 10.125 | -1 | N/A | N/A | 6.934E+05 | 5.671E+05 | 1.030E+06 | 6.954E+05 | 5.682E+05 | 2.503E+06 | 2.450E+06 | 1.752E+04 | 1.605E+05 | 2.111E+05 | 1.408E+05 | 7.362E+04 | 9.048E+05 | 8.888E+05 |
| 410 | 830.949 | 3.185 | -1 | N/A | N/A | 3.103E+05 | 3.007E+06 | 1.844E+06 | 2.714E+06 | 3.138E+06 | 4.735E+06 | 2.393E+06 | 8.516E+04 | 4.903E+05 | 1.815E+05 | 3.098E+05 | 1.111E+06 | 1.091E+06 | 7.467E+05 |
| 421 | 514.951 | 4.135 | -1 | N/A | N/A | 0.000E+00 | 0.000E+00 | 1.062E+06 | 4.291E+05 | 0.000E+00 | 0.000E+00 | 0.000E+00 | 0.000E+00 | 0.000E+00 | 2.624E+05 | 4.250E+05 | 0.000E+00 | 0.000E+00 | 0.000E+00 |
| 423 | 409.180 | 7.399 | -1 | N/A | N/A | 2.591E+05 | 4.025E+06 | 9.660E+05 | 1.219E+06 | 2.380E+06 | 4.570E+06 | 2.411E+06 | 4.299E+04 | 1.940E+06 | 1.622E+05 | 6.575E+05 | 8.814E+05 | 2.701E+06 | 8.316E+05 |
| 425 | 707.607 | 12.056 | -1 | N/A | N/A | 2.190E+06 | 1.928E+06 | 1.482E+06 | 1.297E+06 | 1.054E+06 | 1.128E+06 | 8.563E+05 | 1.798E+05 | 9.327E+04 | 5.022E+04 | 5.585E+04 | 9.626E+04 | 2.887E+04 | 1.323E+05 |
| 428 | 366.633 | 3.310 | -1 | N/A | N/A | 4.713E+03 | 2.632E+04 | 2.185E+06 | 8.875E+06 | 7.843E+03 | 1.865E+04 | 0.000E+00 | 2.360E+03 | 4.479E+03 | 9.170E+05 | 8.872E+06 | 7.843E+03 | 7.174E+03 | 0.000E+00 |
| 430 | 379.862 | 4.127 | -1 | N/A | N/A | 0.000E+00 | 1.311E+05 | 8.957E+05 | 8.953E+05 | 0.000E+00 | 0.000E+00 | 0.000E+00 | 0.000E+00 | 9.770E+04 | 2.911E+05 | 8.953E+05 | 0.000E+00 | 0.000E+00 | 0.000E+00 |
| 436 | 267.137 | 8.329 | -1 | 4.991E+05 | 0.070 | 4.202E+05 | 8.453E+05 | 1.160E+06 | 1.815E+06 | 3.227E+06 | 2.430E+06 | 3.771E+06 | 2.431E+04 | 3.127E+05 | 9.530E+04 | 8.053E+05 | 5.612E+05 | 2.258E+05 | 1.032E+06 |
| 449 | 658.341 | 3.958 | -1 | N/A | N/A | 0.000E+00 | 0.000E+00 | 1.533E+06 | 5.265E+05 | 0.000E+00 | 0.000E+00 | 0.000E+00 | 0.000E+00 | 0.000E+00 | 2.784E+05 | 5.265E+05 | 0.000E+00 | 0.000E+00 | 0.000E+00 |
| 451 | 254.660 | 3.065 | -1 | N/A | N/A | 0.000E+00 | 0.000E+00 | 2.558E+06 | 2.769E+06 | 0.000E+00 | 0.000E+00 | 0.000E+00 | 0.000E+00 | 0.000E+00 | 2.956E+05 | 2.769E+06 | 0.000E+00 | 0.000E+00 | 0.000E+00 |
| 453 | 441.346 | 7.492 | -1 | N/A | N/A | 5.734E+05 | 5.784E+05 | 6.120E+05 | 6.094E+05 | 5.956E+05 | 5.308E+05 | 4.252E+05 | 4.049E+04 | 2.361E+04 | 4.576E+04 | 1.067E+04 | 1.066E+05 | 5.673E+04 | 1.269E+05 |
| 454 | 409.163 | 6.914 | -1 | N/A | N/A | 1.548E+06 | 2.176E+07 | 1.624E+06 | 6.751E+05 | 7.475E+05 | 1.601E+07 | 2.646E+05 | 1.647E+05 | 1.155E+07 | 4.554E+05 | 1.363E+05 | 1.943E+05 | 1.518E+07 | 1.612E+04 |
| 470 | 469.163 | 6.021 | -1 | N/A | N/A | 5.305E+05 | 1.109E+06 | 6.028E+05 | 8.429E+05 | 9.640E+05 | 9.864E+05 | 2.196E+06 | 5.556E+04 | 8.585E+04 | 7.687E+04 | 6.033E+04 | 1.143E+05 | 1.233E+05 | 1.692E+06 |
| 474 | 501.170 | 0.519 | -1 | N/A | N/A | 4.020E+04 | 0.000E+00 | 0.000E+00 | 3.797E+08 | 8.528E+05 | 5.735E+04 | 1.985E+05 | 4.020E+04 | 0.000E+00 | 0.000E+00 | 5.832E+07 | 6.310E+05 | 3.269E+04 | 1.034E+05 |
| 485 | 517.142 | 0.507 | -1 | N/A | N/A | 0.000E+00 | 0.000E+00 | 0.000E+00 | 4.253E+07 | 7.026E+03 | 0.000E+00 | 0.000E+00 | 0.000E+00 | 0.000E+00 | 0.000E+00 | 3.058E+06 | 7.026E+03 | 0.000E+00 | 0.000E+00 |
| 496 | 226.012 | 12.739 | -1 | N/A | N/A | 1.296E+07 | 1.200E+07 | 1.315E+07 | 1.301E+07 | 1.333E+07 | 1.147E+07 | 1.013E+07 | 1.788E+06 | 1.684E+06 | 9.837E+05 | 4.440E+05 | 6.420E+05 | 4.511E+05 | 1.947E+05 |
| 502 | 102.034 | 0.568 | -1 | N/A | N/A | 5.869E+06 | 6.313E+06 | 7.768E+06 | 7.261E+06 | 6.911E+06 | 9.643E+06 | 5.836E+06 | 3.280E+05 | 4.402E+06 | 2.269E+06 | 2.471E+06 | 3.282E+06 | 5.793E+05 | 2.296E+06 |
| 503 | 228.207 | 4.644 | -1 | N/A | N/A | 5.935E+06 | 3.732E+06 | 5.229E+06 | 6.774E+06 | 4.623E+06 | 7.970E+06 | 8.389E+06 | 3.157E+06 | 5.759E+05 | 5.188E+05 | 1.502E+05 | 2.659E+05 | 7.398E+05 | 4.500E+05 |
| 508 | 1011.286 | 0.501 | -1 | N/A | N/A | 0.000E+00 | 0.000E+00 | 0.000E+00 | 2.831E+06 | 0.000E+00 | 0.000E+00 | 0.000E+00 | 0.000E+00 | 0.000E+00 | 0.000E+00 | 1.628E+06 | 0.000E+00 | 0.000E+00 | 0.000E+00 |
| 509 | 385.095 | 3.893 | -1 | N/A | N/A | 1.137E+05 | 7.871E+05 | 0.000E+00 | 1.512E+06 | 1.451E+06 | 1.798E+06 | 6.708E+05 | 4.247E+04 | 6.360E+05 | 0.000E+00 | 1.204E+06 | 4.740E+05 | 8.998E+05 | 6.385E+05 |

| Feature identifiers | | | | | | Average XIC | | | | | | | Standard Err of Average XIC | | | | | | |
|---|---|---|---|---|---|---|---|---|---|---|---|---|---|---|---|---|---|---|---|
| ClusterIdx | m/z | RT [min] | Subnetwork | PPM | MassDiff | TUB00 | 4TUB34 | 4TUB81 | 4TUB93 | 5TUB23 | 5TUB48 | 5TUB83 | TUB00 | 4TUB34 | 4TUB81 | 4TUB93 | 5TUB23 | 5TUB48 | 5TUB83 |
| 525 | 269.210 | 8.072 | -1 | N/A | N/A | 1.257E+05 | 1.861E+05 | 1.294E+06 | 3.563E+07 | 3.218E+06 | 5.996E+06 | 4.255E+06 | 5.015E+04 | 9.911E+04 | 9.599E+05 | 2.432E+07 | 1.528E+06 | 2.715E+06 | 1.275E+06 |
| 538 | 844.952 | 3.465 | -1 | N/A | N/A | 4.325E+05 | 1.690E+06 | 1.107E+06 | 1.538E+06 | 1.313E+06 | 3.090E+06 | 3.031E+06 | 7.923E+04 | 3.533E+05 | 1.885E+05 | 6.478E+05 | 2.685E+05 | 5.967E+05 | 5.632E+05 |
| 543 | 453.188 | 6.313 | -1 | N/A | N/A | 6.177E+05 | 1.433E+06 | 9.967E+05 | 1.042E+06 | 9.867E+05 | 1.348E+06 | 6.700E+06 | 1.598E+05 | 5.427E+04 | 4.344E+04 | 9.109E+04 | 1.089E+05 | 1.867E+05 | 6.088E+06 |
| 547 | 372.717 | 12.740 | -1 | N/A | N/A | 0.000E+00 | 0.000E+00 | 8.167E+05 | 1.215E+05 | 4.133E+05 | 9.288E+05 | 5.506E+06 | 0.000E+00 | 0.000E+00 | 4.100E+05 | 1.215E+05 | 2.921E+05 | 5.155E+05 | 3.631E+06 |
| 549 | 287.114 | 4.597 | -1 | N/A | N/A | 2.048E+05 | 1.568E+06 | 5.228E+05 | 1.505E+06 | 1.114E+06 | 1.758E+06 | 1.168E+06 | 6.209E+04 | 4.545E+05 | 5.779E+04 | 5.107E+05 | 1.155E+05 | 2.052E+05 | 3.607E+05 |
| 551 | 385.294 | 9.364 | -1 | N/A | N/A | 7.341E+05 | 8.374E+05 | 9.985E+05 | 3.717E+06 | 1.242E+06 | 9.773E+05 | 9.698E+05 | 3.431E+04 | 4.116E+04 | 1.256E+05 | 1.351E+06 | 1.994E+05 | 6.050E+04 | 2.351E+04 |
| 557 | 858.957 | 3.760 | -1 | N/A | N/A | 1.177E+06 | 4.428E+05 | 3.702E+05 | 1.252E+06 | 6.304E+05 | 1.356E+06 | 4.176E+06 | 4.771E+05 | 3.481E+04 | 1.056E+05 | 6.454E+05 | 3.092E+05 | 6.346E+05 | 5.653E+05 |
| 563 | 429.319 | 9.342 | -1 | N/A | N/A | 3.979E+05 | 7.359E+05 | 8.149E+05 | 3.128E+06 | 1.066E+06 | 9.076E+05 | 7.724E+05 | 5.549E+04 | 1.556E+05 | 1.458E+05 | 1.107E+06 | 2.565E+05 | 7.446E+04 | 3.717E+04 |
| 566 | 472.175 | 4.688 | -1 | N/A | N/A | 8.384E+05 | 9.439E+05 | 3.430E+04 | 1.033E+06 | 1.161E+06 | 1.664E+06 | 7.380E+05 | 1.713E+05 | 2.619E+05 | 3.430E+04 | 6.499E+05 | 2.722E+05 | 7.204E+05 | 4.603E+05 |
| 569 | 294.074 | 0.519 | -1 | N/A | N/A | 6.414E+03 | 5.976E+06 | 1.447E+05 | 2.728E+06 | 4.459E+06 | 8.898E+06 | 8.218E+06 | 6.414E+03 | 3.102E+06 | 6.580E+03 | 1.212E+06 | 7.796E+05 | 1.497E+06 | 2.593E+06 |
| 575 | 453.189 | 6.022 | -1 | N/A | N/A | 3.642E+05 | 1.010E+06 | 4.350E+05 | 6.060E+05 | 7.790E+05 | 7.534E+05 | 1.931E+06 | 1.005E+05 | 3.455E+03 | 5.014E+04 | 7.015E+04 | 1.168E+05 | 8.734E+04 | 1.527E+06 |
| 586 | 191.086 | 3.681 | -1 | N/A | N/A | 5.795E+04 | 2.307E+06 | 1.744E+06 | 1.442E+07 | 2.059E+04 | 1.510E+04 | 4.857E+03 | 2.684E+04 | 4.669E+05 | 2.374E+04 | 7.983E+06 | 1.794E+04 | 7.611E+03 | 4.857E+03 |
| 592 | 359.104 | 3.680 | -1 | N/A | N/A | 8.567E+03 | 1.529E+06 | 1.192E+06 | 7.226E+06 | 0.000E+00 | 1.367E+03 | 0.000E+00 | 5.446E+03 | 3.312E+05 | 9.584E+04 | 3.133E+06 | 0.000E+00 | 1.367E+03 | 0.000E+00 |
| 593 | 511.165 | 3.612 | -1 | N/A | N/A | 0.000E+00 | 2.514E+06 | 8.776E+05 | 4.042E+06 | 5.592E+05 | 2.281E+06 | 1.946E+06 | 0.000E+00 | 1.037E+06 | 2.025E+05 | 2.074E+06 | 2.480E+05 | 5.228E+05 | 1.596E+06 |
| 594 | 700.315 | 10.020 | -1 | N/A | N/A | 4.424E+04 | 1.410E+06 | 3.544E+05 | 1.020E+06 | 1.246E+06 | 2.990E+06 | 2.541E+06 | 2.675E+04 | 8.055E+05 | 8.752E+04 | 8.530E+05 | 5.671E+05 | 1.002E+06 | 9.684E+05 |
| 597 | 225.149 | 6.492 | -1 | N/A | N/A | 2.217E+06 | 1.036E+07 | 7.884E+05 | 3.051E+06 | 7.035E+05 | 3.970E+06 | 1.765E+06 | 8.270E+05 | 3.283E+06 | 3.984E+05 | 1.319E+06 | 4.222E+05 | 2.289E+06 | 1.161E+06 |
| 599 | 131.049 | 0.515 | -1 | N/A | N/A | 5.797E+06 | 1.120E+07 | 9.388E+05 | 4.304E+06 | 1.327E+07 | 1.592E+07 | 4.089E+05 | 2.481E+06 | 8.465E+06 | 5.632E+05 | 2.321E+06 | 1.522E+06 | 1.449E+07 | 7.390E+04 |
| 601 | 357.262 | 8.098 | -1 | N/A | N/A | 1.768E+03 | 2.025E+04 | 1.293E+05 | 8.505E+06 | 7.417E+05 | 5.831E+06 | 2.883E+06 | 1.768E+03 | 3.086E+03 | 9.107E+04 | 4.582E+06 | 3.016E+05 | 2.817E+06 | 9.297E+05 |
| 602 | 303.138 | 3.688 | -1 | N/A | N/A | 3.834E+03 | 8.225E+05 | 4.905E+05 | 5.090E+06 | 1.936E+03 | 0.000E+00 | 5.180E+03 | 3.834E+03 | 1.216E+05 | 8.787E+04 | 2.720E+06 | 1.936E+03 | 0.000E+00 | 5.180E+03 |
| 605 | 347.222 | 7.583 | -1 | N/A | N/A | 1.866E+06 | 3.817E+06 | 3.192E+06 | 3.283E+06 | 3.081E+06 | 4.539E+06 | 3.254E+06 | 4.598E+05 | 7.842E+05 | 5.226E+05 | 6.048E+05 | 6.342E+05 | 3.721E+05 | 9.175E+05 |
| 606 | 438.143 | 7.111 | -1 | 7.173E+04 | 33.858 | 0.000E+00 | 5.450E+03 | 1.817E+06 | 2.112E+06 | 0.000E+00 | 0.000E+00 | 0.000E+00 | 0.000E+00 | 5.450E+03 | 1.311E+06 | 1.038E+06 | 0.000E+00 | 0.000E+00 | 0.000E+00 |
| 607 | 851.398 | 7.250 | -1 | N/A | N/A | 1.294E+04 | 2.313E+05 | 3.322E+05 | 6.578E+05 | 5.227E+05 | 4.141E+05 | 4.019E+05 | 6.622E+03 | 1.825E+05 | 4.082E+04 | 4.993E+05 | 2.087E+05 | 2.771E+05 | 2.270E+05 |
| 609 | 830.614 | 3.188 | -1 | N/A | N/A | 2.823E+05 | 2.468E+06 | 1.259E+06 | 2.057E+06 | 1.274E+06 | 4.127E+06 | 2.678E+06 | 1.143E+05 | 7.854E+05 | 3.197E+05 | 6.065E+05 | 8.362E+04 | 1.819E+06 | 8.954E+05 |
| 612 | 863.046 | 3.487 | -1 | N/A | N/A | 5.913E+05 | 2.373E+07 | 3.855E+04 | 6.633E+06 | 5.970E+05 | 3.805E+06 | 7.630E+06 | 2.499E+05 | 6.421E+06 | 3.296E+03 | 6.540E+06 | 2.821E+05 | 3.500E+06 | 3.774E+06 |
| 613 | 619.272 | 4.460 | -1 | N/A | N/A | 1.329E+05 | 5.806E+05 | 7.240E+05 | 8.727E+05 | 1.846E+06 | 2.494E+06 | 6.744E+06 | 3.996E+04 | 1.515E+05 | 1.092E+05 | 3.887E+05 | 5.078E+05 | 6.121E+05 | 4.903E+06 |
| 617 | 454.172 | 7.812 | -1 | N/A | N/A | 1.727E+05 | 1.499E+06 | 5.170E+05 | 8.433E+05 | 7.985E+05 | 1.171E+06 | 8.895E+05 | 2.077E+04 | 2.738E+05 | 1.234E+05 | 4.639E+05 | 7.270E+04 | 2.252E+05 | 3.203E+05 |
| 621 | 815.644 | 3.349 | -1 | N/A | N/A | 0.000E+00 | 3.899E+04 | 4.906E+04 | 4.672E+05 | 2.282E+06 | 1.181E+06 | 2.202E+05 | 0.000E+00 | 2.183E+04 | 3.364E+03 | 3.278E+05 | 2.230E+06 | 3.902E+05 | 1.738E+05 |
| 626 | 331.209 | 4.774 | -1 | N/A | N/A | 2.104E+06 | 3.148E+06 | 2.026E+06 | 1.747E+06 | 2.485E+06 | 2.138E+06 | 1.206E+06 | 2.418E+05 | 1.746E+05 | 1.415E+05 | 3.059E+05 | 2.204E+05 | 1.708E+05 | 2.178E+05 |
| 628 | 103.054 | 0.515 | -1 | N/A | N/A | 2.258E+06 | 8.707E+06 | 1.243E+06 | 6.312E+05 | 6.621E+06 | 1.086E+07 | 3.044E+05 | 1.184E+06 | 4.044E+06 | 3.776E+05 | 2.945E+05 | 3.311E+06 | 1.017E+07 | 1.229E+05 |
| 641 | 158.154 | 6.250 | -1 | N/A | N/A | 1.719E+07 | 4.370E+07 | 5.660E+07 | 4.051E+07 | 4.882E+07 | 3.174E+07 | 7.096E+07 | 4.858E+06 | 2.368E+07 | 1.324E+07 | 3.567E+06 | 1.841E+07 | 7.617E+06 | 1.200E+07 |
| 642 | 165.114 | 12.714 | -1 | 5.001E+05 | 55.042 | 3.314E+07 | 3.025E+07 | 2.285E+07 | 2.465E+07 | 1.774E+07 | 1.708E+07 | 1.418E+07 | 3.502E+06 | 8.298E+06 | 7.045E+06 | 7.973E+06 | 1.021E+07 | 1.682E+06 | 2.710E+06 |
| 648 | 425.137 | 6.913 | -1 | N/A | N/A | 1.019E+06 | 1.425E+07 | 1.186E+06 | 4.312E+05 | 5.170E+05 | 9.972E+05 | 1.517E+05 | 9.091E+04 | 7.521E+06 | 2.450E+05 | 6.656E+04 | 1.255E+05 | 9.367E+04 | 1.417E+04 |
| 651 | 267.157 | 4.332 | -1 | N/A | N/A | 3.668E+06 | 1.151E+07 | 1.758E+06 | 3.218E+06 | 3.511E+06 | 2.902E+06 | 2.637E+06 | 1.209E+06 | 2.859E+06 | 9.693E+05 | 2.219E+06 | 4.261E+05 | 9.858E+05 | 1.089E+06 |
| 655 | 290.232 | 4.304 | -1 | N/A | N/A | 1.138E+06 | 6.610E+06 | 1.067E+06 | 1.999E+06 | 1.380E+06 | 9.229E+05 | 5.091E+05 | 5.826E+05 | 1.525E+06 | 5.033E+05 | 1.508E+06 | 5.124E+05 | 2.395E+05 | 9.879E+04 |
| 660 | 391.205 | 10.042 | -1 | N/A | N/A | 4.098E+05 | 8.198E+05 | 1.606E+06 | 8.717E+05 | 6.281E+06 | 1.021E+06 | 7.650E+05 | 7.422E+04 | 7.936E+04 | 5.378E+05 | 6.293E+05 | 2.392E+06 | 3.158E+05 | 6.584E+05 |
| 673 | 870.956 | 4.058 | -1 | N/A | N/A | 6.849E+06 | 2.356E+05 | 2.397E+04 | 2.653E+05 | 2.102E+05 | 4.662E+05 | 2.817E+06 | 1.205E+06 | 1.046E+05 | 1.302E+04 | 1.328E+05 | 1.199E+05 | 4.161E+05 | 1.002E+06 |
| 682 | 425.215 | 9.512 | -1 | N/A | N/A | 2.037E+06 | 9.777E+05 | 9.531E+05 | 1.035E+06 | 9.919E+05 | 1.611E+06 | 1.787E+06 | 6.793E+05 | 1.186E+05 | 6.347E+04 | 1.130E+05 | 1.158E+05 | 2.621E+05 | 2.410E+05 |
| 684 | 612.425 | 0.524 | -1 | N/A | N/A | 1.850E+06 | 6.410E+06 | 3.255E+05 | 2.423E+05 | 3.582E+06 | 3.492E+06 | 2.064E+05 | 8.196E+05 | 1.807E+06 | 7.935E+04 | 7.020E+04 | 3.460E+05 | 3.439E+06 | 1.679E+05 |
| 686 | 231.081 | 0.527 | -1 | 2.092E+05 | 39.978 | 2.683E+06 | 4.820E+06 | 4.227E+03 | 3.934E+04 | 2.179E+07 | 3.450E+05 | 9.344E+03 | 2.432E+05 | 2.159E+06 | 2.133E+03 | 3.934E+04 | 1.353E+07 | 3.398E+05 | 4.014E+03 |
| 688 | 726.463 | 12.425 | -1 | N/A | N/A | 1.597E+06 | 1.351E+06 | 1.006E+06 | 9.114E+05 | 3.623E+06 | 4.027E+06 | 3.540E+06 | 1.757E+05 | 3.306E+05 | 9.675E+04 | 2.327E+05 | 4.471E+05 | 1.129E+06 | 1.414E+06 |
| 690 | 403.233 | 9.514 | -1 | N/A | N/A | 7.186E+05 | 3.848E+05 | 3.897E+05 | 4.056E+05 | 3.685E+05 | 5.560E+05 | 6.187E+05 | 2.194E+05 | 9.803E+04 | 3.152E+04 | 2.701E+04 | 5.457E+04 | 1.638E+05 | 1.264E+05 |
| 694 | 111.020 | 12.539 | -1 | N/A | N/A | 3.545E+05 | 2.724E+05 | 1.989E+05 | 1.860E+05 | 2.409E+05 | 1.345E+05 | 3.316E+05 | 1.212E+05 | 2.997E+04 | 1.662E+04 | 6.489E+04 | 4.863E+04 | 1.437E+04 | 7.220E+04 |
| 695 | 351.102 | 8.964 | -1 | N/A | N/A | 5.905E+05 | 1.378E+06 | 3.967E+05 | 4.936E+05 | 2.098E+06 | 5.857E+05 | 3.934E+05 | 1.028E+05 | 4.753E+05 | 6.056E+04 | 3.738E+04 | 1.427E+06 | 1.604E+05 | 1.085E+05 |
| 700 | 164.051 | 4.799 | -1 | 1.232E+05 | 17.991 | 1.820E+03 | 0.000E+00 | 1.088E+05 | 1.930E+04 | 2.669E+07 | 1.321E+07 | 3.037E+07 | 1.820E+03 | 0.000E+00 | 6.180E+04 | 1.930E+04 | 4.406E+06 | 5.907E+06 | 4.925E+06 |
| 702 | 111.039 | 2.302 | -1 | N/A | N/A | 1.904E+06 | 2.019E+06 | 2.470E+06 | 1.331E+06 | 3.164E+06 | 4.960E+06 | 2.328E+06 | 7.655E+05 | 6.573E+05 | 9.232E+05 | 9.777E+05 | 1.073E+06 | 2.138E+06 | 2.197E+04 |
| 705 | 271.031 | 5.195 | -1 | N/A | N/A | 0.000E+00 | 1.478E+03 | 0.000E+00 | 0.000E+00 | 5.648E+06 | 3.890E+03 | 8.098E+04 | 0.000E+00 | 1.478E+03 | 0.000E+00 | 0.000E+00 | 2.021E+06 | 1.257E+03 | 5.356E+04 |
| 706 | 249.050 | 5.192 | -1 | N/A | N/A | 0.000E+00 | 3.007E+03 | 0.000E+00 | 0.000E+00 | 5.569E+06 | 2.384E+03 | 7.406E+04 | 0.000E+00 | 1.541E+03 | 0.000E+00 | 0.000E+00 | 2.008E+06 | 1.370E+03 | 5.906E+04 |
| 707 | 279.060 | 6.671 | -1 | N/A | N/A | 0.000E+00 | 1.206E+03 | 0.000E+00 | 0.000E+00 | 9.781E+06 | 1.781E+04 | 5.471E+05 | 0.000E+00 | 1.206E+03 | 0.000E+00 | 0.000E+00 | 1.602E+06 | 9.305E+03 | 2.431E+05 |
| 708 | 162.035 | 6.813 | -1 | N/A | N/A | 0.000E+00 | 0.000E+00 | 0.000E+00 | 0.000E+00 | 1.287E+07 | 7.126E+04 | 6.738E+05 | 0.000E+00 | 0.000E+00 | 0.000E+00 | 0.000E+00 | 3.813E+06 | 3.919E+04 | 2.850E+05 |
| 709 | 177.083 | 2.375 | -1 | N/A | N/A | 6.892E+03 | 4.358E+05 | 3.334E+04 | 3.701E+05 | 2.246E+07 | 3.532E+07 | 2.898E+03 | 2.368E+03 | 2.095E+05 | 1.599E+04 | 1.507E+05 | 1.840E+07 | 1.803E+07 | 1.652E+03 |
| 711 | 164.051 | 2.397 | -1 | N/A | N/A | 5.265E+02 | 4.705E+05 | 2.820E+04 | 2.043E+05 | 2.313E+07 | 3.796E+07 | 1.429E+04 | 5.265E+02 | 1.592E+05 | 4.141E+03 | 3.188E+04 | 1.872E+07 | 1.721E+07 | 6.654E+03 |
| 712 | 245.070 | 2.402 | -1 | N/A | N/A | 0.000E+00 | 2.822E+05 | 4.176E+04 | 2.636E+05 | 1.981E+07 | 3.154E+07 | 1.033E+03 | 0.000E+00 | 6.645E+04 | 3.143E+04 | 1.138E+05 | 1.624E+07 | 1.373E+07 | 1.033E+03 |
| 716 | 111.039 | 2.354 | -1 | N/A | N/A | 1.560E+06 | 3.202E+06 | 2.079E+06 | 1.752E+06 | 2.571E+06 | 4.873E+06 | 1.757E+06 | 4.977E+05 | 4.317E+05 | 2.772E+05 | 1.217E+06 | 8.858E+05 | 3.917E+06 | 5.825E+05 |
| 720 | 495.251 | 4.182 | -1 | N/A | N/A | 0.000E+00 | 0.000E+00 | 0.000E+00 | 0.000E+00 | 3.104E+06 | 1.984E+06 | 1.318E+06 | 0.000E+00 | 0.000E+00 | 0.000E+00 | 0.000E+00 | 8.341E+05 | 3.104E+05 | 2.345E+05 |
| 723 | 162.073 | 2.320 | -1 | N/A | N/A | 5.765E+03 | 1.650E+04 | 9.275E+03 | 1.747E+04 | 7.861E+06 | 8.716E+06 | 6.781E+03 | 1.646E+03 | 4.996E+03 | 1.314E+03 | 7.781E+03 | 7.657E+06 | 7.433E+06 | 1.752E+03 |
| 725 | 1347.881 | 12.426 | -1 | N/A | N/A | 7.616E+04 | 2.243E+05 | 8.573E+04 | 1.205E+05 | 2.748E+06 | 3.871E+06 | 3.091E+06 | 1.180E+04 | 1.231E+05 | 2.351E+04 | 5.796E+03 | 6.061E+05 | 2.178E+06 | 2.175E+06 |
| 733 | 297.083 | 6.817 | -1 | N/A | N/A | 0.000E+00 | 3.319E+03 | 0.000E+00 | 0.000E+00 | 1.930E+06 | 1.050E+07 | 4.333E+07 | 0.000E+00 | 3.319E+03 | 0.000E+00 | 0.000E+00 | 4.582E+05 | 2.817E+06 | 1.143E+07 |
| 734 | 471.252 | 7.681 | -1 | N/A | N/A | 0.000E+00 | 3.221E+05 | 1.366E+05 | 5.987E+05 | 1.270E+06 | 7.822E+05 | 1.915E+06 | 0.000E+00 | 1.457E+05 | 1.234E+04 | 4.085E+05 | 4.566E+05 | 9.248E+04 | 5.401E+05 |
| 735 | 597.678 | 0.494 | -1 | N/A | N/A | 1.454E+05 | 2.960E+05 | 0.000E+00 | 9.697E+04 | 5.021E+06 | 1.107E+07 | 2.843E+07 | 2.232E+04 | 1.568E+05 | 0.000E+00 | 5.019E+04 | 2.738E+06 | 4.247E+06 | 1.066E+07 |
| 737 | 701.411 | 12.427 | -1 | N/A | N/A | 6.547E+05 | 1.256E+06 | 7.315E+05 | 8.459E+05 | 3.228E+06 | 3.541E+06 | 3.146E+06 | 1.546E+05 | 2.540E+05 | 2.091E+05 | 8.388E+04 | 4.426E+05 | 8.791E+05 | 1.036E+06 |
| 738 | 253.602 | 7.678 | -1 | N/A | N/A | 3.667E+04 | 7.197E+05 | 4.903E+05 | 8.670E+05 | 1.427E+06 | 1.208E+06 | 1.772E+06 | 4.632E+03 | 2.322E+05 | 4.918E+04 | 3.942E+05 | 3.605E+05 | 1.602E+05 | 3.261E+05 |
| 739 | 749.548 | 12.237 | -1 | N/A | N/A | 3.731E+03 | 2.020E+03 | 1.268E+04 | 1.499E+04 | 2.131E+06 | 8.710E+06 | 2.344E+06 | 3.731E+03 | 2.020E+03 | 8.111E+03 | 1.099E+04 | 1.960E+06 | 5.497E+06 | 1.756E+06 |
| 742 | 287.650 | 8.328 | -1 | N/A | N/A | 1.632E+05 | 2.981E+05 | 2.353E+05 | 5.743E+05 | 1.321E+06 | 8.810E+05 | 1.215E+06 | 4.393E+04 | 1.054E+05 | 4.432E+04 | 3.395E+05 | 4.039E+05 | 5.000E+04 | 2.886E+05 |
| 744 | 247.129 | 0.492 | -1 | 1.279E+05 | 28.031 | 5.759E+04 | 1.171E+04 | 1.654E+03 | 2.349E+04 | 3.671E+05 | 1.377E+07 | 1.816E+06 | 1.739E+04 | 1.404E+03 | 8.317E+02 | 1.288E+04 | 2.954E+05 | 8.449E+06 | 1.805E+06 |
| 748 | 654.456 | 10.203 | -1 | N/A | N/A | 0.000E+00 | 7.084E+03 | 4.481E+03 | 2.255E+03 | 9.796E+05 | 2.118E+06 | 1.721E+06 | 0.000E+00 | 3.787E+03 | 4.481E+03 | 2.255E+03 | 3.250E+05 | 7.280E+05 | 9.086E+05 |
| 752 | 286.520 | 6.796 | -1 | N/A | N/A | 0.000E+00 | 0.000E+00 | 0.000E+00 | 0.000E+00 | 2.001E+06 | 0.000E+00 | 0.000E+00 | 0.000E+00 | 0.000E+00 | 0.000E+00 | 0.000E+00 | 1.094E+06 | 0.000E+00 | 0.000E+00 |
| 753 | 335.125 | 8.963 | -1 | N/A | N/A | 7.624E+05 | 1.648E+06 | 5.024E+05 | 5.562E+05 | 2.805E+06 | 5.522E+05 | 4.803E+05 | 1.403E+05 | 5.652E+05 | 8.116E+04 | 8.464E+04 | 1.979E+06 | 2.654E+05 | 1.441E+05 |
| 754 | 533.934 | 3.609 | -1 | N/A | N/A | 0.000E+00 | 0.000E+00 | 0.000E+00 | 0.000E+00 | 2.373E+06 | 1.435E+06 | 3.015E+06 | 0.000E+00 | 0.000E+00 | 0.000E+00 | 0.000E+00 | 9.198E+05 | 5.670E+05 | 7.644E+05 |
| 762 | 533.282 | 9.675 | -1 | N/A | N/A | 1.779E+05 | 1.418E+06 | 3.501E+05 | 4.159E+05 | 3.592E+05 | 3.204E+05 | 1.162E+05 | 4.892E+04 | 6.275E+05 | 9.643E+04 | 3.857E+04 | 1.156E+05 | 5.527E+04 | 1.096E+04 |
| 763 | 1194.349 | 0.496 | -1 | N/A | N/A | 2.518E+04 | 4.011E+04 | 0.000E+00 | 3.397E+04 | 1.724E+06 | 4.227E+06 | 1.152E+07 | 1.286E+04 | 2.294E+04 | 0.000E+00 | 2.120E+04 | 1.379E+06 | 2.329E+06 | 4.185E+06 |

| Feature identifiers | | | | | | Average XIC | | | | | | | Standard Err of Average XIC | | | | | | |
|---|---|---|---|---|---|---|---|---|---|---|---|---|---|---|---|---|---|---|---|
| ClusterIdx | m/z | RT [min] | Subnetwork | PPM | MassDiff | TUB00 | 4TUB34 | 4TUB81 | 4TUB93 | 5TUB23 | 5TUB48 | 5TUB83 | TUB00 | 4TUB34 | 4TUB81 | 4TUB93 | 5TUB23 | 5TUB48 | 5TUB83 |
| 764 | 616.651 | 0.503 | -1 | N/A | N/A | 1.816E+04 | 1.770E+04 | 0.000E+00 | 0.000E+00 | 1.057E+06 | 3.045E+06 | 7.905E+06 | 1.744E+04 | 7.030E+03 | 0.000E+00 | 0.000E+00 | 7.719E+05 | 1.398E+06 | 2.853E+06 |
| 765 | 223.064 | 6.361 | -1 | N/A | N/A | 1.254E+06 | 6.488E+06 | 5.362E+05 | 1.679E+06 | 1.581E+06 | 9.234E+05 | 7.024E+05 | 2.750E+05 | 1.504E+06 | 1.782E+05 | 1.045E+06 | 4.469E+05 | 1.790E+05 | 2.281E+05 |
| 769 | 498.173 | 7.669 | -1 | N/A | N/A | 3.597E+05 | 7.207E+05 | 4.546E+05 | 5.333E+05 | 7.490E+05 | 1.144E+06 | 1.909E+06 | 7.060E+04 | 2.888E+05 | 7.541E+04 | 9.620E+04 | 1.488E+05 | 2.810E+05 | 1.568E+06 |
| 771 | 143.107 | 1.088 | -1 | N/A | N/A | 2.810E+06 | 2.702E+07 | 1.580E+05 | 6.849E+06 | 9.256E+05 | 2.329E+06 | 1.806E+06 | 2.344E+06 | 1.005E+07 | 4.806E+04 | 6.507E+06 | 4.222E+05 | 2.217E+06 | 1.694E+06 |
| 786 | 819.476 | 8.754 | -1 | 2.994E+01 | 0.025 | 1.245E+05 | 4.135E+05 | 1.083E+05 | 1.487E+06 | 1.230E+05 | 1.995E+05 | 7.168E+04 | 1.948E+04 | 1.338E+05 | 3.071E+03 | 1.377E+06 | 4.331E+04 | 6.619E+04 | 2.332E+04 |
| 794 | 811.308 | 6.909 | -1 | N/A | N/A | 0.000E+00 | 8.101E+05 | 0.000E+00 | 0.000E+00 | 0.000E+00 | 9.259E+05 | 0.000E+00 | 0.000E+00 | 4.919E+05 | 0.000E+00 | 0.000E+00 | 0.000E+00 | 9.259E+05 | 0.000E+00 |
| 797 | 818.299 | 3.374 | -1 | N/A | N/A | 2.323E+03 | 4.425E+05 | 1.131E+03 | 2.476E+04 | 4.339E+05 | 1.576E+06 | 1.372E+06 | 2.323E+03 | 1.086E+05 | 1.131E+03 | 1.288E+04 | 2.686E+05 | 1.164E+06 | 7.956E+05 |
| 806 | 618.984 | 3.690 | -1 | N/A | N/A | 7.667E+03 | 4.937E+05 | 4.778E+03 | 3.997E+04 | 5.673E+05 | 1.932E+06 | 3.109E+06 | 7.667E+03 | 1.792E+05 | 4.778E+03 | 2.385E+04 | 5.147E+05 | 1.738E+06 | 1.402E+06 |
| 808 | 832.302 | 3.693 | -1 | N/A | N/A | 0.000E+00 | 3.760E+05 | 0.000E+00 | 2.268E+04 | 4.394E+05 | 1.232E+06 | 2.055E+06 | 0.000E+00 | 1.454E+05 | 0.000E+00 | 1.178E+04 | 4.117E+05 | 1.103E+06 | 8.658E+05 |
| 814 | 513.189 | 7.230 | -1 | N/A | N/A | 4.998E+05 | 1.156E+06 | 5.271E+05 | 5.928E+05 | 9.877E+05 | 7.917E+05 | 2.492E+06 | 1.186E+05 | 1.741E+05 | 5.851E+04 | 2.170E+05 | 1.263E+05 | 1.873E+05 | 2.034E+06 |
| 815 | 432.241 | 7.420 | -1 | N/A | N/A | 6.381E+03 | 2.542E+05 | 5.184E+04 | 1.622E+04 | 5.272E+04 | 9.493E+05 | 5.092E+05 | 3.363E+03 | 2.290E+05 | 1.948E+04 | 1.946E+03 | 2.124E+04 | 9.176E+05 | 4.898E+05 |
| 818 | 123.079 | 0.380 | -1 | N/A | N/A | 7.083E+07 | 1.181E+07 | 1.765E+07 | 2.009E+07 | 2.036E+07 | 2.016E+07 | 1.223E+08 | 6.278E+07 | 2.747E+06 | 5.413E+06 | 5.512E+06 | 4.261E+06 | 2.115E+06 | 1.049E+08 |
| 822 | 376.087 | 6.788 | -1 | N/A | N/A | 0.000E+00 | 0.000E+00 | 0.000E+00 | 0.000E+00 | 0.000E+00 | 1.875E+06 | 1.061E+07 | 0.000E+00 | 0.000E+00 | 0.000E+00 | 0.000E+00 | 0.000E+00 | 9.031E+05 | 3.352E+06 |
| 827 | 326.112 | 5.859 | -1 | N/A | N/A | 1.782E+04 | 1.365E+04 | 2.529E+03 | 1.881E+04 | 5.695E+05 | 2.504E+06 | 2.276E+06 | 9.182E+03 | 7.363E+03 | 2.529E+03 | 1.263E+04 | 8.120E+04 | 9.419E+05 | 3.171E+05 |
| 828 | 319.064 | 6.836 | -1 | N/A | N/A | 0.000E+00 | 0.000E+00 | 4.798E+04 | 9.691E+03 | 7.705E+04 | 1.404E+06 | 5.490E+06 | 0.000E+00 | 0.000E+00 | 2.472E+04 | 9.691E+03 | 3.915E+04 | 3.672E+05 | 1.191E+06 |
| 831 | 1295.589 | 10.022 | -1 | N/A | N/A | 0.000E+00 | 2.981E+05 | 6.861E+03 | 2.517E+05 | 2.494E+05 | 1.266E+06 | 9.345E+05 | 0.000E+00 | 2.638E+05 | 3.443E+03 | 2.517E+05 | 2.377E+05 | 6.329E+05 | 5.158E+05 |
| 834 | 628.471 | 3.460 | -1 | N/A | N/A | 3.273E+04 | 4.107E+05 | 2.390E+05 | 4.190E+05 | 1.998E+05 | 5.582E+05 | 8.879E+05 | 6.034E+03 | 1.588E+05 | 3.129E+04 | 1.954E+05 | 9.405E+04 | 1.551E+05 | 4.066E+05 |
| 835 | 850.609 | 3.462 | -1 | N/A | N/A | 1.387E+05 | 5.012E+05 | 2.453E+05 | 5.641E+05 | 4.170E+05 | 7.888E+05 | 7.904E+05 | 1.589E+04 | 1.221E+05 | 5.020E+04 | 2.231E+05 | 9.687E+04 | 5.314E+04 | 1.309E+05 |
| 839 | 372.717 | 10.091 | -1 | N/A | N/A | 0.000E+00 | 0.000E+00 | 3.981E+04 | 0.000E+00 | 1.083E+04 | 2.312E+06 | 1.473E+07 | 0.000E+00 | 0.000E+00 | 2.259E+04 | 0.000E+00 | 1.083E+04 | 1.955E+06 | 1.119E+07 |
| 846 | 681.132 | 5.759 | -1 | N/A | N/A | 5.551E+03 | 0.000E+00 | 1.100E+05 | 3.897E+04 | 2.770E+03 | 1.707E+06 | 1.095E+06 | 5.551E+03 | 0.000E+00 | 6.701E+04 | 2.862E+04 | 2.770E+03 | 1.410E+05 | 2.644E+05 |
| 856 | 1194.817 | 11.488 | -1 | N/A | N/A | 5.462E+03 | 1.181E+04 | 3.118E+03 | 0.000E+00 | 1.495E+05 | 4.583E+05 | 2.951E+05 | 5.462E+03 | 6.734E+03 | 3.118E+03 | 0.000E+00 | 8.405E+04 | 2.763E+05 | 2.430E+05 |
| 857 | 804.296 | 3.067 | -1 | N/A | N/A | 0.000E+00 | 9.590E+04 | 0.000E+00 | 1.542E+03 | 1.549E+05 | 1.658E+06 | 3.783E+05 | 0.000E+00 | 3.840E+04 | 0.000E+00 | 1.542E+03 | 8.167E+04 | 9.820E+05 | 2.505E+05 |
| 863 | 353.267 | 9.157 | -1 | N/A | N/A | 4.285E+05 | 3.453E+05 | 3.100E+05 | 2.720E+05 | 3.061E+05 | 5.189E+05 | 5.263E+05 | 1.342E+05 | 5.482E+04 | 5.453E+04 | 1.071E+05 | 6.165E+04 | 1.647E+05 | 1.922E+05 |

| Feature identifiers | | | | XIC | | | | | | | | | | | | | | |
|---|---|---|---|---|---|---|---|---|---|---|---|---|---|---|---|---|---|---|
| ClusterIdx | m/z | RT [min] | Subnetwork | 4TUB34_I | 4TUB34_II | 4TUB34_III | 4TUB81_I | 4TUB81_II | 4TUB81_III | 4TUB93_I | 4TUB93_II | 4TUB93_III | 5TUB23_I | 5TUB23_II | 5TUB23_III | 5TUB48_I | 5TUB48_II | 5TUB48_III |
| 491 | 205.195 | 9.007 | -1 | 5.468E+05 | 3.093E+06 | 2.770E+06 | 4.122E+05 | 6.238E+05 | 1.349E+05 | 8.382E+06 | 1.147E+07 | 8.380E+05 | 2.930E+03 | 4.523E+03 | 5.888E+03 | 0.000E+00 | 2.832E+03 | 1.281E+04 |
| 24 | 399.252 | 8.753 | 58 | 5.214E+07 | 7.733E+07 | 5.841E+07 | 4.343E+07 | 4.200E+07 | 4.237E+07 | 4.273E+07 | 4.397E+07 | 2.092E+08 | 4.139E+07 | 4.492E+07 | 5.518E+07 | 6.397E+07 | 5.154E+07 | 4.523E+07 |
| 44 | 421.232 | 8.753 | -1 | 1.841E+07 | 2.740E+07 | 1.971E+07 | 1.403E+07 | 1.297E+07 | 1.391E+07 | 1.451E+07 | 1.466E+07 | 7.368E+07 | 1.423E+07 | 1.532E+07 | 1.741E+07 | 2.085E+07 | 1.440E+07 | 1.443E+07 |
| 130 | 242.100 | 0.564 | 54 | 1.545E+07 | 1.985E+06 | 2.663E+07 | 1.001E+07 | 1.417E+07 | 1.273E+07 | 8.504E+07 | 9.452E+06 | 6.361E+06 | 4.749E+05 | 4.444E+04 | 8.156E+04 | 2.683E+05 | 5.600E+03 | 0.000E+00 |
| 479 | 242.101 | 0.569 | 54 | 7.183E+05 | 1.214E+06 | 1.232E+07 | 1.891E+06 | 6.839E+06 | 6.452E+06 | 9.083E+07 | 1.047E+07 | 6.853E+06 | 4.749E+05 | 4.444E+04 | 8.156E+04 | 2.683E+05 | 5.600E+03 | 6.198E+03 |
| 478 | 204.087 | 0.529 | 51 | 5.455E+04 | 1.090E+05 | 3.456E+04 | 3.461E+03 | 8.378E+03 | 6.877E+03 | 1.096E+08 | 7.294E+07 | 8.407E+07 | 8.197E+04 | 7.704E+04 | 5.954E+04 | 1.957E+03 | 5.909E+04 | 1.242E+04 |
| 200 | 311.256 | 7.678 | 39 | 2.336E+06 | 6.409E+06 | 8.390E+06 | 4.009E+06 | 4.385E+06 | 3.262E+06 | 5.475E+06 | 3.560E+06 | 1.549E+07 | 1.819E+07 | 8.056E+06 | 1.282E+07 | 1.108E+07 | 1.260E+07 | 9.951E+06 |
| 461 | 311.256 | 7.795 | 39 | 3.118E+05 | 9.747E+05 | 1.292E+06 | 9.141E+05 | 1.634E+06 | 9.371E+05 | 2.496E+06 | 9.351E+05 | 1.257E+06 | 1.697E+07 | 6.711E+06 | 1.075E+07 | 1.033E+07 | 1.049E+07 | 4.843E+06 |
| 536 | 339.290 | 8.561 | 39 | 9.671E+04 | 8.028E+04 | 1.117E+05 | 6.245E+05 | 1.324E+06 | 1.140E+06 | 1.661E+06 | 3.950E+05 | 3.021E+05 | 1.226E+07 | 9.199E+06 | 1.018E+07 | 4.853E+06 | 7.248E+06 | 4.067E+05 |
| 574 | 339.289 | 8.380 | 39 | 1.581E+05 | 2.196E+05 | 4.541E+05 | 7.144E+05 | 7.521E+05 | 9.090E+05 | 5.822E+05 | 2.809E+05 | 1.262E+06 | 2.671E+06 | 2.402E+06 | 2.824E+06 | 9.617E+05 | 1.676E+06 | 7.800E+05 |
| 666 | 188.071 | 1.307 | 27 | 3.424E+03 | 6.882E+03 | 7.024E+03 | 0.000E+00 | 0.000E+00 | 0.000E+00 | 0.000E+00 | 1.560E+03 | 0.000E+00 | 4.401E+03 | 3.642E+03 | 1.926E+03 | 0.000E+00 | 0.000E+00 | 0.000E+00 |
| 6 | 166.086 | 0.517 | 41 | 4.290E+09 | 1.919E+09 | 3.650E+09 | 1.930E+08 | 3.434E+08 | 3.158E+08 | 9.427E+08 | 5.792E+08 | 1.346E+09 | 1.737E+09 | 2.001E+09 | 1.567E+09 | 4.729E+09 | 3.099E+08 | 4.677E+08 |
| 441 | 223.090 | 6.761 | -1 | 2.255E+06 | 6.940E+06 | 5.639E+06 | 2.024E+06 | 1.800E+06 | 1.134E+06 | 1.560E+06 | 1.270E+06 | 5.873E+06 | 1.507E+06 | 1.895E+06 | 1.679E+06 | 1.910E+06 | 1.345E+06 | 3.619E+06 |
| 378 | 391.285 | 10.644 | -1 | 6.254E+05 | 4.669E+05 | 7.317E+05 | 2.678E+05 | 5.660E+05 | 6.059E+05 | 5.185E+05 | 5.528E+05 | 6.289E+05 | 5.728E+05 | 5.940E+05 | 6.350E+05 | 5.678E+05 | 7.792E+05 | 3.601E+05 |
| 12 | 245.096 | 3.603 | 62 | 2.747E+08 | 1.896E+08 | 1.883E+08 | 1.639E+08 | 1.763E+08 | 1.459E+08 | 1.500E+08 | 2.415E+08 | 3.199E+08 | 1.419E+08 | 1.962E+08 | 8.842E+07 | 2.151E+08 | 2.273E+08 | 1.459E+08 |
| 408 | 489.184 | 3.610 | 62 | 5.117E+06 | 2.213E+06 | 2.530E+06 | 1.696E+06 | 1.942E+06 | 1.450E+06 | 1.293E+06 | 3.904E+06 | 6.968E+06 | 1.219E+06 | 2.599E+06 | 3.209E+05 | 3.583E+06 | 3.295E+06 | 1.239E+06 |
| 467 | 250.118 | 7.417 | -1 | 8.566E+05 | 2.515E+05 | 7.449E+05 | 1.331E+06 | 1.234E+06 | 1.268E+06 | 4.595E+05 | 7.218E+05 | 6.458E+05 | 5.035E+05 | 7.195E+05 | 7.655E+05 | 6.697E+05 | 3.615E+05 | 5.214E+05 |
| 462 | 482.321 | 9.064 | 4 | 4.587E+04 | 7.835E+04 | 1.480E+05 | 5.311E+05 | 8.050E+05 | 6.644E+05 | 2.190E+05 | 3.214E+05 | 1.116E+06 | 2.515E+06 | 9.854E+05 | 2.335E+06 | 8.404E+05 | 2.733E+06 | 5.516E+05 |
| 832 | 577.516 | 12.699 | 39 | 0.000E+00 | 0.000E+00 | 7.747E+03 | 0.000E+00 | 6.788E+04 | 0.000E+00 | 7.080E+04 | 6.070E+04 | 8.497E+04 | 7.146E+04 | 1.702E+05 | 2.538E+05 | 2.060E+04 | 9.258E+05 | 6.305E+05 |
| 17 | 454.293 | 8.329 | 4 | 1.578E+07 | 4.219E+07 | 5.951E+07 | 5.359E+07 | 6.378E+07 | 4.715E+07 | 5.619E+07 | 3.082E+07 | 1.326E+08 | 1.841E+08 | 9.021E+07 | 1.347E+08 | 9.171E+07 | 1.193E+08 | 9.067E+07 |
| 89 | 454.293 | 8.190 | 4 | 1.898E+06 | 6.048E+06 | 8.270E+06 | 8.008E+06 | 9.848E+06 | 7.936E+06 | 5.326E+06 | 5.384E+06 | 2.095E+07 | 2.412E+07 | 1.127E+07 | 2.154E+07 | 9.735E+06 | 1.554E+07 | 1.946E+07 |
| 211 | 476.267 | 8.328 | 4 | 9.480E+05 | 2.970E+06 | 3.950E+06 | 3.626E+06 | 4.378E+06 | 3.171E+06 | 3.896E+06 | 2.186E+06 | 9.104E+06 | 1.118E+07 | 6.050E+06 | 8.824E+06 | 6.087E+06 | 7.824E+06 | 6.570E+06 |
| 743 | 907.578 | 8.331 | -1 | 0.000E+00 | 4.880E+04 | 2.346E+05 | 1.172E+05 | 2.360E+05 | 7.365E+04 | 1.613E+05 | 4.224E+04 | 7.952E+05 | 1.252E+06 | 1.751E+05 | 8.828E+05 | 5.280E+05 | 6.148E+05 | 5.112E+05 |
| 138 | 338.342 | 10.765 | 39 | 5.309E+06 | 4.534E+06 | 7.065E+06 | 5.055E+06 | 4.511E+06 | 4.587E+06 | 4.302E+06 | 4.214E+06 | 3.781E+06 | 3.827E+06 | 3.988E+06 | 5.096E+06 | 5.553E+06 | 5.007E+06 | 3.824E+06 |
| 456 | 338.342 | 9.820 | 39 | 1.395E+06 | 1.461E+06 | 1.428E+06 | 1.470E+06 | 1.458E+06 | 1.573E+06 | 1.441E+06 | 1.448E+06 | 1.522E+06 | 1.640E+06 | 1.590E+06 | 1.624E+06 | 1.573E+06 | 2.153E+06 | 1.490E+06 |
| 767 | 688.493 | 12.701 | 4 | 0.000E+00 | 0.000E+00 | 0.000E+00 | 0.000E+00 | 0.000E+00 | 0.000E+00 | 0.000E+00 | 0.000E+00 | 0.000E+00 | 0.000E+00 | 9.173E+05 | 3.254E+06 | 1.544E+05 | 4.183E+06 | 1.177E+07 |
| 30 | 480.309 | 8.382 | 4 | 4.216E+06 | 7.069E+06 | 1.130E+07 | 1.750E+07 | 2.268E+07 | 2.257E+07 | 1.663E+07 | 8.795E+06 | 3.302E+07 | 1.135E+08 | 1.026E+08 | 1.173E+08 | 2.421E+07 | 4.132E+07 | 1.757E+07 |
| 36 | 480.309 | 8.512 | 4 | 1.300E+06 | 2.239E+06 | 3.413E+06 | 1.394E+07 | 1.915E+07 | 1.910E+07 | 9.357E+06 | 3.938E+06 | 1.263E+07 | 4.560E+07 | 4.424E+07 | 5.060E+07 | 1.166E+07 | 2.055E+07 | 1.002E+07 |
| 342 | 377.146 | 3.769 | -1 | 2.223E+05 | 1.294E+06 | 3.977E+05 | 3.072E+06 | 3.607E+06 | 2.673E+06 | 2.447E+06 | 9.775E+05 | 9.832E+06 | 1.799E+06 | 8.622E+05 | 1.260E+06 | 2.178E+05 | 1.124E+06 | 1.439E+06 |
| 473 | 220.118 | 0.532 | 61 | 9.689E+06 | 1.194E+06 | 9.717E+06 | 3.683E+07 | 4.510E+07 | 3.316E+07 | 6.184E+08 | 9.350E+07 | 7.274E+07 | 9.907E+05 | 1.962E+05 | 7.273E+04 | 1.983E+06 | 3.678E+03 | 7.747E+03 |
| 372 | 628.196 | 10.499 | 40 | 1.882E+06 | 1.499E+06 | 1.417E+06 | 1.316E+06 | 1.743E+06 | 1.131E+06 | 9.971E+05 | 1.113E+06 | 1.140E+06 | 1.113E+06 | 1.160E+06 | 1.089E+06 | 1.196E+06 | 1.303E+06 | 1.031E+06 |
| 3 | 432.238 | 7.526 | 24 | 3.587E+08 | 5.056E+08 | 3.741E+08 | 3.390E+08 | 4.757E+08 | 3.564E+08 | 3.801E+08 | 4.288E+08 | 3.760E+08 | 5.429E+08 | 5.087E+08 | 2.887E+08 | 4.716E+08 | 4.145E+08 | 5.568E+08 |
| 161 | 432.238 | 7.249 | 24 | 1.734E+06 | 5.848E+06 | 2.001E+06 | 3.280E+06 | 4.871E+06 | 4.269E+06 | 2.063E+06 | 1.008E+07 | 3.341E+06 | 5.484E+06 | 7.010E+06 | 1.894E+06 | 2.306E+06 | 3.640E+06 | 8.792E+06 |
| 442 | 432.238 | 7.812 | 24 | 3.072E+06 | 1.951E+06 | 1.679E+06 | 1.042E+06 | 1.336E+06 | 1.127E+06 | 7.937E+05 | 2.744E+06 | 9.977E+05 | 1.876E+06 | 2.087E+06 | 1.659E+06 | 2.416E+06 | 1.346E+06 | 2.765E+06 |
| 2 | 415.212 | 7.526 | 24 | 3.492E+08 | 5.272E+08 | 3.706E+08 | 3.295E+08 | 4.850E+08 | 3.509E+08 | 3.729E+08 | 4.566E+08 | 3.878E+08 | 5.702E+08 | 5.582E+08 | 2.876E+08 | 5.122E+08 | 4.148E+08 | 6.072E+08 |
| 78 | 415.212 | 7.248 | 24 | 3.210E+06 | 1.080E+07 | 3.907E+06 | 6.097E+06 | 9.074E+06 | 7.939E+06 | 4.171E+06 | 1.930E+07 | 6.032E+06 | 1.044E+07 | 1.274E+07 | 3.838E+06 | 4.395E+06 | 6.700E+06 | 1.627E+07 |
| 373 | 415.212 | 7.817 | 24 | 9.263E+06 | 4.668E+06 | 3.869E+06 | 1.829E+06 | 2.405E+06 | 1.992E+06 | 1.541E+06 | 5.688E+06 | 2.070E+06 | 3.323E+06 | 3.983E+06 | 4.202E+06 | 4.844E+06 | 3.232E+06 | 6.193E+06 |
| 40 | 327.078 | 8.430 | -1 | 9.186E+05 | 9.646E+05 | 1.304E+06 | 2.196E+07 | 2.094E+07 | 1.473E+07 | 1.327E+06 | 5.111E+06 | 1.418E+07 | 2.421E+06 | 2.409E+06 | 9.023E+05 | 6.915E+05 | 8.319E+05 | 1.177E+06 |
| 435 | 313.143 | 8.965 | -1 | 8.727E+05 | 4.836E+06 | 2.902E+06 | 7.930E+05 | 8.156E+05 | 9.099E+05 | 8.761E+05 | 9.053E+05 | 1.074E+06 | 1.196E+07 | 1.200E+06 | 1.579E+06 | 2.229E+06 | 9.550E+05 | 1.180E+06 |
| 80 | 223.089 | 2.607 | 27 | 3.292E+08 | 4.256E+08 | 1.919E+08 | 6.401E+07 | 5.347E+07 | 5.616E+07 | 1.574E+08 | 3.469E+08 | 1.264E+08 | 1.417E+06 | 8.500E+04 | 1.391E+06 | 7.847E+07 | 4.325E+06 | 5.358E+05 |
| 697 | 223.089 | 2.398 | 27 | 6.305E+07 | 5.308E+07 | 2.834E+07 | 8.385E+06 | 4.125E+06 | 1.073E+07 | 3.229E+07 | 8.113E+07 | 1.318E+07 | 6.664E+07 | 4.149E+08 | 2.794E+09 | 1.111E+09 | 2.097E+09 | 2.824E+09 |
| 798 | 223.089 | 2.491 | 27 | 4.778E+07 | 3.803E+07 | 1.035E+07 | 1.248E+06 | 3.743E+06 | 1.066E+07 | 4.440E+07 | 9.538E+07 | 2.634E+07 | 3.282E+08 | 3.492E+07 | 2.275E+08 | 1.964E+09 | 4.921E+08 | 2.774E+08 |
| 665 | 205.097 | 1.310 | 27 | 5.672E+03 | 1.078E+04 | 2.194E+03 | 3.812E+03 | 0.000E+00 | 4.365E+03 | 1.463E+03 | 0.000E+00 | 0.000E+00 | 4.444E+03 | 0.000E+00 | 0.000E+00 | 0.000E+00 | 0.000E+00 | 0.000E+00 |
| 154 | 457.112 | 3.497 | -1 | 1.506E+07 | 4.050E+07 | 3.655E+07 | 6.228E+06 | 8.312E+06 | 9.839E+06 | 1.878E+07 | 1.532E+07 | 2.874E+07 | 1.420E+07 | 1.299E+07 | 1.499E+07 | 1.965E+07 | 1.940E+07 | 2.179E+07 |
| 29 | 220.118 | 0.526 | 61 | 3.918E+07 | 9.880E+06 | 8.196E+07 | 6.575E+07 | 8.144E+07 | 7.295E+07 | 2.801E+08 | 8.565E+07 | 5.918E+07 | 3.728E+06 | 1.962E+05 | 7.273E+04 | 9.645E+05 | 3.678E+03 | 7.747E+03 |
| 681 | 331.165 | 0.498 | 41 | 1.730E+07 | 4.032E+06 | 1.118E+07 | 0.000E+00 | 3.223E+03 | 5.470E+03 | 0.000E+00 | 0.000E+00 | 2.488E+04 | 4.221E+06 | 5.649E+06 | 3.730E+06 | 1.757E+07 | 1.152E+04 | 3.873E+05 |
| 192 | 343.296 | 6.715 | 23 | 8.946E+06 | 9.317E+06 | 7.099E+06 | 3.691E+06 | 3.576E+06 | 3.687E+06 | 3.404E+06 | 3.500E+06 | 3.415E+06 | 8.390E+06 | 7.406E+06 | 4.464E+06 | 3.608E+06 | 3.177E+06 | 5.032E+06 |
| 589 | 481.400 | 5.241 | 3 | 2.570E+05 | 2.308E+05 | 4.522E+05 | 4.850E+03 | 6.259E+03 | 5.051E+03 | 8.624E+04 | 6.606E+06 | 4.976E+04 | 4.193E+04 | 7.077E+04 | 1.526E+04 | 9.385E+04 | 1.244E+05 | 6.579E+04 |
| 603 | 810.134 | 3.065 | 20 | 6.584E+06 | 1.499E+07 | 3.175E+07 | 1.477E+04 | 1.147E+04 | 7.985E+04 | 7.319E+04 | 5.774E+07 | 1.231E+05 | 1.136E+07 | 3.372E+07 | 1.779E+05 | 4.392E+07 | 1.430E+07 | 6.588E+05 |
| 521 | 716.523 | 12.701 | 4 | 1.037E+05 | 1.906E+05 | 0.000E+00 | 1.241E+06 | 9.144E+05 | 0.000E+00 | 2.460E+06 | 3.239E+05 | 0.000E+00 | 5.553E+05 | 1.418E+06 | 2.866E+05 | 1.292E+07 | 9.327E+06 | 8.621E+05 |
| 572 | 716.525 | 10.404 | 4 | 1.338E+04 | 8.347E+03 | 5.571E+03 | 1.316E+06 | 5.337E+06 | 1.600E+04 | 5.102E+06 | 3.926E+04 | 5.810E+03 | 7.359E+04 | 3.706E+05 | 3.209E+04 | 3.122E+07 | 1.262E+07 | 6.063E+05 |
| 654 | 716.524 | 10.304 | 4 | 3.420E+04 | 1.495E+04 | 1.190E+04 | 8.597E+06 | 7.693E+05 | 2.301E+04 | 1.431E+06 | 2.003E+04 | 4.039E+04 | 7.359E+04 | 5.900E+06 | 6.002E+04 | 2.696E+08 | 2.772E+07 | 8.527E+05 |
| 852 | 716.524 | 10.151 | 4 | 5.979E+04 | 1.219E+04 | 1.985E+04 | 1.018E+06 | 2.920E+05 | 2.608E+04 | 5.543E+04 | 1.686E+05 | 5.417E+04 | 2.872E+05 | 2.989E+05 | 1.407E+05 | 2.147E+07 | 3.632E+08 | 9.616E+05 |
| 524 | 426.262 | 7.583 | 4 | 9.957E+05 | 3.549E+06 | 4.094E+06 | 8.854E+05 | 9.314E+05 | 6.924E+05 | 1.560E+06 | 8.945E+05 | 3.503E+06 | 1.185E+07 | 5.342E+06 | 8.207E+06 | 4.949E+06 | 6.601E+06 | 1.442E+07 |
| 552 | 426.262 | 7.443 | 4 | 9.772E+05 | 5.799E+06 | 6.534E+06 | 7.249E+05 | 6.977E+05 | 4.143E+05 | 1.127E+06 | 1.334E+06 | 4.788E+06 | 1.384E+07 | 7.085E+06 | 7.492E+06 | 1.026E+07 | 1.329E+07 | 2.224E+07 |
| 507 | 718.539 | 12.704 | 4 | 0.000E+00 | 0.000E+00 | 4.669E+06 | 5.659E+06 | 6.960E+05 | 2.558E+05 | 4.310E+06 | 6.576E+05 | 2.854E+06 | 2.350E+06 | 9.149E+05 | 8.486E+06 | 5.745E+06 | 8.012E+07 | 5.506E+07 |
| 523 | 718.539 | 10.302 | 4 | 0.000E+00 | 0.000E+00 | 1.224E+06 | 3.480E+06 | 6.221E+05 | 1.322E+05 | 3.796E+07 | 1.194E+06 | 7.332E+06 | 3.572E+07 | 7.687E+05 | 6.270E+07 | 1.863E+06 | 5.696E+06 | 9.325E+06 |
| 632 | 718.538 | 10.143 | 4 | 0.000E+00 | 0.000E+00 | 3.365E+05 | 7.800E+05 | 2.571E+07 | 3.771E+04 | 1.615E+06 | 2.235E+07 | 1.703E+05 | 1.545E+06 | 1.214E+05 | 4.502E+06 | 1.004E+07 | 4.194E+06 | 6.143E+07 |
| 823 | 771.514 | 12.696 | 4 | 0.000E+00 | 0.000E+00 | 0.000E+00 | 0.000E+00 | 0.000E+00 | 0.000E+00 | 2.042E+05 | 9.950E+04 | 1.897E+05 | 1.073E+05 | 5.439E+05 | 4.442E+05 | 5.647E+04 | 2.155E+05 | 1.495E+05 |
| 346 | 502.290 | 8.380 | 4 | 3.289E+05 | 4.434E+05 | 7.084E+05 | 1.259E+06 | 1.492E+06 | 1.684E+06 | 1.195E+06 | 5.363E+05 | 2.480E+06 | 4.857E+06 | 4.470E+06 | 4.872E+06 | 1.718E+06 | 3.087E+06 | 1.295E+06 |
| 388 | 502.291 | 8.511 | 4 | 3.054E+04 | 1.263E+05 | 1.202E+05 | 9.155E+05 | 1.398E+06 | 1.319E+06 | 5.836E+05 | 2.330E+05 | 8.723E+05 | 2.985E+06 | 3.009E+06 | 3.411E+06 | 8.483E+05 | 1.326E+06 | 5.008E+05 |
| 476 | 979.349 | 0.522 | 91 | 0.000E+00 | 0.000E+00 | 0.000E+00 | 0.000E+00 | 0.000E+00 | 0.000E+00 | 1.955E+08 | 1.647E+08 | 1.481E+08 | 0.000E+00 | 0.000E+00 | 0.000E+00 | 0.000E+00 | 0.000E+00 | 0.000E+00 |
| 488 | 995.318 | 0.510 | 91 | 0.000E+00 | 0.000E+00 | 0.000E+00 | 0.000E+00 | 0.000E+00 | 0.000E+00 | 2.381E+07 | 1.803E+07 | 2.020E+07 | 0.000E+00 | 0.000E+00 | 0.000E+00 | 0.000E+00 | 0.000E+00 | 0.000E+00 |
| 52 | 256.264 | 10.439 | 88 | 2.158E+07 | 1.043E+07 | 8.719E+06 | 1.487E+07 | 1.791E+07 | 1.352E+07 | 1.691E+07 | 1.186E+07 | 8.482E+06 | 7.792E+06 | 1.255E+07 | 1.097E+07 | 1.616E+07 | 1.128E+07 | 6.698E+06 |
| 427 | 284.295 | 10.783 | 88 | 1.609E+06 | 1.042E+06 | 6.856E+05 | 1.110E+06 | 1.627E+06 | 8.283E+05 | 1.903E+06 | 7.480E+05 | 1.022E+06 | 1.717E+06 | 2.407E+06 | 2.572E+06 | 3.016E+06 | 2.642E+06 | 1.237E+06 |
| 62 | 291.096 | 6.457 | 87 | 0.000E+00 | 0.000E+00 | 0.000E+00 | 8.201E+06 | 1.270E+07 | 8.792E+06 | 0.000E+00 | 0.000E+00 | 1.416E+06 | 0.000E+00 | 0.000E+00 | 0.000E+00 | 0.000E+00 | 0.000E+00 | 0.000E+00 |
| 153 | 291.096 | 6.559 | 87 | 0.000E+00 | 0.000E+00 | 0.000E+00 | 4.027E+06 | 6.485E+06 | 4.531E+06 | 0.000E+00 | 0.000E+00 | 6.154E+05 | 7.596E+03 | 0.000E+00 | 0.000E+00 | 0.000E+00 | 0.000E+00 | 0.000E+00 |
| 374 | 748.356 | 4.106 | 84 | 0.000E+00 | 0.000E+00 | 0.000E+00 | 1.347E+06 | 1.349E+06 | 1.270E+06 | 0.000E+00 | 0.000E+00 | 6.552E+05 | 0.000E+00 | 0.000E+00 | 0.000E+00 | 0.000E+00 | 0.000E+00 | 0.000E+00 |
| 384 | 783.875 | 4.128 | 84 | 0.000E+00 | 0.000E+00 | 0.000E+00 | 1.500E+06 | 1.676E+06 | 1.624E+06 | 0.000E+00 | 0.000E+00 | 9.123E+05 | 0.000E+00 | 0.000E+00 | 0.000E+00 | 0.000E+00 | 0.000E+00 | 0.000E+00 |
| 284 | 243.196 | 7.929 | 80 | 1.316E+04 | 6.743E+04 | 9.242E+04 | 2.531E+06 | 3.329E+06 | 2.298E+06 | 2.267E+06 | 3.668E+05 | 7.773E+06 | 9.842E+04 | 7.468E+03 | 1.279E+04 | 1.076E+04 | 8.751E+03 | 1.885E+04 |

| Feature identifiers | | | | XIC | | | | | | | | | | | | | | |
|---|---|---|---|---|---|---|---|---|---|---|---|---|---|---|---|---|---|---|
| ClusterIdx | m/z | RT [min] | Subnetwork | 4TUB34_I | 4TUB34_II | 4TUB34_III | 4TUB81_I | 4TUB81_II | 4TUB81_III | 4TUB93_I | 4TUB93_II | 4TUB93_III | 5TUB23_I | 5TUB23_II | 5TUB23_III | 5TUB48_I | 5TUB48_II | 5TUB48_III |
| 415 | 225.185 | 7.934 | 80 | 1.177E+04 | 4.604E+04 | 5.983E+04 | 1.284E+06 | 1.739E+06 | 1.204E+06 | 1.165E+06 | 2.114E+05 | 4.099E+06 | 5.516E+04 | 4.588E+03 | 1.376E+04 | 1.017E+04 | 4.757E+03 | 2.266E+04 |
| 667 | 176.074 | 0.736 | 79 | 6.233E+07 | 1.433E+08 | 2.296E+08 | 1.906E+04 | 7.572E+04 | 8.925E+04 | 5.455E+05 | 6.792E+05 | 1.184E+05 | 5.876E+07 | 4.773E+07 | 7.313E+07 | 8.721E+07 | 2.387E+05 | 3.928E+06 |
| 668 | 218.121 | 4.865 | 79 | 2.003E+07 | 2.595E+07 | 4.533E+07 | 0.000E+00 | 7.365E+04 | 9.812E+04 | 2.042E+05 | 8.245E+04 | 3.114E+05 | 1.084E+07 | 1.325E+07 | 1.232E+07 | 1.394E+07 | 9.875E+04 | 2.866E+05 |
| 670 | 246.152 | 6.015 | 79 | 2.248E+07 | 1.938E+07 | 3.739E+07 | 5.516E+03 | 2.275E+05 | 1.948E+05 | 5.853E+04 | 4.362E+04 | 3.899E+05 | 1.394E+07 | 1.786E+07 | 1.313E+07 | 1.560E+07 | 1.139E+05 | 7.583E+04 |
| 674 | 232.137 | 5.471 | 79 | 6.937E+06 | 6.902E+06 | 1.117E+07 | 0.000E+00 | 3.239E+04 | 0.000E+00 | 5.440E+03 | 0.000E+00 | 3.912E+04 | 3.406E+06 | 4.733E+06 | 3.286E+06 | 3.864E+06 | 1.088E+04 | 3.339E+04 |
| 675 | 204.105 | 4.140 | 79 | 1.444E+07 | 1.719E+07 | 2.440E+07 | 0.000E+00 | 0.000E+00 | 0.000E+00 | 2.222E+04 | 1.752E+04 | 1.266E+05 | 8.151E+06 | 8.255E+06 | 5.817E+06 | 4.999E+06 | 0.000E+00 | 0.000E+00 |
| 772 | 551.192 | 3.315 | 78 | 9.012E+03 | 7.019E+06 | 6.460E+06 | 0.000E+00 | 0.000E+00 | 0.000E+00 | 0.000E+00 | 2.452E+04 | 2.220E+05 | 0.000E+00 | 2.896E+03 | 5.528E+03 | 1.081E+04 | 0.000E+00 | 1.706E+06 |
| 774 | 615.714 | 3.354 | 78 | 1.506E+04 | 5.803E+06 | 5.248E+06 | 0.000E+00 | 0.000E+00 | 0.000E+00 | 0.000E+00 | 1.154E+04 | 0.000E+00 | 0.000E+00 | 0.000E+00 | 0.000E+00 | 1.384E+04 | 1.631E+04 | 1.776E+06 |
| 776 | 486.670 | 3.240 | 78 | 0.000E+00 | 3.293E+06 | 3.282E+06 | 0.000E+00 | 0.000E+00 | 0.000E+00 | 0.000E+00 | 0.000E+00 | 2.537E+05 | 0.000E+00 | 0.000E+00 | 0.000E+00 | 0.000E+00 | 0.000E+00 | 7.799E+05 |
| 777 | 680.235 | 3.406 | 78 | 0.000E+00 | 3.164E+06 | 2.202E+06 | 0.000E+00 | 0.000E+00 | 0.000E+00 | 0.000E+00 | 1.592E+04 | 0.000E+00 | 0.000E+00 | 0.000E+00 | 0.000E+00 | 1.741E+04 | 0.000E+00 | 1.497E+06 |
| 780 | 357.628 | 3.065 | 78 | 9.842E+03 | 5.650E+06 | 6.838E+06 | 1.030E+04 | 1.431E+04 | 2.368E+04 | 1.040E+05 | 2.167E+05 | 3.014E+05 | 0.000E+00 | 3.919E+05 | 5.659E+04 | 2.241E+04 | 4.383E+04 | 8.373E+05 |
| 46 | 226.158 | 9.291 | 77 | 7.276E+05 | 2.444E+06 | 2.066E+06 | 6.342E+06 | 1.269E+07 | 9.775E+06 | 2.470E+06 | 1.112E+07 | 3.254E+07 | 3.362E+06 | 1.239E+06 | 2.495E+06 | 4.748E+05 | 3.674E+05 | 4.018E+05 |
| 201 | 282.221 | 10.073 | 77 | 2.951E+05 | 3.100E+06 | 7.937E+05 | 7.914E+05 | 4.045E+05 | 4.001E+06 | 1.962E+05 | 1.473E+06 | 5.840E+06 | 3.222E+06 | 1.718E+05 | 7.927E+06 | 5.280E+05 | 3.651E+05 | 2.927E+05 |
| 634 | 282.221 | 10.084 | 77 | 2.951E+05 | 2.028E+06 | 5.762E+05 | 3.827E+05 | 4.045E+05 | 2.667E+06 | 1.962E+05 | 5.331E+05 | 4.467E+06 | 6.347E+06 | 1.718E+05 | 4.052E+06 | 5.280E+05 | 3.651E+05 | 2.927E+05 |
| 150 | 818.299 | 3.185 | 75 | 1.257E+07 | 2.706E+07 | 2.403E+07 | 9.870E+06 | 1.107E+07 | 7.647E+06 | 1.284E+07 | 1.923E+07 | 1.536E+07 | 1.190E+07 | 8.663E+06 | 3.352E+07 | 3.207E+07 | 4.408E+07 | 5.634E+07 |
| 323 | 1226.943 | 3.181 | 75 | 3.044E+06 | 7.199E+06 | 6.909E+06 | 2.606E+06 | 3.009E+06 | 2.340E+06 | 3.582E+06 | 5.760E+06 | 4.513E+06 | 2.233E+06 | 2.430E+06 | 6.851E+06 | 1.047E+07 | 1.231E+07 | 1.050E+07 |
| 843 | 1226.943 | 3.183 | 75 | 3.086E+06 | 5.664E+06 | 4.441E+06 | 2.740E+06 | 1.604E+06 | 1.249E+06 | 2.803E+06 | 3.000E+06 | 4.348E+06 | 1.802E+06 | 2.336E+06 | 3.024E+06 | 3.465E+06 | 1.014E+07 | 1.599E+07 |
| 186 | 135.079 | 7.525 | 74 | 5.436E+06 | 8.229E+06 | 6.189E+06 | 5.739E+06 | 7.253E+06 | 5.314E+06 | 6.110E+06 | 7.109E+06 | 6.676E+06 | 1.011E+07 | 9.447E+06 | 4.573E+06 | 8.051E+06 | 7.128E+06 | 1.014E+07 |
| 319 | 135.079 | 4.463 | 74 | 3.077E+06 | 1.918E+06 | 1.488E+06 | 1.699E+06 | 2.502E+06 | 1.677E+06 | 1.603E+06 | 3.595E+06 | 1.735E+06 | 3.538E+06 | 3.684E+06 | 2.025E+06 | 3.913E+06 | 2.671E+06 | 4.505E+06 |
| 180 | 529.790 | 4.168 | 73 | 0.000E+00 | 0.000E+00 | 0.000E+00 | 6.500E+06 | 5.990E+06 | 4.592E+06 | 0.000E+00 | 0.000E+00 | 5.950E+06 | 1.148E+04 | 0.000E+00 | 0.000E+00 | 0.000E+00 | 0.000E+00 | 0.000E+00 |
| 264 | 465.270 | 4.196 | 73 | 0.000E+00 | 0.000E+00 | 0.000E+00 | 3.715E+06 | 4.400E+06 | 2.835E+06 | 0.000E+00 | 0.000E+00 | 3.844E+06 | 9.629E+03 | 0.000E+00 | 7.600E+03 | 0.000E+00 | 0.000E+00 | 0.000E+00 |
| 48 | 431.565 | 4.119 | 72 | 5.152E+04 | 0.000E+00 | 0.000E+00 | 1.061E+07 | 1.384E+07 | 1.662E+07 | 1.800E+05 | 1.562E+05 | 2.957E+05 | 0.000E+00 | 0.000E+00 | 0.000E+00 | 0.000E+00 | 0.000E+00 | 0.000E+00 |
| 170 | 417.549 | 3.504 | 72 | 9.939E+07 | 2.916E+07 | 2.759E+07 | 4.501E+06 | 6.609E+06 | 1.009E+07 | 1.407E+07 | 1.722E+08 | 8.528E+06 | 4.623E+06 | 1.624E+07 | 5.615E+04 | 1.814E+07 | 1.059E+07 | 1.866E+06 |
| 314 | 396.544 | 3.209 | 72 | 6.741E+05 | 9.150E+06 | 3.433E+06 | 6.379E+06 | 4.465E+06 | 4.932E+06 | 7.465E+06 | 3.522E+07 | 8.466E+06 | 1.201E+05 | 3.073E+06 | 5.373E+05 | 2.224E+04 | 1.178E+06 | 1.090E+06 |
| 117 | 832.302 | 3.461 | 70 | 6.929E+06 | 1.544E+07 | 1.574E+07 | 7.837E+06 | 9.326E+06 | 7.595E+06 | 1.810E+07 | 2.491E+06 | 1.828E+07 | 1.135E+07 | 6.207E+06 | 9.932E+06 | 1.868E+07 | 1.550E+07 | 4.043E+07 |
| 371 | 1247.949 | 3.461 | 70 | 2.358E+06 | 3.917E+06 | 5.272E+06 | 2.830E+06 | 2.653E+06 | 1.939E+06 | 5.815E+06 | 7.737E+05 | 6.398E+06 | 3.611E+06 | 1.947E+06 | 2.975E+06 | 5.694E+06 | 4.720E+06 | 1.387E+07 |
| 512 | 1247.454 | 3.462 | 70 | 1.821E+06 | 4.554E+06 | 4.258E+06 | 2.062E+06 | 1.778E+06 | 1.357E+06 | 4.668E+06 | 4.840E+05 | 5.156E+06 | 3.337E+06 | 1.275E+06 | 2.073E+06 | 5.338E+06 | 3.774E+06 | 6.453E+06 |
| 7 | 271.106 | 8.022 | 69 | 3.281E+07 | 2.469E+07 | 3.708E+07 | 1.007E+08 | 1.382E+08 | 1.308E+08 | 1.050E+07 | 8.624E+07 | 6.802E+07 | 5.829E+05 | 1.798E+05 | 1.076E+05 | 1.389E+05 | 2.141E+04 | 6.240E+04 |
| 45 | 390.122 | 6.709 | 69 | 1.703E+06 | 1.317E+06 | 1.567E+06 | 1.185E+07 | 1.454E+07 | 1.188E+07 | 1.059E+06 | 5.137E+06 | 9.351E+06 | 0.000E+00 | 0.000E+00 | 0.000E+00 | 0.000E+00 | 0.000E+00 | 0.000E+00 |
| 112 | 388.106 | 7.488 | 69 | 1.922E+06 | 8.078E+05 | 1.472E+06 | 4.450E+06 | 5.754E+06 | 5.224E+06 | 5.372E+05 | 5.812E+06 | 3.992E+06 | 0.000E+00 | 0.000E+00 | 0.000E+00 | 0.000E+00 | 0.000E+00 | 0.000E+00 |
| 209 | 414.122 | 6.104 | 69 | 3.362E+06 | 3.670E+06 | 6.328E+06 | 1.899E+06 | 2.560E+06 | 3.048E+06 | 7.431E+05 | 1.557E+07 | 5.611E+06 | 0.000E+00 | 0.000E+00 | 4.837E+03 | 0.000E+00 | 0.000E+00 | 0.000E+00 |
| 663 | 414.122 | 6.419 | 69 | 4.718E+04 | 0.000E+00 | 8.534E+04 | 0.000E+00 | 4.299E+05 | 3.295E+04 | 9.256E+03 | 9.090E+04 | 2.224E+06 | 5.885E+05 | 4.138E+06 | 4.569E+05 | 3.805E+06 | 2.259E+06 | 5.141E+06 |
| 468 | 432.238 | 6.913 | 68 | 2.652E+07 | 2.266E+05 | 1.510E+07 | 1.430E+06 | 1.002E+06 | 5.373E+05 | 3.430E+05 | 3.394E+05 | 4.373E+05 | 5.136E+05 | 2.575E+05 | 5.510E+05 | 3.003E+07 | 6.146E+05 | 4.580E+05 |
| 792 | 404.208 | 6.914 | 68 | 2.000E+07 | 1.272E+05 | 1.086E+07 | 7.923E+05 | 5.989E+05 | 3.474E+05 | 1.897E+05 | 1.519E+05 | 1.367E+05 | 3.744E+05 | 1.363E+05 | 2.614E+05 | 2.225E+07 | 4.431E+05 | 2.192E+05 |
| 793 | 387.181 | 6.913 | 68 | 1.904E+07 | 1.687E+05 | 1.074E+07 | 7.852E+05 | 5.965E+05 | 3.736E+05 | 1.060E+05 | 2.767E+05 | 1.841E+05 | 3.827E+05 | 1.509E+05 | 2.332E+05 | 2.183E+07 | 4.129E+05 | 2.385E+05 |
| 671 | 331.207 | 5.234 | 67 | 1.347E+07 | 2.897E+06 | 1.272E+07 | 2.326E+04 | 8.967E+04 | 2.841E+05 | 1.288E+05 | 3.082E+04 | 1.465E+04 | 2.300E+06 | 5.000E+06 | 4.848E+06 | 2.354E+07 | 2.132E+05 | 8.092E+04 |
| 795 | 317.195 | 4.828 | 67 | 1.197E+06 | 3.266E+05 | 1.306E+06 | 1.884E+04 | 6.556E+04 | 6.358E+04 | 6.278E+04 | 1.691E+05 | 1.358E+05 | 2.711E+05 | 6.970E+05 | 5.663E+05 | 2.701E+06 | 1.919E+05 | 4.120E+05 |
| 84 | 280.188 | 5.797 | 65 | 1.231E+03 | 7.750E+03 | 6.445E+03 | 5.935E+06 | 7.532E+06 | 6.835E+06 | 6.281E+03 | 0.000E+00 | 1.629E+07 | 0.000E+00 | 0.000E+00 | 3.428E+03 | 0.000E+00 | 0.000E+00 | 0.000E+00 |
| 158 | 308.222 | 6.723 | 65 | 0.000E+00 | 0.000E+00 | 0.000E+00 | 3.707E+06 | 5.131E+06 | 4.592E+06 | 3.505E+03 | 0.000E+00 | 7.806E+06 | 0.000E+00 | 0.000E+00 | 0.000E+00 | 0.000E+00 | 0.000E+00 | 0.000E+00 |
| 263 | 693.878 | 3.983 | 64 | 0.000E+00 | 0.000E+00 | 0.000E+00 | 2.399E+06 | 2.680E+06 | 2.631E+06 | 0.000E+00 | 0.000E+00 | 8.184E+05 | 0.000E+00 | 0.000E+00 | 0.000E+00 | 0.000E+00 | 0.000E+00 | 0.000E+00 |
| 392 | 665.367 | 3.986 | 64 | 0.000E+00 | 0.000E+00 | 0.000E+00 | 1.248E+06 | 1.671E+06 | 1.436E+06 | 0.000E+00 | 0.000E+00 | 1.052E+06 | 0.000E+00 | 0.000E+00 | 0.000E+00 | 0.000E+00 | 0.000E+00 | 0.000E+00 |
| 703 | 204.534 | 6.795 | 63 | 5.087E+03 | 0.000E+00 | 0.000E+00 | 0.000E+00 | 0.000E+00 | 0.000E+00 | 0.000E+00 | 0.000E+00 | 0.000E+00 | 1.811E+07 | 1.035E+07 | 9.487E+06 | 0.000E+00 | 0.000E+00 | 0.000E+00 |
| 704 | 193.026 | 6.808 | 63 | 0.000E+00 | 0.000E+00 | 0.000E+00 | 0.000E+00 | 0.000E+00 | 0.000E+00 | 0.000E+00 | 0.000E+00 | 0.000E+00 | 1.990E+07 | 1.048E+07 | 1.099E+07 | 0.000E+00 | 0.000E+00 | 0.000E+00 |
| 88 | 227.084 | 3.606 | 62 | 3.028E+07 | 2.064E+07 | 2.067E+07 | 1.803E+07 | 1.916E+07 | 1.599E+07 | 1.629E+07 | 2.659E+07 | 3.546E+07 | 1.565E+07 | 2.147E+07 | 9.628E+06 | 2.394E+07 | 2.525E+07 | 1.610E+07 |
| 215 | 290.153 | 3.607 | 62 | 1.218E+07 | 7.373E+06 | 7.202E+06 | 6.671E+06 | 7.503E+06 | 5.933E+06 | 5.922E+06 | 1.098E+07 | 1.595E+07 | 5.323E+06 | 7.467E+06 | 3.264E+06 | 9.071E+06 | 9.995E+06 | 6.407E+06 |
| 4 | 182.985 | 12.752 | 60 | 6.356E+08 | 4.292E+08 | 5.436E+08 | 8.219E+08 | 8.035E+08 | 7.858E+08 | 8.000E+08 | 7.799E+08 | 7.802E+08 | 7.630E+08 | 7.630E+08 | 7.396E+08 | 7.356E+08 | 7.128E+08 | 7.297E+08 |
| 5 | 154.990 | 12.767 | 60 | 8.411E+08 | 7.835E+08 | 7.586E+08 | 7.800E+08 | 7.644E+08 | 7.214E+08 | 7.673E+08 | 7.373E+08 | 7.501E+08 | 7.449E+08 | 7.242E+08 | 7.068E+08 | 7.325E+08 | 6.609E+08 | 7.069E+08 |
| 19 | 138.996 | 12.748 | 60 | 1.284E+08 | 1.176E+08 | 1.115E+08 | 1.161E+08 | 1.105E+08 | 1.075E+08 | 1.175E+08 | 1.088E+08 | 1.130E+08 | 1.122E+08 | 1.101E+08 | 1.047E+08 | 1.037E+08 | 1.008E+08 | 1.009E+08 |
| 23 | 205.017 | 12.741 | 60 | 5.651E+07 | 5.066E+07 | 4.623E+07 | 5.318E+07 | 5.021E+07 | 4.644E+07 | 5.155E+07 | 4.525E+07 | 5.010E+07 | 4.651E+07 | 4.887E+07 | 4.417E+07 | 4.710E+07 | 4.168E+07 | 4.199E+07 |
| 49 | 196.017 | 12.766 | 60 | 4.074E+07 | 3.544E+07 | 3.237E+07 | 3.460E+07 | 3.382E+07 | 2.815E+07 | 3.287E+07 | 3.088E+07 | 3.274E+07 | 3.167E+07 | 3.128E+07 | 2.727E+07 | 2.963E+07 | 2.560E+07 | 2.843E+07 |
| 63 | 224.012 | 12.748 | 60 | 3.208E+07 | 2.932E+07 | 2.614E+07 | 2.883E+07 | 2.543E+07 | 2.304E+07 | 2.548E+07 | 2.505E+07 | 2.556E+07 | 2.441E+07 | 2.404E+07 | 2.100E+07 | 2.256E+07 | 1.851E+07 | 2.159E+07 |
| 75 | 195.994 | 12.775 | 60 | 3.231E+07 | 3.208E+07 | 3.205E+07 | 3.185E+07 | 3.118E+07 | 3.039E+07 | 3.270E+07 | 3.170E+07 | 3.146E+07 | 3.054E+07 | 3.104E+07 | 3.106E+07 | 3.052E+07 | 2.840E+07 | 2.991E+07 |
| 20 | 131.534 | 12.743 | 59 | 7.958E+07 | 7.596E+07 | 7.095E+07 | 7.523E+07 | 7.208E+07 | 6.886E+07 | 7.007E+07 | 6.605E+07 | 6.905E+07 | 6.604E+07 | 6.686E+07 | 6.177E+07 | 6.393E+07 | 5.539E+07 | 5.974E+07 |
| 409 | 131.534 | 13.009 | 59 | 1.671E+06 | 8.154E+05 | 1.826E+06 | 1.146E+06 | 5.082E+05 | 3.281E+05 | 3.430E+05 | 1.453E+06 | 2.783E+05 | 4.803E+05 | 1.374E+06 | 3.812E+05 | 2.487E+05 | 5.791E+05 | 3.479E+05 |
| 465 | 131.534 | 13.069 | 59 | 5.047E+05 | 1.592E+06 | 9.309E+05 | 5.180E+05 | 2.283E+05 | 3.350E+05 | 5.856E+05 | 1.191E+05 | 4.170E+05 | 5.413E+05 | 7.007E+05 | 9.531E+04 | 2.618E+05 | 7.478E+05 | 6.862E+05 |
| 696 | 131.534 | 4.507 | 59 | 1.310E+05 | 1.992E+05 | 1.220E+05 | 3.112E+05 | 8.523E+04 | 2.886E+05 | 1.935E+05 | 1.932E+05 | 4.672E+04 | 1.116E+05 | 1.019E+05 | 4.194E+04 | 1.074E+05 | 8.872E+04 | 1.907E+05 |
| 266 | 444.309 | 8.752 | 58 | 3.332E+06 | 5.487E+06 | 3.433E+06 | 2.616E+06 | 2.515E+06 | 2.767E+06 | 2.695E+06 | 2.855E+06 | 1.427E+07 | 2.898E+06 | 2.977E+06 | 3.658E+06 | 4.447E+06 | 2.879E+06 | 3.303E+06 |
| 490 | 350.102 | 0.493 | 57 | 1.773E+04 | 1.697E+07 | 8.865E+06 | 6.541E+05 | 9.840E+04 | 6.533E+04 | 4.111E+06 | 1.109E+07 | 0.000E+00 | 1.358E+07 | 2.245E+07 | 4.316E+06 | 5.895E+04 | 3.791E+07 | 6.817E+06 |
| 495 | 364.118 | 0.634 | 57 | 1.821E+04 | 8.805E+06 | 6.009E+06 | 0.000E+00 | 0.000E+00 | 0.000E+00 | 7.929E+07 | 9.040E+05 | 0.000E+00 | 2.499E+05 | 1.013E+05 | 6.595E+07 | 0.000E+00 | 7.818E+04 | 1.413E+05 |
| 134 | 519.318 | 10.374 | 56 | 1.533E+06 | 3.045E+04 | 4.406E+04 | 5.877E+06 | 1.064E+05 | 4.354E+06 | 2.950E+06 | 8.760E+05 | 3.869E+06 | 2.449E+06 | 1.344E+06 | 2.346E+05 | 2.129E+06 | 2.442E+06 | 2.059E+05 |
| 658 | 519.309 | 10.382 | 56 | 1.662E+06 | 1.813E+06 | 2.007E+06 | 5.685E+06 | 3.437E+06 | 4.498E+06 | 3.051E+06 | 1.335E+06 | 3.979E+06 | 2.715E+06 | 1.411E+06 | 2.251E+06 | 2.446E+06 | 2.561E+06 | 2.490E+06 |
| 498 | 532.098 | 0.493 | 55 | 0.000E+00 | 0.000E+00 | 0.000E+00 | 0.000E+00 | 0.000E+00 | 0.000E+00 | 3.172E+06 | 1.041E+07 | 2.519E+06 | 0.000E+00 | 0.000E+00 | 0.000E+00 | 0.000E+00 | 0.000E+00 | 0.000E+00 |
| 501 | 532.099 | 0.497 | 55 | 0.000E+00 | 0.000E+00 | 0.000E+00 | 0.000E+00 | 0.000E+00 | 0.000E+00 | 1.625E+06 | 1.655E+06 | 1.019E+07 | 0.000E+00 | 0.000E+00 | 0.000E+00 | 0.000E+00 | 0.000E+00 | 0.000E+00 |
| 132 | 711.319 | 4.210 | 53 | 0.000E+00 | 9.464E+04 | 0.000E+00 | 5.162E+06 | 7.074E+06 | 5.027E+06 | 0.000E+00 | 0.000E+00 | 2.729E+06 | 0.000E+00 | 0.000E+00 | 0.000E+00 | 0.000E+00 | 0.000E+00 | 0.000E+00 |
| 282 | 624.289 | 4.196 | 53 | 0.000E+00 | 0.000E+00 | 0.000E+00 | 1.061E+06 | 2.333E+06 | 2.063E+06 | 0.000E+00 | 0.000E+00 | 7.556E+05 | 0.000E+00 | 0.000E+00 | 0.000E+00 | 0.000E+00 | 0.000E+00 | 0.000E+00 |
| 133 | 226.180 | 5.976 | 52 | 6.336E+06 | 5.725E+06 | 5.971E+06 | 5.289E+06 | 4.675E+06 | 5.495E+06 | 5.954E+06 | 6.176E+06 | 5.159E+06 | 5.428E+06 | 5.599E+06 | 6.010E+06 | 6.930E+06 | 5.935E+06 | 5.871E+06 |
| 542 | 384.349 | 10.283 | 52 | 1.420E+06 | 1.533E+06 | 1.480E+06 | 1.838E+06 | 1.837E+06 | 1.388E+06 | 1.669E+06 | 1.622E+06 | 1.443E+06 | 1.512E+06 | 1.566E+06 | 1.414E+06 | 2.606E+06 | 2.541E+06 | 2.510E+06 |
| 131 | 614.463 | 9.741 | 51 | 1.107E+06 | 7.343E+05 | 3.016E+06 | 1.069E+07 | 1.154E+07 | 1.062E+07 | 1.107E+07 | 5.353E+06 | 5.956E+06 | 2.281E+06 | 2.453E+05 | 1.654E+06 | 5.530E+05 | 4.910E+05 | 3.646E+05 |
| 243 | 388.269 | 6.414 | 51 | 6.037E+05 | 3.935E+05 | 6.242E+05 | 6.071E+06 | 5.125E+06 | 3.188E+06 | 3.499E+06 | 1.356E+06 | 2.388E+06 | 3.398E+05 | 7.022E+04 | 2.204E+05 | 1.612E+04 | 1.159E+05 | 1.396E+05 |
| 472 | 479.186 | 0.514 | 51 | 2.080E+03 | 4.241E+04 | 3.048E+05 | 0.000E+00 | 0.000E+00 | 0.000E+00 | 4.081E+09 | 2.734E+09 | 3.298E+09 | 1.994E+06 | 4.827E+06 | 2.140E+06 | 4.091E+04 | 1.165E+05 | 6.751E+05 |

| Feature identifiers | | | | XIC | | | | | | | | | | | | | | |
|---|---|---|---|---|---|---|---|---|---|---|---|---|---|---|---|---|---|---|
| ClusterIdx | m/z | RT [min] | Subnetwork | 4TUB34_I | 4TUB34_II | 4TUB34_III | 4TUB81_I | 4TUB81_II | 4TUB81_III | 4TUB93_I | 4TUB93_II | 4TUB93_III | 5TUB23_I | 5TUB23_II | 5TUB23_III | 5TUB48_I | 5TUB48_II | 5TUB48_III |
| 475 | 957.367 | 0.530 | 51 | 0.000E+00 | 0.000E+00 | 0.000E+00 | 0.000E+00 | 0.000E+00 | 0.000E+00 | 2.436E+08 | 1.264E+08 | 1.800E+08 | 0.000E+00 | 0.000E+00 | 0.000E+00 | 0.000E+00 | 0.000E+00 | 0.000E+00 |
| 477 | 524.246 | 0.489 | 51 | 0.000E+00 | 0.000E+00 | 0.000E+00 | 0.000E+00 | 0.000E+00 | 0.000E+00 | 8.045E+07 | 7.643E+07 | 5.836E+07 | 0.000E+00 | 0.000E+00 | 0.000E+00 | 0.000E+00 | 0.000E+00 | 0.000E+00 |
| 480 | 268.076 | 0.497 | 51 | 0.000E+00 | 0.000E+00 | 0.000E+00 | 0.000E+00 | 0.000E+00 | 0.000E+00 | 5.967E+07 | 4.982E+07 | 4.724E+07 | 0.000E+00 | 0.000E+00 | 0.000E+00 | 0.000E+00 | 0.000E+00 | 0.000E+00 |
| 482 | 498.161 | 0.503 | 51 | 0.000E+00 | 0.000E+00 | 0.000E+00 | 0.000E+00 | 0.000E+00 | 0.000E+00 | 6.276E+07 | 4.260E+07 | 4.744E+07 | 0.000E+00 | 0.000E+00 | 0.000E+00 | 0.000E+00 | 0.000E+00 | 0.000E+00 |
| 483 | 644.267 | 0.510 | 51 | 0.000E+00 | 0.000E+00 | 0.000E+00 | 0.000E+00 | 0.000E+00 | 0.000E+00 | 4.467E+07 | 2.636E+07 | 6.187E+07 | 0.000E+00 | 1.571E+04 | 7.256E+02 | 0.000E+00 | 0.000E+00 | 0.000E+00 |
| 484 | 279.584 | 0.532 | 51 | 0.000E+00 | 0.000E+00 | 0.000E+00 | 0.000E+00 | 0.000E+00 | 0.000E+00 | 5.808E+07 | 4.958E+07 | 4.462E+07 | 0.000E+00 | 0.000E+00 | 0.000E+00 | 0.000E+00 | 0.000E+00 | 0.000E+00 |
| 487 | 276.062 | 0.490 | 51 | 0.000E+00 | 0.000E+00 | 0.000E+00 | 0.000E+00 | 0.000E+00 | 0.000E+00 | 1.755E+07 | 1.324E+07 | 1.216E+07 | 0.000E+00 | 0.000E+00 | 0.000E+00 | 0.000E+00 | 0.000E+00 | 0.000E+00 |
| 492 | 737.248 | 0.522 | 51 | 0.000E+00 | 0.000E+00 | 0.000E+00 | 0.000E+00 | 0.000E+00 | 0.000E+00 | 1.985E+07 | 1.137E+07 | 1.454E+07 | 0.000E+00 | 0.000E+00 | 0.000E+00 | 0.000E+00 | 0.000E+00 | 0.000E+00 |
| 493 | 506.147 | 0.488 | 51 | 0.000E+00 | 0.000E+00 | 0.000E+00 | 0.000E+00 | 0.000E+00 | 0.000E+00 | 1.168E+07 | 8.483E+06 | 9.951E+06 | 0.000E+00 | 0.000E+00 | 0.000E+00 | 0.000E+00 | 0.000E+00 | 0.000E+00 |
| 500 | 287.570 | 0.486 | 51 | 0.000E+00 | 0.000E+00 | 0.000E+00 | 0.000E+00 | 0.000E+00 | 0.000E+00 | 9.619E+06 | 7.806E+06 | 3.319E+06 | 0.000E+00 | 0.000E+00 | 0.000E+00 | 0.000E+00 | 0.000E+00 | 0.000E+00 |
| 687 | 441.187 | 0.508 | 51 | 2.939E+04 | 8.239E+06 | 1.239E+07 | 4.119E+03 | 0.000E+00 | 0.000E+00 | 0.000E+00 | 0.000E+00 | 0.000E+00 | 1.440E+07 | 1.448E+07 | 4.470E+06 | 7.570E+04 | 7.282E+04 | 1.044E+06 |
| 732 | 632.474 | 10.196 | 51 | 1.489E+04 | 1.674E+05 | 2.188E+05 | 5.123E+04 | 7.056E+04 | 1.625E+04 | 4.585E+04 | 1.834E+04 | 1.459E+05 | 3.583E+06 | 8.045E+05 | 1.911E+06 | 3.039E+06 | 3.168E+06 | 8.016E+05 |
| 802 | 860.514 | 9.355 | 51 | 2.804E+04 | 8.345E+04 | 7.202E+04 | 0.000E+00 | 3.354E+04 | 0.000E+00 | 2.152E+05 | 1.958E+04 | 3.267E+04 | 2.082E+06 | 9.687E+05 | 9.297E+05 | 9.476E+05 | 5.907E+05 | 4.152E+05 |
| 168 | 471.307 | 5.923 | 50 | 6.426E+06 | 4.878E+06 | 2.684E+06 | 3.221E+06 | 4.015E+06 | 3.608E+06 | 3.696E+06 | 4.512E+06 | 4.054E+06 | 5.758E+06 | 7.576E+06 | 2.938E+06 | 4.321E+06 | 4.937E+06 | 4.713E+06 |
| 616 | 473.286 | 4.748 | 50 | 3.099E+06 | 3.903E+06 | 3.012E+06 | 9.562E+05 | 3.149E+05 | 4.506E+05 | 5.932E+04 | 2.407E+06 | 2.904E+06 | 7.062E+05 | 1.649E+06 | 2.906E+05 | 2.809E+06 | 1.319E+06 | 2.601E+06 |
| 144 | 393.207 | 10.310 | 49 | 5.319E+06 | 5.471E+06 | 5.037E+06 | 5.207E+06 | 6.366E+06 | 5.531E+06 | 5.667E+06 | 5.711E+06 | 5.583E+06 | 5.650E+06 | 5.929E+06 | 5.726E+06 | 5.856E+06 | 6.088E+06 | 6.042E+06 |
| 176 | 409.183 | 10.451 | 49 | 3.774E+06 | 4.088E+06 | 3.558E+06 | 4.093E+06 | 4.372E+06 | 4.154E+06 | 4.003E+06 | 3.926E+06 | 3.969E+06 | 3.995E+06 | 3.608E+06 | 3.830E+06 | 4.374E+06 | 4.313E+06 | 4.216E+06 |
| 43 | 418.117 | 7.633 | 48 | 2.115E+06 | 2.056E+06 | 1.618E+06 | 1.278E+07 | 1.500E+07 | 1.294E+07 | 1.328E+06 | 2.170E+07 | 1.170E+07 | 0.000E+00 | 0.000E+00 | 0.000E+00 | 0.000E+00 | 0.000E+00 | 0.000E+00 |
| 146 | 551.149 | 6.292 | 48 | 4.586E+06 | 2.061E+05 | 4.694E+06 | 1.638E+06 | 6.245E+06 | 6.365E+06 | 4.005E+05 | 2.394E+06 | 1.018E+07 | 0.000E+00 | 0.000E+00 | 0.000E+00 | 0.000E+00 | 0.000E+00 | 0.000E+00 |
| 363 | 418.117 | 7.570 | 48 | 2.232E+06 | 1.669E+05 | 1.799E+06 | 5.688E+06 | 1.352E+06 | 1.597E+06 | 8.306E+05 | 1.338E+06 | 3.205E+06 | 0.000E+00 | 0.000E+00 | 0.000E+00 | 0.000E+00 | 0.000E+00 | 0.000E+00 |
| 119 | 343.130 | 3.690 | 47 | 8.943E+06 | 1.787E+07 | 1.317E+07 | 1.036E+07 | 1.015E+07 | 9.211E+06 | 8.706E+06 | 6.478E+07 | 1.063E+08 | 3.659E+05 | 5.412E+04 | 6.204E+03 | 2.069E+05 | 4.397E+05 | 0.000E+00 |
| 182 | 537.334 | 9.562 | 47 | 7.850E+04 | 1.063E+04 | 4.458E+04 | 1.096E+07 | 5.085E+06 | 3.289E+06 | 2.675E+04 | 2.075E+05 | 5.393E+06 | 1.291E+04 | 4.160E+03 | 3.184E+04 | 8.612E+03 | 2.537E+04 | 0.000E+00 |
| 231 | 509.303 | 8.961 | 47 | 0.000E+00 | 0.000E+00 | 0.000E+00 | 3.966E+06 | 2.985E+06 | 2.314E+06 | 0.000E+00 | 2.096E+04 | 2.688E+06 | 0.000E+00 | 0.000E+00 | 0.000E+00 | 0.000E+00 | 0.000E+00 | 0.000E+00 |
| 239 | 245.616 | 7.678 | 46 | 1.423E+06 | 4.136E+06 | 5.334E+06 | 2.150E+06 | 2.461E+06 | 2.241E+06 | 3.230E+06 | 2.134E+06 | 9.102E+06 | 1.032E+07 | 4.986E+06 | 7.857E+06 | 6.791E+06 | 7.350E+06 | 6.389E+06 |
| 393 | 266.129 | 7.678 | 46 | 8.498E+05 | 2.569E+06 | 3.640E+06 | 1.573E+06 | 1.749E+06 | 1.335E+06 | 2.270E+06 | 1.456E+06 | 6.150E+06 | 6.379E+06 | 3.018E+06 | 4.611E+06 | 4.644E+06 | 4.112E+06 | 3.915E+06 |
| 429 | 266.129 | 7.807 | 46 | 1.919E+05 | 4.804E+05 | 5.802E+05 | 8.887E+05 | 1.620E+06 | 8.638E+05 | 1.236E+06 | 4.787E+05 | 1.615E+06 | 3.151E+06 | 1.625E+06 | 2.404E+06 | 5.410E+05 | 1.045E+06 | 1.447E+06 |
| 506 | 734.471 | 12.708 | 45 | 6.854E+05 | 5.014E+05 | 0.000E+00 | 5.540E+06 | 4.966E+06 | 1.599E+05 | 3.000E+06 | 8.943E+05 | 5.463E+04 | 1.763E+06 | 4.090E+06 | 8.168E+05 | 4.849E+06 | 4.894E+06 | 7.194E+05 |
| 544 | 734.471 | 10.338 | 45 | 4.309E+04 | 9.605E+04 | 0.000E+00 | 1.830E+06 | 2.396E+07 | 5.667E+04 | 9.070E+06 | 2.984E+05 | 7.290E+03 | 1.735E+06 | 1.955E+07 | 1.656E+05 | 3.078E+06 | 6.662E+06 | 6.636E+05 |
| 550 | 765.459 | 12.703 | 45 | 0.000E+00 | 0.000E+00 | 0.000E+00 | 0.000E+00 | 0.000E+00 | 0.000E+00 | 1.382E+06 | 7.653E+04 | 0.000E+00 | 0.000E+00 | 0.000E+00 | 8.650E+05 | 1.526E+05 | 5.859E+06 | 5.177E+06 |
| 646 | 734.471 | 10.261 | 45 | 1.970E+04 | 3.760E+04 | 0.000E+00 | 3.407E+07 | 4.833E+06 | 5.667E+04 | 2.854E+06 | 1.120E+05 | 0.000E+00 | 2.521E+05 | 1.105E+06 | 1.467E+05 | 6.165E+07 | 1.110E+08 | 4.829E+05 |
| 652 | 762.502 | 12.706 | 45 | 0.000E+00 | 0.000E+00 | 4.565E+05 | 6.959E+05 | 3.877E+04 | 0.000E+00 | 1.011E+06 | 0.000E+00 | 5.653E+05 | 2.430E+05 | 0.000E+00 | 1.313E+06 | 6.287E+05 | 8.794E+06 | 7.441E+06 |
| 370 | 514.353 | 10.382 | 43 | 4.292E+05 | 6.306E+05 | 5.334E+05 | 1.925E+06 | 1.261E+06 | 1.109E+06 | 9.584E+05 | 3.719E+05 | 8.239E+05 | 5.504E+05 | 4.969E+05 | 7.449E+05 | 1.144E+06 | 1.017E+06 | 1.054E+06 |
| 740 | 744.593 | 12.218 | 43 | 8.403E+03 | 0.000E+00 | 0.000E+00 | 4.118E+04 | 0.000E+00 | 7.176E+03 | 0.000E+00 | 0.000E+00 | 1.501E+04 | 1.435E+05 | 0.000E+00 | 5.233E+06 | 3.773E+06 | 0.000E+00 | 9.129E+06 |
| 94 | 247.596 | 7.524 | 42 | 7.832E+06 | 1.228E+07 | 8.229E+06 | 6.354E+06 | 1.123E+07 | 8.960E+06 | 9.456E+06 | 1.090E+07 | 1.043E+07 | 1.262E+07 | 1.266E+07 | 6.419E+06 | 1.171E+07 | 9.730E+06 | 1.367E+07 |
| 110 | 256.601 | 7.527 | 42 | 6.884E+06 | 1.039E+07 | 7.060E+06 | 6.606E+06 | 9.922E+06 | 7.083E+06 | 8.048E+06 | 8.669E+06 | 7.642E+06 | 1.301E+07 | 1.178E+07 | 4.784E+06 | 1.052E+07 | 8.793E+06 | 1.301E+07 |
| 233 | 268.115 | 7.525 | 42 | 4.350E+06 | 4.275E+06 | 3.809E+06 | 3.135E+06 | 4.892E+06 | 3.669E+06 | 2.760E+06 | 3.937E+06 | 3.763E+06 | 4.856E+06 | 4.272E+06 | 2.062E+06 | 4.331E+06 | 3.464E+06 | 4.604E+06 |
| 283 | 554.177 | 10.161 | 40 | 2.211E+06 | 1.785E+06 | 1.754E+06 | 2.037E+06 | 1.960E+06 | 1.614E+06 | 1.494E+06 | 1.487E+06 | 1.244E+06 | 1.909E+06 | 1.732E+06 | 1.407E+06 | 1.655E+06 | 2.250E+06 | 1.498E+06 |
| 545 | 684.203 | 10.983 | 40 | 2.362E+06 | 1.004E+05 | 1.511E+05 | 2.469E+06 | 2.931E+05 | 9.399E+04 | 2.819E+06 | 1.054E+05 | 1.278E+05 | 3.529E+06 | 2.781E+05 | 5.632E+04 | 4.970E+06 | 1.050E+05 | 5.059E+04 |
| 630 | 702.214 | 10.746 | 40 | 1.062E+06 | 7.512E+05 | 5.994E+05 | 7.177E+05 | 7.149E+05 | 5.791E+05 | 5.851E+05 | 5.421E+05 | 5.941E+05 | 6.064E+05 | 5.344E+05 | 3.996E+05 | 6.064E+05 | 5.754E+05 | 4.617E+05 |
| 693 | 463.131 | 9.688 | 40 | 7.641E+05 | 7.570E+05 | 5.871E+05 | 5.199E+05 | 3.415E+05 | 5.086E+05 | 5.195E+05 | 4.457E+05 | 4.977E+05 | 3.896E+05 | 5.509E+05 | 7.250E+05 | 4.808E+05 | 5.161E+05 | 4.489E+05 |
| 821 | 776.232 | 11.006 | 40 | 2.079E+05 | 3.310E+05 | 3.304E+05 | 3.179E+05 | 3.626E+05 | 3.478E+05 | 2.912E+05 | 3.557E+05 | 2.771E+05 | 1.990E+05 | 1.974E+05 | 3.204E+05 | 2.400E+05 | 3.045E+05 | 2.390E+05 |
| 858 | 610.185 | 12.140 | 40 | 1.029E+06 | 7.126E+05 | 6.215E+05 | 1.019E+06 | 5.143E+05 | 6.762E+05 | 7.339E+05 | 7.512E+05 | 8.260E+05 | 9.683E+05 | 9.643E+05 | 3.840E+05 | 1.131E+06 | 1.150E+06 | 5.132E+05 |
| 143 | 228.196 | 7.148 | 39 | 1.339E+06 | 3.055E+06 | 3.552E+06 | 2.161E+06 | 2.677E+06 | 4.482E+06 | 5.402E+06 | 3.306E+06 | 3.259E+06 | 2.300E+06 | 2.613E+06 | 3.618E+06 | 2.167E+06 | 4.263E+06 | 6.064E+06 |
| 230 | 228.196 | 6.462 | 39 | 6.444E+06 | 1.710E+07 | 1.700E+07 | 8.573E+06 | 9.757E+06 | 1.532E+07 | 1.917E+07 | 1.129E+07 | 1.334E+07 | 8.255E+06 | 9.178E+06 | 1.174E+07 | 7.672E+06 | 1.395E+07 | 1.963E+07 |
| 824 | 549.485 | 10.160 | 39 | 0.000E+00 | 0.000E+00 | 0.000E+00 | 2.051E+05 | 1.877E+05 | 0.000E+00 | 4.382E+05 | 3.130E+05 | 9.820E+03 | 9.558E+04 | 2.555E+05 | 3.994E+04 | 5.031E+05 | 2.715E+06 | 4.389E+06 |
| 826 | 549.486 | 12.704 | 39 | 0.000E+00 | 0.000E+00 | 0.000E+00 | 4.657E+05 | 2.008E+05 | 0.000E+00 | 1.274E+06 | 1.573E+05 | 0.000E+00 | 4.038E+04 | 3.046E+05 | 6.755E+05 | 4.456E+05 | 6.118E+06 | 4.085E+06 |
| 840 | 549.486 | 9.853 | 39 | 0.000E+00 | 0.000E+00 | 3.922E+03 | 2.536E+04 | 3.629E+04 | 1.974E+04 | 2.568E+05 | 7.306E+05 | 2.297E+04 | 5.683E+04 | 1.182E+04 | 1.629E+04 | 4.746E+06 | 4.635E+06 | 7.489E+07 |
| 850 | 549.485 | 10.361 | 39 | 0.000E+00 | 0.000E+00 | 0.000E+00 | 8.988E+05 | 6.552E+05 | 0.000E+00 | 6.022E+05 | 5.520E+04 | 0.000E+00 | 3.266E+04 | 1.178E+05 | 1.155E+05 | 3.240E+05 | 5.854E+07 | 2.243E+06 |
| 159 | 480.238 | 3.674 | 37 | 0.000E+00 | 0.000E+00 | 0.000E+00 | 9.914E+06 | 8.780E+06 | 6.546E+06 | 0.000E+00 | 0.000E+00 | 8.530E+06 | 0.000E+00 | 0.000E+00 | 0.000E+00 | 0.000E+00 | 0.000E+00 | 0.000E+00 |
| 332 | 959.468 | 3.674 | 37 | 0.000E+00 | 0.000E+00 | 0.000E+00 | 4.447E+06 | 4.446E+06 | 3.130E+06 | 0.000E+00 | 0.000E+00 | 3.895E+06 | 0.000E+00 | 0.000E+00 | 0.000E+00 | 0.000E+00 | 0.000E+00 | 0.000E+00 |
| 301 | 467.250 | 5.033 | 36 | 0.000E+00 | 0.000E+00 | 0.000E+00 | 4.866E+06 | 3.555E+06 | 1.786E+06 | 0.000E+00 | 0.000E+00 | 3.680E+06 | 0.000E+00 | 0.000E+00 | 0.000E+00 | 0.000E+00 | 0.000E+00 | 0.000E+00 |
| 333 | 933.494 | 5.044 | 36 | 0.000E+00 | 0.000E+00 | 0.000E+00 | 5.479E+06 | 3.765E+06 | 1.752E+06 | 0.000E+00 | 1.345E+03 | 3.797E+06 | 0.000E+00 | 0.000E+00 | 0.000E+00 | 0.000E+00 | 0.000E+00 | 0.000E+00 |
| 420 | 516.785 | 5.231 | 36 | 0.000E+00 | 0.000E+00 | 0.000E+00 | 2.280E+06 | 2.328E+06 | 1.033E+06 | 0.000E+00 | 0.000E+00 | 2.021E+06 | 0.000E+00 | 0.000E+00 | 0.000E+00 | 0.000E+00 | 0.000E+00 | 0.000E+00 |
| 68 | 217.105 | 4.009 | 35 | 5.137E+07 | 7.554E+07 | 5.266E+07 | 3.642E+07 | 3.886E+07 | 1.736E+07 | 3.078E+07 | 3.448E+07 | 5.552E+07 | 2.750E+07 | 3.311E+07 | 2.094E+07 | 3.025E+07 | 2.555E+07 | 4.308E+07 |
| 330 | 234.134 | 4.002 | 35 | 1.025E+07 | 1.444E+07 | 1.027E+07 | 7.068E+06 | 7.440E+06 | 2.970E+06 | 5.559E+06 | 6.547E+06 | 1.055E+07 | 4.838E+06 | 6.109E+06 | 3.632E+06 | 5.563E+06 | 4.574E+06 | 7.246E+06 |
| 431 | 681.651 | 10.701 | 33 | 1.176E+06 | 1.215E+06 | 1.052E+06 | 1.018E+06 | 1.223E+06 | 9.873E+05 | 7.973E+05 | 8.670E+05 | 8.666E+05 | 8.229E+05 | 8.032E+05 | 8.042E+05 | 7.553E+05 | 8.173E+05 | 8.615E+05 |
| 434 | 399.395 | 7.402 | 33 | 1.122E+06 | 1.051E+06 | 1.075E+06 | 1.002E+06 | 1.112E+06 | 9.839E+05 | 9.924E+05 | 1.029E+06 | 9.759E+05 | 1.031E+06 | 1.028E+06 | 1.019E+06 | 9.847E+05 | 8.577E+05 | 9.466E+05 |
| 171 | 278.185 | 3.992 | 32 | 1.347E+07 | 2.079E+07 | 1.719E+07 | 8.352E+06 | 7.438E+06 | 7.793E+06 | 1.239E+07 | 1.341E+07 | 1.426E+07 | 1.027E+07 | 1.227E+07 | 1.202E+07 | 1.385E+07 | 1.570E+07 | 1.637E+07 |
| 437 | 280.186 | 4.202 | 32 | 4.437E+06 | 8.497E+06 | 5.564E+06 | 2.623E+06 | 2.255E+06 | 2.792E+06 | 3.954E+06 | 4.579E+06 | 6.529E+06 | 4.156E+06 | 4.179E+06 | 4.469E+06 | 5.944E+06 | 6.158E+06 | 6.338E+06 |
| 560 | 271.180 | 3.637 | 32 | 2.917E+06 | 4.073E+06 | 2.653E+06 | 1.119E+06 | 9.996E+05 | 9.576E+05 | 1.388E+06 | 2.707E+06 | 3.712E+06 | 3.178E+06 | 3.128E+06 | 3.043E+06 | 3.519E+06 | 3.872E+06 | 3.348E+06 |
| 35 | 255.159 | 6.531 | 31 | 2.239E+07 | 2.144E+07 | 2.099E+07 | 1.994E+07 | 2.164E+07 | 2.327E+07 | 2.210E+07 | 2.283E+07 | 1.971E+07 | 2.193E+07 | 2.396E+07 | 2.015E+07 | 2.313E+07 | 2.510E+07 | 2.654E+07 |
| 99 | 272.185 | 6.531 | 31 | 7.965E+06 | 7.063E+06 | 6.663E+06 | 6.713E+06 | 7.038E+06 | 7.814E+06 | 7.200E+06 | 7.419E+06 | 6.583E+06 | 7.421E+06 | 7.784E+06 | 6.565E+06 | 7.777E+06 | 8.135E+06 | 8.346E+06 |
| 466 | 269.175 | 7.631 | 31 | 1.194E+06 | 4.992E+06 | 4.582E+06 | 2.444E+06 | 3.594E+06 | 6.568E+05 | 6.335E+05 | 2.396E+06 | 7.749E+06 | 5.276E+05 | 1.341E+06 | 1.013E+06 | 1.754E+06 | 1.465E+06 | 6.426E+06 |
| 28 | 297.133 | 7.153 | 30 | 2.328E+07 | 2.279E+07 | 1.902E+07 | 1.547E+07 | 1.692E+07 | 2.332E+07 | 2.269E+07 | 2.803E+07 | 2.572E+07 | 2.120E+07 | 2.492E+07 | 1.450E+07 | 1.959E+07 | 2.677E+07 | 2.524E+07 |
| 38 | 431.207 | 7.153 | 30 | 1.570E+07 | 1.534E+07 | 1.252E+07 | 1.034E+07 | 1.152E+07 | 1.496E+07 | 1.503E+07 | 1.859E+07 | 1.675E+07 | 1.354E+07 | 1.611E+07 | 8.803E+06 | 1.265E+07 | 1.751E+07 | 1.638E+07 |
| 56 | 447.202 | 6.602 | 30 | 1.134E+07 | 1.487E+07 | 1.198E+07 | 7.026E+06 | 6.284E+06 | 9.814E+06 | 1.005E+07 | 1.283E+07 | 1.505E+07 | 8.846E+06 | 9.983E+06 | 7.853E+06 | 1.020E+07 | 1.410E+07 | 1.437E+07 |
| 91 | 448.233 | 7.152 | 30 | 8.122E+06 | 7.970E+06 | 6.343E+06 | 4.904E+06 | 5.139E+06 | 6.721E+06 | 6.976E+06 | 9.200E+06 | 8.584E+06 | 6.971E+06 | 8.210E+06 | 4.306E+06 | 6.313E+06 | 8.566E+06 | 8.119E+06 |
| 102 | 429.191 | 6.409 | 30 | 7.057E+06 | 8.594E+06 | 6.928E+06 | 4.874E+06 | 4.142E+06 | 6.178E+06 | 6.847E+06 | 9.682E+06 | 1.084E+07 | 6.070E+06 | 6.928E+06 | 5.320E+06 | 7.014E+06 | 1.011E+07 | 1.188E+07 |
| 163 | 297.133 | 7.367 | 30 | 6.906E+06 | 5.966E+06 | 4.000E+06 | 3.910E+06 | 4.976E+06 | 5.724E+06 | 5.245E+06 | 6.956E+06 | 5.484E+06 | 5.323E+06 | 6.973E+06 | 3.368E+06 | 5.177E+06 | 6.565E+06 | 6.752E+06 |
| 202 | 448.233 | 7.470 | 30 | 4.128E+06 | 4.361E+06 | 2.987E+06 | 2.520E+06 | 2.590E+06 | 3.382E+06 | 3.078E+06 | 4.326E+06 | 3.948E+06 | 4.546E+06 | 6.103E+06 | 2.590E+06 | 3.759E+06 | 5.636E+06 | 5.586E+06 |

| Feature identifiers | | | | XIC | | | | | | | | | | | | | | |
|---|---|---|---|---|---|---|---|---|---|---|---|---|---|---|---|---|---|---|
| ClusterIdx | m/z | RT [min] | Subnetwork | 4TUB34_I | 4TUB34_II | 4TUB34_III | 4TUB81_I | 4TUB81_II | 4TUB81_III | 4TUB93_I | 4TUB93_II | 4TUB93_III | 5TUB23_I | 5TUB23_II | 5TUB23_III | 5TUB48_I | 5TUB48_II | 5TUB48_III |
| 208 | 431.207 | 7.472 | 30 | 7.441E+06 | 6.788E+06 | 2.711E+06 | 2.682E+06 | 2.852E+06 | 5.661E+06 | 5.543E+06 | 7.484E+06 | 6.110E+06 | 6.344E+06 | 7.723E+06 | 3.836E+06 | 5.781E+06 | 7.692E+06 | 7.045E+06 |
| 213 | 447.202 | 7.074 | 30 | 1.782E+06 | 1.950E+06 | 1.579E+06 | 1.854E+06 | 1.987E+06 | 2.947E+06 | 2.821E+06 | 2.715E+06 | 2.843E+06 | 2.424E+06 | 2.097E+06 | 2.222E+06 | 1.730E+06 | 3.429E+06 | 3.100E+06 |
| 219 | 447.202 | 6.409 | 30 | 3.252E+06 | 3.899E+06 | 2.942E+06 | 2.085E+06 | 1.867E+06 | 2.712E+06 | 2.934E+06 | 4.234E+06 | 4.836E+06 | 2.688E+06 | 2.980E+06 | 2.300E+06 | 2.995E+06 | 4.378E+06 | 5.289E+06 |
| 235 | 297.133 | 6.600 | 30 | 2.666E+06 | 3.520E+06 | 2.741E+06 | 1.772E+06 | 1.497E+06 | 2.370E+06 | 2.571E+06 | 3.472E+06 | 3.770E+06 | 2.299E+06 | 2.459E+06 | 1.995E+06 | 2.499E+06 | 3.874E+06 | 3.723E+06 |
| 245 | 469.184 | 6.601 | 30 | 2.660E+06 | 3.671E+06 | 2.836E+06 | 1.534E+06 | 1.383E+06 | 2.331E+06 | 2.412E+06 | 3.216E+06 | 3.515E+06 | 2.226E+06 | 2.345E+06 | 1.837E+06 | 2.601E+06 | 3.483E+06 | 3.560E+06 |
| 249 | 429.191 | 6.602 | 30 | 2.584E+06 | 3.688E+06 | 2.914E+06 | 1.540E+06 | 1.401E+06 | 2.311E+06 | 2.318E+06 | 3.148E+06 | 3.643E+06 | 2.141E+06 | 2.332E+06 | 1.835E+06 | 2.456E+06 | 3.428E+06 | 3.517E+06 |
| 251 | 297.133 | 6.410 | 30 | 2.496E+06 | 2.995E+06 | 2.432E+06 | 1.552E+06 | 1.347E+06 | 2.319E+06 | 2.381E+06 | 3.315E+06 | 3.846E+06 | 2.157E+06 | 2.266E+06 | 1.865E+06 | 2.502E+06 | 3.616E+06 | 4.276E+06 |
| 262 | 447.202 | 6.882 | 30 | 5.088E+06 | 5.652E+06 | 5.110E+06 | 3.894E+06 | 3.899E+06 | 5.531E+06 | 5.127E+06 | 7.058E+06 | 6.781E+06 | 4.711E+06 | 5.189E+06 | 4.096E+06 | 5.256E+06 | 8.526E+06 | 6.574E+06 |
| 411 | 527.717 | 3.679 | 29 | 0.000E+00 | 0.000E+00 | 0.000E+00 | 1.857E+05 | 6.378E+05 | 1.548E+06 | 2.844E+06 | 6.490E+04 | 0.000E+00 | 0.000E+00 | 5.454E+03 | 0.000E+00 | 0.000E+00 | 3.652E+03 | 0.000E+00 |
| 444 | 791.074 | 3.678 | 29 | 0.000E+00 | 0.000E+00 | 0.000E+00 | 1.370E+05 | 5.351E+05 | 1.100E+06 | 2.513E+06 | 2.773E+04 | 0.000E+00 | 5.181E+03 | 0.000E+00 | 0.000E+00 | 0.000E+00 | 0.000E+00 | 0.000E+00 |
| 355 | 500.380 | 6.151 | 28 | 3.813E+06 | 3.275E+06 | 3.303E+06 | 3.658E+06 | 3.499E+06 | 3.445E+06 | 3.376E+06 | 3.246E+06 | 3.104E+06 | 3.390E+06 | 3.577E+06 | 3.410E+06 | 3.237E+06 | 3.256E+06 | 3.136E+06 |
| 391 | 442.338 | 5.719 | 28 | 2.922E+06 | 2.970E+06 | 2.718E+06 | 2.786E+06 | 2.804E+06 | 2.628E+06 | 2.792E+06 | 2.819E+06 | 2.651E+06 | 2.691E+06 | 2.899E+06 | 2.740E+06 | 2.949E+06 | 2.681E+06 | 2.708E+06 |
| 394 | 309.227 | 4.773 | 28 | 4.304E+06 | 5.219E+06 | 4.764E+06 | 3.638E+06 | 2.907E+06 | 2.795E+06 | 3.981E+06 | 3.415E+06 | 3.391E+06 | 3.102E+06 | 4.246E+06 | 3.754E+06 | 3.650E+06 | 3.166E+06 | 3.539E+06 |
| 446 | 384.296 | 5.267 | 28 | 2.992E+06 | 3.282E+06 | 2.967E+06 | 2.572E+06 | 2.169E+06 | 2.085E+06 | 2.485E+06 | 2.647E+06 | 2.534E+06 | 2.362E+06 | 2.651E+06 | 2.436E+06 | 2.490E+06 | 2.150E+06 | 2.272E+06 |
| 458 | 674.506 | 7.346 | 28 | 1.420E+06 | 1.906E+06 | 8.782E+05 | 2.083E+06 | 2.276E+06 | 2.243E+06 | 2.169E+06 | 1.241E+06 | 2.087E+06 | 1.907E+06 | 5.270E+05 | 6.321E+05 | 1.310E+06 | 9.350E+05 | 2.101E+06 |
| 631 | 790.590 | 8.085 | 28 | 2.164E+06 | 1.443E+06 | 9.216E+05 | 1.551E+06 | 8.131E+05 | 1.036E+06 | 1.041E+06 | 2.401E+06 | 1.570E+06 | 1.417E+06 | 1.531E+06 | 1.014E+06 | 9.536E+05 | 1.334E+06 | 2.092E+06 |
| 635 | 549.416 | 9.554 | 28 | 4.848E+05 | 3.700E+05 | 1.590E+05 | 6.723E+05 | 5.048E+05 | 6.450E+05 | 1.721E+05 | 1.526E+06 | 1.190E+06 | 8.351E+05 | 4.644E+05 | 2.173E+05 | 5.291E+05 | 5.782E+05 | 3.154E+05 |
| 860 | 694.522 | 10.428 | 28 | 3.807E+05 | 2.942E+05 | 4.159E+05 | 1.061E+06 | 2.755E+05 | 4.462E+05 | 5.098E+05 | 3.520E+05 | 4.969E+05 | 2.909E+05 | 3.534E+05 | 3.032E+05 | 2.763E+05 | 1.980E+06 | 6.782E+05 |
| 357 | 206.062 | 2.609 | 27 | 5.952E+07 | 7.761E+07 | 3.513E+07 | 1.139E+07 | 9.598E+06 | 1.007E+07 | 2.894E+07 | 6.437E+07 | 2.312E+07 | 5.582E+05 | 5.938E+04 | 1.125E+06 | 3.827E+07 | 8.745E+05 | 1.853E+06 |
| 440 | 237.106 | 4.092 | 27 | 1.196E+07 | 1.569E+07 | 1.784E+07 | 8.298E+05 | 1.051E+06 | 1.162E+06 | 7.625E+06 | 8.962E+06 | 9.420E+06 | 3.735E+05 | 1.510E+06 | 1.944E+06 | 1.442E+06 | 1.758E+06 | 3.716E+06 |
| 699 | 206.061 | 2.395 | 27 | 1.080E+07 | 6.248E+06 | 3.711E+06 | 7.896E+05 | 7.926E+05 | 2.023E+05 | 1.153E+06 | 1.246E+06 | 1.855E+06 | 1.350E+07 | 2.007E+08 | 1.364E+09 | 1.516E+08 | 1.026E+09 | 1.376E+09 |
| 799 | 206.062 | 2.458 | 27 | 4.014E+06 | 2.058E+06 | 2.480E+06 | 7.896E+05 | 3.131E+05 | 6.762E+05 | 1.534E+06 | 3.315E+06 | 8.917E+05 | 1.597E+08 | 4.710E+07 | 4.052E+07 | 9.584E+08 | 1.332E+08 | 7.505E+07 |
| 448 | 213.144 | 5.071 | 26 | 1.962E+06 | 1.687E+06 | 1.807E+06 | 1.598E+06 | 1.731E+06 | 1.516E+06 | 1.533E+06 | 7.165E+05 | 1.208E+06 | 1.403E+06 | 7.643E+05 | 6.149E+05 | 1.316E+06 | 1.132E+06 | 3.601E+05 |
| 692 | 381.331 | 9.796 | 26 | 1.422E+06 | 5.908E+05 | 6.734E+05 | 4.796E+05 | 6.993E+05 | 4.232E+05 | 4.807E+05 | 4.630E+05 | 4.638E+05 | 6.282E+05 | 9.597E+04 | 9.531E+05 | 5.946E+05 | 6.330E+05 | 4.264E+05 |
| 210 | 445.290 | 4.792 | 25 | 2.983E+06 | 2.502E+06 | 2.694E+06 | 2.581E+06 | 2.680E+06 | 2.748E+06 | 2.592E+06 | 2.662E+06 | 2.525E+06 | 2.701E+06 | 2.770E+06 | 2.812E+06 | 2.947E+06 | 4.070E+06 | 3.848E+06 |
| 227 | 467.300 | 4.849 | 25 | 2.382E+06 | 2.290E+06 | 2.342E+06 | 2.139E+06 | 2.252E+06 | 2.301E+06 | 2.306E+06 | 2.340E+06 | 2.117E+06 | 1.913E+06 | 2.239E+06 | 2.248E+06 | 2.475E+06 | 3.382E+06 | 3.224E+06 |
| 250 | 423.276 | 4.729 | 25 | 2.474E+06 | 1.987E+06 | 2.128E+06 | 2.161E+06 | 2.011E+06 | 2.178E+06 | 2.048E+06 | 2.209E+06 | 1.928E+06 | 2.071E+06 | 1.982E+06 | 2.144E+06 | 2.049E+06 | 2.923E+06 | 2.876E+06 |
| 275 | 401.261 | 4.663 | 25 | 1.933E+06 | 1.358E+06 | 1.551E+06 | 1.913E+06 | 1.587E+06 | 1.733E+06 | 1.664E+06 | 1.638E+06 | 1.843E+06 | 1.579E+06 | 1.630E+06 | 1.722E+06 | 1.746E+06 | 2.313E+06 | 2.170E+06 |
| 307 | 379.249 | 4.593 | 25 | 1.691E+06 | 1.425E+06 | 1.722E+06 | 1.495E+06 | 1.329E+06 | 1.380E+06 | 1.304E+06 | 1.575E+06 | 1.364E+06 | 1.414E+06 | 1.342E+06 | 1.309E+06 | 1.521E+06 | 2.088E+06 | 1.746E+06 |
| 353 | 489.316 | 4.904 | 25 | 1.421E+06 | 1.309E+06 | 1.167E+06 | 1.163E+06 | 1.328E+06 | 1.132E+06 | 1.236E+06 | 1.213E+06 | 1.193E+06 | 1.253E+06 | 1.177E+06 | 1.093E+06 | 1.422E+06 | 2.399E+06 | 2.010E+06 |
| 358 | 505.391 | 10.706 | 25 | 1.820E+06 | 1.918E+06 | 1.856E+06 | 1.363E+06 | 1.355E+06 | 1.442E+06 | 1.331E+06 | 1.244E+06 | 1.498E+06 | 2.585E+06 | 1.695E+06 | 1.813E+06 | 1.612E+06 | 1.393E+06 | 1.496E+06 |
| 402 | 527.403 | 10.694 | 25 | 1.790E+06 | 1.935E+06 | 1.732E+06 | 1.178E+06 | 1.107E+06 | 1.205E+06 | 1.040E+06 | 1.096E+06 | 1.371E+06 | 2.587E+06 | 1.508E+06 | 1.603E+06 | 1.464E+06 | 1.194E+06 | 1.415E+06 |
| 459 | 549.417 | 10.679 | 25 | 1.436E+06 | 1.569E+06 | 1.299E+06 | 5.280E+05 | 5.841E+05 | 6.997E+05 | 4.868E+05 | 5.905E+05 | 8.916E+05 | 2.180E+06 | 1.172E+06 | 1.184E+06 | 1.123E+06 | 7.901E+05 | 1.054E+06 |
| 535 | 463.345 | 9.103 | 25 | 1.100E+06 | 1.990E+06 | 4.130E+05 | 2.042E+06 | 6.504E+05 | 5.958E+05 | 1.241E+06 | 9.236E+05 | 1.574E+06 | 4.167E+05 | 5.573E+05 | 2.680E+05 | 6.507E+05 | 3.836E+05 | 2.431E+05 |
| 540 | 441.330 | 9.123 | 25 | 1.077E+06 | 1.746E+06 | 5.324E+05 | 2.392E+06 | 6.833E+05 | 4.551E+05 | 1.350E+06 | 9.144E+05 | 1.853E+06 | 6.585E+05 | 4.044E+05 | 5.756E+05 | 8.147E+05 | 1.079E+05 | 3.843E+05 |
| 638 | 529.383 | 9.048 | 25 | 1.056E+06 | 1.579E+06 | 1.075E+06 | 9.128E+05 | 2.541E+05 | 2.440E+05 | 2.019E+05 | 4.568E+05 | 8.627E+05 | 2.670E+05 | 4.603E+05 | 2.377E+05 | 7.032E+05 | 4.077E+05 | 2.300E+05 |
| 653 | 507.370 | 9.067 | 25 | 7.657E+05 | 1.800E+06 | 8.991E+05 | 1.020E+06 | 3.582E+05 | 3.181E+05 | 7.787E+05 | 5.917E+05 | 8.923E+05 | 2.615E+05 | 3.979E+05 | 5.342E+05 | 6.774E+05 | 3.723E+05 | 2.893E+05 |
| 662 | 511.328 | 4.955 | 25 | 6.924E+05 | 3.712E+05 | 3.289E+05 | 4.331E+05 | 5.238E+05 | 6.050E+05 | 2.698E+05 | 6.021E+05 | 4.381E+05 | 3.674E+05 | 5.179E+05 | 4.903E+05 | 7.084E+05 | 1.130E+06 | 8.380E+05 |
| 759 | 571.429 | 10.663 | 25 | 1.054E+06 | 1.185E+06 | 1.014E+06 | 3.352E+05 | 4.931E+05 | 4.133E+05 | 2.027E+05 | 3.083E+05 | 4.336E+05 | 1.726E+06 | 8.607E+05 | 8.824E+05 | 8.028E+05 | 2.064E+05 | 6.192E+05 |
| 859 | 533.341 | 5.004 | 25 | 1.524E+05 | 2.478E+05 | 2.360E+05 | 2.118E+05 | 7.883E+04 | 2.908E+05 | 1.467E+05 | 2.177E+05 | 1.691E+05 | 2.027E+05 | 1.041E+05 | 2.438E+05 | 3.090E+05 | 6.516E+05 | 5.508E+05 |
| 9 | 460.270 | 7.525 | 24 | 1.578E+08 | 1.958E+08 | 1.665E+08 | 1.540E+08 | 1.983E+08 | 1.704E+08 | 1.645E+08 | 1.889E+08 | 1.832E+08 | 2.397E+08 | 2.105E+08 | 1.639E+08 | 2.202E+08 | 1.913E+08 | 2.291E+08 |
| 25 | 846.443 | 7.526 | 24 | 2.292E+07 | 5.448E+07 | 2.343E+07 | 1.970E+07 | 4.435E+07 | 2.161E+07 | 2.460E+07 | 3.724E+07 | 2.523E+07 | 6.142E+07 | 5.721E+07 | 1.345E+07 | 4.470E+07 | 3.350E+07 | 6.986E+07 |
| 67 | 446.254 | 7.523 | 24 | 1.118E+07 | 1.337E+07 | 1.159E+07 | 1.097E+07 | 1.453E+07 | 1.200E+07 | 1.218E+07 | 1.366E+07 | 1.286E+07 | 1.704E+07 | 1.609E+07 | 1.058E+07 | 1.439E+07 | 1.300E+07 | 1.632E+07 |
| 81 | 460.270 | 7.249 | 24 | 3.435E+06 | 1.021E+07 | 3.981E+06 | 6.796E+06 | 9.153E+06 | 8.153E+06 | 4.616E+06 | 1.851E+07 | 6.543E+06 | 1.106E+07 | 1.420E+07 | 4.296E+06 | 5.303E+06 | 7.646E+06 | 1.587E+07 |
| 139 | 476.266 | 7.526 | 24 | 6.438E+06 | 7.690E+06 | 7.074E+06 | 5.288E+06 | 7.093E+06 | 6.801E+06 | 5.799E+06 | 6.227E+06 | 5.653E+06 | 7.634E+06 | 6.494E+06 | 4.838E+06 | 7.164E+06 | 5.234E+06 | 7.461E+06 |
| 261 | 397.201 | 7.526 | 24 | 1.936E+06 | 3.746E+06 | 2.524E+06 | 1.771E+06 | 3.218E+06 | 2.571E+06 | 2.319E+06 | 2.637E+06 | 2.510E+06 | 3.729E+06 | 4.114E+06 | 1.954E+06 | 3.514E+06 | 3.260E+06 | 4.003E+06 |
| 272 | 429.191 | 6.884 | 24 | 7.271E+06 | 8.517E+06 | 8.198E+06 | 5.185E+06 | 5.800E+06 | 6.903E+06 | 6.614E+06 | 7.536E+06 | 8.076E+06 | 8.081E+06 | 8.628E+06 | 7.040E+06 | 7.731E+06 | 1.041E+07 | 8.570E+06 |
| 316 | 531.275 | 7.525 | 24 | 1.514E+06 | 4.006E+06 | 1.344E+06 | 1.197E+06 | 3.227E+06 | 1.013E+06 | 1.224E+06 | 2.670E+06 | 1.961E+06 | 4.684E+06 | 3.365E+06 | 5.113E+05 | 3.019E+06 | 1.907E+06 | 5.152E+06 |
| 375 | 462.213 | 6.365 | 24 | 2.660E+06 | 2.628E+06 | 2.362E+06 | 1.240E+06 | 1.270E+06 | 1.135E+06 | 1.435E+06 | 2.102E+06 | 1.690E+06 | 2.182E+06 | 2.873E+06 | 1.481E+06 | 2.216E+06 | 2.543E+06 | 2.018E+06 |
| 418 | 445.186 | 6.364 | 24 | 2.159E+06 | 2.200E+06 | 2.064E+06 | 9.785E+05 | 9.812E+05 | 9.986E+05 | 1.162E+06 | 1.672E+06 | 3.292E+06 | 1.783E+06 | 2.505E+06 | 1.308E+06 | 1.972E+06 | 2.031E+06 | 3.603E+06 |
| 419 | 460.270 | 7.814 | 24 | 3.627E+06 | 2.248E+06 | 1.836E+06 | 1.604E+06 | 1.915E+06 | 1.635E+06 | 1.304E+06 | 3.508E+06 | 1.666E+06 | 3.356E+06 | 3.077E+06 | 3.009E+06 | 3.413E+06 | 1.994E+06 | 4.171E+06 |
| 447 | 450.204 | 8.853 | 24 | 1.247E+06 | 1.089E+06 | 7.144E+05 | 8.742E+05 | 8.287E+05 | 8.219E+05 | 8.992E+05 | 8.188E+05 | 6.990E+05 | 1.502E+06 | 1.454E+06 | 9.737E+05 | 1.522E+06 | 1.446E+06 | 1.147E+06 |
| 469 | 445.186 | 6.211 | 24 | 1.019E+06 | 1.037E+06 | 9.994E+05 | 4.545E+05 | 5.128E+05 | 4.813E+05 | 5.590E+05 | 9.571E+05 | 5.103E+05 | 9.389E+05 | 1.206E+06 | 6.794E+05 | 1.074E+06 | 9.980E+05 | 6.768E+05 |
| 578 | 448.233 | 6.302 | 24 | 6.869E+05 | 5.944E+05 | 6.661E+05 | 4.948E+05 | 3.770E+05 | 2.402E+05 | 4.509E+05 | 5.045E+05 | 4.804E+05 | 4.949E+05 | 5.630E+05 | 5.015E+05 | 6.319E+05 | 6.885E+05 | 3.746E+05 |
| 582 | 448.233 | 6.159 | 24 | 7.412E+05 | 5.813E+05 | 6.434E+05 | 4.642E+05 | 5.031E+05 | 3.595E+05 | 3.787E+05 | 5.004E+05 | 4.280E+05 | 5.878E+05 | 6.805E+05 | 3.818E+05 | 6.626E+05 | 5.995E+05 | 4.113E+05 |
| 636 | 462.213 | 6.211 | 24 | 5.689E+05 | 6.338E+05 | 5.791E+05 | 2.079E+05 | 3.927E+05 | 1.994E+05 | 3.973E+05 | 4.173E+05 | 4.394E+05 | 5.796E+05 | 7.947E+05 | 3.956E+05 | 6.949E+05 | 6.137E+05 | 4.912E+05 |
| 637 | 431.207 | 6.148 | 24 | 6.969E+05 | 5.930E+05 | 6.210E+05 | 4.377E+05 | 3.148E+05 | 3.293E+05 | 4.340E+05 | 5.028E+05 | 3.637E+05 | 5.163E+05 | 6.158E+05 | 4.851E+05 | 5.158E+05 | 5.598E+05 | 4.296E+05 |
| 770 | 474.249 | 6.864 | 24 | 8.382E+05 | 7.154E+05 | 1.481E+06 | 3.374E+05 | 6.222E+05 | 5.790E+05 | 5.546E+05 | 5.260E+05 | 3.202E+05 | 8.711E+05 | 1.343E+06 | 5.558E+05 | 8.728E+05 | 7.666E+05 | 8.137E+05 |
| 790 | 431.207 | 6.021 | 24 | 3.784E+05 | 3.853E+05 | 4.746E+05 | 2.425E+05 | 2.993E+05 | 1.843E+05 | 2.143E+05 | 3.896E+05 | 2.973E+05 | 3.702E+05 | 2.980E+05 | 3.751E+05 | 4.825E+05 | 4.460E+05 | 3.734E+05 |
| 221 | 371.326 | 7.517 | 23 | 4.667E+06 | 4.803E+06 | 4.057E+06 | 2.296E+06 | 1.762E+06 | 2.400E+06 | 2.053E+06 | 2.241E+06 | 2.690E+06 | 5.184E+06 | 4.493E+06 | 3.211E+06 | 2.365E+06 | 1.917E+06 | 1.853E+06 |
| 31 | 328.919 | 12.753 | 22 | 6.648E+07 | 5.840E+07 | 5.778E+07 | 5.484E+07 | 5.595E+07 | 5.468E+07 | 5.184E+07 | 4.909E+07 | 5.203E+07 | 5.222E+07 | 4.960E+07 | 4.515E+07 | 4.586E+07 | 4.300E+07 | 4.477E+07 |
| 39 | 312.943 | 12.757 | 22 | 2.439E+07 | 4.238E+07 | 1.235E+07 | 1.478E+07 | 3.293E+07 | 3.867E+07 | 4.051E+07 | 4.013E+07 | 3.622E+07 | 4.005E+07 | 8.906E+06 | 1.210E+07 | 3.716E+07 | 3.596E+07 | 3.736E+07 |
| 70 | 287.888 | 12.760 | 22 | 3.159E+07 | 2.653E+07 | 2.802E+07 | 2.655E+07 | 2.597E+07 | 2.675E+07 | 2.614E+07 | 2.374E+07 | 2.429E+07 | 2.383E+07 | 2.307E+07 | 2.375E+07 | 2.274E+07 | 2.304E+07 | 2.231E+07 |
| 92 | 313.078 | 6.460 | 22 | 0.000E+00 | 0.000E+00 | 0.000E+00 | 5.992E+06 | 9.146E+06 | 6.921E+06 | 0.000E+00 | 0.000E+00 | 1.094E+06 | 0.000E+00 | 0.000E+00 | 0.000E+00 | 0.000E+00 | 0.000E+00 | 0.000E+00 |
| 187 | 312.943 | 12.757 | 22 | 4.381E+07 | 4.238E+07 | 4.025E+07 | 4.257E+07 | 4.018E+07 | 3.867E+07 | 4.051E+07 | 4.013E+07 | 3.999E+07 | 4.005E+07 | 3.896E+07 | 3.631E+07 | 3.716E+07 | 1.322E+07 | 3.736E+07 |
| 196 | 313.078 | 6.532 | 22 | 0.000E+00 | 0.000E+00 | 0.000E+00 | 2.949E+06 | 4.880E+06 | 3.674E+06 | 0.000E+00 | 0.000E+00 | 7.935E+05 | 0.000E+00 | 0.000E+00 | 0.000E+00 | 0.000E+00 | 0.000E+00 | 0.000E+00 |
| 255 | 313.235 | 8.431 | 22 | 2.802E+06 | 3.294E+06 | 2.140E+06 | 3.596E+06 | 3.295E+06 | 3.095E+06 | 3.299E+06 | 3.595E+06 | 3.899E+06 | 2.762E+06 | 3.376E+06 | 2.417E+06 | 3.185E+06 | 3.598E+06 | 3.821E+06 |
| 558 | 313.235 | 8.075 | 22 | 1.099E+05 | 6.039E+04 | 1.602E+05 | 2.358E+06 | 3.429E+05 | 3.300E+05 | 4.281E+06 | 2.095E+07 | 5.840E+07 | 5.431E+06 | 5.374E+06 | 6.649E+05 | 2.980E+06 | 1.659E+07 | 1.289E+07 |
| 197 | 272.258 | 8.673 | 21 | 3.247E+05 | 7.275E+05 | 7.799E+05 | 4.866E+06 | 4.379E+06 | 3.844E+06 | 7.239E+05 | 1.597E+06 | 9.964E+06 | 1.986E+05 | 1.174E+05 | 1.418E+05 | 1.898E+05 | 1.609E+05 | 1.168E+05 |
| 223 | 300.290 | 9.476 | 21 | 4.838E+05 | 2.721E+05 | 5.694E+05 | 2.633E+06 | 2.290E+06 | 2.717E+06 | 4.999E+05 | 6.757E+05 | 4.904E+06 | 1.090E+06 | 5.515E+05 | 7.865E+05 | 5.908E+05 | 5.939E+05 | 5.695E+05 |

| Feature identifiers | | | | XIC | | | | | | | | | | | | | | |
|---|---|---|---|---|---|---|---|---|---|---|---|---|---|---|---|---|---|---|
| ClusterIdx | m/z | RT [min] | Subnetwork | 4TUB34_I | 4TUB34_II | 4TUB34_III | 4TUB81_I | 4TUB81_II | 4TUB81_III | 4TUB93_I | 4TUB93_II | 4TUB93_III | 5TUB23_I | 5TUB23_II | 5TUB23_III | 5TUB48_I | 5TUB48_II | 5TUB48_III |
| 460 | 406.354 | 10.232 | 21 | 6.215E+05 | 6.644E+05 | 5.990E+05 | 2.894E+05 | 4.839E+05 | 5.454E+05 | 1.953E+05 | 5.907E+05 | 6.624E+05 | 8.242E+05 | 1.158E+06 | 4.583E+05 | 9.072E+05 | 1.032E+06 | 1.084E+06 |
| 157 | 429.146 | 4.153 | 20 | 1.240E+04 | 7.352E+04 | 1.429E+05 | 8.212E+06 | 8.503E+06 | 6.479E+06 | 0.000E+00 | 0.000E+00 | 1.874E+06 | 0.000E+00 | 0.000E+00 | 2.143E+04 | 0.000E+00 | 0.000E+00 | 0.000E+00 |
| 166 | 549.571 | 3.384 | 20 | 1.524E+04 | 0.000E+00 | 0.000E+00 | 2.038E+06 | 3.926E+06 | 8.390E+06 | 1.934E+05 | 0.000E+00 | 7.436E+04 | 0.000E+00 | 0.000E+00 | 0.000E+00 | 0.000E+00 | 0.000E+00 | 0.000E+00 |
| 481 | 549.078 | 3.211 | 20 | 1.576E+06 | 3.111E+04 | 3.970E+04 | 4.032E+05 | 8.146E+04 | 3.677E+05 | 4.687E+07 | 2.738E+06 | 3.420E+04 | 4.257E+05 | 7.242E+05 | 5.473E+05 | 3.397E+04 | 9.688E+05 | 5.240E+05 |
| 520 | 432.026 | 3.379 | 20 | 7.166E+07 | 6.464E+07 | 5.635E+07 | 3.111E+05 | 9.032E+05 | 9.175E+05 | 2.266E+07 | 1.012E+08 | 2.259E+07 | 6.001E+07 | 9.363E+07 | 1.647E+06 | 3.266E+07 | 7.102E+07 | 2.829E+06 |
| 584 | 432.026 | 3.502 | 20 | 1.080E+08 | 1.633E+08 | 1.834E+08 | 7.585E+05 | 1.097E+06 | 7.837E+05 | 5.566E+06 | 2.010E+08 | 9.198E+05 | 1.300E+07 | 6.514E+06 | 8.611E+06 | 1.382E+08 | 2.281E+06 | 1.273E+07 |
| 596 | 405.571 | 3.038 | 20 | 3.263E+06 | 6.151E+05 | 6.213E+07 | 4.460E+04 | 4.063E+04 | 3.442E+04 | 2.109E+05 | 1.015E+08 | 1.945E+05 | 2.139E+07 | 4.326E+07 | 2.283E+05 | 7.490E+07 | 1.390E+06 | 1.593E+05 |
| 600 | 359.104 | 0.595 | 20 | 6.372E+04 | 3.285E+06 | 3.158E+06 | 1.104E+07 | 4.912E+06 | 1.601E+06 | 4.016E+05 | 5.801E+06 | 2.248E+07 | 3.516E+05 | 1.502E+05 | 1.253E+07 | 6.092E+03 | 1.361E+06 | 3.580E+05 |
| 611 | 401.115 | 2.454 | 20 | 4.051E+07 | 1.181E+07 | 2.267E+07 | 1.724E+05 | 6.740E+04 | 2.454E+05 | 7.257E+04 | 2.018E+07 | 1.491E+08 | 2.688E+05 | 1.617E+05 | 3.068E+04 | 2.441E+05 | 5.088E+06 | 1.785E+04 |
| 622 | 411.021 | 3.222 | 20 | 6.908E+05 | 2.343E+07 | 1.268E+07 | 3.728E+04 | 5.506E+04 | 1.522E+05 | 3.992E+05 | 1.462E+07 | 8.525E+05 | 7.326E+05 | 1.096E+06 | 1.132E+07 | 1.817E+05 | 3.270E+05 | 3.653E+06 |
| 315 | 515.353 | 9.549 | 19 | 1.693E+04 | 7.226E+03 | 1.207E+04 | 5.364E+06 | 2.352E+06 | 1.547E+06 | 6.965E+03 | 4.657E+04 | 2.547E+06 | 0.000E+00 | 0.000E+00 | 5.113E+03 | 0.000E+00 | 0.000E+00 | 0.000E+00 |
| 326 | 321.148 | 3.682 | 19 | 2.491E+06 | 5.592E+06 | 4.032E+06 | 3.091E+06 | 2.864E+06 | 2.976E+06 | 2.724E+06 | 2.341E+07 | 4.886E+07 | 9.297E+04 | 0.000E+00 | 0.000E+00 | 8.366E+03 | 4.879E+04 | 0.000E+00 |
| 595 | 366.206 | 3.688 | 19 | 9.108E+05 | 1.695E+06 | 1.280E+06 | 5.866E+05 | 6.606E+05 | 1.168E+06 | 9.908E+05 | 6.797E+06 | 1.604E+07 | 1.036E+04 | 1.501E+04 | 0.000E+00 | 3.991E+04 | 0.000E+00 | 0.000E+00 |
| 494 | 453.189 | 7.153 | 18 | 8.727E+06 | 8.814E+06 | 7.632E+06 | 5.776E+06 | 6.379E+06 | 9.000E+06 | 8.541E+06 | 1.109E+07 | 9.768E+06 | 8.172E+06 | 9.746E+06 | 5.291E+06 | 7.610E+06 | 1.013E+07 | 9.621E+06 |
| 591 | 453.189 | 7.367 | 18 | 5.129E+06 | 4.782E+06 | 3.495E+06 | 2.875E+06 | 3.205E+06 | 3.462E+06 | 3.696E+06 | 4.651E+06 | 4.430E+06 | 4.121E+06 | 4.724E+06 | 2.504E+06 | 5.201E+06 | 5.713E+06 | 4.621E+06 |
| 861 | 453.187 | 7.297 | 18 | 5.129E+06 | 4.365E+06 | 2.610E+06 | 1.069E+05 | 2.185E+05 | 1.168E+05 | 2.677E+05 | 3.128E+05 | 9.158E+04 | 4.088E+05 | 1.858E+05 | 2.299E+05 | 1.254E+05 | 3.434E+05 | 2.070E+07 |
| 21 | 453.169 | 7.511 | 17 | 1.545E+06 | 9.865E+05 | 1.172E+06 | 8.245E+07 | 5.813E+07 | 8.547E+07 | 5.082E+07 | 8.360E+07 | 8.589E+07 | 5.967E+07 | 9.504E+07 | 9.444E+05 | 8.520E+07 | 5.184E+07 | 1.692E+06 |
| 292 | 453.168 | 7.525 | 17 | 8.672E+07 | 9.109E+07 | 8.744E+07 | 8.245E+07 | 9.234E+07 | 7.965E+05 | 8.805E+07 | 1.020E+08 | 8.495E+07 | 9.386E+07 | 9.556E+05 | 8.739E+07 | 9.210E+07 | 5.184E+07 | 9.246E+07 |
| 640 | 453.186 | 7.468 | 17 | 1.262E+06 | 1.251E+06 | 8.872E+05 | 8.557E+07 | 5.828E+05 | 1.131E+06 | 1.023E+06 | 1.004E+06 | 1.240E+06 | 7.642E+05 | 9.893E+05 | 5.887E+05 | 6.516E+05 | 7.752E+05 | 1.407E+06 |
| 59 | 281.138 | 7.525 | 16 | 1.167E+07 | 1.565E+07 | 1.214E+07 | 1.032E+07 | 1.533E+07 | 1.140E+07 | 1.143E+07 | 1.358E+07 | 1.151E+07 | 1.953E+07 | 1.760E+07 | 8.804E+06 | 1.645E+07 | 1.299E+07 | 1.894E+07 |
| 97 | 299.149 | 4.461 | 16 | 1.377E+07 | 8.804E+06 | 7.773E+06 | 7.532E+06 | 1.037E+07 | 8.322E+06 | 7.309E+06 | 1.665E+07 | 7.408E+06 | 1.598E+07 | 1.631E+07 | 9.149E+06 | 1.717E+07 | 1.243E+07 | 2.027E+07 |
| 172 | 299.148 | 7.523 | 16 | 5.198E+06 | 8.530E+06 | 5.708E+06 | 5.539E+06 | 7.403E+06 | 4.880E+06 | 5.615E+06 | 6.612E+06 | 5.217E+06 | 9.837E+06 | 7.874E+06 | 4.195E+06 | 7.569E+06 | 6.048E+06 | 1.033E+07 |
| 296 | 344.206 | 4.461 | 16 | 2.609E+06 | 1.903E+06 | 1.683E+06 | 1.907E+06 | 2.646E+06 | 2.235E+06 | 1.772E+06 | 3.473E+06 | 2.138E+06 | 4.322E+06 | 4.345E+06 | 2.396E+06 | 4.297E+06 | 3.092E+06 | 5.280E+06 |
| 381 | 316.176 | 4.464 | 16 | 1.929E+06 | 1.268E+06 | 1.051E+06 | 1.250E+06 | 1.596E+06 | 1.182E+06 | 1.093E+06 | 2.395E+06 | 1.102E+06 | 2.731E+06 | 2.603E+06 | 1.240E+06 | 2.512E+06 | 1.789E+06 | 2.974E+06 |
| 42 | 473.318 | 10.647 | 15 | 4.263E+06 | 5.335E+06 | 4.700E+06 | 6.559E+06 | 6.570E+06 | 8.083E+06 | 6.749E+06 | 7.575E+06 | 3.498E+06 | 5.174E+06 | 5.914E+06 | 7.044E+06 | 6.873E+06 | 7.499E+06 | 4.164E+06 |
| 72 | 680.481 | 12.424 | 15 | 1.838E+07 | 2.205E+07 | 2.257E+07 | 1.595E+07 | 1.589E+07 | 1.661E+07 | 1.622E+07 | 1.540E+07 | 1.970E+07 | 3.911E+07 | 4.577E+07 | 5.842E+07 | 4.691E+07 | 8.728E+07 | 3.487E+07 |
| 74 | 475.298 | 9.832 | 15 | 2.263E+07 | 1.046E+07 | 1.170E+07 | 1.467E+07 | 1.434E+07 | 1.716E+07 | 2.069E+07 | 2.404E+07 | 9.222E+06 | 6.098E+07 | 5.465E+07 | 6.507E+07 | 4.973E+07 | 7.452E+07 | 1.969E+07 |
| 95 | 492.324 | 9.832 | 15 | 2.010E+07 | 9.400E+06 | 1.076E+07 | 1.333E+07 | 1.308E+07 | 1.520E+07 | 1.818E+07 | 2.104E+07 | 8.238E+06 | 5.509E+07 | 4.803E+07 | 5.787E+07 | 4.485E+07 | 7.147E+07 | 1.776E+07 |
| 96 | 654.333 | 10.021 | 15 | 4.440E+07 | 8.683E+06 | 1.302E+07 | 6.968E+06 | 6.339E+06 | 4.954E+06 | 2.985E+06 | 4.309E+07 | 2.803E+06 | 1.432E+07 | 1.222E+07 | 3.937E+07 | 5.519E+07 | 1.247E+07 | 3.356E+07 |
| 105 | 490.345 | 10.647 | 15 | 1.960E+06 | 2.636E+06 | 2.273E+06 | 3.204E+06 | 3.187E+06 | 3.610E+06 | 3.185E+06 | 3.346E+06 | 1.573E+06 | 1.984E+06 | 2.632E+06 | 3.245E+06 | 2.562E+06 | 3.481E+06 | 2.034E+06 |
| 129 | 663.454 | 12.425 | 15 | 1.129E+07 | 1.391E+07 | 1.386E+07 | 1.019E+07 | 1.017E+07 | 1.039E+07 | 1.082E+07 | 9.802E+06 | 1.241E+07 | 2.407E+07 | 2.855E+07 | 3.761E+07 | 2.996E+07 | 5.738E+07 | 2.114E+07 |
| 141 | 708.511 | 12.424 | 15 | 1.014E+07 | 1.215E+07 | 1.273E+07 | 9.697E+06 | 9.646E+06 | 1.060E+07 | 1.143E+07 | 1.048E+07 | 1.371E+07 | 2.306E+07 | 2.576E+07 | 2.993E+07 | 2.770E+07 | 3.956E+07 | 2.376E+07 |
| 181 | 637.306 | 10.020 | 15 | 2.519E+07 | 5.148E+06 | 7.163E+06 | 4.014E+06 | 3.822E+06 | 2.863E+06 | 1.731E+06 | 2.523E+07 | 1.512E+06 | 8.325E+06 | 7.288E+06 | 2.257E+07 | 2.952E+07 | 4.856E+06 | 1.819E+07 |
| 205 | 518.376 | 10.647 | 15 | 1.188E+06 | 9.080E+05 | 1.121E+06 | 1.570E+06 | 1.780E+06 | 2.034E+06 | 1.911E+06 | 2.099E+06 | 7.780E+05 | 1.239E+06 | 1.481E+06 | 1.926E+06 | 1.575E+06 | 1.949E+06 | 1.240E+06 |
| 273 | 287.141 | 6.170 | 15 | 2.833E+06 | 1.828E+06 | 2.137E+06 | 2.537E+06 | 2.205E+06 | 3.157E+06 | 3.471E+06 | 4.298E+06 | 2.400E+06 | 4.036E+06 | 3.998E+06 | 4.611E+06 | 4.606E+06 | 5.546E+06 | 4.725E+06 |
| 298 | 515.329 | 10.526 | 15 | 1.823E+06 | 1.997E+06 | 1.702E+06 | 1.120E+06 | 1.085E+06 | 1.111E+06 | 8.804E+05 | 1.641E+06 | 1.645E+06 | 3.235E+06 | 3.232E+06 | 2.434E+06 | 3.393E+06 | 5.521E+06 | 3.639E+06 |
| 311 | 474.317 | 10.257 | 15 | 1.114E+06 | 1.699E+06 | 2.933E+06 | 1.902E+06 | 1.382E+06 | 1.210E+06 | 1.658E+06 | 1.340E+06 | 1.568E+06 | 2.879E+06 | 4.084E+06 | 3.292E+06 | 4.821E+06 | 5.939E+06 | 3.303E+06 |
| 317 | 457.288 | 10.396 | 15 | 1.035E+06 | 1.298E+06 | 1.532E+06 | 1.359E+06 | 1.227E+06 | 1.237E+06 | 1.368E+06 | 1.233E+06 | 1.044E+06 | 1.107E+06 | 1.859E+06 | 1.488E+06 | 2.606E+06 | 1.924E+06 | 1.812E+06 |
| 318 | 387.194 | 7.399 | 15 | 1.271E+07 | 1.833E+06 | 4.904E+06 | 2.020E+06 | 1.758E+06 | 1.684E+06 | 7.393E+05 | 4.222E+06 | 1.300E+06 | 2.476E+06 | 2.187E+06 | 6.693E+06 | 1.550E+07 | 2.050E+06 | 3.999E+06 |
| 426 | 682.364 | 10.020 | 15 | 6.783E+06 | 1.391E+06 | 2.098E+06 | 1.278E+06 | 9.531E+05 | 7.215E+05 | 6.008E+05 | 7.333E+06 | 5.525E+05 | 2.276E+06 | 1.862E+06 | 6.970E+06 | 1.346E+07 | 3.938E+06 | 8.722E+06 |
| 433 | 532.356 | 10.526 | 15 | 8.685E+05 | 8.874E+05 | 5.544E+05 | 5.238E+05 | 5.302E+05 | 5.439E+05 | 4.581E+05 | 7.981E+05 | 7.093E+05 | 1.528E+06 | 1.520E+06 | 1.018E+06 | 1.630E+06 | 2.621E+06 | 1.737E+06 |
| 452 | 519.377 | 10.257 | 15 | 5.300E+05 | 8.514E+05 | 1.105E+06 | 6.876E+05 | 5.052E+05 | 5.581E+05 | 5.485E+05 | 7.095E+05 | 8.024E+05 | 1.563E+06 | 1.956E+06 | 1.353E+06 | 1.798E+06 | 1.880E+06 | 1.661E+06 |
| 529 | 520.355 | 9.833 | 15 | 1.982E+06 | 4.608E+05 | 4.884E+05 | 1.681E+05 | 5.071E+05 | 7.321E+05 | 2.537E+06 | 2.543E+06 | 3.905E+05 | 1.382E+07 | 9.411E+06 | 1.277E+07 | 1.921E+07 | 3.377E+07 | 5.725E+06 |
| 726 | 1342.926 | 12.423 | 15 | 4.642E+04 | 1.800E+05 | 3.393E+05 | 2.099E+04 | 2.496E+04 | 6.941E+04 | 7.010E+04 | 5.631E+04 | 1.272E+05 | 1.048E+06 | 1.851E+06 | 3.277E+06 | 2.371E+06 | 7.709E+06 | 8.333E+05 |
| 755 | 949.587 | 9.831 | 15 | 3.424E+04 | 0.000E+00 | 2.644E+03 | 4.302E+03 | 0.000E+00 | 2.250E+04 | 2.777E+04 | 1.044E+05 | 0.000E+00 | 1.109E+06 | 8.006E+05 | 1.070E+06 | 8.165E+05 | 1.615E+06 | 6.757E+04 |
| 783 | 491.374 | 10.264 | 15 | 1.181E+06 | 8.664E+05 | 5.403E+05 | 3.340E+05 | 5.853E+05 | 4.590E+05 | 6.172E+05 | 5.186E+05 | 1.024E+06 | 1.046E+06 | 3.012E+05 | 2.768E+05 | 3.523E+05 | 1.535E+05 | 1.069E+05 |
| 849 | 1370.957 | 12.421 | 15 | 0.000E+00 | 3.966E+04 | 5.935E+03 | 0.000E+00 | 1.877E+04 | 0.000E+00 | 2.085E+04 | 9.846E+03 | 2.480E+04 | 6.977E+05 | 5.632E+05 | 9.312E+05 | 7.270E+05 | 1.242E+06 | 2.943E+05 |
| 585 | 834.093 | 3.504 | 14 | 1.191E+07 | 2.532E+06 | 2.398E+06 | 8.245E+04 | 1.925E+05 | 5.873E+05 | 1.170E+06 | 2.043E+07 | 4.197E+05 | 1.860E+05 | 8.003E+05 | 0.000E+00 | 1.290E+06 | 6.626E+05 | 5.052E+04 |
| 619 | 792.082 | 3.197 | 14 | 0.000E+00 | 3.070E+05 | 7.161E+04 | 4.985E+05 | 1.157E+05 | 2.854E+05 | 4.110E+05 | 2.356E+06 | 7.194E+05 | 0.000E+00 | 1.073E+05 | 6.655E+03 | 0.000E+00 | 1.488E+04 | 0.000E+00 |
| 714 | 542.297 | 4.609 | 13 | 0.000E+00 | 0.000E+00 | 0.000E+00 | 0.000E+00 | 0.000E+00 | 0.000E+00 | 0.000E+00 | 0.000E+00 | 0.000E+00 | 1.104E+06 | 3.834E+06 | 7.258E+06 | 2.477E+06 | 3.606E+06 | 9.312E+05 |
| 717 | 492.763 | 4.165 | 13 | 0.000E+00 | 0.000E+00 | 0.000E+00 | 0.000E+00 | 0.000E+00 | 0.000E+00 | 0.000E+00 | 0.000E+00 | 0.000E+00 | 1.174E+06 | 6.540E+06 | 5.466E+06 | 2.511E+06 | 4.201E+06 | 1.078E+06 |
| 51 | 247.034 | 5.943 | 12 | 4.314E+06 | 1.728E+07 | 9.497E+06 | 1.550E+07 | 1.763E+07 | 1.365E+07 | 1.045E+06 | 0.000E+00 | 7.616E+06 | 0.000E+00 | 0.000E+00 | 8.349E+04 | 0.000E+00 | 0.000E+00 | 0.000E+00 |
| 698 | 247.034 | 6.796 | 12 | 6.493E+05 | 0.000E+00 | 6.343E+05 | 0.000E+00 | 6.426E+03 | 3.862E+03 | 0.000E+00 | 0.000E+00 | 9.236E+04 | 6.481E+08 | 3.021E+08 | 3.166E+08 | 5.491E+04 | 4.160E+04 | 1.933E+05 |
| 718 | 291.023 | 6.253 | 12 | 0.000E+00 | 0.000E+00 | 0.000E+00 | 0.000E+00 | 0.000E+00 | 0.000E+00 | 0.000E+00 | 0.000E+00 | 0.000E+00 | 1.122E+07 | 7.405E+06 | 4.876E+06 | 0.000E+00 | 0.000E+00 | 0.000E+00 |
| 16 | 122.547 | 12.762 | 11 | 1.978E+08 | 2.083E+08 | 1.955E+08 | 1.874E+08 | 1.870E+08 | 1.796E+08 | 1.975E+08 | 2.033E+08 | 1.917E+08 | 2.044E+08 | 1.870E+08 | 1.663E+08 | 1.852E+08 | 1.659E+08 | 1.833E+08 |
| 60 | 122.547 | 14.272 | 11 | 1.193E+07 | 1.737E+06 | 8.846E+06 | 1.530E+07 | 4.033E+06 | 1.316E+07 | 1.593E+06 | 1.674E+06 | 1.948E+06 | 1.406E+06 | 9.965E+06 | 1.661E+06 | 4.651E+06 | 2.129E+06 | 1.556E+06 |
| 103 | 122.547 | 2.944 | 11 | 5.346E+06 | 1.419E+06 | 2.297E+06 | 2.891E+06 | 1.809E+06 | 8.754E+06 | 2.979E+06 | 1.934E+07 | 8.889E+05 | 5.029E+06 | 4.684E+06 | 9.497E+06 | 2.935E+06 | 2.787E+06 | 2.793E+06 |
| 113 | 122.547 | 2.472 | 11 | 3.964E+06 | 1.749E+06 | 5.968E+06 | 5.380E+06 | 4.959E+06 | 3.305E+06 | 7.992E+05 | 1.029E+06 | 2.487E+06 | 2.042E+06 | 3.681E+06 | 3.665E+06 | 6.064E+06 | 8.486E+05 | 2.325E+06 |
| 115 | 122.547 | 13.880 | 11 | 7.644E+06 | 9.807E+05 | 2.426E+06 | 8.480E+06 | 8.349E+06 | 4.692E+06 | 1.660E+06 | 5.008E+06 | 9.966E+06 | 2.178E+06 | 1.840E+06 | 2.475E+06 | 2.144E+06 | 5.482E+06 | 1.284E+07 |
| 120 | 122.547 | 2.832 | 11 | 3.851E+06 | 1.029E+07 | 1.925E+06 | 4.823E+06 | 3.898E+06 | 3.312E+06 | 2.315E+06 | 4.993E+06 | 9.541E+05 | 2.429E+06 | 2.456E+06 | 1.308E+06 | 1.700E+06 | 2.787E+06 | 1.035E+06 |
| 123 | 122.547 | 13.067 | 11 | 4.123E+06 | 1.026E+07 | 1.405E+06 | 1.762E+06 | 4.592E+06 | 2.796E+06 | 9.977E+05 | 2.151E+06 | 3.308E+06 | 4.073E+06 | 1.717E+06 | 3.733E+06 | 9.074E+05 | 6.078E+06 | 2.239E+06 |
| 125 | 122.547 | 2.720 | 11 | 5.364E+06 | 4.973E+06 | 6.931E+06 | 2.885E+05 | 4.810E+06 | 4.945E+06 | 1.403E+07 | 4.115E+06 | 1.658E+06 | 3.845E+06 | 2.456E+06 | 2.685E+06 | 2.713E+06 | 2.133E+06 | 2.303E+06 |
| 127 | 122.547 | 0.660 | 11 | 4.894E+06 | 6.709E+05 | 1.692E+06 | 9.504E+06 | 6.299E+06 | 8.920E+06 | 2.369E+07 | 6.784E+06 | 2.024E+07 | 4.635E+06 | 2.758E+06 | 1.289E+07 | 1.056E+07 | 7.073E+06 | 2.778E+06 |
| 135 | 122.547 | 13.843 | 11 | 5.502E+06 | 1.344E+07 | 5.362E+06 | 3.892E+06 | 8.634E+06 | 2.494E+06 | 4.339E+06 | 1.622E+07 | 5.365E+06 | 2.178E+06 | 2.545E+06 | 2.945E+06 | 2.739E+06 | 8.248E+06 | 2.593E+06 |
| 140 | 122.547 | 2.622 | 11 | 3.606E+06 | 1.022E+07 | 1.534E+06 | 1.998E+06 | 2.323E+06 | 1.873E+06 | 8.471E+05 | 2.807E+06 | 9.897E+05 | 7.197E+06 | 6.281E+05 | 1.877E+06 | 1.049E+06 | 2.173E+06 | 8.938E+05 |
| 145 | 122.547 | 2.598 | 11 | 3.606E+06 | 4.744E+06 | 7.838E+06 | 5.042E+06 | 2.481E+06 | 2.193E+06 | 1.760E+06 | 4.420E+06 | 3.501E+06 | 1.038E+06 | 6.479E+05 | 3.959E+06 | 7.487E+06 | 5.403E+06 | 5.041E+06 |
| 147 | 122.547 | 2.749 | 11 | 2.104E+06 | 1.663E+06 | 3.865E+06 | 1.659E+06 | 1.193E+06 | 1.476E+06 | 2.315E+06 | 1.567E+06 | 9.541E+05 | 2.629E+06 | 2.456E+06 | 4.176E+06 | 1.185E+07 | 2.133E+06 | 2.974E+06 |
| 151 | 122.547 | 13.172 | 11 | 7.100E+06 | 8.777E+06 | 3.980E+06 | 4.062E+06 | 3.153E+06 | 2.614E+06 | 8.457E+06 | 3.487E+06 | 5.404E+06 | 4.169E+06 | 3.827E+06 | 2.551E+06 | 2.475E+06 | 2.978E+06 | 2.239E+06 |
| 156 | 122.547 | 2.528 | 11 | 3.606E+06 | 4.072E+06 | 3.564E+06 | 1.289E+06 | 1.235E+06 | 2.359E+06 | 5.992E+06 | 5.929E+06 | 7.599E+06 | 8.085E+05 | 7.871E+05 | 9.279E+05 | 1.597E+06 | 2.709E+06 | 2.437E+06 |
| 160 | 122.547 | 2.262 | 11 | 2.577E+06 | 6.083E+06 | 1.957E+06 | 2.335E+06 | 1.084E+06 | 3.516E+06 | 3.800E+06 | 6.877E+06 | 4.455E+05 | 4.238E+06 | 3.420E+06 | 2.206E+06 | 3.161E+06 | 1.741E+06 | 3.231E+06 |

| Feature identifiers | | | | XIC | | | | | | | | | | | | | | |
|---|---|---|---|---|---|---|---|---|---|---|---|---|---|---|---|---|---|---|
| ClusterIdx | m/z | RT [min] | Subnetwork | 4TUB34_I | 4TUB34_II | 4TUB34_III | 4TUB81_I | 4TUB81_II | 4TUB81_III | 4TUB93_I | 4TUB93_II | 4TUB93_III | 5TUB23_I | 5TUB23_II | 5TUB23_III | 5TUB48_I | 5TUB48_II | 5TUB48_III |
| 169 | 122.547 | 13.121 | 11 | 2.741E+06 | 1.026E+07 | 8.884E+06 | 3.582E+06 | 4.057E+06 | 1.153E+06 | 2.447E+06 | 1.399E+06 | 4.904E+06 | 1.786E+06 | 4.018E+06 | 4.351E+06 | 2.475E+06 | 6.078E+06 | 2.239E+06 |
| 174 | 122.547 | 2.436 | 11 | 3.109E+06 | 3.609E+06 | 1.965E+06 | 3.283E+06 | 3.670E+06 | 1.901E+06 | 4.927E+06 | 5.893E+06 | 1.121E+06 | 8.085E+05 | 4.034E+06 | 4.109E+06 | 7.418E+06 | 5.472E+06 | 3.662E+06 |
| 177 | 122.547 | 13.217 | 11 | 1.889E+06 | 5.157E+06 | 5.203E+06 | 1.232E+06 | 4.254E+06 | 8.976E+05 | 1.567E+06 | 1.142E+06 | 1.489E+06 | 2.885E+06 | 3.756E+06 | 3.064E+06 | 1.560E+06 | 3.679E+05 | 3.685E+06 |
| 178 | 122.547 | 2.294 | 11 | 8.130E+06 | 2.975E+06 | 2.928E+06 | 8.638E+05 | 3.868E+06 | 1.399E+06 | 5.705E+06 | 9.617E+05 | 1.807E+06 | 1.876E+06 | 5.797E+06 | 4.056E+06 | 2.087E+06 | 2.624E+06 | 1.039E+06 |
| 183 | 122.547 | 1.983 | 11 | 9.968E+05 | 1.809E+06 | 4.025E+05 | 5.721E+05 | 1.315E+06 | 3.476E+06 | 2.760E+06 | 3.717E+06 | 8.132E+05 | 1.559E+06 | 7.845E+05 | 2.727E+06 | 2.661E+06 | 1.184E+06 | 6.480E+06 |
| 185 | 122.547 | 13.651 | 11 | 1.972E+06 | 1.595E+06 | 1.202E+06 | 1.373E+06 | 4.583E+06 | 1.863E+06 | 5.815E+06 | 2.623E+06 | 2.656E+06 | 1.352E+06 | 4.873E+05 | 2.954E+06 | 3.950E+06 | 4.215E+06 | 3.506E+06 |
| 189 | 122.547 | 0.617 | 11 | 2.066E+07 | 1.955E+07 | 1.440E+07 | 6.512E+06 | 7.448E+06 | 1.827E+06 | 8.063E+05 | 2.004E+07 | 5.780E+06 | 3.643E+06 | 1.490E+07 | 3.090E+06 | 1.271E+06 | 6.014E+06 | 1.672E+06 |
| 206 | 122.547 | 1.515 | 11 | 5.384E+05 | 1.052E+06 | 7.630E+05 | 3.404E+06 | 3.963E+06 | 1.353E+06 | 5.440E+05 | 1.472E+06 | 2.795E+06 | 8.601E+05 | 1.022E+06 | 2.938E+06 | 1.009E+06 | 7.161E+05 | 1.232E+06 |
| 207 | 122.547 | 2.189 | 11 | 3.273E+06 | 4.093E+06 | 2.704E+06 | 1.578E+06 | 5.162E+06 | 3.107E+06 | 1.883E+06 | 2.758E+06 | 4.455E+05 | 1.899E+06 | 5.460E+06 | 1.302E+06 | 2.379E+06 | 4.856E+06 | 1.806E+06 |
| 216 | 122.547 | 1.421 | 11 | 4.853E+06 | 1.426E+06 | 7.326E+05 | 9.693E+05 | 2.080E+06 | 1.053E+06 | 1.008E+06 | 3.137E+05 | 2.941E+06 | 1.235E+06 | 9.754E+05 | 1.893E+06 | 5.322E+05 | 1.495E+06 | 2.372E+06 |
| 220 | 122.547 | 2.347 | 11 | 2.160E+06 | 8.575E+06 | 1.319E+06 | 4.356E+06 | 1.482E+06 | 2.297E+06 | 3.334E+06 | 1.731E+06 | 8.881E+05 | 4.856E+06 | 1.368E+06 | 2.303E+06 | 1.348E+07 | 1.734E+06 | 5.672E+06 |
| 248 | 122.547 | 2.158 | 11 | 9.994E+05 | 7.553E+05 | 9.755E+05 | 3.569E+06 | 1.895E+06 | 3.896E+05 | 6.150E+06 | 1.952E+06 | 1.742E+06 | 1.883E+06 | 4.216E+06 | 3.452E+06 | 4.605E+06 | 1.886E+06 | 4.367E+06 |
| 587 | 122.547 | 2.681 | 11 | 1.191E+07 | 4.744E+06 | 3.071E+06 | 5.776E+06 | 1.966E+06 | 4.922E+05 | 2.315E+06 | 2.524E+06 | 9.389E+05 | 7.692E+05 | 2.456E+06 | 9.581E+05 | 2.713E+06 | 2.173E+06 | 7.813E+05 |
| 644 | 122.547 | 2.919 | 11 | 5.346E+06 | 1.291E+07 | 2.297E+06 | 8.458E+06 | 1.613E+06 | 1.786E+06 | 2.979E+06 | 3.043E+06 | 8.889E+05 | 5.029E+06 | 4.684E+06 | 2.040E+06 | 2.935E+06 | 5.077E+07 | 1.149E+06 |
| 645 | 122.547 | 2.389 | 11 | 1.169E+06 | 2.238E+06 | 9.091E+05 | 1.050E+06 | 1.327E+06 | 4.811E+05 | 3.334E+06 | 1.029E+06 | 4.145E+06 | 2.190E+06 | 3.594E+06 | 1.552E+06 | 5.132E+06 | 5.259E+06 | 1.039E+06 |
| 656 | 122.547 | 2.584 | 11 | 5.588E+06 | 1.786E+06 | 2.190E+06 | 6.280E+05 | 8.817E+05 | 2.857E+06 | 2.535E+06 | 7.489E+05 | 1.263E+06 | 7.700E+06 | 5.557E+06 | 1.192E+06 | 3.632E+06 | 1.458E+06 | 6.723E+05 |
| 657 | 122.547 | 2.792 | 11 | 3.851E+06 | 3.748E+06 | 3.408E+06 | 1.496E+06 | 4.396E+05 | 3.828E+06 | 2.315E+06 | 1.626E+06 | 8.576E+06 | 4.608E+06 | 2.456E+06 | 6.718E+05 | 2.713E+06 | 2.570E+06 | 9.452E+06 |
| 710 | 122.547 | 3.154 | 11 | 2.962E+06 | 7.147E+06 | 1.154E+07 | 1.026E+07 | 1.148E+07 | 5.665E+06 | 6.953E+06 | 5.040E+06 | 4.203E+06 | 1.500E+07 | 5.477E+06 | 1.038E+07 | 1.542E+07 | 4.530E+06 | 1.222E+07 |
| 715 | 122.547 | 3.094 | 11 | 2.962E+06 | 6.225E+06 | 1.154E+07 | 1.408E+06 | 2.008E+07 | 5.665E+06 | 6.953E+06 | 9.786E+06 | 4.203E+06 | 1.500E+07 | 9.111E+06 | 5.491E+06 | 6.802E+06 | 1.142E+06 | 4.647E+07 |
| 719 | 122.547 | 2.222 | 11 | 1.023E+06 | 1.338E+06 | 2.438E+06 | 1.578E+06 | 1.482E+06 | 4.233E+05 | 2.428E+06 | 2.124E+06 | 7.342E+05 | 2.503E+06 | 1.408E+06 | 2.662E+06 | 4.229E+06 | 1.854E+06 | 2.432E+06 |
| 773 | 122.547 | 13.630 | 11 | 4.258E+06 | 1.189E+06 | 4.983E+05 | 2.919E+06 | 7.003E+05 | 1.921E+06 | 2.229E+06 | 1.130E+06 | 2.526E+06 | 5.536E+06 | 5.626E+05 | 1.509E+06 | 1.073E+06 | 1.332E+06 | 2.001E+06 |
| 61 | 537.536 | 11.439 | 10 | 4.355E+07 | 5.497E+07 | 5.350E+07 | 5.303E+07 | 5.216E+07 | 5.162E+07 | 3.880E+07 | 4.872E+07 | 4.806E+07 | 4.872E+07 | 3.947E+07 | 4.267E+07 | 4.095E+07 | 5.037E+07 | 3.799E+07 |
| 109 | 565.567 | 11.404 | 10 | 5.024E+07 | 5.595E+07 | 5.327E+07 | 5.149E+07 | 4.802E+07 | 4.121E+07 | 3.955E+07 | 4.748E+07 | 4.820E+07 | 4.524E+07 | 4.613E+07 | 4.716E+07 | 4.626E+07 | 4.956E+07 | 4.827E+07 |
| 295 | 582.594 | 11.445 | 10 | 6.299E+06 | 6.110E+06 | 6.016E+06 | 6.358E+06 | 6.159E+06 | 6.288E+06 | 6.862E+06 | 6.908E+06 | 6.620E+06 | 7.127E+06 | 7.285E+06 | 7.030E+06 | 7.277E+06 | 6.657E+06 | 7.378E+06 |
| 816 | 593.598 | 10.672 | 10 | 5.132E+05 | 1.270E+06 | 1.699E+06 | 6.814E+05 | 5.067E+05 | 1.507E+06 | 1.287E+06 | 6.693E+05 | 5.213E+05 | 8.149E+05 | 1.255E+06 | 7.014E+05 | 1.735E+06 | 7.893E+05 | 7.252E+05 |
| 191 | 608.393 | 9.449 | 9 | 8.296E+06 | 1.152E+07 | 8.829E+06 | 9.771E+06 | 1.029E+07 | 7.511E+06 | 1.326E+07 | 9.286E+06 | 9.834E+06 | 5.046E+06 | 3.971E+06 | 4.148E+06 | 6.041E+06 | 5.301E+06 | 6.027E+06 |
| 313 | 638.404 | 9.642 | 9 | 2.051E+06 | 2.091E+06 | 1.818E+06 | 1.867E+06 | 1.812E+06 | 1.381E+06 | 2.303E+06 | 1.696E+06 | 1.434E+06 | 9.959E+05 | 7.626E+05 | 5.408E+05 | 1.033E+06 | 8.364E+05 | 7.409E+05 |
| 348 | 624.388 | 9.076 | 9 | 1.590E+06 | 1.651E+06 | 1.617E+06 | 1.358E+06 | 1.999E+06 | 1.414E+06 | 2.893E+06 | 1.716E+06 | 1.007E+06 | 9.171E+05 | 8.783E+05 | 7.562E+05 | 1.304E+06 | 9.596E+05 | 8.202E+05 |
| 15 | 824.976 | 3.461 | 8 | 8.487E+07 | 1.811E+08 | 1.839E+08 | 8.016E+07 | 9.843E+07 | 7.359E+07 | 1.922E+08 | 2.769E+07 | 2.147E+08 | 1.421E+08 | 6.926E+07 | 1.078E+08 | 2.143E+08 | 1.720E+08 | 5.759E+08 |
| 18 | 810.973 | 3.183 | 8 | 1.348E+08 | 2.937E+08 | 2.595E+08 | 1.056E+08 | 1.147E+08 | 8.287E+07 | 1.409E+08 | 2.249E+08 | 1.620E+08 | 1.269E+08 | 9.068E+07 | 3.449E+08 | 4.010E+08 | 4.998E+08 | 7.608E+08 |
| 26 | 838.980 | 3.758 | 8 | 4.081E+07 | 9.423E+07 | 1.211E+08 | 3.744E+07 | 4.725E+07 | 3.233E+07 | 1.848E+08 | 2.603E+06 | 1.989E+08 | 1.404E+08 | 5.844E+07 | 2.916E+07 | 1.065E+08 | 4.841E+07 | 3.747E+08 |
| 65 | 1215.954 | 3.181 | 8 | 2.250E+07 | 4.523E+07 | 4.562E+07 | 1.722E+07 | 1.932E+07 | 1.491E+07 | 2.398E+07 | 3.881E+07 | 2.882E+07 | 2.258E+07 | 1.724E+07 | 4.833E+07 | 7.387E+07 | 8.728E+07 | 1.251E+08 |
| 87 | 1236.959 | 3.462 | 8 | 1.655E+07 | 3.532E+07 | 3.640E+07 | 1.664E+07 | 1.817E+07 | 1.318E+07 | 3.554E+07 | 4.322E+06 | 4.404E+07 | 2.611E+07 | 1.289E+07 | 1.863E+07 | 3.850E+07 | 3.296E+07 | 1.013E+08 |
| 121 | 837.625 | 3.464 | 8 | 7.530E+06 | 1.905E+07 | 1.813E+07 | 8.955E+06 | 1.066E+07 | 8.196E+06 | 1.948E+07 | 2.596E+06 | 2.114E+07 | 1.363E+07 | 7.617E+06 | 1.174E+07 | 2.170E+07 | 1.785E+07 | 4.530E+07 |
| 136 | 823.621 | 3.183 | 8 | 1.437E+07 | 3.039E+07 | 2.602E+07 | 1.169E+07 | 1.328E+07 | 9.298E+06 | 1.519E+07 | 2.361E+07 | 1.783E+07 | 1.416E+07 | 1.075E+07 | 3.498E+07 | 4.062E+07 | 4.674E+07 | 6.263E+07 |
| 194 | 618.984 | 3.459 | 8 | 5.362E+06 | 1.182E+07 | 1.312E+07 | 4.642E+06 | 6.453E+06 | 4.869E+06 | 1.302E+07 | 1.393E+06 | 1.525E+07 | 7.512E+06 | 4.042E+06 | 7.076E+06 | 1.406E+07 | 1.076E+07 | 3.894E+07 |
| 204 | 1257.965 | 3.758 | 8 | 8.018E+06 | 1.846E+07 | 2.355E+07 | 7.473E+06 | 8.516E+06 | 6.572E+06 | 3.676E+07 | 1.804E+05 | 3.877E+07 | 2.747E+07 | 1.118E+07 | 4.507E+06 | 1.906E+07 | 1.010E+07 | 7.586E+07 |
| 240 | 852.983 | 4.057 | 8 | 7.066E+06 | 1.720E+07 | 2.588E+07 | 4.648E+06 | 5.656E+06 | 4.062E+06 | 5.058E+07 | 2.568E+05 | 4.773E+07 | 4.901E+07 | 1.740E+07 | 3.429E+06 | 2.038E+07 | 5.919E+06 | 9.408E+07 |
| 244 | 846.306 | 3.758 | 8 | 4.025E+06 | 8.991E+06 | 1.098E+07 | 3.404E+06 | 4.688E+06 | 3.166E+06 | 1.850E+07 | 1.252E+05 | 1.896E+07 | 1.422E+07 | 5.639E+06 | 3.179E+06 | 1.099E+07 | 4.822E+06 | 2.930E+07 |
| 253 | 1069.394 | 4.071 | 8 | 7.278E+06 | 1.598E+07 | 1.998E+07 | 6.585E+06 | 7.572E+06 | 4.336E+06 | 1.597E+07 | 2.373E+06 | 1.290E+07 | 2.047E+07 | 8.093E+06 | 2.593E+07 | 6.237E+07 | 5.481E+07 | 9.344E+07 |
| 257 | 851.628 | 3.759 | 8 | 4.019E+06 | 9.613E+06 | 1.193E+07 | 3.922E+06 | 4.963E+06 | 3.328E+06 | 1.757E+07 | 1.315E+05 | 1.750E+07 | 1.398E+07 | 6.025E+06 | 2.743E+06 | 1.051E+07 | 5.001E+06 | 3.129E+07 |
| 258 | 608.481 | 3.183 | 8 | 5.499E+06 | 1.358E+07 | 1.319E+07 | 4.417E+06 | 4.664E+06 | 3.286E+06 | 5.906E+06 | 1.022E+07 | 7.217E+06 | 5.102E+06 | 3.743E+06 | 1.852E+07 | 2.028E+07 | 2.498E+07 | 3.376E+07 |
| 288 | 1083.397 | 4.311 | 8 | 3.427E+06 | 9.289E+06 | 1.197E+07 | 2.092E+06 | 4.359E+06 | 2.720E+06 | 1.592E+07 | 2.052E+05 | 6.385E+06 | 5.776E+06 | 1.817E+07 | 1.600E+07 | 8.207E+07 | 4.978E+07 | 2.170E+08 |
| 289 | 629.487 | 3.760 | 8 | 2.249E+06 | 6.543E+06 | 8.310E+06 | 2.027E+06 | 3.447E+06 | 2.323E+06 | 1.521E+07 | 4.774E+04 | 1.503E+07 | 9.591E+06 | 2.992E+06 | 2.100E+06 | 7.516E+06 | 3.236E+06 | 2.804E+07 |
| 312 | 1055.391 | 3.820 | 8 | 7.085E+06 | 1.525E+07 | 1.517E+07 | 5.267E+06 | 5.482E+06 | 3.439E+06 | 6.986E+06 | 9.723E+06 | 7.443E+06 | 9.074E+06 | 4.811E+06 | 2.428E+07 | 4.981E+07 | 5.688E+07 | 7.781E+07 |
| 352 | 1234.935 | 3.184 | 8 | 2.272E+06 | 4.039E+06 | 3.261E+05 | 1.566E+06 | 2.293E+06 | 1.435E+06 | 2.342E+06 | 4.246E+06 | 3.037E+06 | 2.376E+06 | 2.019E+06 | 3.822E+06 | 7.208E+06 | 7.740E+06 | 1.034E+07 |
| 406 | 802.298 | 4.071 | 8 | 2.253E+06 | 6.334E+06 | 7.066E+06 | 2.399E+06 | 2.513E+06 | 1.664E+06 | 5.702E+06 | 7.482E+05 | 4.569E+06 | 7.589E+06 | 2.858E+06 | 1.024E+07 | 2.612E+07 | 2.147E+07 | 3.583E+07 |
| 445 | 812.801 | 4.312 | 8 | 1.468E+06 | 4.336E+06 | 5.314E+06 | 1.395E+06 | 8.759E+05 | 1.165E+06 | 7.187E+06 | 5.237E+04 | 5.550E+06 | 2.401E+07 | 8.955E+06 | 9.288E+06 | 4.307E+07 | 2.372E+07 | 1.095E+08 |
| 450 | 1041.387 | 3.604 | 8 | 1.419E+06 | 3.087E+06 | 3.567E+06 | 1.937E+06 | 2.567E+06 | 1.641E+06 | 5.097E+05 | 1.033E+07 | 1.466E+06 | 5.707E+05 | 3.744E+05 | 5.478E+06 | 8.543E+06 | 1.264E+07 | 9.400E+06 |
| 457 | 1255.942 | 3.463 | 8 | 1.405E+06 | 3.187E+06 | 2.466E+06 | 1.948E+06 | 2.363E+06 | 4.250E+05 | 4.131E+06 | 3.991E+05 | 4.565E+06 | 2.891E+06 | 1.268E+06 | 2.136E+06 | 3.671E+06 | 3.607E+06 | 6.151E+06 |
| 497 | 1278.970 | 4.056 | 8 | 1.534E+06 | 2.368E+06 | 4.874E+06 | 5.857E+05 | 7.551E+05 | 3.693E+05 | 1.099E+07 | 0.000E+00 | 9.895E+06 | 9.841E+06 | 3.583E+06 | 5.851E+05 | 3.337E+06 | 1.050E+06 | 1.931E+07 |
| 504 | 1268.955 | 3.761 | 8 | 1.359E+06 | 3.273E+06 | 1.954E+06 | 1.240E+06 | 1.430E+06 | 1.104E+06 | 5.047E+06 | 1.132E+04 | 5.984E+06 | 4.378E+06 | 1.986E+06 | 1.049E+06 | 3.448E+06 | 1.857E+06 | 1.075E+07 |
| 511 | 865.632 | 4.058 | 8 | 4.966E+05 | 1.542E+06 | 2.062E+06 | 3.893E+05 | 3.975E+05 | 3.093E+05 | 4.840E+06 | 0.000E+00 | 4.587E+06 | 4.548E+06 | 1.513E+06 | 1.434E+05 | 1.418E+06 | 4.041E+05 | 8.128E+06 |
| 514 | 639.989 | 4.058 | 8 | 5.746E+05 | 1.353E+06 | 2.083E+06 | 4.042E+05 | 2.935E+05 | 2.895E+05 | 4.477E+06 | 0.000E+00 | 4.568E+06 | 4.782E+06 | 1.429E+06 | 2.622E+05 | 2.017E+06 | 4.261E+05 | 8.847E+06 |
| 515 | 860.310 | 4.057 | 8 | 2.476E+05 | 1.439E+06 | 2.289E+06 | 3.375E+05 | 2.784E+05 | 6.332E+04 | 4.880E+06 | 0.000E+00 | 4.563E+06 | 5.055E+06 | 1.465E+06 | 2.059E+05 | 1.758E+06 | 2.775E+05 | 8.130E+06 |
| 518 | 1276.945 | 3.762 | 8 | 9.840E+05 | 2.446E+06 | 3.006E+06 | 1.133E+06 | 1.246E+06 | 5.752E+05 | 4.404E+06 | 8.877E+03 | 3.838E+06 | 3.451E+06 | 1.504E+06 | 5.378E+05 | 2.045E+06 | 9.330E+05 | 8.080E+06 |
| 519 | 856.952 | 3.759 | 8 | 1.072E+06 | 1.825E+06 | 2.799E+06 | 8.201E+05 | 9.779E+05 | 3.862E+05 | 4.246E+06 | 1.518E+04 | 4.020E+06 | 2.909E+06 | 6.594E+05 | 4.409E+05 | 1.991E+06 | 8.007E+05 | 6.423E+06 |
| 522 | 1090.726 | 4.311 | 8 | 3.648E+05 | 1.217E+06 | 1.715E+06 | 3.082E+05 | 6.641E+05 | 8.836E+04 | 2.167E+06 | 0.000E+00 | 1.816E+06 | 7.882E+06 | 2.799E+06 | 2.757E+06 | 1.105E+07 | 6.789E+06 | 2.470E+07 |
| 531 | 1076.722 | 4.075 | 8 | 4.386E+05 | 2.415E+06 | 2.978E+06 | 1.029E+06 | 9.682E+05 | 6.189E+05 | 2.238E+06 | 3.591E+05 | 1.564E+06 | 3.247E+06 | 1.118E+06 | 4.218E+06 | 1.001E+07 | 8.595E+06 | 1.433E+07 |
| 533 | 1096.048 | 4.311 | 8 | 3.006E+05 | 8.569E+05 | 2.970E+05 | 1.058E+05 | 4.148E+05 | 7.282E+04 | 1.310E+06 | 0.000E+00 | 1.293E+06 | 5.672E+06 | 2.024E+06 | 1.981E+06 | 7.895E+06 | 5.000E+06 | 1.743E+07 |
| 534 | 791.796 | 3.823 | 8 | 1.634E+06 | 4.987E+06 | 4.616E+06 | 1.485E+06 | 5.627E+05 | 6.665E+05 | 1.831E+06 | 2.819E+06 | 2.318E+06 | 2.174E+06 | 1.404E+06 | 8.378E+06 | 1.703E+07 | 1.645E+07 | 2.393E+07 |
| 537 | 810.973 | 3.375 | 8 | 7.033E+06 | 1.010E+07 | 4.254E+06 | 6.898E+04 | 5.185E+04 | 6.968E+04 | 1.753E+06 | 7.891E+04 | 2.434E+06 | 1.302E+07 | 7.353E+05 | 4.465E+06 | 3.027E+06 | 1.024E+07 | 5.124E+07 |
| 559 | 824.976 | 3.689 | 8 | 4.725E+06 | 7.954E+06 | 5.051E+06 | 5.313E+04 | 1.295E+05 | 1.899E+04 | 1.322E+06 | 0.000E+00 | 2.454E+06 | 1.651E+07 | 2.126E+06 | 1.091E+06 | 2.521E+06 | 4.342E+06 | 4.838E+07 |
| 561 | 1289.962 | 4.060 | 8 | 1.003E+05 | 5.319E+05 | 5.577E+05 | 1.345E+04 | 1.002E+05 | 6.214E+04 | 1.730E+06 | 0.000E+00 | 1.558E+06 | 1.847E+06 | 4.367E+05 | 2.270E+04 | 3.640E+05 | 1.264E+05 | 2.327E+06 |
| 567 | 1082.043 | 4.072 | 8 | 4.648E+05 | 1.321E+06 | 1.430E+06 | 6.081E+05 | 3.199E+05 | 3.375E+05 | 1.684E+06 | 8.236E+04 | 1.078E+06 | 1.948E+06 | 7.819E+05 | 2.737E+06 | 6.797E+06 | 5.903E+06 | 8.935E+06 |
| 588 | 1027.384 | 3.427 | 8 | 4.492E+05 | 1.162E+06 | 7.946E+05 | 4.077E+05 | 6.155E+05 | 4.340E+05 | 9.413E+04 | 8.511E+06 | 4.521E+05 | 1.640E+05 | 9.827E+04 | 1.772E+06 | 2.786E+06 | 5.293E+06 | 1.870E+06 |
| 590 | 1013.379 | 3.335 | 8 | 2.843E+05 | 9.668E+05 | 7.091E+05 | 7.559E+04 | 2.500E+04 | 2.026E+04 | 2.498E+04 | 7.393E+06 | 1.582E+04 | 2.511E+04 | 2.800E+04 | 1.231E+06 | 3.832E+06 | 5.934E+06 | 5.302E+06 |
| 610 | 770.790 | 3.427 | 8 | 7.000E+04 | 3.862E+05 | 1.701E+05 | 5.155E+04 | 4.925E+04 | 5.242E+04 | 1.553E+04 | 2.486E+06 | 6.146E+04 | 5.617E+03 | 3.336E+04 | 5.059E+05 | 8.169E+05 | 1.736E+06 | 5.092E+05 |
| 614 | 837.291 | 3.461 | 8 | 3.735E+06 | 1.493E+07 | 1.362E+07 | 3.349E+06 | 7.724E+06 | 6.348E+06 | 1.437E+07 | 2.225E+06 | 1.761E+07 | 1.085E+07 | 5.806E+06 | 4.981E+06 | 1.770E+07 | 1.360E+07 | 3.854E+07 |
| 620 | 760.288 | 3.338 | 8 | 3.645E+04 | 1.920E+05 | 2.654E+04 | 2.486E+04 | 6.162E+03 | 0.000E+00 | 0.000E+00 | 1.858E+06 | 0.000E+00 | 0.000E+00 | 3.881E+03 | 3.380E+05 | 5.977E+05 | 1.289E+06 | 1.326E+06 |

| Feature identifiers | | | | XIC | | | | | | | | | | | | | | |
|---|---|---|---|---|---|---|---|---|---|---|---|---|---|---|---|---|---|---|
| ClusterIdx | m/z | RT [min] | Subnetwork | 4TUB34_I | 4TUB34_II | 4TUB34_III | 4TUB81_I | 4TUB81_II | 4TUB81_III | 4TUB93_I | 4TUB93_II | 4TUB93_III | 5TUB23_I | 5TUB23_II | 5TUB23_III | 5TUB48_I | 5TUB48_II | 5TUB48_III |
| 633 | 1062.719 | 3.822 | 8 | 4.105E+05 | 2.654E+06 | 2.303E+06 | 3.013E+05 | 5.400E+05 | 1.623E+05 | 4.266E+05 | 1.323E+06 | 1.035E+06 | 1.315E+06 | 3.881E+05 | 3.996E+06 | 7.555E+06 | 8.756E+06 | 7.576E+06 |
| 650 | 629.236 | 3.761 | 8 | 2.064E+06 | 2.393E+06 | 8.023E+06 | 1.728E+06 | 2.404E+06 | 1.806E+06 | 9.424E+06 | 5.981E+04 | 1.197E+07 | 7.043E+06 | 3.392E+06 | 1.725E+06 | 4.676E+06 | 2.872E+06 | 2.135E+07 |
| 659 | 813.049 | 4.312 | 8 | 9.655E+05 | 3.479E+06 | 3.283E+06 | 2.873E+05 | 1.395E+06 | 5.705E+05 | 6.138E+06 | 6.487E+04 | 5.164E+06 | 2.154E+07 | 7.501E+06 | 7.932E+06 | 3.740E+07 | 8.778E+06 | 9.608E+07 |
| 664 | 866.987 | 4.353 | 8 | 2.043E+04 | 6.303E+04 | 4.067E+05 | 0.000E+00 | 0.000E+00 | 0.000E+00 | 8.552E+05 | 0.000E+00 | 4.688E+05 | 1.147E+06 | 5.788E+05 | 0.000E+00 | 1.297E+05 | 0.000E+00 | 1.681E+06 |
| 672 | 1299.976 | 4.354 | 8 | 0.000E+00 | 0.000E+00 | 1.216E+04 | 4.742E+03 | 0.000E+00 | 0.000E+00 | 4.510E+04 | 0.000E+00 | 4.444E+04 | 4.328E+04 | 3.180E+04 | 0.000E+00 | 0.000E+00 | 0.000E+00 | 0.000E+00 |
| 676 | 874.315 | 4.356 | 8 | 0.000E+00 | 0.000E+00 | 0.000E+00 | 0.000E+00 | 0.000E+00 | 0.000E+00 | 3.015E+04 | 0.000E+00 | 0.000E+00 | 3.244E+04 | 0.000E+00 | 0.000E+00 | 0.000E+00 | 0.000E+00 | 1.182E+05 |
| 677 | 856.617 | 3.758 | 8 | 9.556E+05 | 2.286E+06 | 2.857E+06 | 8.548E+05 | 9.781E+05 | 6.724E+05 | 1.711E+06 | 1.908E+04 | 4.061E+06 | 2.714E+06 | 4.791E+05 | 6.914E+05 | 1.441E+06 | 1.016E+06 | 7.224E+06 |
| 678 | 879.636 | 4.362 | 8 | 0.000E+00 | 0.000E+00 | 0.000E+00 | 0.000E+00 | 0.000E+00 | 0.000E+00 | 7.638E+03 | 0.000E+00 | 0.000E+00 | 2.914E+04 | 0.000E+00 | 0.000E+00 | 0.000E+00 | 0.000E+00 | 0.000E+00 |
| 680 | 1297.948 | 4.058 | 8 | 4.480E+04 | 3.556E+05 | 5.964E+05 | 5.318E+04 | 7.561E+04 | 4.192E+04 | 1.306E+06 | 0.000E+00 | 1.178E+06 | 6.315E+05 | 3.437E+05 | 1.796E+04 | 2.648E+05 | 6.579E+04 | 3.822E+05 |
| 683 | 975.108 | 4.244 | 8 | 2.121E+04 | 2.981E+05 | 2.847E+05 | 5.455E+04 | 7.795E+04 | 3.733E+04 | 2.281E+05 | 6.188E+03 | 1.750E+05 | 7.124E+05 | 2.650E+05 | 5.006E+05 | 4.347E+06 | 2.418E+06 | 1.028E+07 |
| 685 | 1101.371 | 4.312 | 8 | 1.382E+04 | 3.517E+04 | 1.482E+05 | 4.879E+03 | 2.025E+04 | 6.882E+03 | 2.783E+05 | 0.000E+00 | 7.797E+04 | 1.009E+06 | 2.142E+05 | 2.294E+05 | 1.511E+06 | 7.881E+05 | 1.269E+06 |
| 691 | 822.287 | 4.312 | 8 | 9.517E+03 | 5.977E+04 | 9.205E+04 | 3.358E+04 | 1.511E+04 | 1.285E+04 | 1.985E+05 | 0.000E+00 | 7.795E+04 | 1.188E+06 | 3.480E+05 | 3.563E+05 | 2.245E+06 | 3.669E+05 | 2.854E+06 |
| 730 | 796.969 | 3.070 | 8 | 3.978E+06 | 3.218E+06 | 1.201E+06 | 3.625E+04 | 2.572E+04 | 1.820E+04 | 2.750E+05 | 7.842E+04 | 2.713E+05 | 4.941E+06 | 1.580E+05 | 5.821E+06 | 3.272E+06 | 1.845E+07 | 4.334E+07 |
| 741 | 1068.040 | 3.824 | 8 | 3.890E+05 | 1.075E+06 | 1.228E+06 | 4.207E+05 | 2.911E+05 | 1.338E+05 | 1.319E+05 | 2.527E+05 | 7.106E+05 | 6.295E+05 | 3.813E+05 | 2.257E+06 | 5.415E+06 | 5.633E+06 | 4.283E+06 |
| 746 | 1069.395 | 4.241 | 8 | 1.038E+06 | 1.786E+06 | 1.026E+06 | 1.709E+04 | 2.056E+04 | 0.000E+00 | 2.277E+05 | 0.000E+00 | 1.998E+05 | 1.476E+07 | 1.024E+06 | 1.864E+06 | 4.424E+06 | 8.129E+06 | 6.226E+07 |
| 751 | 1221.446 | 3.186 | 8 | 2.853E+05 | 9.876E+05 | 3.089E+05 | 1.980E+05 | 1.663E+05 | 5.721E+04 | 2.241E+05 | 7.550E+05 | 3.296E+05 | 2.078E+05 | 1.464E+05 | 8.965E+05 | 1.410E+06 | 2.391E+06 | 4.478E+05 |
| 757 | 1055.390 | 3.992 | 8 | 1.090E+06 | 1.271E+06 | 7.971E+05 | 2.591E+04 | 0.000E+00 | 0.000E+00 | 9.417E+04 | 2.053E+04 | 2.038E+05 | 2.014E+06 | 6.957E+04 | 1.319E+06 | 1.545E+06 | 5.173E+06 | 1.604E+07 |
| 760 | 975.359 | 4.241 | 8 | 3.836E+04 | 2.832E+05 | 3.462E+05 | 3.205E+04 | 4.772E+04 | 2.687E+04 | 8.017E+05 | 0.000E+00 | 1.467E+05 | 1.668E+06 | 2.185E+05 | 1.065E+06 | 4.836E+06 | 2.167E+06 | 1.050E+07 |
| 761 | 802.298 | 4.239 | 8 | 3.025E+05 | 2.677E+05 | 6.842E+05 | 0.000E+00 | 6.846E+03 | 0.000E+00 | 4.233E+04 | 0.000E+00 | 1.676E+05 | 8.097E+06 | 5.102E+05 | 9.438E+05 | 2.327E+06 | 4.446E+06 | 3.774E+07 |
| 766 | 618.733 | 3.459 | 8 | 4.502E+06 | 1.114E+07 | 1.112E+07 | 2.077E+06 | 3.690E+06 | 2.624E+06 | 9.177E+06 | 1.442E+06 | 1.190E+07 | 7.124E+06 | 3.932E+06 | 6.235E+06 | 1.188E+07 | 8.746E+06 | 3.156E+07 |
| 768 | 838.979 | 3.988 | 8 | 1.331E+06 | 2.044E+06 | 1.863E+06 | 0.000E+00 | 0.000E+00 | 1.071E+04 | 3.000E+05 | 0.000E+00 | 4.150E+05 | 1.034E+07 | 1.303E+06 | 5.635E+04 | 6.993E+05 | 7.158E+05 | 1.881E+07 |
| 778 | 796.636 | 3.069 | 8 | 2.845E+06 | 4.278E+06 | 1.697E+06 | 2.447E+04 | 1.409E+04 | 8.531E+03 | 1.610E+05 | 7.053E+04 | 2.677E+05 | 1.708E+06 | 2.749E+05 | 2.895E+06 | 2.655E+06 | 1.602E+07 | 2.794E+07 |
| 785 | 1215.954 | 3.392 | 8 | 8.009E+05 | 1.264E+06 | 2.899E+05 | 1.828E+04 | 0.000E+00 | 7.699E+03 | 1.019E+05 | 0.000E+00 | 1.914E+05 | 1.401E+06 | 6.404E+04 | 2.168E+05 | 1.977E+05 | 1.066E+06 | 7.050E+06 |
| 787 | 608.481 | 3.384 | 8 | 2.215E+05 | 7.765E+05 | 3.007E+05 | 0.000E+00 | 0.000E+00 | 8.030E+03 | 3.915E+04 | 0.000E+00 | 8.720E+04 | 8.808E+05 | 3.547E+04 | 1.768E+05 | 2.061E+05 | 9.190E+05 | 4.751E+06 |
| 788 | 1236.958 | 3.690 | 8 | 4.854E+05 | 5.837E+05 | 3.477E+05 | 0.000E+00 | 1.713E+04 | 0.000E+00 | 1.475E+05 | 0.000E+00 | 1.687E+05 | 2.252E+06 | 9.826E+04 | 1.029E+05 | 2.030E+05 | 2.334E+05 | 6.212E+06 |
| 789 | 618.734 | 3.694 | 8 | 1.676E+05 | 4.366E+05 | 4.703E+05 | 0.000E+00 | 0.000E+00 | 0.000E+00 | 6.976E+04 | 0.000E+00 | 4.138E+04 | 1.179E+06 | 5.985E+04 | 7.923E+04 | 1.460E+05 | 1.944E+05 | 5.014E+06 |
| 791 | 792.044 | 3.972 | 8 | 1.850E+05 | 4.216E+05 | 8.765E+04 | 2.424E+04 | 0.000E+00 | 8.969E+03 | 9.548E+04 | 6.182E+04 | 1.643E+04 | 3.676E+05 | 2.971E+04 | 4.164E+05 | 2.944E+05 | 1.913E+06 | 5.770E+06 |
| 796 | 1069.727 | 4.238 | 8 | 9.936E+05 | 1.256E+06 | 6.204E+05 | 0.000E+00 | 1.229E+04 | 1.016E+04 | 2.630E+05 | 0.000E+00 | 3.582E+05 | 1.242E+07 | 9.783E+05 | 1.659E+06 | 3.681E+06 | 7.289E+06 | 5.378E+07 |
| 800 | 824.643 | 3.674 | 8 | 4.336E+06 | 7.029E+06 | 4.580E+06 | 6.232E+04 | 3.206E+04 | 1.289E+04 | 1.084E+06 | 0.000E+00 | 1.954E+06 | 1.508E+07 | 1.153E+06 | 7.891E+05 | 2.000E+06 | 3.712E+06 | 3.989E+07 |
| 801 | 1055.390 | 4.161 | 8 | 4.278E+04 | 7.197E+04 | 1.955E+04 | 0.000E+00 | 0.000E+00 | 0.000E+00 | 2.950E+04 | 0.000E+00 | 2.647E+04 | 3.309E+06 | 1.035E+04 | 1.119E+05 | 7.609E+04 | 1.531E+05 | 1.048E+07 |
| 803 | 791.795 | 4.163 | 8 | 1.203E+05 | 1.017E+05 | 5.565E+04 | 0.000E+00 | 0.000E+00 | 0.000E+00 | 1.962E+04 | 0.000E+00 | 0.000E+00 | 2.431E+06 | 1.313E+04 | 7.909E+04 | 9.430E+04 | 4.753E+05 | 8.059E+06 |
| 804 | 1300.145 | 4.240 | 8 | 9.282E+03 | 4.278E+04 | 1.212E+05 | 2.320E+04 | 3.510E+04 | 5.475E+03 | 3.371E+04 | 0.000E+00 | 5.615E+04 | 1.423E+06 | 5.381E+04 | 1.059E+05 | 6.888E+05 | 1.757E+06 | 7.007E+06 |
| 805 | 810.973 | 3.576 | 8 | 1.790E+05 | 4.402E+05 | 2.683E+04 | 0.000E+00 | 1.911E+04 | 0.000E+00 | 0.000E+00 | 3.122E+04 | 0.000E+00 | 1.646E+06 | 2.204E+04 | 1.168E+04 | 3.193E+04 | 4.128E+04 | 3.219E+06 |
| 807 | 1082.043 | 4.246 | 8 | 1.445E+04 | 5.218E+04 | 1.575E+04 | 0.000E+00 | 0.000E+00 | 0.000E+00 | 0.000E+00 | 0.000E+00 | 0.000E+00 | 1.395E+06 | 7.098E+03 | 1.067E+05 | 1.162E+05 | 6.501E+05 | 4.633E+06 |
| 809 | 629.487 | 3.997 | 8 | 9.846E+04 | 2.873E+05 | 4.429E+04 | 0.000E+00 | 0.000E+00 | 0.000E+00 | 3.308E+04 | 0.000E+00 | 3.070E+04 | 8.745E+05 | 8.029E+04 | 0.000E+00 | 1.359E+04 | 2.671E+04 | 2.249E+06 |
| 812 | 1268.461 | 3.759 | 8 | 1.134E+06 | 2.847E+06 | 3.103E+06 | 1.029E+06 | 1.053E+06 | 9.473E+05 | 5.304E+06 | 5.618E+03 | 5.446E+06 | 1.803E+06 | 1.594E+06 | 8.243E+05 | 2.569E+06 | 5.545E+05 | 9.588E+06 |
| 820 | 1300.134 | 4.241 | 8 | 2.092E+04 | 4.176E+05 | 5.737E+05 | 3.079E+04 | 3.510E+04 | 0.000E+00 | 2.994E+05 | 0.000E+00 | 1.404E+05 | 3.036E+05 | 4.915E+05 | 4.898E+05 | 3.821E+06 | 4.833E+05 | 3.926E+06 |
| 829 | 1137.663 | 4.195 | 8 | 0.000E+00 | 0.000E+00 | 1.939E+04 | 0.000E+00 | 0.000E+00 | 0.000E+00 | 2.466E+04 | 0.000E+00 | 0.000E+00 | 1.835E+05 | 1.002E+04 | 8.080E+04 | 6.487E+05 | 1.543E+05 | 2.491E+06 |
| 830 | 1263.457 | 3.755 | 8 | 4.262E+04 | 1.141E+05 | 1.520E+05 | 0.000E+00 | 0.000E+00 | 3.089E+04 | 2.802E+05 | 0.000E+00 | 2.436E+05 | 1.722E+05 | 4.951E+04 | 0.000E+00 | 2.108E+05 | 0.000E+00 | 3.836E+05 |
| 833 | 1076.722 | 4.246 | 8 | 1.656E+04 | 6.849E+04 | 6.527E+04 | 0.000E+00 | 0.000E+00 | 0.000E+00 | 0.000E+00 | 0.000E+00 | 1.136E+04 | 1.635E+06 | 2.330E+04 | 7.506E+04 | 4.150E+05 | 7.928E+05 | 5.768E+06 |
| 836 | 1063.052 | 3.825 | 8 | 9.870E+05 | 2.286E+06 | 2.333E+06 | 6.600E+05 | 4.649E+05 | 2.567E+05 | 8.339E+05 | 3.586E+05 | 7.481E+05 | 3.591E+05 | 5.286E+05 | 3.643E+06 | 6.833E+06 | 7.079E+06 | 6.347E+06 |
| 838 | 791.796 | 3.996 | 8 | 1.622E+05 | 7.514E+05 | 3.004E+05 | 0.000E+00 | 0.000E+00 | 0.000E+00 | 4.241E+04 | 0.000E+00 | 2.713E+04 | 2.207E+05 | 6.272E+03 | 2.681E+05 | 6.882E+05 | 2.183E+06 | 2.547E+06 |
| 842 | 822.537 | 4.313 | 8 | 8.337E+03 | 1.113E+05 | 1.848E+05 | 1.133E+04 | 9.586E+03 | 9.818E+03 | 4.814E+04 | 0.000E+00 | 1.517E+05 | 4.554E+05 | 1.650E+05 | 5.199E+05 | 1.817E+06 | 6.557E+05 | 4.595E+06 |
| 845 | 1257.962 | 3.997 | 8 | 8.042E+04 | 7.302E+04 | 3.032E+04 | 0.000E+00 | 0.000E+00 | 0.000E+00 | 1.951E+04 | 0.000E+00 | 0.000E+00 | 5.610E+05 | 1.734E+04 | 0.000E+00 | 2.053E+04 | 2.578E+04 | 1.825E+06 |
| 847 | 781.292 | 3.601 | 8 | 5.735E+04 | 5.354E+05 | 6.045E+05 | 2.133E+05 | 3.225E+05 | 2.509E+05 | 8.784E+04 | 3.040E+06 | 5.228E+05 | 6.551E+04 | 4.293E+04 | 6.351E+05 | 2.186E+06 | 3.061E+06 | 1.903E+06 |
| 848 | 811.784 | 4.077 | 8 | 2.591E+04 | 1.739E+05 | 2.527E+05 | 4.803E+04 | 1.333E+04 | 9.660E+03 | 1.829E+05 | 0.000E+00 | 1.509E+05 | 1.534E+05 | 7.915E+04 | 7.572E+04 | 9.731E+05 | 1.229E+06 | 1.496E+06 |
| 851 | 1276.945 | 3.763 | 8 | 9.840E+05 | 2.414E+06 | 6.239E+05 | 1.133E+06 | 1.246E+06 | 5.752E+05 | 4.246E+06 | 8.877E+03 | 3.055E+06 | 3.339E+06 | 1.504E+06 | 5.378E+05 | 1.816E+05 | 1.151E+06 | 8.029E+06 |
| 854 | 1194.947 | 3.069 | 8 | 1.231E+05 | 1.675E+05 | 3.151E+04 | 0.000E+00 | 0.000E+00 | 0.000E+00 | 1.721E+04 | 0.000E+00 | 7.587E+03 | 2.865E+05 | 2.471E+04 | 1.776E+05 | 1.142E+05 | 1.882E+06 | 4.304E+06 |
| 862 | 964.605 | 4.154 | 8 | 1.621E+04 | 0.000E+00 | 0.000E+00 | 0.000E+00 | 0.000E+00 | 0.000E+00 | 1.422E+04 | 0.000E+00 | 0.000E+00 | 3.584E+05 | 0.000E+00 | 1.580E+04 | 6.155E+05 | 9.128E+04 | 2.716E+06 |
| 354 | 828.611 | 3.182 | 7 | 5.090E+06 | 9.188E+06 | 3.997E+06 | 3.106E+06 | 3.551E+06 | 2.487E+06 | 4.325E+06 | 6.283E+06 | 4.239E+06 | 3.568E+06 | 2.590E+06 | 8.435E+06 | 1.068E+07 | 1.232E+07 | 1.598E+07 |
| 360 | 842.615 | 3.460 | 7 | 2.713E+06 | 4.732E+06 | 4.789E+06 | 2.066E+06 | 2.724E+06 | 1.757E+06 | 5.214E+06 | 5.940E+05 | 4.888E+06 | 3.452E+06 | 3.864E+05 | 2.214E+06 | 5.219E+06 | 4.145E+06 | 9.690E+06 |
| 649 | 842.948 | 3.459 | 7 | 2.198E+06 | 4.931E+06 | 4.796E+06 | 1.793E+06 | 1.828E+06 | 1.298E+06 | 4.597E+06 | 6.424E+05 | 4.782E+06 | 2.932E+06 | 1.231E+06 | 2.549E+06 | 4.811E+06 | 3.804E+06 | 1.003E+07 |
| 811 | 828.945 | 3.185 | 7 | 4.016E+06 | 8.341E+06 | 7.958E+06 | 2.855E+06 | 1.488E+06 | 1.580E+06 | 2.963E+06 | 5.144E+06 | 4.136E+06 | 2.692E+06 | 5.311E+05 | 8.402E+06 | 7.950E+06 | 8.538E+06 | 1.495E+07 |
| 22 | 274.274 | 6.821 | 6 | 4.193E+07 | 4.208E+07 | 4.295E+07 | 4.278E+07 | 4.255E+07 | 4.380E+07 | 4.387E+07 | 4.442E+07 | 4.154E+07 | 4.397E+07 | 4.228E+07 | 4.241E+07 | 4.208E+07 | 4.146E+07 | 4.145E+07 |
| 104 | 302.305 | 7.664 | 6 | 8.660E+06 | 9.048E+06 | 9.669E+06 | 9.022E+06 | 9.358E+06 | 9.311E+06 | 9.330E+06 | 9.933E+06 | 9.630E+06 | 1.006E+07 | 9.859E+06 | 1.035E+07 | 9.573E+06 | 1.033E+07 | 1.010E+07 |
| 277 | 330.337 | 8.495 | 6 | 3.854E+06 | 3.960E+06 | 4.013E+06 | 3.918E+06 | 3.826E+06 | 4.332E+06 | 4.078E+06 | 4.209E+06 | 4.339E+06 | 4.290E+06 | 4.097E+06 | 4.347E+06 | 4.222E+06 | 4.206E+06 | 4.150E+06 |
| 341 | 358.368 | 9.237 | 6 | 3.383E+06 | 3.373E+06 | 3.181E+06 | 3.448E+06 | 3.440E+06 | 3.580E+06 | 3.718E+06 | 4.301E+06 | 3.588E+06 | 3.747E+06 | 3.211E+06 | 3.445E+06 | 3.405E+06 | 3.769E+06 | 3.670E+06 |
| 321 | 311.185 | 7.063 | 5 | 1.902E+06 | 2.056E+06 | 2.056E+06 | 2.082E+06 | 1.939E+06 | 1.998E+06 | 2.031E+06 | 2.014E+06 | 1.829E+06 | 1.954E+06 | 2.015E+06 | 2.074E+06 | 1.976E+06 | 2.009E+06 | 2.062E+06 |
| 347 | 423.310 | 10.200 | 5 | 1.887E+06 | 2.439E+06 | 2.802E+06 | 4.210E+05 | 8.999E+06 | 2.860E+06 | 0.000E+00 | 4.173E+06 | 5.576E+06 | 3.287E+07 | 2.275E+06 | 6.121E+06 | 4.104E+05 | 1.602E+05 | 3.901E+05 |
| 661 | 468.369 | 10.166 | 5 | 2.301E+05 | 2.375E+05 | 1.347E+05 | 2.890E+04 | 1.397E+06 | 2.595E+05 | 0.000E+00 | 6.343E+05 | 8.013E+05 | 4.866E+06 | 2.727E+05 | 1.026E+06 | 1.001E+05 | 0.000E+00 | 3.304E+04 |
| 11 | 452.277 | 7.678 | 4 | 4.717E+07 | 1.467E+08 | 1.878E+08 | 1.218E+08 | 1.439E+08 | 1.057E+08 | 1.551E+08 | 7.043E+07 | 3.607E+08 | 4.740E+08 | 2.151E+08 | 3.552E+08 | 2.304E+08 | 2.870E+08 | 2.522E+08 |
| 155 | 474.259 | 7.679 | 4 | 3.215E+06 | 8.124E+06 | 1.050E+07 | 5.417E+06 | 5.823E+06 | 4.557E+06 | 7.239E+06 | 4.797E+06 | 1.734E+07 | 1.990E+07 | 9.843E+06 | 1.498E+07 | 1.256E+07 | 1.451E+07 | 1.236E+07 |
| 238 | 474.258 | 7.808 | 4 | 5.696E+05 | 1.281E+06 | 1.645E+06 | 2.782E+06 | 3.718E+06 | 2.580E+06 | 3.283E+06 | 1.201E+06 | 4.306E+06 | 7.320E+06 | 3.591E+06 | 6.754E+06 | 1.305E+06 | 3.139E+06 | 3.747E+06 |
| 350 | 690.508 | 12.705 | 4 | 4.214E+06 | 5.253E+06 | 2.316E+05 | 3.960E+07 | 2.723E+07 | 1.721E+06 | 2.033E+07 | 6.168E+06 | 1.132E+06 | 9.450E+06 | 2.710E+07 | 5.046E+06 | 3.356E+07 | 4.776E+07 | 6.754E+06 |
| 424 | 511.304 | 8.562 | 4 | 1.198E+05 | 5.299E+04 | 1.619E+05 | 5.230E+05 | 1.781E+06 | 1.382E+06 | 2.364E+06 | 9.156E+05 | 2.386E+05 | 1.742E+07 | 1.285E+07 | 1.439E+07 | 7.126E+06 | 1.050E+07 | 7.197E+05 |
| 486 | 690.508 | 10.348 | 4 | 3.070E+07 | 2.183E+07 | 6.655E+04 | 4.746E+07 | 5.012E+07 | 9.804E+05 | 1.158E+08 | 6.009E+07 | 1.980E+05 | 1.075E+08 | 2.390E+08 | 1.886E+06 | 8.767E+07 | 1.858E+08 | 2.583E+07 |
| 489 | 712.490 | 12.701 | 4 | 1.459E+06 | 2.371E+06 | 4.886E+04 | 1.790E+07 | 1.386E+07 | 8.196E+05 | 9.666E+06 | 2.524E+06 | 2.643E+05 | 4.695E+06 | 1.152E+07 | 2.287E+06 | 1.335E+07 | 1.777E+07 | 2.660E+06 |
| 505 | 712.490 | 10.356 | 4 | 8.873E+05 | 2.963E+06 | 1.590E+04 | 8.826E+06 | 7.002E+07 | 2.219E+05 | 2.637E+07 | 4.271E+06 | 2.933E+04 | 1.357E+07 | 5.666E+07 | 4.372E+05 | 1.909E+07 | 5.101E+07 | 1.630E+07 |
| 510 | 485.288 | 8.510 | 4 | 4.680E+05 | 2.896E+05 | 6.318E+05 | 6.936E+05 | 1.669E+06 | 9.935E+05 | 3.700E+06 | 1.089E+06 | 5.796E+05 | 2.599E+07 | 9.950E+06 | 1.744E+07 | 1.354E+07 | 1.507E+07 | 4.475E+06 |
| 513 | 743.484 | 12.700 | 4 | 0.000E+00 | 0.000E+00 | 1.414E+05 | 5.249E+04 | 0.000E+00 | 0.000E+00 | 3.120E+06 | 5.858E+05 | 8.173E+04 | 3.955E+04 | 0.000E+00 | 2.299E+06 | 8.292E+05 | 1.773E+07 | 1.598E+07 |

| Feature identifiers | | | | XIC | | | | | | | | | | | | | | |
|---|---|---|---|---|---|---|---|---|---|---|---|---|---|---|---|---|---|---|
| ClusterIdx | m/z | RT [min] | Subnetwork | 4TUB34_I | 4TUB34_II | 4TUB34_III | 4TUB81_I | 4TUB81_II | 4TUB81_III | 4TUB93_I | 4TUB93_II | 4TUB93_III | 5TUB23_I | 5TUB23_II | 5TUB23_III | 5TUB48_I | 5TUB48_II | 5TUB48_III |
| 516 | 483.272 | 7.795 | 4 | 4.678E+05 | 9.180E+05 | 1.083E+06 | 8.685E+05 | 1.380E+06 | 4.285E+05 | 2.607E+06 | 1.060E+06 | 9.168E+05 | 1.931E+07 | 7.359E+06 | 1.223E+07 | 1.214E+07 | 1.157E+07 | 4.898E+06 |
| 517 | 740.522 | 12.702 | 4 | 0.000E+00 | 0.000E+00 | 2.023E+06 | 2.325E+06 | 3.544E+05 | 0.000E+00 | 2.248E+06 | 3.560E+05 | 1.345E+06 | 1.189E+06 | 5.252E+05 | 4.472E+06 | 2.618E+06 | 3.169E+07 | 2.358E+07 |
| 532 | 738.526 | 10.287 | 4 | 0.000E+00 | 0.000E+00 | 2.730E+03 | 1.089E+04 | 2.528E+04 | 0.000E+00 | 2.698E+07 | 3.932E+05 | 2.383E+04 | 1.415E+05 | 2.347E+04 | 3.023E+05 | 0.000E+00 | 8.096E+07 | 7.661E+06 |
| 554 | 903.547 | 7.681 | 4 | 1.272E+05 | 1.122E+06 | 1.936E+06 | 5.180E+05 | 5.869E+05 | 2.598E+05 | 7.219E+05 | 3.920E+05 | 4.363E+06 | 6.744E+06 | 1.552E+06 | 3.617E+06 | 2.386E+06 | 3.759E+06 | 2.427E+06 |
| 555 | 467.277 | 8.509 | 4 | 9.168E+04 | 5.754E+04 | 7.293E+04 | 1.899E+05 | 6.725E+05 | 3.885E+05 | 1.339E+06 | 3.010E+05 | 2.129E+05 | 1.089E+07 | 4.197E+06 | 7.269E+06 | 5.564E+06 | 6.093E+06 | 1.791E+06 |
| 562 | 743.484 | 10.306 | 4 | 0.000E+00 | 0.000E+00 | 0.000E+00 | 0.000E+00 | 0.000E+00 | 0.000E+00 | 1.503E+07 | 1.749E+05 | 1.717E+04 | 2.103E+04 | 0.000E+00 | 3.706E+05 | 3.698E+04 | 2.802E+07 | 4.938E+07 |
| 565 | 496.304 | 8.835 | 4 | 1.224E+06 | 9.147E+06 | 1.237E+07 | 0.000E+00 | 0.000E+00 | 0.000E+00 | 8.628E+05 | 3.938E+06 | 1.163E+05 | 7.150E+05 | 2.323E+05 | 6.939E+05 | 2.354E+05 | 1.723E+05 | 1.335E+06 |
| 577 | 483.272 | 7.944 | 4 | 1.437E+04 | 6.214E+04 | 4.133E+04 | 3.274E+05 | 6.157E+05 | 2.956E+05 | 6.461E+05 | 1.666E+04 | 9.721E+04 | 4.922E+06 | 2.120E+06 | 2.200E+06 | 7.943E+05 | 1.852E+06 | 1.746E+06 |
| 581 | 511.304 | 8.710 | 4 | 6.840E+03 | 0.000E+00 | 0.000E+00 | 1.435E+05 | 6.628E+05 | 3.518E+05 | 4.618E+05 | 2.456E+04 | 1.797E+04 | 6.591E+06 | 5.546E+06 | 5.573E+06 | 2.361E+06 | 3.507E+06 | 2.308E+05 |
| 583 | 452.277 | 7.806 | 4 | 8.804E+06 | 1.969E+07 | 2.511E+07 | 4.242E+07 | 5.638E+07 | 3.904E+07 | 4.880E+07 | 1.836E+07 | 6.462E+07 | 1.276E+08 | 5.707E+07 | 1.068E+08 | 2.358E+07 | 5.032E+07 | 6.134E+07 |
| 639 | 690.508 | 10.220 | 4 | 1.076E+06 | 1.227E+06 | 4.537E+05 | 4.441E+08 | 3.365E+08 | 3.632E+06 | 3.020E+07 | 2.473E+06 | 3.714E+05 | 6.925E+06 | 1.489E+07 | 7.387E+05 | 7.055E+07 | 6.321E+08 | 1.682E+08 |
| 643 | 712.490 | 10.268 | 4 | 1.474E+05 | 2.147E+05 | 3.094E+04 | 9.913E+07 | 1.146E+07 | 3.651E+05 | 7.544E+06 | 4.360E+05 | 7.818E+04 | 5.882E+05 | 1.096E+07 | 5.292E+05 | 1.987E+08 | 3.884E+08 | 5.381E+06 |
| 669 | 466.293 | 8.072 | 4 | 1.121E+04 | 4.109E+04 | 7.500E+04 | 2.263E+04 | 9.621E+04 | 1.068E+04 | 3.061E+05 | 2.471E+05 | 5.406E+05 | 5.242E+07 | 2.086E+07 | 1.379E+07 | 3.260E+06 | 2.352E+06 | 4.174E+06 |
| 679 | 466.293 | 8.206 | 4 | 5.734E+03 | 7.978E+03 | 1.250E+04 | 5.726E+03 | 3.714E+04 | 1.738E+04 | 3.251E+04 | 5.385E+04 | 3.497E+04 | 1.486E+07 | 1.061E+07 | 3.667E+06 | 3.533E+05 | 3.182E+05 | 9.433E+05 |
| 701 | 440.278 | 7.956 | 4 | 0.000E+00 | 0.000E+00 | 0.000E+00 | 1.533E+05 | 2.635E+05 | 6.661E+04 | 3.843E+05 | 4.717E+05 | 1.138E+06 | 6.786E+05 | 4.096E+04 | 9.993E+06 | 2.798E+06 | 2.260E+06 | 1.424E+06 |
| 713 | 662.477 | 12.666 | 4 | 0.000E+00 | 0.000E+00 | 6.902E+05 | 1.140E+04 | 1.867E+04 | 2.329E+04 | 0.000E+00 | 4.223E+03 | 2.759E+04 | 0.000E+00 | 8.771E+06 | 5.806E+06 | 1.671E+05 | 1.149E+07 | 1.722E+07 |
| 721 | 533.285 | 8.561 | 4 | 2.816E+04 | 0.000E+00 | 7.727E+03 | 1.849E+05 | 2.434E+05 | 3.191E+05 | 5.946E+05 | 2.366E+05 | 6.606E+04 | 5.287E+06 | 3.785E+06 | 4.219E+06 | 2.074E+06 | 3.063E+06 | 2.508E+05 |
| 722 | 440.278 | 7.818 | 4 | 0.000E+00 | 0.000E+00 | 0.000E+00 | 1.337E+04 | 0.000E+00 | 0.000E+00 | 6.938E+04 | 3.550E+05 | 4.008E+05 | 2.053E+05 | 0.000E+00 | 3.435E+06 | 1.299E+06 | 8.593E+05 | 6.695E+05 |
| 724 | 684.458 | 12.702 | 4 | 0.000E+00 | 0.000E+00 | 2.456E+05 | 0.000E+00 | 0.000E+00 | 0.000E+00 | 0.000E+00 | 0.000E+00 | 0.000E+00 | 0.000E+00 | 4.144E+06 | 2.452E+06 | 6.338E+04 | 3.829E+06 | 7.373E+06 |
| 727 | 465.262 | 7.796 | 4 | 7.131E+04 | 1.868E+05 | 1.328E+05 | 1.380E+05 | 3.604E+05 | 7.234E+04 | 7.000E+05 | 2.169E+05 | 1.460E+05 | 5.412E+06 | 1.881E+06 | 3.432E+06 | 3.363E+06 | 3.141E+06 | 1.386E+06 |
| 728 | 505.254 | 7.795 | 4 | 8.408E+04 | 2.459E+05 | 3.318E+05 | 1.015E+05 | 4.392E+05 | 1.477E+05 | 7.682E+05 | 2.605E+05 | 1.161E+05 | 5.364E+06 | 2.295E+06 | 3.454E+06 | 3.597E+06 | 3.413E+06 | 1.402E+06 |
| 729 | 507.269 | 8.510 | 4 | 4.626E+04 | 1.634E+04 | 6.376E+04 | 1.032E+05 | 4.245E+05 | 2.486E+05 | 6.264E+05 | 1.647E+05 | 2.131E+04 | 5.995E+06 | 2.312E+06 | 4.136E+06 | 3.229E+06 | 3.395E+06 | 9.449E+05 |
| 731 | 493.293 | 8.564 | 4 | 1.014E+04 | 0.000E+00 | 1.867E+04 | 7.456E+04 | 3.518E+05 | 2.327E+05 | 4.199E+05 | 1.257E+05 | 6.392E+04 | 3.868E+06 | 2.577E+06 | 3.006E+06 | 1.486E+06 | 2.275E+06 | 9.280E+04 |
| 736 | 676.492 | 12.707 | 4 | 0.000E+00 | 0.000E+00 | 0.000E+00 | 0.000E+00 | 0.000E+00 | 0.000E+00 | 0.000E+00 | 0.000E+00 | 4.983E+05 | 1.033E+05 | 0.000E+00 | 1.526E+06 | 7.927E+06 | 0.000E+00 | 2.902E+06 |
| 745 | 493.293 | 8.710 | 4 | 0.000E+00 | 0.000E+00 | 0.000E+00 | 2.759E+04 | 1.291E+05 | 5.007E+04 | 1.215E+05 | 9.598E+03 | 0.000E+00 | 2.366E+06 | 1.875E+06 | 1.924E+06 | 7.663E+05 | 1.171E+06 | 1.004E+05 |
| 749 | 500.300 | 7.793 | 4 | 8.164E+03 | 8.973E+04 | 6.260E+04 | 4.919E+04 | 1.025E+05 | 1.475E+04 | 2.660E+05 | 8.006E+04 | 7.286E+04 | 2.614E+06 | 8.677E+05 | 1.362E+06 | 1.358E+06 | 1.571E+06 | 4.060E+05 |
| 756 | 533.285 | 8.715 | 4 | 0.000E+00 | 0.000E+00 | 0.000E+00 | 0.000E+00 | 6.562E+04 | 4.075E+04 | 1.054E+05 | 4.614E+03 | 0.000E+00 | 1.599E+06 | 1.325E+06 | 1.206E+06 | 6.219E+05 | 6.959E+05 | 3.212E+04 |
| 775 | 704.523 | 12.703 | 4 | 0.000E+00 | 3.580E+06 | 3.343E+06 | 1.705E+05 | 0.000E+00 | 0.000E+00 | 0.000E+00 | 0.000E+00 | 0.000E+00 | 3.373E+04 | 1.976E+05 | 5.256E+04 | 2.747E+07 | 1.645E+07 | 2.297E+07 |
| 779 | 704.523 | 10.367 | 4 | 0.000E+00 | 2.737E+07 | 3.711E+07 | 5.012E+05 | 2.939E+04 | 9.374E+03 | 0.000E+00 | 1.566E+04 | 1.598E+04 | 0.000E+00 | 1.311E+05 | 1.646E+04 | 2.820E+05 | 9.473E+06 | 1.729E+08 |
| 781 | 726.506 | 12.701 | 4 | 0.000E+00 | 1.351E+06 | 1.267E+06 | 7.600E+04 | 0.000E+00 | 0.000E+00 | 0.000E+00 | 0.000E+00 | 0.000E+00 | 0.000E+00 | 2.653E+04 | 0.000E+00 | 1.170E+07 | 6.288E+06 | 9.112E+06 |
| 813 | 494.288 | 8.224 | 4 | 3.316E+05 | 3.603E+05 | 1.011E+06 | 2.233E+05 | 4.920E+05 | 3.824E+05 | 9.149E+03 | 5.729E+05 | 2.261E+06 | 3.198E+05 | 1.142E+04 | 1.600E+04 | 3.694E+04 | 1.404E+04 | 5.893E+04 |
| 825 | 664.490 | 9.773 | 4 | 4.372E+04 | 1.774E+04 | 1.251E+04 | 9.875E+04 | 5.920E+04 | 2.025E+04 | 0.000E+00 | 0.000E+00 | 0.000E+00 | 0.000E+00 | 0.000E+00 | 1.310E+05 | 9.168E+07 | 7.679E+06 | 4.218E+05 |
| 837 | 766.559 | 12.703 | 4 | 0.000E+00 | 0.000E+00 | 0.000E+00 | 0.000E+00 | 0.000E+00 | 0.000E+00 | 0.000E+00 | 3.103E+04 | 4.170E+04 | 3.020E+04 | 1.520E+05 | 9.050E+04 | 5.232E+04 | 1.188E+06 | 7.880E+05 |
| 841 | 676.491 | 10.424 | 4 | 0.000E+00 | 0.000E+00 | 0.000E+00 | 0.000E+00 | 0.000E+00 | 0.000E+00 | 0.000E+00 | 0.000E+00 | 2.815E+04 | 0.000E+00 | 0.000E+00 | 5.940E+04 | 1.021E+08 | 4.348E+05 | 2.530E+07 |
| 844 | 698.475 | 10.420 | 4 | 0.000E+00 | 0.000E+00 | 0.000E+00 | 0.000E+00 | 0.000E+00 | 0.000E+00 | 0.000E+00 | 0.000E+00 | 0.000E+00 | 0.000E+00 | 0.000E+00 | 1.839E+04 | 2.370E+07 | 0.000E+00 | 5.870E+06 |
| 853 | 1380.008 | 10.186 | 4 | 0.000E+00 | 0.000E+00 | 0.000E+00 | 2.387E+05 | 6.799E+04 | 0.000E+00 | 0.000E+00 | 0.000E+00 | 0.000E+00 | 0.000E+00 | 0.000E+00 | 0.000E+00 | 5.930E+05 | 1.635E+07 | 0.000E+00 |
| 855 | 1380.006 | 9.800 | 4 | 0.000E+00 | 0.000E+00 | 0.000E+00 | 0.000E+00 | 0.000E+00 | 0.000E+00 | 0.000E+00 | 0.000E+00 | 0.000E+00 | 0.000E+00 | 0.000E+00 | 0.000E+00 | 1.600E+05 | 3.291E+06 | 0.000E+00 |
| 27 | 316.212 | 3.277 | 3 | 2.542E+08 | 3.731E+08 | 3.281E+08 | 1.677E+08 | 1.709E+08 | 1.360E+08 | 2.113E+08 | 2.247E+08 | 1.977E+08 | 1.976E+08 | 2.820E+08 | 2.122E+08 | 2.380E+08 | 2.431E+08 | 2.937E+08 |
| 37 | 430.244 | 4.459 | 3 | 3.261E+07 | 4.052E+07 | 4.124E+07 | 2.699E+07 | 2.800E+07 | 2.340E+07 | 3.179E+07 | 3.335E+07 | 3.046E+07 | 2.610E+07 | 3.327E+07 | 2.690E+07 | 3.283E+07 | 3.422E+07 | 4.018E+07 |
| 55 | 300.200 | 3.938 | 3 | 7.754E+07 | 1.100E+08 | 9.161E+07 | 3.652E+07 | 3.669E+07 | 3.587E+07 | 5.477E+07 | 5.766E+07 | 5.796E+07 | 6.263E+07 | 7.786E+07 | 5.874E+07 | 6.897E+07 | 7.283E+07 | 7.430E+07 |
| 69 | 416.228 | 3.930 | 3 | 3.151E+07 | 3.652E+07 | 3.969E+07 | 2.012E+07 | 1.883E+07 | 1.940E+07 | 2.695E+07 | 3.040E+07 | 2.962E+07 | 2.885E+07 | 3.656E+07 | 3.197E+07 | 3.698E+07 | 4.115E+07 | 2.855E+07 |
| 122 | 243.183 | 3.050 | 3 | 1.982E+07 | 3.078E+07 | 2.485E+07 | 1.217E+07 | 9.732E+06 | 1.292E+07 | 2.299E+07 | 2.315E+07 | 2.432E+07 | 1.422E+07 | 1.972E+07 | 1.938E+07 | 2.514E+07 | 2.855E+07 | 2.642E+07 |
| 124 | 334.178 | 3.942 | 3 | 1.034E+07 | 2.258E+07 | 3.065E+07 | 1.988E+07 | 1.878E+07 | 1.304E+07 | 2.162E+07 | 1.840E+07 | 2.396E+07 | 8.225E+06 | 1.772E+07 | 1.885E+07 | 2.521E+07 | 2.077E+07 | 2.544E+07 |
| 128 | 357.214 | 4.538 | 3 | 1.401E+07 | 1.641E+07 | 1.352E+07 | 8.315E+06 | 1.017E+07 | 9.617E+06 | 1.142E+07 | 1.221E+07 | 1.178E+07 | 1.197E+07 | 1.522E+07 | 9.861E+06 | 1.150E+07 | 1.395E+07 | 1.516E+07 |
| 149 | 350.208 | 4.208 | 3 | 1.849E+07 | 2.596E+07 | 1.859E+07 | 1.325E+07 | 1.163E+07 | 1.240E+07 | 1.478E+07 | 1.738E+07 | 2.105E+07 | 1.866E+07 | 1.881E+07 | 1.630E+07 | 1.895E+07 | 2.271E+07 | 2.383E+07 |
| 167 | 599.391 | 3.938 | 3 | 2.815E+07 | 1.679E+07 | 3.648E+07 | 5.926E+06 | 5.366E+06 | 1.508E+07 | 6.069E+06 | 2.392E+07 | 9.922E+06 | 8.219E+06 | 1.396E+07 | 2.492E+07 | 3.015E+07 | 1.245E+07 | 1.392E+07 |
| 190 | 294.861 | 4.052 | 3 | 3.167E+07 | 4.971E+07 | 1.374E+07 | 1.872E+07 | 1.690E+07 | 1.981E+07 | 2.730E+07 | 2.906E+07 | 3.393E+07 | 3.100E+07 | 3.231E+07 | 3.047E+07 | 3.940E+07 | 3.987E+07 | 4.308E+07 |
| 224 | 302.196 | 0.594 | 3 | 4.455E+07 | 6.167E+07 | 4.507E+07 | 2.338E+07 | 1.748E+07 | 1.818E+07 | 3.235E+07 | 4.602E+07 | 4.731E+07 | 4.480E+07 | 5.575E+07 | 4.915E+07 | 5.983E+07 | 5.844E+07 | 6.751E+07 |
| 246 | 309.181 | 4.412 | 3 | 5.065E+06 | 8.089E+06 | 8.838E+06 | 6.227E+06 | 5.774E+06 | 5.127E+06 | 7.236E+06 | 7.637E+06 | 7.721E+06 | 4.879E+06 | 6.981E+06 | 6.021E+06 | 8.216E+06 | 7.295E+06 | 8.352E+06 |
| 268 | 328.200 | 4.325 | 3 | 8.030E+06 | 1.251E+07 | 7.621E+06 | 5.013E+06 | 4.816E+06 | 5.790E+06 | 7.034E+06 | 8.681E+06 | 9.764E+06 | 8.266E+06 | 7.691E+06 | 8.343E+06 | 9.435E+06 | 1.107E+07 | 1.239E+07 |
| 290 | 272.185 | 3.424 | 3 | 2.421E+07 | 4.232E+07 | 3.741E+07 | 1.293E+07 | 1.118E+07 | 6.586E+06 | 8.917E+06 | 1.754E+07 | 2.441E+07 | 8.935E+06 | 2.338E+07 | 1.334E+07 | 2.635E+07 | 1.719E+07 | 2.528E+07 |
| 293 | 293.191 | 3.577 | 3 | 9.866E+06 | 1.477E+07 | 9.755E+06 | 2.624E+06 | 2.202E+06 | 2.431E+06 | 3.868E+06 | 1.025E+07 | 6.855E+06 | 1.131E+07 | 1.239E+07 | 7.941E+06 | 9.026E+06 | 1.466E+07 | 1.428E+07 |
| 306 | 300.180 | 5.792 | 3 | 3.391E+06 | 4.007E+06 | 3.469E+06 | 2.214E+06 | 2.486E+06 | 1.738E+06 | 2.567E+06 | 2.191E+06 | 2.301E+06 | 2.034E+06 | 3.102E+06 | 2.013E+06 | 2.785E+06 | 2.602E+06 | 3.399E+06 |
| 322 | 713.423 | 4.538 | 3 | 2.843E+06 | 5.423E+06 | 4.561E+06 | 1.856E+06 | 3.399E+06 | 3.162E+06 | 3.836E+06 | 4.372E+06 | 3.809E+06 | 4.166E+06 | 5.199E+06 | 3.390E+06 | 3.871E+06 | 4.872E+06 | 5.093E+06 |
| 328 | 252.166 | 3.641 | 3 | 9.353E+06 | 1.679E+07 | 1.644E+07 | 4.518E+06 | 3.611E+06 | 2.892E+06 | 8.865E+06 | 8.746E+06 | 1.133E+07 | 8.317E+06 | 1.244E+07 | 1.066E+07 | 1.220E+07 | 1.101E+07 | 1.209E+07 |
| 329 | 332.872 | 4.600 | 3 | 9.211E+06 | 1.411E+07 | 9.334E+06 | 5.245E+06 | 4.734E+06 | 5.738E+06 | 7.202E+06 | 8.376E+06 | 9.876E+06 | 9.116E+06 | 8.816E+06 | 8.043E+06 | 9.759E+06 | 1.166E+07 | 1.408E+07 |
| 343 | 471.307 | 6.113 | 3 | 1.869E+06 | 1.697E+06 | 1.102E+06 | 1.088E+06 | 1.298E+06 | 1.254E+06 | 1.202E+06 | 1.380E+06 | 1.090E+06 | 1.752E+06 | 2.015E+06 | 1.088E+06 | 1.440E+06 | 1.557E+06 | 1.459E+06 |
| 361 | 441.788 | 4.079 | 3 | 1.152E+07 | 3.514E+06 | 8.855E+06 | 1.228E+06 | 1.215E+06 | 7.714E+06 | 1.250E+06 | 1.090E+07 | 2.057E+06 | 3.107E+06 | 4.199E+06 | 1.509E+06 | 2.992E+06 | 5.328E+06 | 5.546E+06 |
| 386 | 241.204 | 5.244 | 3 | 7.217E+06 | 7.315E+06 | 1.368E+07 | 1.372E+06 | 1.526E+06 | 1.412E+06 | 3.076E+06 | 1.527E+08 | 2.885E+06 | 2.026E+06 | 3.069E+06 | 3.473E+06 | 4.959E+06 | 4.206E+06 | 5.192E+06 |
| 396 | 469.328 | 4.897 | 3 | 2.548E+06 | 3.202E+06 | 2.688E+06 | 1.360E+06 | 1.286E+06 | 1.118E+06 | 1.938E+06 | 2.073E+06 | 1.825E+06 | 1.954E+06 | 2.418E+06 | 2.131E+06 | 2.114E+06 | 2.304E+06 | 2.427E+06 |
| 401 | 256.850 | 3.715 | 3 | 1.124E+07 | 2.126E+07 | 1.414E+07 | 7.276E+06 | 5.879E+06 | 7.401E+06 | 1.384E+07 | 1.705E+07 | 2.026E+07 | 1.152E+07 | 1.418E+07 | 1.610E+07 | 1.920E+07 | 2.354E+07 | 2.088E+07 |
| 404 | 699.407 | 4.204 | 3 | 4.449E+06 | 5.916E+06 | 4.508E+06 | 2.387E+06 | 2.327E+06 | 2.898E+06 | 3.811E+06 | 4.360E+06 | 4.315E+06 | 4.602E+06 | 4.561E+06 | 3.999E+06 | 4.660E+06 | 5.397E+06 | 6.893E+06 |
| 413 | 292.192 | 4.464 | 3 | 2.025E+07 | 3.363E+07 | 2.279E+07 | 1.058E+07 | 8.954E+05 | 1.052E+07 | 1.617E+07 | 1.924E+07 | 2.600E+07 | 1.685E+07 | 1.686E+07 | 1.742E+07 | 2.429E+07 | 2.360E+07 | 3.365E+07 |
| 439 | 585.375 | 3.578 | 3 | 2.990E+06 | 4.215E+06 | 2.909E+06 | 1.539E+06 | 9.541E+05 | 1.260E+06 | 2.031E+06 | 3.038E+06 | 3.484E+06 | 3.632E+06 | 4.364E+06 | 3.861E+06 | 4.599E+06 | 4.704E+06 | 4.177E+06 |
| 443 | 384.773 | 3.708 | 3 | 6.399E+06 | 1.181E+07 | 8.896E+06 | 4.985E+06 | 4.480E+06 | 5.382E+06 | 8.665E+06 | 1.199E+07 | 1.124E+07 | 6.941E+06 | 9.156E+06 | 1.127E+07 | 9.383E+06 | 5.553E+06 | 4.589E+06 |
| 463 | 293.191 | 3.406 | 3 | 3.229E+06 | 4.567E+06 | 2.826E+06 | 1.667E+06 | 1.399E+06 | 1.696E+06 | 2.182E+06 | 3.238E+06 | 4.104E+06 | 2.254E+06 | 2.042E+06 | 3.438E+06 | 3.866E+06 | 2.228E+06 | 4.745E+06 |
| 499 | 599.391 | 3.791 | 3 | 1.402E+07 | 4.604E+07 | 1.541E+07 | 1.537E+07 | 1.542E+07 | 7.317E+06 | 2.404E+07 | 9.861E+06 | 2.518E+07 | 2.764E+07 | 3.256E+07 | 1.159E+07 | 1.005E+07 | 3.235E+07 | 3.332E+07 |
| 526 | 485.359 | 3.054 | 3 | 3.713E+06 | 6.901E+06 | 5.515E+06 | 2.349E+06 | 1.041E+06 | 1.163E+06 | 4.661E+06 | 5.302E+06 | 5.556E+06 | 3.212E+06 | 4.562E+06 | 4.938E+06 | 6.282E+06 | 7.376E+06 | 7.412E+06 |
| 527 | 441.788 | 4.168 | 3 | 6.316E+06 | 1.608E+07 | 1.507E+07 | 6.963E+06 | 5.916E+06 | 1.100E+06 | 9.405E+06 | 3.286E+06 | 1.153E+07 | 1.095E+07 | 1.180E+07 | 1.134E+07 | 1.252E+07 | 1.307E+07 | 1.253E+07 |
| 530 | 290.193 | 3.898 | 3 | 6.109E+06 | 9.889E+06 | 6.591E+06 | 3.979E+06 | 3.715E+06 | 9.700E+05 | 6.456E+06 | 7.786E+06 | 8.953E+06 | 8.658E+06 | 8.838E+06 | 8.987E+06 | 9.615E+06 | 1.165E+07 | 1.170E+07 |

| Feature identifiers | | | | XIC | | | | | | | | | | | | | | |
|---|---|---|---|---|---|---|---|---|---|---|---|---|---|---|---|---|---|---|
| ClusterIdx | m/z | RT [min] | Subnetwork | 4TUB34_I | 4TUB34_II | 4TUB34_III | 4TUB81_I | 4TUB81_II | 4TUB81_III | 4TUB93_I | 4TUB93_II | 4TUB93_III | 5TUB23_I | 5TUB23_II | 5TUB23_III | 5TUB48_I | 5TUB48_II | 5TUB48_III |
| 553 | 332.207 | 4.839 | 3 | 6.268E+05 | 1.374E+06 | 4.629E+05 | 4.146E+05 | 1.099E+06 | 1.262E+06 | 1.615E+06 | 1.560E+06 | 1.664E+06 | 1.155E+06 | 7.906E+05 | 9.453E+05 | 1.221E+06 | 1.848E+06 | 1.929E+06 |
| 568 | 379.210 | 4.343 | 3 | 3.406E+06 | 5.082E+06 | 2.415E+06 | 4.870E+05 | 7.286E+05 | 6.765E+05 | 2.048E+06 | 3.290E+06 | 3.794E+06 | 3.681E+06 | 2.809E+06 | 2.302E+06 | 3.182E+06 | 4.094E+06 | 5.008E+06 |
| 623 | 343.200 | 3.890 | 3 | 1.803E+06 | 2.716E+06 | 1.524E+06 | 5.009E+05 | 5.095E+05 | 3.941E+05 | 9.889E+05 | 1.953E+06 | 2.431E+06 | 2.117E+06 | 2.487E+06 | 1.953E+06 | 2.304E+06 | 2.656E+06 | 2.792E+06 |
| 624 | 263.684 | 4.125 | 3 | 6.514E+05 | 8.677E+05 | 1.008E+07 | 4.103E+05 | 3.018E+05 | 2.148E+05 | 2.014E+05 | 1.042E+07 | 1.331E+07 | 1.254E+06 | 7.353E+05 | 8.368E+05 | 1.534E+07 | 1.379E+07 | 2.238E+06 |
| 627 | 230.153 | 3.714 | 3 | 2.078E+06 | 4.743E+06 | 5.174E+06 | 2.215E+05 | 5.087E+05 | 3.590E+05 | 1.271E+06 | 2.126E+06 | 2.025E+06 | 8.558E+05 | 2.063E+06 | 3.463E+05 | 3.310E+06 | 2.814E+06 | 3.151E+06 |
| 629 | 347.535 | 4.414 | 3 | 1.785E+06 | 2.874E+06 | 9.293E+05 | 5.295E+05 | 1.976E+05 | 1.597E+05 | 8.271E+05 | 2.002E+06 | 2.479E+06 | 2.007E+06 | 1.330E+06 | 1.692E+06 | 9.283E+05 | 2.783E+06 | 3.316E+06 |
| 747 | 540.426 | 9.375 | 3 | 8.955E+05 | 3.165E+06 | 2.393E+06 | 7.206E+05 | 7.457E+05 | 7.598E+05 | 5.979E+05 | 8.127E+05 | 1.990E+06 | 1.676E+06 | 2.181E+06 | 1.400E+06 | 1.551E+06 | 2.453E+06 | 2.512E+06 |
| 750 | 568.458 | 9.844 | 3 | 1.278E+05 | 1.023E+06 | 8.557E+05 | 5.202E+04 | 1.145E+05 | 1.210E+05 | 9.443E+04 | 1.790E+05 | 6.062E+05 | 1.272E+06 | 1.108E+06 | 1.001E+06 | 8.457E+05 | 2.813E+06 | 8.784E+05 |
| 758 | 263.684 | 4.034 | 3 | 8.854E+06 | 1.500E+07 | 1.399E+06 | 6.955E+05 | 3.370E+05 | 2.148E+05 | 1.685E+06 | 9.985E+05 | 3.241E+06 | 5.741E+05 | 9.253E+05 | 1.139E+07 | 1.293E+06 | 4.307E+05 | 1.688E+07 |
| 782 | 220.150 | 0.536 | 3 | 5.407E+04 | 1.716E+06 | 3.109E+05 | 4.341E+05 | 1.440E+05 | 0.000E+00 | 0.000E+00 | 2.291E+04 | 0.000E+00 | 1.179E+04 | 2.142E+04 | 8.124E+04 | 1.634E+05 | 1.114E+06 | 2.606E+06 |
| 784 | 498.804 | 4.596 | 3 | 1.085E+06 | 3.116E+06 | 2.451E+06 | 8.061E+05 | 5.490E+05 | 8.096E+05 | 1.501E+06 | 1.567E+06 | 2.848E+06 | 2.932E+05 | 9.142E+05 | 1.241E+06 | 2.049E+06 | 2.846E+06 | 2.974E+06 |
| 810 | 294.861 | 4.145 | 3 | 8.328E+06 | 2.400E+07 | 3.941E+07 | 4.531E+06 | 4.877E+06 | 3.070E+06 | 3.824E+06 | 8.364E+06 | 1.711E+07 | 5.142E+06 | 6.890E+06 | 9.898E+06 | 1.756E+07 | 1.225E+07 | 2.064E+07 |
| 817 | 384.772 | 3.711 | 3 | 4.264E+06 | 6.983E+06 | 2.602E+06 | 1.321E+06 | 2.028E+06 | 2.932E+06 | 3.224E+06 | 5.587E+06 | 6.498E+06 | 2.886E+06 | 4.752E+06 | 6.543E+06 | 3.613E+06 | 1.493E+07 | 1.000E+07 |
| 819 | 292.192 | 4.644 | 3 | 1.607E+06 | 8.131E+05 | 8.216E+05 | 8.459E+05 | 4.014E+05 | 2.488E+05 | 4.763E+05 | 6.615E+05 | 9.836E+05 | 9.986E+05 | 6.315E+05 | 6.597E+05 | 2.386E+06 | 2.641E+06 | 1.151E+06 |
| 93 | 256.264 | 9.789 | 2 | 7.530E+06 | 6.759E+06 | 7.139E+06 | 7.786E+06 | 7.977E+06 | 7.446E+06 | 8.080E+06 | 7.861E+06 | 7.095E+06 | 7.253E+06 | 7.876E+06 | 7.228E+06 | 8.762E+06 | 7.605E+06 | 7.129E+06 |
| 142 | 284.295 | 10.325 | 2 | 4.942E+06 | 4.614E+06 | 3.940E+06 | 5.532E+06 | 6.153E+06 | 5.044E+06 | 5.148E+06 | 4.588E+06 | 3.913E+06 | 4.353E+06 | 5.096E+06 | 4.762E+06 | 5.572E+06 | 4.858E+06 | 4.064E+06 |
| 175 | 308.279 | 8.430 | 2 | 4.065E+06 | 4.606E+06 | 3.125E+06 | 4.782E+06 | 4.157E+06 | 4.577E+06 | 4.259E+06 | 5.042E+06 | 5.267E+06 | 3.861E+06 | 4.376E+06 | 3.329E+06 | 4.325E+06 | 4.979E+06 | 5.086E+06 |
| 218 | 482.408 | 9.562 | 2 | 4.240E+06 | 3.689E+06 | 3.558E+06 | 3.135E+06 | 3.246E+06 | 2.992E+06 | 3.236E+06 | 4.340E+06 | 2.876E+06 | 3.684E+06 | 3.352E+06 | 9.019E+06 | 4.846E+06 | 3.931E+06 | 3.858E+06 |
| 226 | 526.434 | 9.538 | 2 | 4.923E+06 | 3.706E+06 | 4.368E+06 | 3.366E+06 | 3.115E+06 | 3.390E+06 | 2.556E+06 | 4.477E+06 | 2.954E+06 | 3.993E+06 | 3.429E+06 | 5.484E+06 | 7.607E+06 | 3.693E+06 | 4.696E+06 |
| 228 | 336.311 | 9.373 | 2 | 3.599E+06 | 3.431E+06 | 3.705E+06 | 4.270E+06 | 3.554E+06 | 3.404E+06 | 4.730E+06 | 1.470E+07 | 9.331E+06 | 4.795E+06 | 4.788E+06 | 3.765E+06 | 3.653E+06 | 4.044E+06 | 4.227E+06 |
| 229 | 291.253 | 8.431 | 2 | 2.928E+06 | 3.393E+06 | 2.213E+06 | 3.343E+06 | 3.110E+06 | 3.395E+06 | 3.296E+06 | 3.577E+06 | 3.940E+06 | 2.871E+06 | 3.239E+06 | 2.376E+06 | 3.168E+06 | 3.715E+06 | 3.941E+06 |
| 241 | 424.364 | 9.342 | 2 | 3.983E+06 | 3.049E+06 | 2.828E+06 | 5.422E+06 | 2.896E+06 | 2.899E+06 | 5.102E+06 | 2.118E+07 | 1.575E+07 | 6.528E+06 | 4.959E+06 | 3.024E+06 | 4.145E+06 | 3.549E+06 | 4.277E+06 |
| 242 | 380.338 | 9.364 | 2 | 4.162E+06 | 3.740E+06 | 3.596E+06 | 5.792E+06 | 3.620E+06 | 3.532E+06 | 5.910E+06 | 2.229E+07 | 1.567E+07 | 6.742E+06 | 5.862E+06 | 3.810E+06 | 4.133E+06 | 4.651E+06 | 4.957E+06 |
| 247 | 352.306 | 8.443 | 2 | 3.148E+06 | 3.213E+06 | 1.686E+06 | 2.649E+06 | 3.111E+06 | 2.946E+06 | 3.039E+06 | 3.570E+06 | 4.091E+06 | 2.909E+06 | 3.354E+06 | 2.161E+06 | 2.848E+06 | 3.797E+06 | 3.781E+06 |
| 269 | 438.382 | 9.584 | 2 | 5.623E+06 | 5.187E+06 | 5.378E+06 | 5.183E+06 | 5.883E+06 | 5.453E+06 | 5.692E+06 | 6.426E+06 | 6.047E+06 | 5.544E+06 | 5.783E+06 | 6.200E+06 | 9.564E+06 | 6.656E+06 | 6.399E+06 |
| 270 | 408.372 | 10.006 | 2 | 4.427E+06 | 3.865E+06 | 4.099E+06 | 3.769E+06 | 4.047E+06 | 3.623E+06 | 3.939E+06 | 5.151E+06 | 4.509E+06 | 3.973E+06 | 4.220E+06 | 4.261E+06 | 5.305E+06 | 4.239E+06 | 4.652E+06 |
| 276 | 570.460 | 9.514 | 2 | 5.683E+06 | 3.310E+06 | 3.945E+06 | 4.119E+06 | 2.305E+06 | 2.701E+06 | 2.088E+06 | 3.727E+06 | 2.334E+06 | 3.123E+06 | 2.735E+06 | 5.339E+06 | 7.104E+06 | 2.982E+06 | 3.904E+06 |
| 278 | 512.417 | 9.299 | 2 | 3.545E+06 | 2.159E+06 | 1.797E+06 | 4.690E+06 | 1.994E+06 | 2.342E+06 | 3.898E+06 | 1.493E+07 | 1.646E+07 | 5.569E+06 | 3.405E+06 | 2.635E+06 | 3.250E+06 | 2.152E+06 | 3.083E+06 |
| 280 | 468.390 | 9.320 | 2 | 3.744E+06 | 2.527E+06 | 2.159E+06 | 5.082E+06 | 2.413E+06 | 2.469E+06 | 4.652E+06 | 1.874E+07 | 1.688E+07 | 6.734E+06 | 4.111E+06 | 2.827E+06 | 3.667E+06 | 2.699E+06 | 3.689E+06 |
| 291 | 366.322 | 8.959 | 2 | 3.731E+06 | 3.581E+06 | 3.493E+06 | 3.431E+06 | 3.484E+06 | 3.344E+06 | 3.329E+06 | 3.383E+06 | 3.660E+06 | 3.434E+06 | 3.458E+06 | 3.292E+06 | 3.477E+06 | 3.444E+06 | 3.457E+06 |
| 294 | 452.398 | 9.985 | 2 | 3.516E+06 | 2.771E+06 | 2.784E+06 | 2.680E+06 | 2.446E+06 | 2.110E+06 | 2.358E+06 | 6.206E+06 | 3.388E+06 | 2.872E+06 | 4.156E+06 | 2.440E+06 | 3.355E+06 | 2.287E+06 | 4.810E+06 |
| 297 | 556.443 | 9.273 | 2 | 2.996E+06 | 1.906E+06 | 1.514E+06 | 4.642E+06 | 1.856E+06 | 1.998E+06 | 3.381E+06 | 1.101E+07 | 1.324E+07 | 4.449E+06 | 2.956E+06 | 2.159E+06 | 2.833E+06 | 1.933E+06 | 2.552E+06 |
| 302 | 247.227 | 8.431 | 2 | 1.306E+06 | 2.286E+06 | 1.766E+06 | 2.058E+06 | 1.808E+06 | 2.111E+06 | 1.811E+06 | 1.620E+06 | 2.367E+06 | 1.485E+06 | 2.026E+06 | 1.643E+06 | 1.975E+06 | 2.330E+06 | 2.304E+06 |
| 303 | 392.338 | 10.385 | 2 | 2.116E+06 | 2.141E+06 | 2.175E+06 | 2.147E+06 | 2.197E+06 | 2.162E+06 | 2.035E+06 | 1.881E+06 | 1.983E+06 | 2.009E+06 | 1.971E+06 | 1.991E+06 | 2.312E+06 | 1.991E+06 | 2.277E+06 |
| 310 | 496.424 | 9.963 | 2 | 5.847E+06 | 4.451E+06 | 4.245E+06 | 2.355E+06 | 3.701E+06 | 2.037E+06 | 2.273E+06 | 7.268E+06 | 3.550E+06 | 3.005E+06 | 4.049E+06 | 2.292E+06 | 5.684E+06 | 2.356E+06 | 5.112E+06 |
| 336 | 364.344 | 10.020 | 2 | 2.780E+06 | 2.608E+06 | 3.400E+06 | 2.515E+06 | 2.337E+06 | 2.324E+06 | 2.216E+06 | 2.808E+06 | 3.482E+06 | 2.569E+06 | 3.703E+06 | 2.624E+06 | 4.758E+06 | 3.594E+06 | 2.915E+06 |
| 337 | 540.449 | 9.937 | 2 | 6.044E+06 | 3.868E+06 | 1.997E+06 | 2.056E+06 | 1.780E+06 | 1.787E+06 | 1.933E+06 | 6.788E+06 | 3.372E+06 | 2.639E+06 | 1.815E+06 | 1.803E+06 | 5.334E+06 | 1.864E+06 | 4.637E+06 |
| 344 | 476.301 | 4.018 | 2 | 3.191E+06 | 3.189E+06 | 3.077E+06 | 2.748E+06 | 2.587E+06 | 2.681E+06 | 2.770E+06 | 3.224E+06 | 3.046E+06 | 3.448E+06 | 2.888E+06 | 2.728E+06 | 2.654E+06 | 2.813E+06 | 2.945E+06 |
| 349 | 440.359 | 8.424 | 2 | 2.355E+06 | 2.265E+06 | 1.453E+06 | 1.541E+06 | 1.672E+06 | 1.623E+06 | 1.785E+06 | 2.207E+06 | 2.317E+06 | 2.044E+06 | 2.287E+06 | 1.231E+06 | 1.827E+06 | 2.134E+06 | 2.253E+06 |
| 359 | 600.469 | 9.251 | 2 | 2.131E+06 | 1.523E+06 | 1.119E+06 | 3.979E+06 | 1.625E+06 | 1.550E+06 | 2.685E+06 | 7.366E+06 | 1.009E+07 | 3.279E+06 | 2.254E+06 | 1.689E+06 | 2.119E+06 | 1.438E+06 | 1.984E+06 |
| 362 | 322.295 | 8.962 | 2 | 2.609E+06 | 2.631E+06 | 2.705E+06 | 2.651E+06 | 2.595E+06 | 2.699E+06 | 2.583E+06 | 2.702E+06 | 2.733E+06 | 2.514E+06 | 2.534E+06 | 2.401E+06 | 2.483E+06 | 2.610E+06 | 2.441E+06 |
| 364 | 614.485 | 9.493 | 2 | 5.423E+06 | 2.344E+06 | 3.174E+06 | 1.718E+06 | 1.445E+06 | 1.885E+06 | 1.365E+06 | 2.797E+06 | 1.510E+06 | 2.274E+06 | 2.016E+06 | 4.659E+06 | 4.578E+06 | 2.377E+06 | 2.692E+06 |
| 366 | 396.333 | 8.432 | 2 | 2.280E+06 | 2.320E+06 | 1.491E+06 | 1.709E+06 | 1.753E+06 | 1.793E+06 | 1.963E+06 | 2.338E+06 | 2.738E+06 | 1.950E+06 | 2.242E+06 | 1.277E+06 | 1.890E+06 | 2.403E+06 | 2.382E+06 |
| 367 | 422.389 | 10.267 | 2 | 2.339E+06 | 1.912E+06 | 2.170E+06 | 1.889E+06 | 1.926E+06 | 1.890E+06 | 1.647E+06 | 1.852E+06 | 1.964E+06 | 2.090E+06 | 2.257E+06 | 1.751E+06 | 2.285E+06 | 2.118E+06 | 2.345E+06 |
| 369 | 436.364 | 10.348 | 2 | 2.261E+06 | 2.252E+06 | 2.228E+06 | 2.235E+06 | 2.438E+06 | 2.235E+06 | 2.164E+06 | 2.208E+06 | 2.316E+06 | 2.177E+06 | 2.300E+06 | 2.154E+06 | 2.316E+06 | 2.308E+06 | 2.340E+06 |
| 377 | 422.347 | 10.141 | 2 | 4.110E+06 | 4.102E+06 | 4.044E+06 | 4.006E+06 | 4.282E+06 | 4.103E+06 | 3.841E+06 | 4.151E+06 | 4.161E+06 | 4.296E+06 | 4.008E+06 | 4.071E+06 | 4.178E+06 | 3.482E+06 | 4.229E+06 |
| 382 | 466.373 | 10.105 | 2 | 3.643E+06 | 3.765E+06 | 1.875E+06 | 2.016E+06 | 3.914E+06 | 3.733E+06 | 2.044E+06 | 3.892E+06 | 4.053E+06 | 4.346E+06 | 3.912E+06 | 1.959E+06 | 3.361E+06 | 2.018E+06 | 3.732E+06 |
| 383 | 319.284 | 9.373 | 2 | 1.598E+06 | 1.708E+06 | 1.627E+06 | 1.775E+06 | 1.439E+06 | 1.310E+06 | 2.117E+06 | 7.441E+06 | 4.517E+06 | 2.117E+06 | 2.195E+06 | 1.636E+06 | 1.583E+06 | 1.826E+06 | 1.780E+06 |
| 387 | 466.415 | 10.248 | 2 | 2.031E+06 | 1.643E+06 | 1.743E+06 | 1.525E+06 | 1.604E+06 | 1.583E+06 | 1.451E+06 | 1.435E+06 | 1.784E+06 | 1.723E+06 | 1.735E+06 | 1.538E+06 | 1.939E+06 | 1.594E+06 | 2.238E+06 |
| 389 | 526.434 | 9.632 | 2 | 3.080E+06 | 2.222E+06 | 2.025E+06 | 1.756E+06 | 1.735E+06 | 1.754E+06 | 1.638E+06 | 2.053E+06 | 2.484E+06 | 2.432E+06 | 1.708E+06 | 2.332E+06 | 3.260E+06 | 1.894E+06 | 2.482E+06 |
| 390 | 468.390 | 9.206 | 2 | 1.824E+06 | 1.432E+06 | 1.431E+06 | 1.143E+06 | 1.188E+06 | 1.241E+06 | 1.074E+06 | 1.671E+06 | 1.038E+06 | 1.419E+06 | 1.396E+06 | 2.003E+06 | 2.034E+06 | 1.472E+06 | 1.650E+06 |
| 399 | 520.331 | 4.142 | 2 | 2.254E+06 | 2.208E+06 | 2.247E+06 | 1.812E+06 | 1.928E+06 | 1.984E+06 | 1.962E+06 | 2.105E+06 | 2.250E+06 | 2.474E+06 | 1.904E+06 | 1.880E+06 | 2.087E+06 | 2.005E+06 | 2.454E+06 |
| 400 | 484.385 | 8.410 | 2 | 2.190E+06 | 2.097E+06 | 1.177E+06 | 1.435E+06 | 1.566E+06 | 1.328E+06 | 1.503E+06 | 1.863E+06 | 1.913E+06 | 1.985E+06 | 2.144E+06 | 1.123E+06 | 1.686E+06 | 1.794E+06 | 1.754E+06 |
| 412 | 512.416 | 9.184 | 2 | 2.449E+06 | 1.471E+06 | 1.623E+06 | 1.102E+06 | 1.078E+06 | 1.263E+06 | 9.344E+05 | 1.878E+06 | 9.351E+05 | 1.471E+06 | 1.358E+06 | 2.441E+06 | 2.287E+06 | 1.487E+06 | 1.653E+06 |
| 414 | 528.411 | 8.398 | 2 | 2.023E+06 | 1.801E+06 | 9.636E+05 | 1.254E+06 | 1.440E+06 | 1.315E+06 | 1.520E+06 | 1.460E+06 | 1.521E+06 | 1.961E+06 | 1.681E+06 | 1.010E+06 | 1.505E+06 | 1.600E+06 | 1.670E+06 |
| 416 | 564.360 | 4.248 | 2 | 1.520E+06 | 1.645E+06 | 1.299E+06 | 1.398E+06 | 1.187E+06 | 1.306E+06 | 1.440E+06 | 1.470E+06 | 1.411E+06 | 1.576E+06 | 1.364E+06 | 1.339E+06 | 1.342E+06 | 1.326E+06 | 1.313E+06 |
| 417 | 644.495 | 9.227 | 2 | 1.445E+06 | 1.016E+06 | 6.837E+05 | 3.106E+06 | 1.328E+06 | 1.121E+06 | 1.777E+06 | 4.299E+06 | 6.650E+06 | 2.186E+06 | 1.387E+06 | 8.102E+05 | 1.333E+06 | 9.600E+05 | 1.237E+06 |
| 422 | 480.390 | 10.316 | 2 | 1.714E+06 | 1.624E+06 | 1.570E+06 | 1.842E+06 | 1.911E+06 | 1.696E+06 | 1.664E+06 | 1.664E+06 | 1.679E+06 | 1.828E+06 | 1.677E+06 | 1.797E+06 | 1.753E+06 | 1.878E+06 | 1.681E+06 |
| 432 | 556.443 | 9.161 | 2 | 2.797E+06 | 1.272E+06 | 1.556E+06 | 8.911E+05 | 8.330E+05 | 1.093E+06 | 7.328E+05 | 1.652E+06 | 7.133E+05 | 1.246E+06 | 1.109E+06 | 2.598E+06 | 2.586E+06 | 1.260E+06 | 1.469E+06 |
| 438 | 584.475 | 9.912 | 2 | 3.245E+06 | 1.599E+06 | 1.255E+06 | 1.666E+06 | 1.094E+06 | 1.187E+06 | 1.282E+06 | 4.042E+06 | 2.985E+06 | 1.890E+06 | 1.261E+06 | 1.138E+06 | 2.161E+06 | 9.260E+05 | 3.426E+06 |
| 455 | 628.501 | 9.881 | 2 | 2.769E+06 | 1.268E+06 | 9.937E+05 | 1.545E+06 | 8.457E+05 | 8.783E+05 | 8.994E+05 | 2.943E+06 | 2.592E+06 | 1.450E+06 | 9.486E+05 | 7.971E+05 | 1.651E+06 | 7.611E+05 | 1.414E+06 |
| 464 | 608.392 | 4.346 | 2 | 1.009E+06 | 1.166E+06 | 1.054E+06 | 8.020E+05 | 1.013E+06 | 9.739E+05 | 9.363E+05 | 1.051E+06 | 9.642E+05 | 1.012E+06 | 1.054E+06 | 8.440E+05 | 7.494E+05 | 9.680E+05 | 9.431E+05 |
| 471 | 352.306 | 8.095 | 2 | 3.758E+05 | 8.836E+04 | 5.647E+05 | 5.195E+06 | 1.398E+06 | 1.097E+06 | 5.421E+06 | 4.510E+07 | 8.256E+07 | 1.029E+07 | 9.914E+06 | 3.104E+06 | 7.361E+06 | 5.030E+07 | 2.768E+07 |
| 528 | 308.279 | 8.075 | 2 | 1.537E+05 | 9.131E+04 | 1.282E+05 | 4.188E+06 | 7.906E+05 | 4.466E+05 | 5.959E+06 | 3.277E+07 | 8.692E+07 | 7.851E+06 | 7.353E+06 | 1.975E+05 | 3.794E+06 | 2.531E+07 | 1.889E+07 |
| 539 | 688.521 | 9.203 | 2 | 9.751E+05 | 7.775E+05 | 4.996E+05 | 2.412E+06 | 9.323E+05 | 8.594E+05 | 1.289E+06 | 2.540E+06 | 4.351E+06 | 1.323E+06 | 9.491E+05 | 0.000E+00 | 0.000E+00 | 6.010E+05 | 0.000E+00 |
| 541 | 363.310 | 9.363 | 2 | 1.108E+06 | 1.006E+06 | 8.562E+05 | 1.398E+06 | 1.014E+06 | 9.403E+05 | 1.462E+06 | 6.341E+06 | 4.493E+06 | 1.669E+06 | 1.488E+06 | 9.538E+05 | 1.071E+06 | 9.994E+05 | 1.122E+06 |
| 546 | 291.253 | 8.075 | 2 | 9.279E+04 | 6.500E+04 | 1.282E+05 | 2.309E+06 | 2.551E+05 | 1.354E+05 | 4.224E+06 | 2.338E+07 | 5.807E+07 | 5.450E+06 | 5.270E+06 | 5.122E+05 | 2.555E+06 | 1.813E+07 | 1.353E+07 |
| 548 | 396.333 | 8.097 | 2 | 1.207E+06 | 3.312E+05 | 3.646E+05 | 4.255E+06 | 2.821E+05 | 6.753E+05 | 4.095E+06 | 3.229E+07 | 5.379E+07 | 8.970E+06 | 9.644E+06 | 3.762E+06 | 8.522E+06 | 5.244E+07 | 2.668E+07 |
| 556 | 732.548 | 9.180 | 2 | 6.838E+05 | 8.231E+05 | 3.960E+05 | 1.841E+06 | 7.809E+05 | 5.741E+05 | 1.011E+06 | 1.459E+06 | 2.756E+06 | 8.213E+05 | 6.605E+05 | 3.597E+05 | 6.864E+05 | 3.037E+05 | 4.812E+05 |
| 564 | 440.359 | 8.096 | 2 | 1.971E+06 | 1.046E+06 | 3.136E+05 | 3.686E+06 | 5.035E+05 | 2.210E+05 | 3.383E+06 | 2.669E+07 | 3.324E+07 | 7.962E+06 | 9.974E+06 | 4.600E+06 | 9.559E+06 | 4.300E+07 | 2.541E+07 |

| Feature identifiers | | | | XIC | | | | | | | | | | | | | | |
|---|---|---|---|---|---|---|---|---|---|---|---|---|---|---|---|---|---|---|
| ClusterIdx | m/z | RT [min] | Subnetwork | 4TUB34_I | 4TUB34_II | 4TUB34_III | 4TUB81_I | 4TUB81_II | 4TUB81_III | 4TUB93_I | 4TUB93_II | 4TUB93_III | 5TUB23_I | 5TUB23_II | 5TUB23_III | 5TUB48_I | 5TUB48_II | 5TUB48_III |
| 570 | 484.385 | 8.094 | 2 | 1.748E+06 | 3.854E+05 | 2.456E+05 | 2.589E+06 | 6.397E+05 | 2.148E+05 | 2.716E+06 | 1.973E+07 | 2.309E+07 | 6.067E+06 | 8.633E+06 | 4.289E+06 | 8.464E+06 | 2.866E+07 | 2.010E+07 |
| 571 | 308.279 | 8.290 | 2 | 6.323E+03 | 2.310E+05 | 2.692E+05 | 5.653E+05 | 2.402E+04 | 1.534E+04 | 9.282E+05 | 5.601E+06 | 8.922E+06 | 8.417E+05 | 7.743E+05 | 7.385E+04 | 6.761E+04 | 3.001E+06 | 1.746E+06 |
| 573 | 528.411 | 8.088 | 2 | 1.156E+06 | 7.778E+04 | 1.429E+05 | 2.056E+06 | 1.684E+05 | 2.026E+05 | 1.773E+06 | 1.497E+07 | 1.479E+07 | 4.212E+06 | 7.188E+06 | 3.886E+06 | 6.488E+06 | 1.704E+07 | 1.445E+07 |
| 576 | 247.227 | 8.064 | 2 | 0.000E+00 | 5.888E+03 | 2.713E+04 | 2.182E+05 | 5.329E+04 | 1.454E+04 | 1.583E+06 | 6.126E+06 | 2.426E+07 | 5.655E+05 | 6.316E+05 | 1.177E+04 | 5.543E+04 | 2.426E+06 | 2.335E+06 |
| 579 | 335.280 | 8.098 | 2 | 9.389E+04 | 3.848E+04 | 5.274E+04 | 1.644E+06 | 1.602E+05 | 1.498E+05 | 1.497E+06 | 1.524E+07 | 2.572E+07 | 3.807E+06 | 3.722E+06 | 4.393E+05 | 1.210E+06 | 1.781E+07 | 1.054E+07 |
| 580 | 572.437 | 8.084 | 2 | 3.969E+05 | 1.923E+05 | 7.358E+04 | 6.285E+05 | 1.641E+05 | 4.464E+04 | 1.307E+06 | 1.028E+07 | 8.165E+06 | 2.921E+06 | 5.608E+06 | 3.132E+06 | 4.635E+06 | 9.195E+06 | 1.019E+07 |
| 598 | 396.333 | 8.315 | 2 | 3.654E+04 | 1.043E+05 | 1.472E+05 | 5.813E+05 | 1.140E+05 | 8.081E+04 | 3.800E+05 | 4.083E+06 | 2.633E+06 | 8.193E+05 | 8.717E+05 | 1.343E+05 | 1.456E+05 | 3.793E+06 | 2.575E+06 |
| 604 | 440.359 | 8.302 | 2 | 6.589E+04 | 1.348E+05 | 1.929E+05 | 2.415E+05 | 5.751E+04 | 6.653E+04 | 1.605E+05 | 3.192E+06 | 4.631E+06 | 6.665E+05 | 8.169E+05 | 2.389E+05 | 5.233E+05 | 3.528E+06 | 2.303E+06 |
| 608 | 672.527 | 9.854 | 2 | 2.229E+06 | 8.442E+05 | 9.024E+05 | 1.326E+06 | 7.677E+05 | 7.671E+05 | 8.657E+05 | 1.881E+06 | 1.962E+06 | 1.077E+06 | 6.377E+05 | 5.494E+05 | 1.229E+06 | 5.074E+05 | 9.309E+05 |
| 615 | 350.328 | 9.687 | 2 | 2.851E+06 | 9.760E+06 | 2.744E+06 | 2.950E+06 | 2.422E+06 | 2.671E+06 | 2.855E+06 | 9.425E+06 | 1.014E+06 | 2.815E+06 | 2.839E+06 | 1.739E+06 | 1.995E+06 | 2.051E+06 | 1.007E+07 |
| 618 | 658.511 | 9.464 | 2 | 4.409E+06 | 1.504E+06 | 2.175E+06 | 9.408E+05 | 4.987E+05 | 8.303E+05 | 5.495E+05 | 1.829E+06 | 1.086E+06 | 1.406E+06 | 1.181E+06 | 3.197E+06 | 3.508E+06 | 1.368E+06 | 1.807E+06 |
| 625 | 702.536 | 9.449 | 2 | 2.971E+06 | 9.969E+05 | 1.284E+06 | 4.249E+05 | 6.464E+05 | 6.749E+05 | 4.377E+05 | 9.787E+05 | 8.498E+05 | 4.651E+05 | 7.907E+05 | 1.919E+06 | 2.504E+06 | 9.031E+05 | 1.015E+06 |
| 647 | 394.355 | 9.697 | 2 | 1.014E+07 | 2.866E+06 | 1.960E+06 | 2.639E+06 | 2.526E+06 | 1.873E+06 | 2.177E+06 | 2.300E+06 | 2.834E+06 | 1.010E+07 | 2.189E+06 | 2.240E+06 | 2.053E+06 | 2.722E+06 | 1.127E+07 |
| 50 | 701.620 | 10.613 | 1 | 9.707E+06 | 9.443E+06 | 9.748E+06 | 8.639E+06 | 9.112E+06 | 8.817E+06 | 6.804E+06 | 7.616E+06 | 6.919E+06 | 8.000E+06 | 6.956E+06 | 6.168E+06 | 6.303E+06 | 6.947E+06 | 6.422E+06 |
| 199 | 419.364 | 7.499 | 1 | 3.007E+06 | 3.118E+06 | 3.016E+06 | 3.519E+06 | 3.053E+06 | 3.015E+06 | 3.142E+06 | 2.864E+06 | 3.008E+06 | 3.293E+06 | 2.974E+06 | 2.876E+06 | 3.000E+06 | 3.050E+06 | 2.507E+06 |
| 689 | 685.624 | 12.053 | 1 | 6.995E+05 | 4.212E+05 | 8.367E+05 | 4.800E+05 | 5.730E+05 | 5.966E+05 | 1.825E+05 | 4.616E+05 | 4.082E+05 | 4.869E+05 | 2.577E+05 | 1.609E+05 | 4.404E+05 | 3.081E+05 | 2.666E+05 |
| 1 | 110.020 | 12.761 | -1 | 1.003E+09 | 1.601E+09 | 1.535E+09 | 1.510E+09 | 1.459E+09 | 1.443E+09 | 1.474E+09 | 1.425E+09 | 1.427E+09 | 1.403E+09 | 1.353E+09 | 1.326E+09 | 1.344E+09 | 1.236E+09 | 1.319E+09 |
| 8 | 437.194 | 7.524 | -1 | 2.588E+08 | 2.893E+08 | 2.536E+08 | 2.528E+08 | 2.845E+08 | 2.534E+08 | 2.594E+08 | 2.857E+08 | 2.673E+08 | 2.959E+08 | 2.881E+08 | 2.404E+08 | 2.882E+08 | 2.631E+08 | 2.997E+08 |
| 10 | 120.081 | 0.519 | -1 | 2.114E+09 | 9.248E+08 | 1.795E+09 | 8.928E+07 | 1.602E+08 | 1.476E+08 | 4.441E+08 | 2.807E+08 | 6.480E+08 | 8.332E+08 | 9.661E+08 | 7.661E+08 | 2.332E+09 | 1.441E+08 | 2.233E+08 |
| 13 | 242.284 | 8.651 | -1 | 3.958E+07 | 2.296E+08 | 1.300E+08 | 2.395E+08 | 2.332E+08 | 2.345E+08 | 2.310E+08 | 2.490E+08 | 2.308E+08 | 2.272E+08 | 2.254E+08 | 2.284E+08 | 2.248E+08 | 2.376E+08 | 2.285E+08 |
| 14 | 851.398 | 7.527 | -1 | 6.010E+07 | 9.206E+07 | 6.078E+07 | 5.936E+07 | 8.436E+07 | 5.779E+07 | 6.626E+07 | 7.620E+07 | 6.352E+07 | 9.941E+07 | 9.368E+07 | 4.873E+07 | 8.699E+07 | 7.293E+07 | 1.014E+08 |
| 32 | 437.194 | 7.249 | -1 | 8.802E+06 | 2.644E+07 | 9.885E+06 | 1.629E+07 | 2.325E+07 | 2.029E+07 | 1.157E+07 | 4.412E+07 | 1.581E+07 | 2.530E+07 | 3.191E+07 | 9.815E+06 | 1.266E+07 | 1.677E+07 | 3.655E+07 |
| 33 | 559.518 | 11.442 | -1 | 1.249E+08 | 1.266E+08 | 1.237E+08 | 1.287E+08 | 1.232E+08 | 1.277E+08 | 1.192E+08 | 1.226E+08 | 1.226E+08 | 1.207E+08 | 1.196E+08 | 1.199E+08 | 1.184E+08 | 1.061E+08 | 1.185E+08 |
| 34 | 867.372 | 7.528 | -1 | 2.642E+07 | 3.215E+07 | 2.780E+07 | 2.763E+07 | 3.288E+07 | 2.626E+07 | 2.904E+07 | 3.078E+07 | 2.694E+07 | 3.787E+07 | 3.280E+07 | 2.447E+07 | 3.146E+07 | 3.112E+07 | 3.508E+07 |
| 41 | 587.549 | 11.407 | -1 | 1.228E+08 | 1.223E+08 | 1.222E+08 | 1.259E+08 | 1.182E+08 | 1.247E+08 | 1.200E+08 | 1.160E+08 | 1.187E+08 | 1.189E+08 | 1.167E+08 | 1.106E+08 | 1.151E+08 | 1.035E+08 | 1.143E+08 |
| 47 | 163.991 | 12.748 | -1 | 1.950E+07 | 1.578E+07 | 1.692E+07 | 1.681E+07 | 1.670E+07 | 1.628E+07 | 1.552E+07 | 1.536E+07 | 1.442E+07 | 1.562E+07 | 1.401E+07 | 1.415E+07 | 1.429E+07 | 1.440E+07 | 1.359E+07 |
| 53 | 243.088 | 4.621 | -1 | 7.856E+06 | 1.644E+07 | 1.639E+07 | 7.113E+06 | 1.060E+07 | 1.191E+07 | 1.607E+07 | 1.238E+07 | 2.376E+07 | 8.620E+06 | 9.478E+06 | 1.027E+07 | 1.369E+07 | 1.410E+07 | 1.491E+07 |
| 54 | 399.209 | 10.100 | -1 | 1.193E+07 | 1.271E+07 | 1.174E+07 | 1.085E+07 | 1.244E+07 | 1.285E+07 | 1.091E+07 | 1.235E+07 | 1.088E+07 | 1.247E+07 | 1.181E+07 | 1.123E+07 | 8.628E+06 | 6.965E+06 | 1.004E+07 |
| 57 | 685.436 | 12.425 | -1 | 1.983E+07 | 2.554E+07 | 2.569E+07 | 1.918E+07 | 1.839E+07 | 2.005E+07 | 1.904E+07 | 1.835E+07 | 2.282E+07 | 4.364E+07 | 4.848E+07 | 6.324E+07 | 5.062E+07 | 8.543E+07 | 3.729E+07 |
| 58 | 250.112 | 5.753 | -1 | 0.000E+00 | 1.524E+04 | 3.662E+03 | 1.259E+07 | 1.517E+07 | 1.035E+07 | 2.578E+05 | 9.319E+04 | 1.861E+07 | 1.570E+05 | 2.217E+05 | 5.698E+03 | 0.000E+00 | 0.000E+00 | 0.000E+00 |
| 64 | 255.073 | 4.542 | -1 | 1.117E+07 | 8.725E+06 | 1.117E+07 | 1.523E+07 | 2.565E+07 | 1.521E+07 | 1.536E+07 | 2.387E+07 | 2.200E+07 | 0.000E+00 | 0.000E+00 | 0.000E+00 | 0.000E+00 | 0.000E+00 | 0.000E+00 |
| 66 | 261.109 | 5.951 | -1 | 1.782E+07 | 5.617E+06 | 1.052E+07 | 1.876E+07 | 9.703E+06 | 2.021E+07 | 2.599E+07 | 1.600E+07 | 1.712E+07 | 3.843E+07 | 2.423E+07 | 2.931E+07 | 1.542E+07 | 1.714E+07 | 1.522E+07 |
| 71 | 267.077 | 3.607 | -1 | 3.129E+07 | 2.135E+07 | 2.080E+07 | 1.766E+07 | 2.017E+07 | 1.552E+07 | 1.642E+07 | 2.955E+07 | 4.121E+07 | 1.426E+07 | 1.990E+07 | 8.924E+06 | 2.530E+07 | 2.798E+07 | 1.878E+07 |
| 73 | 453.168 | 7.250 | -1 | 4.106E+06 | 1.191E+07 | 4.446E+06 | 7.689E+06 | 1.039E+07 | 9.048E+06 | 5.541E+06 | 1.904E+07 | 7.215E+06 | 1.157E+07 | 1.440E+07 | 4.542E+06 | 5.312E+06 | 7.526E+06 | 1.584E+07 |
| 76 | 324.142 | 7.584 | -1 | 9.975E+06 | 1.635E+07 | 7.998E+06 | 8.618E+06 | 1.318E+07 | 7.782E+06 | 8.744E+06 | 7.254E+06 | 1.183E+07 | 7.051E+06 | 1.306E+07 | 6.246E+06 | 1.308E+07 | 1.101E+07 | 1.405E+07 |
| 77 | 372.882 | 12.723 | -1 | 2.683E+06 | 1.120E+06 | 1.463E+06 | 1.486E+06 | 1.528E+06 | 1.425E+06 | 1.780E+06 | 1.294E+06 | 1.598E+06 | 1.485E+06 | 1.098E+06 | 1.108E+06 | 9.736E+05 | 1.140E+06 | 1.966E+06 |
| 79 | 321.131 | 4.460 | -1 | 1.587E+07 | 1.012E+07 | 9.476E+06 | 8.884E+06 | 1.258E+07 | 1.088E+07 | 8.571E+06 | 2.016E+07 | 9.450E+06 | 2.008E+07 | 2.058E+07 | 1.300E+07 | 2.447E+07 | 1.734E+07 | 2.636E+07 |
| 82 | 269.015 | 5.941 | -1 | 3.057E+06 | 1.292E+07 | 7.057E+06 | 1.132E+07 | 1.337E+07 | 1.000E+07 | 7.799E+05 | 0.000E+00 | 5.628E+06 | 0.000E+00 | 0.000E+00 | 3.232E+04 | 0.000E+00 | 0.000E+00 | 0.000E+00 |
| 83 | 230.141 | 12.721 | -1 | 6.534E+06 | 6.611E+06 | 5.516E+06 | 7.567E+06 | 6.292E+06 | 6.878E+06 | 7.214E+06 | 6.851E+06 | 7.109E+06 | 6.426E+06 | 7.632E+06 | 6.911E+06 | 5.998E+06 | 7.000E+06 | 6.020E+06 |
| 85 | 233.079 | 4.825 | -1 | 8.601E+06 | 5.770E+06 | 5.059E+06 | 6.208E+06 | 4.035E+06 | 8.487E+06 | 1.019E+07 | 6.295E+06 | 4.955E+06 | 2.064E+07 | 1.191E+07 | 1.126E+07 | 1.050E+07 | 6.432E+06 | 9.533E+06 |
| 86 | 239.089 | 4.007 | -1 | 4.609E+07 | 5.900E+07 | 4.286E+07 | 3.269E+07 | 3.417E+07 | 1.450E+07 | 2.646E+07 | 3.226E+07 | 4.017E+07 | 2.150E+07 | 2.895E+07 | 1.594E+07 | 2.582E+07 | 1.869E+07 | 2.994E+07 |
| 90 | 332.913 | 12.737 | -1 | 8.527E+06 | 7.954E+06 | 7.791E+06 | 7.578E+06 | 7.313E+06 | 8.062E+06 | 9.519E+06 | 7.254E+06 | 7.137E+06 | 7.379E+06 | 7.474E+06 | 5.350E+06 | 7.146E+06 | 5.872E+06 | 5.196E+06 |
| 98 | 195.102 | 7.496 | -1 | 7.299E+06 | 7.963E+06 | 8.754E+06 | 7.901E+06 | 7.869E+06 | 1.039E+07 | 8.084E+06 | 8.925E+06 | 9.956E+06 | 7.796E+06 | 9.487E+06 | 9.670E+06 | 9.215E+06 | 9.850E+06 | 7.385E+06 |
| 100 | 440.099 | 7.644 | -1 | 7.618E+05 | 1.033E+06 | 8.027E+05 | 6.103E+06 | 7.039E+06 | 6.568E+06 | 7.521E+05 | 1.004E+07 | 6.021E+06 | 0.000E+00 | 0.000E+00 | 0.000E+00 | 0.000E+00 | 0.000E+00 | 4.393E+03 |
| 101 | 235.169 | 8.489 | -1 | 1.652E+07 | 4.258E+07 | 5.595E+07 | 1.487E+07 | 1.402E+07 | 1.062E+07 | 1.283E+07 | 1.674E+07 | 3.872E+07 | 1.530E+07 | 2.271E+07 | 1.560E+07 | 3.023E+07 | 1.687E+07 | 3.080E+07 |
| 106 | 670.153 | 5.750 | -1 | 6.706E+04 | 0.000E+00 | 0.000E+00 | 1.154E+06 | 4.720E+06 | 8.136E+06 | 2.482E+06 | 5.450E+06 | 2.559E+06 | 5.188E+05 | 2.193E+03 | 1.412E+06 | 4.735E+07 | 6.474E+07 | 5.366E+07 |
| 107 | 183.150 | 4.641 | -1 | 6.892E+06 | 1.039E+07 | 6.570E+06 | 9.673E+06 | 1.285E+07 | 9.956E+06 | 1.443E+07 | 1.369E+07 | 1.304E+07 | 8.888E+06 | 1.043E+07 | 8.856E+06 | 1.434E+07 | 1.554E+07 | 1.863E+07 |
| 108 | 281.088 | 8.127 | -1 | 2.689E+06 | 1.257E+06 | 2.271E+06 | 4.383E+06 | 6.562E+06 | 6.088E+06 | 3.602E+05 | 1.371E+07 | 3.696E+06 | 0.000E+00 | 0.000E+00 | 0.000E+00 | 0.000E+00 | 0.000E+00 | 0.000E+00 |
| 111 | 262.165 | 4.000 | -1 | 2.577E+07 | 3.661E+07 | 2.752E+07 | 1.999E+07 | 2.143E+07 | 9.250E+06 | 1.656E+07 | 1.932E+07 | 2.899E+07 | 1.558E+07 | 1.910E+07 | 1.175E+07 | 1.690E+07 | 1.364E+07 | 1.699E+07 |
| 114 | 244.191 | 5.975 | -1 | 6.499E+06 | 5.723E+06 | 5.979E+06 | 5.283E+06 | 4.595E+06 | 5.672E+06 | 5.985E+06 | 6.109E+06 | 5.116E+06 | 5.366E+06 | 5.584E+06 | 5.937E+06 | 6.668E+06 | 6.001E+06 | 6.051E+06 |
| 116 | 228.232 | 9.984 | -1 | 9.983E+06 | 6.164E+06 | 5.443E+06 | 7.335E+06 | 1.155E+07 | 7.873E+06 | 1.045E+07 | 7.555E+06 | 5.889E+06 | 4.127E+06 | 8.117E+06 | 5.643E+06 | 9.231E+06 | 5.790E+06 | 4.642E+06 |
| 118 | 219.175 | 8.820 | -1 | 6.059E+06 | 5.867E+06 | 5.441E+06 | 5.988E+06 | 5.905E+06 | 6.473E+06 | 6.299E+06 | 5.980E+06 | 6.247E+06 | 6.410E+06 | 6.050E+06 | 6.376E+06 | 6.268E+06 | 6.524E+06 | 6.425E+06 |
| 126 | 296.219 | 7.149 | -1 | 1.498E+06 | 3.095E+06 | 3.970E+06 | 2.412E+06 | 2.731E+06 | 5.226E+06 | 5.741E+06 | 3.727E+06 | 3.594E+06 | 2.751E+06 | 2.899E+06 | 4.041E+06 | 2.317E+06 | 4.685E+06 | 6.721E+06 |
| 137 | 431.183 | 10.115 | -1 | 5.387E+06 | 5.744E+06 | 5.105E+06 | 5.741E+06 | 5.915E+06 | 5.810E+06 | 5.344E+06 | 5.484E+06 | 5.675E+06 | 5.071E+06 | 5.750E+06 | 5.106E+06 | 4.898E+06 | 3.707E+06 | 5.683E+06 |
| 148 | 319.187 | 7.588 | -1 | 5.950E+06 | 9.618E+06 | 4.722E+06 | 4.900E+06 | 7.436E+06 | 4.491E+06 | 4.801E+06 | 3.815E+06 | 6.831E+06 | 4.007E+06 | 7.200E+06 | 3.230E+06 | 7.272E+06 | 6.121E+06 | 8.442E+06 |
| 152 | 707.875 | 4.385 | -1 | 0.000E+00 | 3.991E+04 | 8.342E+03 | 4.223E+06 | 5.706E+06 | 4.708E+06 | 7.484E+03 | 0.000E+00 | 4.877E+06 | 0.000E+00 | 0.000E+00 | 0.000E+00 | 0.000E+00 | 0.000E+00 | 0.000E+00 |
| 162 | 174.999 | 12.765 | -1 | 1.134E+07 | 1.099E+07 | 1.039E+07 | 9.765E+06 | 9.267E+06 | 8.528E+06 | 1.115E+07 | 1.034E+07 | 1.088E+07 | 1.066E+07 | 9.761E+06 | 8.456E+06 | 9.672E+06 | 8.650E+06 | 8.624E+06 |
| 164 | 338.147 | 5.707 | -1 | 3.074E+03 | 8.681E+03 | 3.455E+04 | 1.630E+05 | 3.362E+06 | 3.659E+06 | 3.036E+03 | 6.321E+04 | 1.199E+06 | 0.000E+00 | 0.000E+00 | 0.000E+00 | 0.000E+00 | 0.000E+00 | 0.000E+00 |
| 165 | 269.210 | 8.432 | -1 | 4.346E+06 | 6.847E+06 | 5.381E+06 | 7.390E+06 | 6.880E+06 | 6.776E+06 | 5.989E+06 | 8.941E+06 | 7.091E+06 | 4.423E+06 | 5.689E+06 | 4.829E+06 | 5.689E+06 | 6.744E+06 | 7.227E+06 |
| 173 | 451.246 | 9.774 | -1 | 1.309E+06 | 3.604E+06 | 3.278E+06 | 9.179E+06 | 4.974E+06 | 5.836E+06 | 4.701E+06 | 1.313E+06 | 8.371E+06 | 2.880E+06 | 1.428E+06 | 2.269E+06 | 2.938E+06 | 3.321E+06 | 4.120E+06 |
| 179 | 723.608 | 10.615 | -1 | 4.501E+06 | 3.952E+06 | 3.677E+06 | 3.263E+06 | 3.091E+06 | 3.593E+06 | 2.540E+06 | 2.614E+06 | 2.669E+06 | 2.951E+06 | 2.822E+06 | 2.174E+06 | 2.036E+06 | 2.117E+06 | 2.261E+06 |
| 184 | 434.186 | 7.527 | -1 | 1.889E+06 | 4.482E+06 | 1.956E+06 | 1.254E+06 | 3.714E+06 | 1.541E+06 | 2.428E+06 | 3.256E+06 | 1.964E+06 | 5.118E+06 | 5.158E+06 | 1.372E+06 | 3.156E+06 | 2.963E+06 | 5.135E+06 |
| 188 | 270.279 | 10.247 | -1 | 3.312E+06 | 3.008E+06 | 3.090E+06 | 3.614E+06 | 3.957E+06 | 3.906E+06 | 3.800E+06 | 3.592E+06 | 2.427E+06 | 2.279E+06 | 3.207E+06 | 3.012E+06 | 3.573E+06 | 3.127E+06 | 2.389E+06 |
| 193 | 242.284 | 7.258 | -1 | 9.235E+06 | 9.065E+06 | 7.449E+06 | 6.648E+06 | 6.053E+06 | 4.885E+06 | 4.314E+06 | 4.488E+06 | 3.709E+06 | 3.425E+06 | 3.288E+06 | 2.819E+06 | 2.923E+06 | 2.655E+06 | 2.394E+06 |
| 195 | 312.326 | 10.525 | -1 | 2.587E+06 | 1.619E+06 | 1.479E+06 | 2.513E+06 | 3.260E+06 | 2.613E+06 | 2.961E+06 | 2.613E+06 | 1.389E+06 | 1.350E+06 | 2.438E+06 | 2.235E+06 | 3.074E+06 | 3.262E+06 | 1.382E+06 |
| 198 | 499.165 | 7.526 | -1 | 4.363E+06 | 6.541E+06 | 4.679E+06 | 4.377E+06 | 5.443E+06 | 4.443E+06 | 5.145E+06 | 6.026E+06 | 5.146E+06 | 7.000E+06 | 6.652E+06 | 3.938E+06 | 6.736E+06 | 4.975E+06 | 6.963E+06 |
| 203 | 122.547 | 1.761 | -1 | 3.126E+06 | 1.922E+06 | 5.477E+06 | 1.948E+06 | 1.177E+06 | 2.690E+06 | 1.503E+06 | 2.503E+06 | 1.749E+06 | 7.427E+05 | 1.016E+06 | 9.553E+05 | 2.711E+06 | 6.220E+05 | 1.413E+06 |
| 212 | 409.201 | 10.114 | -1 | 3.985E+06 | 3.030E+06 | 3.375E+06 | 4.668E+06 | 4.575E+06 | 4.277E+06 | 4.168E+06 | 4.185E+06 | 3.995E+06 | 4.296E+06 | 4.320E+06 | 4.027E+06 | 5.096E+06 | 5.843E+06 | 5.462E+06 |
| 214 | 270.279 | 9.930 | -1 | 7.106E+06 | 6.432E+06 | 5.841E+06 | 6.314E+06 | 5.561E+06 | 5.952E+06 | 5.318E+06 | 5.573E+06 | 5.127E+06 | 4.602E+06 | 4.635E+06 | 4.899E+06 | 5.332E+06 | 5.931E+06 | 4.366E+06 |
| 217 | 266.248 | 6.949 | -1 | 0.000E+00 | 0.000E+00 | 0.000E+00 | 9.227E+06 | 3.648E+06 | 4.153E+06 | 0.000E+00 | 0.000E+00 | 2.059E+06 | 0.000E+00 | 0.000E+00 | 0.000E+00 | 0.000E+00 | 0.000E+00 | 0.000E+00 |

| Feature identifiers | | | | XIC | | | | | | | | | | | | | | |
|---|---|---|---|---|---|---|---|---|---|---|---|---|---|---|---|---|---|---|
| ClusterIdx | m/z | RT [min] | Subnetwork | 4TUB34_I | 4TUB34_II | 4TUB34_III | 4TUB81_I | 4TUB81_II | 4TUB81_III | 4TUB93_I | 4TUB93_II | 4TUB93_III | 5TUB23_I | 5TUB23_II | 5TUB23_III | 5TUB48_I | 5TUB48_II | 5TUB48_III |
| 222 | 495.300 | 10.647 | -1 | 7.111E+05 | 7.837E+05 | 1.031E+06 | 1.310E+06 | 1.318E+06 | 1.728E+06 | 1.401E+06 | 1.430E+06 | 7.729E+05 | 9.106E+05 | 1.107E+06 | 1.465E+06 | 1.111E+06 | 1.461E+06 | 7.560E+05 |
| 225 | 330.300 | 9.151 | -1 | 4.160E+05 | 1.233E+06 | 1.875E+06 | 2.304E+06 | 2.678E+06 | 2.609E+06 | 2.189E+06 | 2.115E+06 | 9.420E+06 | 1.059E+05 | 3.611E+04 | 1.524E+05 | 1.925E+04 | 0.000E+00 | 9.719E+03 |
| 232 | 250.178 | 6.460 | -1 | 5.899E+06 | 1.657E+07 | 1.698E+07 | 8.318E+06 | 9.192E+06 | 1.516E+07 | 1.915E+07 | 1.109E+07 | 1.311E+07 | 7.923E+06 | 9.187E+06 | 1.127E+07 | 7.795E+06 | 1.356E+07 | 1.963E+07 |
| 234 | 555.250 | 4.014 | -1 | 0.000E+00 | 0.000E+00 | 0.000E+00 | 1.181E+06 | 3.007E+06 | 2.790E+06 | 0.000E+00 | 0.000E+00 | 8.193E+05 | 0.000E+00 | 0.000E+00 | 0.000E+00 | 0.000E+00 | 0.000E+00 | 0.000E+00 |
| 236 | 149.988 | 12.784 | -1 | 1.102E+07 | 7.763E+06 | 8.377E+06 | 8.258E+06 | 8.194E+06 | 7.816E+06 | 8.181E+06 | 8.248E+06 | 7.971E+06 | 8.071E+06 | 7.754E+06 | 8.678E+06 | 7.475E+06 | 8.270E+06 | 7.474E+06 |
| 237 | 640.348 | 4.374 | -1 | 0.000E+00 | 4.674E+04 | 1.599E+04 | 2.720E+06 | 2.090E+06 | 2.733E+06 | 2.579E+04 | 1.741E+04 | 3.553E+06 | 0.000E+00 | 2.163E+04 | 8.712E+03 | 3.790E+04 | 0.000E+00 | 3.834E+04 |
| 252 | 157.084 | 0.657 | -1 | 4.170E+07 | 3.494E+07 | 2.608E+07 | 1.574E+07 | 1.384E+07 | 1.094E+07 | 8.229E+06 | 1.086E+07 | 1.504E+07 | 3.992E+07 | 2.771E+07 | 1.734E+07 | 1.933E+07 | 1.594E+07 | 1.477E+07 |
| 254 | 111.039 | 13.492 | -1 | 3.470E+06 | 4.653E+05 | 9.390E+05 | 1.134E+06 | 1.896E+06 | 1.779E+06 | 4.317E+06 | 1.957E+06 | 1.958E+06 | 1.868E+06 | 1.984E+06 | 2.300E+06 | 1.245E+06 | 1.779E+06 | 1.560E+06 |
| 256 | 297.241 | 9.395 | -1 | 2.886E+06 | 3.287E+06 | 4.950E+06 | 3.010E+06 | 2.762E+06 | 3.091E+06 | 2.973E+06 | 6.585E+06 | 4.850E+06 | 3.229E+06 | 3.342E+06 | 3.149E+06 | 2.919E+06 | 3.042E+06 | 2.925E+06 |
| 259 | 249.185 | 8.504 | -1 | 2.958E+06 | 4.710E+06 | 5.445E+06 | 3.407E+06 | 3.589E+06 | 3.018E+06 | 3.275E+06 | 3.144E+06 | 4.818E+06 | 2.676E+06 | 3.669E+06 | 3.382E+06 | 3.778E+06 | 3.378E+06 | 3.917E+06 |
| 260 | 290.269 | 6.924 | -1 | 2.388E+06 | 2.373E+06 | 2.493E+06 | 1.991E+06 | 2.402E+06 | 2.438E+06 | 2.240E+06 | 2.323E+06 | 2.134E+06 | 2.147E+06 | 2.311E+06 | 2.333E+06 | 1.885E+06 | 2.376E+06 | 2.052E+06 |
| 265 | 535.153 | 8.517 | -1 | 0.000E+00 | 2.716E+05 | 2.662E+05 | 7.455E+05 | 3.082E+06 | 2.535E+06 | 4.078E+05 | 7.012E+06 | 0.000E+00 | 0.000E+00 | 0.000E+00 | 0.000E+00 | 0.000E+00 | 0.000E+00 | 0.000E+00 |
| 267 | 453.168 | 7.817 | -1 | 8.395E+06 | 5.863E+06 | 5.011E+06 | 2.687E+06 | 1.779E+06 | 2.654E+06 | 1.848E+06 | 7.072E+06 | 2.166E+06 | 3.612E+06 | 4.723E+06 | 4.772E+06 | 5.521E+06 | 3.052E+06 | 6.729E+06 |
| 271 | 283.104 | 5.343 | -1 | 1.208E+06 | 1.723E+06 | 2.552E+06 | 1.349E+06 | 4.445E+06 | 2.987E+06 | 3.243E+06 | 8.398E+06 | 2.885E+06 | 0.000E+00 | 0.000E+00 | 0.000E+00 | 0.000E+00 | 0.000E+00 | 0.000E+00 |
| 274 | 537.311 | 10.526 | -1 | 2.105E+06 | 2.273E+06 | 1.923E+06 | 1.288E+06 | 1.212E+06 | 1.430E+06 | 9.855E+05 | 1.967E+06 | 1.853E+06 | 3.484E+06 | 3.843E+06 | 2.623E+06 | 3.123E+06 | 4.464E+06 | 3.469E+06 |
| 279 | 149.024 | 6.761 | -1 | 5.111E+06 | 1.500E+07 | 1.316E+07 | 4.892E+06 | 4.258E+06 | 2.955E+06 | 3.601E+06 | 3.145E+06 | 1.329E+07 | 3.958E+06 | 4.236E+06 | 4.328E+06 | 4.543E+06 | 3.539E+06 | 9.138E+06 |
| 281 | 467.147 | 6.891 | -1 | 5.299E+06 | 8.311E+06 | 8.173E+06 | 4.575E+06 | 6.007E+06 | 5.654E+06 | 5.969E+06 | 5.122E+06 | 6.432E+06 | 7.507E+06 | 8.055E+06 | 6.583E+06 | 5.990E+06 | 8.993E+06 | 6.757E+06 |
| 285 | 408.309 | 10.166 | -1 | 1.858E+06 | 1.968E+06 | 3.975E+06 | 4.651E+06 | 3.037E+06 | 1.596E+06 | 1.389E+06 | 6.233E+05 | 5.130E+05 | 1.913E+06 | 2.001E+06 | 1.969E+06 | 2.177E+06 | 3.948E+06 | 2.017E+06 |
| 286 | 291.195 | 8.162 | -1 | 1.313E+06 | 1.887E+06 | 2.061E+06 | 2.807E+06 | 2.840E+06 | 2.266E+06 | 2.710E+06 | 2.619E+06 | 2.802E+06 | 1.896E+06 | 2.621E+06 | 2.473E+06 | 2.256E+06 | 3.358E+06 | 3.093E+06 |
| 287 | 573.110 | 8.520 | -1 | 0.000E+00 | 1.117E+05 | 1.023E+05 | 4.534E+05 | 2.258E+06 | 1.839E+06 | 2.112E+05 | 4.284E+06 | 0.000E+00 | 0.000E+00 | 0.000E+00 | 0.000E+00 | 0.000E+00 | 0.000E+00 | 0.000E+00 |
| 299 | 437.194 | 8.754 | -1 | 4.979E+06 | 5.655E+06 | 3.912E+06 | 2.847E+06 | 2.680E+06 | 3.012E+06 | 2.825E+06 | 2.384E+06 | 1.047E+07 | 3.248E+06 | 2.794E+06 | 3.863E+06 | 4.008E+06 | 2.935E+06 | 3.212E+06 |
| 300 | 604.812 | 4.484 | -1 | 0.000E+00 | 2.163E+04 | 0.000E+00 | 8.639E+05 | 1.594E+06 | 1.324E+06 | 0.000E+00 | 0.000E+00 | 6.122E+05 | 0.000E+00 | 0.000E+00 | 0.000E+00 | 0.000E+00 | 0.000E+00 | 0.000E+00 |
| 304 | 365.137 | 8.808 | -1 | 8.199E+06 | 1.074E+07 | 9.651E+06 | 7.512E+06 | 9.738E+06 | 8.028E+06 | 7.646E+06 | 8.820E+06 | 9.968E+06 | 8.487E+06 | 8.945E+06 | 7.342E+06 | 9.085E+06 | 7.843E+06 | 6.964E+06 |
| 305 | 480.273 | 3.605 | -1 | 0.000E+00 | 1.485E+04 | 0.000E+00 | 7.019E+05 | 4.002E+06 | 5.793E+05 | 0.000E+00 | 0.000E+00 | 3.714E+05 | 0.000E+00 | 0.000E+00 | 0.000E+00 | 3.290E+04 | 0.000E+00 | 0.000E+00 |
| 308 | 659.288 | 10.020 | -1 | 1.096E+07 | 2.280E+06 | 3.518E+06 | 1.982E+06 | 1.643E+06 | 1.195E+06 | 6.968E+05 | 1.135E+07 | 6.476E+05 | 3.837E+06 | 3.264E+06 | 1.147E+07 | 1.972E+07 | 7.589E+06 | 1.120E+07 |
| 309 | 165.076 | 4.458 | -1 | 2.881E+06 | 1.975E+06 | 1.555E+06 | 1.827E+06 | 2.509E+06 | 2.150E+06 | 1.595E+06 | 3.829E+06 | 1.674E+06 | 3.766E+06 | 3.799E+06 | 2.122E+06 | 3.804E+06 | 2.894E+06 | 4.368E+06 |
| 320 | 341.267 | 9.373 | -1 | 2.259E+06 | 2.299E+06 | 2.164E+06 | 2.689E+06 | 2.245E+06 | 2.134E+06 | 2.986E+06 | 1.001E+07 | 6.382E+06 | 2.806E+06 | 3.367E+06 | 2.220E+06 | 2.297E+06 | 2.265E+06 | 2.563E+06 |
| 324 | 270.315 | 9.426 | -1 | 5.602E+06 | 1.239E+07 | 8.743E+06 | 1.209E+07 | 6.571E+06 | 8.624E+06 | 7.487E+06 | 9.793E+06 | 4.055E+06 | 1.331E+07 | 9.454E+06 | 1.977E+07 | 1.116E+07 | 2.509E+07 | 9.021E+06 |
| 325 | 326.378 | 9.409 | -1 | 7.261E+06 | 5.864E+06 | 7.211E+06 | 3.730E+06 | 3.738E+06 | 3.665E+06 | 3.418E+06 | 3.949E+06 | 5.132E+06 | 7.170E+06 | 4.584E+06 | 7.511E+06 | 5.326E+06 | 4.361E+06 | 4.058E+06 |
| 327 | 131.049 | 0.513 | -1 | 3.829E+07 | 1.680E+07 | 3.367E+07 | 1.280E+05 | 2.928E+06 | 2.836E+06 | 6.879E+06 | 9.783E+05 | 1.119E+07 | 1.224E+07 | 1.875E+07 | 1.486E+07 | 3.072E+07 | 1.612E+06 | 4.304E+06 |
| 331 | 260.165 | 6.499 | -1 | 9.551E+06 | 8.223E+06 | 1.438E+07 | 1.267E+06 | 1.571E+06 | 1.409E+06 | 1.397E+06 | 1.931E+06 | 1.673E+06 | 5.731E+06 | 9.382E+06 | 6.421E+06 | 7.427E+06 | 1.718E+06 | 1.797E+06 |
| 334 | 469.184 | 6.409 | -1 | 1.480E+06 | 1.846E+06 | 1.542E+06 | 1.007E+06 | 9.779E+05 | 1.439E+06 | 1.521E+06 | 2.216E+06 | 2.363E+06 | 1.179E+06 | 1.408E+06 | 1.212E+06 | 1.533E+06 | 2.231E+06 | 2.598E+06 |
| 335 | 469.163 | 6.314 | -1 | 2.282E+06 | 2.015E+06 | 1.979E+06 | 1.595E+06 | 1.605E+06 | 1.236E+06 | 1.566E+06 | 1.887E+06 | 1.462E+06 | 1.816E+06 | 2.056E+06 | 1.632E+06 | 2.148E+06 | 2.499E+06 | 1.641E+06 |
| 338 | 499.780 | 3.714 | -1 | 0.000E+00 | 0.000E+00 | 0.000E+00 | 3.407E+06 | 3.730E+06 | 2.638E+06 | 0.000E+00 | 0.000E+00 | 4.296E+06 | 0.000E+00 | 0.000E+00 | 0.000E+00 | 0.000E+00 | 0.000E+00 | 0.000E+00 |
| 339 | 615.581 | 10.604 | -1 | 1.458E+07 | 1.497E+07 | 1.513E+07 | 1.601E+07 | 1.530E+07 | 1.575E+07 | 1.542E+07 | 1.572E+07 | 1.538E+07 | 1.592E+07 | 1.494E+07 | 1.475E+07 | 1.382E+07 | 1.362E+07 | 1.421E+07 |
| 340 | 354.553 | 5.749 | -1 | 6.778E+03 | 0.000E+00 | 0.000E+00 | 1.242E+05 | 1.061E+06 | 1.806E+06 | 5.594E+05 | 1.219E+06 | 6.496E+05 | 5.902E+04 | 0.000E+00 | 2.891E+05 | 1.117E+07 | 1.351E+07 | 1.177E+07 |
| 345 | 236.142 | 5.355 | -1 | 2.494E+06 | 4.555E+06 | 3.094E+06 | 2.470E+06 | 3.080E+06 | 1.755E+06 | 2.217E+06 | 1.741E+06 | 2.742E+06 | 1.632E+06 | 2.960E+06 | 1.799E+06 | 3.097E+06 | 2.845E+06 | 3.606E+06 |
| 351 | 451.173 | 6.881 | -1 | 1.431E+06 | 7.372E+06 | 7.386E+06 | 3.801E+06 | 5.115E+06 | 4.677E+06 | 5.218E+06 | 4.292E+06 | 5.939E+06 | 7.978E+06 | 8.019E+06 | 6.329E+06 | 1.380E+06 | 8.132E+06 | 6.098E+06 |
| 356 | 310.173 | 6.844 | -1 | 1.466E+06 | 3.081E+06 | 1.301E+06 | 1.004E+06 | 1.379E+06 | 1.435E+06 | 1.396E+06 | 1.697E+06 | 1.359E+06 | 1.365E+06 | 3.345E+06 | 1.235E+06 | 1.502E+06 | 3.114E+06 | 1.579E+06 |
| 365 | 519.261 | 9.835 | -1 | 4.167E+06 | 1.822E+06 | 2.005E+06 | 2.589E+06 | 2.638E+06 | 3.102E+06 | 3.671E+06 | 4.102E+06 | 1.637E+06 | 1.122E+07 | 9.302E+06 | 1.004E+07 | 6.655E+06 | 8.005E+06 | 2.951E+06 |
| 368 | 273.167 | 4.221 | -1 | 7.320E+06 | 8.688E+06 | 6.126E+06 | 5.400E+06 | 5.409E+06 | 4.352E+06 | 5.615E+06 | 5.228E+06 | 5.748E+06 | 5.055E+06 | 5.392E+06 | 4.524E+06 | 5.036E+06 | 4.757E+06 | 6.253E+06 |
| 376 | 288.181 | 4.136 | -1 | 5.335E+06 | 7.118E+06 | 3.793E+06 | 2.451E+06 | 3.913E+06 | 1.971E+06 | 3.462E+06 | 3.195E+06 | 3.778E+06 | 3.944E+06 | 5.121E+06 | 3.121E+06 | 5.568E+06 | 4.175E+06 | 5.821E+06 |
| 379 | 496.301 | 10.257 | -1 | 6.295E+05 | 1.113E+06 | 2.060E+06 | 8.203E+05 | 8.670E+05 | 7.168E+05 | 5.965E+05 | 9.557E+05 | 1.029E+06 | 1.612E+06 | 2.215E+06 | 2.141E+06 | 2.083E+06 | 1.651E+06 | 1.664E+06 |
| 380 | 497.280 | 9.835 | -1 | 3.560E+06 | 1.211E+06 | 1.872E+06 | 2.200E+06 | 2.316E+06 | 2.654E+06 | 3.224E+06 | 3.726E+06 | 9.866E+05 | 1.194E+07 | 9.367E+06 | 1.179E+07 | 9.342E+06 | 1.462E+07 | 3.537E+06 |
| 385 | 188.069 | 0.519 | -1 | 4.515E+07 | 1.088E+07 | 3.044E+07 | 1.179E+05 | 2.185E+06 | 2.010E+06 | 2.166E+04 | 9.354E+04 | 5.867E+04 | 1.423E+07 | 1.258E+07 | 9.504E+06 | 3.976E+07 | 3.505E+04 | 2.277E+06 |
| 395 | 721.506 | 12.424 | -1 | 2.526E+06 | 3.285E+06 | 3.394E+06 | 2.422E+06 | 2.182E+06 | 2.558E+06 | 2.576E+06 | 2.309E+06 | 3.133E+06 | 5.358E+06 | 6.235E+06 | 7.239E+06 | 6.597E+06 | 1.036E+07 | 5.555E+06 |
| 397 | 459.263 | 10.315 | -1 | 7.454E+05 | 4.844E+05 | 3.692E+05 | 6.941E+05 | 4.986E+05 | 7.460E+05 | 5.802E+05 | 8.053E+05 | 5.922E+05 | 6.555E+05 | 8.894E+05 | 5.960E+05 | 7.191E+05 | 1.153E+06 | 1.035E+06 |
| 398 | 445.291 | 10.157 | -1 | 1.075E+06 | 1.398E+06 | 1.552E+06 | 5.552E+05 | 5.436E+06 | 1.637E+06 | 3.590E+03 | 2.543E+06 | 3.484E+06 | 1.913E+07 | 1.219E+06 | 3.405E+06 | 2.579E+05 | 1.904E+04 | 3.042E+05 |
| 403 | 763.517 | 12.425 | -1 | 2.274E+06 | 3.112E+06 | 2.923E+06 | 2.093E+06 | 2.292E+06 | 2.230E+06 | 2.233E+06 | 2.065E+06 | 2.725E+06 | 4.923E+06 | 6.028E+06 | 6.766E+06 | 6.023E+06 | 1.018E+07 | 5.277E+06 |
| 405 | 528.788 | 4.417 | -1 | 0.000E+00 | 0.000E+00 | 0.000E+00 | 8.550E+05 | 1.585E+06 | 8.796E+05 | 0.000E+00 | 0.000E+00 | 7.533E+05 | 0.000E+00 | 5.164E+04 | 0.000E+00 | 0.000E+00 | 0.000E+00 | 0.000E+00 |
| 407 | 455.335 | 10.125 | -1 | 8.855E+05 | 4.433E+05 | 3.726E+05 | 1.446E+06 | 7.585E+05 | 8.859E+05 | 7.746E+05 | 8.899E+05 | 4.218E+05 | 5.763E+05 | 4.368E+05 | 6.915E+05 | 2.603E+06 | 4.017E+06 | 8.879E+05 |
| 410 | 830.949 | 3.185 | -1 | 2.085E+06 | 3.757E+06 | 3.179E+06 | 1.854E+06 | 2.153E+06 | 1.525E+06 | 2.105E+06 | 3.120E+06 | 2.916E+06 | 2.311E+06 | 1.766E+06 | 5.338E+06 | 5.046E+06 | 6.450E+06 | 2.710E+06 |
| 421 | 514.951 | 4.135 | -1 | 0.000E+00 | 0.000E+00 | 0.000E+00 | 6.040E+05 | 1.513E+06 | 1.069E+06 | 8.376E+03 | 0.000E+00 | 1.279E+06 | 0.000E+00 | 0.000E+00 | 0.000E+00 | 0.000E+00 | 0.000E+00 | 0.000E+00 |
| 423 | 409.180 | 7.399 | -1 | 7.717E+06 | 1.144E+06 | 3.213E+06 | 1.226E+06 | 1.005E+06 | 6.677E+05 | 4.126E+05 | 2.522E+06 | 7.222E+05 | 1.558E+06 | 1.441E+06 | 4.141E+06 | 9.928E+06 | 1.293E+06 | 2.489E+06 |
| 425 | 707.607 | 12.056 | -1 | 1.898E+06 | 1.784E+06 | 2.103E+06 | 1.579E+06 | 1.455E+06 | 1.412E+06 | 1.387E+06 | 1.194E+06 | 1.310E+06 | 1.163E+06 | 1.137E+06 | 8.618E+05 | 1.086E+06 | 1.184E+06 | 1.115E+06 |
| 428 | 366.633 | 3.310 | -1 | 2.754E+04 | 1.802E+04 | 3.339E+04 | 3.750E+05 | 3.347E+06 | 2.833E+06 | 6.560E+03 | 0.000E+00 | 2.662E+07 | 0.000E+00 | 2.353E+04 | 0.000E+00 | 7.021E+03 | 3.174E+04 | 1.720E+04 |
| 430 | 379.862 | 4.127 | -1 | 0.000E+00 | 3.221E+05 | 7.119E+04 | 3.161E+05 | 1.137E+06 | 1.234E+06 | 0.000E+00 | 0.000E+00 | 2.686E+06 | 0.000E+00 | 0.000E+00 | 0.000E+00 | 0.000E+00 | 0.000E+00 | 0.000E+00 |
| 436 | 267.137 | 8.329 | -1 | 2.844E+05 | 8.862E+05 | 1.365E+06 | 1.196E+06 | 1.304E+06 | 9.796E+05 | 1.381E+06 | 6.881E+05 | 3.375E+06 | 4.183E+06 | 2.240E+06 | 3.258E+06 | 2.256E+06 | 2.878E+06 | 2.157E+06 |
| 449 | 658.341 | 3.958 | -1 | 0.000E+00 | 0.000E+00 | 0.000E+00 | 1.893E+06 | 1.721E+06 | 9.852E+05 | 0.000E+00 | 0.000E+00 | 1.580E+06 | 0.000E+00 | 0.000E+00 | 0.000E+00 | 0.000E+00 | 0.000E+00 | 0.000E+00 |
| 451 | 254.660 | 3.065 | -1 | 0.000E+00 | 0.000E+00 | 0.000E+00 | 3.035E+06 | 2.622E+06 | 2.017E+06 | 0.000E+00 | 0.000E+00 | 8.306E+06 | 0.000E+00 | 0.000E+00 | 0.000E+00 | 0.000E+00 | 0.000E+00 | 0.000E+00 |
| 453 | 441.346 | 7.492 | -1 | 6.204E+05 | 5.761E+05 | 5.387E+05 | 6.325E+05 | 6.790E+05 | 5.245E+05 | 6.293E+05 | 6.060E+05 | 5.928E+05 | 5.266E+05 | 4.556E+05 | 8.048E+05 | 5.496E+05 | 6.183E+05 | 4.245E+05 |
| 454 | 409.163 | 6.914 | -1 | 4.021E+07 | 5.057E+05 | 2.456E+07 | 2.406E+06 | 1.637E+06 | 8.291E+05 | 4.080E+05 | 8.557E+05 | 7.617E+05 | 1.118E+06 | 4.609E+05 | 6.634E+05 | 4.637E+07 | 1.010E+06 | 6.484E+05 |
| 470 | 469.163 | 6.021 | -1 | 1.024E+06 | 1.023E+06 | 1.281E+06 | 6.828E+05 | 6.765E+05 | 4.491E+05 | 7.502E+05 | 9.562E+05 | 8.223E+05 | 9.219E+05 | 1.180E+06 | 7.904E+05 | 1.031E+06 | 1.174E+06 | 7.538E+05 |
| 474 | 501.170 | 0.519 | -1 | 0.000E+00 | 0.000E+00 | 0.000E+00 | 0.000E+00 | 0.000E+00 | 0.000E+00 | 4.950E+08 | 3.374E+08 | 3.067E+08 | 4.187E+05 | 2.096E+06 | 4.376E+04 | 0.000E+00 | 1.132E+05 | 5.884E+04 |
| 485 | 517.142 | 0.507 | -1 | 0.000E+00 | 0.000E+00 | 0.000E+00 | 0.000E+00 | 0.000E+00 | 0.000E+00 | 4.786E+07 | 3.727E+07 | 4.247E+07 | 0.000E+00 | 2.108E+04 | 0.000E+00 | 0.000E+00 | 0.000E+00 | 0.000E+00 |
| 496 | 226.012 | 12.739 | -1 | 8.642E+06 | 1.340E+07 | 1.395E+07 | 1.426E+07 | 1.401E+07 | 1.119E+07 | 1.289E+07 | 1.230E+07 | 1.383E+07 | 1.347E+07 | 1.437E+07 | 1.216E+07 | 1.209E+07 | 1.059E+07 | 1.173E+07 |
| 502 | 102.034 | 0.568 | -1 | 1.492E+07 | 4.151E+05 | 3.601E+06 | 1.104E+07 | 3.409E+06 | 8.856E+06 | 2.488E+06 | 8.536E+06 | 1.076E+07 | 8.638E+06 | 1.153E+07 | 5.620E+05 | 8.540E+06 | 9.889E+06 | 1.050E+07 |
| 503 | 228.207 | 4.644 | -1 | 3.246E+06 | 4.879E+06 | 3.070E+06 | 4.571E+06 | 6.253E+06 | 4.864E+06 | 6.980E+06 | 6.860E+06 | 6.482E+06 | 4.386E+06 | 5.154E+06 | 4.330E+06 | 7.031E+06 | 7.450E+06 | 9.430E+06 |
| 508 | 1011.286 | 0.501 | -1 | 0.000E+00 | 0.000E+00 | 0.000E+00 | 0.000E+00 | 0.000E+00 | 0.000E+00 | 6.084E+06 | 1.066E+06 | 1.344E+06 | 0.000E+00 | 0.000E+00 | 0.000E+00 | 0.000E+00 | 0.000E+00 | 0.000E+00 |
| 509 | 385.095 | 3.893 | -1 | 0.000E+00 | 2.046E+06 | 3.153E+05 | 0.000E+00 | 0.000E+00 | 0.000E+00 | 3.890E+06 | 6.464E+05 | 0.000E+00 | 2.041E+06 | 5.134E+05 | 1.799E+06 | 0.000E+00 | 2.764E+06 | 2.630E+06 |

| Feature identifiers | | | | XIC | | | | | | | | | | | | | | |
|---|---|---|---|---|---|---|---|---|---|---|---|---|---|---|---|---|---|---|
| ClusterIdx | m/z | RT [min] | Subnetwork | 4TUB34_I | 4TUB34_II | 4TUB34_III | 4TUB81_I | 4TUB81_II | 4TUB81_III | 4TUB93_I | 4TUB93_II | 4TUB93_III | 5TUB23_I | 5TUB23_II | 5TUB23_III | 5TUB48_I | 5TUB48_II | 5TUB48_III |
| 525 | 269.210 | 8.072 | -1 | 7.102E+04 | 1.038E+05 | 3.834E+05 | 3.201E+06 | 5.286E+05 | 1.516E+05 | 6.147E+06 | 1.686E+07 | 8.387E+07 | 3.819E+06 | 5.513E+06 | 3.218E+05 | 5.674E+05 | 8.583E+06 | 8.837E+06 |
| 538 | 844.952 | 3.465 | -1 | 1.013E+06 | 2.205E+06 | 1.851E+06 | 1.191E+06 | 1.383E+06 | 7.462E+05 | 2.278E+06 | 2.468E+05 | 2.088E+06 | 1.713E+06 | 8.025E+05 | 1.423E+06 | 2.697E+06 | 2.310E+06 | 4.262E+06 |
| 543 | 453.188 | 6.313 | -1 | 1.534E+06 | 1.418E+06 | 1.348E+06 | 1.078E+06 | 9.829E+05 | 9.293E+05 | 9.768E+05 | 1.222E+06 | 9.268E+05 | 1.158E+06 | 7.843E+05 | 1.018E+06 | 1.436E+06 | 1.619E+06 | 9.899E+05 |
| 547 | 372.717 | 12.740 | -1 | 0.000E+00 | 0.000E+00 | 0.000E+00 | 1.289E+06 | 1.161E+06 | 0.000E+00 | 3.645E+05 | 0.000E+00 | 0.000E+00 | 3.684E+04 | 9.884E+05 | 2.146E+05 | 1.006E+06 | 1.781E+06 | 0.000E+00 |
| 549 | 287.114 | 4.597 | -1 | 6.610E+05 | 2.069E+06 | 1.975E+06 | 4.261E+05 | 5.163E+05 | 6.260E+05 | 1.110E+06 | 8.857E+05 | 2.518E+06 | 9.620E+05 | 1.038E+06 | 1.340E+06 | 1.349E+06 | 1.990E+06 | 1.935E+06 |
| 551 | 385.294 | 9.364 | -1 | 8.700E+05 | 8.865E+05 | 7.556E+05 | 1.239E+06 | 9.402E+05 | 8.161E+05 | 1.271E+06 | 5.934E+06 | 3.946E+06 | 1.503E+06 | 1.373E+06 | 8.507E+05 | 1.046E+06 | 8.567E+05 | 1.029E+06 |
| 557 | 858.957 | 3.760 | -1 | 5.056E+05 | 4.376E+05 | 3.854E+05 | 2.604E+05 | 5.813E+05 | 2.688E+05 | 2.161E+06 | 4.000E+03 | 1.591E+06 | 1.245E+06 | 3.816E+05 | 2.645E+05 | 1.289E+06 | 2.911E+05 | 2.486E+06 |
| 563 | 429.319 | 9.342 | -1 | 9.915E+05 | 7.619E+05 | 4.545E+05 | 1.106E+06 | 6.539E+05 | 6.851E+05 | 1.050E+06 | 4.830E+06 | 3.503E+06 | 1.501E+06 | 1.083E+06 | 6.133E+05 | 1.052E+06 | 8.031E+05 | 8.680E+05 |
| 566 | 472.175 | 4.688 | -1 | 7.627E+05 | 1.460E+06 | 6.089E+05 | 1.029E+05 | 0.000E+00 | 0.000E+00 | 8.655E+05 | 2.233E+06 | 0.000E+00 | 8.043E+05 | 9.828E+05 | 1.695E+06 | 2.721E+05 | 2.040E+06 | 2.681E+06 |
| 569 | 294.074 | 0.519 | -1 | 4.846E+04 | 1.052E+07 | 7.356E+06 | 1.331E+05 | 1.451E+05 | 1.559E+05 | 3.292E+05 | 3.628E+06 | 4.227E+06 | 3.458E+06 | 3.923E+06 | 5.995E+06 | 6.849E+06 | 8.031E+06 | 1.181E+07 |
| 575 | 453.189 | 6.022 | -1 | 1.012E+06 | 1.003E+06 | 1.014E+06 | 4.933E+05 | 4.766E+05 | 3.352E+05 | 4.859E+05 | 7.289E+05 | 6.031E+05 | 7.229E+05 | 1.003E+06 | 6.107E+05 | 7.863E+05 | 8.856E+05 | 5.884E+05 |
| 586 | 191.086 | 3.681 | -1 | 1.535E+06 | 3.148E+06 | 2.238E+06 | 1.731E+06 | 1.712E+06 | 1.790E+06 | 9.305E+05 | 1.377E+07 | 2.856E+07 | 5.632E+04 | 5.440E+03 | 0.000E+00 | 2.096E+04 | 2.433E+04 | 0.000E+00 |
| 592 | 359.104 | 3.680 | -1 | 1.135E+06 | 2.187E+06 | 1.264E+06 | 1.382E+06 | 1.120E+06 | 1.075E+06 | 9.984E+05 | 9.733E+06 | 1.095E+07 | 0.000E+00 | 0.000E+00 | 0.000E+00 | 0.000E+00 | 4.102E+03 | 0.000E+00 |
| 593 | 511.165 | 3.612 | -1 | 4.582E+06 | 1.601E+06 | 1.358E+06 | 5.419E+05 | 1.242E+06 | 8.491E+05 | 2.763E+05 | 4.420E+06 | 7.429E+06 | 4.380E+05 | 1.036E+06 | 2.033E+05 | 2.882E+06 | 2.721E+06 | 1.239E+06 |
| 594 | 700.315 | 10.020 | -1 | 3.009E+06 | 4.401E+05 | 7.801E+05 | 5.289E+05 | 2.551E+05 | 2.792E+05 | 1.824E+05 | 2.726E+06 | 1.518E+05 | 7.169E+05 | 6.426E+05 | 2.380E+06 | 4.918E+06 | 1.553E+06 | 2.498E+06 |
| 597 | 225.149 | 6.492 | -1 | 5.252E+06 | 9.335E+06 | 1.649E+07 | 1.579E+06 | 4.779E+05 | 3.082E+05 | 5.985E+05 | 3.434E+06 | 5.120E+06 | 2.837E+05 | 1.548E+06 | 2.790E+05 | 8.373E+06 | 6.836E+05 | 2.852E+06 |
| 599 | 131.049 | 0.515 | -1 | 2.810E+07 | 1.910E+06 | 3.590E+06 | 1.280E+05 | 2.021E+06 | 6.672E+05 | 6.390E+05 | 3.667E+06 | 8.604E+06 | 1.600E+07 | 1.308E+07 | 1.074E+07 | 4.486E+07 | 2.662E+06 | 2.498E+05 |
| 601 | 357.262 | 8.098 | -1 | 2.599E+04 | 1.934E+04 | 1.541E+04 | 3.115E+05 | 3.911E+04 | 3.743E+04 | 2.577E+05 | 9.171E+06 | 1.609E+07 | 5.894E+05 | 1.323E+06 | 3.124E+05 | 7.631E+05 | 1.050E+07 | 6.234E+06 |
| 602 | 303.138 | 3.688 | -1 | 6.038E+05 | 1.024E+06 | 8.396E+05 | 6.442E+05 | 4.875E+05 | 3.398E+05 | 4.541E+05 | 4.943E+06 | 9.873E+06 | 0.000E+00 | 5.808E+03 | 0.000E+00 | 0.000E+00 | 0.000E+00 | 0.000E+00 |
| 605 | 347.222 | 7.583 | -1 | 3.629E+06 | 5.260E+06 | 2.562E+06 | 2.821E+06 | 4.223E+06 | 2.530E+06 | 2.954E+06 | 2.440E+06 | 4.456E+06 | 2.810E+06 | 4.289E+06 | 2.142E+06 | 4.756E+06 | 3.814E+06 | 5.047E+06 |
| 606 | 438.143 | 7.111 | -1 | 1.635E+04 | 0.000E+00 | 0.000E+00 | 4.437E+06 | 5.890E+05 | 4.257E+05 | 2.095E+05 | 2.344E+06 | 3.782E+06 | 0.000E+00 | 0.000E+00 | 0.000E+00 | 0.000E+00 | 0.000E+00 | 0.000E+00 |
| 607 | 851.398 | 7.250 | -1 | 5.520E+04 | 5.962E+05 | 4.242E+04 | 2.640E+05 | 4.052E+05 | 3.275E+05 | 1.077E+05 | 1.655E+06 | 2.110E+05 | 6.950E+05 | 7.659E+05 | 1.074E+05 | 2.826E+04 | 2.624E+05 | 9.517E+05 |
| 609 | 830.614 | 3.188 | -1 | 9.975E+05 | 3.681E+06 | 2.726E+06 | 1.725E+06 | 1.405E+06 | 6.468E+05 | 9.174E+05 | 2.987E+06 | 2.266E+06 | 1.380E+06 | 1.332E+06 | 1.109E+06 | 1.080E+06 | 3.930E+06 | 7.372E+06 |
| 612 | 863.046 | 3.487 | -1 | 3.657E+07 | 1.701E+07 | 1.761E+07 | 3.697E+04 | 4.488E+04 | 3.380E+04 | 6.906E+04 | 1.971E+07 | 1.173E+05 | 8.128E+05 | 9.407E+05 | 3.758E+04 | 1.080E+07 | 5.764E+05 | 4.047E+04 |
| 613 | 619.272 | 4.460 | -1 | 8.761E+05 | 4.909E+05 | 3.747E+05 | 5.767E+05 | 9.372E+05 | 6.580E+05 | 4.842E+05 | 1.650E+06 | 4.839E+05 | 2.253E+06 | 2.449E+06 | 8.368E+05 | 2.509E+06 | 1.426E+06 | 3.547E+06 |
| 617 | 454.172 | 7.812 | -1 | 2.034E+06 | 1.331E+06 | 1.131E+06 | 3.810E+05 | 7.633E+05 | 4.067E+05 | 5.106E+05 | 1.760E+06 | 2.595E+05 | 8.679E+05 | 6.532E+05 | 8.745E+05 | 1.321E+06 | 7.288E+05 | 1.465E+06 |
| 621 | 815.644 | 3.349 | -1 | 0.000E+00 | 7.550E+04 | 4.148E+04 | 4.590E+04 | 5.579E+04 | 4.551E+04 | 1.594E+05 | 1.122E+06 | 1.196E+05 | 8.323E+04 | 2.053E+04 | 6.743E+06 | 1.718E+06 | 1.403E+06 | 4.224E+05 |
| 626 | 331.209 | 4.774 | -1 | 2.821E+06 | 3.418E+06 | 3.205E+06 | 2.304E+06 | 1.936E+06 | 1.839E+06 | 1.834E+06 | 2.228E+06 | 1.179E+06 | 2.103E+06 | 2.867E+06 | 2.485E+06 | 2.356E+06 | 2.256E+06 | 1.801E+06 |
| 628 | 103.054 | 0.515 | -1 | 1.644E+07 | 6.879E+06 | 2.798E+06 | 6.817E+05 | 1.961E+06 | 1.085E+06 | 2.742E+05 | 4.039E+05 | 1.215E+06 | 6.697E+05 | 1.211E+07 | 7.081E+06 | 3.120E+07 | 1.057E+06 | 3.283E+05 |
| 641 | 158.154 | 6.250 | -1 | 3.948E+07 | 8.666E+07 | 4.956E+06 | 8.307E+07 | 4.385E+07 | 4.289E+07 | 3.390E+07 | 4.149E+07 | 4.614E+07 | 8.269E+07 | 4.437E+07 | 1.939E+07 | 2.298E+07 | 4.691E+07 | 2.533E+07 |
| 642 | 165.114 | 12.714 | -1 | 4.356E+07 | 3.218E+07 | 1.501E+07 | 3.680E+07 | 1.414E+07 | 1.761E+07 | 1.594E+07 | 1.745E+07 | 4.057E+07 | 3.581E+07 | 1.695E+07 | 4.574E+05 | 1.516E+07 | 2.043E+07 | 1.565E+07 |
| 648 | 425.137 | 6.913 | -1 | 2.606E+07 | 2.787E+05 | 1.642E+07 | 1.590E+06 | 1.224E+06 | 7.435E+05 | 3.039E+05 | 5.286E+05 | 4.612E+05 | 7.512E+05 | 3.218E+05 | 4.779E+05 | 2.871E+07 | 7.624E+05 | 4.483E+05 |
| 651 | 267.157 | 4.332 | -1 | 6.216E+06 | 1.603E+07 | 1.227E+07 | 3.648E+06 | 1.189E+06 | 4.382E+05 | 5.723E+05 | 1.455E+06 | 7.626E+06 | 4.033E+06 | 3.832E+06 | 2.666E+06 | 3.151E+06 | 1.084E+06 | 4.472E+06 |
| 655 | 290.232 | 4.304 | -1 | 3.660E+06 | 8.759E+06 | 7.411E+06 | 2.074E+06 | 5.794E+05 | 5.489E+05 | 4.668E+05 | 5.145E+05 | 5.014E+06 | 9.331E+05 | 2.402E+06 | 8.040E+05 | 7.322E+05 | 6.376E+05 | 1.399E+06 |
| 660 | 391.205 | 10.042 | -1 | 6.699E+05 | 8.497E+05 | 9.398E+05 | 1.910E+06 | 2.348E+06 | 5.611E+05 | 1.480E+05 | 3.417E+05 | 2.125E+06 | 1.106E+07 | 3.664E+06 | 4.121E+06 | 1.610E+06 | 9.220E+05 | 5.298E+05 |
| 673 | 870.956 | 4.058 | -1 | 5.159E+04 | 2.414E+05 | 4.139E+05 | 0.000E+00 | 4.476E+04 | 2.715E+04 | 3.872E+05 | 0.000E+00 | 4.087E+05 | 4.154E+05 | 2.151E+05 | 0.000E+00 | 1.023E+05 | 0.000E+00 | 1.296E+06 |
| 682 | 425.215 | 9.512 | -1 | 1.064E+06 | 1.126E+06 | 7.432E+05 | 9.830E+05 | 8.314E+05 | 1.045E+06 | 9.053E+05 | 9.392E+05 | 1.260E+06 | 8.283E+05 | 9.315E+05 | 1.216E+06 | 1.126E+06 | 2.025E+06 | 1.682E+06 |
| 684 | 612.425 | 0.524 | -1 | 9.200E+06 | 3.026E+06 | 7.003E+06 | 1.787E+05 | 3.467E+05 | 4.511E+05 | 3.723E+05 | 2.233E+05 | 1.313E+05 | 3.403E+06 | 4.251E+06 | 3.093E+06 | 1.037E+07 | 3.125E+04 | 7.431E+04 |
| 686 | 231.081 | 0.527 | -1 | 5.388E+05 | 6.468E+06 | 7.452E+06 | 6.842E+03 | 0.000E+00 | 5.838E+03 | 0.000E+00 | 1.180E+05 | 0.000E+00 | 8.372E+06 | 8.156E+06 | 4.885E+07 | 1.025E+06 | 3.280E+03 | 7.084E+03 |
| 688 | 726.463 | 12.425 | -1 | 7.078E+05 | 1.541E+06 | 1.805E+06 | 1.190E+06 | 9.656E+05 | 8.617E+05 | 8.487E+05 | 5.434E+05 | 1.342E+06 | 2.975E+06 | 3.413E+06 | 4.481E+06 | 3.539E+06 | 6.180E+06 | 2.362E+06 |
| 690 | 403.233 | 9.514 | -1 | 2.816E+05 | 5.808E+05 | 2.921E+05 | 3.340E+05 | 3.921E+05 | 4.431E+05 | 4.496E+05 | 3.565E+05 | 4.107E+05 | 3.508E+05 | 2.841E+05 | 4.706E+05 | 3.410E+05 | 8.775E+05 | 4.495E+05 |
| 694 | 111.020 | 12.539 | -1 | 2.186E+05 | 3.222E+05 | 2.763E+05 | 2.151E+05 | 2.159E+05 | 1.656E+05 | 2.807E+05 | 2.154E+05 | 6.181E+04 | 3.378E+05 | 2.000E+05 | 1.850E+05 | 1.374E+05 | 1.083E+05 | 1.578E+05 |
| 695 | 351.102 | 8.964 | -1 | 4.925E+05 | 2.120E+06 | 1.523E+06 | 2.848E+05 | 4.124E+05 | 4.928E+05 | 4.706E+05 | 4.435E+05 | 5.667E+05 | 4.949E+06 | 5.332E+05 | 8.130E+05 | 9.050E+05 | 4.532E+05 | 3.988E+05 |
| 700 | 164.051 | 4.799 | -1 | 0.000E+00 | 0.000E+00 | 0.000E+00 | 3.483E+04 | 6.008E+04 | 2.316E+05 | 0.000E+00 | 0.000E+00 | 5.790E+04 | 2.715E+07 | 1.884E+07 | 3.408E+07 | 1.457E+06 | 1.807E+07 | 2.011E+07 |
| 702 | 111.039 | 2.302 | -1 | 2.521E+06 | 2.821E+06 | 7.162E+05 | 1.677E+06 | 1.423E+06 | 4.311E+06 | 2.472E+05 | 3.282E+06 | 4.627E+05 | 1.025E+06 | 4.093E+06 | 4.374E+06 | 4.176E+06 | 1.712E+06 | 8.992E+06 |
| 705 | 271.031 | 5.195 | -1 | 4.433E+03 | 0.000E+00 | 0.000E+00 | 0.000E+00 | 0.000E+00 | 0.000E+00 | 0.000E+00 | 0.000E+00 | 0.000E+00 | 5.814E+06 | 2.067E+06 | 9.063E+06 | 2.176E+03 | 3.155E+03 | 6.338E+03 |
| 706 | 249.050 | 5.192 | -1 | 3.924E+03 | 0.000E+00 | 5.098E+03 | 0.000E+00 | 0.000E+00 | 0.000E+00 | 0.000E+00 | 0.000E+00 | 0.000E+00 | 5.765E+06 | 1.997E+06 | 8.944E+06 | 4.745E+03 | 0.000E+00 | 2.407E+03 |
| 707 | 279.060 | 6.671 | -1 | 0.000E+00 | 0.000E+00 | 3.618E+03 | 0.000E+00 | 0.000E+00 | 0.000E+00 | 0.000E+00 | 0.000E+00 | 0.000E+00 | 1.281E+07 | 7.360E+06 | 9.174E+06 | 0.000E+00 | 2.205E+04 | 3.139E+04 |
| 708 | 162.035 | 6.813 | -1 | 0.000E+00 | 0.000E+00 | 0.000E+00 | 0.000E+00 | 0.000E+00 | 0.000E+00 | 0.000E+00 | 0.000E+00 | 0.000E+00 | 2.050E+07 | 9.025E+06 | 9.098E+06 | 3.662E+04 | 1.495E+05 | 2.768E+04 |
| 709 | 177.083 | 2.375 | -1 | 2.955E+05 | 8.478E+05 | 1.641E+05 | 6.524E+04 | 1.944E+04 | 1.533E+04 | 6.704E+05 | 2.432E+05 | 1.967E+05 | 1.675E+05 | 8.249E+06 | 5.898E+07 | 4.271E+05 | 4.486E+07 | 6.066E+07 |
| 711 | 164.051 | 2.397 | -1 | 7.591E+05 | 4.429E+05 | 2.096E+05 | 3.635E+04 | 2.285E+04 | 2.540E+04 | 1.662E+05 | 2.677E+05 | 1.791E+05 | 4.652E+05 | 8.653E+06 | 6.027E+07 | 4.937E+06 | 4.607E+07 | 6.287E+07 |
| 712 | 245.070 | 2.402 | -1 | 2.963E+05 | 3.896E+05 | 1.607E+05 | 1.041E+05 | 1.743E+04 | 3.737E+03 | 1.023E+05 | 4.833E+05 | 2.051E+05 | 6.129E+05 | 6.705E+06 | 5.210E+07 | 5.839E+06 | 3.603E+07 | 5.276E+07 |
| 716 | 111.039 | 2.354 | -1 | 2.521E+06 | 4.002E+06 | 3.082E+06 | 1.677E+06 | 1.950E+06 | 2.611E+06 | 2.472E+05 | 4.161E+06 | 8.475E+05 | 1.025E+06 | 4.093E+06 | 2.596E+06 | 1.251E+06 | 1.270E+07 | 6.675E+05 |
| 720 | 495.251 | 4.182 | -1 | 0.000E+00 | 0.000E+00 | 0.000E+00 | 0.000E+00 | 0.000E+00 | 0.000E+00 | 0.000E+00 | 0.000E+00 | 0.000E+00 | 1.557E+06 | 4.418E+06 | 3.337E+06 | 2.406E+06 | 2.167E+06 | 1.379E+06 |
| 723 | 162.073 | 2.320 | -1 | 2.608E+04 | 1.420E+04 | 9.237E+03 | 1.054E+04 | 6.647E+03 | 1.064E+04 | 1.620E+04 | 3.154E+04 | 4.677E+03 | 1.769E+05 | 2.318E+05 | 2.317E+07 | 9.060E+05 | 1.666E+06 | 2.358E+07 |
| 725 | 1347.881 | 12.426 | -1 | 5.313E+04 | 1.566E+05 | 4.632E+05 | 3.872E+04 | 1.097E+05 | 1.087E+05 | 1.110E+05 | 1.195E+05 | 1.310E+05 | 1.884E+06 | 2.444E+06 | 3.916E+06 | 2.448E+06 | 8.147E+06 | 1.018E+06 |
| 733 | 297.083 | 6.817 | -1 | 0.000E+00 | 0.000E+00 | 9.956E+03 | 0.000E+00 | 0.000E+00 | 0.000E+00 | 0.000E+00 | 0.000E+00 | 0.000E+00 | 1.947E+06 | 1.127E+06 | 2.714E+06 | 4.869E+06 | 1.316E+07 | 1.347E+07 |
| 734 | 471.252 | 7.681 | -1 | 4.399E+04 | 5.364E+05 | 3.860E+05 | 1.422E+05 | 1.547E+05 | 1.131E+05 | 3.139E+05 | 7.795E+04 | 1.404E+06 | 2.054E+06 | 4.730E+05 | 1.283E+06 | 6.046E+05 | 9.158E+05 | 8.263E+05 |
| 735 | 597.678 | 0.494 | -1 | 1.648E+03 | 5.369E+05 | 3.495E+05 | 0.000E+00 | 0.000E+00 | 0.000E+00 | 1.230E+05 | 1.679E+05 | 0.000E+00 | 3.241E+06 | 1.040E+07 | 1.427E+06 | 5.647E+06 | 1.944E+07 | 8.118E+06 |
| 737 | 701.411 | 12.427 | -1 | 8.097E+05 | 1.269E+06 | 1.689E+06 | 5.596E+05 | 4.873E+05 | 1.148E+06 | 7.156E+05 | 1.003E+06 | 8.196E+05 | 2.602E+06 | 3.000E+06 | 4.083E+06 | 3.199E+06 | 5.205E+06 | 2.218E+06 |
| 738 | 253.602 | 7.678 | -1 | 2.682E+05 | 8.512E+05 | 1.040E+06 | 5.740E+05 | 4.930E+05 | 4.038E+05 | 7.269E+05 | 2.651E+05 | 1.609E+06 | 2.117E+06 | 9.006E+05 | 1.263E+06 | 1.101E+06 | 1.523E+06 | 9.994E+05 |
| 739 | 749.548 | 12.237 | -1 | 0.000E+00 | 0.000E+00 | 6.060E+03 | 2.778E+04 | 0.000E+00 | 1.026E+04 | 8.563E+03 | 0.000E+00 | 3.641E+04 | 3.140E+05 | 3.185E+04 | 6.047E+06 | 7.226E+06 | 1.881E+04 | 1.889E+07 |
| 742 | 287.650 | 8.328 | -1 | 1.025E+05 | 3.277E+05 | 4.641E+05 | 1.866E+05 | 3.238E+05 | 1.955E+05 | 2.525E+05 | 2.173E+05 | 1.253E+06 | 2.107E+06 | 7.678E+05 | 1.087E+06 | 7.967E+05 | 9.697E+05 | 8.767E+05 |
| 744 | 247.129 | 0.492 | -1 | 1.353E+04 | 8.946E+03 | 1.266E+04 | 2.332E+03 | 0.000E+00 | 2.631E+03 | 4.611E+04 | 2.287E+04 | 1.496E+03 | 7.271E+04 | 7.062E+04 | 9.579E+05 | 7.681E+05 | 2.962E+07 | 1.092E+07 |
| 748 | 654.456 | 10.203 | -1 | 0.000E+00 | 8.304E+03 | 1.295E+04 | 0.000E+00 | 1.344E+04 | 0.000E+00 | 0.000E+00 | 0.000E+00 | 6.766E+03 | 1.574E+06 | 4.544E+05 | 9.106E+05 | 2.656E+06 | 3.020E+06 | 6.769E+05 |
| 752 | 286.520 | 6.796 | -1 | 0.000E+00 | 0.000E+00 | 0.000E+00 | 0.000E+00 | 0.000E+00 | 0.000E+00 | 0.000E+00 | 0.000E+00 | 0.000E+00 | 4.176E+06 | 1.113E+06 | 7.142E+05 | 0.000E+00 | 0.000E+00 | 0.000E+00 |
| 753 | 335.125 | 8.963 | -1 | 5.276E+05 | 2.340E+06 | 2.076E+06 | 3.441E+05 | 5.507E+05 | 6.126E+05 | 6.338E+05 | 3.872E+05 | 6.478E+05 | 6.761E+06 | 7.131E+05 | 9.409E+05 | 1.078E+06 | 2.260E+05 | 3.528E+05 |
| 754 | 533.934 | 3.609 | -1 | 0.000E+00 | 0.000E+00 | 0.000E+00 | 0.000E+00 | 0.000E+00 | 0.000E+00 | 0.000E+00 | 0.000E+00 | 0.000E+00 | 7.249E+05 | 3.905E+06 | 2.488E+06 | 2.542E+06 | 1.092E+06 | 6.697E+05 |
| 762 | 533.282 | 9.675 | -1 | 7.134E+05 | 2.669E+06 | 8.702E+05 | 5.417E+05 | 2.347E+05 | 2.741E+05 | 3.401E+05 | 4.416E+05 | 4.661E+05 | 1.309E+05 | 5.052E+05 | 4.415E+05 | 4.169E+05 | 3.189E+05 | 2.255E+05 |
| 763 | 1194.349 | 0.496 | -1 | 2.599E+03 | 3.597E+04 | 8.176E+04 | 0.000E+00 | 0.000E+00 | 0.000E+00 | 7.292E+04 | 2.900E+04 | 0.000E+00 | 4.689E+05 | 4.479E+06 | 2.239E+05 | 1.889E+06 | 8.886E+06 | 1.907E+06 |

| Feature identifiers | | | | XIC | | | | | | | | | | | | | | |
|---|---|---|---|---|---|---|---|---|---|---|---|---|---|---|---|---|---|---|
| ClusterIdx | m/z | RT [min] | Subnetwork | 4TUB34_I | 4TUB34_II | 4TUB34_III | 4TUB81_I | 4TUB81_II | 4TUB81_III | 4TUB93_I | 4TUB93_II | 4TUB93_III | 5TUB23_I | 5TUB23_II | 5TUB23_III | 5TUB48_I | 5TUB48_II | 5TUB48_III |
| 764 | 616.651 | 0.503 | -1 | 2.393E+04 | 2.550E+04 | 3.666E+03 | 0.000E+00 | 0.000E+00 | 0.000E+00 | 0.000E+00 | 0.000E+00 | 0.000E+00 | 5.567E+05 | 2.571E+06 | 4.168E+04 | 1.033E+06 | 5.732E+06 | 2.370E+06 |
| 765 | 223.064 | 6.361 | -1 | 4.426E+06 | 9.415E+06 | 5.624E+06 | 8.577E+05 | 5.088E+05 | 2.423E+05 | 5.485E+05 | 7.213E+05 | 3.767E+06 | 1.329E+06 | 2.450E+06 | 9.649E+05 | 9.341E+05 | 6.082E+05 | 1.228E+06 |
| 769 | 498.173 | 7.669 | -1 | 6.765E+05 | 2.440E+05 | 1.241E+06 | 3.322E+05 | 5.921E+05 | 4.395E+05 | 4.435E+05 | 4.308E+05 | 7.255E+05 | 8.036E+05 | 9.751E+05 | 4.685E+05 | 1.535E+06 | 1.298E+06 | 5.989E+05 |
| 771 | 143.107 | 1.088 | -1 | 8.383E+06 | 4.287E+07 | 2.980E+07 | 2.211E+05 | 1.893E+05 | 6.367E+04 | 1.456E+05 | 5.411E+05 | 1.986E+07 | 9.368E+05 | 1.651E+06 | 1.888E+05 | 1.299E+05 | 9.392E+04 | 6.762E+06 |
| 786 | 819.476 | 8.754 | -1 | 2.699E+05 | 6.809E+05 | 2.897E+05 | 1.127E+05 | 1.024E+05 | 1.098E+05 | 1.356E+05 | 8.402E+04 | 4.241E+06 | 5.615E+04 | 1.087E+05 | 2.041E+05 | 3.301E+05 | 1.155E+05 | 1.529E+05 |
| 794 | 811.308 | 6.909 | -1 | 1.699E+06 | 0.000E+00 | 7.316E+05 | 0.000E+00 | 0.000E+00 | 0.000E+00 | 0.000E+00 | 0.000E+00 | 0.000E+00 | 0.000E+00 | 0.000E+00 | 0.000E+00 | 2.778E+06 | 0.000E+00 | 0.000E+00 |
| 797 | 818.299 | 3.374 | -1 | 3.613E+05 | 6.575E+05 | 3.087E+05 | 3.392E+03 | 0.000E+00 | 0.000E+00 | 3.100E+04 | 0.000E+00 | 4.328E+04 | 9.472E+05 | 3.985E+04 | 3.146E+05 | 2.427E+05 | 5.908E+05 | 3.895E+06 |
| 806 | 618.984 | 3.690 | -1 | 2.585E+05 | 8.455E+05 | 3.773E+05 | 0.000E+00 | 0.000E+00 | 1.433E+04 | 3.740E+04 | 0.000E+00 | 8.251E+04 | 1.597E+06 | 4.852E+04 | 5.666E+04 | 2.283E+05 | 1.590E+05 | 5.409E+06 |
| 808 | 832.302 | 3.693 | -1 | 1.432E+05 | 6.435E+05 | 3.414E+05 | 0.000E+00 | 0.000E+00 | 0.000E+00 | 2.849E+04 | 0.000E+00 | 3.955E+04 | 1.263E+06 | 4.588E+04 | 9.659E+03 | 1.766E+05 | 8.193E+04 | 3.436E+06 |
| 814 | 513.189 | 7.230 | -1 | 9.617E+05 | 1.002E+06 | 1.503E+06 | 4.400E+05 | 6.383E+05 | 5.030E+05 | 9.863E+05 | 2.375E+05 | 5.547E+05 | 1.099E+06 | 1.129E+06 | 7.357E+05 | 6.834E+05 | 1.156E+06 | 5.352E+05 |
| 815 | 432.241 | 7.420 | -1 | 1.464E+04 | 3.598E+04 | 7.120E+05 | 4.552E+04 | 8.828E+04 | 2.171E+04 | 1.737E+04 | 1.887E+04 | 1.243E+04 | 3.180E+04 | 9.519E+04 | 3.116E+04 | 2.784E+06 | 1.157E+04 | 5.194E+04 |
| 818 | 123.079 | 0.380 | -1 | 1.025E+07 | 8.021E+06 | 1.715E+07 | 8.392E+06 | 1.742E+07 | 2.714E+07 | 1.690E+07 | 3.082E+07 | 1.254E+07 | 2.553E+07 | 2.363E+07 | 1.191E+07 | 1.661E+07 | 1.994E+07 | 2.393E+07 |
| 822 | 376.087 | 6.788 | -1 | 0.000E+00 | 0.000E+00 | 0.000E+00 | 0.000E+00 | 0.000E+00 | 0.000E+00 | 0.000E+00 | 0.000E+00 | 0.000E+00 | 0.000E+00 | 0.000E+00 | 0.000E+00 | 1.708E+05 | 3.245E+06 | 2.210E+06 |
| 827 | 326.112 | 5.859 | -1 | 4.081E+03 | 8.751E+03 | 2.813E+04 | 0.000E+00 | 7.587E+03 | 0.000E+00 | 0.000E+00 | 1.361E+04 | 4.283E+04 | 6.002E+05 | 4.160E+05 | 6.922E+05 | 7.455E+05 | 3.968E+06 | 2.797E+06 |
| 828 | 319.064 | 6.836 | -1 | 0.000E+00 | 0.000E+00 | 0.000E+00 | 6.163E+04 | 8.230E+04 | 0.000E+00 | 0.000E+00 | 0.000E+00 | 2.907E+04 | 4.369E+04 | 3.237E+04 | 1.551E+05 | 6.696E+05 | 1.790E+06 | 1.752E+06 |
| 831 | 1295.589 | 10.022 | -1 | 8.242E+05 | 0.000E+00 | 7.004E+04 | 1.081E+04 | 9.774E+03 | 0.000E+00 | 0.000E+00 | 7.552E+05 | 0.000E+00 | 0.000E+00 | 2.368E+04 | 7.246E+05 | 2.510E+06 | 4.397E+05 | 8.489E+05 |
| 834 | 628.471 | 3.460 | -1 | 1.013E+05 | 5.036E+05 | 6.272E+05 | 1.764E+05 | 2.705E+05 | 2.701E+05 | 7.562E+05 | 7.932E+04 | 4.215E+05 | 3.627E+05 | 3.687E+04 | 1.999E+05 | 7.370E+05 | 2.493E+05 | 6.882E+05 |
| 835 | 850.609 | 3.462 | -1 | 3.154E+05 | 7.314E+05 | 4.568E+05 | 2.610E+05 | 3.233E+05 | 1.516E+05 | 7.033E+05 | 1.273E+05 | 8.616E+05 | 6.091E+05 | 3.422E+05 | 2.995E+05 | 7.470E+05 | 7.250E+05 | 8.943E+05 |
| 839 | 372.717 | 10.091 | -1 | 0.000E+00 | 0.000E+00 | 0.000E+00 | 7.820E+04 | 4.124E+04 | 0.000E+00 | 0.000E+00 | 0.000E+00 | 0.000E+00 | 0.000E+00 | 3.250E+04 | 0.000E+00 | 6.200E+06 | 7.225E+05 | 1.466E+04 |
| 846 | 681.132 | 5.759 | -1 | 0.000E+00 | 0.000E+00 | 0.000E+00 | 7.519E+03 | 8.656E+04 | 2.361E+05 | 1.166E+04 | 9.619E+04 | 9.061E+03 | 0.000E+00 | 0.000E+00 | 8.311E+03 | 1.479E+06 | 1.964E+06 | 1.678E+06 |
| 856 | 1194.817 | 11.488 | -1 | 0.000E+00 | 1.211E+04 | 2.332E+04 | 9.355E+03 | 0.000E+00 | 0.000E+00 | 0.000E+00 | 0.000E+00 | 0.000E+00 | 7.096E+04 | 6.008E+04 | 3.175E+05 | 2.407E+05 | 1.007E+06 | 1.272E+05 |
| 857 | 804.296 | 3.067 | -1 | 1.109E+05 | 1.536E+05 | 2.317E+04 | 0.000E+00 | 0.000E+00 | 0.000E+00 | 0.000E+00 | 0.000E+00 | 4.625E+03 | 2.773E+05 | 0.000E+00 | 1.873E+05 | 1.527E+05 | 1.319E+06 | 3.503E+06 |
| 863 | 353.267 | 9.157 | -1 | 4.282E+05 | 2.417E+05 | 3.659E+05 | 3.534E+05 | 2.017E+05 | 3.750E+05 | 3.521E+05 | 4.040E+05 | 6.004E+04 | 1.844E+05 | 3.495E+05 | 3.842E+05 | 3.491E+05 | 8.482E+05 | 3.593E+05 |

| Feature identifiers | | | | | | | | | | |
|---|---|---|---|---|---|---|---|---|---|---|
| ClusterIdx | m/z | RT [min] | Subnetwork | 5TUB83_I | 5TUB83_II | 5TUB83_III | TUB00_I | TUB00_II | TUB00_III | blank |
| 491 | 205.195 | 9.007 | -1 | 0.000E+00 | 2.331E+03 | 5.940E+03 | 3.542E+04 | 1.502E+04 | 6.823E+03 | 0.000E+00 |
| 24 | 399.252 | 8.753 | 58 | 7.936E+07 | 4.705E+07 | 4.789E+07 | 4.698E+07 | 3.999E+07 | 3.998E+07 | 7.304E+07 |
| 44 | 421.232 | 8.753 | -1 | 1.628E+07 | 1.410E+07 | 1.371E+07 | 1.501E+07 | 1.341E+07 | 1.338E+07 | 1.895E+07 |
| 130 | 242.100 | 0.564 | 54 | 4.082E+03 | 3.498E+04 | 0.000E+00 | 1.495E+07 | 1.085E+07 | 1.561E+07 | 0.000E+00 |
| 479 | 242.101 | 0.569 | 54 | 4.082E+03 | 3.498E+04 | 0.000E+00 | 1.131E+07 | 8.268E+06 | 9.263E+06 | 0.000E+00 |
| 478 | 204.087 | 0.529 | 51 | 5.970E+04 | 3.139E+04 | 1.946E+04 | 3.244E+04 | 1.630E+04 | 5.981E+04 | 1.701E+04 |
| 200 | 311.256 | 7.678 | 39 | 2.227E+07 | 1.198E+07 | 1.633E+07 | 9.888E+05 | 6.488E+05 | 7.492E+05 | 0.000E+00 |
| 461 | 311.256 | 7.795 | 39 | 5.428E+07 | 8.565E+06 | 3.725E+07 | 1.767E+05 | 2.418E+05 | 2.632E+05 | 0.000E+00 |
| 536 | 339.290 | 8.561 | 39 | 2.197E+07 | 5.895E+06 | 1.814E+07 | 4.257E+05 | 1.051E+05 | 1.628E+05 | 0.000E+00 |
| 574 | 339.289 | 8.380 | 39 | 2.897E+06 | 1.419E+06 | 2.679E+06 | 4.913E+05 | 4.658E+05 | 4.882E+05 | 0.000E+00 |
| 666 | 188.071 | 1.307 | 27 | 7.673E+03 | 0.000E+00 | 0.000E+00 | 2.370E+08 | 1.642E+08 | 2.067E+08 | 0.000E+00 |
| 6 | 166.086 | 0.517 | 41 | 3.426E+08 | 3.518E+08 | 1.352E+08 | 1.341E+09 | 1.195E+09 | 1.708E+09 | 8.715E+03 |
| 441 | 223.090 | 6.761 | -1 | 1.846E+06 | 3.325E+06 | 1.212E+06 | 1.075E+06 | 2.847E+06 | 1.704E+06 | 1.226E+05 |
| 378 | 391.285 | 10.644 | -1 | 6.468E+05 | 7.229E+05 | 4.104E+05 | 5.226E+05 | 5.191E+05 | 5.654E+05 | 1.624E+06 |
| 12 | 245.096 | 3.603 | 62 | 2.078E+08 | 1.434E+08 | 1.313E+08 | 3.035E+07 | 2.410E+07 | 2.779E+07 | 1.025E+04 |
| 408 | 489.184 | 3.610 | 62 | 3.567E+06 | 1.120E+06 | 1.061E+06 | 0.000E+00 | 0.000E+00 | 0.000E+00 | 0.000E+00 |
| 467 | 250.118 | 7.417 | -1 | 5.811E+05 | 2.330E+05 | 5.050E+05 | 9.358E+05 | 5.942E+05 | 5.142E+05 | 2.030E+07 |
| 462 | 482.321 | 9.064 | 4 | 5.010E+06 | 8.710E+05 | 1.600E+06 | 6.045E+04 | 8.513E+04 | 7.802E+04 | 0.000E+00 |
| 832 | 577.516 | 12.699 | 39 | 2.119E+05 | 2.473E+06 | 3.084E+06 | 0.000E+00 | 0.000E+00 | 0.000E+00 | 0.000E+00 |
| 17 | 454.293 | 8.329 | 4 | 2.250E+08 | 9.321E+07 | 1.381E+08 | 2.588E+07 | 2.149E+07 | 2.133E+07 | 1.456E+05 |
| 89 | 454.293 | 8.190 | 4 | 1.785E+07 | 9.482E+06 | 1.145E+07 | 2.124E+06 | 1.639E+06 | 1.437E+06 | 1.708E+04 |
| 211 | 476.267 | 8.328 | 4 | 1.366E+07 | 6.280E+06 | 8.886E+06 | 1.581E+06 | 1.474E+06 | 1.396E+06 | 0.000E+00 |
| 743 | 907.578 | 8.331 | -1 | 2.597E+06 | 5.594E+05 | 9.667E+05 | 0.000E+00 | 1.161E+04 | 0.000E+00 | 0.000E+00 |
| 138 | 338.342 | 10.765 | 39 | 8.137E+06 | 3.725E+06 | 4.187E+06 | 4.956E+06 | 4.383E+06 | 4.695E+06 | 2.315E+07 |
| 456 | 338.342 | 9.820 | 39 | 1.947E+06 | 1.960E+06 | 1.646E+06 | 1.510E+06 | 1.427E+06 | 1.560E+06 | 4.806E+06 |
| 767 | 688.493 | 12.701 | 4 | 6.690E+04 | 2.137E+07 | 6.845E+06 | 0.000E+00 | 0.000E+00 | 0.000E+00 | 0.000E+00 |
| 30 | 480.309 | 8.382 | 4 | 1.179E+08 | 3.314E+07 | 9.094E+07 | 1.402E+07 | 1.065E+07 | 1.182E+07 | 3.652E+03 |
| 36 | 480.309 | 8.512 | 4 | 4.445E+07 | 1.920E+07 | 2.995E+07 | 4.777E+06 | 3.655E+06 | 3.208E+06 | 0.000E+00 |
| 342 | 377.146 | 3.769 | -1 | 1.043E+06 | 7.210E+05 | 8.310E+05 | 9.537E+05 | 3.110E+05 | 9.452E+05 | 1.002E+04 |
| 473 | 220.118 | 0.532 | 61 | 1.361E+05 | 4.491E+05 | 1.672E+04 | 3.414E+06 | 1.772E+07 | 4.351E+07 | 0.000E+00 |
| 372 | 628.196 | 10.499 | 40 | 1.175E+06 | 1.120E+06 | 1.150E+06 | 2.387E+06 | 2.238E+06 | 1.697E+06 | 7.961E+07 |
| 3 | 432.238 | 7.526 | 24 | 1.052E+09 | 2.448E+08 | 4.101E+08 | 2.071E+08 | 2.148E+08 | 2.628E+08 | 9.714E+06 |
| 161 | 432.238 | 7.249 | 24 | 7.179E+06 | 9.364E+05 | 5.366E+06 | 1.416E+06 | 1.218E+06 | 1.285E+06 | 1.272E+04 |
| 442 | 432.238 | 7.812 | 24 | 3.629E+06 | 1.022E+06 | 1.334E+06 | 4.110E+05 | 6.889E+05 | 4.522E+05 | 1.434E+05 |
| 2 | 415.212 | 7.526 | 24 | 1.203E+09 | 2.373E+08 | 4.167E+08 | 1.909E+08 | 2.003E+08 | 2.468E+08 | 4.071E+06 |
| 78 | 415.212 | 7.248 | 24 | 1.403E+07 | 2.176E+06 | 1.006E+07 | 2.753E+06 | 2.309E+06 | 2.220E+06 | 2.788E+04 |
| 373 | 415.212 | 7.817 | 24 | 5.664E+06 | 1.992E+06 | 2.201E+06 | 1.269E+06 | 9.623E+05 | 9.703E+05 | 1.548E+05 |
| 40 | 327.078 | 8.430 | -1 | 1.044E+06 | 8.540E+05 | 6.374E+05 | 6.463E+05 | 6.546E+05 | 8.003E+05 | 2.917E+06 |
| 435 | 313.143 | 8.965 | -1 | 1.290E+06 | 9.045E+05 | 6.842E+05 | 1.267E+06 | 8.584E+05 | 1.654E+06 | 1.998E+06 |
| 80 | 223.089 | 2.607 | 27 | 6.007E+05 | 1.614E+04 | 2.423E+03 | 1.292E+04 | 1.056E+04 | 1.206E+04 | 1.036E+04 |
| 697 | 223.089 | 2.398 | 27 | 8.994E+05 | 7.103E+06 | 7.278E+06 | 1.254E+04 | 3.811E+04 | 4.368E+04 | 1.851E+04 |
| 798 | 223.089 | 2.491 | 27 | 1.178E+07 | 1.735E+05 | 4.615E+05 | 1.047E+04 | 3.388E+04 | 4.368E+04 | 7.080E+03 |
| 665 | 205.097 | 1.310 | 27 | 4.024E+03 | 0.000E+00 | 0.000E+00 | 2.954E+08 | 2.057E+08 | 2.553E+08 | 0.000E+00 |
| 154 | 457.112 | 3.497 | -1 | 1.322E+07 | 1.123E+07 | 9.988E+06 | 8.747E+06 | 7.355E+06 | 8.512E+06 | 0.000E+00 |
| 29 | 220.118 | 0.526 | 61 | 1.361E+05 | 4.491E+05 | 1.672E+04 | 4.579E+07 | 2.897E+07 | 4.506E+07 | 0.000E+00 |
| 681 | 331.165 | 0.498 | 41 | 2.890E+04 | 7.340E+03 | 0.000E+00 | 3.034E+06 | 2.525E+06 | 3.910E+06 | 0.000E+00 |
| 192 | 343.296 | 6.715 | 23 | 3.433E+06 | 2.987E+06 | 5.180E+06 | 3.814E+06 | 3.826E+06 | 7.181E+06 | 5.287E+06 |
| 589 | 481.400 | 5.241 | 3 | 1.777E+06 | 1.993E+04 | 1.587E+04 | 2.004E+04 | 8.438E+03 | 1.037E+05 | 0.000E+00 |
| 603 | 810.134 | 3.065 | 20 | 7.073E+07 | 2.449E+07 | 6.070E+07 | 9.709E+03 | 3.853E+05 | 1.828E+05 | 0.000E+00 |
| 521 | 716.523 | 12.701 | 4 | 1.317E+07 | 2.714E+06 | 1.680E+07 | 0.000E+00 | 0.000E+00 | 0.000E+00 | 0.000E+00 |
| 572 | 716.525 | 10.404 | 4 | 2.614E+07 | 6.623E+06 | 2.348E+05 | 0.000E+00 | 0.000E+00 | 0.000E+00 | 4.363E+06 |
| 654 | 716.524 | 10.304 | 4 | 6.770E+07 | 7.650E+07 | 2.348E+05 | 0.000E+00 | 0.000E+00 | 0.000E+00 | 1.720E+06 |
| 852 | 716.524 | 10.151 | 4 | 3.897E+08 | 1.558E+07 | 2.699E+05 | 0.000E+00 | 8.918E+03 | 0.000E+00 | 7.740E+04 |
| 524 | 426.262 | 7.583 | 4 | 1.239E+07 | 6.768E+06 | 7.163E+06 | 9.896E+05 | 7.997E+05 | 7.260E+05 | 0.000E+00 |
| 552 | 426.262 | 7.443 | 4 | 2.018E+07 | 1.162E+07 | 1.562E+07 | 3.294E+06 | 2.861E+06 | 3.009E+06 | 0.000E+00 |
| 507 | 718.539 | 12.704 | 4 | 1.744E+07 | 2.299E+07 | 1.241E+08 | 0.000E+00 | 0.000E+00 | 0.000E+00 | 0.000E+00 |
| 523 | 718.539 | 10.302 | 4 | 1.945E+07 | 4.143E+06 | 1.303E+07 | 0.000E+00 | 0.000E+00 | 0.000E+00 | 0.000E+00 |
| 632 | 718.538 | 10.143 | 4 | 2.759E+07 | 9.699E+07 | 1.566E+07 | 0.000E+00 | 0.000E+00 | 0.000E+00 | 0.000E+00 |
| 823 | 771.514 | 12.696 | 4 | 3.304E+05 | 8.376E+06 | 6.821E+06 | 0.000E+00 | 0.000E+00 | 0.000E+00 | 0.000E+00 |
| 346 | 502.290 | 8.380 | 4 | 4.916E+06 | 2.473E+06 | 4.593E+06 | 8.869E+05 | 7.012E+05 | 7.978E+05 | 0.000E+00 |
| 388 | 502.291 | 8.511 | 4 | 1.724E+06 | 1.144E+06 | 1.294E+06 | 2.864E+05 | 2.297E+05 | 1.289E+05 | 0.000E+00 |
| 476 | 979.349 | 0.522 | 91 | 0.000E+00 | 0.000E+00 | 0.000E+00 | 0.000E+00 | 0.000E+00 | 0.000E+00 | 0.000E+00 |
| 488 | 995.318 | 0.510 | 91 | 0.000E+00 | 0.000E+00 | 0.000E+00 | 0.000E+00 | 0.000E+00 | 0.000E+00 | 0.000E+00 |
| 52 | 256.264 | 10.439 | 88 | 1.161E+07 | 7.707E+06 | 7.796E+06 | 1.632E+07 | 1.528E+07 | 1.582E+07 | 7.131E+04 |
| 427 | 284.295 | 10.783 | 88 | 3.386E+06 | 1.194E+06 | 1.702E+06 | 1.363E+06 | 1.145E+06 | 1.346E+06 | 0.000E+00 |
| 62 | 291.096 | 6.457 | 87 | 0.000E+00 | 0.000E+00 | 0.000E+00 | 0.000E+00 | 0.000E+00 | 0.000E+00 | 0.000E+00 |
| 153 | 291.096 | 6.559 | 87 | 0.000E+00 | 0.000E+00 | 0.000E+00 | 0.000E+00 | 0.000E+00 | 0.000E+00 | 0.000E+00 |
| 374 | 748.356 | 4.106 | 84 | 0.000E+00 | 0.000E+00 | 0.000E+00 | 0.000E+00 | 0.000E+00 | 0.000E+00 | 0.000E+00 |
| 384 | 783.875 | 4.128 | 84 | 0.000E+00 | 0.000E+00 | 0.000E+00 | 0.000E+00 | 0.000E+00 | 0.000E+00 | 0.000E+00 |
| 284 | 243.196 | 7.929 | 80 | 0.000E+00 | 0.000E+00 | 6.100E+03 | 1.761E+04 | 3.366E+04 | 1.600E+04 | 0.000E+00 |

| Feature identifiers | | | | | | | | | | |
|---|---|---|---|---|---|---|---|---|---|---|
| ClusterIdx | m/z | RT [min] | Subnetwork | 5TUB83_I | 5TUB83_II | 5TUB83_III | TUB00_I | TUB00_II | TUB00_III | blank |
| 415 | 225.185 | 7.934 | 80 | 5.811E+03 | 1.028E+04 | 6.091E+03 | 3.627E+04 | 1.180E+04 | 2.086E+04 | 7.434E+03 |
| 667 | 176.074 | 0.736 | 79 | 8.425E+04 | 3.903E+05 | 8.460E+04 | 1.151E+08 | 8.287E+07 | 7.681E+07 | 4.430E+03 |
| 668 | 218.121 | 4.865 | 79 | 4.528E+04 | 2.452E+05 | 2.705E+04 | 1.720E+07 | 1.466E+07 | 1.367E+07 | 0.000E+00 |
| 670 | 246.152 | 6.015 | 79 | 2.392E+04 | 3.746E+04 | 8.207E+03 | 2.000E+07 | 1.976E+07 | 2.020E+07 | 0.000E+00 |
| 674 | 232.137 | 5.471 | 79 | 0.000E+00 | 5.802E+03 | 0.000E+00 | 4.367E+06 | 4.785E+06 | 4.519E+06 | 0.000E+00 |
| 675 | 204.105 | 4.140 | 79 | 0.000E+00 | 6.044E+04 | 0.000E+00 | 7.482E+06 | 8.148E+06 | 8.669E+06 | 0.000E+00 |
| 772 | 551.192 | 3.315 | 78 | 0.000E+00 | 4.211E+05 | 0.000E+00 | 0.000E+00 | 3.422E+05 | 3.149E+05 | 0.000E+00 |
| 774 | 615.714 | 3.354 | 78 | 0.000E+00 | 6.341E+05 | 0.000E+00 | 0.000E+00 | 8.486E+04 | 9.294E+04 | 0.000E+00 |
| 776 | 486.670 | 3.240 | 78 | 0.000E+00 | 1.363E+05 | 0.000E+00 | 0.000E+00 | 1.137E+05 | 1.186E+05 | 0.000E+00 |
| 777 | 680.235 | 3.406 | 78 | 0.000E+00 | 6.404E+05 | 0.000E+00 | 0.000E+00 | 2.726E+04 | 6.511E+04 | 0.000E+00 |
| 780 | 357.628 | 3.065 | 78 | 0.000E+00 | 1.704E+05 | 0.000E+00 | 1.543E+04 | 1.112E+05 | 1.829E+05 | 0.000E+00 |
| 46 | 226.158 | 9.291 | 77 | 8.327E+05 | 2.603E+05 | 1.349E+05 | 7.907E+05 | 4.377E+05 | 5.840E+05 | 2.542E+05 |
| 201 | 282.221 | 10.073 | 77 | 3.257E+05 | 6.867E+04 | 1.353E+05 | 1.699E+06 | 6.075E+05 | 7.471E+05 | 1.700E+05 |
| 634 | 282.221 | 10.084 | 77 | 2.746E+05 | 6.867E+04 | 1.353E+05 | 1.096E+06 | 6.075E+05 | 5.470E+05 | 1.700E+05 |
| 150 | 818.299 | 3.185 | 75 | 2.877E+07 | 2.330E+07 | 1.375E+07 | 3.202E+06 | 7.604E+05 | 1.709E+06 | 0.000E+00 |
| 323 | 1226.943 | 3.181 | 75 | 7.468E+06 | 5.780E+06 | 6.865E+05 | 6.091E+05 | 4.815E+05 | 3.825E+05 | 0.000E+00 |
| 843 | 1226.943 | 3.183 | 75 | 5.937E+06 | 4.922E+06 | 2.817E+06 | 7.132E+05 | 2.787E+05 | 2.614E+05 | 0.000E+00 |
| 186 | 135.079 | 7.525 | 74 | 2.181E+07 | 3.765E+06 | 7.078E+06 | 2.925E+06 | 2.905E+06 | 3.702E+06 | 4.673E+04 |
| 319 | 135.079 | 4.463 | 74 | 9.914E+06 | 1.829E+06 | 3.898E+06 | 8.065E+05 | 9.642E+05 | 9.628E+05 | 2.544E+04 |
| 180 | 529.790 | 4.168 | 73 | 0.000E+00 | 0.000E+00 | 0.000E+00 | 0.000E+00 | 0.000E+00 | 0.000E+00 | 0.000E+00 |
| 264 | 465.270 | 4.196 | 73 | 0.000E+00 | 0.000E+00 | 0.000E+00 | 0.000E+00 | 0.000E+00 | 0.000E+00 | 0.000E+00 |
| 48 | 431.565 | 4.119 | 72 | 0.000E+00 | 0.000E+00 | 0.000E+00 | 1.025E+04 | 5.458E+04 | 0.000E+00 | 0.000E+00 |
| 170 | 417.549 | 3.504 | 72 | 1.374E+07 | 1.468E+07 | 1.705E+07 | 1.702E+07 | 5.403E+07 | 5.464E+07 | 0.000E+00 |
| 314 | 396.544 | 3.209 | 72 | 0.000E+00 | 2.957E+06 | 7.839E+03 | 8.158E+07 | 6.711E+07 | 5.274E+07 | 0.000E+00 |
| 117 | 832.302 | 3.461 | 70 | 3.890E+07 | 3.419E+07 | 2.160E+07 | 4.785E+06 | 2.007E+06 | 2.842E+06 | 0.000E+00 |
| 371 | 1247.949 | 3.461 | 70 | 9.959E+06 | 9.673E+06 | 6.910E+06 | 1.544E+06 | 1.142E+06 | 9.851E+05 | 0.000E+00 |
| 512 | 1247.454 | 3.462 | 70 | 1.012E+07 | 8.004E+06 | 6.431E+06 | 5.712E+05 | 4.591E+05 | 7.542E+05 | 0.000E+00 |
| 7 | 271.106 | 8.022 | 69 | 1.978E+04 | 1.524E+04 | 2.235E+04 | 1.329E+04 | 1.357E+04 | 0.000E+00 | 9.999E+03 |
| 45 | 390.122 | 6.709 | 69 | 0.000E+00 | 0.000E+00 | 0.000E+00 | 0.000E+00 | 0.000E+00 | 0.000E+00 | 0.000E+00 |
| 112 | 388.106 | 7.488 | 69 | 0.000E+00 | 0.000E+00 | 0.000E+00 | 0.000E+00 | 0.000E+00 | 0.000E+00 | 0.000E+00 |
| 209 | 414.122 | 6.104 | 69 | 0.000E+00 | 0.000E+00 | 0.000E+00 | 0.000E+00 | 0.000E+00 | 0.000E+00 | 0.000E+00 |
| 663 | 414.122 | 6.419 | 69 | 2.427E+06 | 1.280E+06 | 2.692E+05 | 2.352E+05 | 0.000E+00 | 0.000E+00 | 0.000E+00 |
| 468 | 432.238 | 6.913 | 68 | 1.435E+05 | 2.117E+05 | 1.581E+05 | 1.185E+06 | 7.575E+05 | 8.522E+05 | 0.000E+00 |
| 792 | 404.208 | 6.914 | 68 | 6.177E+04 | 1.052E+05 | 5.710E+04 | 7.447E+05 | 4.518E+05 | 6.468E+05 | 0.000E+00 |
| 793 | 387.181 | 6.913 | 68 | 5.373E+04 | 7.533E+04 | 6.646E+04 | 7.321E+05 | 4.702E+05 | 6.067E+05 | 0.000E+00 |
| 671 | 331.207 | 5.234 | 67 | 1.045E+04 | 4.481E+04 | 1.015E+04 | 1.202E+07 | 1.069E+07 | 1.598E+07 | 0.000E+00 |
| 795 | 317.195 | 4.828 | 67 | 6.163E+04 | 6.533E+04 | 1.644E+05 | 7.016E+05 | 7.194E+05 | 1.072E+06 | 0.000E+00 |
| 84 | 280.188 | 5.797 | 65 | 5.465E+03 | 0.000E+00 | 0.000E+00 | 1.248E+04 | 9.219E+03 | 1.440E+04 | 0.000E+00 |
| 158 | 308.222 | 6.723 | 65 | 0.000E+00 | 0.000E+00 | 0.000E+00 | 2.468E+04 | 0.000E+00 | 4.872E+03 | 0.000E+00 |
| 263 | 693.878 | 3.983 | 64 | 0.000E+00 | 0.000E+00 | 0.000E+00 | 0.000E+00 | 0.000E+00 | 0.000E+00 | 0.000E+00 |
| 392 | 665.367 | 3.986 | 64 | 0.000E+00 | 0.000E+00 | 0.000E+00 | 0.000E+00 | 0.000E+00 | 0.000E+00 | 0.000E+00 |
| 703 | 204.534 | 6.795 | 63 | 1.520E+05 | 0.000E+00 | 2.991E+05 | 0.000E+00 | 0.000E+00 | 0.000E+00 | 0.000E+00 |
| 704 | 193.026 | 6.808 | 63 | 2.615E+05 | 7.857E+03 | 3.433E+05 | 0.000E+00 | 0.000E+00 | 0.000E+00 | 0.000E+00 |
| 88 | 227.084 | 3.606 | 62 | 2.310E+07 | 1.602E+07 | 1.461E+07 | 2.968E+06 | 2.549E+06 | 2.666E+06 | 0.000E+00 |
| 215 | 290.153 | 3.607 | 62 | 9.643E+06 | 5.831E+06 | 5.195E+06 | 5.126E+05 | 3.941E+05 | 4.095E+05 | 0.000E+00 |
| 4 | 182.985 | 12.752 | 60 | 7.080E+08 | 6.936E+08 | 7.014E+08 | 6.827E+08 | 1.932E+08 | 6.588E+08 | 8.100E+08 |
| 5 | 154.990 | 12.767 | 60 | 6.891E+08 | 6.495E+08 | 6.848E+08 | 8.701E+08 | 8.905E+08 | 8.899E+08 | 8.451E+08 |
| 19 | 138.996 | 12.748 | 60 | 1.032E+08 | 9.468E+07 | 1.011E+08 | 1.328E+08 | 1.367E+08 | 1.373E+08 | 1.384E+08 |
| 23 | 205.017 | 12.741 | 60 | 4.318E+07 | 4.017E+07 | 4.155E+07 | 5.777E+07 | 5.690E+07 | 5.732E+07 | 3.072E+07 |
| 49 | 196.017 | 12.766 | 60 | 2.584E+07 | 2.407E+07 | 2.715E+07 | 3.976E+07 | 4.424E+07 | 4.384E+07 | 4.360E+07 |
| 63 | 224.012 | 12.748 | 60 | 1.941E+07 | 1.824E+07 | 1.896E+07 | 3.480E+07 | 3.605E+07 | 3.288E+07 | 1.959E+07 |
| 75 | 195.994 | 12.775 | 60 | 3.178E+07 | 2.849E+07 | 2.950E+07 | 3.297E+07 | 3.346E+07 | 3.497E+07 | 3.898E+07 |
| 20 | 131.534 | 12.743 | 59 | 5.569E+07 | 5.736E+07 | 5.639E+07 | 8.483E+07 | 8.470E+07 | 8.484E+07 | 6.287E+07 |
| 409 | 131.534 | 13.009 | 59 | 3.211E+05 | 9.489E+05 | 7.700E+05 | 4.135E+05 | 1.103E+06 | 1.172E+06 | 1.050E+06 |
| 465 | 131.534 | 13.069 | 59 | 2.331E+05 | 2.466E+05 | 6.338E+05 | 9.243E+05 | 6.386E+05 | 3.151E+05 | 1.490E+06 |
| 696 | 131.534 | 4.507 | 59 | 9.572E+04 | 1.481E+05 | 1.433E+05 | 2.360E+05 | 1.791E+05 | 3.146E+05 | 5.186E+05 |
| 266 | 444.309 | 8.752 | 58 | 3.520E+06 | 2.637E+06 | 3.158E+06 | 2.504E+06 | 2.505E+06 | 2.036E+06 | 4.700E+06 |
| 490 | 350.102 | 0.493 | 57 | 1.844E+06 | 4.134E+07 | 2.837E+03 | 6.170E+06 | 3.809E+06 | 5.408E+06 | 0.000E+00 |
| 495 | 364.118 | 0.634 | 57 | 0.000E+00 | 1.908E+04 | 0.000E+00 | 3.574E+05 | 3.304E+05 | 1.231E+05 | 0.000E+00 |
| 134 | 519.318 | 10.374 | 56 | 2.339E+05 | 1.605E+06 | 1.787E+06 | 2.263E+05 | 4.262E+06 | 8.620E+04 | 0.000E+00 |
| 658 | 519.309 | 10.382 | 56 | 2.083E+06 | 1.736E+06 | 1.898E+06 | 3.250E+06 | 4.318E+06 | 2.708E+06 | 0.000E+00 |
| 498 | 532.098 | 0.493 | 55 | 0.000E+00 | 0.000E+00 | 0.000E+00 | 0.000E+00 | 0.000E+00 | 0.000E+00 | 0.000E+00 |
| 501 | 532.099 | 0.497 | 55 | 0.000E+00 | 0.000E+00 | 0.000E+00 | 0.000E+00 | 0.000E+00 | 0.000E+00 | 0.000E+00 |
| 132 | 711.319 | 4.210 | 53 | 0.000E+00 | 0.000E+00 | 0.000E+00 | 0.000E+00 | 0.000E+00 | 0.000E+00 | 0.000E+00 |
| 282 | 624.289 | 4.196 | 53 | 0.000E+00 | 0.000E+00 | 0.000E+00 | 0.000E+00 | 0.000E+00 | 0.000E+00 | 0.000E+00 |
| 133 | 226.180 | 5.976 | 52 | 5.470E+06 | 6.561E+06 | 5.109E+06 | 6.176E+06 | 5.040E+06 | 7.024E+06 | 7.661E+06 |
| 542 | 384.349 | 10.283 | 52 | 2.809E+06 | 2.790E+06 | 2.110E+06 | 1.482E+06 | 1.371E+06 | 1.444E+06 | 4.630E+06 |
| 131 | 614.463 | 9.741 | 51 | 1.377E+06 | 6.628E+05 | 5.789E+05 | 1.586E+05 | 4.654E+04 | 1.109E+05 | 8.973E+05 |
| 243 | 388.269 | 6.414 | 51 | 7.853E+04 | 1.068E+05 | 1.697E+04 | 1.002E+04 | 1.121E+04 | 9.699E+03 | 1.571E+04 |
| 472 | 479.186 | 0.514 | 51 | 1.927E+04 | 1.685E+05 | 4.232E+04 | 1.769E+06 | 0.000E+00 | 0.000E+00 | 0.000E+00 |

| Feature identifiers | | | | | | | | | | |
|---|---|---|---|---|---|---|---|---|---|---|
| ClusterIdx | m/z | RT [min] | Subnetwork | 5TUB83_I | 5TUB83_II | 5TUB83_III | TUB00_I | TUB00_II | TUB00_III | blank |
| 475 | 957.367 | 0.530 | 51 | 0.000E+00 | 0.000E+00 | 0.000E+00 | 0.000E+00 | 0.000E+00 | 0.000E+00 | 0.000E+00 |
| 477 | 524.246 | 0.489 | 51 | 0.000E+00 | 5.430E+03 | 0.000E+00 | 0.000E+00 | 0.000E+00 | 0.000E+00 | 0.000E+00 |
| 480 | 268.076 | 0.497 | 51 | 0.000E+00 | 0.000E+00 | 0.000E+00 | 0.000E+00 | 0.000E+00 | 0.000E+00 | 0.000E+00 |
| 482 | 498.161 | 0.503 | 51 | 0.000E+00 | 0.000E+00 | 0.000E+00 | 2.306E+04 | 1.871E+03 | 0.000E+00 | 0.000E+00 |
| 483 | 644.267 | 0.510 | 51 | 0.000E+00 | 0.000E+00 | 0.000E+00 | 0.000E+00 | 0.000E+00 | 0.000E+00 | 0.000E+00 |
| 484 | 279.584 | 0.532 | 51 | 0.000E+00 | 0.000E+00 | 0.000E+00 | 0.000E+00 | 0.000E+00 | 0.000E+00 | 0.000E+00 |
| 487 | 276.062 | 0.490 | 51 | 0.000E+00 | 0.000E+00 | 0.000E+00 | 0.000E+00 | 0.000E+00 | 0.000E+00 | 0.000E+00 |
| 492 | 737.248 | 0.522 | 51 | 0.000E+00 | 0.000E+00 | 0.000E+00 | 0.000E+00 | 0.000E+00 | 0.000E+00 | 0.000E+00 |
| 493 | 506.147 | 0.488 | 51 | 0.000E+00 | 0.000E+00 | 0.000E+00 | 0.000E+00 | 0.000E+00 | 0.000E+00 | 0.000E+00 |
| 500 | 287.570 | 0.486 | 51 | 0.000E+00 | 0.000E+00 | 0.000E+00 | 0.000E+00 | 0.000E+00 | 0.000E+00 | 0.000E+00 |
| 687 | 441.187 | 0.508 | 51 | 1.460E+06 | 1.012E+05 | 1.505E+05 | 9.920E+05 | 2.859E+05 | 9.019E+05 | 0.000E+00 |
| 732 | 632.474 | 10.196 | 51 | 3.153E+06 | 9.709E+05 | 1.455E+06 | 0.000E+00 | 0.000E+00 | 7.474E+03 | 0.000E+00 |
| 802 | 860.514 | 9.355 | 51 | 3.750E+06 | 1.059E+06 | 1.581E+06 | 0.000E+00 | 0.000E+00 | 0.000E+00 | 0.000E+00 |
| 168 | 471.307 | 5.923 | 50 | 3.570E+06 | 3.623E+06 | 2.908E+06 | 2.232E+06 | 3.387E+06 | 4.243E+06 | 0.000E+00 |
| 616 | 473.286 | 4.748 | 50 | 4.312E+05 | 2.213E+06 | 5.039E+05 | 1.114E+05 | 1.347E+06 | 2.022E+06 | 0.000E+00 |
| 144 | 393.207 | 10.310 | 49 | 5.696E+06 | 6.367E+06 | 5.857E+06 | 5.574E+06 | 5.184E+06 | 5.238E+06 | 9.920E+06 |
| 176 | 409.183 | 10.451 | 49 | 4.595E+06 | 4.030E+06 | 4.425E+06 | 3.762E+06 | 4.129E+06 | 3.718E+06 | 8.719E+06 |
| 43 | 418.117 | 7.633 | 48 | 0.000E+00 | 0.000E+00 | 0.000E+00 | 0.000E+00 | 0.000E+00 | 0.000E+00 | 0.000E+00 |
| 146 | 551.149 | 6.292 | 48 | 0.000E+00 | 0.000E+00 | 0.000E+00 | 0.000E+00 | 0.000E+00 | 0.000E+00 | 0.000E+00 |
| 363 | 418.117 | 7.570 | 48 | 0.000E+00 | 0.000E+00 | 0.000E+00 | 0.000E+00 | 0.000E+00 | 0.000E+00 | 0.000E+00 |
| 119 | 343.130 | 3.690 | 47 | 3.548E+04 | 3.651E+04 | 1.010E+04 | 4.073E+04 | 4.267E+05 | 1.213E+06 | 0.000E+00 |
| 182 | 537.334 | 9.562 | 47 | 0.000E+00 | 0.000E+00 | 0.000E+00 | 7.085E+03 | 8.899E+03 | 3.830E+03 | 3.991E+04 |
| 231 | 509.303 | 8.961 | 47 | 0.000E+00 | 0.000E+00 | 0.000E+00 | 0.000E+00 | 0.000E+00 | 0.000E+00 | 0.000E+00 |
| 239 | 245.616 | 7.678 | 46 | 1.342E+07 | 7.341E+06 | 9.946E+06 | 6.299E+05 | 2.482E+05 | 4.909E+05 | 0.000E+00 |
| 393 | 266.129 | 7.678 | 46 | 7.255E+06 | 4.237E+06 | 5.701E+06 | 1.871E+05 | 1.963E+05 | 1.801E+05 | 0.000E+00 |
| 429 | 266.129 | 7.807 | 46 | 1.714E+06 | 1.027E+06 | 9.597E+05 | 4.942E+04 | 2.414E+04 | 3.685E+04 | 0.000E+00 |
| 506 | 734.471 | 12.708 | 45 | 1.023E+07 | 3.305E+06 | 2.097E+07 | 0.000E+00 | 0.000E+00 | 0.000E+00 | 0.000E+00 |
| 544 | 734.471 | 10.338 | 45 | 7.246E+06 | 1.023E+07 | 5.820E+05 | 0.000E+00 | 0.000E+00 | 0.000E+00 | 0.000E+00 |
| 550 | 765.459 | 12.703 | 45 | 4.394E+05 | 7.087E+06 | 1.932E+06 | 0.000E+00 | 0.000E+00 | 0.000E+00 | 0.000E+00 |
| 646 | 734.471 | 10.261 | 45 | 1.344E+08 | 6.401E+07 | 4.933E+05 | 0.000E+00 | 0.000E+00 | 0.000E+00 | 0.000E+00 |
| 652 | 762.502 | 12.706 | 45 | 2.092E+06 | 2.766E+06 | 7.771E+06 | 0.000E+00 | 0.000E+00 | 0.000E+00 | 0.000E+00 |
| 370 | 514.353 | 10.382 | 43 | 1.372E+06 | 7.676E+05 | 6.720E+05 | 9.752E+05 | 1.064E+06 | 7.337E+05 | 0.000E+00 |
| 740 | 744.593 | 12.218 | 43 | 4.589E+06 | 0.000E+00 | 2.234E+07 | 0.000E+00 | 0.000E+00 | 0.000E+00 | 4.028E+04 |
| 94 | 247.596 | 7.524 | 42 | 2.388E+07 | 5.585E+06 | 1.068E+07 | 4.570E+06 | 4.992E+06 | 5.060E+06 | 0.000E+00 |
| 110 | 256.601 | 7.527 | 42 | 2.170E+07 | 5.433E+06 | 8.799E+06 | 3.413E+06 | 3.439E+06 | 5.043E+06 | 0.000E+00 |
| 233 | 268.115 | 7.525 | 42 | 5.580E+06 | 1.369E+06 | 3.942E+06 | 2.366E+06 | 2.220E+06 | 2.919E+06 | 1.129E+04 |
| 283 | 554.177 | 10.161 | 40 | 1.611E+06 | 2.366E+06 | 1.867E+06 | 2.499E+06 | 2.405E+06 | 2.161E+06 | 4.732E+07 |
| 545 | 684.203 | 10.983 | 40 | 5.163E+06 | 2.786E+05 | 1.265E+05 | 2.924E+05 | 9.823E+04 | 1.206E+05 | 1.871E+04 |
| 630 | 702.214 | 10.746 | 40 | 5.102E+05 | 5.389E+05 | 5.098E+05 | 1.266E+06 | 1.012E+06 | 9.409E+05 | 5.634E+07 |
| 693 | 463.131 | 9.688 | 40 | 5.570E+05 | 2.810E+05 | 6.474E+05 | 6.173E+05 | 6.446E+05 | 6.249E+05 | 4.657E+06 |
| 821 | 776.232 | 11.006 | 40 | 3.132E+05 | 2.980E+05 | 1.713E+05 | 6.580E+05 | 6.168E+05 | 5.868E+05 | 4.303E+07 |
| 858 | 610.185 | 12.140 | 40 | 1.147E+06 | 1.079E+06 | 8.867E+05 | 7.377E+05 | 5.053E+05 | 8.637E+05 | 1.396E+08 |
| 143 | 228.196 | 7.148 | 39 | 4.835E+06 | 9.450E+06 | 3.478E+06 | 1.836E+06 | 1.412E+06 | 1.871E+06 | 1.589E+04 |
| 230 | 228.196 | 6.462 | 39 | 1.424E+07 | 2.496E+07 | 8.955E+06 | 8.760E+06 | 6.653E+06 | 9.114E+06 | 1.667E+07 |
| 824 | 549.485 | 10.160 | 39 | 4.550E+06 | 7.065E+07 | 1.241E+07 | 0.000E+00 | 0.000E+00 | 0.000E+00 | 0.000E+00 |
| 826 | 549.486 | 12.704 | 39 | 1.221E+06 | 7.671E+06 | 6.545E+06 | 0.000E+00 | 0.000E+00 | 0.000E+00 | 0.000E+00 |
| 840 | 549.486 | 9.853 | 39 | 5.064E+06 | 1.388E+06 | 1.940E+08 | 0.000E+00 | 0.000E+00 | 0.000E+00 | 0.000E+00 |
| 850 | 549.485 | 10.361 | 39 | 2.365E+06 | 4.623E+06 | 6.915E+06 | 0.000E+00 | 0.000E+00 | 0.000E+00 | 0.000E+00 |
| 159 | 480.238 | 3.674 | 37 | 0.000E+00 | 0.000E+00 | 0.000E+00 | 0.000E+00 | 0.000E+00 | 0.000E+00 | 0.000E+00 |
| 332 | 959.468 | 3.674 | 37 | 0.000E+00 | 0.000E+00 | 0.000E+00 | 0.000E+00 | 0.000E+00 | 0.000E+00 | 0.000E+00 |
| 301 | 467.250 | 5.033 | 36 | 0.000E+00 | 0.000E+00 | 0.000E+00 | 0.000E+00 | 0.000E+00 | 0.000E+00 | 0.000E+00 |
| 333 | 933.494 | 5.044 | 36 | 0.000E+00 | 0.000E+00 | 0.000E+00 | 0.000E+00 | 0.000E+00 | 0.000E+00 | 0.000E+00 |
| 420 | 516.785 | 5.231 | 36 | 0.000E+00 | 0.000E+00 | 0.000E+00 | 0.000E+00 | 0.000E+00 | 0.000E+00 | 0.000E+00 |
| 68 | 217.105 | 4.009 | 35 | 3.752E+07 | 4.220E+07 | 1.350E+07 | 1.051E+07 | 4.854E+07 | 5.403E+07 | 1.436E+05 |
| 330 | 234.134 | 4.002 | 35 | 6.917E+06 | 7.741E+06 | 2.121E+06 | 1.970E+06 | 9.539E+06 | 9.971E+06 | 1.006E+04 |
| 431 | 681.651 | 10.701 | 33 | 8.928E+05 | 7.454E+05 | 8.192E+05 | 1.322E+06 | 1.235E+06 | 1.122E+06 | 3.643E+06 |
| 434 | 399.395 | 7.402 | 33 | 9.842E+05 | 9.559E+05 | 9.321E+05 | 1.109E+06 | 1.096E+06 | 1.043E+06 | 4.001E+06 |
| 171 | 278.185 | 3.992 | 32 | 1.294E+07 | 1.029E+07 | 9.939E+06 | 7.268E+06 | 8.507E+06 | 1.008E+07 | 0.000E+00 |
| 437 | 280.186 | 4.202 | 32 | 4.315E+06 | 4.811E+06 | 4.147E+06 | 1.912E+06 | 3.194E+06 | 3.233E+06 | 0.000E+00 |
| 560 | 271.180 | 3.637 | 32 | 3.998E+06 | 2.373E+06 | 2.955E+06 | 1.133E+06 | 1.988E+06 | 4.377E+06 | 0.000E+00 |
| 35 | 255.159 | 6.531 | 31 | 3.573E+07 | 2.263E+07 | 2.110E+07 | 1.685E+07 | 1.886E+07 | 1.973E+07 | 1.111E+07 |
| 99 | 272.185 | 6.531 | 31 | 1.243E+07 | 7.500E+06 | 6.757E+06 | 5.855E+06 | 6.359E+06 | 6.549E+06 | 7.678E+06 |
| 466 | 269.175 | 7.631 | 31 | 8.728E+06 | 3.517E+06 | 3.789E+05 | 3.181E+05 | 2.268E+06 | 9.266E+05 | 8.584E+03 |
| 28 | 297.133 | 7.153 | 30 | 6.289E+06 | 2.225E+07 | 1.698E+07 | 1.599E+07 | 1.852E+07 | 2.062E+07 | 0.000E+00 |
| 38 | 431.207 | 7.153 | 30 | 4.146E+06 | 1.393E+07 | 1.068E+07 | 1.117E+07 | 1.292E+07 | 1.369E+07 | 0.000E+00 |
| 56 | 447.202 | 6.602 | 30 | 2.547E+06 | 1.176E+07 | 1.007E+07 | 8.560E+06 | 1.254E+07 | 1.190E+07 | 0.000E+00 |
| 91 | 448.233 | 7.152 | 30 | 2.599E+06 | 6.543E+06 | 5.002E+06 | 5.053E+06 | 6.302E+06 | 6.664E+06 | 0.000E+00 |
| 102 | 429.191 | 6.409 | 30 | 1.877E+06 | 8.657E+06 | 7.688E+06 | 5.289E+06 | 7.151E+06 | 7.297E+06 | 0.000E+00 |
| 163 | 297.133 | 7.367 | 30 | 3.440E+06 | 4.948E+06 | 3.365E+06 | 2.992E+06 | 4.057E+06 | 4.738E+06 | 0.000E+00 |
| 202 | 448.233 | 7.470 | 30 | 2.388E+06 | 3.907E+06 | 3.245E+06 | 2.069E+06 | 3.134E+06 | 3.488E+06 | 0.000E+00 |

| Feature identifiers | | | | | | | | | | |
|---|---|---|---|---|---|---|---|---|---|---|
| **ClusterIdx** | **m/z** | **RT [min]** | **Subnetwork** | **5TUB83_I** | **5TUB83_II** | **5TUB83_III** | **TUB00_I** | **TUB00_II** | **TUB00_III** | **blank** |
| 208 | 431.207 | 7.472 | 30 | 1.263E+06 | 5.340E+06 | 2.564E+06 | 1.828E+06 | 3.400E+06 | 3.557E+06 | 0.000E+00 |
| 213 | 447.202 | 7.074 | 30 | 9.207E+05 | 2.600E+06 | 2.489E+06 | 1.394E+06 | 1.693E+06 | 1.716E+06 | 0.000E+00 |
| 219 | 447.202 | 6.409 | 30 | 9.462E+05 | 3.801E+06 | 3.341E+06 | 2.510E+06 | 3.360E+06 | 3.401E+06 | 0.000E+00 |
| 235 | 297.133 | 6.600 | 30 | 6.100E+05 | 2.959E+06 | 2.634E+06 | 1.727E+06 | 2.681E+06 | 2.833E+06 | 0.000E+00 |
| 245 | 469.184 | 6.601 | 30 | 6.144E+05 | 2.882E+06 | 2.526E+06 | 1.780E+06 | 2.880E+06 | 2.540E+06 | 0.000E+00 |
| 249 | 429.191 | 6.602 | 30 | 6.186E+05 | 2.823E+06 | 2.422E+06 | 1.937E+06 | 2.859E+06 | 2.844E+06 | 0.000E+00 |
| 251 | 297.133 | 6.410 | 30 | 6.298E+05 | 3.219E+06 | 2.655E+06 | 1.799E+06 | 2.434E+06 | 2.487E+06 | 0.000E+00 |
| 262 | 447.202 | 6.882 | 30 | 8.730E+06 | 5.992E+06 | 4.650E+06 | 4.254E+06 | 5.046E+06 | 5.140E+06 | 0.000E+00 |
| 411 | 527.717 | 3.679 | 29 | 0.000E+00 | 0.000E+00 | 0.000E+00 | 9.976E+04 | 2.119E+03 | 0.000E+00 | 0.000E+00 |
| 444 | 791.074 | 3.678 | 29 | 0.000E+00 | 0.000E+00 | 0.000E+00 | 6.173E+04 | 0.000E+00 | 0.000E+00 | 0.000E+00 |
| 355 | 500.380 | 6.151 | 28 | 3.208E+06 | 3.155E+06 | 3.323E+06 | 4.259E+06 | 3.927E+06 | 3.817E+06 | 2.137E+07 |
| 391 | 442.338 | 5.719 | 28 | 2.659E+06 | 2.619E+06 | 2.709E+06 | 2.973E+06 | 2.718E+06 | 2.932E+06 | 2.226E+07 |
| 394 | 309.227 | 4.773 | 28 | 2.971E+06 | 2.745E+06 | 2.575E+06 | 3.117E+06 | 2.825E+06 | 4.085E+06 | 1.320E+07 |
| 446 | 384.296 | 5.267 | 28 | 2.214E+06 | 2.071E+06 | 1.999E+06 | 2.619E+06 | 2.119E+06 | 3.104E+06 | 1.952E+07 |
| 458 | 674.506 | 7.346 | 28 | 3.811E+05 | 8.297E+05 | 1.056E+06 | 2.413E+06 | 2.140E+06 | 2.056E+06 | 5.791E+06 |
| 631 | 790.590 | 8.085 | 28 | 1.345E+06 | 9.232E+05 | 1.291E+06 | 2.280E+06 | 2.213E+06 | 2.148E+06 | 6.480E+06 |
| 635 | 549.416 | 9.554 | 28 | 4.760E+05 | 3.744E+05 | 5.257E+05 | 6.818E+05 | 7.922E+05 | 1.139E+06 | 1.503E+07 |
| 860 | 694.522 | 10.428 | 28 | 1.429E+06 | 5.014E+05 | 5.777E+05 | 1.889E+05 | 3.851E+05 | 2.234E+05 | 1.676E+07 |
| 357 | 206.062 | 2.609 | 27 | 1.099E+06 | 2.449E+03 | 0.000E+00 | 0.000E+00 | 0.000E+00 | 0.000E+00 | 0.000E+00 |
| 440 | 237.106 | 4.092 | 27 | 4.512E+04 | 2.559E+04 | 4.365E+04 | 2.199E+04 | 6.594E+04 | 3.905E+04 | 9.337E+03 |
| 699 | 206.061 | 2.395 | 27 | 2.469E+05 | 4.243E+05 | 1.748E+05 | 0.000E+00 | 0.000E+00 | 0.000E+00 | 0.000E+00 |
| 799 | 206.062 | 2.458 | 27 | 7.934E+05 | 4.243E+05 | 1.218E+05 | 0.000E+00 | 1.764E+03 | 0.000E+00 | 0.000E+00 |
| 448 | 213.144 | 5.071 | 26 | 1.064E+06 | 1.031E+06 | 5.735E+05 | 2.727E+06 | 2.396E+06 | 2.147E+06 | 3.527E+07 |
| 692 | 381.331 | 9.796 | 26 | 2.395E+05 | 8.562E+05 | 7.044E+05 | 3.224E+06 | 1.245E+06 | 1.001E+06 | 9.007E+07 |
| 210 | 445.290 | 4.792 | 25 | 2.207E+06 | 3.031E+06 | 2.791E+06 | 3.112E+06 | 2.691E+06 | 2.847E+06 | 5.822E+06 |
| 227 | 467.300 | 4.849 | 25 | 2.322E+06 | 2.404E+06 | 2.210E+06 | 2.238E+06 | 2.398E+06 | 2.117E+06 | 5.869E+06 |
| 250 | 423.276 | 4.729 | 25 | 1.993E+06 | 2.258E+06 | 2.064E+06 | 2.525E+06 | 2.128E+06 | 2.280E+06 | 6.354E+06 |
| 275 | 401.261 | 4.663 | 25 | 1.654E+06 | 1.774E+06 | 1.704E+06 | 1.901E+06 | 1.879E+06 | 1.831E+06 | 6.557E+06 |
| 307 | 379.249 | 4.593 | 25 | 1.337E+06 | 1.372E+06 | 1.473E+06 | 1.836E+06 | 1.468E+06 | 1.622E+06 | 7.357E+06 |
| 353 | 489.316 | 4.904 | 25 | 1.219E+06 | 1.436E+06 | 1.241E+06 | 1.408E+06 | 1.342E+06 | 1.278E+06 | 5.437E+06 |
| 358 | 505.391 | 10.706 | 25 | 1.564E+06 | 1.272E+06 | 1.075E+06 | 1.301E+06 | 1.117E+06 | 1.430E+06 | 4.342E+06 |
| 402 | 527.403 | 10.694 | 25 | 1.296E+06 | 1.039E+06 | 9.715E+05 | 1.056E+06 | 1.051E+06 | 1.252E+06 | 3.156E+06 |
| 459 | 549.417 | 10.679 | 25 | 8.355E+05 | 6.805E+05 | 7.342E+05 | 8.515E+05 | 5.435E+05 | 9.833E+05 | 2.381E+06 |
| 535 | 463.345 | 9.103 | 25 | 3.579E+05 | 3.391E+05 | 3.184E+05 | 1.459E+05 | 2.444E+05 | 2.660E+05 | 4.257E+05 |
| 540 | 441.330 | 9.123 | 25 | 3.297E+05 | 2.230E+05 | 3.295E+05 | 2.501E+05 | 3.076E+05 | 2.025E+05 | 6.855E+05 |
| 638 | 529.383 | 9.048 | 25 | 3.615E+05 | 2.250E+05 | 1.662E+05 | 2.110E+05 | 4.355E+05 | 1.822E+05 | 1.801E+05 |
| 653 | 507.370 | 9.067 | 25 | 3.106E+05 | 2.444E+05 | 2.593E+05 | 2.162E+05 | 4.432E+05 | 1.868E+05 | 1.942E+05 |
| 662 | 511.328 | 4.955 | 25 | 5.946E+05 | 3.489E+05 | 5.250E+05 | 4.451E+05 | 6.966E+05 | 5.102E+05 | 4.930E+06 |
| 759 | 571.429 | 10.663 | 25 | 4.691E+05 | 4.200E+05 | 5.294E+05 | 4.165E+05 | 4.070E+05 | 7.886E+05 | 1.449E+06 |
| 859 | 533.341 | 5.004 | 25 | 3.188E+05 | 2.372E+05 | 2.313E+05 | 4.024E+05 | 3.384E+05 | 3.032E+05 | 4.887E+06 |
| 9 | 460.270 | 7.525 | 24 | 3.254E+08 | 1.468E+08 | 2.089E+08 | 1.142E+08 | 1.201E+08 | 1.262E+08 | 9.832E+06 |
| 25 | 846.443 | 7.526 | 24 | 2.306E+08 | 7.563E+06 | 3.202E+07 | 6.811E+06 | 6.134E+06 | 1.078E+07 | 0.000E+00 |
| 67 | 446.254 | 7.523 | 24 | 1.913E+07 | 9.370E+06 | 1.307E+07 | 6.528E+06 | 7.385E+06 | 9.382E+06 | 3.425E+04 |
| 81 | 460.270 | 7.249 | 24 | 1.464E+07 | 2.745E+06 | 1.138E+07 | 3.050E+06 | 2.666E+06 | 2.594E+06 | 9.004E+04 |
| 139 | 476.266 | 7.526 | 24 | 1.015E+07 | 4.161E+06 | 6.291E+06 | 5.430E+06 | 4.914E+06 | 4.850E+06 | 7.880E+04 |
| 261 | 397.201 | 7.526 | 24 | 9.127E+06 | 9.992E+05 | 3.298E+06 | 1.295E+06 | 1.281E+06 | 1.194E+06 | 0.000E+00 |
| 272 | 429.191 | 6.884 | 24 | 3.191E+07 | 7.639E+06 | 5.498E+06 | 4.715E+06 | 5.317E+06 | 6.334E+06 | 0.000E+00 |
| 316 | 531.275 | 7.525 | 24 | 2.770E+07 | 2.969E+05 | 1.608E+06 | 2.642E+05 | 3.754E+05 | 3.198E+05 | 0.000E+00 |
| 375 | 462.213 | 6.365 | 24 | 3.110E+07 | 1.277E+06 | 9.498E+05 | 7.031E+05 | 9.353E+05 | 1.434E+06 | 0.000E+00 |
| 418 | 445.186 | 6.364 | 24 | 2.614E+07 | 1.087E+06 | 8.096E+05 | 7.126E+05 | 7.970E+05 | 1.259E+06 | 0.000E+00 |
| 419 | 460.270 | 7.814 | 24 | 5.613E+06 | 1.830E+06 | 2.533E+06 | 5.226E+05 | 6.190E+05 | 6.372E+05 | 1.659E+05 |
| 447 | 450.204 | 8.853 | 24 | 2.298E+06 | 8.411E+05 | 7.889E+05 | 5.327E+05 | 6.714E+05 | 7.023E+05 | 0.000E+00 |
| 469 | 445.186 | 6.211 | 24 | 9.977E+06 | 4.596E+05 | 2.857E+05 | 2.571E+05 | 4.991E+05 | 5.798E+05 | 0.000E+00 |
| 578 | 448.233 | 6.302 | 24 | 3.715E+06 | 2.966E+05 | 1.688E+05 | 2.340E+05 | 1.705E+05 | 4.021E+05 | 0.000E+00 |
| 582 | 448.233 | 6.159 | 24 | 4.874E+06 | 4.067E+05 | 2.461E+05 | 2.447E+05 | 3.142E+05 | 4.490E+05 | 0.000E+00 |
| 636 | 462.213 | 6.211 | 24 | 6.499E+06 | 3.450E+05 | 1.177E+05 | 2.220E+05 | 2.610E+05 | 2.186E+05 | 0.000E+00 |
| 637 | 431.207 | 6.148 | 24 | 4.407E+06 | 3.748E+05 | 2.274E+05 | 1.636E+05 | 2.968E+05 | 2.757E+05 | 0.000E+00 |
| 770 | 474.249 | 6.864 | 24 | 1.000E+07 | 7.080E+05 | 3.441E+05 | 2.340E+05 | 3.220E+05 | 4.797E+05 | 0.000E+00 |
| 790 | 431.207 | 6.021 | 24 | 3.045E+06 | 2.280E+05 | 1.889E+05 | 1.375E+05 | 2.013E+05 | 3.042E+05 | 0.000E+00 |
| 221 | 371.326 | 7.517 | 23 | 1.561E+06 | 2.022E+06 | 2.702E+06 | 2.282E+06 | 2.568E+06 | 3.859E+06 | 4.749E+06 |
| 31 | 328.919 | 12.753 | 22 | 4.571E+07 | 4.142E+07 | 4.366E+07 | 8.157E+07 | 7.603E+07 | 7.295E+07 | 1.580E+08 |
| 39 | 312.943 | 12.757 | 22 | 3.140E+07 | 2.226E+07 | 3.427E+07 | 4.540E+07 | 1.156E+07 | 4.844E+07 | 6.384E+07 |
| 70 | 287.888 | 12.760 | 22 | 2.496E+07 | 2.118E+07 | 2.201E+07 | 4.072E+07 | 3.519E+07 | 3.553E+07 | 7.729E+07 |
| 92 | 313.078 | 6.460 | 22 | 0.000E+00 | 0.000E+00 | 0.000E+00 | 0.000E+00 | 0.000E+00 | 0.000E+00 | 0.000E+00 |
| 187 | 312.943 | 12.757 | 22 | 3.521E+07 | 3.345E+07 | 2.809E+07 | 4.540E+07 | 4.805E+07 | 4.661E+07 | 6.384E+07 |
| 196 | 313.078 | 6.532 | 22 | 0.000E+00 | 0.000E+00 | 0.000E+00 | 0.000E+00 | 0.000E+00 | 0.000E+00 | 0.000E+00 |
| 255 | 313.235 | 8.431 | 22 | 6.446E+06 | 3.052E+06 | 2.371E+06 | 1.552E+06 | 2.227E+06 | 2.365E+06 | 3.582E+06 |
| 558 | 313.235 | 8.075 | 22 | 6.021E+06 | 8.762E+06 | 4.052E+06 | 6.010E+04 | 8.592E+04 | 8.345E+04 | 1.640E+05 |
| 197 | 272.258 | 8.673 | 21 | 1.985E+05 | 1.292E+05 | 1.835E+05 | 1.058E+05 | 5.406E+04 | 1.501E+05 | 7.675E+04 |
| 223 | 300.290 | 9.476 | 21 | 6.882E+05 | 5.786E+05 | 6.162E+05 | 1.152E+05 | 2.191E+05 | 2.704E+05 | 1.283E+06 |

| Feature identifiers | | | | | | | | | | |
|---|---|---|---|---|---|---|---|---|---|---|
| ClusterIdx | m/z | RT [min] | Subnetwork | 5TUB83_I | 5TUB83_II | 5TUB83_III | TUB00_I | TUB00_II | TUB00_III | blank |
| 460 | 406.354 | 10.232 | 21 | 2.908E+06 | 6.533E+05 | 6.806E+05 | 2.890E+05 | 5.062E+05 | 5.380E+05 | 1.679E+06 |
| 157 | 429.146 | 4.153 | 20 | 0.000E+00 | 0.000E+00 | 0.000E+00 | 0.000E+00 | 0.000E+00 | 0.000E+00 | 0.000E+00 |
| 166 | 549.571 | 3.384 | 20 | 0.000E+00 | 0.000E+00 | 0.000E+00 | 0.000E+00 | 0.000E+00 | 0.000E+00 | 0.000E+00 |
| 481 | 549.078 | 3.211 | 20 | 0.000E+00 | 2.504E+05 | 2.346E+04 | 5.778E+06 | 3.665E+06 | 2.765E+06 | 0.000E+00 |
| 520 | 432.026 | 3.379 | 20 | 1.486E+07 | 8.033E+07 | 2.154E+07 | 3.654E+06 | 3.077E+07 | 5.610E+06 | 0.000E+00 |
| 584 | 432.026 | 3.502 | 20 | 1.394E+08 | 1.147E+07 | 1.052E+08 | 1.609E+07 | 1.156E+07 | 4.876E+07 | 7.556E+04 |
| 596 | 405.571 | 3.038 | 20 | 9.396E+07 | 4.655E+07 | 3.774E+06 | 3.313E+04 | 4.811E+04 | 2.450E+05 | 0.000E+00 |
| 600 | 359.104 | 0.595 | 20 | 2.034E+04 | 5.363E+05 | 3.390E+03 | 4.193E+06 | 7.843E+05 | 2.701E+06 | 0.000E+00 |
| 611 | 401.115 | 2.454 | 20 | 4.067E+05 | 4.457E+05 | 3.057E+05 | 2.088E+04 | 5.685E+04 | 5.881E+04 | 0.000E+00 |
| 622 | 411.021 | 3.222 | 20 | 2.939E+05 | 8.851E+05 | 2.158E+05 | 6.453E+07 | 2.683E+06 | 4.327E+06 | 0.000E+00 |
| 315 | 515.353 | 9.549 | 19 | 0.000E+00 | 0.000E+00 | 0.000E+00 | 0.000E+00 | 0.000E+00 | 0.000E+00 | 0.000E+00 |
| 326 | 321.148 | 3.682 | 19 | 0.000E+00 | 6.517E+03 | 0.000E+00 | 2.740E+04 | 1.092E+05 | 1.023E+05 | 0.000E+00 |
| 595 | 366.206 | 3.688 | 19 | 0.000E+00 | 0.000E+00 | 0.000E+00 | 0.000E+00 | 0.000E+00 | 3.139E+04 | 0.000E+00 |
| 494 | 453.189 | 7.153 | 18 | 2.689E+06 | 8.202E+06 | 6.413E+06 | 6.357E+06 | 7.060E+06 | 7.629E+06 | 0.000E+00 |
| 591 | 453.189 | 7.367 | 18 | 4.008E+06 | 5.000E+06 | 1.695E+06 | 2.528E+06 | 3.124E+06 | 3.744E+06 | 0.000E+00 |
| 861 | 453.187 | 7.297 | 18 | 1.320E+07 | 6.202E+06 | 5.060E+06 | 1.402E+06 | 3.221E+06 | 3.836E+06 | 0.000E+00 |
| 21 | 453.169 | 7.511 | 17 | 6.046E+07 | 7.784E+07 | 6.631E+07 | 7.070E+07 | 1.046E+06 | 3.654E+07 | 2.551E+06 |
| 292 | 453.168 | 7.525 | 17 | 1.077E+08 | 7.562E+07 | 6.631E+07 | 7.199E+07 | 7.302E+07 | 7.677E+07 | 3.198E+06 |
| 640 | 453.186 | 7.468 | 17 | 0.000E+00 | 8.640E+05 | 8.760E+07 | 4.251E+05 | 1.293E+06 | 8.255E+05 | 0.000E+00 |
| 59 | 281.138 | 7.525 | 16 | 3.997E+07 | 8.079E+06 | 1.369E+07 | 5.931E+06 | 5.857E+06 | 7.646E+06 | 6.538E+04 |
| 97 | 299.149 | 4.461 | 16 | 4.352E+07 | 8.335E+06 | 1.694E+07 | 3.671E+06 | 4.078E+06 | 4.633E+06 | 5.319E+03 |
| 172 | 299.148 | 7.523 | 16 | 2.019E+07 | 3.470E+06 | 6.333E+06 | 2.863E+06 | 2.133E+06 | 3.286E+06 | 5.433E+03 |
| 296 | 344.206 | 4.461 | 16 | 1.328E+07 | 2.200E+06 | 4.511E+06 | 9.161E+05 | 1.112E+06 | 1.413E+06 | 2.599E+04 |
| 381 | 316.176 | 4.464 | 16 | 6.726E+06 | 1.232E+06 | 2.675E+06 | 6.693E+05 | 7.671E+05 | 8.021E+05 | 1.937E+04 |
| 42 | 473.318 | 10.647 | 15 | 6.491E+06 | 4.412E+06 | 6.431E+06 | 7.030E+06 | 6.398E+06 | 4.902E+06 | 0.000E+00 |
| 72 | 680.481 | 12.424 | 15 | 8.952E+07 | 2.569E+07 | 3.525E+07 | 2.364E+07 | 2.183E+07 | 1.829E+07 | 8.893E+07 |
| 74 | 475.298 | 9.832 | 15 | 8.631E+07 | 2.577E+07 | 5.014E+07 | 1.680E+07 | 1.585E+07 | 1.914E+07 | 9.849E+06 |
| 95 | 492.324 | 9.832 | 15 | 8.642E+07 | 2.319E+07 | 4.486E+07 | 1.612E+07 | 1.467E+07 | 1.762E+07 | 1.767E+07 |
| 96 | 654.333 | 10.021 | 15 | 3.934E+07 | 1.630E+07 | 1.563E+07 | 1.630E+06 | 1.360E+06 | 6.977E+05 | 7.323E+04 |
| 105 | 490.345 | 10.647 | 15 | 3.202E+06 | 1.592E+06 | 2.785E+06 | 3.529E+06 | 2.642E+06 | 2.280E+06 | 0.000E+00 |
| 129 | 663.454 | 12.425 | 15 | 5.998E+07 | 1.744E+07 | 2.278E+07 | 1.385E+07 | 1.293E+07 | 1.126E+07 | 4.528E+07 |
| 141 | 708.511 | 12.424 | 15 | 4.356E+07 | 1.971E+07 | 2.577E+07 | 9.619E+06 | 1.085E+07 | 8.892E+06 | 1.827E+07 |
| 181 | 637.306 | 10.020 | 15 | 1.537E+07 | 7.582E+06 | 8.315E+06 | 8.683E+05 | 6.198E+05 | 4.492E+05 | 0.000E+00 |
| 205 | 518.376 | 10.647 | 15 | 1.900E+06 | 1.001E+06 | 1.813E+06 | 1.747E+06 | 1.371E+06 | 1.177E+06 | 0.000E+00 |
| 273 | 287.141 | 6.170 | 15 | 9.993E+06 | 3.923E+06 | 4.901E+06 | 2.364E+06 | 2.668E+06 | 2.694E+06 | 0.000E+00 |
| 298 | 515.329 | 10.526 | 15 | 6.284E+06 | 2.642E+06 | 2.220E+06 | 6.141E+05 | 1.161E+06 | 1.188E+06 | 0.000E+00 |
| 311 | 474.317 | 10.257 | 15 | 5.375E+06 | 2.199E+06 | 2.261E+06 | 1.665E+06 | 1.640E+06 | 1.183E+06 | 0.000E+00 |
| 317 | 457.288 | 10.396 | 15 | 2.526E+06 | 1.389E+06 | 1.586E+06 | 8.520E+05 | 1.099E+06 | 6.436E+05 | 0.000E+00 |
| 318 | 387.194 | 7.399 | 15 | 5.939E+06 | 3.800E+06 | 1.562E+06 | 7.692E+05 | 2.867E+05 | 5.254E+05 | 1.453E+05 |
| 426 | 682.364 | 10.020 | 15 | 1.025E+07 | 6.200E+06 | 6.034E+06 | 2.031E+05 | 2.096E+05 | 6.642E+04 | 0.000E+00 |
| 433 | 532.356 | 10.526 | 15 | 2.786E+06 | 9.830E+05 | 7.993E+05 | 3.467E+05 | 4.827E+05 | 5.670E+05 | 0.000E+00 |
| 452 | 519.377 | 10.257 | 15 | 1.455E+06 | 7.213E+05 | 9.550E+05 | 7.072E+05 | 7.882E+05 | 6.652E+05 | 0.000E+00 |
| 529 | 520.355 | 9.833 | 15 | 4.244E+07 | 1.033E+07 | 2.211E+07 | 6.910E+05 | 6.512E+05 | 8.063E+05 | 2.111E+06 |
| 726 | 1342.926 | 12.423 | 15 | 8.357E+06 | 2.557E+05 | 8.408E+05 | 1.997E+04 | 1.799E+05 | 4.302E+04 | 4.965E+05 |
| 755 | 949.587 | 9.831 | 15 | 3.424E+06 | 1.414E+05 | 4.346E+05 | 1.074E+04 | 3.975E+03 | 1.016E+04 | 5.332E+04 |
| 783 | 491.374 | 10.264 | 15 | 2.653E+05 | 4.115E+05 | 2.670E+05 | 5.030E+05 | 2.886E+05 | 6.505E+05 | 2.304E+06 |
| 849 | 1370.957 | 12.421 | 15 | 2.569E+06 | 7.417E+04 | 2.239E+05 | 0.000E+00 | 0.000E+00 | 1.445E+04 | 0.000E+00 |
| 585 | 834.093 | 3.504 | 14 | 1.730E+05 | 1.983E+05 | 9.229E+05 | 1.406E+06 | 7.070E+06 | 6.729E+06 | 0.000E+00 |
| 619 | 792.082 | 3.197 | 14 | 0.000E+00 | 4.669E+04 | 0.000E+00 | 1.160E+07 | 9.051E+06 | 6.853E+06 | 0.000E+00 |
| 714 | 542.297 | 4.609 | 13 | 5.299E+04 | 1.847E+04 | 5.793E+04 | 0.000E+00 | 0.000E+00 | 0.000E+00 | 0.000E+00 |
| 717 | 492.763 | 4.165 | 13 | 3.470E+04 | 7.809E+04 | 2.390E+04 | 0.000E+00 | 0.000E+00 | 0.000E+00 | 0.000E+00 |
| 51 | 247.034 | 5.943 | 12 | 0.000E+00 | 0.000E+00 | 0.000E+00 | 0.000E+00 | 0.000E+00 | 0.000E+00 | 0.000E+00 |
| 698 | 247.034 | 6.796 | 12 | 6.305E+06 | 1.108E+06 | 1.377E+07 | 0.000E+00 | 0.000E+00 | 0.000E+00 | 0.000E+00 |
| 718 | 291.023 | 6.253 | 12 | 6.249E+04 | 0.000E+00 | 1.237E+05 | 0.000E+00 | 0.000E+00 | 0.000E+00 | 0.000E+00 |
| 16 | 122.547 | 12.762 | 11 | 1.666E+08 | 1.667E+08 | 1.663E+08 | 2.029E+08 | 2.204E+08 | 2.129E+08 | 1.593E+08 |
| 60 | 122.547 | 14.272 | 11 | 1.714E+07 | 2.733E+06 | 3.296E+06 | 5.466E+06 | 5.436E+06 | 1.174E+06 | 3.449E+06 |
| 103 | 122.547 | 2.944 | 11 | 2.806E+06 | 1.617E+07 | 1.117E+06 | 6.941E+06 | 3.539E+06 | 1.704E+06 | 4.988E+06 |
| 113 | 122.547 | 2.472 | 11 | 1.719E+06 | 9.921E+05 | 1.905E+06 | 6.434E+06 | 4.707E+06 | 7.779E+06 | 5.231E+06 |
| 115 | 122.547 | 13.880 | 11 | 6.910E+05 | 9.355E+06 | 1.473E+06 | 8.832E+06 | 1.201E+07 | 1.449E+07 | 3.157E+06 |
| 120 | 122.547 | 2.832 | 11 | 2.810E+06 | 1.800E+06 | 1.397E+06 | 2.292E+06 | 3.539E+06 | 4.032E+06 | 7.710E+06 |
| 123 | 122.547 | 13.067 | 11 | 1.156E+06 | 2.785E+06 | 1.755E+06 | 2.160E+06 | 4.552E+06 | 4.970E+06 | 1.256E+07 |
| 125 | 122.547 | 2.720 | 11 | 4.006E+06 | 5.196E+06 | 5.294E+06 | 5.668E+06 | 3.978E+06 | 8.186E+06 | 5.043E+06 |
| 127 | 122.547 | 0.660 | 11 | 1.150E+06 | 1.291E+06 | 4.637E+06 | 6.327E+06 | 1.540E+07 | 2.148E+06 | 1.972E+06 |
| 135 | 122.547 | 13.843 | 11 | 6.910E+05 | 1.323E+06 | 1.543E+07 | 1.060E+07 | 3.034E+06 | 4.601E+06 | 9.359E+06 |
| 140 | 122.547 | 2.622 | 11 | 1.520E+06 | 5.970E+06 | 3.095E+06 | 7.817E+06 | 3.613E+06 | 8.244E+06 | 5.043E+06 |
| 145 | 122.547 | 2.598 | 11 | 6.550E+06 | 1.477E+06 | 5.469E+05 | 1.574E+06 | 4.600E+06 | 3.347E+06 | 1.966E+06 |
| 147 | 122.547 | 2.749 | 11 | 2.810E+06 | 2.709E+06 | 2.194E+06 | 1.494E+06 | 1.130E+07 | 1.944E+06 | 4.356E+06 |
| 151 | 122.547 | 13.172 | 11 | 5.884E+06 | 2.714E+06 | 4.443E+05 | 7.519E+06 | 2.506E+06 | 2.666E+06 | 1.977E+06 |
| 156 | 122.547 | 2.528 | 11 | 3.900E+06 | 1.982E+06 | 1.971E+06 | 3.495E+06 | 1.697E+06 | 5.953E+06 | 4.033E+06 |
| 160 | 122.547 | 2.262 | 11 | 1.794E+06 | 1.886E+06 | 1.552E+06 | 2.465E+06 | 1.756E+06 | 2.411E+06 | 2.957E+06 |

| Feature identifiers | | | | | | | | | | |
|---|---|---|---|---|---|---|---|---|---|---|
| ClusterIdx | m/z | RT [min] | Subnetwork | 5TUB83_I | 5TUB83_II | 5TUB83_III | TUB00_I | TUB00_II | TUB00_III | blank |
| 169 | 122.547 | 13.121 | 11 | 2.693E+06 | 5.104E+05 | 6.665E+06 | 5.250E+05 | 2.506E+06 | 8.845E+05 | 2.237E+06 |
| 174 | 122.547 | 2.436 | 11 | 2.112E+06 | 1.374E+06 | 1.563E+06 | 7.531E+05 | 2.894E+06 | 7.335E+05 | 5.594E+06 |
| 177 | 122.547 | 13.217 | 11 | 1.086E+06 | 3.256E+06 | 4.443E+05 | 5.830E+05 | 1.714E+06 | 3.075E+06 | 2.924E+06 |
| 178 | 122.547 | 2.294 | 11 | 2.028E+06 | 3.047E+06 | 2.259E+06 | 2.593E+06 | 1.572E+06 | 6.936E+06 | 5.537E+06 |
| 183 | 122.547 | 1.983 | 11 | 6.996E+05 | 1.026E+06 | 7.484E+05 | 2.697E+06 | 3.583E+06 | 5.961E+05 | 4.829E+06 |
| 185 | 122.547 | 13.651 | 11 | 3.933E+06 | 1.089E+06 | 4.290E+05 | 6.130E+06 | 7.771E+06 | 5.144E+06 | 1.027E+06 |
| 189 | 122.547 | 0.617 | 11 | 5.964E+06 | 9.218E+06 | 2.078E+06 | 3.634E+06 | 2.527E+06 | 1.293E+07 | 9.196E+06 |
| 206 | 122.547 | 1.515 | 11 | 1.384E+06 | 1.171E+06 | 7.984E+05 | 2.051E+06 | 2.903E+06 | 1.256E+06 | 1.577E+06 |
| 207 | 122.547 | 2.189 | 11 | 2.160E+06 | 1.222E+06 | 1.017E+06 | 3.636E+06 | 2.555E+06 | 4.152E+06 | 2.634E+06 |
| 216 | 122.547 | 1.421 | 11 | 4.487E+05 | 6.794E+05 | 1.116E+06 | 1.924E+06 | 5.624E+05 | 8.982E+05 | 2.701E+06 |
| 220 | 122.547 | 2.347 | 11 | 2.659E+06 | 5.995E+05 | 1.912E+06 | 3.144E+06 | 2.370E+06 | 4.721E+06 | 1.488E+06 |
| 248 | 122.547 | 2.158 | 11 | 2.160E+06 | 1.222E+06 | 1.699E+06 | 3.210E+06 | 2.555E+06 | 2.604E+06 | 5.926E+06 |
| 587 | 122.547 | 2.681 | 11 | 6.354E+06 | 2.709E+06 | 5.469E+05 | 1.636E+06 | 4.939E+06 | 1.325E+06 | 9.287E+06 |
| 644 | 122.547 | 2.919 | 11 | 1.455E+06 | 1.850E+06 | 6.582E+06 | 1.147E+07 | 1.102E+07 | 7.433E+05 | 1.720E+06 |
| 645 | 122.547 | 2.389 | 11 | 3.425E+06 | 2.534E+06 | 8.909E+05 | 3.484E+06 | 4.703E+06 | 1.605E+06 | 2.612E+06 |
| 656 | 122.547 | 2.584 | 11 | 1.436E+06 | 9.071E+05 | 5.469E+05 | 2.400E+06 | 5.190E+06 | 3.347E+06 | 2.164E+06 |
| 657 | 122.547 | 2.792 | 11 | 2.810E+06 | 1.440E+06 | 4.763E+06 | 1.241E+06 | 3.539E+06 | 1.944E+06 | 2.908E+06 |
| 710 | 122.547 | 3.154 | 11 | 1.373E+07 | 4.954E+06 | 3.316E+07 | 1.406E+07 | 1.934E+07 | 7.843E+06 | 4.077E+06 |
| 715 | 122.547 | 3.094 | 11 | 2.003E+06 | 4.954E+06 | 3.284E+06 | 6.076E+06 | 3.096E+06 | 1.704E+06 | 1.943E+06 |
| 719 | 122.547 | 2.222 | 11 | 2.337E+06 | 2.830E+06 | 4.943E+05 | 4.092E+06 | 1.270E+07 | 2.109E+06 | 4.847E+06 |
| 773 | 122.547 | 13.630 | 11 | 6.212E+06 | 1.617E+06 | 1.376E+06 | 2.981E+06 | 5.253E+05 | 1.359E+06 | 3.219E+06 |
| 61 | 537.536 | 11.439 | 10 | 4.754E+07 | 5.238E+07 | 5.121E+07 | 6.641E+07 | 6.104E+07 | 5.547E+07 | 4.031E+05 |
| 109 | 565.567 | 11.404 | 10 | 4.775E+07 | 4.338E+07 | 5.192E+07 | 6.876E+07 | 6.462E+07 | 5.854E+07 | 2.458E+05 |
| 295 | 582.594 | 11.445 | 10 | 8.075E+06 | 7.849E+06 | 8.395E+06 | 6.015E+06 | 7.187E+06 | 5.323E+06 | 0.000E+00 |
| 816 | 593.598 | 10.672 | 10 | 1.203E+06 | 5.481E+05 | 5.442E+05 | 1.338E+04 | 4.955E+07 | 1.250E+07 | 0.000E+00 |
| 191 | 608.393 | 9.449 | 9 | 3.908E+06 | 4.972E+06 | 3.423E+06 | 5.117E+06 | 4.161E+06 | 2.520E+06 | 5.933E+04 |
| 313 | 638.404 | 9.642 | 9 | 9.968E+05 | 6.631E+05 | 6.979E+05 | 1.033E+06 | 9.649E+05 | 6.599E+05 | 0.000E+00 |
| 348 | 624.388 | 9.076 | 9 | 6.437E+05 | 7.979E+05 | 7.316E+05 | 1.071E+06 | 7.714E+05 | 5.209E+05 | 6.261E+04 |
| 15 | 824.976 | 3.461 | 8 | 5.327E+08 | 3.726E+08 | 2.668E+08 | 4.985E+07 | 3.201E+07 | 3.652E+07 | 0.000E+00 |
| 18 | 810.973 | 3.183 | 8 | 3.564E+08 | 2.497E+08 | 1.490E+08 | 3.637E+07 | 1.988E+07 | 2.174E+07 | 0.000E+00 |
| 26 | 838.980 | 3.758 | 8 | 7.366E+08 | 4.538E+08 | 4.250E+08 | 2.784E+08 | 1.865E+08 | 2.234E+08 | 0.000E+00 |
| 65 | 1215.954 | 3.181 | 8 | 5.164E+07 | 3.773E+07 | 2.402E+07 | 6.456E+06 | 3.395E+06 | 4.013E+06 | 0.000E+00 |
| 87 | 1236.959 | 3.462 | 8 | 8.637E+07 | 6.797E+07 | 5.057E+07 | 1.061E+07 | 7.222E+06 | 6.931E+06 | 0.000E+00 |
| 121 | 837.625 | 3.464 | 8 | 4.496E+07 | 3.588E+07 | 2.474E+07 | 4.765E+06 | 3.241E+06 | 3.619E+06 | 0.000E+00 |
| 136 | 823.621 | 3.183 | 8 | 3.329E+07 | 2.674E+07 | 1.670E+07 | 3.589E+06 | 1.941E+06 | 2.064E+06 | 0.000E+00 |
| 194 | 618.984 | 3.459 | 8 | 3.520E+07 | 2.768E+07 | 1.632E+07 | 2.783E+06 | 1.623E+06 | 1.873E+06 | 0.000E+00 |
| 204 | 1257.965 | 3.758 | 8 | 1.371E+08 | 8.451E+07 | 9.125E+07 | 5.487E+07 | 4.026E+07 | 4.580E+07 | 0.000E+00 |
| 240 | 852.983 | 4.057 | 8 | 2.475E+08 | 1.384E+08 | 2.091E+08 | 3.508E+08 | 2.211E+08 | 2.590E+08 | 0.000E+00 |
| 244 | 846.306 | 3.758 | 8 | 5.679E+07 | 4.223E+07 | 3.696E+07 | 2.763E+07 | 1.810E+07 | 1.011E+07 | 0.000E+00 |
| 253 | 1069.394 | 4.071 | 8 | 4.218E+07 | 3.186E+07 | 2.035E+07 | 3.271E+06 | 1.612E+06 | 1.874E+06 | 0.000E+00 |
| 257 | 851.628 | 3.759 | 8 | 5.707E+07 | 3.992E+07 | 3.620E+07 | 2.482E+07 | 1.809E+07 | 2.019E+07 | 0.000E+00 |
| 258 | 608.481 | 3.183 | 8 | 1.625E+07 | 1.209E+07 | 6.533E+06 | 1.143E+06 | 5.938E+05 | 6.086E+05 | 0.000E+00 |
| 288 | 1083.397 | 4.311 | 8 | 2.333E+08 | 1.578E+08 | 1.417E+08 | 6.807E+07 | 4.488E+07 | 5.269E+07 | 0.000E+00 |
| 289 | 629.487 | 3.760 | 8 | 6.236E+07 | 3.657E+07 | 3.340E+07 | 2.088E+07 | 1.242E+07 | 1.511E+07 | 0.000E+00 |
| 312 | 1055.391 | 3.820 | 8 | 1.276E+07 | 9.427E+06 | 5.954E+06 | 7.773E+05 | 1.093E+05 | 1.197E+05 | 0.000E+00 |
| 352 | 1234.935 | 3.184 | 8 | 4.610E+06 | 4.296E+06 | 2.858E+06 | 4.754E+05 | 1.184E+05 | 4.180E+05 | 0.000E+00 |
| 406 | 802.298 | 4.071 | 8 | 1.923E+07 | 1.416E+07 | 9.031E+06 | 2.560E+05 | 3.318E+05 | 4.260E+05 | 0.000E+00 |
| 445 | 812.801 | 4.312 | 8 | 1.363E+08 | 8.678E+07 | 7.122E+07 | 2.957E+07 | 2.023E+07 | 2.424E+07 | 0.000E+00 |
| 450 | 1041.387 | 3.604 | 8 | 1.470E+06 | 3.651E+05 | 2.077E+05 | 2.424E+04 | 3.811E+04 | 3.240E+04 | 0.000E+00 |
| 457 | 1255.942 | 3.463 | 8 | 6.944E+06 | 6.367E+06 | 5.001E+06 | 1.337E+06 | 9.030E+05 | 6.295E+05 | 0.000E+00 |
| 497 | 1278.970 | 4.056 | 8 | 5.150E+07 | 2.708E+07 | 4.155E+07 | 7.647E+07 | 5.067E+07 | 5.871E+07 | 0.000E+00 |
| 504 | 1268.955 | 3.761 | 8 | 1.001E+07 | 1.288E+07 | 1.340E+07 | 9.243E+06 | 6.551E+06 | 7.521E+06 | 0.000E+00 |
| 511 | 865.632 | 4.058 | 8 | 2.127E+07 | 1.255E+07 | 1.722E+07 | 2.562E+07 | 1.779E+07 | 2.156E+07 | 0.000E+00 |
| 514 | 639.989 | 4.058 | 8 | 2.811E+07 | 1.619E+07 | 2.286E+07 | 3.581E+07 | 2.265E+07 | 2.573E+07 | 0.000E+00 |
| 515 | 860.310 | 4.057 | 8 | 2.191E+07 | 1.269E+07 | 1.779E+07 | 2.751E+07 | 1.925E+07 | 2.095E+07 | 0.000E+00 |
| 518 | 1276.945 | 3.762 | 8 | 1.174E+07 | 8.285E+06 | 7.286E+06 | 4.612E+06 | 5.038E+06 | 5.489E+06 | 0.000E+00 |
| 519 | 856.952 | 3.759 | 8 | 6.218E+06 | 6.990E+06 | 6.386E+06 | 5.482E+06 | 3.343E+06 | 6.351E+06 | 0.000E+00 |
| 522 | 1090.726 | 4.311 | 8 | 3.004E+07 | 2.005E+07 | 1.834E+07 | 9.224E+06 | 5.773E+06 | 7.124E+06 | 0.000E+00 |
| 531 | 1076.722 | 4.075 | 8 | 3.370E+06 | 4.522E+06 | 1.864E+06 | 5.223E+04 | 5.793E+04 | 1.484E+05 | 0.000E+00 |
| 533 | 1096.048 | 4.311 | 8 | 1.912E+07 | 1.333E+07 | 1.322E+07 | 6.143E+06 | 4.154E+06 | 5.045E+06 | 0.000E+00 |
| 534 | 791.796 | 3.823 | 8 | 4.228E+06 | 3.611E+06 | 1.757E+06 | 1.521E+05 | 7.472E+03 | 4.182E+04 | 0.000E+00 |
| 537 | 810.973 | 3.375 | 8 | 3.240E+07 | 1.078E+07 | 7.751E+06 | 1.097E+04 | 1.131E+04 | 4.927E+04 | 0.000E+00 |
| 559 | 824.976 | 3.689 | 8 | 4.585E+07 | 1.716E+07 | 1.141E+07 | 1.926E+05 | 8.993E+04 | 4.617E+05 | 0.000E+00 |
| 561 | 1289.962 | 4.060 | 8 | 7.533E+06 | 4.670E+06 | 6.870E+06 | 1.096E+07 | 7.661E+06 | 8.908E+06 | 0.000E+00 |
| 567 | 1082.043 | 4.072 | 8 | 4.616E+05 | 3.173E+06 | 1.023E+06 | 0.000E+00 | 0.000E+00 | 0.000E+00 | 0.000E+00 |
| 588 | 1027.384 | 3.427 | 8 | 1.773E+05 | 2.272E+05 | 1.051E+05 | 0.000E+00 | 0.000E+00 | 0.000E+00 | 0.000E+00 |
| 590 | 1013.379 | 3.335 | 8 | 7.690E+05 | 2.867E+05 | 1.061E+05 | 3.578E+05 | 2.923E+05 | 2.236E+05 | 0.000E+00 |
| 610 | 770.790 | 3.427 | 8 | 3.972E+04 | 2.128E+04 | 2.058E+04 | 1.731E+04 | 0.000E+00 | 0.000E+00 | 0.000E+00 |
| 614 | 837.291 | 3.461 | 8 | 3.612E+07 | 1.600E+07 | 2.150E+07 | 3.614E+06 | 1.708E+06 | 3.099E+06 | 0.000E+00 |
| 620 | 760.288 | 3.338 | 8 | 5.655E+04 | 3.779E+04 | 4.141E+04 | 3.406E+04 | 1.374E+04 | 4.755E+03 | 0.000E+00 |

| Feature identifiers | | | | | | | | | | |
|---|---|---|---|---|---|---|---|---|---|---|
| ClusterIdx | m/z | RT [min] | Subnetwork | 5TUB83_I | 5TUB83_II | 5TUB83_III | TUB00_I | TUB00_II | TUB00_III | blank |
| 633 | 1062.719 | 3.822 | 8 | 1.884E+06 | 1.491E+06 | 5.733E+05 | 0.000E+00 | 5.999E+04 | 0.000E+00 | 0.000E+00 |
| 650 | 629.236 | 3.761 | 8 | 4.556E+07 | 3.007E+07 | 2.686E+07 | 1.166E+07 | 6.142E+06 | 7.298E+06 | 0.000E+00 |
| 659 | 813.049 | 4.312 | 8 | 1.192E+08 | 7.646E+07 | 5.976E+07 | 2.248E+07 | 1.662E+07 | 2.217E+07 | 0.000E+00 |
| 664 | 866.987 | 4.353 | 8 | 8.286E+06 | 3.176E+06 | 7.602E+06 | 6.531E+07 | 4.263E+07 | 5.076E+07 | 0.000E+00 |
| 672 | 1299.976 | 4.354 | 8 | 4.950E+05 | 2.345E+05 | 9.076E+05 | 1.194E+07 | 7.803E+06 | 8.638E+06 | 0.000E+00 |
| 676 | 874.315 | 4.356 | 8 | 6.712E+05 | 5.996E+04 | 3.158E+05 | 5.918E+06 | 3.548E+06 | 4.332E+06 | 0.000E+00 |
| 677 | 856.617 | 3.758 | 8 | 1.116E+07 | 8.817E+06 | 7.696E+06 | 9.771E+06 | 6.129E+06 | 6.647E+06 | 0.000E+00 |
| 678 | 879.636 | 4.362 | 8 | 3.357E+05 | 0.000E+00 | 2.701E+05 | 4.593E+06 | 2.870E+06 | 3.763E+06 | 0.000E+00 |
| 680 | 1297.948 | 4.058 | 8 | 4.835E+06 | 3.124E+06 | 3.766E+06 | 6.280E+06 | 4.773E+06 | 5.487E+06 | 0.000E+00 |
| 683 | 975.108 | 4.244 | 8 | 1.397E+07 | 7.368E+06 | 6.501E+06 | 2.099E+06 | 2.650E+05 | 8.177E+05 | 0.000E+00 |
| 685 | 1101.371 | 4.312 | 8 | 3.584E+06 | 2.033E+06 | 2.232E+06 | 1.946E+06 | 1.176E+06 | 1.520E+06 | 0.000E+00 |
| 691 | 822.287 | 4.312 | 8 | 6.129E+06 | 3.621E+06 | 2.778E+06 | 1.341E+06 | 1.071E+06 | 1.054E+06 | 0.000E+00 |
| 730 | 796.969 | 3.070 | 8 | 1.397E+07 | 3.344E+06 | 1.387E+06 | 0.000E+00 | 0.000E+00 | 0.000E+00 | 0.000E+00 |
| 741 | 1068.040 | 3.824 | 8 | 1.070E+06 | 8.525E+05 | 4.415E+05 | 0.000E+00 | 0.000E+00 | 0.000E+00 | 0.000E+00 |
| 746 | 1069.395 | 4.241 | 8 | 4.743E+07 | 1.593E+07 | 1.368E+07 | 1.822E+04 | 1.095E+04 | 1.333E+05 | 0.000E+00 |
| 751 | 1221.446 | 3.186 | 8 | 1.015E+06 | 6.657E+05 | 1.955E+05 | 0.000E+00 | 0.000E+00 | 0.000E+00 | 0.000E+00 |
| 757 | 1055.390 | 3.992 | 8 | 6.015E+06 | 2.315E+06 | 1.375E+06 | 5.675E+03 | 0.000E+00 | 1.525E+04 | 0.000E+00 |
| 760 | 975.359 | 4.241 | 8 | 1.412E+07 | 7.465E+06 | 6.456E+06 | 1.861E+06 | 1.025E+06 | 1.575E+06 | 0.000E+00 |
| 761 | 802.298 | 4.239 | 8 | 3.190E+07 | 1.005E+07 | 8.111E+06 | 3.736E+04 | 1.795E+04 | 1.740E+05 | 0.000E+00 |
| 766 | 618.733 | 3.459 | 8 | 3.034E+07 | 2.182E+07 | 8.783E+06 | 1.455E+06 | 3.996E+05 | 1.205E+06 | 0.000E+00 |
| 768 | 838.979 | 3.988 | 8 | 2.640E+07 | 6.074E+06 | 1.279E+07 | 1.455E+05 | 3.447E+04 | 4.398E+05 | 0.000E+00 |
| 778 | 796.636 | 3.069 | 8 | 1.205E+07 | 3.057E+06 | 3.739E+05 | 0.000E+00 | 0.000E+00 | 0.000E+00 | 0.000E+00 |
| 785 | 1215.954 | 3.392 | 8 | 3.713E+06 | 7.417E+05 | 9.125E+05 | 0.000E+00 | 0.000E+00 | 0.000E+00 | 0.000E+00 |
| 787 | 608.481 | 3.384 | 8 | 3.538E+06 | 9.815E+05 | 5.488E+05 | 0.000E+00 | 0.000E+00 | 0.000E+00 | 0.000E+00 |
| 788 | 1236.958 | 3.690 | 8 | 6.261E+06 | 1.869E+06 | 2.138E+06 | 0.000E+00 | 0.000E+00 | 1.459E+04 | 0.000E+00 |
| 789 | 618.734 | 3.694 | 8 | 4.937E+06 | 1.508E+06 | 1.331E+06 | 0.000E+00 | 9.486E+03 | 1.268E+04 | 0.000E+00 |
| 791 | 792.044 | 3.972 | 8 | 2.684E+06 | 6.977E+05 | 3.904E+05 | 0.000E+00 | 0.000E+00 | 0.000E+00 | 0.000E+00 |
| 796 | 1069.727 | 4.238 | 8 | 4.010E+07 | 1.402E+07 | 1.215E+07 | 6.179E+04 | 5.142E+04 | 3.774E+05 | 0.000E+00 |
| 800 | 824.643 | 3.674 | 8 | 4.127E+07 | 1.456E+07 | 1.335E+07 | 6.870E+04 | 4.660E+04 | 4.277E+05 | 0.000E+00 |
| 801 | 1055.390 | 4.161 | 8 | 5.622E+06 | 4.277E+05 | 7.031E+05 | 0.000E+00 | 0.000E+00 | 0.000E+00 | 0.000E+00 |
| 803 | 791.795 | 4.163 | 8 | 4.248E+06 | 8.989E+05 | 8.955E+05 | 0.000E+00 | 0.000E+00 | 0.000E+00 | 0.000E+00 |
| 804 | 1300.145 | 4.240 | 8 | 1.005E+07 | 5.741E+06 | 5.245E+06 | 3.712E+04 | 6.722E+04 | 3.535E+04 | 0.000E+00 |
| 805 | 810.973 | 3.576 | 8 | 1.755E+06 | 1.986E+05 | 8.813E+04 | 0.000E+00 | 0.000E+00 | 0.000E+00 | 0.000E+00 |
| 807 | 1082.043 | 4.246 | 8 | 4.012E+06 | 1.427E+06 | 1.097E+06 | 0.000E+00 | 0.000E+00 | 3.931E+03 | 0.000E+00 |
| 809 | 629.487 | 3.997 | 8 | 3.964E+06 | 6.253E+05 | 1.497E+06 | 0.000E+00 | 0.000E+00 | 1.854E+04 | 0.000E+00 |
| 812 | 1268.461 | 3.759 | 8 | 1.546E+07 | 9.089E+06 | 1.164E+07 | 7.186E+06 | 5.146E+06 | 6.615E+06 | 0.000E+00 |
| 820 | 1300.134 | 4.241 | 8 | 1.533E+06 | 1.277E+06 | 2.312E+06 | 1.353E+06 | 9.763E+05 | 8.338E+05 | 0.000E+00 |
| 829 | 1137.663 | 4.195 | 8 | 3.770E+06 | 1.768E+06 | 1.824E+06 | 1.986E+05 | 3.648E+04 | 5.040E+04 | 0.000E+00 |
| 830 | 1263.457 | 3.755 | 8 | 2.997E+06 | 2.148E+06 | 6.992E+05 | 5.205E+05 | 6.087E+05 | 9.078E+05 | 0.000E+00 |
| 833 | 1076.722 | 4.246 | 8 | 5.104E+06 | 1.496E+06 | 1.516E+06 | 0.000E+00 | 0.000E+00 | 1.508E+04 | 0.000E+00 |
| 836 | 1063.052 | 3.825 | 8 | 1.372E+06 | 8.108E+05 | 2.998E+05 | 3.458E+04 | 0.000E+00 | 0.000E+00 | 0.000E+00 |
| 838 | 791.796 | 3.996 | 8 | 2.458E+06 | 1.007E+06 | 2.747E+05 | 0.000E+00 | 0.000E+00 | 1.025E+04 | 0.000E+00 |
| 842 | 822.537 | 4.313 | 8 | 3.739E+06 | 2.955E+06 | 3.008E+06 | 8.911E+05 | 8.461E+05 | 5.901E+05 | 0.000E+00 |
| 845 | 1257.962 | 3.997 | 8 | 5.580E+05 | 6.659E+05 | 1.158E+06 | 0.000E+00 | 0.000E+00 | 2.467E+04 | 0.000E+00 |
| 847 | 781.292 | 3.601 | 8 | 2.799E+05 | 9.613E+04 | 4.287E+04 | 5.910E+03 | 9.221E+03 | 0.000E+00 | 0.000E+00 |
| 848 | 811.784 | 4.077 | 8 | 6.898E+05 | 3.571E+05 | 2.074E+05 | 0.000E+00 | 0.000E+00 | 0.000E+00 | 0.000E+00 |
| 851 | 1276.945 | 3.763 | 8 | 6.814E+06 | 7.178E+06 | 3.324E+06 | 4.813E+06 | 2.961E+06 | 5.265E+06 | 0.000E+00 |
| 854 | 1194.947 | 3.069 | 8 | 1.031E+06 | 1.607E+05 | 4.625E+04 | 0.000E+00 | 0.000E+00 | 0.000E+00 | 0.000E+00 |
| 862 | 964.605 | 4.154 | 8 | 2.385E+06 | 5.929E+05 | 4.282E+05 | 0.000E+00 | 0.000E+00 | 0.000E+00 | 0.000E+00 |
| 354 | 828.611 | 3.182 | 7 | 8.443E+06 | 5.838E+06 | 2.018E+06 | 1.383E+06 | 6.448E+05 | 8.019E+05 | 0.000E+00 |
| 360 | 842.615 | 3.460 | 7 | 8.117E+06 | 7.925E+06 | 5.881E+06 | 1.796E+06 | 1.175E+06 | 6.260E+05 | 0.000E+00 |
| 649 | 842.948 | 3.459 | 7 | 1.020E+07 | 7.981E+06 | 1.731E+06 | 1.298E+06 | 6.385E+05 | 2.969E+05 | 0.000E+00 |
| 811 | 828.945 | 3.185 | 7 | 4.221E+06 | 6.235E+06 | 3.993E+06 | 1.224E+06 | 7.405E+05 | 7.636E+05 | 0.000E+00 |
| 22 | 274.274 | 6.821 | 6 | 4.278E+07 | 4.242E+07 | 4.177E+07 | 4.410E+07 | 4.305E+07 | 4.543E+07 | 8.587E+07 |
| 104 | 302.305 | 7.664 | 6 | 1.075E+07 | 1.004E+07 | 1.066E+07 | 9.045E+06 | 8.729E+06 | 8.524E+06 | 1.872E+07 |
| 277 | 330.337 | 8.495 | 6 | 4.706E+06 | 4.109E+06 | 4.338E+06 | 3.923E+06 | 3.830E+06 | 3.955E+06 | 1.307E+07 |
| 341 | 358.368 | 9.237 | 6 | 4.287E+06 | 3.128E+06 | 3.562E+06 | 3.709E+06 | 3.266E+06 | 3.441E+06 | 2.572E+07 |
| 321 | 311.185 | 7.063 | 5 | 2.008E+06 | 2.025E+06 | 2.077E+06 | 1.871E+06 | 1.874E+06 | 2.040E+06 | 3.486E+06 |
| 347 | 423.310 | 10.200 | 5 | 2.363E+06 | 1.172E+05 | 0.000E+00 | 6.941E+05 | 1.372E+06 | 1.847E+06 | 0.000E+00 |
| 661 | 468.369 | 10.166 | 5 | 1.281E+05 | 0.000E+00 | 0.000E+00 | 7.233E+04 | 1.386E+05 | 1.224E+05 | 0.000E+00 |
| 11 | 452.277 | 7.678 | 4 | 5.062E+08 | 2.729E+08 | 3.593E+08 | 2.163E+07 | 1.492E+07 | 1.778E+07 | 2.271E+04 |
| 155 | 474.259 | 7.679 | 4 | 2.254E+07 | 1.380E+07 | 1.836E+07 | 1.300E+06 | 9.427E+05 | 1.034E+06 | 0.000E+00 |
| 238 | 474.258 | 7.808 | 4 | 5.013E+06 | 3.180E+06 | 2.714E+06 | 1.327E+05 | 4.313E+04 | 1.627E+05 | 0.000E+00 |
| 350 | 690.508 | 12.705 | 4 | 7.693E+07 | 3.198E+07 | 2.613E+08 | 0.000E+00 | 0.000E+00 | 0.000E+00 | 0.000E+00 |
| 424 | 511.304 | 8.562 | 4 | 4.305E+07 | 8.348E+06 | 3.403E+07 | 6.003E+05 | 4.380E+05 | 4.447E+05 | 0.000E+00 |
| 486 | 690.508 | 10.348 | 4 | 1.192E+08 | 6.975E+07 | 1.343E+08 | 0.000E+00 | 0.000E+00 | 0.000E+00 | 0.000E+00 |
| 489 | 712.490 | 12.701 | 4 | 3.141E+07 | 1.285E+07 | 8.874E+07 | 0.000E+00 | 0.000E+00 | 0.000E+00 | 0.000E+00 |
| 505 | 712.490 | 10.356 | 4 | 4.196E+07 | 1.848E+07 | 2.118E+07 | 0.000E+00 | 0.000E+00 | 0.000E+00 | 0.000E+00 |
| 510 | 485.288 | 8.510 | 4 | 4.590E+07 | 1.055E+07 | 3.174E+07 | 9.263E+05 | 7.068E+05 | 6.182E+05 | 0.000E+00 |
| 513 | 743.484 | 12.700 | 4 | 2.431E+06 | 3.001E+07 | 8.769E+06 | 0.000E+00 | 0.000E+00 | 0.000E+00 | 0.000E+00 |

| Feature identifiers | | | | | | | | | | |
|---|---|---|---|---|---|---|---|---|---|---|
| ClusterIdx | m/z | RT [min] | Subnetwork | 5TUB83_I | 5TUB83_II | 5TUB83_III | TUB00_I | TUB00_II | TUB00_III | blank |
| 516 | 483.272 | 7.795 | 4 | 3.897E+07 | 9.762E+06 | 2.440E+07 | 4.616E+05 | 1.330E+05 | 2.436E+05 | 0.000E+00 |
| 517 | 740.522 | 12.702 | 4 | 7.320E+06 | 1.174E+07 | 3.639E+07 | 0.000E+00 | 0.000E+00 | 0.000E+00 | 0.000E+00 |
| 532 | 738.526 | 10.287 | 4 | 9.790E+07 | 8.702E+07 | 5.792E+06 | 0.000E+00 | 0.000E+00 | 6.463E+03 | 0.000E+00 |
| 554 | 903.547 | 7.681 | 4 | 8.297E+06 | 2.431E+06 | 5.922E+06 | 0.000E+00 | 0.000E+00 | 0.000E+00 | 0.000E+00 |
| 555 | 467.277 | 8.509 | 4 | 1.983E+07 | 4.321E+06 | 1.357E+07 | 4.228E+05 | 2.965E+05 | 1.591E+05 | 0.000E+00 |
| 562 | 743.484 | 10.306 | 4 | 5.185E+06 | 5.130E+07 | 1.270E+08 | 0.000E+00 | 0.000E+00 | 0.000E+00 | 0.000E+00 |
| 565 | 496.304 | 8.835 | 4 | 2.701E+05 | 2.955E+05 | 1.860E+05 | 5.819E+05 | 1.101E+06 | 6.290E+05 | 0.000E+00 |
| 577 | 483.272 | 7.944 | 4 | 3.985E+06 | 1.327E+06 | 1.602E+06 | 2.709E+04 | 2.495E+04 | 2.382E+04 | 3.388E+04 |
| 581 | 511.304 | 8.710 | 4 | 1.189E+07 | 2.387E+06 | 8.503E+06 | 8.260E+04 | 5.045E+04 | 3.100E+04 | 0.000E+00 |
| 583 | 452.277 | 7.806 | 4 | 8.918E+07 | 4.819E+07 | 4.223E+07 | 4.521E+06 | 2.667E+06 | 3.067E+06 | 0.000E+00 |
| 639 | 690.508 | 10.220 | 4 | 1.014E+08 | 1.080E+08 | 6.333E+06 | 8.570E+03 | 0.000E+00 | 0.000E+00 | 0.000E+00 |
| 643 | 712.490 | 10.268 | 4 | 5.071E+08 | 2.073E+08 | 1.542E+06 | 0.000E+00 | 0.000E+00 | 0.000E+00 | 0.000E+00 |
| 669 | 466.293 | 8.072 | 4 | 3.100E+07 | 1.363E+07 | 2.771E+07 | 1.597E+07 | 1.485E+07 | 1.443E+07 | 7.039E+03 |
| 679 | 466.293 | 8.206 | 4 | 4.041E+06 | 2.280E+06 | 2.694E+06 | 3.098E+06 | 2.252E+06 | 1.850E+06 | 1.264E+04 |
| 701 | 440.278 | 7.956 | 4 | 2.496E+06 | 1.139E+06 | 1.588E+06 | 0.000E+00 | 0.000E+00 | 0.000E+00 | 0.000E+00 |
| 713 | 662.477 | 12.666 | 4 | 1.434E+05 | 3.388E+07 | 5.365E+06 | 0.000E+00 | 1.198E+04 | 0.000E+00 | 0.000E+00 |
| 721 | 533.285 | 8.561 | 4 | 8.893E+06 | 2.406E+06 | 7.463E+06 | 5.600E+04 | 7.198E+04 | 5.612E+04 | 0.000E+00 |
| 722 | 440.278 | 7.818 | 4 | 7.857E+05 | 4.126E+05 | 4.825E+05 | 0.000E+00 | 0.000E+00 | 0.000E+00 | 0.000E+00 |
| 724 | 684.458 | 12.702 | 4 | 0.000E+00 | 1.391E+07 | 1.804E+06 | 0.000E+00 | 0.000E+00 | 0.000E+00 | 0.000E+00 |
| 727 | 465.262 | 7.796 | 4 | 1.079E+07 | 2.501E+06 | 6.646E+06 | 9.879E+04 | 2.324E+04 | 2.793E+04 | 0.000E+00 |
| 728 | 505.254 | 7.795 | 4 | 1.102E+07 | 2.796E+06 | 7.018E+06 | 4.000E+04 | 1.916E+04 | 1.384E+04 | 0.000E+00 |
| 729 | 507.269 | 8.510 | 4 | 9.909E+06 | 2.520E+06 | 7.298E+06 | 9.305E+04 | 6.250E+04 | 9.654E+04 | 0.000E+00 |
| 731 | 493.293 | 8.564 | 4 | 6.892E+06 | 1.747E+06 | 5.764E+06 | 9.420E+04 | 3.640E+04 | 2.125E+04 | 0.000E+00 |
| 736 | 676.492 | 12.707 | 4 | 5.281E+05 | 0.000E+00 | 3.630E+06 | 0.000E+00 | 0.000E+00 | 0.000E+00 | 0.000E+00 |
| 745 | 493.293 | 8.710 | 4 | 4.367E+06 | 7.531E+05 | 2.922E+06 | 1.428E+04 | 0.000E+00 | 4.600E+03 | 0.000E+00 |
| 749 | 500.300 | 7.793 | 4 | 4.508E+06 | 1.158E+06 | 2.919E+06 | 2.592E+04 | 0.000E+00 | 7.995E+03 | 0.000E+00 |
| 756 | 533.285 | 8.715 | 4 | 2.781E+06 | 5.828E+05 | 1.905E+06 | 1.309E+04 | 0.000E+00 | 0.000E+00 | 0.000E+00 |
| 775 | 704.523 | 12.703 | 4 | 5.399E+06 | 1.249E+07 | 2.334E+06 | 0.000E+00 | 0.000E+00 | 0.000E+00 | 0.000E+00 |
| 779 | 704.523 | 10.367 | 4 | 1.766E+07 | 7.788E+07 | 3.272E+06 | 0.000E+00 | 0.000E+00 | 0.000E+00 | 0.000E+00 |
| 781 | 726.506 | 12.701 | 4 | 2.462E+06 | 5.296E+06 | 4.447E+05 | 0.000E+00 | 0.000E+00 | 0.000E+00 | 0.000E+00 |
| 813 | 494.288 | 8.224 | 4 | 3.089E+04 | 2.154E+04 | 0.000E+00 | 0.000E+00 | 0.000E+00 | 2.230E+03 | 0.000E+00 |
| 825 | 664.490 | 9.773 | 4 | 2.730E+08 | 9.008E+07 | 7.686E+05 | 0.000E+00 | 0.000E+00 | 7.555E+03 | 0.000E+00 |
| 837 | 766.559 | 12.703 | 4 | 5.836E+05 | 3.757E+05 | 8.572E+06 | 0.000E+00 | 0.000E+00 | 0.000E+00 | 0.000E+00 |
| 841 | 676.491 | 10.424 | 4 | 7.418E+06 | 1.995E+04 | 3.110E+07 | 0.000E+00 | 0.000E+00 | 0.000E+00 | 0.000E+00 |
| 844 | 698.475 | 10.420 | 4 | 2.955E+05 | 0.000E+00 | 6.416E+06 | 0.000E+00 | 0.000E+00 | 0.000E+00 | 0.000E+00 |
| 853 | 1380.008 | 10.186 | 4 | 3.208E+07 | 6.376E+05 | 0.000E+00 | 0.000E+00 | 0.000E+00 | 0.000E+00 | 0.000E+00 |
| 855 | 1380.006 | 9.800 | 4 | 6.259E+06 | 1.825E+05 | 0.000E+00 | 0.000E+00 | 0.000E+00 | 0.000E+00 | 0.000E+00 |
| 27 | 316.212 | 3.277 | 3 | 2.425E+08 | 1.659E+08 | 1.543E+08 | 1.549E+08 | 1.339E+08 | 2.325E+08 | 1.644E+04 |
| 37 | 430.244 | 4.459 | 3 | 4.239E+07 | 3.154E+07 | 2.451E+07 | 2.661E+07 | 2.387E+07 | 2.834E+07 | 8.297E+03 |
| 55 | 300.200 | 3.938 | 3 | 5.567E+07 | 4.126E+07 | 4.330E+07 | 3.901E+07 | 3.489E+07 | 5.202E+07 | 3.426E+04 |
| 69 | 416.228 | 3.930 | 3 | 3.722E+07 | 2.770E+07 | 2.029E+07 | 2.402E+07 | 2.185E+07 | 2.997E+07 | 0.000E+00 |
| 122 | 243.183 | 3.050 | 3 | 3.477E+07 | 1.662E+07 | 2.074E+07 | 1.613E+07 | 1.534E+07 | 2.654E+07 | 0.000E+00 |
| 124 | 334.178 | 3.942 | 3 | 1.982E+07 | 2.057E+07 | 1.317E+07 | 1.886E+07 | 1.332E+07 | 7.917E+06 | 0.000E+00 |
| 128 | 357.214 | 4.538 | 3 | 1.424E+07 | 1.047E+07 | 8.935E+06 | 7.641E+06 | 7.702E+06 | 9.265E+06 | 0.000E+00 |
| 149 | 350.208 | 4.208 | 3 | 1.780E+07 | 1.428E+07 | 1.598E+07 | 1.023E+07 | 1.142E+07 | 1.573E+07 | 0.000E+00 |
| 167 | 599.391 | 3.938 | 3 | 2.398E+07 | 7.193E+06 | 7.810E+06 | 5.578E+06 | 4.261E+06 | 7.748E+06 | 0.000E+00 |
| 190 | 294.861 | 4.052 | 3 | 8.752E+06 | 1.317E+07 | 2.586E+07 | 1.657E+07 | 2.043E+07 | 9.003E+06 | 0.000E+00 |
| 224 | 302.196 | 0.594 | 3 | 4.887E+07 | 3.721E+07 | 3.747E+07 | 3.033E+07 | 3.294E+07 | 5.085E+07 | 7.632E+03 |
| 246 | 309.181 | 4.412 | 3 | 9.935E+06 | 6.789E+06 | 5.524E+06 | 7.298E+06 | 5.918E+06 | 4.371E+06 | 0.000E+00 |
| 268 | 328.200 | 4.325 | 3 | 6.878E+06 | 8.380E+06 | 8.083E+06 | 3.860E+06 | 5.152E+06 | 5.965E+06 | 0.000E+00 |
| 290 | 272.185 | 3.424 | 3 | 1.373E+07 | 1.638E+07 | 7.601E+06 | 6.252E+06 | 1.227E+07 | 1.816E+07 | 6.200E+03 |
| 293 | 293.191 | 3.577 | 3 | 8.303E+06 | 5.309E+06 | 6.671E+06 | 3.586E+06 | 4.507E+06 | 1.115E+07 | 0.000E+00 |
| 306 | 300.180 | 5.792 | 3 | 2.265E+06 | 2.234E+06 | 1.105E+06 | 1.356E+06 | 1.610E+06 | 2.119E+06 | 0.000E+00 |
| 322 | 713.423 | 4.538 | 3 | 4.929E+06 | 3.541E+06 | 3.182E+06 | 2.475E+06 | 2.803E+06 | 3.176E+06 | 0.000E+00 |
| 328 | 252.166 | 3.641 | 3 | 8.477E+06 | 3.959E+06 | 4.370E+06 | 7.407E+06 | 4.325E+06 | 4.369E+06 | 0.000E+00 |
| 329 | 332.872 | 4.600 | 3 | 5.922E+06 | 8.444E+06 | 7.750E+06 | 3.837E+06 | 6.104E+06 | 5.583E+06 | 0.000E+00 |
| 343 | 471.307 | 6.113 | 3 | 1.334E+06 | 1.184E+06 | 1.092E+06 | 8.252E+05 | 1.048E+06 | 1.163E+06 | 0.000E+00 |
| 361 | 441.788 | 4.079 | 3 | 1.842E+06 | 2.587E+06 | 8.304E+06 | 1.384E+06 | 5.076E+05 | 6.140E+05 | 0.000E+00 |
| 386 | 241.204 | 5.244 | 3 | 4.518E+07 | 1.632E+06 | 1.667E+06 | 2.197E+06 | 2.292E+06 | 4.303E+06 | 2.593E+06 |
| 396 | 469.328 | 4.897 | 3 | 2.273E+06 | 1.420E+06 | 1.642E+06 | 1.495E+06 | 1.290E+06 | 2.315E+06 | 0.000E+00 |
| 401 | 256.850 | 3.715 | 3 | 1.730E+07 | 1.387E+07 | 1.508E+07 | 9.267E+06 | 1.043E+07 | 1.348E+07 | 0.000E+00 |
| 404 | 699.407 | 4.204 | 3 | 4.417E+06 | 3.391E+06 | 3.852E+06 | 1.962E+06 | 2.514E+06 | 4.159E+06 | 0.000E+00 |
| 413 | 292.192 | 4.464 | 3 | 7.025E+05 | 2.062E+07 | 1.507E+07 | 5.420E+05 | 8.300E+05 | 3.171E+06 | 0.000E+00 |
| 439 | 585.375 | 3.578 | 3 | 4.271E+06 | 3.029E+06 | 3.578E+06 | 1.522E+06 | 2.063E+06 | 3.672E+06 | 0.000E+00 |
| 443 | 384.773 | 3.708 | 3 | 8.895E+06 | 1.256E+06 | 1.608E+06 | 6.422E+06 | 7.002E+06 | 7.004E+06 | 0.000E+00 |
| 463 | 293.191 | 3.406 | 3 | 4.317E+06 | 3.233E+06 | 3.538E+06 | 1.429E+06 | 2.190E+06 | 1.383E+06 | 0.000E+00 |
| 499 | 599.391 | 3.791 | 3 | 9.283E+06 | 1.875E+07 | 1.945E+07 | 1.570E+07 | 1.418E+07 | 2.154E+07 | 0.000E+00 |
| 526 | 485.359 | 3.054 | 3 | 7.513E+06 | 4.215E+06 | 5.021E+06 | 2.725E+06 | 2.552E+06 | 5.039E+06 | 0.000E+00 |
| 527 | 441.788 | 4.168 | 3 | 7.305E+06 | 8.382E+06 | 3.337E+06 | 6.356E+06 | 6.952E+06 | 7.760E+06 | 7.852E+03 |
| 530 | 290.193 | 3.898 | 3 | 9.943E+06 | 6.750E+06 | 7.143E+06 | 4.376E+06 | 4.289E+06 | 6.086E+06 | 2.856E+04 |

| Feature identifiers | | | | | | | | | | |
|---|---|---|---|---|---|---|---|---|---|---|
| ClusterIdx | m/z | RT [min] | Subnetwork | 5TUB83_I | 5TUB83_II | 5TUB83_III | TUB00_I | TUB00_II | TUB00_III | blank |
| 553 | 332.207 | 4.839 | 3 | 2.571E+06 | 1.976E+06 | 1.498E+06 | 1.129E+06 | 1.347E+06 | 1.023E+06 | 4.412E+04 |
| 568 | 379.210 | 4.343 | 3 | 2.384E+06 | 3.169E+06 | 3.339E+06 | 2.755E+05 | 9.198E+05 | 2.269E+06 | 0.000E+00 |
| 623 | 343.200 | 3.890 | 3 | 2.851E+06 | 1.609E+06 | 1.742E+06 | 6.984E+05 | 1.314E+06 | 2.114E+06 | 0.000E+00 |
| 624 | 263.684 | 4.125 | 3 | 9.385E+05 | 1.281E+07 | 8.086E+06 | 4.106E+05 | 7.839E+05 | 7.175E+05 | 8.765E+03 |
| 627 | 230.153 | 3.714 | 3 | 1.631E+06 | 1.147E+06 | 2.155E+05 | 7.168E+05 | 8.300E+05 | 4.496E+05 | 0.000E+00 |
| 629 | 347.535 | 4.414 | 3 | 1.647E+06 | 2.105E+06 | 1.893E+06 | 1.517E+05 | 5.270E+05 | 1.030E+06 | 0.000E+00 |
| 747 | 540.426 | 9.375 | 3 | 4.904E+06 | 1.072E+06 | 6.789E+05 | 4.945E+05 | 1.016E+06 | 8.717E+05 | 4.009E+04 |
| 750 | 568.458 | 9.844 | 3 | 4.868E+06 | 5.860E+05 | 5.370E+05 | 6.942E+04 | 7.841E+04 | 7.909E+04 | 0.000E+00 |
| 758 | 263.684 | 4.034 | 3 | 3.466E+05 | 8.790E+05 | 9.935E+05 | 2.266E+05 | 5.249E+05 | 9.077E+05 | 0.000E+00 |
| 782 | 220.150 | 0.536 | 3 | 3.513E+06 | 1.286E+06 | 3.624E+04 | 0.000E+00 | 1.879E+04 | 9.822E+04 | 0.000E+00 |
| 784 | 498.804 | 4.596 | 3 | 6.187E+05 | 2.042E+06 | 1.034E+06 | 3.158E+05 | 7.196E+05 | 1.020E+06 | 0.000E+00 |
| 810 | 294.861 | 4.145 | 3 | 2.432E+07 | 2.675E+07 | 7.933E+06 | 6.136E+06 | 2.325E+06 | 2.218E+07 | 0.000E+00 |
| 817 | 384.772 | 3.711 | 3 | 2.744E+06 | 6.160E+06 | 7.309E+06 | 1.950E+06 | 1.600E+06 | 2.220E+06 | 0.000E+00 |
| 819 | 292.192 | 4.644 | 3 | 7.525E+06 | 9.152E+05 | 3.340E+05 | 3.618E+05 | 1.389E+07 | 1.282E+06 | 7.604E+04 |
| 93 | 256.264 | 9.789 | 2 | 7.871E+06 | 8.413E+06 | 7.071E+06 | 7.429E+06 | 6.864E+06 | 7.110E+06 | 9.662E+06 |
| 142 | 284.295 | 10.325 | 2 | 3.735E+06 | 4.118E+06 | 3.940E+06 | 4.718E+06 | 4.924E+06 | 4.115E+06 | 7.963E+06 |
| 175 | 308.279 | 8.430 | 2 | 8.561E+06 | 3.914E+06 | 3.234E+06 | 2.481E+06 | 3.479E+06 | 3.233E+06 | 1.971E+07 |
| 218 | 482.408 | 9.562 | 2 | 4.362E+06 | 3.212E+06 | 3.576E+06 | 2.957E+06 | 2.867E+06 | 2.860E+06 | 1.813E+06 |
| 226 | 526.434 | 9.538 | 2 | 4.967E+06 | 3.494E+06 | 3.845E+06 | 2.765E+06 | 3.205E+06 | 2.913E+06 | 3.028E+06 |
| 228 | 336.311 | 9.373 | 2 | 4.194E+06 | 3.589E+06 | 3.633E+06 | 3.475E+06 | 3.389E+06 | 3.452E+06 | 1.979E+07 |
| 229 | 291.253 | 8.431 | 2 | 6.615E+06 | 2.684E+06 | 2.468E+06 | 1.600E+06 | 2.493E+06 | 2.377E+06 | 6.362E+06 |
| 241 | 424.364 | 9.342 | 2 | 3.746E+06 | 3.307E+06 | 3.117E+06 | 2.469E+06 | 2.377E+06 | 2.362E+06 | 5.964E+06 |
| 242 | 380.338 | 9.364 | 2 | 4.381E+06 | 4.043E+06 | 3.801E+06 | 3.213E+06 | 3.154E+06 | 3.328E+06 | 1.145E+07 |
| 247 | 352.306 | 8.443 | 2 | 5.963E+06 | 2.767E+06 | 2.262E+06 | 1.301E+06 | 2.239E+06 | 2.417E+06 | 2.351E+07 |
| 269 | 438.382 | 9.584 | 2 | 6.744E+06 | 5.377E+06 | 6.258E+06 | 5.265E+06 | 4.815E+06 | 4.659E+06 | 2.776E+06 |
| 270 | 408.372 | 10.006 | 2 | 4.095E+06 | 3.582E+06 | 3.836E+06 | 3.393E+06 | 3.368E+06 | 3.092E+06 | 1.228E+07 |
| 276 | 570.460 | 9.514 | 2 | 4.624E+06 | 3.188E+06 | 4.762E+06 | 2.392E+06 | 2.566E+06 | 2.270E+06 | 1.805E+06 |
| 278 | 512.417 | 9.299 | 2 | 2.338E+06 | 2.285E+06 | 1.731E+06 | 1.541E+06 | 1.524E+06 | 1.602E+06 | 2.941E+06 |
| 280 | 468.390 | 9.320 | 2 | 2.807E+06 | 2.533E+06 | 2.323E+06 | 1.764E+06 | 1.858E+06 | 1.841E+06 | 3.215E+06 |
| 291 | 366.322 | 8.959 | 2 | 3.526E+06 | 3.250E+06 | 3.257E+06 | 3.670E+06 | 3.623E+06 | 3.484E+06 | 1.777E+07 |
| 294 | 452.398 | 9.985 | 2 | 2.667E+06 | 2.097E+06 | 2.596E+06 | 2.245E+06 | 2.149E+06 | 2.265E+06 | 9.800E+06 |
| 297 | 556.443 | 9.273 | 2 | 2.315E+06 | 1.901E+06 | 1.572E+06 | 1.225E+06 | 1.237E+06 | 1.312E+06 | 2.576E+06 |
| 302 | 247.227 | 8.431 | 2 | 4.044E+06 | 1.978E+06 | 1.155E+06 | 1.178E+06 | 1.231E+06 | 1.084E+06 | 3.117E+06 |
| 303 | 392.338 | 10.385 | 2 | 2.259E+06 | 2.210E+06 | 2.340E+06 | 2.149E+06 | 1.978E+06 | 2.199E+06 | 1.119E+07 |
| 310 | 496.424 | 9.963 | 2 | 2.118E+06 | 2.226E+06 | 2.305E+06 | 1.989E+06 | 1.881E+06 | 1.959E+06 | 6.873E+06 |
| 336 | 364.344 | 10.020 | 2 | 3.908E+06 | 3.629E+06 | 3.640E+06 | 2.276E+06 | 2.174E+06 | 2.129E+06 | 1.961E+07 |
| 337 | 540.449 | 9.937 | 2 | 1.976E+06 | 1.589E+06 | 3.774E+06 | 1.543E+06 | 1.513E+06 | 1.527E+06 | 4.075E+06 |
| 344 | 476.301 | 4.018 | 2 | 2.680E+06 | 2.653E+06 | 2.469E+06 | 2.674E+06 | 2.885E+06 | 3.007E+06 | 1.199E+07 |
| 349 | 440.359 | 8.424 | 2 | 3.150E+06 | 1.517E+06 | 1.447E+06 | 8.706E+05 | 1.584E+06 | 1.629E+06 | 2.192E+05 |
| 359 | 600.469 | 9.251 | 2 | 1.607E+06 | 1.581E+06 | 1.174E+06 | 9.646E+05 | 8.456E+05 | 9.898E+05 | 2.129E+06 |
| 362 | 322.295 | 8.962 | 2 | 2.520E+06 | 2.497E+06 | 2.565E+06 | 2.934E+06 | 2.901E+06 | 2.679E+06 | 4.028E+07 |
| 364 | 614.485 | 9.493 | 2 | 3.552E+06 | 2.521E+06 | 2.168E+06 | 1.478E+06 | 1.635E+06 | 1.552E+06 | 2.589E+06 |
| 366 | 396.333 | 8.432 | 2 | 3.775E+06 | 1.844E+06 | 1.491E+06 | 1.019E+06 | 1.655E+06 | 1.661E+06 | 1.211E+06 |
| 367 | 422.389 | 10.267 | 2 | 1.916E+06 | 1.499E+06 | 1.816E+06 | 1.789E+06 | 1.744E+06 | 1.847E+06 | 4.591E+06 |
| 369 | 436.364 | 10.348 | 2 | 2.243E+06 | 2.419E+06 | 2.236E+06 | 2.282E+06 | 2.259E+06 | 2.235E+06 | 9.132E+06 |
| 377 | 422.347 | 10.141 | 2 | 4.730E+06 | 4.162E+06 | 2.478E+06 | 3.888E+06 | 4.003E+06 | 3.891E+06 | 1.421E+07 |
| 382 | 466.373 | 10.105 | 2 | 2.024E+06 | 1.936E+06 | 1.889E+06 | 3.505E+06 | 1.923E+06 | 3.393E+06 | 1.124E+07 |
| 383 | 319.284 | 9.373 | 2 | 1.670E+06 | 1.592E+06 | 1.637E+06 | 1.571E+06 | 1.367E+06 | 1.482E+06 | 5.544E+06 |
| 387 | 466.415 | 10.248 | 2 | 1.276E+06 | 1.353E+06 | 1.599E+06 | 1.526E+06 | 1.433E+06 | 1.367E+06 | 3.273E+06 |
| 389 | 526.434 | 9.632 | 2 | 1.889E+06 | 1.749E+06 | 1.685E+06 | 1.722E+06 | 1.682E+06 | 1.717E+06 | 3.483E+06 |
| 390 | 468.390 | 9.206 | 2 | 1.822E+06 | 1.266E+06 | 1.411E+06 | 1.066E+06 | 1.102E+06 | 9.945E+05 | 5.620E+04 |
| 399 | 520.331 | 4.142 | 2 | 1.949E+06 | 1.691E+06 | 1.923E+06 | 1.922E+06 | 1.901E+06 | 1.905E+06 | 1.256E+07 |
| 400 | 484.385 | 8.410 | 2 | 2.455E+06 | 1.302E+06 | 1.105E+06 | 3.098E+05 | 1.451E+06 | 1.396E+06 | 1.648E+05 |
| 412 | 512.416 | 9.184 | 2 | 1.992E+06 | 1.466E+06 | 1.393E+06 | 1.014E+06 | 1.069E+06 | 1.005E+06 | 1.585E+05 |
| 414 | 528.411 | 8.398 | 2 | 2.157E+06 | 9.848E+05 | 1.103E+06 | 3.879E+05 | 1.182E+06 | 1.140E+06 | 8.946E+04 |
| 416 | 564.360 | 4.248 | 2 | 1.062E+06 | 1.213E+06 | 1.365E+06 | 1.459E+06 | 1.522E+06 | 1.511E+06 | 9.939E+06 |
| 417 | 644.495 | 9.227 | 2 | 9.478E+05 | 1.016E+06 | 8.327E+05 | 6.893E+05 | 5.624E+05 | 4.699E+05 | 1.659E+06 |
| 422 | 480.390 | 10.316 | 2 | 1.577E+06 | 1.749E+06 | 1.618E+06 | 1.871E+06 | 1.685E+06 | 1.742E+06 | 7.393E+06 |
| 432 | 556.443 | 9.161 | 2 | 2.080E+06 | 1.283E+06 | 1.179E+06 | 8.470E+05 | 8.169E+05 | 8.894E+05 | 1.683E+05 |
| 438 | 584.475 | 9.912 | 2 | 8.135E+05 | 1.040E+06 | 1.155E+06 | 1.085E+06 | 9.492E+05 | 9.609E+05 | 2.794E+06 |
| 455 | 628.501 | 9.881 | 2 | 1.484E+06 | 6.928E+05 | 9.945E+05 | 9.161E+05 | 4.183E+05 | 7.927E+05 | 2.452E+06 |
| 464 | 608.392 | 4.346 | 2 | 5.456E+05 | 9.977E+05 | 8.885E+05 | 8.701E+05 | 9.984E+05 | 1.160E+06 | 7.244E+06 |
| 471 | 352.306 | 8.095 | 2 | 1.509E+07 | 2.061E+07 | 1.172E+07 | 1.026E+05 | 7.277E+04 | 1.145E+05 | 1.346E+05 |
| 528 | 308.279 | 8.075 | 2 | 9.091E+06 | 1.282E+07 | 5.597E+06 | 4.876E+04 | 2.139E+04 | 3.972E+04 | 2.338E+05 |
| 539 | 688.521 | 9.203 | 2 | 6.451E+05 | 6.727E+05 | 0.000E+00 | 1.790E+05 | 1.176E+05 | 3.590E+05 | 9.283E+05 |
| 541 | 363.310 | 9.363 | 2 | 9.017E+05 | 7.693E+05 | 9.165E+05 | 8.955E+05 | 8.828E+05 | 8.577E+05 | 1.234E+06 |
| 546 | 291.253 | 8.075 | 2 | 6.357E+06 | 9.304E+06 | 3.956E+06 | 1.079E+04 | 1.935E+04 | 1.822E+04 | 1.737E+05 |
| 548 | 396.333 | 8.097 | 2 | 1.669E+07 | 2.000E+07 | 1.285E+07 | 1.890E+05 | 3.154E+04 | 1.360E+05 | 1.756E+05 |
| 556 | 732.548 | 9.180 | 2 | 2.950E+05 | 4.911E+05 | 3.208E+04 | 1.481E+05 | 1.458E+05 | 1.286E+05 | 5.757E+05 |
| 564 | 440.359 | 8.096 | 2 | 1.746E+07 | 1.943E+07 | 1.210E+07 | 1.546E+05 | 2.196E+05 | 2.252E+05 | 1.514E+05 |

| Feature identifiers | | | | | | | | | | |
|---|---|---|---|---|---|---|---|---|---|---|
| ClusterIdx | m/z | RT [min] | Subnetwork | 5TUB83_I | 5TUB83_II | 5TUB83_III | TUB00_I | TUB00_II | TUB00_III | blank |
| 570 | 484.385 | 8.094 | 2 | 1.538E+07 | 1.522E+07 | 8.966E+06 | 8.931E+04 | 1.047E+05 | 2.735E+04 | 9.399E+04 |
| 571 | 308.279 | 8.290 | 2 | 8.369E+05 | 1.115E+06 | 5.281E+05 | 6.677E+04 | 2.018E+05 | 1.991E+05 | 2.546E+05 |
| 573 | 528.411 | 8.088 | 2 | 1.202E+07 | 1.075E+07 | 5.648E+06 | 4.634E+04 | 4.333E+04 | 4.562E+04 | 8.600E+04 |
| 576 | 247.227 | 8.064 | 2 | 8.772E+05 | 7.020E+05 | 9.723E+04 | 7.208E+03 | 3.786E+03 | 0.000E+00 | 1.289E+05 |
| 579 | 335.280 | 8.098 | 2 | 5.628E+06 | 7.744E+06 | 4.449E+06 | 2.362E+04 | 6.046E+03 | 1.169E+04 | 0.000E+00 |
| 580 | 572.437 | 8.084 | 2 | 8.869E+06 | 7.188E+06 | 3.395E+06 | 1.274E+04 | 3.117E+04 | 1.815E+04 | 1.807E+04 |
| 598 | 396.333 | 8.315 | 2 | 1.481E+06 | 1.839E+06 | 1.050E+06 | 7.812E+04 | 7.948E+04 | 4.223E+04 | 6.932E+04 |
| 604 | 440.359 | 8.302 | 2 | 1.461E+06 | 1.503E+06 | 7.949E+05 | 1.774E+05 | 1.993E+05 | 4.695E+04 | 1.366E+05 |
| 608 | 672.527 | 9.854 | 2 | 1.027E+06 | 4.664E+05 | 6.901E+05 | 8.085E+05 | 4.995E+05 | 4.978E+05 | 1.659E+06 |
| 615 | 350.328 | 9.687 | 2 | 2.096E+06 | 1.874E+06 | 1.733E+06 | 2.660E+06 | 2.053E+06 | 9.429E+06 | 4.210E+07 |
| 618 | 658.511 | 9.464 | 2 | 2.255E+06 | 1.754E+06 | 1.194E+06 | 6.342E+05 | 7.515E+05 | 9.288E+05 | 3.990E+05 |
| 625 | 702.536 | 9.449 | 2 | 1.148E+06 | 9.978E+05 | 7.578E+05 | 5.550E+05 | 5.351E+05 | 5.145E+05 | 1.262E+06 |
| 647 | 394.355 | 9.697 | 2 | 2.812E+06 | 1.914E+06 | 1.745E+06 | 2.179E+06 | 2.273E+06 | 2.411E+06 | 2.230E+07 |
| 50 | 701.620 | 10.613 | 1 | 7.008E+06 | 6.691E+06 | 7.010E+06 | 1.206E+07 | 1.049E+07 | 9.879E+06 | 4.248E+07 |
| 199 | 419.364 | 7.499 | 1 | 2.563E+06 | 2.629E+06 | 2.811E+06 | 3.441E+06 | 3.355E+06 | 3.148E+06 | 8.219E+06 |
| 689 | 685.624 | 12.053 | 1 | 5.115E+05 | 3.299E+05 | 5.545E+05 | 1.495E+06 | 7.935E+05 | 4.618E+05 | 1.054E+07 |
| 1 | 110.020 | 12.761 | -1 | 1.286E+09 | 1.221E+09 | 1.236E+09 | 1.187E+09 | 8.075E+08 | 1.233E+09 | 1.046E+09 |
| 8 | 437.194 | 7.524 | -1 | 3.620E+08 | 2.180E+08 | 2.631E+08 | 1.875E+08 | 1.911E+08 | 2.108E+08 | 7.365E+06 |
| 10 | 120.081 | 0.519 | -1 | 1.606E+08 | 1.651E+08 | 6.540E+07 | 6.281E+08 | 5.654E+08 | 8.128E+08 | 2.116E+04 |
| 13 | 242.284 | 8.651 | -1 | 2.371E+08 | 2.271E+08 | 2.331E+08 | 1.097E+08 | 7.587E+07 | 2.319E+08 | 3.668E+08 |
| 14 | 851.398 | 7.527 | -1 | 1.761E+08 | 3.832E+07 | 7.391E+07 | 3.053E+07 | 3.073E+07 | 4.059E+07 | 0.000E+00 |
| 32 | 437.194 | 7.249 | -1 | 3.193E+07 | 5.690E+06 | 2.520E+07 | 6.894E+06 | 6.552E+06 | 6.336E+06 | 5.134E+04 |
| 33 | 559.518 | 11.442 | -1 | 1.098E+08 | 1.193E+08 | 1.148E+08 | 1.236E+08 | 1.207E+08 | 1.235E+08 | 4.453E+04 |
| 34 | 867.372 | 7.528 | -1 | 4.311E+07 | 2.143E+07 | 3.037E+07 | 1.745E+07 | 1.838E+07 | 2.230E+07 | 0.000E+00 |
| 41 | 587.549 | 11.407 | -1 | 1.080E+08 | 1.130E+08 | 1.126E+08 | 1.120E+08 | 1.196E+08 | 1.219E+08 | 5.719E+04 |
| 47 | 163.991 | 12.748 | -1 | 1.467E+07 | 1.408E+07 | 1.392E+07 | 1.717E+07 | 1.812E+07 | 1.671E+07 | 4.397E+06 |
| 53 | 243.088 | 4.621 | -1 | 1.323E+07 | 1.063E+07 | 9.511E+06 | 5.153E+06 | 3.964E+06 | 4.867E+06 | 0.000E+00 |
| 54 | 399.209 | 10.100 | -1 | 5.985E+06 | 7.950E+06 | 8.991E+06 | 1.017E+07 | 1.124E+07 | 1.174E+07 | 3.093E+06 |
| 57 | 685.436 | 12.425 | -1 | 8.807E+07 | 2.977E+07 | 3.822E+07 | 2.361E+07 | 2.236E+07 | 1.889E+07 | 5.421E+07 |
| 58 | 250.112 | 5.753 | -1 | 4.791E+04 | 3.629E+04 | 1.362E+04 | 2.909E+05 | 8.389E+04 | 1.095E+05 | 1.273E+04 |
| 64 | 255.073 | 4.542 | -1 | 0.000E+00 | 0.000E+00 | 0.000E+00 | 0.000E+00 | 0.000E+00 | 0.000E+00 | 0.000E+00 |
| 66 | 261.109 | 5.951 | -1 | 1.489E+07 | 2.815E+07 | 3.382E+07 | 2.898E+07 | 2.768E+07 | 2.900E+07 | 6.941E+06 |
| 71 | 267.077 | 3.607 | -1 | 2.790E+07 | 1.631E+07 | 1.351E+07 | 2.672E+06 | 2.148E+06 | 2.533E+06 | 0.000E+00 |
| 73 | 453.168 | 7.250 | -1 | 1.332E+07 | 2.597E+06 | 1.106E+07 | 2.970E+06 | 3.006E+06 | 2.923E+06 | 1.079E+04 |
| 76 | 324.142 | 7.584 | -1 | 1.070E+07 | 1.133E+07 | 3.799E+06 | 2.443E+06 | 7.074E+06 | 6.553E+06 | 0.000E+00 |
| 77 | 372.882 | 12.723 | -1 | 1.017E+06 | 9.810E+05 | 2.283E+06 | 3.288E+06 | 2.563E+06 | 1.640E+06 | 5.011E+06 |
| 79 | 321.131 | 4.460 | -1 | 5.135E+07 | 1.172E+07 | 2.233E+07 | 4.092E+06 | 4.584E+06 | 5.389E+06 | 0.000E+00 |
| 82 | 269.015 | 5.941 | -1 | 0.000E+00 | 0.000E+00 | 0.000E+00 | 0.000E+00 | 0.000E+00 | 0.000E+00 | 0.000E+00 |
| 83 | 230.141 | 12.721 | -1 | 6.624E+06 | 6.711E+06 | 4.556E+06 | 5.766E+06 | 5.707E+06 | 6.089E+06 | 5.120E+06 |
| 85 | 233.079 | 4.825 | -1 | 8.762E+06 | 1.162E+07 | 1.563E+07 | 1.428E+07 | 1.203E+07 | 1.692E+07 | 2.507E+05 |
| 86 | 239.089 | 4.007 | -1 | 2.632E+07 | 3.444E+07 | 8.996E+06 | 8.169E+06 | 4.131E+07 | 4.576E+07 | 5.274E+04 |
| 90 | 332.913 | 12.737 | -1 | 6.435E+06 | 6.126E+06 | 5.712E+06 | 9.766E+06 | 1.010E+07 | 9.339E+06 | 1.532E+07 |
| 98 | 195.102 | 7.496 | -1 | 6.613E+06 | 1.051E+07 | 8.592E+06 | 7.022E+06 | 8.219E+06 | 7.917E+06 | 5.830E+06 |
| 100 | 440.099 | 7.644 | -1 | 0.000E+00 | 0.000E+00 | 0.000E+00 | 0.000E+00 | 0.000E+00 | 0.000E+00 | 0.000E+00 |
| 101 | 235.169 | 8.489 | -1 | 1.910E+07 | 2.252E+07 | 1.269E+07 | 1.199E+07 | 2.169E+07 | 1.323E+07 | 8.183E+06 |
| 106 | 670.153 | 5.750 | -1 | 4.685E+07 | 1.772E+07 | 4.036E+07 | 2.159E+06 | 4.771E+05 | 5.793E+05 | 0.000E+00 |
| 107 | 183.150 | 4.641 | -1 | 1.553E+07 | 1.836E+07 | 1.763E+07 | 6.223E+06 | 6.813E+06 | 2.732E+07 | 0.000E+00 |
| 108 | 281.088 | 8.127 | -1 | 0.000E+00 | 0.000E+00 | 0.000E+00 | 0.000E+00 | 0.000E+00 | 0.000E+00 | 0.000E+00 |
| 111 | 262.165 | 4.000 | -1 | 2.184E+07 | 2.448E+07 | 5.246E+06 | 4.699E+06 | 2.309E+07 | 2.539E+07 | 2.930E+04 |
| 114 | 244.191 | 5.975 | -1 | 5.452E+06 | 6.388E+06 | 5.154E+06 | 6.531E+06 | 5.263E+06 | 7.101E+06 | 9.602E+06 |
| 116 | 228.232 | 9.984 | -1 | 4.853E+06 | 5.409E+06 | 4.139E+06 | 9.490E+06 | 8.506E+06 | 7.932E+06 | 6.911E+04 |
| 118 | 219.175 | 8.820 | -1 | 6.395E+06 | 7.072E+06 | 5.798E+06 | 5.864E+06 | 5.728E+06 | 5.796E+06 | 4.692E+06 |
| 126 | 296.219 | 7.149 | -1 | 5.632E+06 | 9.831E+06 | 4.035E+06 | 1.962E+06 | 1.525E+06 | 2.076E+06 | 0.000E+00 |
| 137 | 431.183 | 10.115 | -1 | 3.896E+06 | 4.198E+06 | 4.917E+06 | 5.001E+06 | 5.094E+06 | 5.130E+06 | 1.803E+06 |
| 148 | 319.187 | 7.588 | -1 | 7.257E+06 | 6.367E+06 | 1.977E+06 | 1.429E+06 | 4.193E+06 | 3.854E+06 | 1.082E+04 |
| 152 | 707.875 | 4.385 | -1 | 0.000E+00 | 0.000E+00 | 0.000E+00 | 0.000E+00 | 0.000E+00 | 0.000E+00 | 0.000E+00 |
| 162 | 174.999 | 12.765 | -1 | 7.399E+06 | 8.488E+06 | 9.003E+06 | 1.172E+07 | 1.244E+07 | 1.097E+07 | 8.145E+06 |
| 164 | 338.147 | 5.707 | -1 | 0.000E+00 | 0.000E+00 | 0.000E+00 | 0.000E+00 | 0.000E+00 | 0.000E+00 | 0.000E+00 |
| 165 | 269.210 | 8.432 | -1 | 1.030E+07 | 5.692E+06 | 3.837E+06 | 3.884E+06 | 4.335E+06 | 3.795E+06 | 5.961E+06 |
| 173 | 451.246 | 9.774 | -1 | 2.609E+06 | 2.468E+06 | 1.957E+06 | 3.264E+06 | 4.250E+06 | 2.697E+06 | 0.000E+00 |
| 179 | 723.608 | 10.615 | -1 | 2.546E+06 | 2.220E+06 | 2.133E+06 | 4.774E+06 | 4.594E+06 | 4.063E+06 | 1.391E+07 |
| 184 | 434.186 | 7.527 | -1 | 1.663E+07 | 1.317E+06 | 3.203E+06 | 5.305E+05 | 5.797E+05 | 1.112E+06 | 0.000E+00 |
| 188 | 270.279 | 10.247 | -1 | 3.406E+06 | 2.639E+06 | 2.815E+06 | 4.260E+06 | 3.827E+06 | 3.839E+06 | 1.502E+06 |
| 193 | 242.284 | 7.258 | -1 | 2.284E+06 | 2.348E+06 | 2.210E+06 | 1.690E+07 | 1.490E+07 | 1.199E+07 | 1.614E+07 |
| 195 | 312.326 | 10.525 | -1 | 2.521E+06 | 1.584E+06 | 2.080E+06 | 2.503E+06 | 2.454E+06 | 2.167E+06 | 2.960E+06 |
| 198 | 499.165 | 7.526 | -1 | 1.178E+07 | 3.277E+06 | 5.885E+06 | 2.388E+06 | 2.661E+06 | 3.230E+06 | 0.000E+00 |
| 203 | 122.547 | 1.761 | -1 | 6.216E+05 | 2.440E+06 | 5.770E+05 | 1.851E+06 | 3.366E+06 | 4.494E+06 | 3.095E+06 |
| 212 | 409.201 | 10.114 | -1 | 5.691E+06 | 5.365E+06 | 4.546E+06 | 4.117E+06 | 4.187E+06 | 3.967E+06 | 9.893E+06 |
| 214 | 270.279 | 9.930 | -1 | 5.583E+06 | 5.277E+06 | 4.923E+06 | 8.786E+06 | 7.495E+06 | 7.655E+06 | 1.555E+07 |
| 217 | 266.248 | 6.949 | -1 | 0.000E+00 | 0.000E+00 | 0.000E+00 | 2.040E+04 | 1.082E+04 | 2.235E+04 | 0.000E+00 |

| Feature identifiers | | | | | | | | | | |
|---|---|---|---|---|---|---|---|---|---|---|
| ClusterIdx | m/z | RT [min] | Subnetwork | 5TUB83_I | 5TUB83_II | 5TUB83_III | TUB00_I | TUB00_II | TUB00_III | blank |
| 222 | 495.300 | 10.647 | -1 | 1.303E+06 | 7.520E+05 | 1.224E+06 | 1.353E+06 | 1.029E+06 | 9.765E+05 | 0.000E+00 |
| 225 | 330.300 | 9.151 | -1 | 0.000E+00 | 2.009E+03 | 0.000E+00 | 7.730E+05 | 9.211E+05 | 9.636E+05 | 0.000E+00 |
| 232 | 250.178 | 6.460 | -1 | 1.374E+07 | 2.461E+07 | 8.802E+06 | 8.124E+06 | 6.482E+06 | 8.427E+06 | 1.159E+07 |
| 234 | 555.250 | 4.014 | -1 | 0.000E+00 | 0.000E+00 | 0.000E+00 | 0.000E+00 | 0.000E+00 | 0.000E+00 | 0.000E+00 |
| 236 | 149.988 | 12.784 | -1 | 8.116E+06 | 6.935E+06 | 7.287E+06 | 8.744E+06 | 1.024E+07 | 9.377E+06 | 5.710E+05 |
| 237 | 640.348 | 4.374 | -1 | 0.000E+00 | 7.855E+04 | 1.189E+04 | 0.000E+00 | 0.000E+00 | 0.000E+00 | 0.000E+00 |
| 252 | 157.084 | 0.657 | -1 | 1.553E+07 | 1.624E+07 | 1.449E+07 | 1.442E+07 | 2.305E+07 | 2.735E+07 | 2.341E+05 |
| 254 | 111.039 | 13.492 | -1 | 2.339E+06 | 1.497E+06 | 1.396E+06 | 2.058E+06 | 4.474E+06 | 8.625E+05 | 7.630E+05 |
| 256 | 297.241 | 9.395 | -1 | 2.692E+06 | 3.116E+06 | 2.633E+06 | 3.473E+06 | 2.613E+06 | 2.910E+06 | 1.782E+07 |
| 259 | 249.185 | 8.504 | -1 | 3.637E+06 | 3.626E+06 | 2.902E+06 | 2.587E+06 | 2.859E+06 | 2.375E+06 | 1.907E+06 |
| 260 | 290.269 | 6.924 | -1 | 2.305E+06 | 2.201E+06 | 2.134E+06 | 2.560E+06 | 2.375E+06 | 2.559E+06 | 5.585E+06 |
| 265 | 535.153 | 8.517 | -1 | 0.000E+00 | 0.000E+00 | 0.000E+00 | 0.000E+00 | 0.000E+00 | 0.000E+00 | 0.000E+00 |
| 267 | 453.168 | 7.817 | -1 | 5.995E+06 | 2.804E+06 | 2.858E+06 | 1.686E+06 | 1.435E+06 | 1.568E+06 | 8.421E+04 |
| 271 | 283.104 | 5.343 | -1 | 0.000E+00 | 0.000E+00 | 0.000E+00 | 0.000E+00 | 0.000E+00 | 0.000E+00 | 0.000E+00 |
| 274 | 537.311 | 10.526 | -1 | 4.868E+06 | 2.147E+06 | 1.379E+06 | 5.852E+05 | 1.299E+06 | 1.425E+06 | 0.000E+00 |
| 279 | 149.024 | 6.761 | -1 | 4.752E+06 | 8.269E+06 | 3.015E+06 | 2.610E+06 | 6.280E+06 | 3.807E+06 | 1.149E+06 |
| 281 | 467.147 | 6.891 | -1 | 3.457E+07 | 6.251E+06 | 3.467E+06 | 3.437E+06 | 4.073E+06 | 5.663E+06 | 0.000E+00 |
| 285 | 408.309 | 10.166 | -1 | 4.758E+05 | 1.095E+06 | 2.203E+06 | 7.368E+05 | 1.134E+06 | 1.391E+06 | 8.029E+05 |
| 286 | 291.195 | 8.162 | -1 | 2.681E+06 | 2.217E+06 | 1.817E+06 | 1.366E+06 | 1.121E+06 | 3.336E+05 | 9.301E+04 |
| 287 | 573.110 | 8.520 | -1 | 0.000E+00 | 0.000E+00 | 0.000E+00 | 0.000E+00 | 0.000E+00 | 0.000E+00 | 1.845E+04 |
| 299 | 437.194 | 8.754 | -1 | 4.492E+06 | 2.853E+06 | 2.754E+06 | 2.737E+06 | 2.570E+06 | 2.451E+06 | 2.758E+06 |
| 300 | 604.812 | 4.484 | -1 | 0.000E+00 | 0.000E+00 | 0.000E+00 | 0.000E+00 | 0.000E+00 | 0.000E+00 | 0.000E+00 |
| 304 | 365.137 | 8.808 | -1 | 7.728E+06 | 7.730E+06 | 6.516E+06 | 1.162E+07 | 8.838E+06 | 1.304E+07 | 3.152E+07 |
| 305 | 480.273 | 3.605 | -1 | 0.000E+00 | 0.000E+00 | 0.000E+00 | 0.000E+00 | 0.000E+00 | 0.000E+00 | 0.000E+00 |
| 308 | 659.288 | 10.020 | -1 | 1.882E+07 | 9.500E+06 | 6.395E+06 | 2.279E+05 | 3.041E+05 | 1.635E+05 | 0.000E+00 |
| 309 | 165.076 | 4.458 | -1 | 1.028E+07 | 1.938E+06 | 4.063E+06 | 9.145E+05 | 1.076E+06 | 1.046E+06 | 1.782E+04 |
| 320 | 341.267 | 9.373 | -1 | 1.909E+06 | 2.243E+06 | 2.068E+06 | 2.001E+06 | 1.904E+06 | 2.112E+06 | 2.684E+06 |
| 324 | 270.315 | 9.426 | -1 | 2.120E+07 | 9.776E+06 | 1.791E+07 | 1.158E+07 | 1.597E+07 | 1.322E+07 | 2.067E+06 |
| 325 | 326.378 | 9.409 | -1 | 3.798E+06 | 3.460E+06 | 3.640E+06 | 3.696E+06 | 4.787E+06 | 3.933E+06 | 6.572E+06 |
| 327 | 131.049 | 0.513 | -1 | 3.039E+06 | 3.056E+06 | 3.075E+05 | 1.167E+07 | 1.031E+07 | 1.515E+07 | 1.317E+03 |
| 331 | 260.165 | 6.499 | -1 | 1.700E+06 | 1.747E+06 | 1.221E+06 | 8.717E+06 | 9.222E+06 | 9.400E+06 | 0.000E+00 |
| 334 | 469.184 | 6.409 | -1 | 5.520E+05 | 1.888E+06 | 1.594E+06 | 1.029E+06 | 1.657E+06 | 1.555E+06 | 0.000E+00 |
| 335 | 469.163 | 6.314 | -1 | 2.583E+07 | 1.332E+06 | 7.841E+05 | 6.519E+05 | 8.688E+05 | 1.393E+06 | 0.000E+00 |
| 338 | 499.780 | 3.714 | -1 | 0.000E+00 | 0.000E+00 | 0.000E+00 | 0.000E+00 | 0.000E+00 | 0.000E+00 | 0.000E+00 |
| 339 | 615.581 | 10.604 | -1 | 1.307E+07 | 1.315E+07 | 1.354E+07 | 6.812E+03 | 1.773E+06 | 1.589E+07 | 0.000E+00 |
| 340 | 354.553 | 5.749 | -1 | 9.848E+06 | 4.149E+06 | 8.791E+06 | 3.941E+05 | 3.391E+04 | 1.473E+05 | 0.000E+00 |
| 345 | 236.142 | 5.355 | -1 | 3.368E+06 | 2.287E+06 | 5.195E+05 | 5.119E+05 | 1.224E+06 | 9.583E+05 | 0.000E+00 |
| 351 | 451.173 | 6.881 | -1 | 4.204E+07 | 5.608E+06 | 3.306E+06 | 2.873E+06 | 3.299E+06 | 4.925E+06 | 0.000E+00 |
| 356 | 310.173 | 6.844 | -1 | 2.832E+06 | 1.502E+06 | 1.156E+06 | 1.157E+06 | 1.446E+06 | 1.393E+06 | 8.916E+03 |
| 365 | 519.261 | 9.835 | -1 | 8.543E+06 | 3.399E+06 | 7.367E+06 | 2.824E+06 | 2.769E+06 | 3.568E+06 | 1.453E+06 |
| 368 | 273.167 | 4.221 | -1 | 5.112E+06 | 4.653E+06 | 3.920E+06 | 3.457E+06 | 4.860E+06 | 6.959E+06 | 4.681E+05 |
| 376 | 288.181 | 4.136 | -1 | 4.067E+06 | 4.198E+06 | 1.704E+06 | 1.114E+06 | 2.016E+06 | 2.309E+06 | 1.888E+04 |
| 379 | 496.301 | 10.257 | -1 | 1.536E+06 | 8.770E+05 | 9.721E+05 | 1.016E+06 | 1.024E+06 | 8.352E+05 | 0.000E+00 |
| 380 | 497.280 | 9.835 | -1 | 1.762E+07 | 5.405E+06 | 1.041E+07 | 2.791E+06 | 2.519E+06 | 2.985E+06 | 2.953E+06 |
| 385 | 188.069 | 0.519 | -1 | 1.953E+05 | 1.164E+06 | 1.022E+05 | 9.859E+06 | 1.143E+07 | 1.384E+07 | 0.000E+00 |
| 395 | 721.506 | 12.424 | -1 | 9.890E+06 | 4.642E+06 | 5.646E+06 | 2.449E+06 | 2.571E+06 | 2.593E+06 | 2.475E+06 |
| 397 | 459.263 | 10.315 | -1 | 1.462E+06 | 6.573E+05 | 5.080E+05 | 5.446E+05 | 5.887E+05 | 6.703E+05 | 0.000E+00 |
| 398 | 445.291 | 10.157 | -1 | 2.441E+05 | 2.527E+04 | 0.000E+00 | 5.880E+05 | 5.756E+05 | 1.005E+06 | 6.944E+04 |
| 403 | 763.517 | 12.425 | -1 | 1.006E+07 | 4.127E+06 | 5.716E+06 | 2.673E+06 | 2.833E+06 | 2.444E+06 | 1.787E+06 |
| 405 | 528.788 | 4.417 | -1 | 0.000E+00 | 0.000E+00 | 0.000E+00 | 0.000E+00 | 0.000E+00 | 0.000E+00 | 0.000E+00 |
| 407 | 455.335 | 10.125 | -1 | 4.215E+06 | 1.752E+06 | 1.384E+06 | 6.604E+05 | 6.998E+05 | 7.201E+05 | 1.562E+06 |
| 410 | 830.949 | 3.185 | -1 | 2.319E+06 | 3.722E+06 | 1.139E+06 | 4.715E+05 | 1.822E+05 | 2.771E+05 | 0.000E+00 |
| 421 | 514.951 | 4.135 | -1 | 0.000E+00 | 0.000E+00 | 0.000E+00 | 0.000E+00 | 0.000E+00 | 0.000E+00 | 0.000E+00 |
| 423 | 409.180 | 7.399 | -1 | 3.854E+06 | 2.406E+06 | 9.731E+05 | 3.070E+05 | 1.733E+05 | 2.970E+05 | 1.732E+05 |
| 425 | 707.607 | 12.056 | -1 | 8.316E+05 | 6.404E+05 | 1.097E+06 | 2.546E+06 | 2.057E+06 | 1.967E+06 | 5.917E+06 |
| 428 | 366.633 | 3.310 | -1 | 0.000E+00 | 0.000E+00 | 0.000E+00 | 0.000E+00 | 7.295E+03 | 6.844E+03 | 0.000E+00 |
| 430 | 379.862 | 4.127 | -1 | 0.000E+00 | 0.000E+00 | 0.000E+00 | 0.000E+00 | 0.000E+00 | 0.000E+00 | 0.000E+00 |
| 436 | 267.137 | 8.329 | -1 | 5.711E+06 | 2.192E+06 | 3.411E+06 | 3.941E+05 | 4.688E+05 | 3.977E+05 | 0.000E+00 |
| 449 | 658.341 | 3.958 | -1 | 0.000E+00 | 0.000E+00 | 0.000E+00 | 0.000E+00 | 0.000E+00 | 0.000E+00 | 0.000E+00 |
| 451 | 254.660 | 3.065 | -1 | 0.000E+00 | 0.000E+00 | 0.000E+00 | 0.000E+00 | 0.000E+00 | 0.000E+00 | 0.000E+00 |
| 453 | 441.346 | 7.492 | -1 | 1.897E+05 | 6.251E+05 | 4.609E+05 | 4.947E+05 | 6.292E+05 | 5.964E+05 | 7.342E+06 |
| 454 | 409.163 | 6.914 | -1 | 2.801E+05 | 2.813E+05 | 2.324E+05 | 1.753E+06 | 1.222E+06 | 1.669E+06 | 0.000E+00 |
| 470 | 469.163 | 6.021 | -1 | 5.580E+06 | 5.520E+05 | 4.561E+05 | 4.298E+05 | 5.404E+05 | 6.215E+05 | 0.000E+00 |
| 474 | 501.170 | 0.519 | -1 | 1.619E+05 | 3.930E+05 | 4.039E+04 | 1.206E+05 | 0.000E+00 | 0.000E+00 | 0.000E+00 |
| 485 | 517.142 | 0.507 | -1 | 0.000E+00 | 0.000E+00 | 0.000E+00 | 0.000E+00 | 0.000E+00 | 0.000E+00 | 0.000E+00 |
| 496 | 226.012 | 12.739 | -1 | 1.049E+07 | 1.008E+07 | 9.816E+06 | 1.536E+07 | 9.467E+06 | 1.407E+07 | 1.721E+07 |
| 502 | 102.034 | 0.568 | -1 | 6.039E+06 | 9.707E+06 | 1.760E+06 | 5.504E+06 | 5.580E+06 | 6.524E+06 | 2.702E+06 |
| 503 | 228.207 | 4.644 | -1 | 7.542E+06 | 8.549E+06 | 9.076E+06 | 2.551E+06 | 3.009E+06 | 1.224E+07 | 6.584E+03 |
| 508 | 1011.286 | 0.501 | -1 | 0.000E+00 | 0.000E+00 | 0.000E+00 | 0.000E+00 | 0.000E+00 | 0.000E+00 | 0.000E+00 |
| 509 | 385.095 | 3.893 | -1 | 2.814E+04 | 1.948E+06 | 3.637E+04 | 1.511E+05 | 2.894E+04 | 1.611E+05 | 0.000E+00 |

| Feature identifiers | | | | | | | | | | |
|---|---|---|---|---|---|---|---|---|---|---|
| **ClusterIdx** | **m/z** | **RT [min]** | **Subnetwork** | **5TUB83_I** | **5TUB83_II** | **5TUB83_III** | **TUB00_I** | **TUB00_II** | **TUB00_III** | **blank** |
| 525 | 269.210 | 8.072 | -1 | 4.820E+06 | 6.127E+06 | 1.819E+06 | 1.826E+05 | 1.687E+05 | 2.569E+04 | 3.891E+05 |
| 538 | 844.952 | 3.465 | -1 | 4.149E+06 | 2.591E+06 | 2.352E+06 | 5.760E+05 | 4.188E+05 | 3.025E+05 | 0.000E+00 |
| 543 | 453.188 | 6.313 | -1 | 1.887E+07 | 7.811E+05 | 4.442E+05 | 4.212E+05 | 4.976E+05 | 9.343E+05 | 0.000E+00 |
| 547 | 372.717 | 12.740 | -1 | 2.932E+06 | 9.117E+05 | 1.267E+07 | 0.000E+00 | 0.000E+00 | 0.000E+00 | 0.000E+00 |
| 549 | 287.114 | 4.597 | -1 | 1.888E+06 | 8.365E+05 | 7.782E+05 | 3.257E+05 | 1.691E+05 | 1.197E+05 | 0.000E+00 |
| 551 | 385.294 | 9.364 | -1 | 9.896E+05 | 9.229E+05 | 9.968E+05 | 7.510E+05 | 6.681E+05 | 7.833E+05 | 3.705E+05 |
| 557 | 858.957 | 3.760 | -1 | 5.195E+06 | 4.090E+06 | 3.242E+06 | 2.705E+05 | 1.372E+06 | 1.888E+06 | 0.000E+00 |
| 563 | 429.319 | 9.342 | -1 | 8.407E+05 | 7.638E+05 | 7.128E+05 | 4.452E+05 | 4.612E+05 | 2.873E+05 | 4.798E+05 |
| 566 | 472.175 | 4.688 | -1 | 4.093E+05 | 1.647E+06 | 1.576E+05 | 5.276E+05 | 8.690E+05 | 1.118E+06 | 0.000E+00 |
| 569 | 294.074 | 0.519 | -1 | 1.030E+07 | 1.129E+07 | 3.064E+06 | 0.000E+00 | 1.924E+04 | 0.000E+00 | 0.000E+00 |
| 575 | 453.189 | 6.022 | -1 | 4.984E+06 | 4.412E+05 | 3.668E+05 | 1.738E+05 | 4.034E+05 | 5.153E+05 | 0.000E+00 |
| 586 | 191.086 | 3.681 | -1 | 0.000E+00 | 0.000E+00 | 1.457E+04 | 1.304E+04 | 5.493E+04 | 1.059E+05 | 5.183E+03 |
| 592 | 359.104 | 3.680 | -1 | 0.000E+00 | 0.000E+00 | 0.000E+00 | 0.000E+00 | 7.025E+03 | 1.868E+04 | 0.000E+00 |
| 593 | 511.165 | 3.612 | -1 | 5.139E+06 | 3.491E+05 | 3.507E+05 | 0.000E+00 | 0.000E+00 | 0.000E+00 | 0.000E+00 |
| 594 | 700.315 | 10.020 | -1 | 4.458E+06 | 1.824E+06 | 1.342E+06 | 9.626E+04 | 2.902E+04 | 7.434E+03 | 0.000E+00 |
| 597 | 225.149 | 6.492 | -1 | 6.387E+05 | 4.088E+06 | 5.694E+05 | 6.648E+05 | 3.488E+06 | 2.496E+06 | 1.982E+05 |
| 599 | 131.049 | 0.515 | -1 | 3.664E+05 | 5.527E+05 | 3.075E+05 | 1.522E+06 | 5.753E+06 | 1.012E+07 | 1.317E+03 |
| 601 | 357.262 | 8.098 | -1 | 2.710E+06 | 4.573E+06 | 1.366E+06 | 5.305E+03 | 0.000E+00 | 0.000E+00 | 0.000E+00 |
| 602 | 303.138 | 3.688 | -1 | 0.000E+00 | 0.000E+00 | 1.554E+04 | 0.000E+00 | 0.000E+00 | 1.150E+04 | 0.000E+00 |
| 605 | 347.222 | 7.583 | -1 | 4.464E+06 | 3.845E+06 | 1.455E+06 | 9.863E+05 | 2.538E+06 | 2.075E+06 | 0.000E+00 |
| 606 | 438.143 | 7.111 | -1 | 0.000E+00 | 0.000E+00 | 0.000E+00 | 0.000E+00 | 0.000E+00 | 0.000E+00 | 0.000E+00 |
| 607 | 851.398 | 7.250 | -1 | 7.859E+05 | 0.000E+00 | 4.199E+05 | 0.000E+00 | 2.185E+04 | 1.696E+04 | 0.000E+00 |
| 609 | 830.614 | 3.188 | -1 | 4.439E+06 | 1.515E+06 | 2.079E+06 | 4.761E+05 | 8.030E+04 | 2.904E+05 | 0.000E+00 |
| 612 | 863.046 | 3.487 | -1 | 1.334E+07 | 4.981E+05 | 9.055E+06 | 1.080E+05 | 7.229E+05 | 9.432E+05 | 0.000E+00 |
| 613 | 619.272 | 4.460 | -1 | 1.645E+07 | 7.043E+05 | 3.074E+06 | 6.316E+04 | 1.340E+05 | 2.016E+05 | 0.000E+00 |
| 617 | 454.172 | 7.812 | -1 | 1.530E+06 | 5.477E+05 | 5.912E+05 | 1.369E+05 | 2.088E+05 | 1.725E+05 | 1.105E+04 |
| 621 | 815.644 | 3.349 | -1 | 5.646E+05 | 8.800E+04 | 7.836E+03 | 0.000E+00 | 0.000E+00 | 0.000E+00 | 0.000E+00 |
| 626 | 331.209 | 4.774 | -1 | 1.170E+06 | 8.481E+05 | 1.600E+06 | 1.939E+06 | 1.793E+06 | 2.580E+06 | 5.723E+06 |
| 628 | 103.054 | 0.515 | -1 | 2.158E+05 | 5.472E+05 | 1.501E+05 | 8.771E+05 | 4.615E+06 | 1.284E+06 | 7.817E+03 |
| 641 | 158.154 | 6.250 | -1 | 8.351E+07 | 4.696E+07 | 8.242E+07 | 2.435E+07 | 7.924E+06 | 1.930E+07 | 3.501E+07 |
| 642 | 165.114 | 12.714 | -1 | 1.910E+07 | 1.370E+07 | 9.748E+06 | 3.049E+07 | 2.885E+07 | 4.008E+07 | 1.257E+07 |
| 648 | 425.137 | 6.913 | -1 | 1.235E+05 | 1.677E+05 | 1.641E+05 | 1.200E+06 | 9.139E+05 | 9.424E+05 | 0.000E+00 |
| 651 | 267.157 | 4.332 | -1 | 9.644E+05 | 4.682E+06 | 2.265E+06 | 1.313E+06 | 5.319E+06 | 4.371E+06 | 1.257E+05 |
| 655 | 290.232 | 4.304 | -1 | 3.142E+05 | 6.346E+05 | 5.785E+05 | 2.576E+05 | 2.239E+06 | 9.169E+05 | 6.177E+03 |
| 660 | 391.205 | 10.042 | -1 | 2.080E+06 | 1.704E+05 | 4.477E+04 | 5.547E+05 | 3.095E+05 | 3.650E+05 | 0.000E+00 |
| 673 | 870.956 | 4.058 | -1 | 4.084E+06 | 8.384E+05 | 3.529E+06 | 9.218E+06 | 5.280E+06 | 6.048E+06 | 0.000E+00 |
| 682 | 425.215 | 9.512 | -1 | 1.420E+06 | 1.700E+06 | 2.241E+06 | 3.367E+06 | 1.617E+06 | 1.129E+06 | 2.341E+08 |
| 684 | 612.425 | 0.524 | -1 | 4.908E+04 | 5.419E+05 | 2.831E+04 | 2.261E+06 | 2.711E+05 | 3.020E+06 | 0.000E+00 |
| 686 | 231.081 | 0.527 | -1 | 1.737E+04 | 5.472E+03 | 5.191E+03 | 2.365E+06 | 2.524E+06 | 3.161E+06 | 3.576E+03 |
| 688 | 726.463 | 12.425 | -1 | 6.342E+06 | 1.815E+06 | 2.461E+06 | 1.802E+06 | 1.741E+06 | 1.247E+06 | 3.785E+06 |
| 690 | 403.233 | 9.514 | -1 | 8.666E+05 | 4.518E+05 | 5.379E+05 | 1.067E+06 | 7.747E+05 | 3.137E+05 | 2.289E+08 |
| 694 | 111.020 | 12.539 | -1 | 4.607E+05 | 3.231E+05 | 2.110E+05 | 5.843E+05 | 1.728E+05 | 3.063E+05 | 1.772E+06 |
| 695 | 351.102 | 8.964 | -1 | 5.934E+05 | 3.664E+05 | 2.204E+05 | 6.092E+05 | 4.037E+05 | 7.585E+05 | 4.912E+05 |
| 700 | 164.051 | 4.799 | -1 | 2.164E+07 | 3.868E+07 | 3.080E+07 | 0.000E+00 | 5.461E+03 | 0.000E+00 | 0.000E+00 |
| 702 | 111.039 | 2.302 | -1 | 2.306E+06 | 2.306E+06 | 2.372E+06 | 5.802E+05 | 1.900E+06 | 3.232E+06 | 4.944E+06 |
| 705 | 271.031 | 5.195 | -1 | 2.031E+04 | 3.487E+04 | 1.878E+05 | 0.000E+00 | 0.000E+00 | 0.000E+00 | 0.000E+00 |
| 706 | 249.050 | 5.192 | -1 | 1.143E+04 | 1.863E+04 | 1.921E+05 | 0.000E+00 | 0.000E+00 | 0.000E+00 | 0.000E+00 |
| 707 | 279.060 | 6.671 | -1 | 5.913E+05 | 1.056E+05 | 9.442E+05 | 0.000E+00 | 0.000E+00 | 0.000E+00 | 0.000E+00 |
| 708 | 162.035 | 6.813 | -1 | 8.067E+05 | 1.272E+05 | 1.087E+06 | 0.000E+00 | 0.000E+00 | 0.000E+00 | 0.000E+00 |
| 709 | 177.083 | 2.375 | -1 | 2.974E+03 | 5.720E+03 | 0.000E+00 | 1.153E+04 | 5.382E+03 | 3.759E+03 | 1.460E+04 |
| 711 | 164.051 | 2.397 | -1 | 6.025E+03 | 2.745E+04 | 9.378E+03 | 0.000E+00 | 0.000E+00 | 1.579E+03 | 0.000E+00 |
| 712 | 245.070 | 2.402 | -1 | 0.000E+00 | 0.000E+00 | 3.100E+03 | 0.000E+00 | 0.000E+00 | 0.000E+00 | 0.000E+00 |
| 716 | 111.039 | 2.354 | -1 | 5.926E+05 | 2.306E+06 | 2.372E+06 | 5.802E+05 | 1.900E+06 | 2.201E+06 | 5.101E+06 |
| 720 | 495.251 | 4.182 | -1 | 1.739E+06 | 1.286E+06 | 9.289E+05 | 0.000E+00 | 0.000E+00 | 0.000E+00 | 0.000E+00 |
| 723 | 162.073 | 2.320 | -1 | 7.183E+03 | 9.594E+03 | 3.566E+03 | 5.787E+03 | 2.902E+03 | 8.605E+03 | 0.000E+00 |
| 725 | 1347.881 | 12.426 | -1 | 7.436E+06 | 7.466E+05 | 1.091E+06 | 7.701E+04 | 9.616E+04 | 5.532E+04 | 4.222E+05 |
| 733 | 297.083 | 6.817 | -1 | 4.986E+07 | 2.110E+07 | 5.904E+07 | 0.000E+00 | 0.000E+00 | 0.000E+00 | 0.000E+00 |
| 734 | 471.252 | 7.681 | -1 | 2.697E+06 | 8.787E+05 | 2.168E+06 | 0.000E+00 | 0.000E+00 | 0.000E+00 | 0.000E+00 |
| 735 | 597.678 | 0.494 | -1 | 4.621E+07 | 2.975E+07 | 9.342E+06 | 1.569E+05 | 1.770E+05 | 1.023E+05 | 0.000E+00 |
| 737 | 701.411 | 12.427 | -1 | 5.183E+06 | 1.797E+06 | 2.458E+06 | 5.326E+05 | 9.618E+05 | 4.696E+05 | 1.528E+06 |
| 738 | 253.602 | 7.678 | -1 | 2.412E+06 | 1.344E+06 | 1.559E+06 | 3.417E+04 | 3.019E+04 | 4.564E+04 | 0.000E+00 |
| 739 | 749.548 | 12.237 | -1 | 5.783E+06 | 1.125E+04 | 1.238E+06 | 0.000E+00 | 0.000E+00 | 1.119E+04 | 0.000E+00 |
| 742 | 287.650 | 8.328 | -1 | 1.778E+06 | 8.229E+05 | 1.043E+06 | 2.393E+05 | 1.632E+05 | 8.709E+04 | 0.000E+00 |
| 744 | 247.129 | 0.492 | -1 | 2.051E+04 | 5.427E+06 | 0.000E+00 | 4.486E+04 | 3.592E+04 | 9.198E+04 | 5.457E+02 |
| 748 | 654.456 | 10.203 | -1 | 3.528E+06 | 6.445E+05 | 9.920E+05 | 0.000E+00 | 0.000E+00 | 0.000E+00 | 0.000E+00 |
| 752 | 286.520 | 6.796 | -1 | 0.000E+00 | 0.000E+00 | 0.000E+00 | 0.000E+00 | 0.000E+00 | 0.000E+00 | 0.000E+00 |
| 753 | 335.125 | 8.963 | -1 | 7.653E+05 | 3.739E+05 | 3.016E+05 | 8.271E+05 | 4.936E+05 | 9.666E+05 | 6.921E+05 |
| 754 | 533.934 | 3.609 | -1 | 3.267E+06 | 4.195E+06 | 1.583E+06 | 0.000E+00 | 0.000E+00 | 0.000E+00 | 0.000E+00 |
| 762 | 533.282 | 9.675 | -1 | 9.508E+04 | 1.317E+05 | 1.219E+05 | 1.461E+05 | 1.137E+05 | 2.740E+05 | 0.000E+00 |
| 763 | 1194.349 | 0.496 | -1 | 1.861E+07 | 1.183E+07 | 4.125E+06 | 4.428E+03 | 4.871E+04 | 2.241E+04 | 0.000E+00 |

| Feature identifiers | | | | | | | | | | |
|---|---|---|---|---|---|---|---|---|---|---|
| **ClusterIdx** | **m/z** | **RT [min]** | **Subnetwork** | **5TUB83_I** | **5TUB83_II** | **5TUB83_III** | **TUB00_I** | **TUB00_II** | **TUB00_III** | **blank** |
| 764 | 616.651 | 0.503 | -1 | 1.283E+07 | 7.946E+06 | 2.943E+06 | 0.000E+00 | 1.447E+03 | 5.303E+04 | 0.000E+00 |
| 765 | 223.064 | 6.361 | -1 | 6.565E+05 | 1.119E+06 | 3.324E+05 | 7.121E+05 | 1.444E+06 | 1.606E+06 | 5.300E+04 |
| 769 | 498.173 | 7.669 | -1 | 5.030E+06 | 6.141E+05 | 8.253E+04 | 2.696E+05 | 3.105E+05 | 4.989E+05 | 0.000E+00 |
| 771 | 143.107 | 1.088 | -1 | 1.024E+05 | 5.194E+06 | 1.211E+05 | 4.306E+05 | 7.497E+06 | 5.017E+05 | 1.638E+05 |
| 786 | 819.476 | 8.754 | -1 | 1.112E+05 | 3.046E+04 | 7.337E+04 | 1.582E+05 | 9.074E+04 | 1.245E+05 | 8.298E+04 |
| 794 | 811.308 | 6.909 | -1 | 0.000E+00 | 0.000E+00 | 0.000E+00 | 0.000E+00 | 0.000E+00 | 0.000E+00 | 0.000E+00 |
| 797 | 818.299 | 3.374 | -1 | 2.961E+06 | 5.166E+05 | 6.369E+05 | 0.000E+00 | 0.000E+00 | 6.970E+03 | 0.000E+00 |
| 806 | 618.984 | 3.690 | -1 | 5.908E+06 | 1.840E+06 | 1.578E+06 | 0.000E+00 | 0.000E+00 | 2.300E+04 | 0.000E+00 |
| 808 | 832.302 | 3.693 | -1 | 3.782E+06 | 1.308E+06 | 1.076E+06 | 0.000E+00 | 0.000E+00 | 0.000E+00 | 0.000E+00 |
| 814 | 513.189 | 7.230 | -1 | 6.553E+06 | 6.648E+05 | 2.580E+05 | 4.627E+05 | 3.154E+05 | 7.213E+05 | 0.000E+00 |
| 815 | 432.241 | 7.420 | -1 | 1.489E+06 | 1.063E+04 | 2.833E+04 | 1.141E+04 | 0.000E+00 | 7.730E+03 | 4.819E+04 |
| 818 | 123.079 | 0.380 | -1 | 1.688E+07 | 3.321E+08 | 1.809E+07 | 3.239E+06 | 1.963E+08 | 1.300E+07 | 3.881E+05 |
| 822 | 376.087 | 6.788 | -1 | 1.731E+07 | 6.925E+06 | 7.606E+06 | 0.000E+00 | 0.000E+00 | 0.000E+00 | 0.000E+00 |
| 827 | 326.112 | 5.859 | -1 | 2.141E+06 | 2.880E+06 | 1.807E+06 | 0.000E+00 | 3.058E+04 | 2.287E+04 | 0.000E+00 |
| 828 | 319.064 | 6.836 | -1 | 6.034E+06 | 3.209E+06 | 7.227E+06 | 0.000E+00 | 0.000E+00 | 0.000E+00 | 0.000E+00 |
| 831 | 1295.589 | 10.022 | -1 | 1.966E+06 | 4.403E+05 | 3.973E+05 | 0.000E+00 | 0.000E+00 | 0.000E+00 | 0.000E+00 |
| 834 | 628.471 | 3.460 | -1 | 1.601E+06 | 8.697E+05 | 1.928E+05 | 3.468E+04 | 2.144E+04 | 4.207E+04 | 0.000E+00 |
| 835 | 850.609 | 3.462 | -1 | 5.973E+05 | 1.040E+06 | 7.339E+05 | 1.129E+05 | 1.357E+05 | 1.676E+05 | 0.000E+00 |
| 839 | 372.717 | 10.091 | -1 | 3.667E+07 | 7.509E+06 | 0.000E+00 | 0.000E+00 | 0.000E+00 | 0.000E+00 | 0.000E+00 |
| 846 | 681.132 | 5.759 | -1 | 1.487E+06 | 5.917E+05 | 1.208E+06 | 1.665E+04 | 0.000E+00 | 0.000E+00 | 0.000E+00 |
| 856 | 1194.817 | 11.488 | -1 | 7.792E+05 | 1.619E+04 | 9.004E+04 | 0.000E+00 | 1.639E+04 | 0.000E+00 | 1.392E+06 |
| 857 | 804.296 | 3.067 | -1 | 8.677E+05 | 2.268E+05 | 4.048E+04 | 0.000E+00 | 0.000E+00 | 0.000E+00 | 0.000E+00 |
| 863 | 353.267 | 9.157 | -1 | 8.462E+05 | 5.509E+05 | 1.817E+05 | 6.966E+05 | 2.840E+05 | 3.049E+05 | 1.354E+07 |
